# Multi-proxy analyses of a mid-15^th^ century ‘Middle Iron Age’ Bantu-speaker palaeo-faecal specimen elucidates the configuration of the ‘ancestral’ sub-Saharan African intestinal microbiome

**DOI:** 10.1101/817692

**Authors:** Riaan F. Rifkin, Surendra Vikram, Jean-Baptiste Ramond, Alba Rey-Iglesia, Tina B. Brand, Guillaume Porraz, Aurore Val, Grant Hall, Stephan Woodborne, Matthieu Le Bailly, Marnie Potgieter, Simon J. Underdown, Jessica E. Koopman, Don A. Cowan, Yves Van de Peer, Eske Willerslev, Anders J. Hansen

## Abstract

The archaeological incidence of ancient human faecal material provides a rare opportunity to explore the taxonomic composition and metabolic capacity of the ancestral human intestinal microbiome (IM). Following the recovery of a single desiccated palaeo-faecal specimen from Bushman Rock Shelter in Limpopo Province, South Africa, we applied a multi-proxy analytical protocol to the sample. Our results indicate that the distal IM of the Neolithic ‘Middle Iron Age’ (*c.* AD 1485) Bantu-speaking individual exhibits features indicative of a largely mixed forager-agro-pastoralist diet. Subsequent comparison with the IMs of the Tyrolean Iceman (Ötzi) and contemporary Hadza hunter-gatherers, Malawian agro-pastoralists and Italians, reveals that this IM precedes recent adaptation to ‘Western’ diets, including the consumption of coffee, tea, chocolate, citrus and soy, and the use of antibiotics, analgesics and also exposure to various toxic environmental pollutants. Our analyses reveal some of the causes and means by which current human IMs are likely to have responded to recent dietary changes, prescription medications and environmental pollutants, providing rare insight into human IM evolution following the advent of the Neolithic *c*. 12,000 years ago.

## INTRODUCTION

The human gastrointestinal tract (GI) harbours a dynamic population of bacteria, archaea, fungi, protozoa and viruses; the intestinal microbiota. This collection of microorganisms, comprising the human intestinal microbiome (IM) (*1*) performs critical functions in digestion, development, behaviour and immunity (*2, 3*). Modifications of the core IM composition (dysbiosis) have been associated with the pathogenesis of inflammatory diseases and infections (*3, 4*), including autoimmune and allergic diseases, obesity, inflammatory bowel disease and diabetes (*5*). Despite its clinical importance, the factors that contribute to changes in IM taxonomic composition and functionality are not entirely understood (*6, 7*). This is attributed to the fact that most of what is known about the human IM is based on contemporary industrialised and ‘traditional’ human societies (*8–10*). In evolutionary terms, our species have subsisted by hunting and gathering for >90% of our existence (*11*). Evidence derived from the analyses of the IMs of traditional societies, including the Tanzanian Hadza hunter-gatherers (*8*), the Venezuelan Yanomami Amerindians (*5*), the BaAka Pygmies in the Central African Republic (*12*) and the Arctic Inuit (*13*) are thus widely viewed as representing ‘snapshots’ of ancient human IM composition. However, as exposure to Western diets, medicines and microbes cannot be excluded, one must be cautious about the use of these ethnographic cohorts as proxies for prehistoric human IMs (*14*).

The transformation of the IMs of present-day humans to their current ‘modernised’ state commenced millennia ago, with the advent of the Neolithic, which, at *c.* 12,000 years ago (ya), resulted in the first major human dietary transition (*15*). But precisely how our IMs changed following the advent of the Neolithic, and the Industrial Revolution after *c.* AD 1760, remains ambiguous (*16–18*). In this regard, the analyses of ancient human IMs provide a unique view into the co-evolution of microbes and human hosts, host microbial ecology and changing human IM-related health states through time (*2*). Indeed, over the past 15 million years, multiple lineages of intestinal bacterial taxa arose via co-speciation with African hominins and non-human primates, *i.e.*, chimpanzees, bonobos and gorillas (*19*). The departure of behaviourally ‘fully-modern’ *Homo sapiens* from Africa *c.* 75,000 years ago (kya) resulted in the global dispersal of our species (*20*). Significantly, various microbes accompanied these human dispersals ‘out of Africa’ (*21, 22*). Since the ancestral human IM is estimated to comprise a taxonomically and metabolically more diverse array of microbes than those found in contemporary societies (*6, 10*), the IMs of pre-Clovis North Americas (*23*), pre-Columbian Puerto Rican Amerindians (*24*) and pre-Columbian Andeans (*25*) represent more accurate indications of ancient human IM composition. These studies have provided significant insight into the structure, function and evolution of the human IM, highlighting the influence of dietary changes on the intestinal microbial ecology of contemporary humans (*2, 6, 7*). These have also provided essential baseline information for understanding the evolutionary processes implicated in the taxonomic configuration and metabolic capacity of both healthy and dysbiotic human IMs.

Despite the fact that African populations are not underrepresented in studies concerning ‘traditional’ human IMs (*8, 12, 26*), there is, currently, no information concerning the taxonomic diversity and metabolic capacity of ancestral (*i.e.*, archaeologically-derived) African IMs. To gain insight into the ancient African human IM, the prehistoric incidence of intestinal parasites, pathogenic microbes and antibiotic resistance genes, we performed shotgun metagenomic sequencing of a prehistoric (pre-colonial) faecal specimen recovered from a Middle Iron Age (*c.* AD 1485) context at Bushman Rock Shelter (BRS) in Limpopo Province, South Africa. Comparison with ancient (Ötzi), traditional (Hadza and Malawian) and contemporary ‘Western’ (Italian) IM datasets indicate that the IM of the BRS individual represents a unique taxonomic and metabolic configuration observed in neither contemporary African, nor European, populations (see Methods).

## RESULTS AND DISCUSSION

### Specimen provenience and ancient subsistence reconstruction

The palaeo-faecal specimen was recovered *in situ* from an exposed stratigraphic section at BRS (*27*) (Fig. 1a). This large dolomitic rock-shelter is situated on the edge of the Great Escarpment in the Drakensberg chain. The occupation level from which the specimen derives comprises the uppermost archaeological unit of the rock-shelter designated ‘Angel’ in ‘Block A’. Layer 1 (‘Angel’) relates to the arrival of Bantu-speaking Iron Age agro-pastoralists in the region after *c.* 1,800 ya (Fig. S1). This occupation therefore reflects the advent of the Neolithic in South Africa, which entailed the introduction of domesticated taxa such as sorghum (*Sorghum bicolor*), cattle (*Bos taurus*) and various other Iron Age-related species and cultural practices (*e.g.*, ceramic and iron-smelting technologies) into the region (*28*). All the preceding archaeological layers at BRS are representative of occupations by Holocene (*e.g.*, the Oakhurst techno-complex at ∼10 kya), Terminal Pleistocene (the Robberg techno-complex at ∼20 kya) and Pleistocene (*i.e.*, the Pietersburg techno-complex ∼80 kya) hunter-gatherers (*27, 29*).

**Fig. 1.**
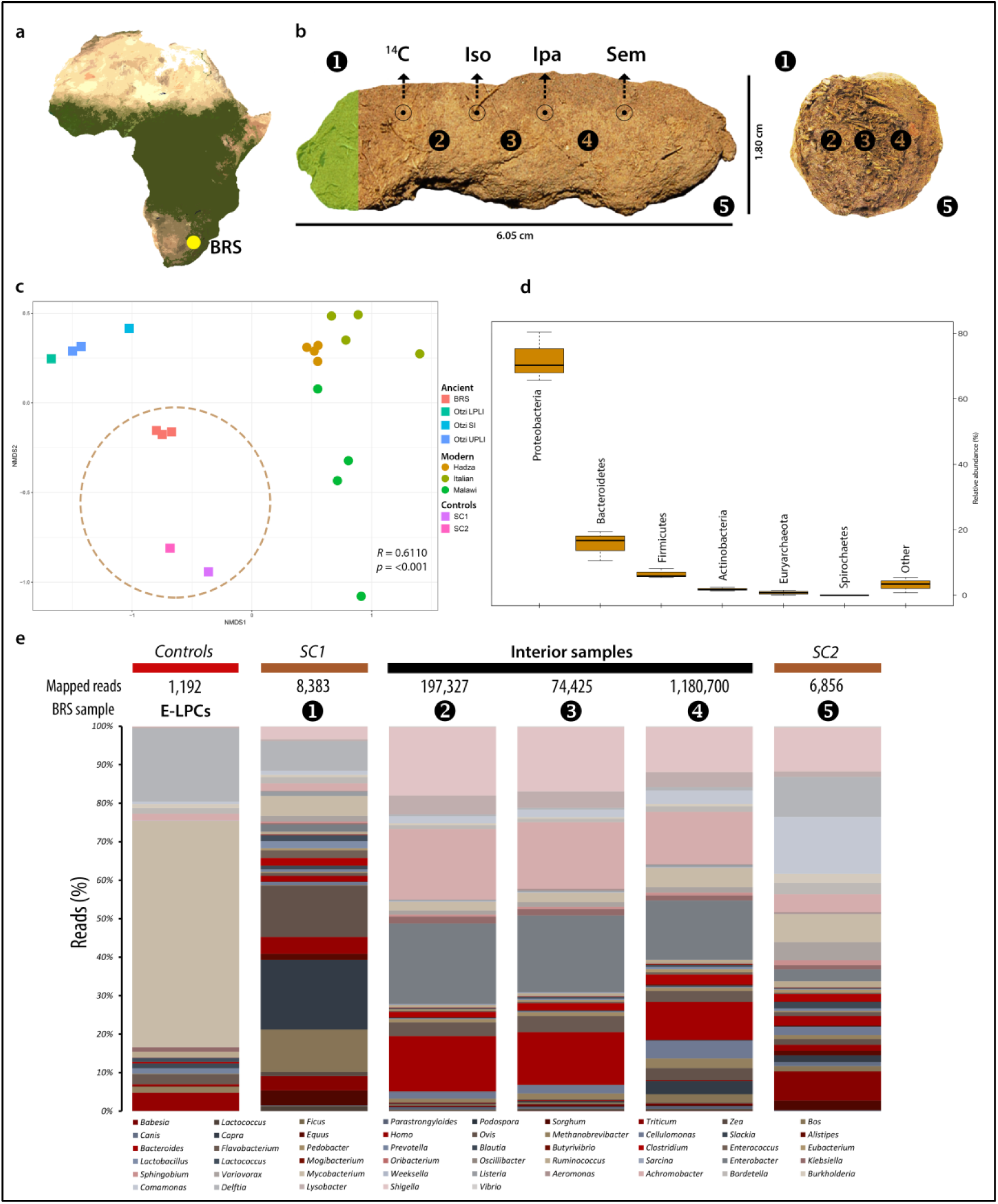
Provenience of, sub-sampling protocol applied to and microbial taxa detected in the BRS palaeo-faecal specimen. **A**) The location of Bushman Rock Shelter (BRS) in Limpopo Province, South Africa, **B**) Lateral (left) and cross-sectional (right) views of the specimen indicating the sub-sampling protocol applied to facilitate DNA extraction, including ‘sedimentary control’ sample 1 (‘SC1’), faecal samples 2, 3and 4 and sedimentary control sample 5 (SC2), ^14^C AMS dating (*^14^C*), isotope analyses (*Iso*), intestinal parasitic analyses (*Ipa*), scanning electron microscopy (*Sem*) and the preservation of a voucher sample (indicated in green shading), **C**) Non-metric multi-dimensional scaling (NMDS) plot comparing the taxonomic community structure (by weighted Bray-Curtis dissimilarity analysis) of the BRS specimen (*i.e*., BRS2, BRS3 and BRS4) and the sediment controls (SC1 and SC2) with the ancient (Ötzi) (indicated as SI ‘small intestine’, LPLI ‘lower part of the lower intestine’ and UPLI ‘upper part of the lower intestine’), traditional (Hadza and Malawian) and modern (Italian) IM datasets (taxa were filtered to the occurrence of >3 in at least 20% of the samples resulting in the inclusion of 371 taxa) (*R* = 0.6110 indicates ANOSIM analysis which revealed significant differences (*p* = <0.001) between the ancient and modern IM samples), **D**) Box-and-whisker plot indicating the relative abundance of intestinal bacterial phyla detected in the BRS specimen (*i.e*., BRS2, BRS3 and BRS4) (‘other’ comprises phyla with <0.6% relative abundance), **E**) Bar-chart providing an overview of all environmental, commensal and pathogenic genera identified in the BRS specimen (BRS2, BRS3 and BRS4) and information concerning the DNA extraction and library preparation negative controls (E-LPCs) and modern and ancient sedimentary controls (SC1 and SC2) (data derived from Table 1, Table 2 and Table S1) (see Methods).

Following recovery of the specimen using latex gloves and stainless-steel forceps (whilst wearing a biologically-impervious body-suit and surgical face-mask) it was sealed in a sterile plastic ziplock bag and stored at ∼4°C. Sediment control sample BRS1 (‘SC1’) was collected from the surface of the rock-shelter (∼25 cm above the specimen) and BRS5 (‘SC2’) from the levels preceding the Iron Age (∼25 cm below the specimen) in the Oakhurst occupation dated to *c*. 10 kya (*27, 29*). These were used as ‘controls’ to assess ecological and faecal community composition for biological plausibility and also the likelihood of sedimentary aDNA (sedaDNA) leaching. Sub-sampling was performed in ancient DNA (aDNA) laboratories at the Centre for GeoGenetics, University of Copenhagen (Denmark), applying established protocols (*30*) (Fig. S2). Prior to sub-sampling, the outer surface or cortex (∼5mm) of the specimen was removed with a scalpel and excluded from further analyses, primarily as it was in contact with surrounding sediment and could therefore have been contaminated by soil-derived microbes (Fig. S2). To address within-sample variability, three faecal sub-samples (*i.e.*, BRS2, BRS3 and BRS4) were taken from different sites within the specimen. From the remaining one-third of the specimen, four sub-samples were taken for radiocarbon (^14^C) dating, isotopic analysis, and microscopic intestinal parasitic and scanning electron microscopy (SEM) analyses. One-sixth of the specimen was preserved (at −20°C) as a voucher sample (Fig. 1b).

To ascertain whether the palaeo-faecal specimen derives from a human individual, all other potential source species were eliminated. Given the limited number (1,967) of aDNA sequence reads mapped to *H. sapiens*, likely due to the removal of the exterior cortex prior to sub-sampling (in which most human-derived (nuclear and mitochondrial) DNA would be expected to reside), metagenome assembly could not be performed. Morphologically, the specimen resembles several candidate species (*31*), although no DNA sequence reads for indigenous felids (*e.g.*, leopard (*Panthera pardus*), caracal (*Caracal caracal*) *etc.*), mustelids (honey badger (*Mellivora capensis*), polecat (*Ictonyx striatus*) *etc.*), jackal (*Canis mesomelas*) or domestic dogs (*C. lupus familiaris*), and none for the indigenous primates, *i.e.*, vervet monkeys (*Cercopithecus aethiops*) or baboon (*Papio ursinus*), were detected. Reads related to non-human primates, *i.e.*, *Pan troglodytes* (the common chimpanzee) and *Macaca mulatta* (rhesus macaque) are likely the result of false-positive identifications, as these taxa do not currently occur in the region, nor would they have in the past (*32*). The incidence of statistically-significant (*i.e.*, verified ancient) C-T *p*-values for the 1,967 reads mapped to *H. sapiens* supports the conclusion that the faecal specimen derives from a human individual (Table S1).

To confirm the association of the faecal specimen with the archaeological context from which it was recovered, two direct radiocarbon (^14^C) Accelerator Mass Spectrometry (AMS) dates were generated from two sub-samples taken from within the specimen (Fig. 1b) (see Methods). The dates of 470 ± 44 years before present (BP) (IT-C-1020) and 460 ± 35 years BP (IT-C-1077) indicate that the sample was deposited *c.* AD 1485 (Table S2). This date falls within the South African Middle Iron Age (spanning AD 1300-1840), follows the demise of the nearby Kingdom of Zimbabwe at *c.* AD 1450 (*33*) and closely precedes first contact with European seafarers in AD 1488 (*34*).

Prior to the identification of environmental and subsistence-related taxa, all exotic taxa, including kiwi (*Apteryx*), carp (*Cyprinus*), salmon (*Oncorhynchus*), pig (*Sus*), chicken (*Gallus*) and rice (*Oryza*) were identified and excluded from further analyses. The evaluation of taxa present in the DNA extraction (*n* = 1) and library preparation (*n* = 1) negative controls (E-LPCs) indicated that instances of environmental contamination were restricted largely to taxa widely cited as either ‘contaminants’ or as derived from false-positive identifications (*35, 36*). The authenticity of microbial and macrobial sequence-derived taxa was determined by statistical aDNA sequence damage estimation (*37*), comparison to E-LPCs, DNA read-length characteristics and ecological conformity. Using high-quality filtered reads for DNA damage estimation analyses with PMDtools (*37*), this process facilitated the validation of forty-seven taxa represented by 1,470,662 reads as ancient (Fig. 2, Table 1, Table 2) (Table S1) (see Methods). Subject to the availability of sufficient numbers of high-quality ‘mappable’ aDNA sequence reads, we employed mapDamage (*38, 39*) to validate the authenticity of taxa by determining the incidence of C-T and G-A substitution rates at the 5’-ends and 3’-ends of strands (see Methods). DNA damage analyses of sequence reads derived from the genera *Bacteroides* and *Shigella* (Fig. 2b, c) and also *B. taurus* and *S. bicolor* (Fig. 2d, e) indicate that the nucleotide composition at the ends of the analysed reads exhibits the typical pattern expected for ancient DNA (*38, 39*).

**Fig. 2.**
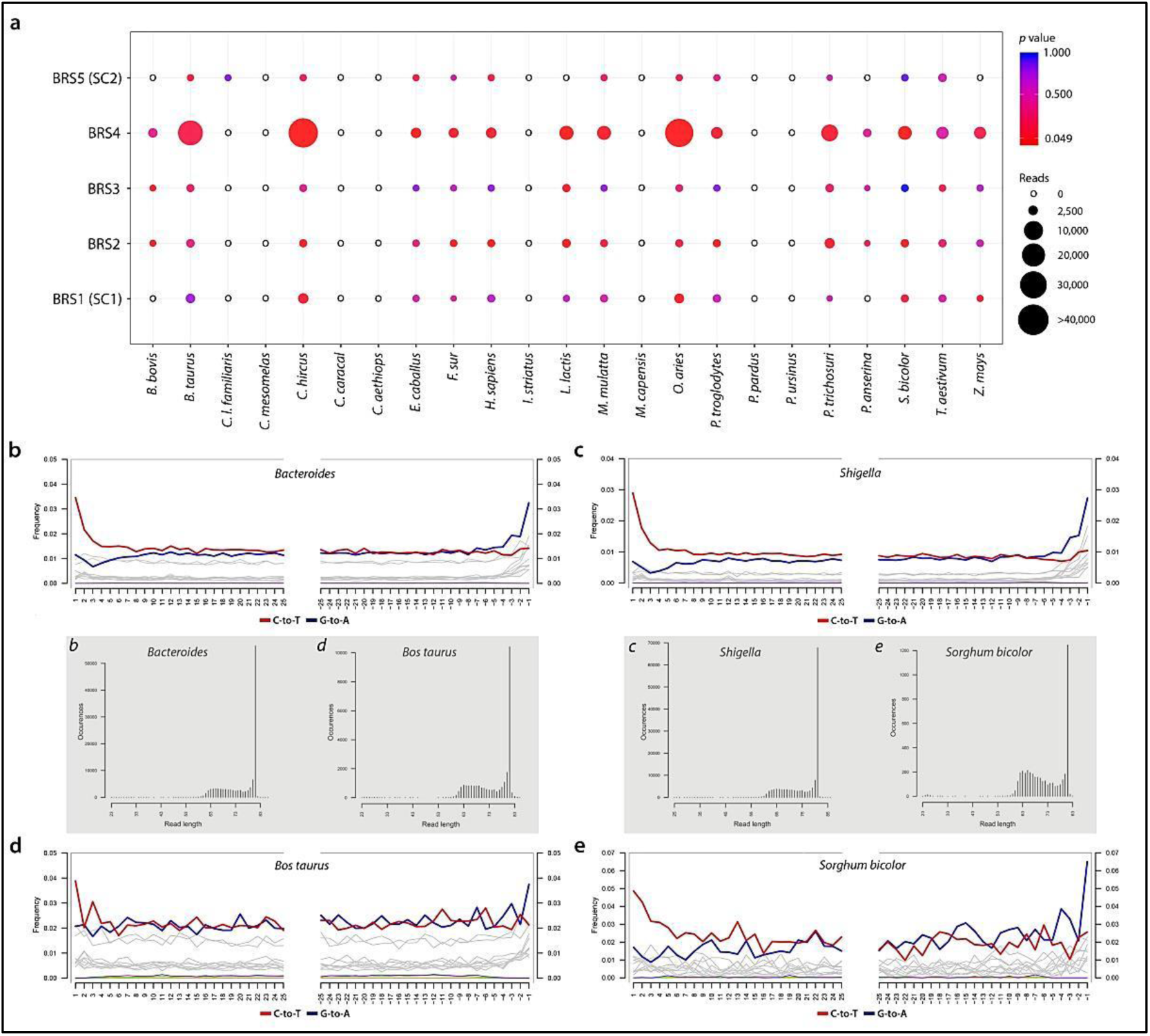
DNA damage estimation analyses and authentication of environmental and subsistence-related taxa detected in the BRS palaeo-faecal specimen. **A**) Dot-plot indicating the occurrence of statistically-significant C-T *p*-values calculated for environmental- and subsistence-related taxa detected in BRS (1 (SC1), BRS2, BRS3, BRS4 and BRS5 (SC2)) (circle sizes and colours represent mapped read-counts and *p*-value significance) and ancient DNA fragmentation patterns shown within the first 25 bp from read ends for the genera **B**) *Bacteroides* and **C**) *Shigella* and the species **D**) *Bos taurus* and E) *Sorghum bicolor* (fragment size distributions for each taxon is indicated in the grey inset and labelled *b*, *c*, *d* and *e*) (Table S1) (Fig. S3).

**Table 1.**
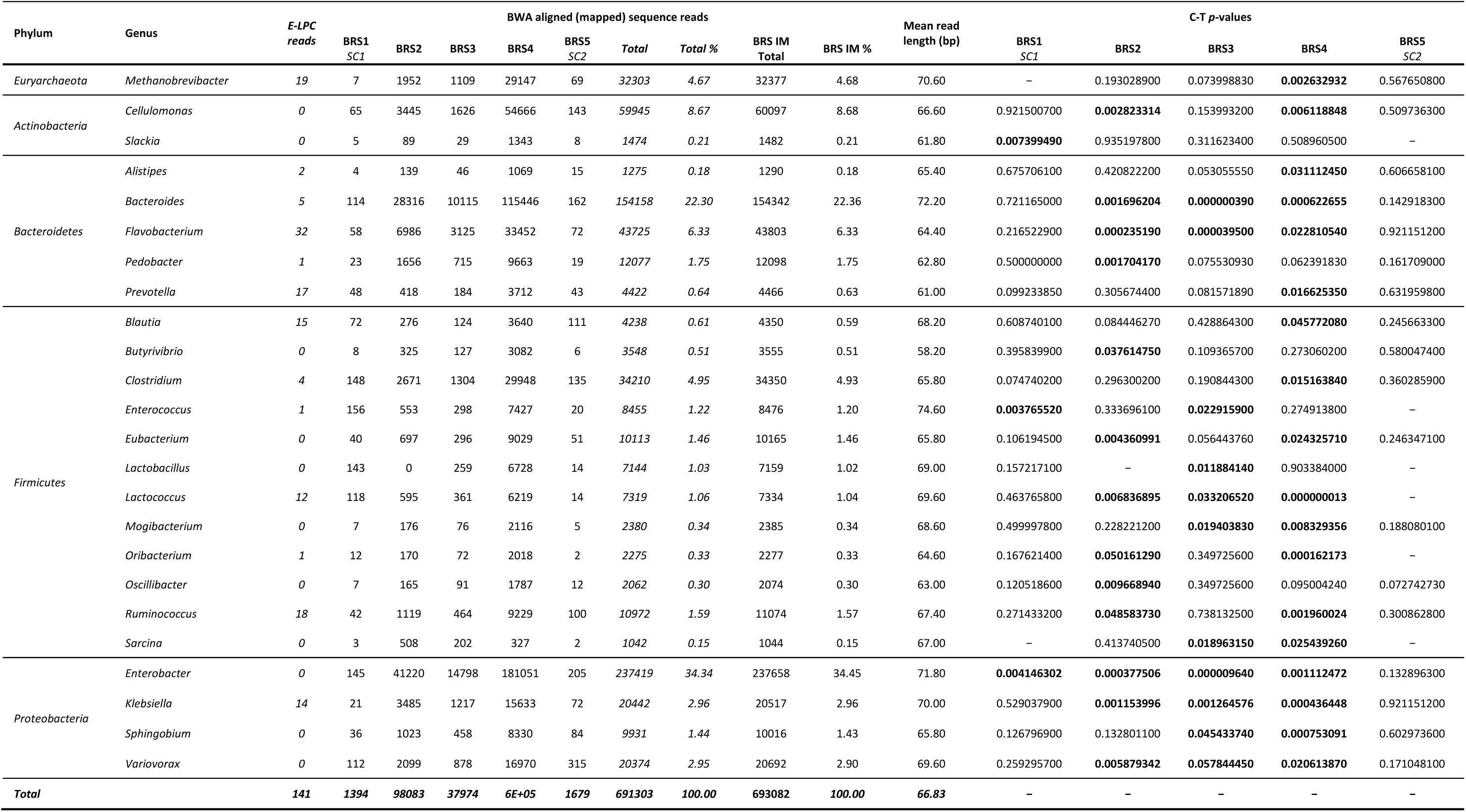
DNA sequence reads for twenty-four authenticated commensal IM taxa detected in the BRS palaeo-faecal specimen. Statistically-significant (*i.e.*, verified ancient) C-T *p*-values are indicated in bold. BWA mapping was performed using high-quality filtered reads for DNA damage estimation analyses using PMDtools (‘C-T *p*- values’) (see Methods). Additional read-length information for individual taxa is provided in Table S5.

**Table 2.**
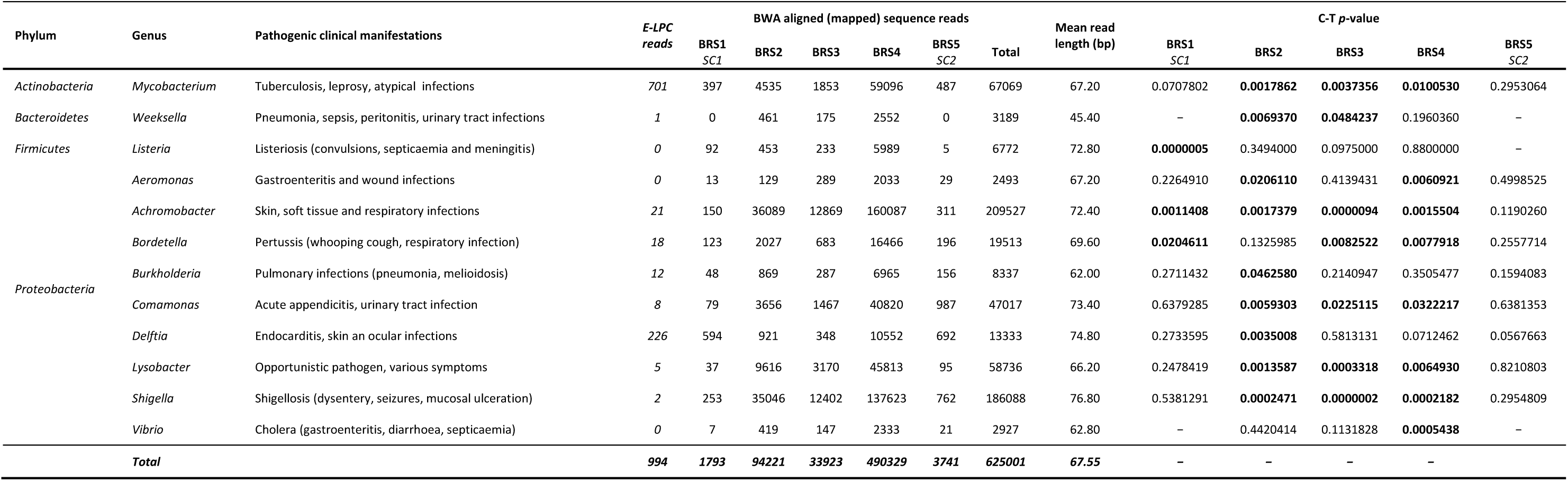
DNA sequence reads for twelve authenticated pathogenic taxa detected in the BRS palaeo-faecal specimen. Statistically-significant (*i.e.*, verified ancient) C-T *p*-values are indicated in bold text. BWA mapping was performed using high-quality filtered reads for DNA damage estimation analyses using PMDtools (‘C-T *p*- values’) (see Methods). Additional read-length information for individual taxa is provided in Table S5.

In dietary terms, the Bantu-speaker agro-pastoralist diet comprised not only domesticated animal and plant taxa, but also various hunted and gathered indigenous species, including antelope, fish, plants and fruits (*40*). The presence of subsistence-related reads derived from sorghum (*S. bicolor*), cluster figs (*Ficus sur*), goat (*Capra hircus*), sheep (*Ovis aries*) and beef (*B. taurus*) are indicative of taxa that were consumed shortly before (*i.e.*, 24 to 36 hours) stool deposition by the BRS individual. Based on bulk untreated *δ*^13^C and *δ*^15^N values obtained from isotopic analyses, the individual had a predominantly C4-based meal with a minor C3-based contribution (see Methods). This concurs with the aDNA evidence indicating the presence of sorghum, wild figs and beef. Sorghum is a C4 plant with published *δ*^13^C values of −14.0 to −11.0‰, and cattle are grazers (*i.e.*, C4 consumers). The *δ*^13^C values (−16.79‰) are higher than −18‰, which is considered the threshold for a predominantly terrestrial diet. These do not however preclude the occasional consumption of freshwater resources, including fish, given the close proximity (∼1.5 km) of the shelter to the perennial Ohrigstad River (*27*) (Table S3) (Fig. S4). The incidence of cattle (*B. taurus*), cattle-specific microbes, *i.e.*, *Lactococcus lactis* (a component in fermented milks) and *Babesia bovis* (the causative agent of babesiosis) are also representative of a Bantu-speaker pastoralist subsistence economy. The non-authenticated incidence of *Podospora anserina* (a dung-colonising fungus) is interpreted as symptomatic of post-depositional saprophytic processes.

### Identifying commensal intestinal microbiota

It is estimated that the human IM harbours ∼150 to ∼400 IM species (*41*), most of which belong to the phyla *Actinobacteria*, *Bacteroidetes*, *Firmicutes* and *Proteobacteria* (*6*). The variability of microbial taxonomic abundance, however, influences the identification of the common core IM (*42*). As many microbes are capable of transient integration into the IM, where they influence the composition and metabolic activity of resident IM communities (*43*), essentially environmentally-derived genera, *i.e.*, *Bacillus*, *Dietzia*, *Microbacterium*, *Paracoccus*, *Pseudomonas*, *Staphylococcus* and *Streptomyces* were omitted from further analyses.

On the basis of taxa detectable at sequencing depth, metagenomic comparison of the shotgun reads with the National Center for Biotechnology Information (NCBI) BLASTn non-redundant nucleotide (*nt*) database using MEGAN Community Edition (CE) v6.10.10 and the Burrows-Wheeler Aligner (BWA) facilitated the identification of 691,303 reads (1.48% of all sequence reads) representing thirty-six ancient commensal IM genera. Subsequent statistical DNA damage estimation resulted in the elimination of 12 non-authenticated bacterial genera (*i.e.*, 29,012 reads) from the dataset, including *Bifidobacterium*, *Coprococcus*, *Dorea*, *Faecalibacterium*, *Mollicutes*, *Neisseria*, *Parabacteroides*, *Phascolarctobacterium*, *Romboutsia*, *Roseburia*, *Ruminiclostridium*, *Tissierellia* and *Treponema* (see Methods) (Table S4). Based on the remaining authenticated 693,082 sequence reads exhibiting an average read-length of 66.83 base-pairs (bp), twenty-four ancient IM taxa (Table 1) were identified. It is of interest to note that, whereas the BRS1 ‘surface control’ sample (SC1) yielded authenticated reads derived from ancient microbial IM taxa (*i.e.*, *Enterobacter*, *Enterococcus* and *Slackia*), the much older BRS5 control (SC2) (*i.e.*, the Oakhurst occupation dated to *c.* 10 kya), did not (Table 1, 2) (Table S1). The IM of the BRS individual (*i.e.*, including only the interior sub-samples BRS2, BRS3 and BRS4) was determined to be dominated by the two phyla *Proteobacteria* (41.73%) and *Bacteroidetes* (31.25 %), followed by *Firmicutes* (13.44%), *Actinobacteria* (8.89%) and *Euryarchaeota* (4.68%) (Table 1) (Fig. 1d). At genus level, the bulk (71.82%) of reads was ascribed to *Enterobacter* (34.45%), *Bacteroides* (22.36%), *Cellulomonas* (8.68%) and *Flavobacterium* (6.33%). In addition to *Clostridium* (4.93%) and *Methanobrevibacter* (4.68%), all other genera exhibit <5% relative abundance. We note that the use of ‘relative abundance’ as a measure of taxonomic representation has been a standard means by which differences in taxonomic composition or ‘abundance’ in IM datasets is analysed, verified and compared, and that various notable IM studies have adhered to the use of ‘relative abundance’ as standard analytical protocol (*6, 8, 9,12, 24, 25, 44*). In addition, while cognisant of the compositional complexity of microbiome samples (*44*) and of the possible influence of the fragmented nature of ancient microbial DNA on taxonomic classification (*45*), we note that ancient DNA damage have been revealed to exert a minimal influence on species detection and on the ‘relative abundance’ of IM taxa in simulated ancient and modern datasets (*46*).

### Identification of ancient pathogenic microbial taxa

There is an estimated 1,407 recognised species of human pathogens (*47*), many of which influence not only health and immune responses (*48*), but also cognitive development (*49*) and social behaviour (*50*). On the basis of taxa detectable at sequencing depth, metagenomic comparison of the shotgun reads with the NCBI BLASTn non-redundant nucleotide (*nt*) database using MEGAN and BWA facilitated the identification of 625,001 ‘mappable’ reads (1.34% of all sequence reads) representing twelve ancient pathogenic taxa (Table 2). Only authenticated ancient taxa were retained (exhibiting an average read-length of 67.55 bp), the authenticity of which was determined by assessing the incidence of statistically-significant (*p* = <0.05) C-T substitutions at the 5’ ends of sequence reads (Table 2).

The occurrence of authenticated ancient reads homologous to *Listeria* and restricted to BRS1 (SC1), suggests that this taxon, although ancient, is most likely environmental (*i.e.*, sedimentary) and that it does not derive from the faecal specimen. The incidence of authenticated DNA reads for *Mycobacterium* in BRS2, BRS3 and BRS4, and not in SC1 or SC2, is indicative of the general environmental (*i.e.*, sedimentary) presence of this genus. However, although not a known member of the intestinal microbiota, and given that not all species are pathogenic, the presence of authenticated ‘ancient’ reads derived from *Mycobacterium* within the faecal specimen cannot be precluded as symptomatic of an infection. Some species, particularly *M. avium*, is known to invade intestinal epithelial cells and has been implicated in ulcerative colitis (*51*). Similarly, the incidence of authenticated reads for the genera *Comamonas*, *Lysobacter* and *Shigella* in BRS2, BRS3 and BRS4, *Aeromonas* in BRS2 and BRS4 and *Vibrio* in BRS4, confirms that these taxa derive from the faecal specimen and are, consequently, representative of an ancient gastro-intestinal infection. *Achromobacter* occurs in all sub-samples except for the ancient (*c.* 10 kya) control (BRS5/SC2) which did not yield any authenticated microbial pathogenic taxa. The notable abundance of commensal (41.73 %) (Table 1) and pathogenic (87.67%) (Table 2) members of the *Proteobacteria* in the BRS IM are of interest as it has been established that an increase in *Proteobacteria* is indicative of IM dysbiosis and metabolic disease (*52*). Compared to primary human IM phyla, *i.e.*, *Actinobacteria, Bacteroidetes* and *Firmicutes*, the relative abundance of *Proteobacteria* in the IM is, however, highly variable. While an increase in the abundance of *Proteobacteria*, especially members of the *Enterobacteriaceae* (*i.e.*, *Klebsiella*, *Salmonella* and *Shigella*) (*53*) is a feature of bacterial dysbiosis, the human IM also contains members of commensal *Proteobacteria*, *i.e.*, *Enterobacter*, *Klebsiella*, *Sphingobium* and *Variovorax.* Under ‘healthy’ conditions, the relative abundance of *Proteobacteria* in the human IM can increase to ∼45% without observable clinical implications (*51*).

Microscopic analysis aimed to determine the presence of intestinal parasites, namely helminths and protozoa, did not yield conclusive results (see Methods). Although this concurs with the aDNA results, the analyses of a single sub-sample might not have been sufficient to detect intestinal parasitic remains. Conversely, not all members of a population would necessarily be infected by intestinal parasites, possibly because of either natural resistance or limited exposure to contaminant sources. Similarly, while SEM analyses did not result in the detection of parasitic remains, it did facilitate the recognition of degraded plant fragments, and also of desiccated bacterial cells and saprophytic organisms, the latter of which likely represent both ancient and modern organisms, respectively (Fig. S5).

### Ancient and modern IM taxonomic comparisons

In terms of taxonomic composition, the ancient samples (BRS and Ötzi) exhibit spatial separation from the ‘traditional’ (Hadza, Malawian) and modern (Italian) comparative cohorts. Hierarchical clustering using complete-linkage based on Spearman’s correlations, produced a clear separation of ancient and modern populations. ANOSIM analysis revealed significant differences between the ancient and modern IM samples (*R* = 0.9098; *p* = <0.001) for 371 taxa (Fig. S6) (see Methods). As stated, and bearing in mind the compositional complexity of IM samples (*44*) and the conceivable influence of fragmented DNA on taxonomic classification (*45*), ancient DNA damage result in minor differences in species detection and in comparisons concerning the ‘relative abundance’ of microbial taxa identified in ancient and modern IM datasets (*46*). As with weighted Bray-Curtis analysis based on the relative abundance of all identified IM taxa (Fig. 1c), we note that un-weighted Bray-Curtis analyses based on the ‘presence-absence’ of IM taxa exhibits correspondingly clear differences between the sedimentary controls (SC1 and SC2) and the ancient (*i.e.*, BRS and Ötzi), ‘traditional’ (Hadza, Malawian) and modern (Italian) comparative IM cohorts (ANOSIM *R* = 0.8361; *p* = <0.001) (Fig. S7). With regards potential contamination derived from the surrounding archaeological sedimentary matrix in the BRS palaeo-faecal specimen, comparison of the incidence of the 24 authenticated ancient IM taxa (Table 1) indicate that the surrounding sedimentary matrix (BRS1 ‘SC1’ and BRS5 ‘SC2’) and the DNA extraction and library preparation negative controls (E-LPCs) are not significant sources of microbial taxa identified in the palaeo-faecal specimen (BRS2, BRS3 and BRS4) (Fig. S8).

Metagenomic comparison of all analysed shotgun reads revealed that IM of the BRS individual (*i.e.*, BRS2, BRS3 and BRS4) is characterised by a *Firmicutes*/*Bacteroidetes* (F/B) ratio significantly skewed towards *Bacteroidetes* (at 31.25%), as opposed to *Firmicutes* (at 13.44%) (Fig. 1d) (Table 1). The F/B ratio (*54*) is widely considered as significant in human IM composition, with dysbiosis associated with inflammation, obesity, and metabolic diseases (*55*). Although this significance is controversial (*56*), we note that the BRS F/B ratio (0.4) does not resemble those reported for modern ‘traditional’ Bantu-speaking Africans in Burkina Faso (2.8) (*57*) nor that calculated here for the East African Hadza (2.6) (*8*). This can likely be attributed to the fact that ‘traditional’ diets rich in starches (*e.g.*, potatoes, yams and sweet potatoes) have been shown to increase the F/B ratio, including increases in relative abundance of *Firmicutes* and enzymatic pathways and metabolites involved in lipid metabolism (*58*).

The IMs of modern humans have furthermore been stated to comprise one of three ‘enterotypes’, based on prevailing genera, *i.e.*, *Bacteroides*, *Prevotella* or *Ruminococcus* (*59*). Some taxa relate to long-term diets, such as *Bacteroides*, which is associated with the consumption of large amounts of protein and animal fat, and *Prevotella*, which is indicative of a high plant-derived carbohydrate intake (*60*). Similarly, *Ruminococcus* prevails in individuals who consume significant amounts of polyunsaturated fats, *e.g.*, marine fish, vegetable oils and nuts and seeds. The enterotypic composition of the BRS IM diverges from that reported for ‘traditional’ Africans (*8, 12, 15, 26, 57, 61*). In relation to *Ruminococcus* (1.57%) and *Prevotella* (0.63%), the BRS IM is characterised by a predominance of *Bacteroides* (22.36%) (*i.e.*, ‘Enterotype 1’) which concurs with a diet rich in protein and animal fat and which lends support our interpretation of the BRS *Bacteroidetes*-dominated F/B ratio. While this corresponds to data reported for the West African BaAka (*12*), it differs from the IM taxonomic composition reported for modern African cohorts, including the Tanzanian Hadza (*8*) and children in Burkina Faso (*57*), which exhibits higher abundance of *Prevotella* (Fig. 1e). The sizable incidence of *Flavobacterium* (6.33%) in the BRS IM likely relates to the fact that members of this genus are resistant to dietary phenolic compounds derived from largely ‘medicinal’ plant taxa, including phenolic acids, flavonoids, tannins, curcuminoids, coumarins, lignans, quinones *etc*. (*62*). This genus also occurs in the IMs of non-human primates, including baboons (*P. ursinus*) and gorillas (*Gorilla gorilla*) (*63*). In relation to its substantial presence in the BRS IM, members of this genus might also have played a role in the elimination of aflatoxins present in milk, cheese, grains and figs. *Methanobrevibacter* (4.68%) is the most abundant archaeon in the human IM (*64*). Besides consuming fermentation products produced by saccharolytic bacteria, archaeal methanogenesis also improves the efficiency of polysaccharide fermentation.

The BRS IM furthermore exhibits enrichment towards *Cellulomonas* (8.68%) which degrades cellulose (*65*), *Clostridium* (4.93%) which is essential for IM resistance to infection and dysbiosis (*66*) and *Pedobacter* (1.75%) and *Prevotella* (0.63%) which, resembling non-pathogenic *Treponema*, are cellulose and xylan hydrolyzers (*8*). *Alistipes* (0.18%) is associated with a protein-rich diet and involved in amino acid fermentation (*61*) and *Butyrivibrio* (0.51%) ferments sugars, cellodextrins and cellulose (*67*). Since the antiquity of *Treponema* (*Spirochaetes*) in the BRS IM could not be verified, we cannot substantiate the premise that *Treponema* is inherently characteristic of all ‘traditional’ IMs (*8, 9*). Less abundant taxa, *i.e.*, *Ruminococcus* (1.57%), *Eubacterium* (1.46%) and *Enterococcus* (1.20%) are implicated in the digestion of starches and vitamin synthesis. *Sarcina* (0.15%) synthesizes microbial cellulose and occurs in high numbers amongst yam-farming African Pygmy hunter-gatherers and traditional populations in Papua New Guinea (*68*).

### Ancient and modern IM metabolic comparisons

Despite having highly divergent IM taxonomic compositions, functional gene profiles are relatively similar amongst different contemporary human populations. Accordingly, microbial community (*i.e.*, taxonomic) composition does not afford a thorough understanding of microbial IM community function (*i.e.*, metabolic capacity) (*69*). To ascertain statistically-significant differences between the metabolic capacities of ancient (pre-industrial) and modern ‘Westernised’ human IMs, we explored and compared the functional IM capacities of two ancient (BRS and Ötzi), two traditional (Hadza and Malawian) and one modern ‘Western’ (Italian) human cohorts (see Methods). This was performed only for the 24 ancient authenticated IM taxa (Table 1). ANOSIM analysis revealed significant difference (*R* = 0.5395; *p* = 0.001) in the metabolic capacity of 24 authenticated ancient taxa for the ancient and modern cohorts. Spearman’s correlation was performed on the taxa linked to the KEGG categories (*i.e.*, 1,487 KO gene categories linked to specific IM taxa). Although not contiguous to the faecal specimen, the considerable differences in the incidence of particular KEGG Orthology (KO) genes in BRS1 (SC1) and BRS5 (SC2) (the younger surface-derived and the ancient Oakhurst (*c*. 10 kya) sediment samples), and BRS2, BRS3 and BRS4, eliminates the sedimentary matrix as a source for the greater proportion of KO genes identified in the faecal specimen (Fig. 3). The differential distribution of the 24 authenticated ancient IM taxonomic categories (Table 1), particularly in terms of the taxa detected in SC1 and SC2 *vs*. those detected in the BRS specimen (BRS2, BRS3 and BRS4) (*R* = 0.8361; *p* = <0.001), lends support to this conclusion (Fig. S7). In SC1 (BRS1), only three taxa (*i.e.*, *Enterobacter*, *Enterococcus* and *Slackia*) could be authenticated as ancient. No authenticated ancient taxa were recovered from SC2 (BRS5).

**Fig. 3.**
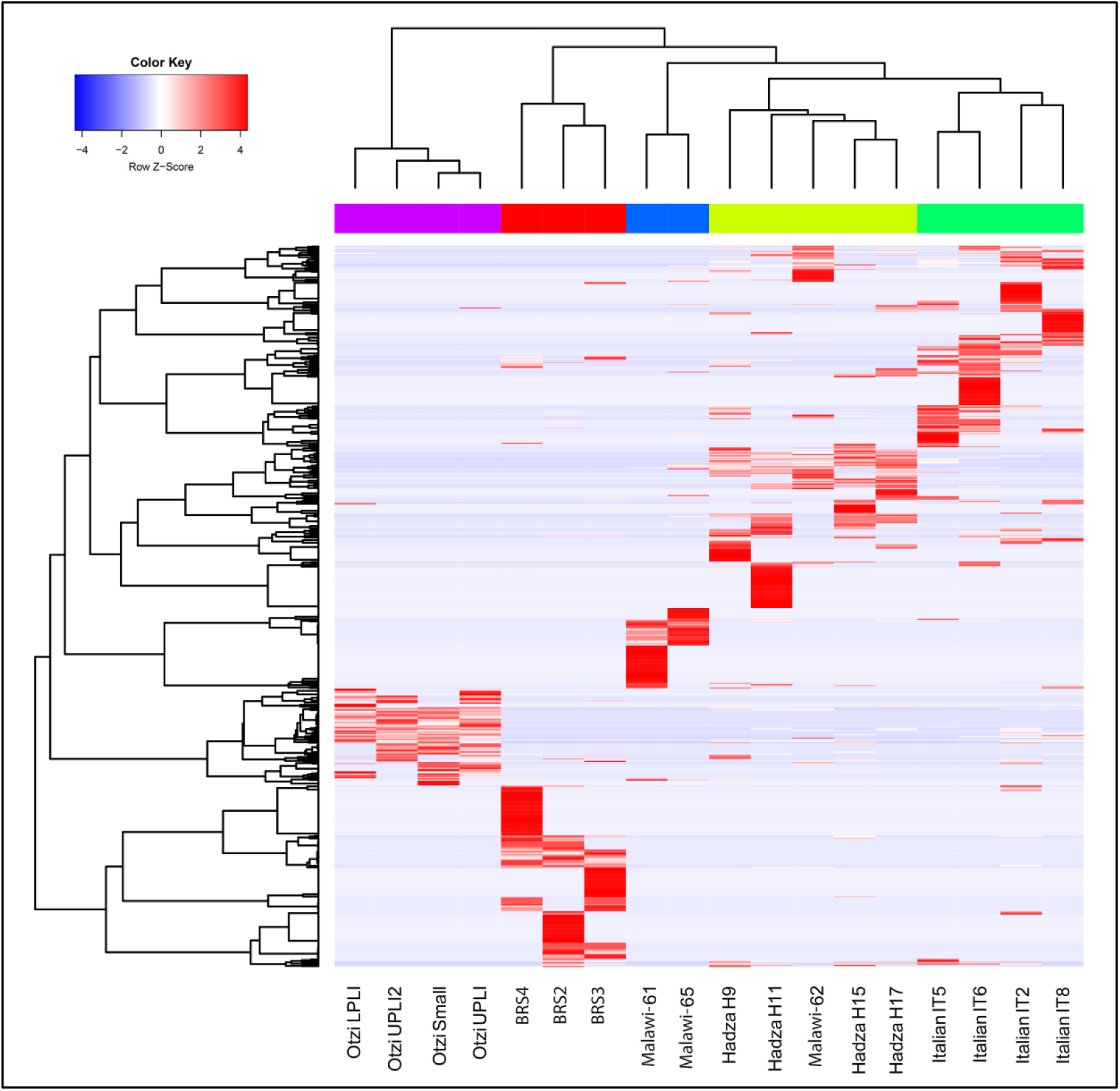
Functional (metabolic) comparison of the ancient (BRS and Ötzi), ethnographic (Hadza and Malawian) and contemporary (Italian) faecal-derived human IMs based on KO-gene analyses for the twenty-four ancient authenticated IM taxa listed in Table 1 (Table S10). In SC1 (BRS1), only three taxa (*i.e.*, *Enterobacter*, *Enterococcus* and *Slackia*) could be authenticated as ancient, and no authenticated ancient taxa were recovered from SC2 (BRS5). The heat-map based on Spearman’s correlation coefficients comparing differences in functionality for the BRS IM (*i.e.*, BRS2, BRS3 and BRS4) with the ancient, traditional and contemporary IM datasets.

ANOSIM analysis revealed significant differences (*R* = 0.4840; *p* = <0.001) in the metabolic capacity of the ancient and modern comparative cohorts (Fig. 4a). Based on the analyses of the KO gene categories occurring only in the 24 authenticated ancient taxa (*i.e.*, 1,487) (Table 1), 72 taxa-specific KO genes are identified as unique to the BRS IM (Fig. 4b). Metagenomic comparison of the shotgun reads with the BLASTx NCBI non redundant protein (*nr*) database using DIAMOND v0.8.36.98 and MEGAN CE v6.10.10 revealed that 117 taxa-specific KO genes (7.86%) are shared between all (*i.e.*, ancient and modern) IM cohorts. While this is indicative of the relative temporal stability of a core commensal human IM community, it also reflects the variable (adaptable) and transient nature of human commensal IM community composition and metabolic capacity (*3, 39, 70*). This responsive adaptability is also echoed by the variable co-abundance of metabolic pathways identified in the BRS IM and the ancient and modern cohorts (Fig. 4c) (Table S7, S8, S9).

**Fig. 4.**
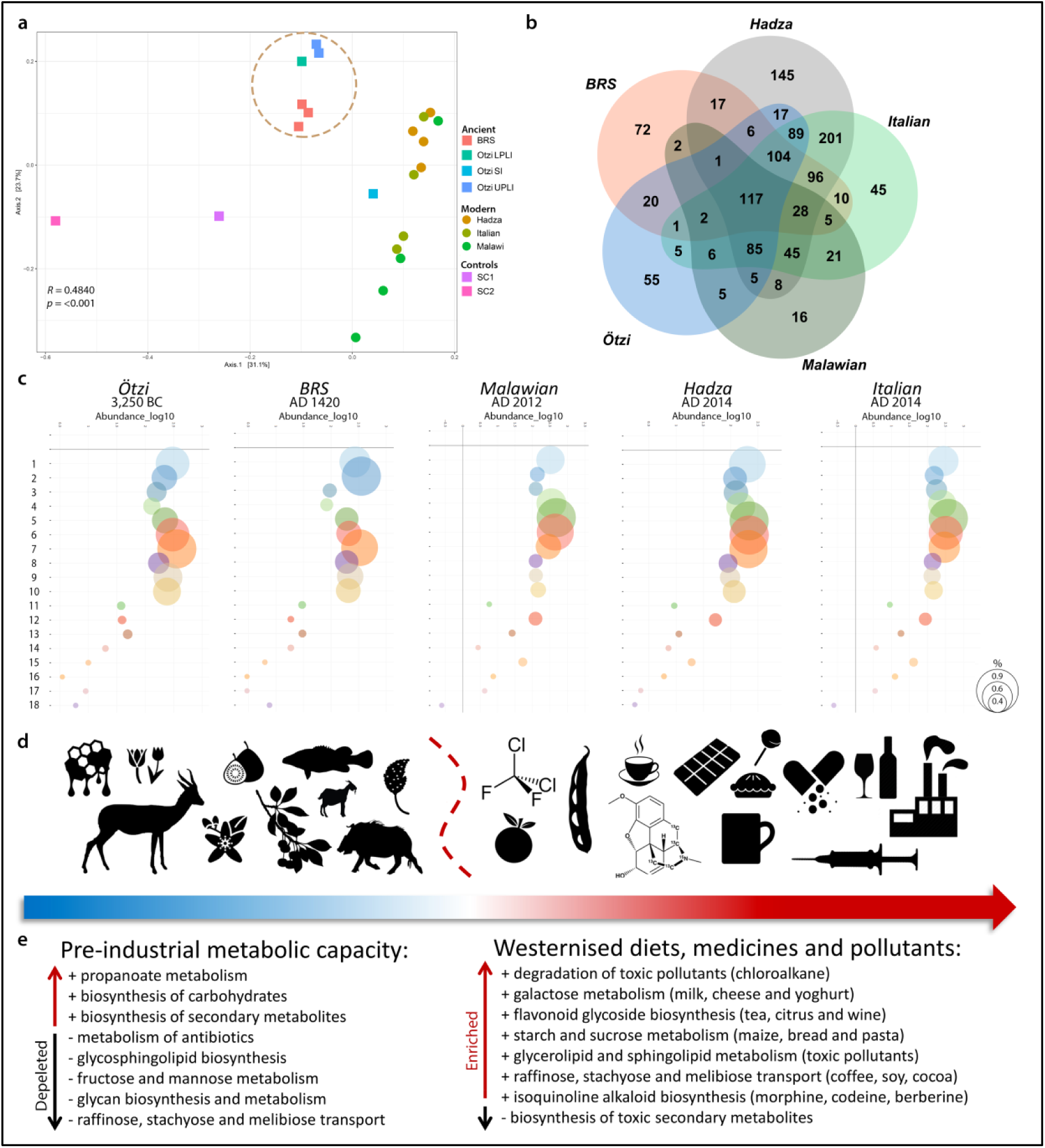
Graphic summary of dietary- and environmentally-induced differences in the metabolic capacities of the ancient and modern IM datasets analysed in this study. **A**) Principal coordinates analysis (PCoA) comparison of the metabolic (functional) capacity of the BRS specimen (*i.e.*, BRS2, BRS3 and BRS4) and the sediment controls (SC1 and SC2) with the ancient (Ötzi) (SI ‘small intestine’, LPLI ‘lower part of the lower intestine’ and UPLI ‘upper part of the lower intestine’), traditional (Hadza and Malawian) and modern (Italian) IM datasets (KEGG categories were filtered for occurrence of >3 in at least 20% of the samples), **B**) Venn diagram indicating the relative abundance of IM taxa-linked KO genes identified in the ancient (*i.e.*, BRS2, BRS3, BRS4 and Ötzi), traditional (Hadza and Malawian) and modern (Italian) comparative cohorts, calculated as based on the 24 authenticated ancient IM taxa indicated in Table 1, **C**) Bubble-charts indicating the co-abundance (log_10_) of eighteen (labelled ‘1’ to ‘18’) metabolic IM capacities for the ancient, traditional and modern IM cohorts (bubble sizes are representative of the relative abundance of KEGG categories (see scale on right) and comprise 1) glycolysis/gluconeogenesis, 2) citrate cycle, 3) fructose/mannose metabolism, 4) galactose metabolism, 5) starch/sucrose metabolism, 6) amino sugar and nucleotide sugar metabolism, 7) pyruvate metabolism, 8) glyoxylate/dicarboxylate metabolism, 9) propanoate metabolism, 10) butanoate metabolism, 11) synthesis and degradation of ketone bodies, 12) sphingolipid metabolism, 13) biosynthesis of unsaturated fatty acids, 14) n-glycan biosynthesis, 15) glycosphingolipid biosynthesis (-globo), 16) glycosphingolipid biosynthesis (-ganglio), 17) chloroalkane/ chloroalkene degradation and 18) naphthalene degradation) (Table S7), **D**) Dissimilarities in ancient and modern IM metabolic capacities are related to recent (historical) changes in human dietary composition and exposure to toxic environmental pollutants (as indicated by the icons and the blue and red arrow) and **E**) Differences in IM metabolic capacities are contrasted in terms of the up- and down-regulation of IM metabolic capacities as an ‘ancient’ *vs*. ‘modern’ comparative summary (see Methods).

In our comparison of the functional IM capacities of the two ancient (BRS and Ötzi), two traditional (Hadza and Malawian) and one modern ‘Western’ (Italian) human IM cohorts, we also focussed our analyses only on the 24 authenticated ancient IM taxa as indicated in Table 1 (Table S8, S9, S10). In relation to the modern (Italian, Hadza and Malawian) and ancient (Ötzi) datasets, the BRS IM (*i.e.*, BRS2, BRS3 and BRS4) appears to exhibit enrichment of KO genes implicated in the biosynthesis of secondary metabolites, including K00163 (Kruskal-Wallis value (*H*) = 14.151 and *p*-value (*p*) = 0.002), K00164 (*H* = 14.096, *p* = 0.002), K00163 (*H* = 15.812, *p* = 0.014), K00600 (*H* = 11.243, *p* = 0.004), K00568 (*H* = 11.706, *p* = 0.003) and K00457 (*H* = 14.762, *p* = 0.002) (Table S10). K00568 and K00457 are also implicated in the biosynthesis of terpenoid-quinones. The capacity to biosynthesise toxic secondary metabolites (*e.g.*, polyketides, isoprenoids, aromatics (phenylpropanoids) or alkaloids) is essential when dietary sources comprise largely natural unprocessed foods. The BRS IM also exhibits enrichment of genes implicated in glyoxylate and dicarboxylate metabolism (K03781: *H* = 15.275, *p* = 0.018 and K00600) and the citric acid cycle (CAC) (K00164: *H* = 15.650, *q* = 0.015 and K02274: *H* = 15.928, *p* = 0.014). Glyoxylate and dicarboxylate glyoxylate are involved in the biosynthesis of carbohydrates, and CAC facilitates the release of energy from dietary carbohydrates, proteins and fats. Genes involved in glycolysis and gluconeogenesis (K00163: *H* = 15.812, *p* = 0.015) (pyruvate dehydrogenase E1 component) are also enriched.

The BRS IM exhibits depletion of KO genes involved in raffinose, stachyose and melibiose transport (*e.g.*, K10119: *H* = 15.934, *p* = 0.015, K10118: *H* = 15.640, *p* = 0.015 and K10117: *H* = 16.383, *p* = 0.011) (Table S8). Soybeans are primary dietary sources of raffinose and stachyose, and melibiose occurs in coffee, cacao and processed soy (*71*). Deglycosylation by intestinal epithelial cell beta-glucosidases is a critical step in the metabolism of dietary flavonoid glycosides derived specifically from tea, citrus and wine (K05349: *H* = 15.068, *p* = 0.019). The BRS IM also contrasts with the modern cohort in terms of the depletion of genes involved in glycan (sugar-chain) biosynthesis and metabolism (alpha-mannosidase) (K01206: *H* = 14.443, *p* = 0.025) (beta-galactosidase) (K01190: *H* = 15.777, *p* = 0.015). While modern oligosaccharides are derived largely from processed ‘table sugar’ comprising mainly sucrose and fructose, foremost natural sources of sugar, comprising fruits and honey, would not have been consistently available for consumption. Correspondingly, KO genes involved in starch and sucrose metabolism (K00705: *H* = 15.318, *q* = 0.018 and K00975: *H* = 15.438, *p* = 0.017) (starch phosphorylase) (K00688: *H* = 13.496, *p* = 0.035) and fructose and mannose metabolism (K01193: *H* = 14.468, *p* = 0.025) (6-phosphofructokinase 1) (K00847: *H* = 16.009, *p* = 0.014) (fructokinase) are also depleted.

Whereas the Hadza IM is enriched in genes involved in fructose and mannose metabolism, the Italian IM is enriched in genes involved in the metabolism of simple sugars *e.g.*, glucose, galactose and sucrose (*26*). KO genes involved in the degradation of toxic pollutants (*i.e.*, chloroalkane, chloroalkene and naphthalene) and butanoate metabolism (K04072: *H* = 13.468, *p* = 0.036 and K00128: *H* = 15.970, *p* = 0.013) are also depleted in the BRS IM. This is noteworthy, as there are no known natural sources of chlorinated paraffins (CPs) (*72*). CPs, including chloro-alkanes (C_10-13_), are widely used in the production of refrigerants, solvents, plasticisers and fire-retarding agents (*73*). Naphthalene (C_10_H_8_), a polycyclic aromatic hydrocarbon, is derived from petroleum distillation and used in the manufacture of plastics, resins, fuels and insecticides. In addition, the depletion of KO genes involved in galactose, glycerolipid and sphingolipid metabolism and glycosphingolipid biosynthesis (K07407: *H* = 14.352, *p* = 0.026) (alpha-galactosidase) (K01190) (beta-galactosidase) is significant as it provides insight into the influence of exposure to modern environmental pollutants on IM composition and metabolic capacity.

Although it is challenging to infer ancient diet from ancient IM data, there is a growing understanding of the role of dietary choices on IM composition (*2*). The enrichment, in the BRS IM, of genes serving specific metabolic processes, including the biosynthesis of secondary metabolites, xenobiotic biodegradation, the metabolism of terpenoids and polyketides, propionate and butanoate and lysine degradation and the synthesis of ketone bodies is noteworthy as it suggest that the BRS metabolic profile is indicative of a diet rich in unprocessed natural resources, conceivably comprising medicinal plant substances (given the capacity for biosynthesising secondary metabolites and biodegrading xenobiotics), and encompassing irregular dietary intake. In addition, and given the enrichment of KO genes implicated in the synthesis and degradation of ketone bodies (*i.e.*, K00626), the metabolic profile of the BRS individual approximates that induced by a ketogenic diet (*74*), characterised by high-fat, adequate-protein and low-carbohydrate dietary consumption and accompanied by prolonged exercise and periods of low dietary intake or unintentional ‘fasting’ (Fig. 4d) (Table S9). Whether this metabolic profile resembles that generally referred to as a ‘palaeo-diet’ is unclear, as this would have entailed the exclusion of dairy, grains and legumes, nutritional categories which did indeed form part of the BRS diet. The BRS IM, and also that of Ötzi, are furthermore characterized by depletion of KO genes involved in the metabolism of antibiotics (*e.g.*, K11358: *H* = 15.320, *p* = 0.017), including aminocoumarin antibiotics and the metabolism of isoquinoline alkaloids, including the opiates morphine and codeine, as well as the antibiotic berberine.

In contrast to the ancient IMs, the modern (Hadza, Malawian and Italian) IM is characterized by enrichment of KO genes involved in raffinose, stachyose and melibiose transport (*e.g.*, K10119, K10118 and K10117) indicative of a diet comprising soy, coffee, cacao and dietary flavonoid glycosides derived from tea, citrus and wine (K05349) (Table S10). This group also exhibits enrichment in genes concerning galactose metabolism, *i.e.*, K07407 (*H* =14.352, *p* = 0.0259) (alpha-galactosidase), K00849 (*H* = 15.553, *p* = 0.016) (galactokinase) and K00965 (*H* = 15.694, *p* = 0.015) (UDPglucose--hexose-1-phosphate uridylyltransferase). Galactose is metabolized from milk sugar (lactose, a disaccharide glucose and galactose), the primary dietary source of which is milk and yogurt. KO genes involved in starch and sucrose (K00705) and amino sugar and nucleotide sugar metabolism (K00965 and K00849: *H* = 15.553, *p* = 0.016) are also enriched, and so are genes involved in glycine, serine, threonine and methane and antibiotic metabolism. The enrichment of genes involved in glycerolipid and sphingolipid metabolism and glycosphingolipid biosynthesis (*e.g.*, K01190 and K07407) (alpha-galactosidase) likely reflects the impact of modern environmental pollutants on IM composition and metabolic capacity. It would be of interest to determine whether this does in fact represent a ‘population-wide’ functional response to exposure to toxic compounds ubiquitous in modern industrialized urban environments

In summary, significant differences between the ancient and modern IM metabolic capacity comprise the ability of the modern IMs to metabolise opiates (morphine, codeine) and antibiotics (berberine), raffinose, stachyose and melibiose indicative of a diet comprising soy, coffee, cacao, dietary flavonoid glycosides derived from tea, citrus and wine and glycerolipid and sphingolipid metabolism. Since these compounds were not present at the time of deposition of the BRS specimen, our results document the evolutionary influence of dietary changes, medicinal treatments and environmental pollutants on the IM taxonomic composition and metabolic capacity of contemporary human populations (Fig. 4e) (Table S8, S9).

### Antibiotic-resistance genes

Our results furthermore confirm reports of antibiotic-resistance genes (ARGs) previously recovered from ethnographic cohorts and archaeological faecal samples (*5, 25, 75*). Following analysis of the ancient and modern resistomes using Resistome Analyser (https://github.com/cdeanj/resistomeanalyzer) (see Methods), we identified a total of 15 functional ARGs, four of which occurs in the BRS IM *i.e.*, BRS4. These include the prokaryotic protein synthesis elongation factor Tu (EF-Tu) (*tufA* and *tufB*), flouro-quinolene-resistant DNA topoisomerase (*parE*) and daptomycin-resistant *rpoB* (Table S11) (Fig. S9). Several bacterial taxa (*e.g.*, *Escherichia coli*, *Staphylococcus aureus* and *Streptomyces collinus*) have duplicate genes for the gene encoding EF-Tu (*tufA* and *tufB*) which confers resistance to the antibiotic kirromycin. These genes are also present in the Ötzi (sample UPLI), Hadza (H9 and H11) and Italian (IT6) datasets. The daptomycin-resistant *rpoB* gene encodes the *β* subunit bacterial RNA polymerase and is the site of mutations that confer resistance to the daptomycin antibacterial agents. Daptomycin-resistant *rpoB* is present in the Ötzi (SI, LPLI and UPLI) and Hadza (H11) datasets, but absent from the Malawian and Italian cohorts. Certain mutations in the RNA polymerase *β* subunit have been found to reduce the susceptibility of methicillin-resistant *S. aureus* (MRSA) for the antibiotics daptomycin and vancomycin (*76*). Several ARGs are limited in occurrence to the modern comparative cohorts. Some, including *gyrA* which confers fluoroquinolone resistance to *Neisseria gonorrhoeae* and *Ureaplasma urealyticum*, triclosan resistance to *Salmonella enterica*, and *pbp4B*, a penicillin-binding protein and the target of *β*-lactam antibiotics and *marA*, are shared only with the Hadza cohort. *Pbp2*, a point mutation in *N. meningitidis* which confers resistance to *β*-lactam, and a penicillin-binding protein found in *Streptococcus pneumoniae*, also occurs only in the Hadza dataset. *Cat*, which confers resistance to broad-spectrum phenicol antibiotics by antibiotic inactivation, occurs only in the Italian cohort. *Cat* has also been detected amongst isolated Amerindians (*5*).

## CONCLUSIONS

In this study, we performed a comprehensive analysis of an ancient palaeo-faecal specimen derived from a 15^th^ century Iron Age (Neolithic) South African Bantu-speaking hunter-agro-pastoralist. Although representative of the IM composition, metabolic capacity and ARG configuration of the distal (*i.e.*, the colon including the cecum, rectum and anal canal) IM of a single human individual, following particular dietary consumption and excreted at a single point in time, the characterisation of an authenticated ancient African Bantu-speaker IM is an important step towards understanding the ancestral (*i.e.*, pre-colonial African) state of the human IM. Our analyses designate a diet atypical of what is generally expected from a Neolithic (Iron Age) IM (*17*), instead comprising taxa indicative of a mixed forager-agro-pastoralist diet, supporting the role of dietary habits in shaping human IM composition.

It must be emphasised that the Neolithic of South Africa is different from the ‘classic’ Near-Eastern Neolithic, as foraging and hunting did play a prominent role in the subsistence configuration of southern African Iron Age communities (*40*). It must also be emphasised that the samples derived from the interior of the BRS specimen (*i.e.*, BRS2, BRS3 and BRS4) ‘clusters’ with the ancient comparative samples (*i.e.*, those derived from Ötzi) in terms of both taxonomic composition (ANOSIM analysis revealed significant differences between the ancient and modern IM samples (*R* = 0.8361; *p* = <0.001) for 731 taxa (Fig. S7) and metabolic capacity (ANOSIM analysis revealed significant differences (*R* = 0.4840; *p* = <0.001) (Fig. 4a), and not with the sedimentary controls (SC1 and SC2) or the modern comparative (*i.e.*, Hadza, Malawian and Italian) cohorts. In contrasting contemporary (Hadza, Malawian and Italian) and ancient (BRS and Ötzi) human IM taxonomic composition and metabolic capacity, it is evident that the changes brought about by modern human dietary composition, exposure to toxic pollutants and the excessive use of antibiotics, almost certainly resulted in positive selection for bacterial taxa involved in specific metabolic IM activities (*69, 77*). While this does not correlate directly with geography, it does exhibit a temporal trend towards the selection of KO genes in direct response to a number of specific changes in human dietary behaviour and environmental interaction and modification.

The IM of the BRS individual represents a unique taxonomic and metabolic configuration not observed in either contemporary African or European populations. Several studies have found that IM composition differs between Western urbanized and indigenous rural populations, and that these dissimilarities frequently correlate with dietary characteristics. In this instance, the diet of the BRS individual, based on hunting, foraging and also agricultural and pastoral resources, differs from the typical Western diet comprising preservatives and food-enhancers, as well as coffee, chocolate, soy, wine and citrus. In terms of modern human hygiene-practices, it has been suggested that regular contact with ‘old friends’ (including both pathogenic and commensal environmental bacteria) is significantly diminished in Western countries (*78*) and that, given our extensive evolutionary history with microbes (*19*), this diminishes the capacity of the modern human IM to mediate allergic reactions and autoimmune and inflammatory diseases. It is evident that the ubiquitous use of antibiotics has altered the properties of formerly commensal bacteria and of the human IM (*79*). We therefore hypothesise that, by compelling commensal IM residents to respond to the introduction of antibiotics prescribed for pathogenic taxa, artificially-introduced dysbiosis has significantly modulated the pathogenic potential of commensal taxa, resulting in long-term deleterious impacts on optimal human IM functioning.

The IM of the BRS individual also provides evidence for recent human IM adaptation to environmental pollutants. The emergence of xenobiotic degradation pathways involved in naphthalene, chloroalkane and chloroalkene, benzoate and xylene degradation is likely a population-wide functional response of the IM to exposure to toxic and foreign compounds that are ubiquitous in industrialized urban environments. The respective enrichment and depletion of several KO genes implicated in the metabolism of morphine and codeine, as well as additives and supplements including pyruvate, l-arginine and beta-alanine, are also indicative of the adaptive capacity of the human IM. Given these modern influences, the contemporary human IM appears to be predisposed towards shifting to a state of dysbiosis. Such altered states of equilibrium frequently result in the pathogenesis of inflammatory diseases and infections, including autoimmune and allergic diseases, obesity and diabetes. The IM of the BRS individual also corroborates the premise that ARGs are a feature of the human IM, regardless of exposure to currently-available commercial antibiotics.

In conclusion, the large number of taxonomically (92.63%) and metabolically (88.51%) unassigned reads in the BRS palaeo-faecal specimen analysed here, granting that this might, to some extent, be a result of aDNA damage and the inability to ‘map’ all reads to existing comparative sequences (*44, 45*), is suggestive of substantial unknown IM taxonomic diversity and metabolic functionality (Table S12). In the future, the identification of these taxa and metabolic capacities might have significant implications for identifying health risks specific to the sub-Saharan African Bantu-speaker population which has increased in prevalence with the adoption of Western diets, medical treatments and exposure to modern pollutants. Given that sub-Saharan Africans living outside Africa exhibits a high prevalence of complex diseases, the comparison of ancient African IM data to those of modern Africans might facilitate not only retrospective disease diagnosis, but also the identification of IM-related risk factors that contribute to the onset of certain diseases.

## METHODS

### Accelerator mass spectrometry (AMS) dating

Two sub-samples derived from the interior of the specimen were subjected to accelerator mass spectrometry (AMS) dating. The samples were pre-treated using the standard acid-base-acid approach (*80*) performed at 70°C. Carbon was oxidised using off-line combustion in the presence of excess CuO and Ag, and the resulting CO_2_ was reduced to graphite through Fe reduction at 600°C (*81*). The graphite was measured at the iThemba LABS AMS facility using Oxalic Acid II and Coal as the reference and background, respectively. We report ^14^C ages in conventional radio-carbon years BP (*i.e.*, before present refers to AD 1950).

### Dietary isotope analyses

To investigate the dietary composition of the BRS individual, one sub-sample derived from the interior of the specimen were subjected to isotopic analyses. The sample was homogenised using a mortar and pestle and then divided in to three sub-samples. The first was left untreated and the second was subjected to a lipid extraction process using a 2:1 chloroform/ethanol solution to remove any lipids present (*82*). The sample was covered with 25 ml of the solution and the mixture agitated in an ultra-sonic bath for 15 minutes and then left overnight. The solvent was then decanted and the sample dried at 70°C prior to weighing for analysis. The third sample was covered with 25 ml 1% hydrochloric acid (HCl) to remove inorganic carbonates (*78*), agitated for 15 min and left overnight. The acid was then decanted and the sample repeatedly washed (6 times) with distilled water before drying at 70°C. Aliquots of the samples weighing between 0.80 mg and 0.90 mg were weighed using a Mettler Toledo MX5 micro-balance. The weighed material was placed in tin capsules that had been pre-cleaned in toluene. All the samples were run in triplicate. Samples for isotopic analyses were combusted at 1020 °C using an elemental analyser (Flash EA 1112 Series) coupled to a Delta V Plus stable light isotope ratio mass spectrometer via a ConFlo IV system (Thermo Fischer, Bremen, Germany), housed at the Stable Isotope Laboratory, University of Pretoria. Two laboratory running standards (Merck Gel: δ^13^C = −20.26‰, δ^15^N=7.89‰, C%=41.28, N%=15.29 and DL-Valine: δ^13^C = −10.57‰, δ^15^N=-6.15‰, C%=55.50, N%=11.86) and a blank sample were run after every 11 unknown samples. Data corrections were performed using the values obtained for the Merck Gel during each run and the values for the DL-Valine standard provide the ± error/precision for each run. The precision for the BRS analyses was > 0.04‰ and 0.05‰ for nitrogen and carbon respectively. These running standards are calibrated against international standards, *i.e.*, National Institute of Standards and Technology (NIST): NIST 1557b (bovine liver), NIST 2976 (muscle tissue) and NIST 1547 (peach leaves). All results are referenced to Vienna Pee-Dee Belemnite for carbon isotope values, and to air for nitrogen isotope values. Results are expressed in delta notation using a per mille scale using the standard equation ‘δX(‰) = [(R_sample_/R_standard_)-1]’ where X= ^15^N or ^13^C and R represents ^15^N/^14^N or ^13^C/^12^C respectively.

### Intestinal parasite detection

In addition to genomic taxonomic profiling, we also performed microscopic analysis to determine the incidence of intestinal parasitic helminths and protozoa. The extraction protocols applied in palaeo-parasitology used to extract parasitic markers (*i.e.*, eggs or oocysts) typically entails rehydration, homogenisation and micro-sieving (*83*). The sub-sample (∼5 g) was placed in a rehydration solution comprising 50 ml 0.5% TSP solution and 50 ml 5% glycerinated water for seven days, after which it was ground and passed through an ultrasonic bath for 1 minute. The sample was then filtered in a sieving column comprising mesh sizes of 315 µm, 160 µm, 50 µm and 25 µm in aperture diameter. Because of the typical size of most intestinal parasite eggs range from 30 µm to 160 µm long and 15 µm to 90 µm wide, only the two last sieves (*i.e.*, 50 µm and 25 µm) were subjected to microscopic analyses.

### Scanning electron microscopy (SEM)

We immobilised 0.5 g palaeo-faecal material on double-sided carbon tape (SPI supplies). Excess loose particles were blown off with compressed argon gas and coated with gold using an Emitech K450X sputter coater (Quorum Technologies, UK). SEM images were acquired on a Zeiss Ultra Plus Field Emission Scanning Electron microscopes (Carl Zeiss, Oberkochen, Germany), at an accelerating voltage of 1kV.

### Ancient DNA extraction and library preparation

All pre-PCR amplification steps were carried out in aDNA laboratories at the Centre for GeoGenetics applying established aDNA protocols (*84*). Extractions for shotgun metagenome sequencing were carried out using a phenol-chloroform- and kit-based extraction protocol optimized for ancient sedimentary and faecal samples. This entailed dissolving a total of ∼16 g of palaeo-faecal matter in 40 ml digestion buffer. Seven libraries, comprising BRS1 (SC1), BRS2, BRS3, BRS4, BRS5 (SC2) and two negative controls (*i.e.*, an extraction and library preparation control referred to as ‘E-LPCs’) were constructed using the NEBNext DNA Library Prep Master Mix for 454 (E6070) and sequenced on an Illumina HiSeq 2500 platform at the Danish National High-Throughput DNA Sequencing Centre. The libraries were sequenced twice to improve DNA recovery, producing 14.563 Gbp of data (17,979,669 reads) for BRS1, 29.201 Gbp (36,050,926 reads) for BRS2, 0.709 Gbp (8,754,692 reads) for BRS3, 95.323 Gbp (117,683,541 reads) for BRS4 and 9.54 Gbp (110,485,402 reads) for BRS5. The merged E-LPCs yielded 2.946 Gbp of data (3,637,328 reads) (Table S13).

### Sequence processing and microbial taxonomic profiling

The potential for retrieving ancient IM data from palaeo-faecal remains is confounded by technical and biological variables (*14, 44, 45*). In technical terms, the detection of ˃90 microbial genera in DNA extraction and library preparation controls suggest that reagent and laboratory contamination can influence sequence-based IM analyses (*84, 85, 86*). The choice of DNA extraction protocols can also impact metagenomic compositional profiles (*87*). In biological terms, IM research generally focuses on the microbial community of the large intestine as expressed in stools, despite the fact that the 6.5 meter human digestive tract consists of three organs, *i.e.*, the stomach, small intestine and large intestine (*6*). Since microbial communities change along the length of the GI, differences exist between oral, intestinal and faecal taxonomic profiles in both modern (*88*) and ancient (*25*) instances. Moreover, and besides the influence on IM taxonomic composition of diet (*57*), age (*61*), seasonal variation (*89*) and host immuno-modulation (*90*), stool consistency also influence IM taxonomic composition (*91*). Differences in taxonomic composition between the cores and cortices of specimens have also been documented, with larger proportions of soil-derived taxa present in the cortices (*24*). In post-depositional terms, the retrieval of ancient IM data is confounded by on-going microbial activity and also environmental contamination (*92, 93, 94*). Instances of reverse contamination, *i.e.*, from a faecal specimen into the surrounding sediment, are also probable as the exchange of microbes between a stool and the surrounding sediment would certainly occur.

With these concerns in mind, taxonomic profiling was preceded by several data pre-processing steps. First, raw sequence reads were processed to remove all Illumina PhiX spikes, human reads and all exact duplicate reads present in the extraction (*n* = 1) and library preparation (*n* = 1) negative controls (E-LPCs) using BBDuk (*95*). Second, barcodes, adapters, reads shorter than 25 base-pairs (bp) and ‘quality score’ <25 were removed from the dataset (*96, 97*) using AdapterRemoval V2 (*98*). Taxonomic binning was then performed via comparison of the shotgun reads with the BLASTn v2.2.31+ NCBI *nt* database (*99*). Taxa were identified using MEGAN CE v6.10.10 (*97*) by using the weighted lowest common ancestor (‘wlca’) option and the default percent-to-cover value setting (‘80’) with parameter values set as follows: min. bit score: 50, expect value (*e*-value): 1.0*e*-10, top percent: 10, min. support: 10 and min complexity: 0.45. Species identifications were based on significant hits (bit score ≥50) and on MEGAN parameters established at ‘identities’: 97%, ‘positives’: 100% and no (0%) ‘gaps’. Comparisons of BRS IM sequence reads with those derived from other (comparative) IMs was performed by the subsampling in of the reads to the lowest number of reads present in any sample (*i.e*., BRS1 (SC1) with 56,682 filtered sequence reads).

### Ancient DNA authentication

Molecular damage following death is a standard feature of all aDNA molecules. The accumulation of deaminated cytosine (uracil) within the overhanging ends of aDNA templates results in increasing cytosine (C) to thymine (T) misincorporation rates toward read starts, with matching guanine (G) to adenine (A) misincorporations increasing toward read ends in double-stranded library preparations (*100, 101*). MapDamage (*38, 39*) is widely used to determine the incidence of cytosine (C) to thymine (T) and guanine (G) to adenine (A) substitution rates at the 5’-ends and 3’-ends of strands. MapDamage is not however optimised for ancient samples lacking high genome coverage that would permit the identification of all possible misincorporations (*101*), and DNA damage patterns cannot be calculated for taxa with insufficient read counts, *i.e.*, <150 (*101, 102*). In addition, DNA fragmentation rates vary according to environmental conditions and the types of organisms involved (*103, 104*), often resulting in ‘alternative’ damage patterns (*105–107*). We therefore also validated the antiquity of putatively ancient taxa by a statistical method that compares post-mortem damage patterns indicative of the cytosine to thymine (C-T) substitutions at the 5’ ends of sequence reads (*37, 108*). High-quality filtered reads were aligned to comparative genomes (Table S13) using BWA (-n 0.02; −l 1024) (*98*) and duplicate sequence reads were removed using the Picard tools script MarkDuplicates (https://broadinstitute.github.io/picard/). Resulting alignments were used to perform statistical DNA damage estimation analyses which entailed the calculation of goodness-of-fit *p*-values (*p* = <0.05) indicative of significant cytosine to thymine (C-T) substitutions at the 5’ ends of sequence reads using PMDtools (https://omictools.com/pmdtools-tool) (*37*).

The authentication of aDNA sequence reads was furthermore based on the comparison of reads derived from negative (*i.e.*, extraction and library preparation) controls (*84, 93, 94, 100, 101*). In addition to the detection of >90 microbial taxa derived from reagents and laboratory contamination (*85, 86*), the probability that negative controls can become cross-contaminated during sample processing (*103*) complicates the authentication process. To establish the antiquity of microbial taxa occurring in the E-LPCs (*i.e.*, *Arthrobacter*, *Blautia*, *Klebsiella*, *Lactobacillus*, *Prevotella* and *Ruminococcus*), we compared the read yields in the BRS specimen with those in the E-LPCs. Since aDNA sequences are shorter than those derived from modern organisms, most frequently via contamination (*104–108*), DNA read length was also used as criteria in the authentication process. This criterion is particularly relevant in instances that exclude the use of bleach for the removal of contaminants (*84*), as is the case here. Lastly, evaluating ecological conformity, *i.e.*, excluding DNA reads that are derived from either non-indigenous taxa, foreign contaminants or false-positive identifications, such as *Apteryx* (kiwi), *Cyprinus* (carp), *Oncorhynchus* (salmon) and *Oryza* (rice) was used to assess taxonomic community composition for biological plausibility (*93, 94*). The authenticity (*i.e.*, prehistoric provenience) of microbial and macrobial sequence-derived taxa was therefore evaluated according to 1) validating the existence of C-T and G-A substitution rates, 2) statistical aDNA sequence damage estimation, 3) comparison to negative DNA E-LPCs, 4) DNA read length characteristics and 5) ecological conformity.

### Functional metabolic profiling

To discern the metabolic pathways associated with the dietary and environmental factors characteristic of the Bantu-speaking forager-agro-pastoralist in question, we identified reads assigned to functional genes in the shotgun metagenome dataset. To ascertain the presence of microbial groups either positively or negatively correlated with specific metabolic profiles, we determined functional categories based on DIAMOND v0.36.98 BLASTx comparisons to the NCBI non-redundant protein database. The GI accessions were used to identify the Kyoto Encyclopaedia of Genes and Genomes (KEGG) orthologies (https://www.genome.jp/kegg/) in MEGAN CE v6.10.10 (97) by using the weighted lowest common ancestor (‘wlca’) option and the default percent-to-cover value setting (‘80’) with the parameters as follows: ‘*e*-value cut-off’: 1.0*e*-4, ‘top percent’:10, ‘min support’:10, ‘min complexity’:0.45 and ‘identity’: 97%. Comparisons of KEGG orthologies was performed following the subsampling to the lowest number of reads for any sample (*i.e.*, BRS1 (SC1) with 56,682 sequence reads).

### Comparative IM datasets

To gain insight into the taxonomic and functional (metabolic) differences between the BRS IM (*n* = 4) and other ancient and modern (including ‘traditional’) IMs, we compared BRS with data derived from the shotgun metagenome sequencing of Malawian agro-pastoralists (*51*) (MG-RAST http://metagenomics.anl.gov/ accession number ‘qiime:621’), Tanzanian Hadza hunter-gatherers (*26*) (NCBI SRA SRP056480 Bioproject ID PRJNA278393), contemporary Italians (*26*) and the Copper Age (dated to *c*. 3,250 BC) Alpine (Tyrolean) Iceman (Ötzi) (*109*) (European Nucleotide Archive accession number ERP012908). For comparison, four metagenome samples (comprising two males and two females) were randomly selected from each of the comparative cohorts (*n* = 12).

### Detecting antibiotic-resistance genes

The resistome is defined as the complete set of antibiotic resistant genes (ARGs) presented in a microbial community, which is important for understanding the proliferation of pathogen antibiotic resistance (*26*). Sequence reads were aligned to the MEGARes database (*110*) using the Burrows-Wheeler Alignment tool (BWA) (*111*). The BRS IM resistome was analysed using Resistome Analyser (https://github.com/cdeanj/resistomeanalyzer) by applying the default threshold of ‘80’ to determine the gene significance and in order to decrease false positive gene identifications. Relative abundance for each of the resistance genes was calculated and sub-sampled to the lowest number of sequences (*i.e.,* the Ötzi ‘small intestine’ (Ötzi_SI_F) comprising 2,377 sequences) in any sample (*i.e.*, 28,524 sequences for a total of twelve samples). Because of low numbers of gene assignments, seven samples were excluded from the analysis.

### Statistical analyses

Statistical analysis of the ancient and modern IM samples was performed by filtering the taxa and KEGG orthologies for >3 occurrences in at least 20% of the samples. The data was Hellinger-transformed and the Bray-Curtis dissimilarity matrix was used for the *vegdist* and *anosim* functions in the VEGAN (https://www.rdocumentation.org/packages/vegan/versions/2.4-2) package of R. The ordination plots were generated in the Phyloseq package (*112*) of R. Functional (metabolic) group significance tests were performed in Qiime v1.9.1 package (*113*) for the ancient and modern comparative cohorts and only gene categories with significant *p*-value (*p* ˂ 0.05) were included in the analyses. The *p**-***values were corrected using false discovery rate (FDR) (*114*) to corrected *p* values (*q*) and Kruskal-Wallis values (*H*) to determine the significance of differences between samples. Hierarchical clustering using complete-linkage based on Spearman’s correlations was performed and visualised in R using ‘gplots’ (https://www.rdocumentation.org/packages/gplots/versions/3.0.1). Heat-maps were generated using Spearman’s correlation and complete linkage method for microbial taxa and antibiotic resistance genes. Taxa were filtered for occurrence of >3 in at least 20% of the samples, and ARG data was filtered for the occurrence of >5 in at least 20% samples. Heat-map for the KEGG orthologies linked to 24 taxa (1,487 categories) was produced by using Spearman’s correlation and Ward-linkage method.

## Acknowledgements

We thank Clemens Weiß (University of Tübingen), Fredrik Seersholm (University of Copenhagen) and Pedro Lebre (University of Pretoria) for informative discussions and analytical support.

## Author contributions

RFR, AJH, JBR and SV conceived the study and composed the manuscript. SV performed the bioinformatic and statistical analyses and RFR and SV generated the figures. JBR, ARI, TBB and RFR performed aDNA extractions, library preparations and sequencing. GP and AV provided access to the analysed specimen. GH performed the isotope analyses, and SW performed the 14C dating. MLB carried out intestinal parasitic analyses. MP and RFR performed SEM analyses. EW, DAC and YVDP provided funding and access to analytical fascilities. AJH, JBR and RFR conceived the analytical protocol and AJH supervised the experimental protocol. All authors contributed to the completion of the final manuscript.

## SUPPLEMENTARY MATERIALS

### Supplementary Figures

**Fig. S1.**
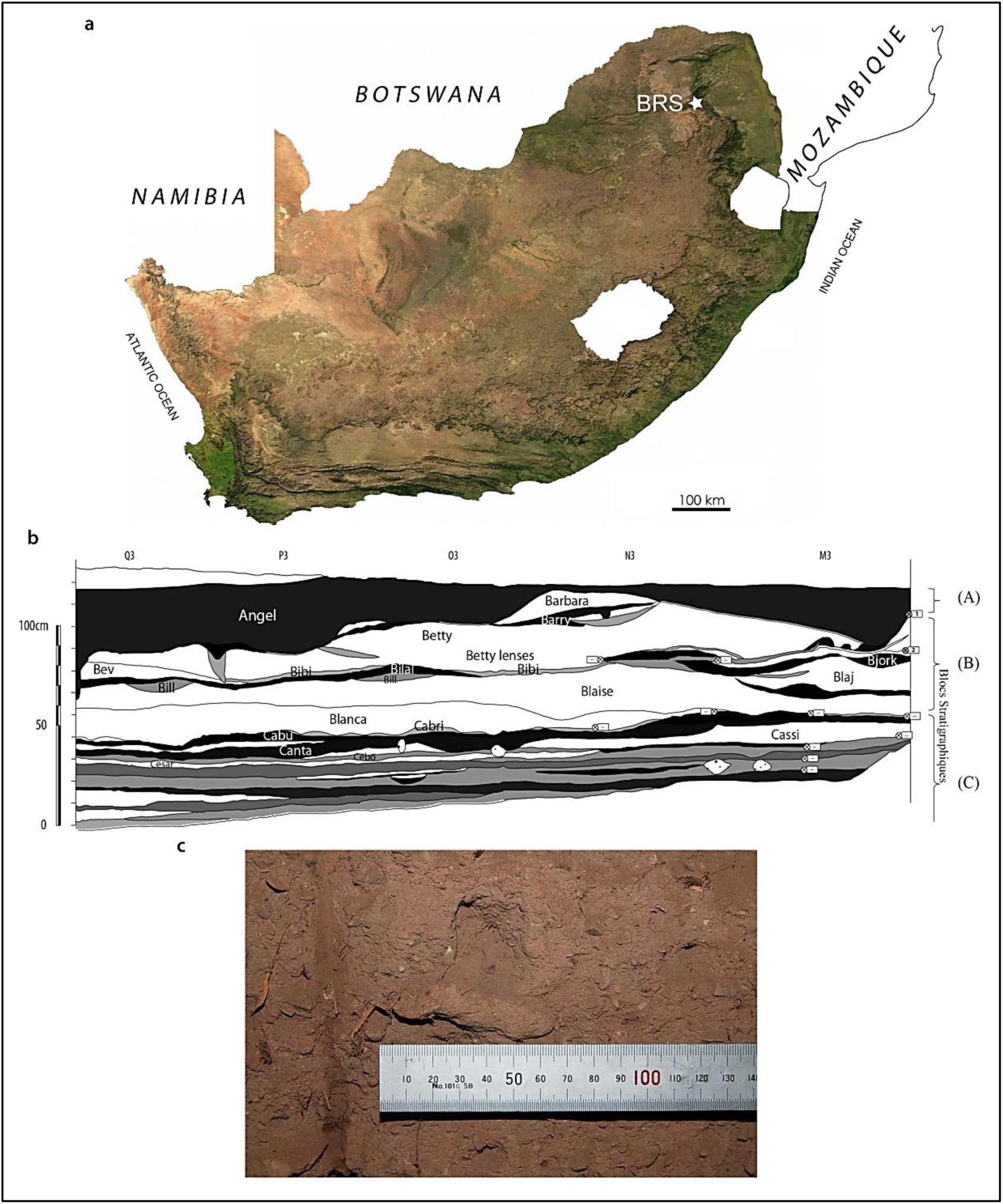
The provenience of the BRS palaeo-faecal specimen. The location of Bushman Rock Shelter (BRS) in Limpopo Province, South Africa (**A**), stratigraphic profile of the Iron Age occupation level from which the specimen derives and which comprise the upper layer of the rock-shelter (Layer 1) situated in excavation ‘Block A’ and designated ‘Angel’ (**B**) and the palaeo-faecal specimen *in situ* in the exposed (excavated) section prior to removal from the deposit (**C**).

**Fig. S2.**
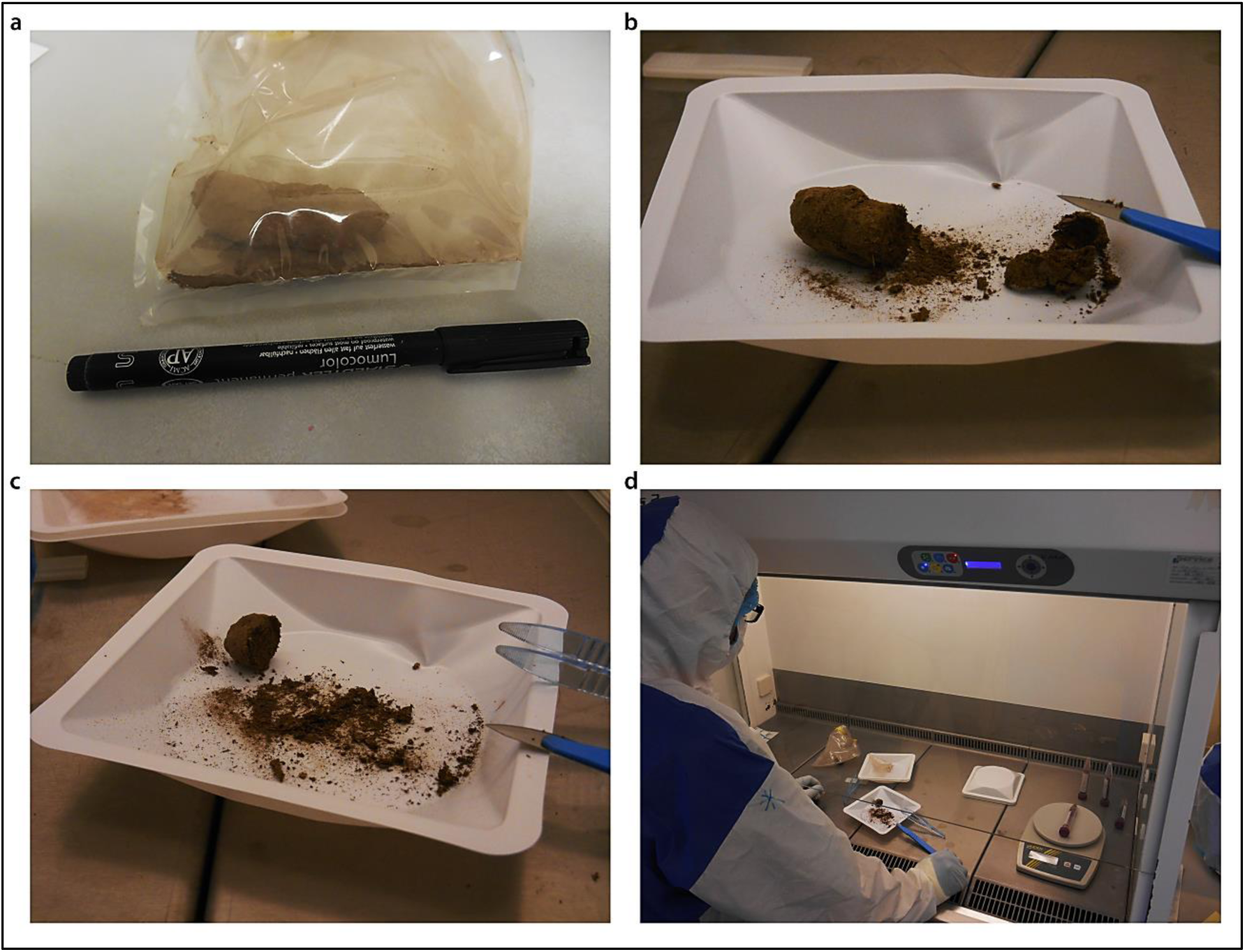
Processing (sub-sampling and aDNA extraction) protocol applied to the BRS palaeo-faecal specimen. The frozen specimen was first removed from the sealed packaging (**A**), after which the outer surface or cortex (∼5mm) was removed with a scalpel (**B**) and subsequently sub-sampled for radiocarbon (14C) dating and isotopic and microscopic intestinal parasitic analyses, preserving one-sixth of the specimen (at −20°C) as a voucher sample (**C**). Sub-sampling and aDNA extraction and library preparation was performed in the ‘clean’ ancient DNA laboratories at the Centre for GeoGenetics, University of Copenhagen (Denmark) (**D**).

**Fig. S3.**
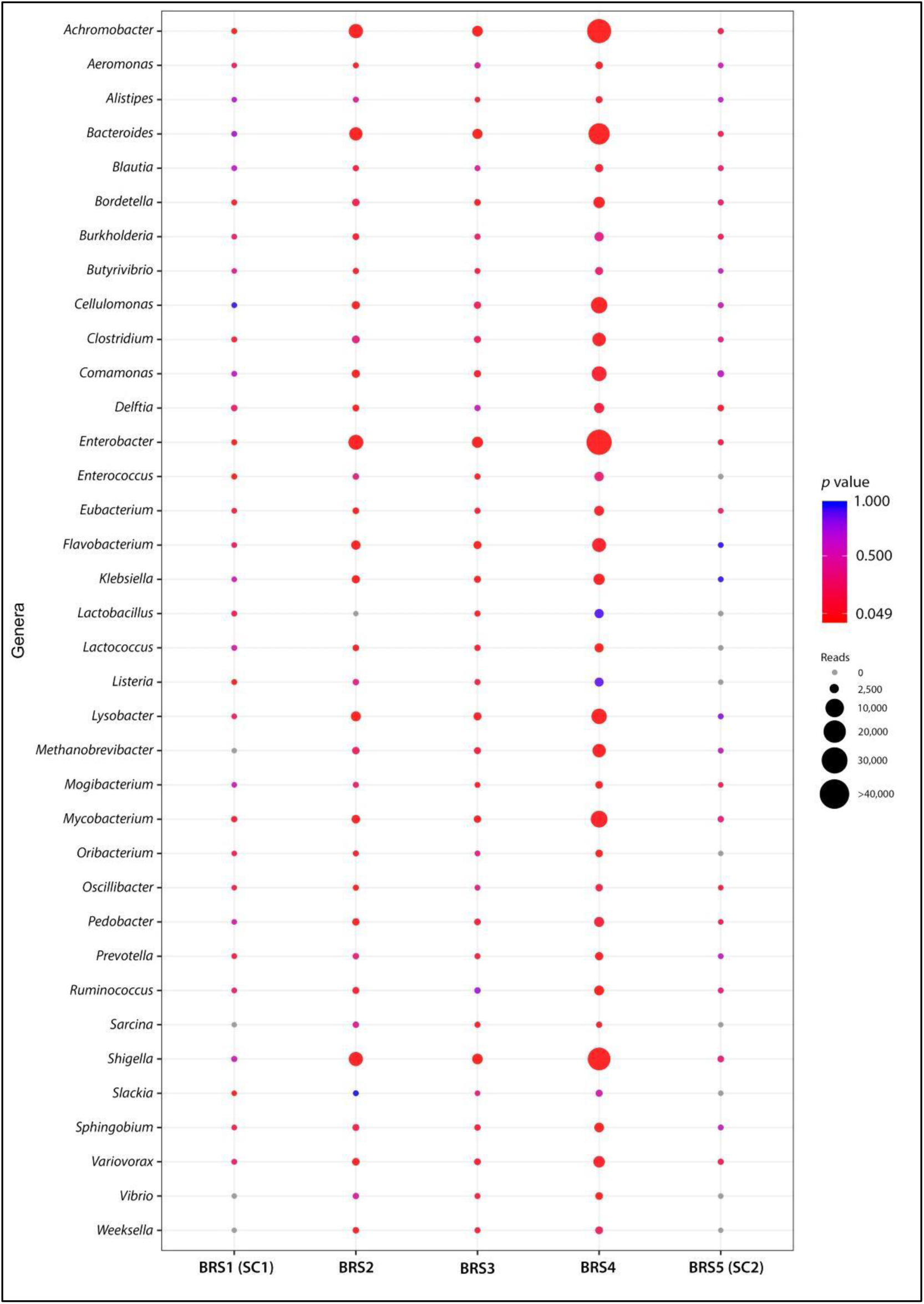
Dot-plot based on the alignment of high-quality DNA sequence reads indicating the occurrence of statistically-significant C-T *p*-values calculated for commensal and pathogenic taxa detected in the BRS specimen (BRS2, BRS3 and BRS4) and the sedimentary controls (BRS1 or ‘SC1’ and BRS5 or ‘SC2’). Circle sizes and colours represent mapped read counts and *p*-value significance, respectively (see legend and scale).

**Fig. S4.**
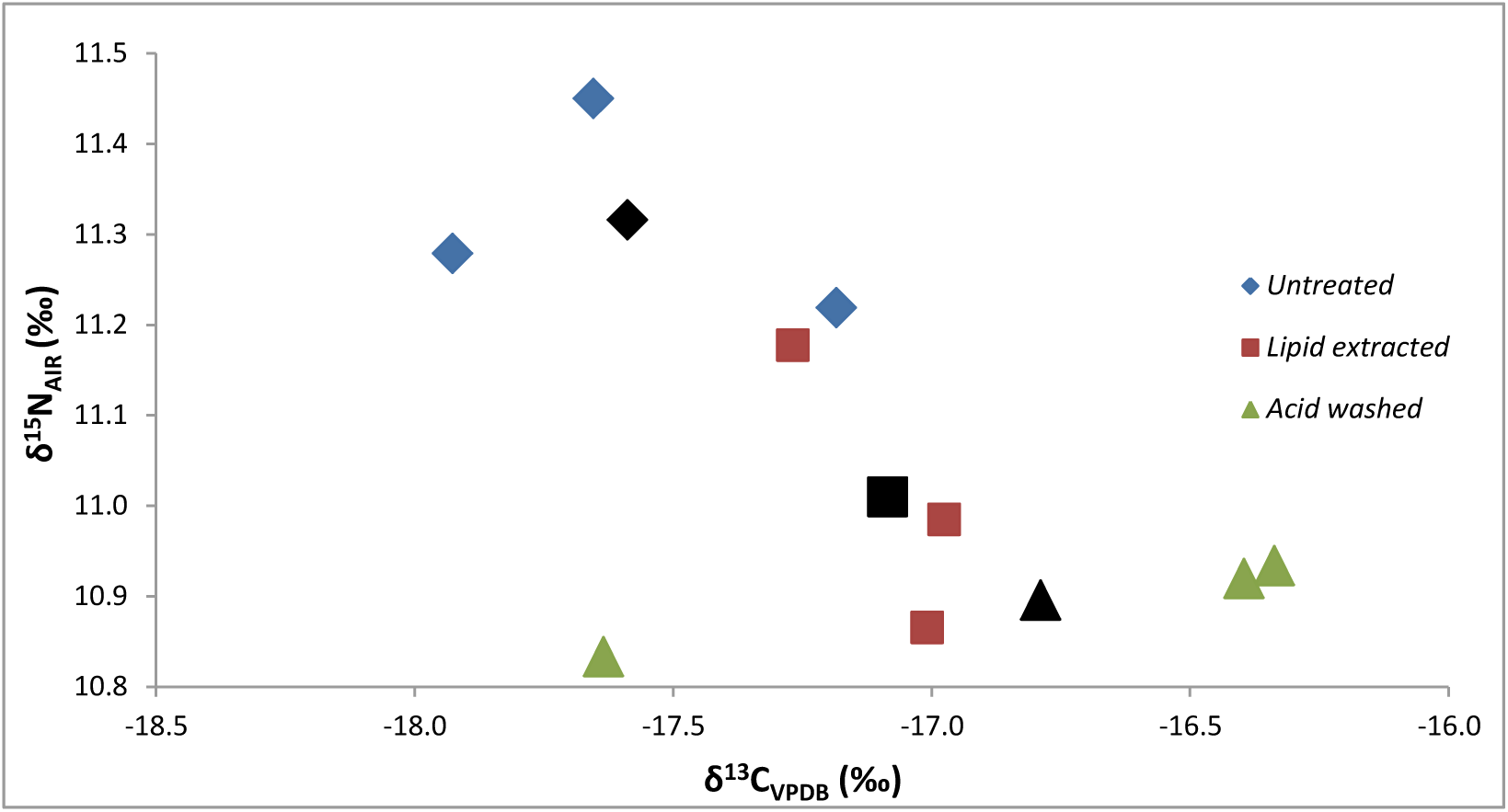
Biplot of δ^13^C and δ^15^N stable isotope ratios obtained for the BRS specimen. The various pre-treatments (*i.e.*, the acid wash and lipid extraction) had some effect on both δ^13^C and δ^15^N values. The lipid extraction (2:1 chloroform/ethanol) removed traces of lipids with carbon isotope values becoming less negative, but nevertheless suggesting a mixed C4- (mostly) and C3-based meal. The results for the solvent residue reflect a geological signal most likely from the shelter’s sediments. Average values for ‘untreated’, ‘lipid extracted’ and ‘acid washed’ are indicated in corresponding black markers.

**Fig. S5.**
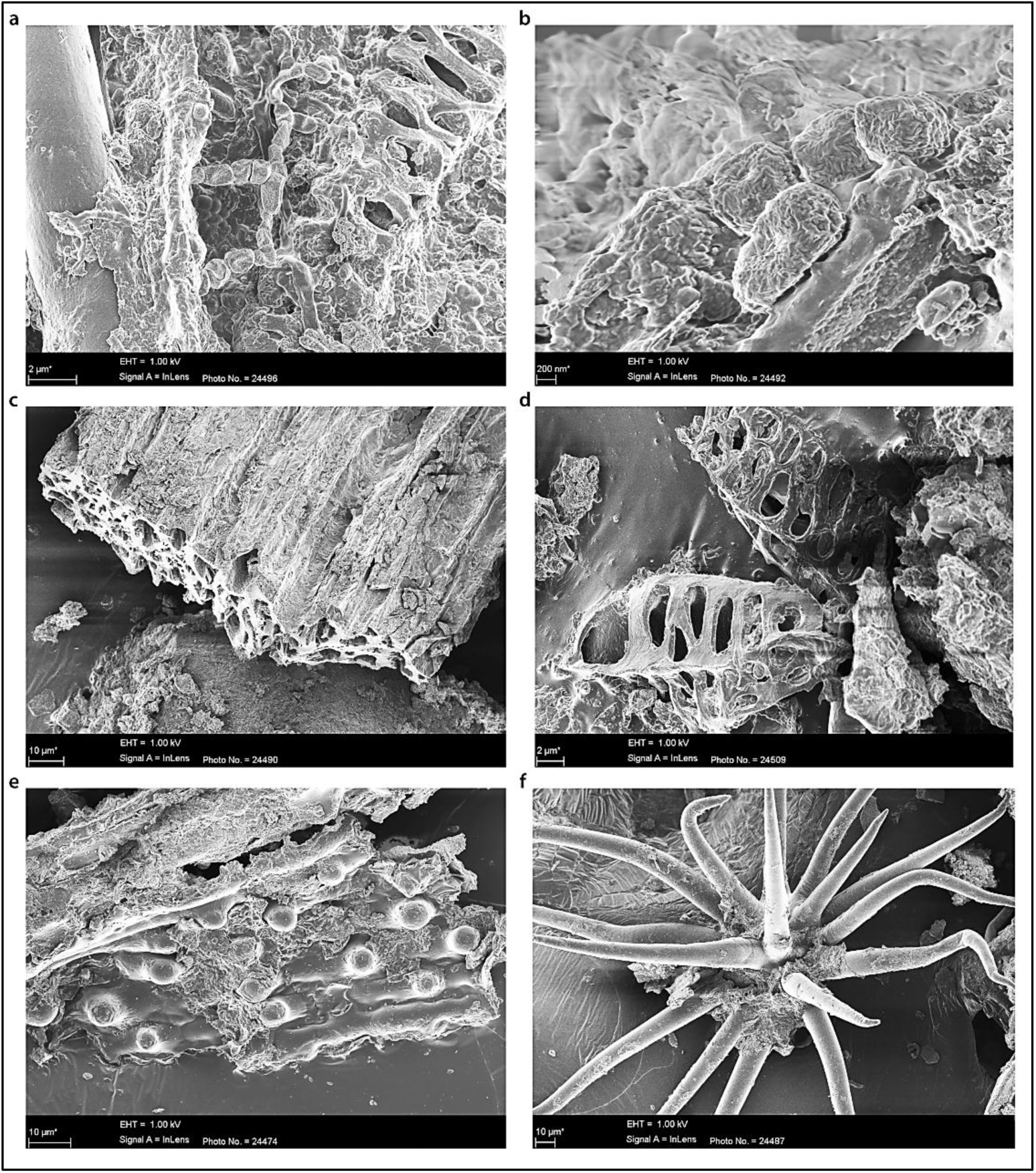
Although scanning electron microscopy (SEM) analyses did not result in the detection of parasitic remains (*e.g.*, eggs), it did result in the detection of desiccated bacterial cells (**A** and **B**), degraded plant fragments (**C**, **D** and **E**) and unidentified saprophytic organisms (**F**).

**Fig. S6.**
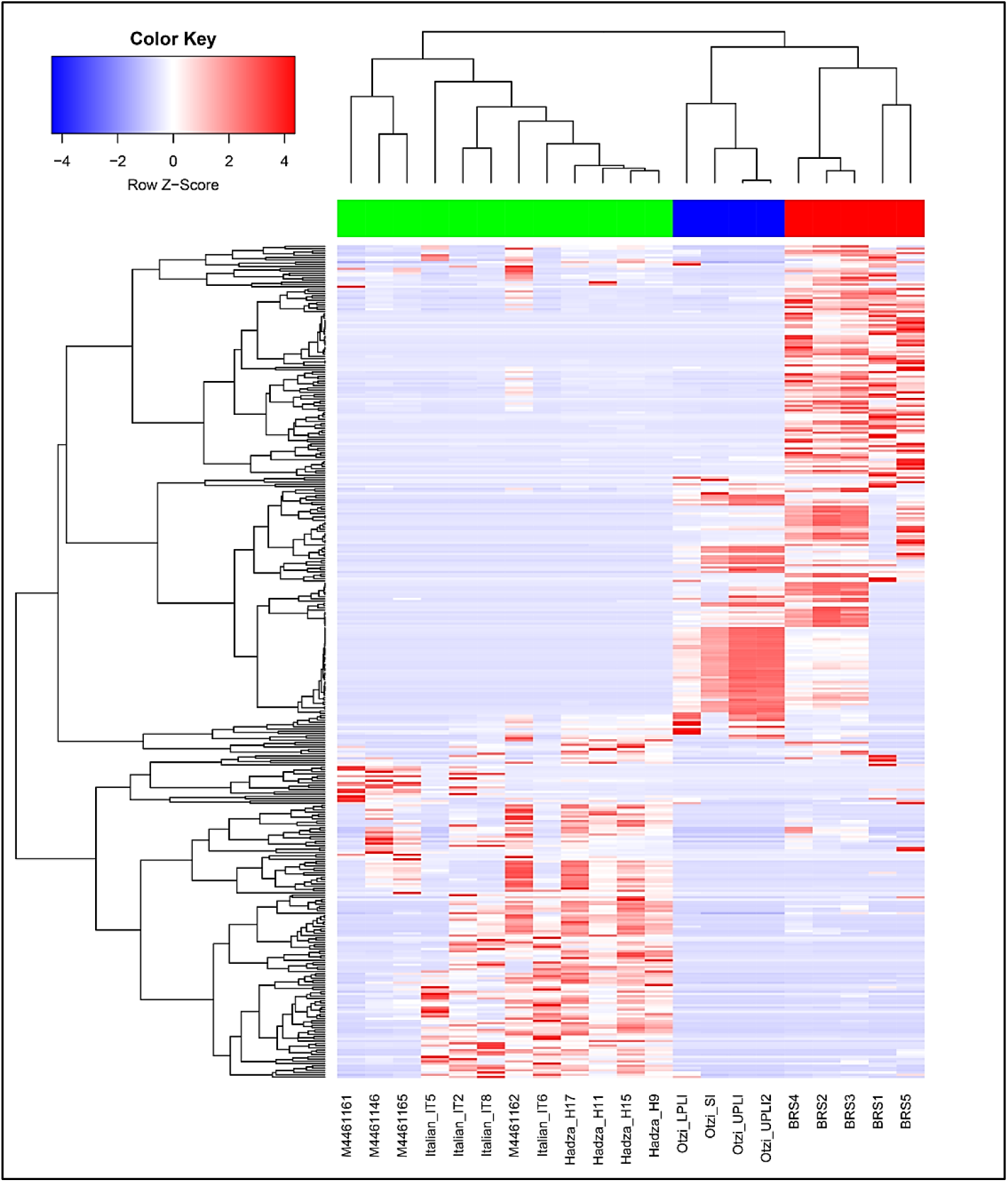
Heat-map comparing differences in general taxonomic community structure for BRS (1, 2, 3, 4 and 5) with the modern (Italian), ethnographic (Hadza and Malawian) and ancient (Ötzi) IM datasets. Hierarchical clustering using complete linkage based on Spearman’s correlation, produced a clear separation between ancient (BRS and Ötzi) and modern (Italian, Hadza and Malawian) populations. ANOSIM analysis revealed significant differences between the ancient and modern IM samples (*R* = 0.6111; *p* = < 0.05) in the taxonomic categories of 731 taxa. Taxa were filtered for occurrence of >3 in at least 20% of the samples.

**Fig. S7.**
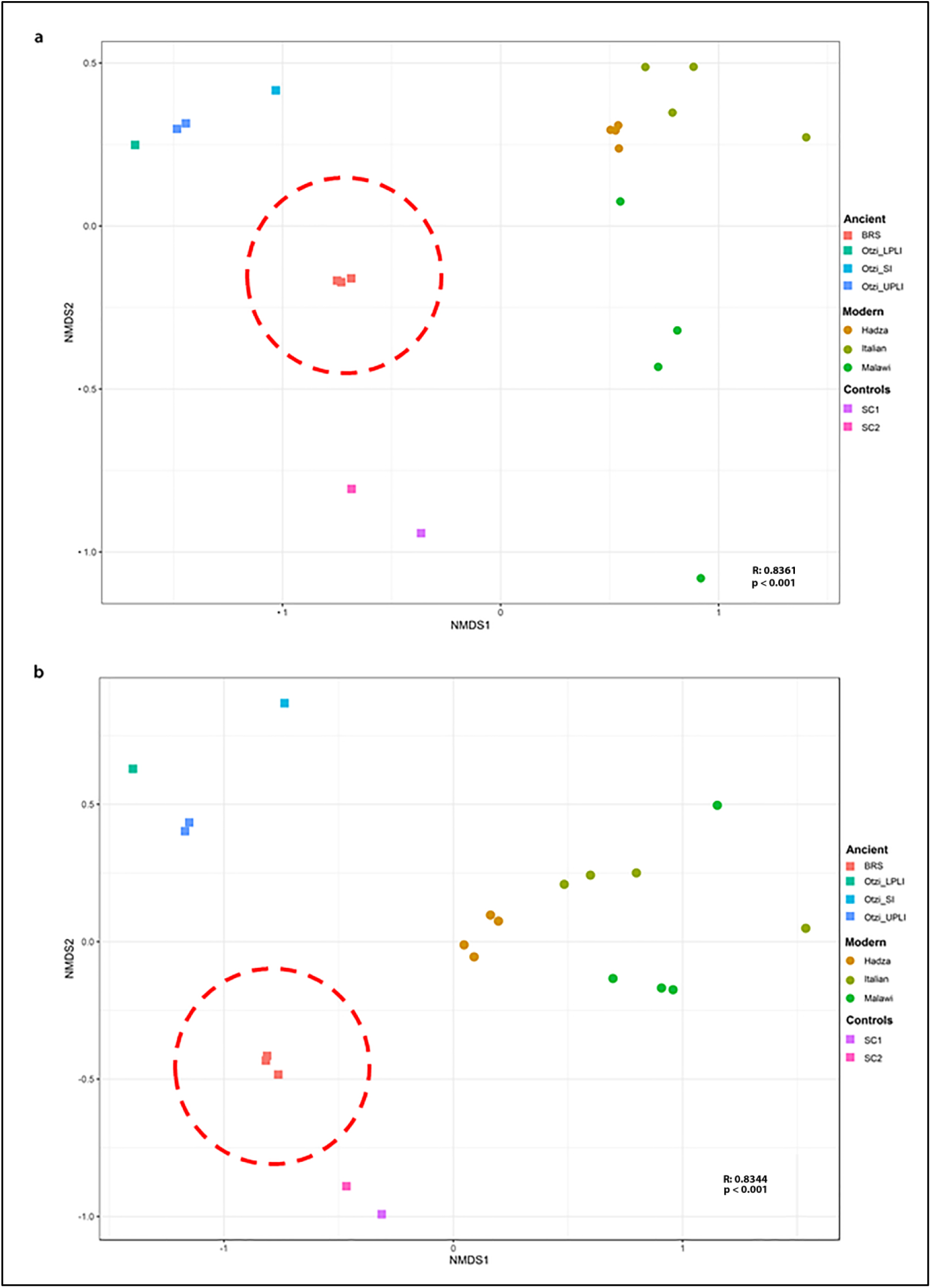
Weighted (**A**) and un-weighted (**B**) Bray-Curtis non-metric multi-dimensional scaling (NMDS) plots comparing the use of ‘relative abundance’ (indicated in **A**) and ‘presence-absence’ data (indicated in **B**) as measures of taxonomic representation in the BRS specimen and in the modern (Italian), ethnographic (Hadza and Malawian) and ancient (Ötzi) IM datasets. Weighted (**A**) (as shown Fig. 1C and based on the ‘relative abundance’ of identified IM taxa) (ANOSIM *R* = 0.6111; *p* = <0.001) and un-weighted (**B**) Bray-Curtis analysis (based on ‘presence-absence’ of identified IM taxa) exhibit corresponding differences between the ancient and modern IM cohorts (ANOSIM *R* = 0.8166; *p* = <0.001).

**Fig. S8.**
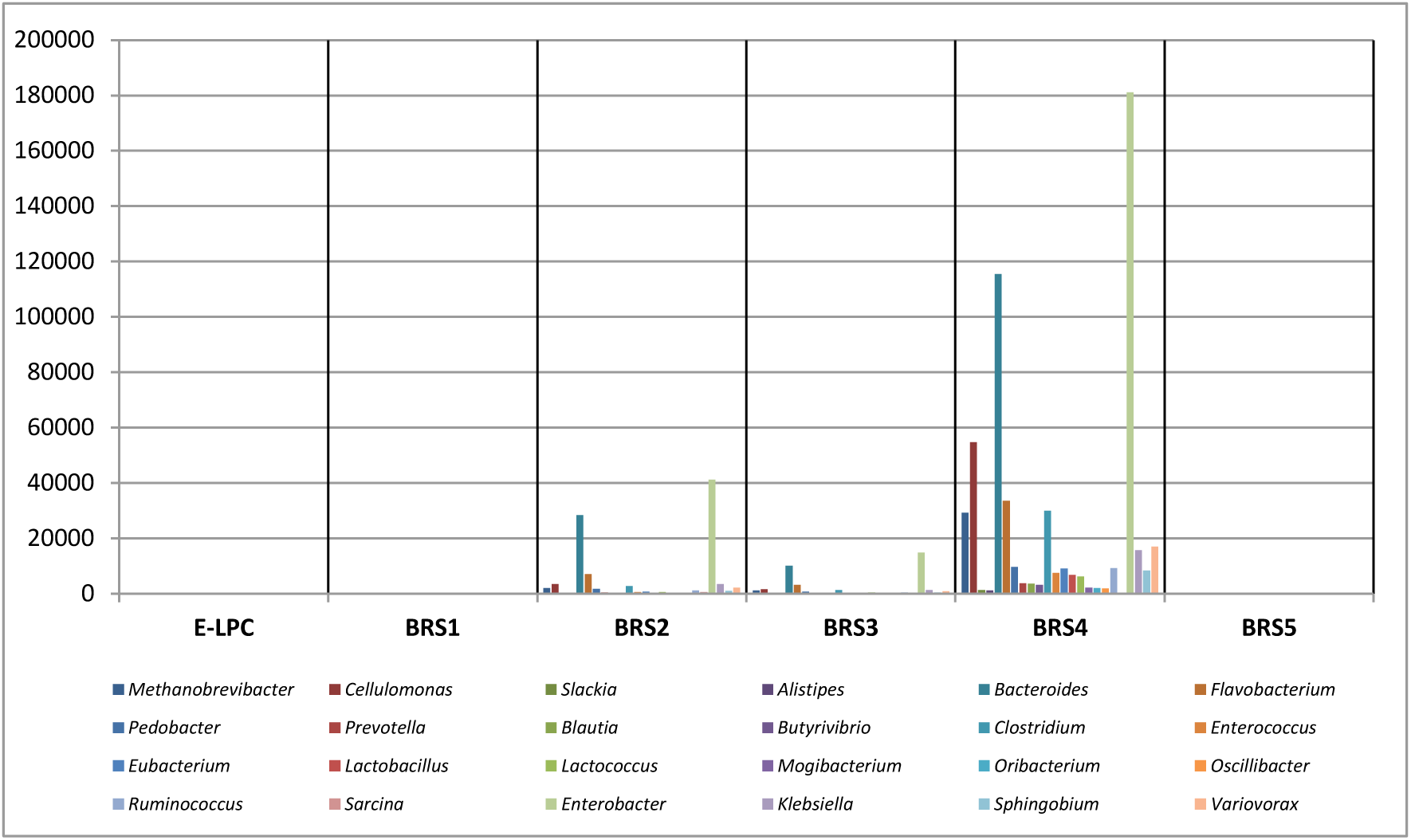
Bar-plot demonstrating that the surrounding sedimentary matrix (*i.e.*, samples BRS1 ‘SC1’ and BRS5 ‘SC2’) and the DNA extraction (*n* = 1) and library preparation (*n* = 1) negative controls (‘E-LPCs’) are not significant sources of the twenty-four authenticated ancient microbial taxa identified in the palaeo-faecal specimen (*i.e.*, BRS2, BRS3 and BRS4). Reverse contamination, *i.e.*, from the palaeo-faecal specimen into the surrounding sediment, is most likely responsible for the incidence of the very low numbers of reads for IM-specific taxa in the surrounding sedimentary matrix (*i.e.*, BRS1 and BRS5).

**Fig. S9.**
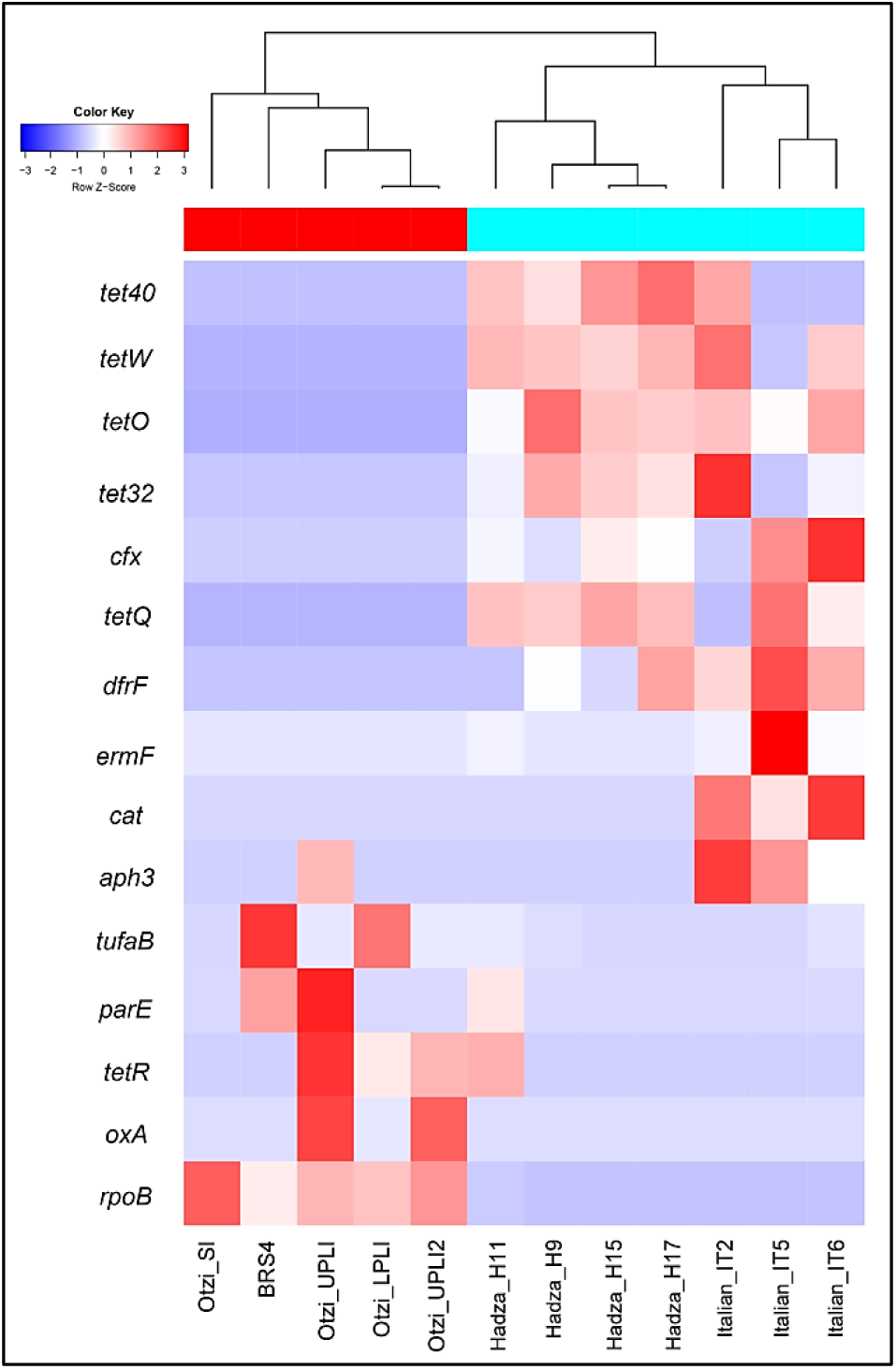
Heat-map indicating the presence of fifteen functional ARGs identified in the analysed datasets, four of which occurs in the BRS IM, including the prokaryotic protein synthesis elongation factor Tu (EF-Tu) (*tufA* and *tufB*), flouro-quinolene-resistant DNA topoisomerase (*parE*) and daptomycin-resistant *rpoB*. ARG categories were filtered for occurrence of >5 in at least 20% of samples.

### Supplementary Tables

**Table S1.** Mapped DNA sequence reads for environmental- and subsistence-related taxa detected in this study. Statistically-significant (*i.e.*, verified ancient) C-T *p*-values are indicated in bold text. Analyses were performed using high-quality filtered read alignments against NCBI reference genomes. DNA damage estimation analyses was performed using PMDtools (‘C-T *p*-values’).

**Table S2.** Information concerning the two direct radiocarbon (^14^C) Accelerator Mass Spectrometry (AMS) dates generated from two sub-samples taken from within the BRS palaeo-faecal specimen.

**Table S3a.** Processing protocol and results for isotope analyses of samples derived from the BRS specimen indicating the relative proportions of C3 and C4 dietary contributions.

**Table S3b.** Results for isotope analyses (Merck standard) of samples derived from the BRS specimen indicating the relative proportions of C3 and C4 dietary contributions.

**Table S3c.** Results for isotope analyses (DL-Valine standard) of samples derived from the BRS specimen indicating the relative proportions of C3 and C4 dietary contributions.

**Table S4.** Assignment of abundance of bacterial taxonomic categories to the BRS and ancient (Ötzi) and modern (Italian, Hadza and Malawian) comparative samples based on *p*-value (*p* = <0.05) designation. Group significance assignments were obtained via Qiime v1.9.1.

**Table S5.** Information concerning DNA sequence read-length distribution for taxa identified in this study. Read-length distributions were calculated from BWA alignments and BAM files.

**Table S6.** Relative abundance of eighteen significant KEGG pathways detected in BRS and the ancient (Ötzi) and modern (Italian, Hadza and Malawian) comparative IM cohorts.

**Table S7.** Enrichment and depletion of KO metabolic gene categories in the BRS and ancient (Ötzi) and modern (Italian, Hadza and Malawian) comparative IM sample cohorts based on *p*-value (*p* = <0.05) designation. Group significance analyses were performed using Qiime v1.9.1.

**Table S8.** Enrichment and depletion of KO metabolic gene categories in the ancient (BRS and Ötzi) and modern (Italian, Hadza and Malawian) comparative IM sample cohorts based on false discovery rate (FDR) corrected *p*- values (*q* = <0.05).

**Table S9.** Enrichment and depletion of KO metabolic gene categories in the ancient and modern comparative IM sample cohort as calculated for the twenty-four authenticated ancient IM taxa.

**Table S10.** Comparison of relative abundance of antibiotic resistance genes in the BRS and the ancient (Ötzi) and modern (Italian, Hadza and Malawian) comparative IM cohorts.

**Table S11.** Raw and filtered high-quality sequence read counts as related to the BRS and the ancient (Ötzi) and modern (Italian, Hadza and Malawian) comparative IM datasets.

**Table S12.** Information concerning the comparative NCBI genomes used during this study.

## REFERENCES

1. J. Lederberg, A. T. McCray, ‘Ome sweet’ omics: A genealogical treasury of words. Scientist 15, 8 (2010).

2. C. Warinner, C. Speller, M. J. Collins, C. M. Lewis, Ancient human microbiomes. J Hum Evol. 79, 125–136 (2015).

3. E. Thursby, N. Juge, Introduction to the human gut microbiota. Biochem J. 474, 1823–1836 (2017).

4. L. W. van den Elsen, H. C. Poyntz, L. S. Weyrich, W. Young, E. E. Forbes-Blom, Embracing the gut microbiota: The new frontier for inflammatory and infectious diseases. Clin Transl Immunology 6, doi: 10.1038/cti.2016.91 (2017).

5. J. C. Clemente. E. C. Pehrsson, M. J. Blaser, K. Sandhu, Z. Gao, B. Wang, M. Magris, G. Hidalgo, M Contreras, Ó. Noya-Alarcón, O. Lander, J. McDonald, M. Cox, J. Walter, P. L. Oh, J. F. Ruiz, S. Rodriguez, N. Shen, S. J. Song, J. Metcalf, R. Knight, G. Dantas, M. G. Dominguez-Bello, The microbiome of uncontacted Amerindians. Sci Adv. 1, doi: 10.1126/sciadv.1500183 (2015).

6. E. R. Davenport, J. G. Sanders, S. J. Song, K. R. Amato, A. G. Clark, R. Knight, The human microbiome in evolution. BMC Biol. 15, doi: 10.1186/s12915-017-0454-7 (2017).

7. M. J. Blaser, The past and future biology of the human microbiome in an age of extinctions. Cell 172, 1173–1177 (2018).

8. S. L. Schnorr, M. Candela, S. Rampelli, M. Centanni, C. Consolandi, G. Basaglia, S. Turroni, E. Biagi, C. Peano, M. Severgnini, J. Fiori, R. Gotti, G. De Bellis, D. Luiselli, P. Brigidi, A. Mabulla, F. Marlowe, A. G. Henry, A. N. Crittenden, Gut microbiome of the Hadza hunter-gatherers. Nat Commun. 5, doi: 10.1038/ncomms4654 (2014).

9. A. J. Obregon-Tito, R. Y. Tito, J. Metcalf, K. Sankaranarayanan, J. C. Clemente, L. K. Ursell, Z. Zech Xu, W. Van Treuren, R. Knight, P. M. Gaffney, P. Spicer, P. Lawson, L. Marin-Reyes, O. Trujillo-Villarroel, M. Foster, E. Guija-Poma, L. Troncoso-Corzo, C. Warinner, A. T. Ozga, C. M. Lewis, Subsistence strategies in traditional societies distinguish gut microbiomes. Nat Commun. 6505, doi: 10.1038/ncomms7505 (2015).

10. C. Warinner, C. M. Lewis, Microbiome and health in past and present human populations. Am Anthrop. 117, 740–741 (2015).

11. R. Lee, R. H. Daly, Cambridge encyclopaedia of hunters and gatherers. Cambridge, Cambridge University Press ISBN 9780521609197 (1999).

12. A. Gomez, K. J. Petrzelkova, M. B. Burns, C. J. Yeoman, A. R. Amato, K. Vlckova, D. Modry, A. Todd, C. A. Jost Robinson, M. J. Remis, M. G. Torralba, E. Morton, J. D. Umaña, F. Carbonero, H. R. Gaskins, K. E. Nelson, B. A. Wilson, R. M. Stumpf, B. A. White, S. R. Leigh, Gut microbiome of coexisting BaAka Pygmies and Bantu reflects gradients of traditional subsistence patterns. Cell Rep. 14, 2142–2153 (2016).

13. C. Girard, N. Tromas, M. Amyot, B. J. Shapiro, Gut microbiome of the Canadian Arctic Inuit. mSphere 2, doi: 10.1128/mSphere.00297-16 (2017).

14. A. Y. Voigt, P. I. Costea, J. R. Kultima, S. S. Li, G. Zeller, S. Sunagawa, P. Bork, Temporal and technical variability of human gut metagenomes. Genome Biol. 16, doi: 10.1186/s13059-015-0639-8 (2015).

15. J. Walter, R. Ley, The human gut microbiome: Ecology and recent evolutionary changes. Annu Rev Microbiol. 65, 411–429 (2011).

16. M. J. Blaser, S. Falkow, What are the consequences of the disappearing human microbiota? Nat Rev Microbiol. 7, 887–894 (2009).

17. C. J. Adler, K. Dobney, L. S. Weyrich, J. Kaidonis, A. W. Walker, W. Haak, C. J. Bradshaw, G. Townsend, A. Soltysiak, K. W. Alt, J. Parkhill, A. Cooper, Sequencing ancient calcified dental plaque shows changes in oral microbiota with dietary shifts of the Neolithic and Industrial revolutions. Nat Genet. 45, 450–455 (2013).

18. S. Quercia, M. Candela, C. Giuliani, S. Turroni, D. Luiselli, S. Rampelli, P. Brigidi, C. Franceschi, M. G. Bacalini, P. Garagnani, C. Pirazzini, From lifetime to evolution: Timescales of human gut microbiota adaptation. Front Microbiol. 5, doi: 10.3389/fmicb.2014.00587 (2014).

19. A. H. Moeller, A. Caro-Quintero, D. Mjungu, A. V. Georgiev, E. V. Lonsdorf, M. N. Muller, A. E. Pusey, M. Peeters, B. H. Hahn, H. Ochman, Cospeciation of gut microbiota with hominids. Science 353, 380–382 (2016).

20. H. Reyes-Centeno, K. Harvati, G. Jäger, Tracking modern human population history from linguistic and cranial phenotype. Sci Rep. 6, doi: 10.1038/srep36645 (2016).

21. C. Houldcroft, J. B. Ramond, R. F. Rifkin, S. J. Underdown, Migrating microbes: What pathogens can tell us about population movements and human evolution. Ann Hum Biol. 44, 397–407 (2017).

22. B. Linz, F. Balloux, Y. Moodley, A. Manica, H. Liu, P. Roumagnac, D. Falush, C. Stamer, F. Prugnolle, S. W. van der Merwe, Y. Yamaoka, D. Y. Graham, E. Perez-Trallero, T. Wadstrom, S. Suerbaum, M. Achtman, An African origin for the intimate association between humans and *Helicobacter pylori*. Nature 445, 915–918 (2007).

23. M. T. Gilbert, D. L. Jenkins, A. Götherstrom, N. Naveran, J. J. Sanchez, M. Hofreiter, P. F. Thomsen, J. Binladen, T. F. Higham, R. M. Yohe, R. Parr, L. S. Cummings, E. Willerslev, DNA from pre-Clovis human coprolites in Oregon, North America. Science 320, 786–789 (2008).

24. R. J. Cano, J. Rivera-Perez, G. A. Toranzos, T. M. Santiago-Rodriguez, Y. M. Narganes-Storde, L. Chanlatte-Baik, E. García-Roldán, L. Bunkley-Williams, S. E. Massey, Paleomicrobiology: Revealing fecal microbiomes of ancient indigenous cultures. PLoS One 9, doi: 10.1371/journal.pone.0106833 (2014).

25. T. M. Santiago-Rodriguez, G. Fornaciari, S. Luciani, S. E. Dowd, G. A. Toranzos, I. Marota, R. J. Cano, Gut microbiome of an 11^th^ century A.D. Pre-Columbian Andean mummy. PLoS One 10, doi: 10.1371/journal.pone.0138135 (2015).

26. S. Rampelli, S. L. Schnorr, C. Consolandi, S. Turroni, M. Severgnini, C. Peano, P. Brigidi, A. N. Crittenden, A. G. Henry, M. Candela, Metagenome sequencing of the Hadza hunter-gatherer gut microbiota. Curr Biol. 25, 1682–1693 (2015).

27. G. Porraz, A. Val, L. Dayet, P. de la Pena, K. Douze, C. E. Miller, M. Murungi, C. Tribolo, V. Schmid, C. Sievers, Bushman Rock Shelter (Limpopo, South Africa): A perspective from the edge of the Highveld. S Afr Archaeol Bull. 70, 166–179 (2015).

28. T. Russell, F. Silva, J. Steele, Modelling the spread of farming in the Bantu-speaking regions of Africa: an archaeology-based phylogeography. PLoS One 9, doi: 10.1371/journal.pone.0087854 (2014).

29. G. Porraz, A. Val, C. Tribolo, N. Mercier, P. de la Peña, M. M. Haaland, M. Igreja, C. E. Miller, V. Schmid, The MIS5 Pietersburg at ‘28’ Bushman Rock Shelter, Limpopo Province, South Africa. PLoS One 13, doi: https://doi.org/10.1371/journal.pone.0202853 (2018).

30. J. R. Wood, J. M. Wilmshurst, A protocol for subsampling Late Quaternary coprolites for multi-proxy analysis. Quat Sci Rev. 138, 1–5 (2014).

31. C. Walker, Signs of the wild: A field guide to the spoor and signs of the mammals of southern Africa. Cape Town: Struik Nature (2015).

32. R. D. Estes, The behavior guide to African mammals: Including hoofed mammals, carnivores, primates. Berkeley, University of California Press pp. 1–611 (2012).

33. T. N. Huffman, Mapungubwe and Great Zimbabwe: The origin and spread of social complexity in southern Africa. J Anthropol Archaeol. 28, 37–54 (2007).

34. M. Samuelson, Rendering the Cape-as-port: Sea-mountain, Cape of Storms/Good Hope, Adamastor and local-world literary formations. J S Afr Stud. 42, 523–537 (2016).

35. M. G. Campana, N. Robles García, F. J. Rühli, N. Tuross, False positives complicate ancient pathogen identifications using high-throughput shotgun sequencing. BMC Res Notes 7, doi: 10.1186/1756-0500-7-111 (2014).

36. E. Willerslev, L. Davison, M. Moora, M. Zobel, E. Coissac, M. E. Edwards, E. D. Lorenzen, M. Vestergård, G. Gussarova, J. Haile, J. Craine, L. Gielly, S. Boessenkool, L. S. Epp, P. B. Pearman, R. Cheddadi, D. Murray, K. A. Bråthen, N. Yoccoz, H. Binney, C. Cruaud, P. Wincker, T. Goslar, I. Greve Alsos, E. Bellemain, A. Krag Brysting, R. Elven, J. H. Sønstebø, J. Murton, A. Sher, M. Rasmussen, R. Rønn, T. Mourier, A. Cooper, J. Austin, P. Möller, D. Froese, G. Zazula, F. Pompanon, D. Rioux, V. Niderkorn, A. Tikhonov, G. Savvinov, R. G. Roberts, R. D. E. MacPhee, M. T. P. Gilbert, K. H. Kjær, L. Orlando, C. Brochmann, P. Taberlet, Fifty thousand years of Arctic vegetation and megafaunal diet. Nature 506, 47–51 (2014).

37. C. J. Weiß, M. Dannemann, K. Prüfer, H. A. Burbano, Contesting the presence of wheat in the British Isles 8,000 years ago by assessing ancient DNA authenticity from low-coverage data. Elife 4, doi: 10.7554/eLife.10005 (2015).

38. A. Ginolhac, M. Rasmussen, M. T. P. Gilbert, E. Willerslev, L. Orlando, mapDamage: Testing for damage patterns in ancient DNA sequences. Bioinformatics 27, 2153–2155 (2011).

39. H. Jónsson, A. Ginolhac, M. Schubert, P. L. Johnson, L. Orlando, mapDamage 2.0: Fast approximate Bayesian estimates of ancient DNA damage parameters. Bioinformatics 29, 1682–1684 (2013).

40. S. Badenhorst, Intensive hunting during the Iron Age of Southern Africa. J Hum Palaeoecol. 20, 41–45 (2015).

41. J. Lloyd-Price, G. Abu-Ali, C. Huttenhower, The healthy human microbiome. Genome Med. 8, doi: https://doi.org/10.1186/s13073-016-0307-y (2016).

42. J. Qin, R. Li, J. Raes, M. Arumugam, K. Solvsten Burgdorf, C. Manichanh, T. Nielsen, N. Pons, F. Levenez, T. Yamada, D. R. Mende, J. Li, J. Xu, S. Li, D. Li, J. Cao, B. Wang, H. Liang, H. Zheng, Y. Xie, J. Tap, P. Lepage, M. Bertalan, J-. M. Batto, T. Hansen, D. Le Paslier, A. Linneberg, H. B. Nielsen, E. Pelletier, P. Renault, T. Sicheritz-Ponten, K. Turner, H. Zhu, C. Yu, S. Li, M. Jian, Y. Zhou, Y. Li, X. Zhang, S. Li, N. Qin, H. Yang, J. Wang, S. Brunak, J. Doré, F. Guarner, K. Kristiansen, O. Pedersen, J. Parkhill, J. Weissenbach, MetaHIT Consortium, P. Bork, S. D. Ehrlich, J. Wang, A human gut microbial gene catalog established by metagenomic sequencing. Nature 464, 59–65 (2010).

43. M. Derrien, J. E. van Hylckama Vlieg, Fate, activity, and impact of ingested bacteria within the human gut microbiota. Trends Microbiol. 23, 354–366 (2015).

44. G. B. Gloor, J. M. Macklaim, V. Pawlowsky-Glahn, J. J. Egozcue, Microbiome datasets are compositional: And this is not optional. Front Microbiol. 8, doi: https://doi.org/10.3389/fmicb.2017.02224 (2017).

45. L. Kistler, R. Ware, O. Smith, M. Collins, R. G. Allaby, A new model for ancient DNA decay based on paleogenomic meta-analysis. Nucleic Acids Res. 45, 6310–6320 (2017).

46. I. M. Velsko, L. A. F. Frantz, A. Herbig, G. Larson, C. Warinner, Selection of appropriate metagenome taxonomic classifiers for ancient microbiome research. mSystems 3, doi: 10.1128/mSystems.00080-18 (2018).

47. M. E. J. Woolhouse, Where do emerging pathogens come from? Microbe 1, 511–515 (2006).

48. Y. Nédélec, J. Sanz, G. Baharian, Z. A. Szpiech, A. Pacis, A. Dumaine, J. C. Grenier, A. Freiman, A. J. Sams, S. Hebert, A. Pagé Sabourin, F. Luca, R. Blekhman, R. D. Hernandez, R. Pique-Regi, J. Tung, V. Yotova, L. B. Barreiro, Genetic ancestry and natural selection drive population differences in immune responses to pathogens. Cell 167, 657–669 (2017).

49. S. E. Kessler, T. R. Bonnell, R. W. Byrne, C. A. Chapman, Selection to outsmart the germs: The evolution of disease recognition and social cognition. J Hum Evol. 108, 92–109 (2017).

50. R. Thornhill, C. L. Fincher, The parasite-stress theory of values and sociality. London: Springer pp. 1–440 (2014).

51. E. S. Pierce, Could *Mycobacterium avium* subspecies *paratuberculosis* cause Crohn’s disease, ulcerative colitis…and colorectal cancer? BMC Infect. Agents Cancer 13, doi: https://doi.org/10.1186/s13027-017-0172-3 (2018).

52. N. R. Shin, T. W. Whon. J. W. Bae, *Proteobacteria*: Microbial signature of dysbiosis in gut microbiota. Trends Biotechnol. 33, 496–503 (2015).

53. E. R. Hughes, M. G. Winter, B. A. Buerkop, L. Spiga, T. Furtado de Carvalho, W. Zhu, C. C. Gillis, L. Büttner, M. P. Smoot, C. L. Behrendt, S. Cherry, R. L. Santos, L. V. Hooper, S. E. Winter, Microbial respiration and formate oxidation as metabolic signatures of inflammation-associated dysbiosis. Cell Host Microbe. 21, 208–219 (2017).

54. R. E. Ley, P. J. Turnbaugh, S. Klein, J. I. Gordon, Microbial ecology: Human gut microbes associated with obesity. Nature 444, 1022–1023 (2006).

55. J. C. Clemente, L. K. Ursell, L. W. Parfrey, R. Knight, The impact of the gut microbiota on human health: An integrative view. Cell 148, 1258–1270 (2012).

56. J. R. Marchesi, D. H. Adams, F. Fava, G. D. Hermes, G. M. Hirschfield, G. Hold, M. N. Quraishi, L. Kinross, H. Smidt, K. M. Tuohy, L. V. Thomas, E. G. Zoetendal, A. Hart, The gut microbiota and host health: A new clinical frontier. Gut 65, 330–339 (2016).

57. C. De Filippo, D. Cavalieri, M. Di Paola, M. Ramazzotti, J. B. Poullet, S. Massart, S. Collini, G. Pieraccini, P. Lionetti, Impact of diet in shaping gut microbiota revealed by a comparative study in children from Europe and rural Africa. Proc Natl Acad Sci U S A. 107, 14691–14696 (2010).

58. T. V. Maier, M. Lucio, L. H. Lee, N. C. VerBerkmoes, C. J. Brislawn, J. Bernhardt, R. Lamendella, J. E. McDermott, N. Bergeron, S. S. Heinzmann, J. T. Morton, A. González, G. Ackermann, R. Knight, K. Riedel, R. M. Krauss, P. Schmitt-Kopplin, J. K. Jansson, Impact of dietary resistant starch on the human gut microbiome, metaproteome, and metabolome. MBio 8, doi: 10.1128/mBio.01343-17 (2017).

59. M. Arumugam, J. Raes, E. Pelletier, D. Le Paslier, T. Yamada, D. R. Mende, G. R. Fernandes, J. Tap, T. Bruls, J-. M. Batto, M. Bertalan, N. Borruel, F. Casellas, L. Fernandez, L. Gautier, T. Hansen, M. Hattori, T. Hayashi, M. Kleerebezem, K. Kurokawa, M. Leclerc, F. Levenez, C. Manichanh, H. B. Nielsen, T. Nielsen, N. Pons, J. Poulain, J. Qin, T. Sicheritz-Ponten, S. Tims, D. Torrents, E. Ugarte, E. G. Zoetendal, J. Wang, F. Guarner, O. Pedersen, W. M. de Vos, S. Brunak, J. Doré, MetaHIT Consortium, J. Weissenbach, S. D. Ehrlich, P. Bork, Enterotypes of the human gut microbiome. Nature 473, 174–180 (2011).

60. G. D. Wu, J. Chen, C. Hoffmann, K. Bittinger, Y. Y. Chen, S. A. Keilbaugh, M. Bewtra, D. Knights, W. A. Walters, R. Knight, R. Sinha, E. Gilroy, K. Gupta, R. Baldassano, L. Nessel, H. Li, F. D. Bushman, J. D. Lewis, Linking long-term dietary patterns with gut microbial enterotypes Science 334, 105–108 (2011).

61. T. Yatsunenko, F. E. Rey, M. J. Manary, I. Trehan, M. G. Dominguez-Bello, M. Contreras, M. Magris, G. Hidalgo, R. N. Baldassano, A. P. Anokhin, A. C. Heath, B. Warner, J. Reeder, J. Kuczynski, J. G. Caporaso, C. A. Lozupone, C. Lauber, J. C. Clemente, D. Knights, R. Knight, J. I. Gordon, Human gut microbiome viewed across age and geography. Nature 486, 222–227 (2012).

62. M. J. Rodríguez-Vaquero, M. R. Alberto, M. C. Manca de Nadra, Antibacterial effect of phenolic compounds from different wines. Food Control 18, 93–101 (2007).

63. E. A. McKenney, M. Ashwell, J. E. Lambert, V. Fellner, Fecal microbial diversity and putative function in captive western lowland gorillas (*Gorilla gorilla gorilla*), common chimpanzees (*Pan troglodytes*), Hamadryas baboons (*Papio hamadryas*) and binturongs (*Arctictis binturong*). Integr Zool. 9, 557–569 (2014).

64. B. S. Samuel, E. E. Hansen, J. K. Manchester, P. M. Coutinho, B. Henrissat, R. Fulton, P. Latreille, K. Kim, R. K. Wilson, J. I. Gordon, Genomic and metabolic adaptations of *Methanobrevibacter smithii* to the human gut. Proc Natl Acad Sci U S A. 104, 10643–10648 (2007).

65. T. Chen, W. Long, C. Zhang, S. Liu, L. Zhao, B. R. Hamaker, Fiber-utilizing capacity varies in *Prevotella*-versus *Bacteroides*-dominated gut microbiota. Sci. Rep. 7, doi: 10.1038/s41598-017-02995-4 (2017).

66. L. R. Lopetuso, F. Scaldaferri, V. Petito, A. Gasbarrini, Commensal *Clostridia*: Leading players in the maintenance of gut homeostasis. Gut Pathog. 5, doi: 10.1186/1757-4749-5-23 (2013).

67. C. Chassard, E. Delmas, C. Robert, A. Bernalier-Donadille, The cellulose-degrading microbial community of the human gut varies according to the presence or absence of methanogens. FEMS Microbiol Ecol. 74, 205–213 (2010).

68. E. R. Morton, J. Lynch, A. Froment, S. Lafosse, E. Heyer, M. Przeworski, R. Blekhman, L. Ségurel, Variation in rural African gut microbiota is strongly correlated with colonization by *Entamoeba* and subsistence. PLoS Genet. 11, doi: 10.1371/journal.pgen.1005658 (2015).

69. C. A. Lozupone, J. I. Stombaugh, J. I. Gordon, J. K. Jansson, R. Knight, Diversity, stability and resilience of the human gut microbiota. Nature 489, 220–230 (2012).

70. P. Vangay, A. J. Johnson, T. L. Ward, G. A. Al-Ghalith, R. R. Shields-Cutler, B. M. Hillmann, S. K. Lucas, L. K. Beura, E. A. Thompson, L. M. Till, R. Batres, B. Paw, S. L. Pergament, P. Saenyakul, M. Xiong, A. D. Kim, G. Kim, D. Masopust, D. Knight, US immigration westernizes the human gut microbiome. Cell 175, doi: 10.1016/j.cell.2018.10.029 (2018).

71. K. Tomita, T. Nagura, Y. Okuhara, H. Nakajima-Adachi, N. Shigematsu, T. Aritsuka, S. Kaminogawa, S. Hachimura, Dietary melibiose regulates *th* cell response and enhances the induction of oral tolerance. Biosci Biotechnol Biochem. 71, 2774–2280 (2007).

72. H. Fiedler, Short-chain chlorinated paraffins: Production, use and international regulations. In: Boer J. (ed.) Chlorinated paraffins. The handbook of environmental chemistry. Berlin, Springer pp. 1–41 (2010).

73. J. Glüge, Z. Wang, C. Bogdal, M. Scheringer, K. Hungerbühler, Global production, use, and emission volumes of short-chain chlorinated paraffins: A minimum scenario. Sci Total Environ. 573, 1132–46 (2016).

74. A. Paoli, A. Bianco, K. A. Grimaldi, A. Lodi, G. Bosco, Long term successful weight loss with a combination biphasic ketogenic Mediterranean diet and Mediterranean diet maintenance protocol. Nutrients 5, 5205–5217 (2013).

75. V. M. D’Costa, C. E. King, L. Kalan, M. Morar, W. W. Sung, C. Schwarz, D. Froese, G. Zazula, F. Calmels, R. Debruyne, G. B. Golding, H. N. Poinar, G. D. Wright, Antibiotic resistance is ancient. Nature 477, 457–461 (2011).

76. R. Kaur, V. Gautam, P. Ray, G. Singh, L. Singhal, R. Tiwari, Daptomycin susceptibility of methicillin resistant *Staphylococcus aureus* (MRSA) Indian J Med Res. 136, 676–677 (2012).

77. K. Ye, F. Gao, D. Wang, O. Bar-Yosef, A. Keinan, Dietary adaptation of FADS genes in Europe varied across time and geography. Nat Ecol Evol. 1, doi: 10.1038/s41559-017-0167 (2017).

78. G. A. W. Rook, L. R. Brunet, Microbes, immunoregulation, and the gut. Gut 54, 317–320 (2005).

79. B. Stecher, L. Maier, W. D. Hardt, ‘Blooming’ in the gut: How dysbiosis might contribute to pathogen evolution. Nat Rev Microbiol. 11, 277–284 (2013).

80. M. Stuiver, P. D. Quay, Atmospheric ^14^C changes resulting from fossil fuel CO^2^ release and cosmic ray flux variability. Earth and Planetary Science 53, 349–362 (1981).

81. M. Nemec. L. Wacker, H. Gäggeler, Optimization of the graphitization process at AGE-1. Radiocarbon 52, 1380–1393 (2010).

82. S. Woodborne, D. Huchzermeyer, D. Govender, D. J. Pienaar, G. Hall, J. G. Myburgh, A. R. Deacon, J. Venter, N. Lubcker, Ecosystem change and the Olifants River crocodile mass mortality events. Ecosphere 3, doi: http://dx.doi.org/10.1890/ES12-00170.1 (2012).

83. B. Dufour, M. Le Bailly, Testing new parasite egg extraction methods in paleoparasitology and an attempt at quantification. Int J Paleopathol. 3, 199–203 (2013).

84. C. Warinner, A. Herbig, A. Mann, J. A. Fellows Yates, C. L. Weiß, H. A. Burbano, L. Orlando, J. Krause, A robust framework for microbial archaeology. Annu Rev Genomics Hum Genet. 18, 321–356 (2017).

85. S. J. Salter, M. J. Cox, E. M. Turek, S. T. Calus, W. O. Cookson, M. F. Moffatt, P. Turner, J. Parkhill, N. J. Loman, A. W. Walker, Reagent and laboratory contamination can critically impact sequence-based microbiome analyses. BMC Biol. 12, doi: 10.1186/s12915-014-0087-z (2014).

86. A. P. Lauder, A. M. Roche, S. Sherrill-Mix, A. Bailey, A. L. Laughlin, K. Bittinger, R. Leite, M. A. Elovitz, S. Parry, F. D. Bushman, Comparison of placenta samples with contamination controls does not provide evidence for a distinct placenta microbiota. Microbiome 4, doi: https://doi.org/10.1186/s40168-016-0172-3 (2016).

87. P. I. Costea, G. Zeller, S. Sunagawa, E. Pelletier, A. Alberti, F. Levenez, M. Tramontano, M. Driessen, R. Hercog, F. Jung, J. Roat Kultima, M. R. Hayward, L. P. Coelho, E. Allen-Vercoe, L. Bertrand, M. Blaut, J. R. M. Brown, T. Carton, S, Cools-Portier, M. Daigneault, M. Derrien, A. Druesne, W. M. de Vos, B. B. Finlay, H. J. Flint, F. Guarner, M. Hattori, H. Heilig, R. A. Luna, J. van Hylckama Vlieg, J. Junick, I. Klymiuk, P. Langella, E. Le Chatelier, V. Mai, C. Manichanh, J. C. Martin, C. Mery, H. Morita, P. W. O’Toole, C. Orvain, K. Raosaheb Patil, J. Penders, S. Persson, N. Pons, M. Popova, A. Salonen, D. Saulnier, K. P. Scott, B. Singh, K. Slezak, P. Veiga, J. Versalovic, L. Zhao, E. G. Zoetendal, S. D. Ehrlich, J. Dore, P. Bork, Towards standards for human fecal sample processing in metagenomic studies. Nat Biotechnol. 35, 1069–1076 (2017).

88. J. C. Stearns, M. D. J. Lynch, D. B. Senadheera, H. C. Tenenbaum, M. B. Goldberg, D. G. Cvitkovitch, K. Croitoru, G. Moreno-Hagelsieb, J. D. Neufeld, Bacterial biogeography of the human digestive tract. Sci. Rep. 1, doi: 10.1038/srep00170 (2011).

89. S. A. Smits, C. G. Gonzalez, J. Leach, A. Manjurano, E. D. Sonnenburg, J. S. Lichtman, J. Changalucha, M. G. Dominguez-Bello, G. Reid, R. Knight, J. E. Elias, J. L. Sonnenburg, Seasonal cycling in the gut microbiome of the Hadza hunter-gatherers of Tanzania. Science 6353, 802–806 (2017).

90. G. P. Donaldson, S. M. Lee, S. K. Mazmanian, Gut biogeography of the bacterial microbiota. Nat Rev Microbiol. 14, 20–32 (2016).

91. D. Vandeputte, G. Falony, S. Vieira-Silva, R. Y. Tito, M. Joossens, J. Raes, Stool consistency is strongly associated with gut microbiota richness and composition, enterotypes and bacterial growth rates. BMJ Gut 65, 57–62 (2016).

92. S. J. Ott, M. Musfeldt, K. N. Timmis, J. Hampe, D. F. Wenderoth, S. Schreiber, *In vitro* alterations of intestinal bacterial microbiota in fecal samples during storage. Diagn Microbiol Infect Dis. 50, 237–245 (2004).

93. R. Y. Tito, S. Macmil, G. Wiley, F. Najar, L. Cleeland, C. Qu, P. Wang, F. Romagne, S. Leonard, A. Jiménez Ruiz, K. Reinhard, B. A. Roe, C. M. Lewis Jr., Phylotyping and functional analysis of two ancient human microbiomes. PLoS One 3, doi: 10.1371/journal.pone.0003703 (2008).

94. R. Y. Tito, D. Knights, J. Metcalf, A. J. Obregon-Tito, L. Cleeland, F. Najar, B. Roe, K. Reinhard, K. Sobolik, S. Belknap, M. Foster, P. Spicer, R. Knight, C. M. Lewis Jr., Insights from characterizing extinct human gut microbiomes. PLoS One 12, doi: https://doi.org/10.1371/journal.pone.0051146 (2012).

95. B. Bushnell, BBMap. http://sourceforge.net/projects/bbmap/ (2014).

96. S. Sawyer, J. Krause, K. Guschanski, V. Savolainen, S. Pääbo, Temporal patterns of nucleotide misincorporations and DNA fragmentation in ancient DNA. PLoS One 7, doi: 10.1371/journal.pone.0034131 (2012).

97. J. Dabney, M. Knapp, I. Glocke, M. Gansauge, A. Weihmann, B. Nickel, C. Valdiosera, N. García, S. Pääbo, J. Arsuag, M. Meyer, Complete mitochondrial genome sequence of a Middle Pleistocene cave bear reconstructed from ultrashort DNA fragments. Proc Natl Acad Sci U S A. 110, 15758–15763 (2013).

98. M. Schubert, S. Lindgreen, L. Orlando, AdapterRemoval v2: Rapid adapter trimming, identification, and read merging. BMC Res Notes. 9, doi: 10.1186/s13104-016-1900-2 (2016).

99. S. F. Altschul, W. Gish, W. Miller, E. W. Myers, D. J. Lipman, Basic local alignment search tool. J Mol Biol. 215, 403–410 (1990).

100. D. H. Huson, S. Beier, I. Flade, A. Górska, M. El-Hadidi, S. Mitra, H. Ruscheweyh, R. Tappu, MEGAN Community Edition: Interactive exploration and analysis of large-scale microbiome sequencing data. PLoS Comput Biol. 12, doi: 10.1371/journal.pcbi.1004957 (2016).

101. A. W. Briggs, U. Stenzel, P. L. Johnson, R. E. Green, J. Kelso, K. Prüfer, M. Meyer, J. Krause, M. T. Ronan, M. Lachmann, S. Pääbo, Patterns of damage in genomic DNA sequences from a Neandertal. Proc Natl Acad Sci U S A. 104, 14616–14621 (2007).

102. F. V. Seersholm, M. W. Pedersen, M. J. Søe, H. Shokry, S. S. Mak, A. Ruter, M. Raghavan, W. Fitzhugh, K. H. Kjær, E. Willerslev, M. Meldgaard, C. M. O. Kapel, A. J. Hansen DNA evidence of bowhead whale exploitation by Greenlandic Paleo-Inuit 4,000 years ago. Nat Commun. 7, doi: 10.1038/ncomms13389 (2016).

103. Å. J. Vågene, A. Herbig, M. G. Campana, N. M. Robles García, C. Warinner, S. Sabin, M. A. Spyrou, A. Andrades Valtueña, D. Huson, N. Tuross, K. I. Bos, J. Krause, *Salmonella enterica* genomes from victims of a major sixteenth-century epidemic in Mexico. Nat Ecol Evol. 2, 520–528 (2017).

104. J. C. Tackney, B. A. Potter, J. Raff, M. Powers, W. S. Watkins, D. Warner, J. D. Reuther, J. D. Irish, D. H. O’Rourke, Two contemporaneous mitogenomes from terminal Pleistocene burials in eastern Beringia. Proc Natl Acad Sci U S A. 112, doi: 10.1073/pnas.1511903112 (2015).

105. J. Dabney, M. Meyer, S. Pääbo, Ancient DNA damage. Mol Ecol Res. 17, doi: 10.1111/1755-0998.12595 (2017).

106. C. L. Weiß, V. J. Schuenemann, J. Devos, G. Shirsekar, E. Reiter, B. A. Gould, J. R. Stinchcombe, J. Krause, H. A. Burbano, Temporal patterns of damage and decay kinetics of DNA retrieved from plant herbarium specimens. R Soc Open Sci. 3, doi: 10.1098/rsos.160239 (2016).

107. H. Malmström, E. M. Svensson, M. T. P. Gilbert, E. Willerslev, A. Götherström, G. Holmlund, More on contamination: The use of asymmetric molecular behavior to identify authentic ancient human DNA. Mol Biol Evol. 24, 998–1004 (2007).

108. K. Prüfer, U. Stenzel, M. Hofreiter, S. Pääbo, J. Kelso, R. E. Green, Computational challenges in the analysis of ancient DNA. Genome Biol. 11, doi: 10.1186/gb-2010-11-5-r47 (2010).

109. G. A. Lugli, C. Milani, L. Mancabelli, F. Turroni, C. Ferrario, S. Duranti, D. van Sinderen, M Ventura, Ancient bacteria of the Ötzi’s microbiome: A genomic tale from the Copper Age. Microbiome 5, doi: 10.1186/s40168-016-0221-y (2017).

110. S. M. Lakin, C. Dean, N. R. Noyes, A. Dettenwanger, A. S. Ross, E. Doster, P. Rovira, Z. Abdo, K. L. Jones, J. Ruiz, K. E. Belk, P. S. Morley, C. Boucher, MEGARes: An antimicrobial resistance database for high throughput sequencing. Nucleic Acids Res. 45, doi: 10.1093/nar/gkw1009 (2017).

111. H. Li, R. Durbin, Fast and accurate short read alignment with Burrows-Wheeler transform. Bioinformatics 25, 1754–1760 (2009).

112. P. J. McMurdie, S. Holmes, phyloseq: an R package for reproducible interactive analysis and graphics of microbiome census data. PLoS One 8, doi: 10.1371/journal.pone.0061217 (2013).

113. J. G. Caporaso, J. Kuczynski, J. Stombaugh, K. Bittinger, F. D. Bushman, E. K. Costello, N. Fierer, A. G. Pena, J. K Goodrich, J. I. Gordon, G. A. Huttley, S. T. Kelley, D. Knights, J. E. Koenig, R. E Ley, C. A. Lozupone, D. McDonald, B. D. Muegge, M. Pirrung, J. Reeder, J. R. Sevinsky, P. J. Turnbaugh, W. A. Walters, J. Widmann, T. Yatsunenko, J. Zaneveld, R. Knight, QIIME allows analysis of high-throughput community sequencing data. Nat Methods 7, doi: 10.1038/nmeth.f.303 (2010).

114. L. Jiang, A. Amir, J. T. Morton, R. Heller, E. Arias-Castro, R. Knight, Discrete false-discovery rate improves identification of differentially abundant microbes. mSystems 2, doi: 10.1128/mSystems.00092-17 (2017).

