## Supplementary Tables for "Multi-proxy analyses of a mid-15^th^ century ‘Middle Iron Age’ Bantu-speaker palaeo-faecal specimen elucidates the configuration of the ‘ancestral’ sub-Saharan African intestinal microbiome"

**Supplementary Table 1.** Mapped DNA sequence reads for environmental- and subsistence-related taxa detected in this study. Statistically-significant ( *i.e* ., verified ancient) C-T *p* -values are indicated in bold text. Analyses were performed using high-quality filtered read alignments against NCBI reference genomes. DNA damage estimation analyses was performed using PMDtools ('C-T *p* -values').

| Class | Taxon | Common name | *E-LPC reads* | BWA aligned (mapped) sequence reads | | | | | | Mean read length (bp) | C-T *p* -value | | | | |
|---|---|---|---|---|---|---|---|---|---|---|---|---|---|---|---|
| | | | | BRS1 | BRS2 | BRS3 | BRS4 | BRS5 | Total | | BRS1 | BRS2 | BRS3 | BRS4 | BRS5 |
| *Aconoidasida* | *Babesia bovis* | Protozoan parasite | *0* | 0 | *59* | *45* | *820* | 0 | 924 | 43.8 | – | **0.0273139** | **0.0314666** | 0.3810030 | – |
| *Bacilli* | *Lactococcus lactis* | Bacterium | *0* | 100 | *541* | *340* | *4818* | 0 | 5799 | 70.6 | 0.5582629 | **0.0027567** | **0.0355133** | **0.0000015** | – |
| *Dicotyledonae* | *Ficus sur* | Cluster fig | *0* | 6 | *179* | *59* | *1171* | 2 | 1417 | 67.0 | 0.3958356 | **0.0120650** | 0.5833718 | **0.0334960** | 0.5000000 |
| *Secernentea* | *Parastrongyloides trichosuri* | Nematode parasite | *0* | 11 | *1281* | *447* | *8481* | 13 | 10233 | 65.0 | 0.5337832 | **0.0253805** | 0.2410555 | 0.0528887 | 0.5085527 |
| *Sordariomycetes* | *Podospora anserina* | Ascomycete fungus | *0* | 1 | *16* | *6* | *466* | 2 | 491 | 61.4 | – | 0.1799564 | 0.5000000 | 0.3874920 | – |
| | *Sorghum bicolor* | Sorghum or durra | *0* | 297 | *371* | *221* | *4359* | 166 | 5414 | 70.2 | 0.1020672 | **0.0066557** | 0.9053457 | **0.0000108** | 0.8410222 |
| *Monocotyledonae* | *Triticum aestivum* | Common wheat | *0* | 290 | *335* | *153* | *2826* | 505 | 4109 | 64.0 | 0.4611596 | 0.2830501 | 0.1544875 | 0.4358290 | 0.5196183 |
| | *Zea mays* | Maize or corn | *1* | 81 | *205* | *122* | *2661* | 0 | 3070 | 54.8 | 0.0718288 | 0.5123976 | 0.6440315 | 0.1234000 | – |
| | *Bos taurus* | Domestic cattle | *0* | 847 | *459* | *281* | *26215* | 91 | 27893 | 62.8 | 0.7214745 | 0.3237143 | 0.2069171 | 0.1183520 | 0.0844617 |
| | *Canis lupus familiaris* | Domestic dog | *0* | 0 | *0* | *0* | *0* | 71 | 71 | 51.4 | – | – | – | – | 0.7881283 |
| | *Canis mesomelas* | Black-backed jackal | *0* | 0 | *0* | *0* | *0* | 0 | 0 | – | – | – | – | – | – |
| | *Capra hircus* | Domestic goat | *0* | 1390 | *325* | *238* | *40118* | 117 | 42188 | 63.4 | 0.0580973 | **0.0132379** | 0.3843594 | **0.0000900** | 0.1955150 |
| | *Caracal caracal* | Caracal or rooikat | *0* | 0 | *0* | *0* | *0* | 0 | 0 | – | – | – | – | – | – |
| | *Cercopithecus aethiops* | Vervet monkey | *0* | 0 | *0* | *0* | *0* | 0 | 0 | – | – | – | – | – | – |
| | *Equus caballus* | Modern horse | *0* | 126 | *170* | *68* | *1395* | 79 | 1838 | 51.2 | 0.4765005 | 0.3077120 | 0.7492095 | **0.0007678** | 0.1313032 |
| *Mammalia* | *Homo sapiens* | Human | *56* | 335 | *270* | *117* | *1580* | 104 | 2462 | 61.2 | 0.6122221 | **0.0058446** | 0.7521352 | **0.0366582** | 0.1393800 |
| | *Ictonyx striatus* | Striped polecat | *0* | 0 | *0* | *0* | *0* | 0 | 0 | – | – | – | – | – | – |
| | *Macaca mulatta* | Rhesus macaque | *0* | 296 | *247* | *114* | *4593* | 96 | 5346 | 58.0 | 0.4855835 | 0.1070172 | 0.7493560 | **0.0269014** | 0.2207390 |
| | *Mellivora capensis* | Honey badger | *0* | 0 | *0* | *0* | *0* | 0 | 0 | – | – | – | – | – | – |
| | *Ovis aries* | Domestic sheep | *0* | 1026 | *289* | *206* | *36490* | 100 | 38111 | 62.6 | **0.0226299** | 0.1273375 | 0.2981717 | **0.0086748** | 0.1452299 |
| | *Pan troglodytes* | Chimpanzee | *0* | 390 | *276* | *111* | *2346* | 90 | 3213 | 60.6 | 0.5325693 | **0.0025637** | 0.7734403 | **0.0398303** | 0.3383619 |
| | *Panthera pardus* | Leopard | *0* | 0 | *0* | *0* | *0* | 0 | 0 | – | – | – | – | – | – |
| | *Papio ursinus* | Chachma baboon | *0* | 0 | *0* | *0* | *0* | 0 | 0 | – | – | – | – | – | – |
| | ***Total*** | | ***57*** | ***5196*** | ***5023*** | ***2528*** | ***138339*** | ***1436*** | ***152579*** | ***60.5*** | **–** | **–** | **–** | **–** | |

**Supplementary Table 2.** Information concerning the two direct radiocarbon (14C) Accelerator Mass Spectrometry (AMS) dates generated from two sub-samples taken from within the BRS palaeo-faecal specimen.

| Sample | Lab # | 13C | pMC | pMC_err | conv_age | age_err |
|---|---|---|---|---|---|---|
| BRS1 | IT-C-1020 | -16.79 | 94.37 | 0.524 | 470.2 | 44.1 |
| BRS2 | IT-C-1077 | -16.79 | 94.39 | 0.416 | 460.2 | 35.0 |

**Supplementary Table 3a.** Processing protocol and results for isotope analyses of samples derived from the BRS specimen indicating the relative proportions of C3 and C4 dietary contributions.

| Sample | Weight | N peak | 15N corr | %N | C peak | 13C corr | %C | C/N |
|---|---|---|---|---|---|---|---|---|
| BRS COP 1 | 0.82 | 12.66 | 11.28 | 1.52% | 19.49 | -17.93 | 22.46% | 17.20 |
| BRS COP 2 | 0.89 | 13.58 | 11.22 | 1.53% | 20.84 | -17.18 | 22.27% | 16.95 |
| BRS COP 3 | 0.84 | 13.36 | 11.45 | 1.58% | 22.03 | -17.65 | 24.78% | 18.29 |
| BRS LX 1 | 0.81 | 13.86 | 10.87 | 1.71% | 21.89 | -17.01 | 25.53% | 17.44 |
| BRS LX 2 | 0.84 | 14.16 | 10.98 | 1.69% | 21.30 | -16.98 | 23.95% | 16.57 |
| BRS LX 3 | 0.90 | 14.15 | 11.18 | 1.58% | 21.69 | -17.27 | 22.85% | 16.92 |
| BRS Acid 1 | 0.89 | 11.72 | 10.93 | 1.26% | 23.78 | -16.34 | 25.22% | 23.39 |
| BRS Acid 2 | 0.86 | 10.74 | 10.83 | 1.17% | 24.36 | -17.63 | 26.73% | 26.69 |
| BRS Acid 3 | 0.85 | 11.35 | 10.92 | 1.26% | 23.14 | -16.40 | 25.63% | 23.74 |
| BRS CS 1 | 1.04 | 22.14 | 11.10 | 2.23% | 39.95 | -23.08 | 36.27% | 18.96 |
| BRS CS 2 | 0.88 | 18.93 | 11.12 | 2.21% | 33.38 | -23.16 | 35.84% | 18.88 |
| BRS CS 3 | 1.03 | 21.77 | 11.05 | 2.21% | 39.73 | -23.60 | 36.52% | 19.26 |

**Supplementary Table 3b.** Results for isotope analyses *(Merck* standard) of samples derived from the BRS specimen indicating the relative proportions of C3 and C4 dietary contributions.

| | Weight | N peak | N | C peak | C |
|---|---|---|---|---|---|
| *Merck* | 0.22 | 29.65 | 7.82 | 9.43 | -20.28 |
| *Merck* | 0.41 | 56.33 | 7.95 | 17.96 | -20.23 |
| *Merck* | 0.60 | 82.27 | 7.97 | 26.34 | -20.26 |
| *Merck* | 0.24 | 34.14 | 7.81 | 10.24 | -20.27 |
| *Merck* | 0.40 | 57.99 | 7.92 | 17.83 | -20.28 |
| *Merck* | 0.60 | 89.53 | 7.78 | 26.13 | -20.22 |
| *Merck* | 0.23 | 36.75 | 7.91 | 10.33 | -20.27 |
| *Merck* | 0.41 | 61.18 | 7.98 | 18.12 | -20.28 |
| *Merck* | 0.64 | 92.49 | 7.95 | 28.44 | -20.25 |
| *Merck* | 0.64 | 92.85 | 7.80 | 28.68 | -20.26 |
| | | | **7.89** | | **-20.26** |
| | | | **0.08** | | **0.02** |

**Supplementary Table 3c.** Results for isotope analyses (*DL-Valine* standard) of samples derived from the BRS specimen indicating the relative proportions of C3 and C4 dietary contributions.

| | Weight | N peak | N | C peak | C |
|---|---|---|---|---|---|
| *DL-Valine* | 0.23 | 24.84 | -6.10 | 12.09 | -10.56 |
| *DL-Valine* | 0.41 | 44.35 | -6.22 | 21.92 | -10.53 |
| *DL-Valine* | 0.63 | 66.39 | -6.15 | 33.12 | -10.68 |
| *DL-Valine* | 0.21 | 24.24 | -6.13 | 11.07 | -10.51 |
| *DL-Valine* | 0.44 | 50.16 | -6.09 | 23.54 | -10.54 |
| *DL-Valine* | 0.60 | 68.98 | -6.21 | 32.72 | -10.56 |
| *DL-Valine* | 0.23 | 29.22 | -6.19 | 12.15 | -10.61 |
| *DL-Valine* | 0.42 | 49.46 | -6.12 | 22.52 | -10.63 |
| *DL-Valine* | 0.62 | 70.56 | -6.16 | 33.30 | -10.55 |
| *DL-Valine* | 0.62 | 70.17 | -6.13 | 32.95 | -10.53 |
| | | | **-6.15** | | **-10.57** |
| | | | **0.04** | | **0.05** |

**Supplementary Table 4.** Assignment of abundance of bacterial taxonomic categories to the BRS and ancient (Ötzi) and modern (Italian, Hadza and Malawian) comparative samples based on *p* -value (*p* = <0.05) designation. Group significance assignments were obtained via Qiime v1.9.1.

| OTU | Test-Statistic | *P* | BRS_mean | Otzi-UPLI_mean | Otzi-SI_mean | Otzi-LPLI_mean | Hadza_mean | Malawi_mean | Italian_mean | Taxonomy |
|---|---|---|---|---|---|---|---|---|---|---|
| ID251698 | 17.978824 | 0.006285 | 1.00 | 20.50 | 26.00 | 5.00 | 0.00 | 0.00 | 0.00 | k__Bacteria; p__Proteobacteria; c__Gammaproteobacteria; o__Pseudomonadales; f__Pseudomonadaceae; g__Pseudomonas; s__(Pseudomonas) |
| ID47877 | 17.978824 | 0.006285 | 3.00 | 70.50 | 94.00 | 12.00 | 0.00 | 0.00 | 0.00 | k__Bacteria; p__Proteobacteria; c__Gammaproteobacteria; o__Pseudomonadales; f__Pseudomonadaceae; g__Pseudomonas; s__Pseudomonas amygdali |
| ID40215 | 17.978673 | 0.006286 | 2.00 | 17.00 | 5.00 | 2.00 | 0.00 | 0.00 | 0.00 | k__Bacteria; p__Proteobacteria; c__Gammaproteobacteria; o__Pseudomonadales; f__Moraxellaceae; g__Acinetobacter; s__Acinetobacter junii |
| ID162481 | 17.968966 | 0.006310 | 0.00 | 260.00 | 191.00 | 9.00 | 0.00 | 0.00 | 0.00 | k__Eukaryota; p__Basidiomycota; c__Microbotryomycetes; o__(Microbotryomycetes); f__(Microbotryomycetes); g__(Microbotryomycetes); s__(Microbotryomycetes) |
| ID288795 | 17.968966 | 0.006310 | 0.00 | 501.00 | 248.00 | 8.00 | 0.00 | 0.00 | 0.00 | k__Eukaryota; p__Basidiomycota; c__Microbotryomycetes; o__Microbotryales; f__Microbotryaceae; g__Microbotryum; s__Microbotryum lychnidis-dioicae |
| ID5286 | 17.968966 | 0.006310 | 0.00 | 218.50 | 148.00 | 6.00 | 0.00 | 0.00 | 0.00 | k__Eukaryota; p__Basidiomycota; c__Microbotryomycetes; o__Sporidiobolales; f__Sporidiobolaceae; g__Rhodotorula; s__Rhodotorula toruloides |
| ID65741 | 17.936620 | 0.006393 | 3.33 | 6.00 | 8.00 | 1.00 | 0.00 | 0.00 | 0.00 | k__Bacteria; p__Proteobacteria; c__Gammaproteobacteria; o__Pseudomonadales; f__Pseudomonadaceae; g__Pseudomonas; s__Pseudomonas knackmussii |
| ID351095 | 17.929961 | 0.006410 | 70.67 | 1.00 | 0.00 | 1.00 | 0.00 | 0.00 | 0.00 | k__Bacteria; p__Firmicutes; c__Bacilli; o__Bacillales; f__Bacillaceae; g__Paucisalibacillus; s__Paucisalibacillus globulus |
| ID109871 | 17.929961 | 0.006410 | 7.33 | 1.00 | 0.00 | 1.00 | 0.00 | 0.00 | 0.00 | k__Eukaryota; p__Chytridiomycota; c__Chytridiomycetes; o__Rhizophydiales; f__(Batrachochytrium); g__Batrachochytrium; s__Batrachochytrium dendrobatidis |
| ID40214 | 17.915592 | 0.006447 | 4.67 | 346.50 | 1.00 | 106.00 | 0.00 | 0.00 | 0.00 | k__Bacteria; p__Proteobacteria; c__Gammaproteobacteria; o__Pseudomonadales; f__Moraxellaceae; g__Acinetobacter; s__Acinetobacter johnsonii |
| ID1439940 | 17.915592 | 0.006447 | 13.00 | 3.00 | 5.00 | 1.00 | 0.00 | 0.00 | 0.00 | k__Bacteria; p__Proteobacteria; c__Gammaproteobacteria; o__Pseudomonadales; f__Pseudomonadaceae; g__Pseudomonas; s__Pseudomonas sp. BAY1663 |
| ID157784 | 17.915592 | 0.006447 | 15.00 | 4.00 | 5.00 | 1.00 | 0.00 | 0.00 | 0.00 | k__Bacteria; p__Proteobacteria; c__Gammaproteobacteria; o__Pseudomonadales; f__Pseudomonadaceae; g__Pseudomonas; s__Pseudomonas thermotolerans |
| ID706570 | 17.914692 | 0.006449 | 591.33 | 1.00 | 1.00 | 1.00 | 0.00 | 0.00 | 0.00 | k__Bacteria; p__Proteobacteria; c__Gammaproteobacteria; o__Pseudomonadales; f__Pseudomonadaceae; g__Pseudomonas; s__Pseudomonas flexibilis |
| ID5061 | 17.914692 | 0.006449 | 8.00 | 1.00 | 1.00 | 1.00 | 0.00 | 0.00 | 0.00 | k__Eukaryota; p__Ascomycota; c__Eurotiomycetes; o__Eurotiales; f__Aspergillaceae; g__Aspergillus; s__Aspergillus niger |
| ID5073 | 17.914692 | 0.006449 | 19.00 | 1.00 | 1.00 | 1.00 | 0.00 | 0.00 | 0.00 | k__Eukaryota; p__Ascomycota; c__Eurotiomycetes; o__Eurotiales; f__Aspergillaceae; g__Penicillium; s__(Penicillium) |
| ID37914 | 17.906736 | 0.006470 | 26.00 | 1.00 | 0.00 | 1.00 | 0.00 | 0.00 | 0.00 | k__Bacteria; p__Actinobacteria <phylum>; c__Actinobacteria; o__Corynebacteriales; f__Dietziaceae; g__Dietzia; s__(Dietzia) |
| ID558169 | 17.906736 | 0.006470 | 67.67 | 1.00 | 0.00 | 1.00 | 0.00 | 0.00 | 0.00 | k__Bacteria; p__Firmicutes; c__Bacilli; o__Bacillales; f__Bacillaceae; g__Lentibacillus; s__Lentibacillus jeotgali |
| ID694055 | 17.906736 | 0.006470 | 19.00 | 1.00 | 0.00 | 1.00 | 0.00 | 0.00 | 0.00 | k__Bacteria; p__Firmicutes; c__Bacilli; o__Bacillales; f__Bacillaceae; g__Ornithinibacillus; s__Ornithinibacillus contaminans |
| ID403957 | 17.906736 | 0.006470 | 17.33 | 1.00 | 0.00 | 1.00 | 0.00 | 0.00 | 0.00 | k__Bacteria; p__Firmicutes; c__Bacilli; o__Bacillales; f__Bacillaceae; g__Virgibacillus; s__Virgibacillus sp. SK37 |
| ID1637974 | 17.906736 | 0.006470 | 50.67 | 1.00 | 0.00 | 1.00 | 0.00 | 0.00 | 0.00 | k__Bacteria; p__Firmicutes; c__Bacilli; o__(Bacilli bacterium VT-13-104); f__(Bacilli bacterium VT-13-104); g__(Bacilli bacterium VT-13-104); s__Bacilli bacterium VT-13-104 |
| ID746691 | 17.906736 | 0.006470 | 9.00 | 0.00 | 0.00 | 1.00 | 0.00 | 0.00 | 0.00 | k__Bacteria; p__Firmicutes; c__Bacilli; o__Bacillales; f__Bacillaceae; g__Oceanobacillus; s__Oceanobacillus kimchii |
| ID225453 | 17.894737 | 0.006501 | 29.00 | 1.00 | 0.00 | 0.00 | 0.00 | 0.00 | 0.00 | k__Bacteria; p__Actinobacteria <phylum>; c__Actinobacteria; o__Micrococcales; f__Micrococcaceae; g__Yaniella; s__Yaniella halotolerans |
| ID171693 | 17.894737 | 0.006501 | 21.33 | 1.00 | 0.00 | 0.00 | 0.00 | 0.00 | 0.00 | k__Bacteria; p__Firmicutes; c__Bacilli; o__Bacillales; f__Bacillaceae; g__Oceanobacillus; s__Oceanobacillus picturae |
| ID1482 | 17.894737 | 0.006501 | 17.33 | 1.00 | 0.00 | 0.00 | 0.00 | 0.00 | 0.00 | k__Bacteria; p__Firmicutes; c__Bacilli; o__Bacillales; f__Bacillaceae; g__Virgibacillus; s__Virgibacillus halodenitrificans |
| ID396024 | 17.894737 | 0.006501 | 93.33 | 1.00 | 0.00 | 0.00 | 0.00 | 0.00 | 0.00 | k__Eukaryota; p__Ascomycota; c__Eurotiomycetes; o__Eurotiales; f__Aspergillaceae; g__Aspergillus; s__Aspergillus ruber |
| ID587753 | 17.894614 | 0.006501 | 2.00 | 200.00 | 317.00 | 28.00 | 0.00 | 0.00 | 0.00 | k__Bacteria; p__Proteobacteria; c__Gammaproteobacteria; o__Pseudomonadales; f__Pseudomonadaceae; g__Pseudomonas; s__Pseudomonas chlororaphis |
| ID440512 | 17.894614 | 0.006501 | 3.33 | 9.50 | 22.00 | 1.00 | 0.00 | 0.00 | 0.00 | k__Bacteria; p__Proteobacteria; c__Gammaproteobacteria; o__Pseudomonadales; f__Pseudomonadaceae; g__Pseudomonas; s__Pseudomonas sp. Chol1 |
| ID330 | 17.889085 | 0.006515 | 5.33 | 3.50 | 4.00 | 1.00 | 0.00 | 0.00 | 0.00 | k__Bacteria; p__Proteobacteria; c__Gammaproteobacteria; o__Pseudomonadales; f__Pseudomonadaceae; g__Pseudomonas; s__Pseudomonas pseudoalcaligenes |
| ID466088 | 17.884021 | 0.006529 | 208.00 | 3.50 | 0.00 | 1.00 | 0.00 | 0.00 | 0.00 | k__Bacteria; p__Proteobacteria; c__Gammaproteobacteria; o__Pseudomonadales; f__Moraxellaceae; g__Acinetobacter; s__Acinetobacter sp. Ver3 |
| ID359337 | 17.882353 | 0.006533 | 25.33 | 0.00 | 0.00 | 0.00 | 0.00 | 0.00 | 0.00 | k__Bacteria; p__Proteobacteria; c__Gammaproteobacteria; o__Cellvibrionales; f__Cellvibrionaceae; g__Marinimicrobium; s__(Marinimicrobium) |
| ID76853 | 17.882353 | 0.006533 | 19.33 | 0.00 | 0.00 | 0.00 | 0.00 | 0.00 | 0.00 | k__Bacteria; p__Firmicutes; c__Bacilli; o__Bacillales; f__Planococcaceae; g__Solibacillus; s__Solibacillus silvestris |
| ID407035 | 17.875862 | 0.006550 | 52.00 | 0.00 | 0.00 | 1.00 | 0.00 | 0.00 | 0.00 | k__Bacteria; p__Firmicutes; c__Bacilli; o__Bacillales; f__Staphylococcaceae; g__Salinicoccus; s__Salinicoccus halodurans |
| ID43263 | 17.866197 | 0.006575 | 10.33 | 7.00 | 9.00 | 1.00 | 0.00 | 0.00 | 0.00 | k__Bacteria; p__Proteobacteria; c__Gammaproteobacteria; o__Pseudomonadales; f__Pseudomonadaceae; g__Pseudomonas; s__Pseudomonas alcaligenes |
| ID574962 | 17.866197 | 0.006575 | 3.67 | 17.50 | 30.00 | 3.00 | 0.00 | 0.00 | 0.00 | k__Bacteria; p__Proteobacteria; c__Gammaproteobacteria; o__Pseudomonadales; f__Pseudomonadaceae; g__Pseudomonas; s__Pseudomonas taeanensis |
| ID1381596 | 17.843478 | 0.006635 | 32.00 | 0.00 | 0.00 | 0.00 | 0.00 | 0.00 | 0.00 | k__Bacteria; p__Proteobacteria; c__Gammaproteobacteria; o__Cellvibrionales; f__Cellvibrionaceae; g__Marinimicrobium; s__Marinimicrobium sp. LS-A18 |
| ID225326 | 17.843478 | 0.006635 | 454.67 | 0.00 | 0.00 | 0.00 | 0.00 | 0.00 | 0.00 | k__Bacteria; p__Actinobacteria <phylum>; c__Actinobacteria; o__Corynebacteriales; f__Corynebacteriaceae; g__Corynebacterium; s__Corynebacterium halotolerans |
| ID37928 | 17.843478 | 0.006635 | 21.00 | 0.00 | 0.00 | 0.00 | 0.00 | 0.00 | 0.00 | k__Bacteria; p__Actinobacteria <phylum>; c__Actinobacteria; o__Micrococcales; f__Micrococcaceae; g__Arthrobacter; s__Arthrobacter crystallopoietes |
| ID1354277 | 17.843478 | 0.006635 | 7.67 | 0.00 | 0.00 | 0.00 | 0.00 | 0.00 | 0.00 | k__Bacteria; p__Firmicutes; c__Bacilli; o__Bacillales; f__Bacillaceae; g__Virgibacillus; s__Virgibacillus sp. CM-4 |
| ID418756 | 17.843478 | 0.006635 | 13.67 | 0.00 | 0.00 | 0.00 | 0.00 | 0.00 | 0.00 | k__Bacteria; p__Firmicutes; c__Bacilli; o__Bacillales; f__Staphylococcaceae; g__Salinicoccus; s__Salinicoccus albus |
| ID53412 | 17.822275 | 0.006692 | 3.67 | 4.00 | 6.00 | 1.00 | 0.00 | 0.00 | 0.00 | k__Bacteria; p__Proteobacteria; c__Gammaproteobacteria; o__Pseudomonadales; f__Pseudomonadaceae; g__Pseudomonas; s__Pseudomonas resinovorans |
| ID353 | 17.802353 | 0.006746 | 4.33 | 3.50 | 7.00 | 1.00 | 0.00 | 0.00 | 0.00 | k__Bacteria; p__Proteobacteria; c__Gammaproteobacteria; o__Pseudomonadales; f__Pseudomonadaceae; g__Azotobacter; s__Azotobacter chroococcum |
| ID300 | 17.634232 | 0.007214 | 9.33 | 8.00 | 13.00 | 1.00 | 0.00 | 0.00 | 0.00 | k__Bacteria; p__Proteobacteria; c__Gammaproteobacteria; o__Pseudomonadales; f__Pseudomonadaceae; g__Pseudomonas; s__Pseudomonas mendocina |
| ID203192 | 17.521201 | 0.007547 | 4.00 | 4.50 | 5.00 | 1.00 | 0.00 | 0.00 | 0.00 | k__Bacteria; p__Proteobacteria; c__Gammaproteobacteria; o__Pseudomonadales; f__Pseudomonadaceae; g__Pseudomonas; s__Pseudomonas chloritidismutans |
| ID374 | 17.066038 | 0.009044 | 4.00 | 8.00 | 2.00 | 3.00 | 0.75 | 0.00 | 1.00 | k__Bacteria; p__Proteobacteria; c__Alphaproteobacteria; o__Rhizobiales; f__Bradyrhizobiaceae; g__Bradyrhizobium; s__(Bradyrhizobium) |
| ID251701 | 16.919214 | 0.009585 | 2.00 | 46.00 | 73.00 | 7.00 | 0.25 | 0.00 | 0.00 | k__Bacteria; p__Proteobacteria; c__Gammaproteobacteria; o__Pseudomonadales; f__Pseudomonadaceae; g__Pseudomonas; s__Pseudomonas syringae group genomosp. 3 |
| ID1496 | 16.895431 | 0.009675 | 4.00 | 7.50 | 2.00 | 437.00 | 28.75 | 11.00 | 137.50 | k__Bacteria; p__Firmicutes; c__Clostridia; o__Clostridiales; f__Peptostreptococcaceae; g__Clostridioides; s__Clostridioides difficile |
| ID1236 | 16.861160 | 0.009807 | 925.00 | 411.50 | 478.00 | 102.00 | 39.25 | 22.75 | 6.50 | k__Bacteria; p__Proteobacteria; c__Gammaproteobacteria; o__(Gammaproteobacteria); f__(Gammaproteobacteria); g__(Gammaproteobacteria); s__(Gammaproteobacteria) |
| ID294 | 16.845652 | 0.009868 | 8.67 | 1403.00 | 1656.00 | 173.00 | 0.00 | 0.00 | 0.25 | k__Bacteria; p__Proteobacteria; c__Gammaproteobacteria; o__Pseudomonadales; f__Pseudomonadaceae; g__Pseudomonas; s__Pseudomonas fluorescens |
| ID33882 | 16.828261 | 0.009936 | 117.67 | 2.00 | 1.00 | 5.00 | 0.75 | 0.00 | 1.00 | k__Bacteria; p__Actinobacteria <phylum>; c__Actinobacteria; o__Micrococcales; f__Microbacteriaceae; g__Microbacterium; s__(Microbacterium) |
| ID29465 | 16.806122 | 0.010023 | 1.33 | 2.00 | 2.00 | 14.00 | 14.75 | 616.00 | 4.25 | k__Bacteria; p__Firmicutes; c__Negativicutes; o__Veillonellales; f__Veillonellaceae; g__Veillonella; s__(Veillonella) |
| ID338 | 16.789130 | 0.010090 | 61.67 | 3.50 | 4.00 | 1.00 | 0.75 | 0.00 | 1.00 | k__Bacteria; p__Proteobacteria; c__Gammaproteobacteria; o__Xanthomonadales; f__Xanthomonadaceae; g__Xanthomonas; s__(Xanthomonas) |
| ID85007 | 16.716429 | 0.010384 | 56.33 | 4.00 | 1.00 | 2.00 | 0.75 | 3.50 | 1.00 | k__Bacteria; p__Actinobacteria <phylum>; c__Actinobacteria; o__Corynebacteriales; f__(Corynebacteriales); g__(Corynebacteriales); s__(Corynebacteriales) |
| ID1504823 | 16.685658 | 0.010511 | 0.33 | 0.00 | 0.00 | 1.00 | 10.00 | 0.00 | 45.25 | k__Bacteria; p__(bacterium LF-3); c__(bacterium LF-3); o__(bacterium LF-3); f__(bacterium LF-3); g__(bacterium LF-3); s__bacterium LF-3 |
| ID796942 | 16.614286 | 0.010810 | 1.00 | 0.00 | 0.00 | 1.00 | 6.00 | 0.00 | 2.75 | k__Bacteria; p__Firmicutes; c__Clostridia; o__Clostridiales; f__Lachnospiraceae; g__Stomatobaculum; s__Stomatobaculum longum |
| ID32033 | 16.603604 | 0.010856 | 230.33 | 2.00 | 1.00 | 1.00 | 0.75 | 0.00 | 1.00 | k__Bacteria; p__Proteobacteria; c__Gammaproteobacteria; o__Xanthomonadales; f__Xanthomonadaceae; g__(Xanthomonadaceae); s__(Xanthomonadaceae) |
| ID2070 | 16.587000 | 0.010927 | 10.00 | 2.50 | 1.00 | 1.00 | 0.75 | 0.00 | 1.00 | k__Bacteria; p__Actinobacteria <phylum>; c__Actinobacteria; o__Pseudonocardiales; f__Pseudonocardiaceae; g__(Pseudonocardiaceae); s__(Pseudonocardiaceae) |
| ID470565 | 16.568182 | 0.011008 | 0.00 | 0.00 | 0.00 | 1.00 | 15.25 | 24.75 | 1.50 | k__Bacteria; p__Bacteroidetes; c__Bacteroidia; o__Bacteroidales; f__Prevotellaceae; g__Prevotella; s__Prevotella histicola |
| ID1301 | 16.436731 | 0.011592 | 10.67 | 6.00 | 12.00 | 102.00 | 33.25 | 527.25 | 42.00 | k__Bacteria; p__Firmicutes; c__Bacilli; o__Lactobacillales; f__Streptococcaceae; g__Streptococcus; s__(Streptococcus) |
| ID186818 | 16.334293 | 0.012068 | 50.33 | 1.00 | 1.00 | 2.00 | 0.75 | 0.00 | 1.00 | k__Bacteria; p__Firmicutes; c__Bacilli; o__Bacillales; f__Planococcaceae; g__(Planococcaceae); s__(Planococcaceae) |
| ID1608996 | 16.255814 | 0.012445 | 0.67 | 10.00 | 17.00 | 4.00 | 0.00 | 0.00 | 0.00 | k__Bacteria; p__Proteobacteria; c__Gammaproteobacteria; o__Pseudomonadales; f__Pseudomonadaceae; g__Pseudomonas; s__Pseudomonas helleri |
| ID33069 | 16.255814 | 0.012445 | 0.67 | 61.00 | 100.00 | 10.00 | 0.00 | 0.00 | 0.00 | k__Bacteria; p__Proteobacteria; c__Gammaproteobacteria; o__Pseudomonadales; f__Pseudomonadaceae; g__Pseudomonas; s__Pseudomonas viridiflava |
| ID930166 | 16.234839 | 0.012547 | 0.67 | 31.50 | 40.00 | 4.00 | 0.00 | 0.00 | 0.00 | k__Bacteria; p__Proteobacteria; c__Gammaproteobacteria; o__Pseudomonadales; f__Pseudomonadaceae; g__Pseudomonas; s__Pseudomonas brassicacearum |
| ID1179778 | 16.234839 | 0.012547 | 0.67 | 31.00 | 63.00 | 6.00 | 0.00 | 0.00 | 0.00 | k__Bacteria; p__Proteobacteria; c__Gammaproteobacteria; o__Pseudomonadales; f__Pseudomonadaceae; g__Pseudomonas; s__Pseudomonas sp. M47T1 |
| ID36746 | 16.234839 | 0.012547 | 0.67 | 69.50 | 106.00 | 12.00 | 0.00 | 0.00 | 0.00 | k__Bacteria; p__Proteobacteria; c__Gammaproteobacteria; o__Pseudomonadales; f__Pseudomonadaceae; g__Pseudomonas; s__Pseudomonas cichorii |
| ID50340 | 16.234839 | 0.012547 | 0.67 | 32.00 | 47.00 | 6.00 | 0.00 | 0.00 | 0.00 | k__Bacteria; p__Proteobacteria; c__Gammaproteobacteria; o__Pseudomonadales; f__Pseudomonadaceae; g__Pseudomonas; s__Pseudomonas fuscovaginae |

| | | | | | | | | | | |
|---|---|---|---|---|---|---|---|---|---|---|
| ID147545 | 16.161168 | 0.012915 | 13.33 | 1.50 | 1.00 | 1.00 | 0.75 | 0.00 | 0.00 | k__Eukaryota; p__Ascomycota; c__Eurotiomycetes; o__(Eurotiomycetes); f__(Eurotiomycetes); g__(Eurotiomycetes); s__(Eurotiomycetes) |
| ID83816 | 16.132780 | 0.013059 | 173.67 | 0.00 | 0.00 | 0.00 | 0.75 | 0.00 | 1.00 | k__Archaea; p__Euryarchaeota; c__Methanobacteria; o__Methanobacteriales; f__Methanobacteriaceae; g__Methanobrevibacter; s__Methanobrevibacter ruminantium |
| ID287 | 16.099065 | 0.013232 | 91.33 | 171.00 | 195.00 | 28.00 | 0.75 | 0.00 | 0.75 | k__Bacteria; p__Proteobacteria; c__Gammaproteobacteria; o__Pseudomonadales; f__Pseudomonadaceae; g__Pseudomonas; s__Pseudomonas aeruginosa |
| ID45667 | 16.085106 | 0.013304 | 5.67 | 1.00 | 0.00 | 1.00 | 0.75 | 0.00 | 1.00 | k__Bacteria; p__Firmicutes; c__Bacilli; o__Bacillales; f__Bacillaceae; g__Halobacillus; s__(Halobacillus) |
| ID182709 | 16.085106 | 0.013304 | 205.33 | 1.00 | 0.00 | 1.00 | 0.75 | 0.00 | 1.00 | k__Bacteria; p__Firmicutes; c__Bacilli; o__Bacillales; f__Bacillaceae; g__Oceanobacillus; s__(Oceanobacillus) |
| ID45669 | 16.085106 | 0.013304 | 147.00 | 1.00 | 0.00 | 1.00 | 0.75 | 0.00 | 1.00 | k__Bacteria; p__Firmicutes; c__Bacilli; o__Bacillales; f__Staphylococcaceae; g__Salinicoccus; s__(Salinicoccus) |
| ID572511 | 16.077465 | 0.013344 | 4.33 | 1.00 | 1.00 | 7.00 | 92.75 | 51.25 | 526.00 | k__Bacteria; p__Firmicutes; c__Clostridia; o__Clostridiales; f__Lachnospiraceae; g__Blautia; s__(Blautia) |
| ID22 | 16.074649 | 0.013359 | 8.33 | 1.50 | 2.00 | 1.00 | 0.75 | 0.00 | 1.00 | k__Bacteria; p__Proteobacteria; c__Gammaproteobacteria; o__Alteromonadales; f__Shewanellaceae; g__Shewanella; s__(Shewanella) |
| ID12916 | 16.070813 | 0.013379 | 14.00 | 4.00 | 3.00 | 2.00 | 0.75 | 0.00 | 0.25 | k__Bacteria; p__Proteobacteria; c__Betaproteobacteria; o__Burkholderiales; f__Comamonadaceae; g__Acidovorax; s__(Acidovorax) |
| ID470 | 16.057998 | 0.013446 | 40.67 | 95.50 | 6.00 | 31.00 | 0.75 | 0.00 | 0.75 | k__Bacteria; p__Proteobacteria; c__Gammaproteobacteria; o__Pseudomonadales; f__Moraxellaceae; g__Acinetobacter; s__Acinetobacter baumannii |
| ID909768 | 16.055449 | 0.013459 | 35.00 | 59.50 | 2.00 | 19.00 | 0.75 | 0.25 | 1.00 | k__Bacteria; p__Proteobacteria; c__Gammaproteobacteria; o__Pseudomonadales; f__Moraxellaceae; g__Acinetobacter; s__(Acinetobacter) |
| ID135621 | 16.055449 | 0.013459 | 119.33 | 50.00 | 71.00 | 13.00 | 0.75 | 0.25 | 1.00 | k__Bacteria; p__Proteobacteria; c__Gammaproteobacteria; o__Pseudomonadales; f__Pseudomonadaceae; g__(Pseudomonadaceae); s__(Pseudomonadaceae) |
| ID451864 | 16.055449 | 0.013459 | 8.33 | 62.50 | 42.00 | 4.00 | 0.75 | 0.25 | 1.00 | k__Eukaryota; p__(Eukaryota); c__(Eukaryota); o__(Eukaryota); f__(Eukaryota); g__(Eukaryota); s__(Eukaryota) |
| ID716546 | 16.055449 | 0.013459 | 65.00 | 16.00 | 34.00 | 2.00 | 0.75 | 0.25 | 1.00 | k__Eukaryota; p__Ascomycota; c__(Ascomycota); o__(Ascomycota); f__(Ascomycota); g__(Ascomycota); s__(Ascomycota) |
| ID1232447 | 16.041420 | 0.013533 | 0.33 | 0.00 | 0.00 | 1.00 | 4.25 | 0.00 | 6.25 | k__Bacteria; p__Firmicutes; c__Clostridia; o__Clostridiales; f__(Clostridiales bacterium VE202-09); g__(Clostridiales bacterium VE202-09); s__Clostridiales bacterium VE202-09 |
| ID226910 | 16.026316 | 0.013613 | 0.33 | 24.00 | 40.00 | 4.00 | 0.00 | 0.00 | 0.00 | k__Bacteria; p__Proteobacteria; c__Gammaproteobacteria; o__Pseudomonadales; f__Pseudomonadaceae; g__Pseudomonas; s__Pseudomonas batumici |
| ID1488328 | 16.026316 | 0.013613 | 0.33 | 17.00 | 31.00 | 4.00 | 0.00 | 0.00 | 0.00 | k__Bacteria; p__Proteobacteria; c__Gammaproteobacteria; o__Pseudomonadales; f__Pseudomonadaceae; g__Pseudomonas; s__Pseudomonas sp. Ant30-3 |
| ID35841 | 16.017798 | 0.013659 | 6.67 | 1.00 | 1.00 | 1.00 | 0.75 | 0.00 | 1.00 | k__Bacteria; p__Firmicutes; c__Bacilli; o__Bacillales; f__Bacillaceae; g__Bacillus; s__Bacillus thermoamylovorans |
| ID136841 | 16.009346 | 0.013704 | 37.67 | 22.50 | 34.00 | 5.00 | 0.75 | 0.00 | 0.75 | k__Bacteria; p__Proteobacteria; c__Gammaproteobacteria; o__Pseudomonadales; f__Pseudomonadaceae; g__Pseudomonas; s__(Pseudomonas) |
| ID359110 | 16.002920 | 0.013738 | 0.33 | 148.50 | 166.00 | 19.00 | 0.00 | 0.00 | 0.00 | k__Bacteria; p__Proteobacteria; c__Gammaproteobacteria; o__Pseudomonadales; f__Pseudomonadaceae; g__Pseudomonas; s__Pseudomonas extremaustralis |
| ID75612 | 16.002920 | 0.013738 | 0.33 | 114.50 | 152.00 | 15.00 | 0.00 | 0.00 | 0.00 | k__Bacteria; p__Proteobacteria; c__Gammaproteobacteria; o__Pseudomonadales; f__Pseudomonadaceae; g__Pseudomonas; s__Pseudomonas mandelii |
| ID243924 | 16.002920 | 0.013738 | 0.33 | 58.50 | 90.00 | 10.00 | 0.00 | 0.00 | 0.00 | k__Bacteria; p__Proteobacteria; c__Gammaproteobacteria; o__Pseudomonadales; f__Pseudomonadaceae; g__Pseudomonas; s__Pseudomonas lutea |
| ID1506582 | 16.002920 | 0.013738 | 0.33 | 38.00 | 60.00 | 8.00 | 0.00 | 0.00 | 0.00 | k__Bacteria; p__Proteobacteria; c__Gammaproteobacteria; o__Pseudomonadales; f__Pseudomonadaceae; g__Pseudomonas; s__Pseudomonas sp. 11/12A |
| ID911242 | 16.002920 | 0.013738 | 0.33 | 42.00 | 66.00 | 8.00 | 0.00 | 0.00 | 0.00 | k__Bacteria; p__Proteobacteria; c__Gammaproteobacteria; o__Pseudomonadales; f__Pseudomonadaceae; g__Pseudomonas; s__Pseudomonas sp. CFII64 |
| ID1463662 | 16.002920 | 0.013738 | 0.33 | 88.50 | 80.00 | 7.00 | 0.00 | 0.00 | 0.00 | k__Bacteria; p__Proteobacteria; c__Gammaproteobacteria; o__Pseudomonadales; f__Pseudomonadaceae; g__Pseudomonas; s__Pseudomonas sp. CHM02 |
| ID1674920 | 16.002920 | 0.013738 | 0.33 | 149.00 | 175.00 | 17.00 | 0.00 | 0.00 | 0.00 | k__Bacteria; p__Proteobacteria; c__Gammaproteobacteria; o__Pseudomonadales; f__Pseudomonadaceae; g__Pseudomonas; s__Pseudomonas sp. KG01 |
| ID1397282 | 16.002920 | 0.013738 | 0.33 | 38.00 | 61.00 | 4.00 | 0.00 | 0.00 | 0.00 | k__Bacteria; p__Proteobacteria; c__Gammaproteobacteria; o__Pseudomonadales; f__Pseudomonadaceae; g__Pseudomonas; s__Pseudomonas sp. PH1b |
| ID1470593 | 16.002920 | 0.013738 | 0.33 | 69.00 | 94.00 | 7.00 | 0.00 | 0.00 | 0.00 | k__Bacteria; p__Proteobacteria; c__Gammaproteobacteria; o__Pseudomonadales; f__Pseudomonadaceae; g__Pseudomonas; s__Pseudomonas sp. RIT357 |
| ID1415630 | 16.002920 | 0.013738 | 0.33 | 86.50 | 115.00 | 11.00 | 0.00 | 0.00 | 0.00 | k__Bacteria; p__Proteobacteria; c__Gammaproteobacteria; o__Pseudomonadales; f__Pseudomonadaceae; g__Pseudomonas; s__Pseudomonas sp. TKP |
| ID29438 | 16.002920 | 0.013738 | 0.33 | 61.50 | 33.00 | 6.00 | 0.00 | 0.00 | 0.00 | k__Bacteria; p__Proteobacteria; c__Gammaproteobacteria; o__Pseudomonadales; f__Pseudomonadaceae; g__Pseudomonas; s__Pseudomonas savastanoi |
| ID69277 | 16.000000 | 0.013754 | 6.67 | 1.00 | 1.00 | 1.00 | 0.75 | 0.00 | 1.00 | k__Bacteria; p__Proteobacteria; c__Alphaproteobacteria; o__Rhizobiales; f__Phyllobacteriaceae; g__(Phyllobacteriaceae); s__(Phyllobacteriaceae) |
| ID135614 | 16.000000 | 0.013754 | 30.33 | 1.00 | 1.00 | 1.00 | 0.75 | 0.00 | 1.00 | k__Bacteria; p__Proteobacteria; c__Gammaproteobacteria; o__Xanthomonadales; f__(Xanthomonadales); g__(Xanthomonadales); s__(Xanthomonadales) |
| ID28056 | 16.000000 | 0.013754 | 146.67 | 1.00 | 1.00 | 1.00 | 0.75 | 0.00 | 1.00 | k__Bacteria; p__Actinobacteria <phylum>; c__Actinobacteria; o__Micromonosporales; f__Micromonosporaceae; g__(Micromonosporaceae); s__(Micromonosporaceae) |
| ID129337 | 16.000000 | 0.013754 | 8.67 | 1.00 | 1.00 | 1.00 | 0.75 | 0.00 | 1.00 | k__Bacteria; p__Firmicutes; c__Bacilli; o__Bacillales; f__Bacillaceae; g__Geobacillus; s__(Geobacillus) |
| ID122355 | 15.967105 | 0.013932 | 0.33 | 18.50 | 19.00 | 6.00 | 0.00 | 0.00 | 0.00 | k__Bacteria; p__Proteobacteria; c__Gammaproteobacteria; o__Pseudomonadales; f__Pseudomonadaceae; g__Pseudomonas; s__Pseudomonas psychrophila |
| ID279813 | 15.917098 | 0.014206 | 6.00 | 0.50 | 0.00 | 0.00 | 0.00 | 0.00 | 0.00 | k__Bacteria; p__Firmicutes; c__Bacilli; o__Bacillales; f__Bacillaceae; g__Thalassobacillus; s__Thalassobacillus devorans |
| ID316 | 15.911704 | 0.014236 | 1292.67 | 56.00 | 80.00 | 9.00 | 0.25 | 0.00 | 0.25 | k__Bacteria; p__Proteobacteria; c__Gammaproteobacteria; o__Pseudomonadales; f__Pseudomonadaceae; g__Pseudomonas; s__Pseudomonas stutzeri |
| ID586413 | 15.889655 | 0.014359 | 19.00 | 0.50 | 0.00 | 0.00 | 0.00 | 0.00 | 0.00 | k__Bacteria; p__Firmicutes; c__Bacilli; o__Bacillales; f__Bacillaceae; g__Oceanobacillus; s__Oceanobacillus manasiensis |
| ID254759 | 15.867299 | 0.014484 | 5.00 | 1.00 | 0.00 | 1.00 | 0.25 | 0.00 | 0.00 | k__Bacteria; p__Firmicutes; c__Bacilli; o__Bacillales; f__Bacillaceae; g__Bacillus; s__Bacillus fordii |
| ID303 | 15.798683 | 0.014876 | 14.67 | 206.50 | 216.00 | 70.00 | 0.75 | 0.00 | 0.50 | k__Bacteria; p__Proteobacteria; c__Gammaproteobacteria; o__Pseudomonadales; f__Pseudomonadaceae; g__Pseudomonas; s__Pseudomonas putida |
| ID136849 | 15.798683 | 0.014876 | 26.00 | 1070.00 | 1483.00 | 264.00 | 0.75 | 0.00 | 0.50 | k__Bacteria; p__Proteobacteria; c__Gammaproteobacteria; o__Pseudomonadales; f__Pseudomonadaceae; g__Pseudomonas; s__(Pseudomonas) |
| ID13687 | 15.754294 | 0.015135 | 20.67 | 2.00 | 1.00 | 5.00 | 0.75 | 0.00 | 0.75 | k__Bacteria; p__Proteobacteria; c__Alphaproteobacteria; o__Sphingomonadales; f__Sphingomonadaceae; g__Sphingomonas; s__(Sphingomonas) |
| ID2742 | 15.754294 | 0.015135 | 20.00 | 5.00 | 8.00 | 1.00 | 0.75 | 0.00 | 0.75 | k__Bacteria; p__Proteobacteria; c__Gammaproteobacteria; o__Alteromonadales; f__Alteromonadaceae; g__Marinobacter; s__(Marinobacter) |
| ID2745 | 15.754294 | 0.015135 | 23.67 | 2.00 | 3.00 | 1.00 | 0.75 | 0.00 | 0.75 | k__Bacteria; p__Proteobacteria; c__Gammaproteobacteria; o__Oceanospirillales; f__Halomonadaceae; g__Halomonas; s__(Halomonas) |
| ID1827 | 15.754294 | 0.015135 | 160.00 | 4.00 | 1.00 | 10.00 | 0.75 | 0.00 | 0.75 | k__Bacteria; p__Actinobacteria <phylum>; c__Actinobacteria; o__Corynebacteriales; f__Nocardiaceae; g__Rhodococcus; s__(Rhodococcus) |
| ID147550 | 15.739276 | 0.015224 | 13.33 | 2.50 | 4.00 | 1.00 | 0.75 | 0.00 | 0.75 | k__Eukaryota; p__Ascomycota; c__Sordariomycetes; o__(Sordariomycetes); f__(Sordariomycetes); g__(Sordariomycetes); s__(Sordariomycetes) |
| ID40324 | 15.726087 | 0.015302 | 42.00 | 5.00 | 3.00 | 1.00 | 0.75 | 0.00 | 0.25 | k__Bacteria; p__Proteobacteria; c__Gammaproteobacteria; o__Xanthomonadales; f__Xanthomonadaceae; g__Stenotrophomonas; s__Stenotrophomonas maltophilia |
| ID136843 | 15.694581 | 0.015491 | 5.33 | 1383.50 | 1747.00 | 223.00 | 0.25 | 0.00 | 0.50 | k__Bacteria; p__Proteobacteria; c__Gammaproteobacteria; o__Pseudomonadales; f__Pseudomonadaceae; g__Pseudomonas; s__(Pseudomonas) |
| ID578833 | 15.694581 | 0.015491 | 49.00 | 7.00 | 8.00 | 2.00 | 0.50 | 0.00 | 0.25 | k__Bacteria; p__Proteobacteria; c__Gammaproteobacteria; o__Pseudomonadales; f__Pseudomonadaceae; g__Pseudomonas; s__(Pseudomonas) |
| ID468 | 15.651980 | 0.015749 | 13.67 | 7.00 | 1.00 | 4.00 | 0.75 | 0.25 | 1.00 | k__Bacteria; p__Proteobacteria; c__Gammaproteobacteria; o__Pseudomonadales; f__Moraxellaceae; g__(Moraxellaceae); s__(Moraxellaceae) |
| ID469 | 15.624302 | 0.015919 | 542.00 | 1088.50 | 25.00 | 448.00 | 1.00 | 0.25 | 1.00 | k__Bacteria; p__Proteobacteria; c__Gammaproteobacteria; o__Pseudomonadales; f__Moraxellaceae; g__Acinetobacter; s__(Acinetobacter) |
| ID285 | 15.557673 | 0.016336 | 27.67 | 2.50 | 5.00 | 1.00 | 0.25 | 0.00 | 0.25 | k__Bacteria; p__Proteobacteria; c__Betaproteobacteria; o__Burkholderiales; f__Comamonadaceae; g__Comamonas; s__Comamonas testosteroni |
| ID46228 | 15.493142 | 0.016749 | 1.00 | 0.00 | 0.00 | 3.00 | 56.75 | 17.75 | 175.00 | k__Bacteria; p__Firmicutes; c__Clostridia; o__Clostridiales; f__Ruminococcaceae; g__Ruminococcus; s__Ruminococcus lactaris |
| ID29580 | 15.422819 | 0.017211 | 2.67 | 51.00 | 4.00 | 102.00 | 0.50 | 0.00 | 0.50 | k__Bacteria; p__Proteobacteria; c__Betaproteobacteria; o__Burkholderiales; f__Oxalobacteraceae; g__Janthinobacterium; s__(Janthinobacterium) |
| ID68287 | 15.398707 | 0.017372 | 5.33 | 2.00 | 2.00 | 1.00 | 0.75 | 0.00 | 0.50 | k__Bacteria; p__Proteobacteria; c__Alphaproteobacteria; o__Rhizobiales; f__Phyllobacteriaceae; g__Mesorhizobium; s__(Mesorhizobium) |
| ID333367 | 15.361061 | 0.017627 | 0.33 | 0.00 | 0.00 | 1.00 | 12.50 | 5.00 | 24.75 | k__Bacteria; p__Firmicutes; c__Clostridia; o__Clostridiales; f__Lachnospiraceae; g__Lachnoclostridium; s__[Clostridium] asparagiforme |
| ID698769 | 15.360000 | 0.017634 | 30.00 | 1.00 | 0.00 | 1.00 | 0.50 | 0.00 | 0.00 | k__Bacteria; p__Firmicutes; c__Bacilli; o__Bacillales; f__Bacillaceae; g__Virgibacillus; s__Virgibacillus alimentarius |
| ID305719 | 15.356757 | 0.017656 | 0.33 | 0.00 | 0.00 | 1.00 | 44.00 | 24.75 | 4.25 | k__Bacteria; p__Bacteroidetes; c__Bacteroidia; o__Bacteroidales; f__Prevotellaceae; g__Prevotella; s__Prevotella baroniae |
| ID2053 | 15.353457 | 0.017679 | 9.67 | 3.00 | 0.00 | 1.00 | 0.75 | 0.00 | 0.75 | k__Bacteria; p__Actinobacteria <phylum>; c__Actinobacteria; o__Corynebacteriales; f__Gordoniaceae; g__Gordonia; s__(Gordonia) |
| ID265 | 15.352815 | 0.017683 | 88.00 | 2.50 | 1.00 | 3.00 | 0.75 | 0.00 | 0.50 | k__Bacteria; p__Proteobacteria; c__Alphaproteobacteria; o__Rhodobacterales; f__Rhodobacteraceae; g__Paracoccus; s__(Paracoccus) |
| ID1663 | 15.337904 | 0.017785 | 186.33 | 1.00 | 3.00 | 2.00 | 0.75 | 0.00 | 0.75 | k__Bacteria; p__Actinobacteria <phylum>; c__Actinobacteria; o__Micrococcales; f__Micrococcaceae; g__Arthrobacter; s__(Arthrobacter) |
| ID712 | 15.323621 | 0.017884 | 3.33 | 1.50 | 2.00 | 20.00 | 6.25 | 17.75 | 1.50 | k__Bacteria; p__Proteobacteria; c__Gammaproteobacteria; o__Pasteurellales; f__Pasteurellaceae; g__(Pasteurellaceae); s__(Pasteurellaceae) |
| ID59753 | 15.320274 | 0.017907 | 8.00 | 3.50 | 1.00 | 1.00 | 0.75 | 0.00 | 0.75 | k__Bacteria; p__Proteobacteria; c__Gammaproteobacteria; o__Oceanospirillales; f__Alcanivoracaceae; g__Alcanivorax; s__(Alcanivorax) |
| ID411486 | 15.300268 | 0.018046 | 0.67 | 0.00 | 0.00 | 2.00 | 22.25 | 10.25 | 60.75 | k__Bacteria; p__Firmicutes; c__Clostridia; o__Clostridiales; f__Clostridiaceae; g__Clostridium; s__Clostridium sp. M62/1 |
| ID5052 | 15.290634 | 0.018113 | 240.67 | 4.00 | 2.00 | 1.00 | 0.50 | 0.00 | 0.25 | k__Eukaryota; p__Ascomycota; c__Eurotiomycetes; o__Eurotiales; f__Aspergillaceae; g__Aspergillus; s__(Aspergillus) |
| ID665950 | 15.276961 | 0.018209 | 0.00 | 0.00 | 0.00 | 1.00 | 1.75 | 0.00 | 7.25 | k__Bacteria; p__Firmicutes; c__Clostridia; o__Clostridiales; f__Lachnospiraceae; g__(Lachnospiraceae bacterium 3_1_46FAA); s__Lachnospiraceae bacterium 3_1_46FAA |
| ID136846 | 15.275495 | 0.018219 | 54.33 | 4.50 | 6.00 | 1.00 | 0.50 | 0.00 | 0.25 | k__Bacteria; p__Proteobacteria; c__Gammaproteobacteria; o__Pseudomonadales; f__Pseudomonadaceae; g__Pseudomonas; s__(Pseudomonas) |

| | | | | | | | | | | |
|---|---|---|---|---|---|---|---|---|---|---|
| ID28126 | 15.256988 | 0.018349 | 0.33 | 0.00 | 0.00 | 1.00 | 79.50 | 50.75 | 11.25 | k__Bacteria; p__Bacteroidetes; c__Bacteroidia; o__Bacteroidales; f__Prevotellaceae; g__Prevotella; s__Prevotella buccae |
| ID77095 | 15.243243 | 0.018447 | 0.33 | 0.00 | 0.00 | 1.00 | 85.75 | 65.75 | 10.00 | k__Bacteria; p__Bacteroidetes; c__Bacteroidia; o__Bacteroidales; f__Prevotellaceae; g__Prevotella; s__Prevotella bryantii |
| ID77768 | 15.212353 | 0.018668 | 0.33 | 0.00 | 0.00 | 1.00 | 43.00 | 26.50 | 3.75 | k__Bacteria; p__Bacteroidetes; c__Bacteroidia; o__Bacteroidales; f__Prevotellaceae; g__Prevotella; s__Prevotella albensis |
| ID80865 | 15.196262 | 0.018784 | 10.00 | 2.00 | 1.00 | 1.00 | 0.25 | 0.00 | 0.25 | k__Bacteria; p__Proteobacteria; c__Betaproteobacteria; o__Burkholderiales; f__Comamonadaceae; g__Delftia; s__(Delftia) |
| ID363265 | 15.171582 | 0.018963 | 0.67 | 1.00 | 0.00 | 16.00 | 1399.00 | 507.75 | 27.00 | k__Bacteria; p__Bacteroidetes; c__Bacteroidia; o__Bacteroidales; f__Prevotellaceae; g__Prevotella; s__Prevotella stercorea |
| ID1542 | 15.131323 | 0.019260 | 0.33 | 45.00 | 285.00 | 16.00 | 0.75 | 0.00 | 1.25 | k__Bacteria; p__Firmicutes; c__Clostridia; o__Clostridiales; f__Clostridiaceae; g__Clostridium; s__Clostridium novyi |
| ID641107 | 15.131323 | 0.019260 | 0.33 | 16.00 | 5.00 | 12.00 | 0.75 | 0.00 | 1.25 | k__Bacteria; p__Firmicutes; c__Clostridia; o__Clostridiales; f__Clostridiaceae; g__Clostridium; s__Clostridium sp. DL-VIII |
| ID91623 | 15.113250 | 0.019394 | 0.33 | 130.50 | 32.00 | 63.00 | 0.75 | 0.00 | 1.00 | k__Bacteria; p__Firmicutes; c__Clostridia; o__Clostridiales; f__Clostridiaceae; g__Clostridium; s__Clostridium akagii |
| ID37659 | 15.113250 | 0.019394 | 0.33 | 2572.00 | 11.00 | 17.00 | 0.75 | 0.00 | 1.00 | k__Bacteria; p__Firmicutes; c__Clostridia; o__Clostridiales; f__Clostridiaceae; g__Clostridium; s__Clostridium algidicarnis |
| ID217159 | 15.113250 | 0.019394 | 0.33 | 22.50 | 4.00 | 15.00 | 0.75 | 0.00 | 1.00 | k__Bacteria; p__Firmicutes; c__Clostridia; o__Clostridiales; f__Clostridiaceae; g__Clostridium; s__Clostridium carboxidivorans |
| ID319475 | 15.113250 | 0.019394 | 0.33 | 19.00 | 4.00 | 13.00 | 0.75 | 0.00 | 1.00 | k__Bacteria; p__Firmicutes; c__Clostridia; o__Clostridiales; f__Clostridiaceae; g__Clostridium; s__Clostridium lundense |
| ID1354301 | 15.113250 | 0.019394 | 0.33 | 24.00 | 4.00 | 18.00 | 0.75 | 0.00 | 1.00 | k__Bacteria; p__Firmicutes; c__Clostridia; o__Clostridiales; f__Clostridiaceae; g__Clostridium; s__Clostridium sp. BL8 |
| ID136845 | 15.090974 | 0.019561 | 1.67 | 17.00 | 12.00 | 7.00 | 0.75 | 0.00 | 0.50 | k__Bacteria; p__Proteobacteria; c__Gammaproteobacteria; o__Pseudomonadales; f__Pseudomonadaceae; g__Pseudomonas; s__(Pseudomonas) |
| ID1042156 | 15.073221 | 0.019695 | 3.67 | 0.00 | 0.00 | 1.00 | 5.25 | 1.25 | 19.25 | k__Bacteria; p__Firmicutes; c__Clostridia; o__Clostridiales; f__Clostridiaceae; g__Clostridium; s__Clostridium sp. SY8519 |
| ID28117 | 15.071844 | 0.019705 | 2.00 | 0.00 | 0.00 | 1.00 | 8.75 | 2.50 | 279.50 | k__Bacteria; p__Bacteroidetes; c__Bacteroidia; o__Bacteroidales; f__Rikenellaceae; g__Alistipes; s__Alistipes putredinis |
| ID39496 | 15.067316 | 0.019740 | 1.00 | 0.50 | 0.00 | 5.00 | 27.75 | 5.25 | 48.75 | k__Bacteria; p__Firmicutes; c__Clostridia; o__Clostridiales; f__Eubacteriaceae; g__Eubacterium; s__Eubacterium ventriosum |
| ID1487921 | 15.059250 | 0.019801 | 0.33 | 15.50 | 4.00 | 14.00 | 0.75 | 0.00 | 1.00 | k__Bacteria; p__Firmicutes; c__Clostridia; o__Clostridiales; f__Clostridiaceae; g__Clostridium; s__Clostridium sp. HMP27 |
| ID655338 | 15.059250 | 0.019801 | 0.33 | 10.00 | 6.00 | 9.00 | 0.75 | 0.00 | 1.00 | k__Bacteria; p__Firmicutes; c__Clostridia; o__Clostridiales; f__Clostridiaceae; g__Fervidicella; s__Fervidicella metallireducens |
| ID1226324 | 15.058363 | 0.019808 | 0.67 | 0.00 | 0.00 | 2.00 | 43.75 | 30.75 | 761.50 | k__Bacteria; p__Firmicutes; c__Clostridia; o__Clostridiales; f__Lachnospiraceae; g__Blautia; s__Blautia sp. KLE 1732 |
| ID724 | 15.050474 | 0.019868 | 0.67 | 1.00 | 2.00 | 16.00 | 4.50 | 27.50 | 1.25 | k__Bacteria; p__Proteobacteria; c__Gammaproteobacteria; o__Pasteurellales; f__Pasteurellaceae; g__Haemophilus; s__(Haemophilus) |
| ID1352 | 15.039084 | 0.019955 | 1.67 | 1.00 | 1.00 | 1.00 | 4.75 | 86.25 | 21.75 | k__Bacteria; p__Firmicutes; c__Bacilli; o__Lactobacillales; f__Enterococcaceae; g__Enterococcus; s__Enterococcus faecium |
| ID166486 | 15.010629 | 0.020174 | 0.67 | 1.00 | 0.00 | 15.00 | 133.50 | 23.00 | 407.50 | k__Bacteria; p__Firmicutes; c__Clostridia; o__Clostridiales; f__Lachnospiraceae; g__Roseburia; s__Roseburia intestinalis |
| ID1232459 | 14.985071 | 0.020373 | 0.33 | 0.00 | 0.00 | 1.00 | 18.75 | 1.00 | 20.50 | k__Bacteria; p__Firmicutes; c__Clostridia; o__Clostridiales; f__Oscillospiraceae; g__(Oscillospiraceae bacterium VE202-24); s__Oscillospiraceae bacterium VE202-24 |
| ID72274 | 14.977230 | 0.020435 | 22.33 | 30.50 | 30.00 | 7.00 | 2.25 | 1.00 | 1.00 | k__Bacteria; p__Proteobacteria; c__Gammaproteobacteria; o__Pseudomonadales; f__(Pseudomonadales); g__(Pseudomonadales); s__(Pseudomonadales) |
| ID375288 | 14.967621 | 0.020510 | 9.67 | 1.00 | 1.00 | 4.00 | 117.75 | 42.25 | 165.50 | k__Bacteria; p__Bacteroidetes; c__Bacteroidia; o__Bacteroidales; f__Porphyromonadaceae; g__Parabacteroides; s__(Parabacteroides) |
| ID1531 | 14.935448 | 0.020765 | 1.67 | 1.00 | 0.00 | 2.00 | 30.25 | 13.25 | 124.50 | k__Bacteria; p__Firmicutes; c__Clostridia; o__Clostridiales; f__Lachnospiraceae; g__Lachnoclostridium; s__[Clostridium] clostridioforme |
| ID563835 | 14.930180 | 0.020807 | 5.33 | 1.00 | 1.00 | 5.00 | 1.00 | 0.00 | 1.00 | k__Bacteria; p__Bacteroidetes; c__Chitinophagia; o__Chitinophagales; f__Chitinophagaceae; g__(Chitinophagaceae); s__(Chitinophagaceae) |
| ID469614 | 14.924918 | 0.020849 | 0.33 | 0.00 | 0.00 | 0.00 | 1.75 | 0.00 | 7.00 | k__Bacteria; p__Firmicutes; c__Erysipelotrichia; o__Erysipelotrichales; f__Erysipelotrichaceae; g__(Erysipelotrichaceae bacterium 6_1_45); s__Erysipelotrichaceae bacterium 6_1_45 |
| ID649724 | 14.916825 | 0.020914 | 0.33 | 0.00 | 0.00 | 1.00 | 32.00 | 3.25 | 31.00 | k__Bacteria; p__Firmicutes; c__Clostridia; o__Clostridiales; f__Clostridiaceae; g__Clostridium; s__Clostridium sp. ATCC BAA-442 |
| ID405946 | 14.896552 | 0.021077 | 25.00 | 0.00 | 0.00 | 0.00 | 0.00 | 0.00 | 0.25 | k__Bacteria; p__Firmicutes; c__Bacilli; o__Bacillales; f__Bacillaceae; g__Oceanobacillus; s__Oceanobacillus caeni |
| ID222 | 14.894118 | 0.021096 | 6.00 | 1.50 | 2.00 | 1.00 | 0.75 | 0.00 | 0.75 | k__Bacteria; p__Proteobacteria; c__Betaproteobacteria; o__Burkholderiales; f__Alcaligenaceae; g__Achromobacter; s__(Achromobacter) |
| ID80864 | 14.886872 | 0.021155 | 141.33 | 19.50 | 11.00 | 35.00 | 0.75 | 0.75 | 1.00 | k__Bacteria; p__Proteobacteria; c__Betaproteobacteria; o__Burkholderiales; f__Comamonadaceae; g__(Comamonadaceae); s__(Comamonadaceae) |
| ID1883 | 14.876776 | 0.021237 | 166.67 | 3.00 | 2.00 | 5.00 | 2.50 | 7.50 | 2.00 | k__Bacteria; p__Actinobacteria <phylum>; c__Actinobacteria; o__Streptomycetales; f__Streptomycetaceae; g__Streptomyces; s__(Streptomyces) |
| ID197 | 14.851770 | 0.021441 | 0.00 | 0.00 | 0.00 | 1.00 | 8.50 | 0.75 | 3.00 | k__Bacteria; p__Proteobacteria; c__Epsilonproteobacteria; o__Campylobacterales; f__Campylobacteraceae; g__Campylobacter; s__Campylobacter jejuni |
| ID1307 | 14.846667 | 0.021483 | 0.33 | 1.00 | 1.00 | 2.00 | 28.50 | 4.25 | 13.50 | k__Bacteria; p__Firmicutes; c__Bacilli; o__Lactobacillales; f__Streptococcaceae; g__Streptococcus; s__Streptococcus suis |
| ID501571 | 14.840888 | 0.021531 | 0.67 | 1.00 | 1.00 | 1.00 | 35.00 | 12.50 | 16.00 | k__Bacteria; p__Firmicutes; c__Clostridia; o__Clostridiales; f__Clostridiaceae; g__Butyricicoccus; s__Butyricicoccus pullicaecorum |
| ID31977 | 14.795945 | 0.021905 | 1.00 | 1.00 | 1.00 | 1.00 | 4.00 | 19.00 | 2.75 | k__Bacteria; p__Firmicutes; c__Negativicutes; o__Veillonellales; f__Veillonellaceae; g__(Veillonellaceae); s__(Veillonellaceae) |
| ID836 | 14.791704 | 0.021940 | 5.33 | 1.00 | 1.00 | 1.00 | 44.75 | 20.00 | 5.50 | k__Bacteria; p__Bacteroidetes; c__Bacteroidia; o__Bacteroidales; f__Porphyromonadaceae; g__Porphyromonas; s__(Porphyromonas) |
| ID90964 | 14.786471 | 0.021984 | 31.67 | 1.00 | 1.00 | 1.00 | 2.25 | 0.25 | 1.00 | k__Bacteria; p__Firmicutes; c__Bacilli; o__Bacillales; f__Staphylococcaceae; g__(Staphylococcaceae); s__(Staphylococcaceae) |
| ID12968 | 14.775773 | 0.022074 | 0.00 | 0.00 | 0.00 | 1.00 | 14.25 | 3.00 | 0.00 | k__Eukaryota; p__(Blastocystis); c__(Blastocystis); o__(Blastocystis); f__(Blastocystis); g__Blastocystis; s__Blastocystis hominis |
| ID74426 | 14.758497 | 0.022221 | 0.33 | 0.00 | 0.00 | 1.00 | 10.50 | 21.25 | 49.50 | k__Bacteria; p__Actinobacteria <phylum>; c__Coriobacteriia; o__Coriobacteriales; f__Coriobacteriaceae; g__Collinsella; s__Collinsella aerofaciens |
| ID36745 | 14.706062 | 0.022670 | 0.33 | 13.50 | 6.00 | 8.00 | 1.25 | 0.00 | 1.50 | k__Bacteria; p__Firmicutes; c__Clostridia; o__Clostridiales; f__Clostridiaceae; g__Clostridium; s__Clostridium saccharoperbutylacetonicum |
| ID237 | 14.700134 | 0.022722 | 258.00 | 17.00 | 8.00 | 4.00 | 7.00 | 5.50 | 1.00 | k__Bacteria; p__Bacteroidetes; c__Flavobacteriia; o__Flavobacteriales; f__Flavobacteriaceae; g__Flavobacterium; s__(Flavobacterium) |
| ID658656 | 14.678241 | 0.022912 | 0.33 | 0.00 | 0.00 | 1.00 | 2.75 | 0.00 | 8.25 | k__Bacteria; p__Firmicutes; c__Clostridia; o__Clostridiales; f__Lachnospiraceae; g__(Lachnospiraceae bacterium 6_1_37FAA); s__Lachnospiraceae bacterium 6_1_37FAA |
| ID186803 | 14.676316 | 0.022929 | 110.33 | 9.00 | 3.00 | 116.00 | 861.00 | 315.50 | 1442.25 | k__Bacteria; p__Firmicutes; c__Clostridia; o__Clostridiales; f__Lachnospiraceae; g__(Lachnospiraceae); s__(Lachnospiraceae) |
| ID1763 | 14.668750 | 0.022996 | 27.00 | 6.00 | 2.00 | 5.00 | 0.75 | 2.00 | 1.00 | k__Bacteria; p__Actinobacteria <phylum>; c__Actinobacteria; o__Corynebacteriales; f__Mycobacteriaceae; g__Mycobacterium; s__(Mycobacterium) |
| ID39490 | 14.659647 | 0.023076 | 1.67 | 0.00 | 0.00 | 3.00 | 78.75 | 8.75 | 199.25 | k__Bacteria; p__Firmicutes; c__Clostridia; o__Clostridiales; f__Eubacteriaceae; g__Eubacterium; s__Eubacterium ramulus |
| ID1501 | 14.655773 | 0.023110 | 0.33 | 40.00 | 11.00 | 31.00 | 1.00 | 0.00 | 1.25 | k__Bacteria; p__Firmicutes; c__Clostridia; o__Clostridiales; f__Clostridiaceae; g__Clostridium; s__Clostridium pasteurianum |
| ID1513 | 14.650381 | 0.023157 | 0.33 | 21.00 | 4.00 | 21.00 | 1.00 | 0.00 | 1.50 | k__Bacteria; p__Firmicutes; c__Clostridia; o__Clostridiales; f__Clostridiaceae; g__Clostridium; s__Clostridium tetani |
| ID1224 | 14.647368 | 0.023184 | 492.33 | 343.00 | 420.00 | 119.00 | 32.75 | 25.00 | 14.00 | k__Bacteria; p__Proteobacteria; c__(Proteobacteria); o__(Proteobacteria); f__(Proteobacteria); g__(Proteobacteria); s__(Proteobacteria) |
| ID1321777 | 14.641051 | 0.023240 | 0.33 | 42.00 | 12.00 | 24.00 | 1.00 | 0.00 | 1.00 | k__Bacteria; p__Firmicutes; c__Clostridia; o__Clostridiales; f__(Clostridiales bacterium oral taxon 876); g__(Clostridiales bacterium oral taxon 876); s__Clostridiales bacterium oral taxon 876 |
| ID1578 | 14.605752 | 0.023555 | 6.33 | 4.50 | 13.00 | 5.00 | 14.50 | 236.50 | 6.75 | k__Bacteria; p__Firmicutes; c__Bacilli; o__Lactobacillales; f__Lactobacillaceae; g__Lactobacillus; s__(Lactobacillus) |
| ID1843491 | 14.595588 | 0.023646 | 2.00 | 1.50 | 1.00 | 4.00 | 97.00 | 28.25 | 6.25 | k__Bacteria; p__Firmicutes; c__Negativicutes; o__Selenomonadales; f__Selenomonadaceae; g__(Selenomonadaceae); s__(Selenomonadaceae) |
| ID1602168 | 14.593324 | 0.023667 | 0.33 | 0.00 | 0.00 | 1.00 | 69.25 | 45.75 | 4.75 | k__Bacteria; p__Bacteroidetes; c__Bacteroidia; o__Bacteroidales; f__Prevotellaceae; g__Prevotella; s__Prevotella sp. P4-65 |
| ID450746 | 14.584305 | 0.023748 | 0.33 | 1.50 | 1.00 | 8.00 | 10.50 | 1.50 | 23.50 | k__Bacteria; p__Firmicutes; c__Erysipelotrichia; o__Erysipelotrichales; f__Erysipelotrichaceae; g__Coprobacillus; s__Coprobacillus sp. 8_1_38FAA |
| ID360807 | 14.557726 | 0.023990 | 0.67 | 1.00 | 0.00 | 9.00 | 343.25 | 26.00 | 475.50 | k__Bacteria; p__Firmicutes; c__Clostridia; o__Clostridiales; f__Lachnospiraceae; g__Roseburia; s__Roseburia inulinivorans |
| ID102106 | 14.531333 | 0.024232 | 1.67 | 0.50 | 0.00 | 1.00 | 51.00 | 305.75 | 262.00 | k__Bacteria; p__Actinobacteria <phylum>; c__Coriobacteriia; o__Coriobacteriales; f__Coriobacteriaceae; g__Collinsella; s__(Collinsella) |
| ID1232443 | 14.502183 | 0.024503 | 0.00 | 0.00 | 0.00 | 1.00 | 4.50 | 0.00 | 8.25 | k__Bacteria; p__Firmicutes; c__Clostridia; o__Clostridiales; f__(Clostridiales bacterium VE202-13); g__(Clostridiales bacterium VE202-13); s__Clostridiales bacterium VE202-13 |
| ID1550024 | 14.498815 | 0.024534 | 0.33 | 0.00 | 0.00 | 1.00 | 16.50 | 2.50 | 49.75 | k__Bacteria; p__Firmicutes; c__Clostridia; o__Clostridiales; f__Ruminococcaceae; g__Ruthenibacterium; s__Ruthenibacterium lactatiformans |
| ID1540257 | 14.484017 | 0.024673 | 0.33 | 25.00 | 9.00 | 17.00 | 1.25 | 0.00 | 1.00 | k__Bacteria; p__Firmicutes; c__Clostridia; o__Clostridiales; f__Clostridiaceae; g__Clostridium; s__Clostridium sp. KNHs214 |
| ID61171 | 14.483618 | 0.024677 | 0.33 | 0.00 | 0.00 | 1.00 | 15.50 | 2.00 | 10.75 | k__Bacteria; p__Firmicutes; c__Erysipelotrichia; o__Erysipelotrichales; f__Erysipelotrichaceae; g__Holdemania; s__Holdemania filiformis |
| ID1380 | 14.447475 | 0.025018 | 2.67 | 1.00 | 0.00 | 1.00 | 5.25 | 26.25 | 3.25 | k__Bacteria; p__Actinobacteria <phylum>; c__Coriobacteriia; o__Coriobacteriales; f__Atopobiaceae; g__Atopobium; s__(Atopobium) |
| ID28216 | 14.443548 | 0.025055 | 39.33 | 17.00 | 23.00 | 13.00 | 4.75 | 4.75 | 2.00 | k__Bacteria; p__Proteobacteria; c__Betaproteobacteria; o__(Betaproteobacteria); f__(Betaproteobacteria); g__(Betaproteobacteria); s__(Betaproteobacteria) |
| ID5204 | 14.438250 | 0.025106 | 1.00 | 246.50 | 130.00 | 12.00 | 0.75 | 0.00 | 0.50 | k__Eukaryota; p__Basidiomycota; c__(Basidiomycota); o__(Basidiomycota); f__(Basidiomycota); g__(Basidiomycota); s__(Basidiomycota) |
| ID970 | 14.435484 | 0.025132 | 3.00 | 1.00 | 1.00 | 3.00 | 38.50 | 11.50 | 5.25 | k__Bacteria; p__Firmicutes; c__Negativicutes; o__Selenomonadales; f__Selenomonadaceae; g__Selenomonas; s__(Selenomonas) |
| ID1519 | 14.433962 | 0.025147 | 0.33 | 17.00 | 4.00 | 18.00 | 1.25 | 0.00 | 1.25 | k__Bacteria; p__Firmicutes; c__Clostridia; o__Clostridiales; f__Clostridiaceae; g__Clostridium; s__Clostridium tyrobutyricum |
| ID768507 | 14.413662 | 0.025342 | 90.33 | 1.00 | 1.00 | 1.00 | 7.25 | 2.25 | 1.00 | k__Bacteria; p__Bacteroidetes; c__Cytophagia; o__Cytophagales; f__(Cytophagales); g__(Cytophagales); s__(Cytophagales) |

| | | | | | | | | | | |
|---|---|---|---|---|---|---|---|---|---|---|
| ID53443 | 14.404167 | 0.025433 | 0.67 | 0.00 | 0.00 | 2.00 | 12.75 | 3.25 | 24.75 | k__Bacteria; p__Firmicutes; c__Clostridia; o__Clostridiales; f__Lachnospiraceae; g__Blautia; s__Blautia hydrogenotrophica |
| ID85021 | 14.398649 | 0.025487 | 6.67 | 1.00 | 0.00 | 3.00 | 0.75 | 0.00 | 0.50 | k__Bacteria; p__Actinobacteria <phylum>; c__Actinobacteria; o__Micrococcales; f__Intrasporangiaceae; g__(Intrasporangiaceae); s__(Intrasporangiaceae) |
| ID451871 | 14.384250 | 0.025626 | 41.00 | 1.00 | 2.00 | 1.00 | 0.75 | 0.00 | 0.50 | k__Eukaryota; p__Ascomycota; c__Eurotiomycetes; o__(Eurotiomycetes); f__(Eurotiomycetes); g__(Eurotiomycetes); s__(Eurotiomycetes) |
| ID75682 | 14.377803 | 0.025689 | 24.67 | 13.00 | 5.00 | 44.00 | 3.75 | 1.25 | 3.25 | k__Bacteria; p__Proteobacteria; c__Betaproteobacteria; o__Burkholderiales; f__Oxalobacteraceae; g__(Oxalobacteraceae); s__(Oxalobacteraceae) |
| ID652706 | 14.376337 | 0.025703 | 2.00 | 0.00 | 0.00 | 1.00 | 10.50 | 3.50 | 6.00 | k__Bacteria; p__Firmicutes; c__Clostridia; o__Clostridiales; f__Lachnospiraceae; g__Oribacterium; s__Oribacterium sp. oral taxon 078 |
| ID658089 | 14.344011 | 0.026021 | 0.33 | 0.00 | 0.00 | 1.00 | 9.50 | 1.50 | 28.00 | k__Bacteria; p__Firmicutes; c__Clostridia; o__Clostridiales; f__Lachnospiraceae; g__(Lachnospiraceae bacterium 5_1_63FAA); s__Lachnospiraceae bacterium 5_1_63FAA |
| ID258 | 14.340249 | 0.026058 | 30.00 | 1.00 | 0.00 | 1.00 | 0.75 | 0.00 | 0.25 | k__Bacteria; p__Bacteroidetes; c__Sphingobacteriia; o__Sphingobacteriales; f__Sphingobacteriaceae; g__Sphingobacterium; s__Sphingobacterium spiritivorum |
| ID83676 | 14.340249 | 0.026058 | 6.33 | 1.00 | 0.00 | 1.00 | 0.75 | 0.00 | 0.75 | k__Bacteria; p__Actinobacteria <phylum>; c__Actinobacteria; o__Streptosporangiales; f__Nocardiopsaceae; g__(Nocardiopsaceae); s__(Nocardiopsaceae) |
| ID1679169 | 14.340249 | 0.026058 | 176.67 | 0.00 | 0.00 | 1.00 | 0.75 | 0.00 | 0.75 | k__Bacteria; p__Firmicutes; c__Bacilli; o__Bacillales; f__Bacillaceae; g__Bacillus; s__Bacillus sp. FJAT-27916 |
| ID161536 | 14.340249 | 0.026058 | 106.67 | 1.00 | 0.00 | 1.00 | 0.75 | 0.00 | 0.25 | k__Bacteria; p__Firmicutes; c__Bacilli; o__Bacillales; f__Bacillaceae; g__Ornithinibacillus; s__Ornithinibacillus californiensis |
| ID1372 | 14.340249 | 0.026058 | 9.67 | 1.00 | 0.00 | 1.00 | 0.75 | 0.00 | 0.75 | k__Bacteria; p__Firmicutes; c__Bacilli; o__Bacillales; f__Planococcaceae; g__Planococcus; s__(Planococcus) |
| ID1519439 | 14.313109 | 0.026328 | 4.67 | 0.50 | 0.00 | 9.00 | 453.50 | 71.25 | 184.00 | k__Bacteria; p__Firmicutes; c__Clostridia; o__Clostridiales; f__Oscillospiraceae; g__Oscillibacter; s__Oscillibacter sp. ER4 |
| ID119060 | 14.293230 | 0.026527 | 21.00 | 21.50 | 29.00 | 10.00 | 1.00 | 1.25 | 1.00 | k__Bacteria; p__Proteobacteria; c__Betaproteobacteria; o__Burkholderiales; f__Burkholderiaceae; g__(Burkholderiaceae); s__(Burkholderiaceae) |
| ID626940 | 14.289194 | 0.026567 | 5.33 | 0.50 | 0.00 | 2.00 | 568.25 | 26.75 | 198.25 | k__Bacteria; p__Firmicutes; c__Negativicutes; o__Acidaminococcales; f__Acidaminococcaceae; g__Phascolarctobacterium; s__Phascolarctobacterium succinatutens |
| ID841 | 14.282698 | 0.026633 | 1.67 | 1.00 | 1.00 | 14.00 | 99.00 | 18.50 | 130.50 | k__Bacteria; p__Firmicutes; c__Clostridia; o__Clostridiales; f__Lachnospiraceae; g__Roseburia; s__(Roseburia) |
| ID909932 | 14.273313 | 0.026728 | 1.67 | 2.00 | 1.00 | 3.00 | 19.25 | 31.00 | 10.75 | k__Bacteria; p__Firmicutes; c__Negativicutes; o__(Negativicutes); f__(Negativicutes); g__(Negativicutes); s__(Negativicutes) |
| ID517 | 14.246809 | 0.026997 | 9.00 | 1.00 | 1.00 | 1.00 | 0.75 | 0.00 | 0.75 | k__Bacteria; p__Proteobacteria; c__Betaproteobacteria; o__Burkholderiales; f__Alcaligenaceae; g__Bordetella; s__(Bordetella) |
| ID1706371 | 14.246809 | 0.026997 | 408.00 | 1.00 | 1.00 | 1.00 | 0.75 | 0.00 | 0.75 | k__Bacteria; p__Proteobacteria; c__Gammaproteobacteria; o__Cellvibrionales; f__Cellvibrionaceae; g__(Cellvibrionaceae); s__(Cellvibrionaceae) |
| ID85025 | 14.246809 | 0.026997 | 14.00 | 1.00 | 1.00 | 1.00 | 0.75 | 0.00 | 0.75 | k__Bacteria; p__Actinobacteria <phylum>; c__Actinobacteria; o__Corynebacteriales; f__Nocardiaceae; g__(Nocardiaceae); s__(Nocardiaceae) |
| ID2013 | 14.246809 | 0.026997 | 206.00 | 1.00 | 1.00 | 1.00 | 0.75 | 0.00 | 0.75 | k__Bacteria; p__Actinobacteria <phylum>; c__Actinobacteria; o__Streptosporangiales; f__Nocardiopsaceae; g__Nocardiopsis; s__(Nocardiopsis) |
| ID1404 | 14.246809 | 0.026997 | 30.00 | 1.00 | 1.00 | 1.00 | 0.75 | 0.00 | 0.75 | k__Bacteria; p__Firmicutes; c__Bacilli; o__Bacillales; f__Bacillaceae; g__Bacillus; s__Bacillus megaterium |
| ID1304 | 14.242290 | 0.027044 | 0.33 | 0.00 | 0.00 | 1.00 | 1.00 | 62.50 | 4.00 | k__Bacteria; p__Firmicutes; c__Bacilli; o__Lactobacillales; f__Streptococcaceae; g__Streptococcus; s__Streptococcus salivarius |
| ID649756 | 14.238377 | 0.027084 | 0.33 | 0.50 | 0.00 | 1.00 | 21.25 | 3.50 | 78.75 | k__Bacteria; p__Firmicutes; c__Clostridia; o__Clostridiales; f__Lachnospiraceae; g__Anaerostipes; s__Anaerostipes hadrus |
| ID665956 | 14.236364 | 0.027104 | 0.67 | 0.00 | 0.00 | 1.00 | 23.00 | 4.00 | 107.25 | k__Bacteria; p__Firmicutes; c__Clostridia; o__Clostridiales; f__Ruminococcaceae; g__Subdoligranulum; s__Subdoligranulum sp. 4_3_54A2FAA |
| ID1392487 | 14.234232 | 0.027126 | 17.33 | 0.00 | 0.00 | 1.00 | 5.75 | 1.50 | 4.75 | k__Bacteria; p__Firmicutes; c__Clostridia; o__Clostridiales; f__Eubacteriaceae; g__Eubacterium; s__Eubacterium sp. AB3007 |
| ID39486 | 14.228667 | 0.027184 | 0.67 | 0.50 | 0.00 | 3.00 | 54.75 | 16.75 | 122.25 | k__Bacteria; p__Firmicutes; c__Clostridia; o__Clostridiales; f__Lachnospiraceae; g__Dorea; s__Dorea formicigenerans |
| ID84107 | 14.217040 | 0.027303 | 1.67 | 1.00 | 1.00 | 1.00 | 6.75 | 73.50 | 18.00 | k__Bacteria; p__Actinobacteria <phylum>; c__Coriobacteriia; o__Coriobacteriales; f__Coriobacteriaceae; g__(Coriobacteriaceae); s__(Coriobacteriaceae) |
| ID976 | 14.203248 | 0.027446 | 474.00 | 10.00 | 2.00 | 18.00 | 162.25 | 116.25 | 31.00 | k__Bacteria; p__Bacteroidetes; c__(Bacteroidetes); o__(Bacteroidetes); f__(Bacteroidetes); g__(Bacteroidetes); s__(Bacteroidetes) |
| ID310514 | 14.201591 | 0.027464 | 0.33 | 0.00 | 0.00 | 1.00 | 61.00 | 108.75 | 4.25 | k__Bacteria; p__Bacteroidetes; c__Bacteroidia; o__Bacteroidales; f__Prevotellaceae; g__Prevotella; s__Prevotella multisaccharivorax |
| ID1268 | 14.190882 | 0.027575 | 199.33 | 1.00 | 1.00 | 2.00 | 0.75 | 1.50 | 1.00 | k__Bacteria; p__Actinobacteria <phylum>; c__Actinobacteria; o__Micrococcales; f__Micrococcaceae; g__(Micrococcaceae); s__(Micrococcaceae) |
| ID57493 | 14.176166 | 0.027729 | 9.33 | 0.50 | 0.00 | 1.00 | 0.00 | 0.00 | 0.25 | k__Bacteria; p__Actinobacteria <phylum>; c__Actinobacteria; o__Micrococcales; f__Micrococcaceae; g__Kocuria; s__(Kocuria) |
| ID180164 | 14.116119 | 0.028366 | 0.67 | 0.50 | 0.00 | 2.00 | 24.25 | 10.00 | 60.00 | k__Bacteria; p__Firmicutes; c__Clostridia; o__Clostridiales; f__Lachnospiraceae; g__Blautia; s__Blautia schinkii |
| ID1232453 | 14.114929 | 0.028378 | 0.33 | 0.00 | 0.00 | 1.00 | 23.25 | 3.75 | 21.25 | k__Bacteria; p__Firmicutes; c__Clostridia; o__Clostridiales; f__(Clostridiales bacterium VE202-21); g__(Clostridiales bacterium VE202-21); s__Clostridiales bacterium VE202-21 |
| ID2759 | 14.109375 | 0.028438 | 8.00 | 24.50 | 19.00 | 3.00 | 5.00 | 6.50 | 1.00 | k__Eukaryota; p__(Eukaryota); c__(Eukaryota); o__(Eukaryota); f__(Eukaryota); g__(Eukaryota); s__(Eukaryota) |
| ID39488 | 14.091090 | 0.028635 | 2.33 | 0.50 | 0.00 | 4.00 | 67.75 | 44.00 | 314.25 | k__Bacteria; p__Firmicutes; c__Clostridia; o__Clostridiales; f__Eubacteriaceae; g__Eubacterium; s__[Eubacterium] hallii |
| ID28130 | 14.088894 | 0.028659 | 0.33 | 0.00 | 0.00 | 1.00 | 46.50 | 7.50 | 4.00 | k__Bacteria; p__Bacteroidetes; c__Bacteroidia; o__Bacteroidales; f__Prevotellaceae; g__Prevotella; s__Prevotella disiens |
| ID31971 | 14.085626 | 0.028694 | 0.33 | 0.00 | 0.00 | 1.00 | 6.25 | 1.00 | 13.50 | k__Bacteria; p__Firmicutes; c__Erysipelotrichia; o__Erysipelotrichales; f__Erysipelotrichaceae; g__([Eubacterium] dolichum); s__[Eubacterium] dolichum |
| ID1462527 | 14.080000 | 0.028755 | 11.33 | 1.00 | 0.00 | 1.00 | 0.25 | 0.00 | 0.25 | k__Bacteria; p__Firmicutes; c__Bacilli; o__Bacillales; f__Bacillaceae; g__Oceanobacillus; s__Oceanobacillus jeddahense |
| ID1491 | 14.054084 | 0.029038 | 2.33 | 207.50 | 76.00 | 267.00 | 10.25 | 5.50 | 7.50 | k__Bacteria; p__Firmicutes; c__Clostridia; o__Clostridiales; f__Clostridiaceae; g__Clostridium; s__Clostridium botulinum |
| ID1235792 | 14.038104 | 0.029213 | 0.67 | 0.00 | 0.00 | 1.00 | 6.25 | 1.50 | 9.50 | k__Bacteria; p__Firmicutes; c__Clostridia; o__Clostridiales; f__Lachnospiraceae; g__(Lachnospiraceae bacterium M18-1); s__Lachnospiraceae bacterium M18-1 |
| ID328814 | 14.011585 | 0.029507 | 0.67 | 0.00 | 0.00 | 1.00 | 13.50 | 1.75 | 30.50 | k__Bacteria; p__Bacteroidetes; c__Bacteroidia; o__Bacteroidales; f__Rikenellaceae; g__Alistipes; s__Alistipes shahii |
| ID1602169 | 14.004848 | 0.029582 | 0.00 | 0.00 | 0.00 | 1.00 | 53.00 | 23.25 | 4.00 | k__Bacteria; p__Bacteroidetes; c__Bacteroidia; o__Bacteroidales; f__Prevotellaceae; g__Prevotella; s__Prevotella sp. P4-76 |
| ID29361 | 13.994181 | 0.029701 | 1.33 | 0.00 | 0.00 | 2.00 | 16.00 | 4.00 | 36.75 | k__Bacteria; p__Firmicutes; c__Clostridia; o__Clostridiales; f__Lachnospiraceae; g__Tyzzerella; s__Tyzzerella nexilis |
| ID33039 | 13.988420 | 0.029766 | 0.33 | 1.00 | 0.00 | 5.00 | 11.50 | 4.00 | 151.50 | k__Bacteria; p__Firmicutes; c__Clostridia; o__Clostridiales; f__Lachnospiraceae; g__Blautia; s__[Ruminococcus] torques |
| ID2159 | 13.981967 | 0.029838 | 12.67 | 1.00 | 1.00 | 1.00 | 1.25 | 0.25 | 1.25 | k__Archaea; p__Euryarchaeota; c__Methanobacteria; o__Methanobacteriales; f__Methanobacteriaceae; g__(Methanobacteriaceae); s__(Methanobacteriaceae) |
| ID658655 | 13.941095 | 0.030301 | 0.33 | 0.50 | 0.00 | 1.00 | 8.25 | 2.00 | 18.25 | k__Bacteria; p__Firmicutes; c__Clostridia; o__Clostridiales; f__Lachnospiraceae; g__(Lachnospiraceae bacterium 1_4_56FAA); s__Lachnospiraceae bacterium 1_4_56FAA |
| ID32008 | 13.935886 | 0.030361 | 6.33 | 8.50 | 11.00 | 3.00 | 0.75 | 1.00 | 0.75 | k__Bacteria; p__Proteobacteria; c__Betaproteobacteria; o__Burkholderiales; f__Burkholderiaceae; g__Burkholderia; s__(Burkholderia) |
| ID86661 | 13.887857 | 0.030914 | 13.67 | 4.00 | 1.00 | 6.00 | 0.75 | 1.25 | 1.25 | k__Bacteria; p__Firmicutes; c__Bacilli; o__Bacillales; f__Bacillaceae; g__Bacillus; s__(Bacillus) |
| ID292800 | 13.853864 | 0.031312 | 0.33 | 1.00 | 0.00 | 5.00 | 32.25 | 5.75 | 28.25 | k__Bacteria; p__Firmicutes; c__Clostridia; o__Clostridiales; f__(Flavonifractor); g__Flavonifractor; s__Flavonifractor plautii |
| ID674529 | 13.825269 | 0.031650 | 0.33 | 0.00 | 0.00 | 1.00 | 234.25 | 116.50 | 55.50 | k__Bacteria; p__Bacteroidetes; c__Bacteroidia; o__Bacteroidales; f__Bacteroidaceae; g__Bacteroides; s__Bacteroides faecis |
| ID410072 | 13.819794 | 0.031715 | 0.33 | 1.00 | 0.00 | 3.00 | 52.00 | 13.25 | 296.75 | k__Bacteria; p__Firmicutes; c__Clostridia; o__Clostridiales; f__Lachnospiraceae; g__Coprococcus; s__Coprococcus comes |
| ID457412 | 13.803324 | 0.031912 | 0.33 | 0.00 | 0.00 | 1.00 | 35.00 | 12.75 | 170.75 | k__Bacteria; p__Firmicutes; c__Clostridia; o__Clostridiales; f__Ruminococcaceae; g__Ruminococcus; s__Ruminococcus sp. 5_1_39BFAA |
| ID85015 | 13.777778 | 0.032219 | 32.67 | 1.00 | 0.00 | 1.00 | 0.75 | 0.00 | 0.50 | k__Bacteria; p__Actinobacteria <phylum>; c__Actinobacteria; o__Propionibacteriales; f__Nocardioidaceae; g__(Nocardioidaceae); s__(Nocardioidaceae) |
| ID484508 | 13.777778 | 0.032219 | 24.00 | 1.00 | 0.00 | 1.00 | 0.75 | 0.00 | 0.50 | k__Bacteria; p__Firmicutes; c__Bacilli; o__Bacillales; f__Bacillaceae; g__Ornithinibacillus; s__(Ornithinibacillus) |
| ID84406 | 13.777778 | 0.032219 | 98.67 | 1.00 | 0.00 | 1.00 | 0.75 | 0.00 | 0.50 | k__Bacteria; p__Firmicutes; c__Bacilli; o__Bacillales; f__Bacillaceae; g__Virgibacillus; s__(Virgibacillus) |
| ID1284775 | 13.766841 | 0.032352 | 0.00 | 0.00 | 0.00 | 1.00 | 31.25 | 7.25 | 1.50 | k__Bacteria; p__Bacteroidetes; c__Bacteroidia; o__Bacteroidales; f__Prevotellaceae; g__Prevotella; s__Prevotella sp. S7-1-8 |
| ID186650 | 13.757009 | 0.032471 | 5.00 | 1.00 | 1.00 | 1.00 | 0.50 | 0.00 | 0.25 | k__Bacteria; p__Proteobacteria; c__Alphaproteobacteria; o__Rhizobiales; f__Methylobacteriaceae; g__Microvirga; s__(Microvirga) |
| ID1760 | 13.756804 | 0.032474 | 451.00 | 14.50 | 4.00 | 26.00 | 11.75 | 51.25 | 14.00 | k__Bacteria; p__Actinobacteria <phylum>; c__Actinobacteria; o__(Actinobacteria); f__(Actinobacteria); g__(Actinobacteria); s__(Actinobacteria) |
| ID283 | 13.742739 | 0.032646 | 42.00 | 1.00 | 1.00 | 1.00 | 0.75 | 0.00 | 0.50 | k__Bacteria; p__Proteobacteria; c__Betaproteobacteria; o__Burkholderiales; f__Comamonadaceae; g__Comamonas; s__(Comamonas) |
| ID2425 | 13.742739 | 0.032646 | 11.33 | 1.00 | 1.00 | 1.00 | 0.75 | 0.00 | 0.50 | k__Bacteria; p__Proteobacteria; c__Gammaproteobacteria; o__Cellvibrionales; f__Cellvibrionaceae; g__Teredinibacter; s__(Teredinibacter) |
| ID68 | 13.742739 | 0.032646 | 63.00 | 1.00 | 1.00 | 1.00 | 0.75 | 0.00 | 0.50 | k__Bacteria; p__Proteobacteria; c__Gammaproteobacteria; o__Xanthomonadales; f__Xanthomonadaceae; g__Lysobacter; s__(Lysobacter) |
| ID40323 | 13.742739 | 0.032646 | 38.00 | 1.00 | 1.00 | 1.00 | 0.75 | 0.00 | 0.50 | k__Bacteria; p__Proteobacteria; c__Gammaproteobacteria; o__Xanthomonadales; f__Xanthomonadaceae; g__Stenotrophomonas; s__(Stenotrophomonas) |
| ID1817 | 13.742739 | 0.032646 | 21.33 | 1.00 | 1.00 | 1.00 | 0.75 | 0.00 | 0.50 | k__Bacteria; p__Actinobacteria <phylum>; c__Actinobacteria; o__Corynebacteriales; f__Nocardiaceae; g__Nocardia; s__(Nocardia) |
| ID5042 | 13.742739 | 0.032646 | 42.33 | 1.00 | 1.00 | 1.00 | 0.50 | 0.00 | 0.25 | k__Eukaryota; p__Ascomycota; c__Eurotiomycetes; o__Eurotiales; f__(Eurotiales); g__(Eurotiales); s__(Eurotiales) |
| ID1131492 | 13.742739 | 0.032646 | 74.67 | 1.00 | 1.00 | 1.00 | 0.75 | 0.00 | 0.50 | k__Eukaryota; p__Ascomycota; c__Eurotiomycetes; o__Eurotiales; f__Aspergillaceae; g__(Aspergillaceae); s__(Aspergillaceae) |
| ID1378168 | 13.728905 | 0.032815 | 1.67 | 0.00 | 0.00 | 1.00 | 29.50 | 6.50 | 20.25 | k__Bacteria; p__Firmicutes; c__(Firmicutes bacterium ASF500); o__(Firmicutes bacterium ASF500); f__(Firmicutes bacterium ASF500); g__(Firmicutes bacterium ASF500); s__Firmicutes bacterium |
| ID1730 | 13.702446 | 0.033143 | 2.33 | 1.00 | 1.00 | 1.00 | 12.00 | 5.75 | 15.50 | k__Bacteria; p__Firmicutes; c__Clostridia; o__Clostridiales; f__Eubacteriaceae; g__Eubacterium; s__(Eubacterium) |

| | | | | | | | | | | |
|---|---|---|---|---|---|---|---|---|---|---|
| ID1232439 | 13.693571 | 0.033253 | 0.33 | 0.00 | 0.00 | 1.00 | 9.25 | 2.00 | 8.25 | k__Bacteria; p__Firmicutes; c__Clostridia; o__Clostridiales; f__(Clostridiales bacterium VE202-03); g__(Clostridiales bacterium VE202-03); s__Clostridiales bacterium VE202-03 |
| ID171552 | 13.663130 | 0.033634 | 3.33 | 1.00 | 1.00 | 2.00 | 81.75 | 42.75 | 10.50 | k__Bacteria; p__Bacteroidetes; c__Bacteroidia; o__Bacteroidales; f__Prevotellaceae; g__(Prevotellaceae); s__(Prevotellaceae) |
| ID715989 | 13.636095 | 0.033976 | 4.00 | 3.50 | 14.00 | 1.00 | 0.75 | 1.00 | 0.75 | k__Eukaryota; p__Ascomycota; c__(Ascomycota); o__(Ascomycota); f__(Ascomycota); g__(Ascomycota); s__(Ascomycota) |
| ID1385985 | 13.634043 | 0.034002 | 102.33 | 0.00 | 0.00 | 1.00 | 0.75 | 0.00 | 0.50 | k__Bacteria; p__Bacteroidetes; c__Sphingobacteriia; o__Sphingobacteriales; f__Sphingobacteriaceae; g__Sphingobacterium; s__Sphingobacterium paucimobilis |
| ID1839 | 13.634043 | 0.034002 | 44.00 | 1.00 | 0.00 | 1.00 | 0.25 | 0.00 | 0.50 | k__Bacteria; p__Actinobacteria <phylum>; c__Actinobacteria; o__Propionibacteriales; f__Nocardioidaceae; g__Nocardioides; s__(Nocardioides) |
| ID1506553 | 13.601762 | 0.034415 | 5.00 | 1.00 | 1.00 | 5.00 | 38.25 | 28.00 | 83.75 | k__Bacteria; p__Firmicutes; c__Clostridia; o__Clostridiales; f__Lachnospiraceae; g__Lachnoclostridium; s__(Lachnoclostridium) |
| ID301301 | 13.547003 | 0.035127 | 0.33 | 1.00 | 0.00 | 11.00 | 91.00 | 10.00 | 81.75 | k__Bacteria; p__Firmicutes; c__Clostridia; o__Clostridiales; f__Lachnospiraceae; g__Roseburia; s__Roseburia hominis |
| ID78346 | 13.539566 | 0.035224 | 0.00 | 0.00 | 0.00 | 0.00 | 0.00 | 38.50 | 4.25 | k__Bacteria; p__Actinobacteria <phylum>; c__Actinobacteria; o__Bifidobacteriales; f__Bifidobacteriaceae; g__Bifidobacterium; s__Bifidobacterium ruminantium |
| ID35832 | 13.535340 | 0.035280 | 0.33 | 0.00 | 0.00 | 1.00 | 2.25 | 0.00 | 63.25 | k__Bacteria; p__Proteobacteria; c__Deltaproteobacteria; o__Desulfovibrionales; f__Desulfovibrionaceae; g__Bilophila; s__(Bilophila) |
| ID817 | 13.522467 | 0.035450 | 4.67 | 1.00 | 1.00 | 5.00 | 173.00 | 134.25 | 276.25 | k__Bacteria; p__Bacteroidetes; c__Bacteroidia; o__Bacteroidales; f__Bacteroidaceae; g__Bacteroides; s__Bacteroides fragilis |
| ID131567 | 13.500659 | 0.035740 | 95.33 | 136.00 | 91.00 | 77.00 | 48.25 | 55.00 | 24.75 | k__(null); p__(null); c__(null); o__(null); f__(null); g__(null); s__(null) |
| ID265975 | 13.494667 | 0.035820 | 4.00 | 1.00 | 0.00 | 1.00 | 28.00 | 8.75 | 7.25 | k__Bacteria; p__Firmicutes; c__Clostridia; o__Clostridiales; f__Lachnospiraceae; g__Oribacterium; s__(Oribacterium) |
| ID729 | 13.467273 | 0.036187 | 0.00 | 1.00 | 1.00 | 36.00 | 6.50 | 47.00 | 1.50 | k__Bacteria; p__Proteobacteria; c__Gammaproteobacteria; o__Pasteurellales; f__Pasteurellaceae; g__Haemophilus; s__Haemophilus parainfluenzae |
| ID28127 | 13.463665 | 0.036236 | 0.33 | 0.00 | 0.00 | 1.00 | 45.50 | 20.25 | 3.75 | k__Bacteria; p__Bacteroidetes; c__Bacteroidia; o__Bacteroidales; f__Prevotellaceae; g__Prevotella; s__Prevotella buccalis |
| ID202611 | 13.447471 | 0.036456 | 106.00 | 0.00 | 0.00 | 1.00 | 0.25 | 0.00 | 0.25 | k__Bacteria; p__Firmicutes; c__Bacilli; o__Bacillales; f__Bacillaceae; g__Lysinibacillus; s__Lysinibacillus odysseyi |
| ID80840 | 13.444493 | 0.036496 | 73.33 | 32.50 | 28.00 | 32.00 | 7.00 | 8.75 | 7.25 | k__Bacteria; p__Proteobacteria; c__Betaproteobacteria; o__Burkholderiales; f__(Burkholderiales); g__(Burkholderiales); s__(Burkholderiales) |
| ID1235797 | 13.443649 | 0.036507 | 2.67 | 0.50 | 0.00 | 4.00 | 68.50 | 16.25 | 30.75 | k__Bacteria; p__Firmicutes; c__Clostridia; o__Clostridiales; f__Oscillospiraceae; g__Oscillibacter; s__Oscillibacter sp. 1-3 |
| ID742723 | 13.442251 | 0.036526 | 0.67 | 0.00 | 0.00 | 1.00 | 2.50 | 0.00 | 4.50 | k__Bacteria; p__Firmicutes; c__Clostridia; o__Clostridiales; f__Lachnospiraceae; g__(Lachnospiraceae bacterium 2_1_46FAA); s__Lachnospiraceae bacterium 2_1_46FAA |
| ID1338009 | 13.440000 | 0.036557 | 18.67 | 0.00 | 0.00 | 0.00 | 0.75 | 0.00 | 0.50 | k__Bacteria; p__Bacteroidetes; c__Sphingobacteriia; o__Sphingobacteriales; f__Sphingobacteriaceae; g__Sphingobacterium; s__Sphingobacterium sp. IITKGP-BTPF85 |
| ID77133 | 13.427313 | 0.036730 | 13.67 | 7.00 | 30.00 | 4.00 | 41.75 | 8.75 | 17.50 | k__Bacteria; p__(uncultured bacterium); c__(uncultured bacterium); o__(uncultured bacterium); f__(uncultured bacterium); g__(uncultured bacterium); s__uncultured bacterium |
| ID208479 | 13.394595 | 0.037181 | 0.67 | 1.00 | 0.00 | 1.00 | 18.50 | 4.75 | 36.50 | k__Bacteria; p__Firmicutes; c__Clostridia; o__Clostridiales; f__Lachnospiraceae; g__Lachnoclostridium; s__[Clostridium] bolteae |
| ID2172 | 13.364754 | 0.037596 | 48.33 | 0.00 | 1.00 | 1.00 | 3.00 | 0.75 | 6.00 | k__Archaea; p__Euryarchaeota; c__Methanobacteria; o__Methanobacteriales; f__Methanobacteriaceae; g__Methanobrevibacter; s__(Methanobrevibacter) |
| ID906 | 13.362162 | 0.037632 | 1.67 | 1.00 | 1.00 | 1.00 | 9.75 | 224.50 | 31.75 | k__Bacteria; p__Firmicutes; c__Negativicutes; o__Veillonellales; f__Veillonellaceae; g__Megasphaera; s__(Megasphaera) |
| ID742722 | 13.353808 | 0.037749 | 0.33 | 0.00 | 0.00 | 1.00 | 4.75 | 18.50 | 49.50 | k__Bacteria; p__Actinobacteria <phylum>; c__Coriobacteriia; o__Coriobacteriales; f__Coriobacteriaceae; g__Collinsella; s__Collinsella sp. 4_8_47FAA |
| ID1165092 | 13.350546 | 0.037795 | 2.33 | 0.00 | 0.00 | 1.00 | 9.75 | 2.00 | 2.25 | k__Bacteria; p__Firmicutes; c__Clostridia; o__Clostridiales; f__Lachnospiraceae; g__(Lachnospiraceae bacterium JC7); s__Lachnospiraceae bacterium JC7 |
| ID28118 | 13.349462 | 0.037810 | 0.67 | 1.00 | 0.00 | 1.00 | 45.25 | 7.00 | 80.25 | k__Bacteria; p__Bacteroidetes; c__Bacteroidia; o__Bacteroidales; f__Odoribacteraceae; g__Odoribacter; s__Odoribacter splanchnicus |
| ID1095771 | 13.337950 | 0.037972 | 0.33 | 0.00 | 0.00 | 1.00 | 54.00 | 16.75 | 93.50 | k__Bacteria; p__Firmicutes; c__Clostridia; o__Clostridiales; f__Ruminococcaceae; g__Ruminococcus; s__Ruminococcus sp. JC304 |
| ID1507 | 13.330909 | 0.038072 | 0.67 | 1.00 | 0.00 | 1.00 | 14.00 | 2.50 | 31.75 | k__Bacteria; p__Firmicutes; c__Clostridia; o__Clostridiales; f__Clostridiaceae; g__Clostridium; s__Clostridium sp. ATCC 29733 |
| ID59732 | 13.322316 | 0.038194 | 71.67 | 101.50 | 2.00 | 74.00 | 4.00 | 3.00 | 1.00 | k__Bacteria; p__Bacteroidetes; c__Flavobacteriia; o__Flavobacteriales; f__Flavobacteriaceae; g__Chryseobacterium; s__(Chryseobacterium) |
| ID228604 | 13.299132 | 0.038524 | 0.33 | 0.00 | 0.00 | 1.00 | 18.25 | 8.25 | 2.75 | k__Bacteria; p__Bacteroidetes; c__Bacteroidia; o__Bacteroidales; f__Prevotellaceae; g__Prevotella; s__Prevotella salivae |
| ID189330 | 13.287611 | 0.038689 | 2.33 | 1.00 | 1.00 | 3.00 | 43.00 | 44.25 | 93.50 | k__Bacteria; p__Firmicutes; c__Clostridia; o__Clostridiales; f__Lachnospiraceae; g__Dorea; s__(Dorea) |
| ID154046 | 13.229443 | 0.039534 | 2.00 | 1.00 | 1.00 | 5.00 | 41.00 | 18.50 | 56.75 | k__Bacteria; p__Firmicutes; c__Clostridia; o__Clostridiales; f__Clostridiaceae; g__Hungatella; s__Hungatella hathewayi |
| ID106588 | 13.190941 | 0.040102 | 2.33 | 2.00 | 0.00 | 12.00 | 47.50 | 11.25 | 40.50 | k__Bacteria; p__Firmicutes; c__Clostridia; o__Clostridiales; f__(Pseudoflavonifractor); g__Pseudoflavonifractor; s__Pseudoflavonifractor capillosus |
| ID165697 | 13.185185 | 0.040188 | 7.00 | 1.00 | 1.00 | 1.00 | 0.50 | 0.00 | 0.50 | k__Bacteria; p__Proteobacteria; c__Alphaproteobacteria; o__Sphingomonadales; f__Sphingomonadaceae; g__Sphingopyxis; s__(Sphingopyxis) |
| ID1243664 | 13.185185 | 0.040188 | 15.67 | 0.50 | 0.00 | 1.00 | 0.75 | 0.00 | 0.75 | k__Bacteria; p__Firmicutes; c__Bacilli; o__Bacillales; f__Bacillaceae; g__Bacillus; s__Bacillus massiliogorillae |
| ID1519438 | 13.184651 | 0.040196 | 0.67 | 0.00 | 0.00 | 1.00 | 5.00 | 1.50 | 10.75 | k__Bacteria; p__Firmicutes; c__Clostridia; o__Clostridiales; f__Eubacteriaceae; g__Eubacterium; s__Eubacterium sp. ER2 |
| ID541000 | 13.183759 | 0.040209 | 8.67 | 3.50 | 1.00 | 8.00 | 92.25 | 55.50 | 337.50 | k__Bacteria; p__Firmicutes; c__Clostridia; o__Clostridiales; f__Ruminococcaceae; g__(Ruminococcaceae); s__(Ruminococcaceae) |
| ID43305 | 13.177207 | 0.040307 | 1.67 | 0.00 | 0.00 | 1.00 | 10.00 | 2.00 | 5.25 | k__Bacteria; p__Firmicutes; c__Clostridia; o__Clostridiales; f__Lachnospiraceae; g__Butyrivibrio; s__Butyrivibrio proteoclasticus |
| ID545501 | 13.145228 | 0.040787 | 11.33 | 1.00 | 0.00 | 1.00 | 0.50 | 0.00 | 0.50 | k__Bacteria; p__Firmicutes; c__Bacilli; o__Bacillales; f__Bacillaceae; g__Oceanobacillus; s__Oceanobacillus oncorhynchi |
| ID351094 | 13.145228 | 0.040787 | 49.33 | 1.00 | 0.00 | 1.00 | 0.50 | 0.00 | 0.50 | k__Bacteria; p__Firmicutes; c__Bacilli; o__Bacillales; f__Bacillaceae; g__Paucisalibacillus; s__(Paucisalibacillus) |
| ID28135 | 13.130822 | 0.041005 | 0.33 | 0.00 | 0.00 | 1.00 | 39.75 | 13.75 | 2.75 | k__Bacteria; p__Bacteroidetes; c__Bacteroidia; o__Bacteroidales; f__Prevotellaceae; g__Prevotella; s__Prevotella oris |
| ID1226322 | 13.129269 | 0.041028 | 0.33 | 0.00 | 0.00 | 1.00 | 30.25 | 6.75 | 28.00 | k__Bacteria; p__Firmicutes; c__Clostridia; o__Clostridiales; f__Oscillospiraceae; g__Oscillibacter; s__Oscillibacter sp. KLE 1728 |
| ID386414 | 13.129018 | 0.041032 | 0.33 | 1.00 | 0.00 | 12.00 | 84.75 | 36.25 | 5.50 | k__Bacteria; p__Bacteroidetes; c__Bacteroidia; o__Bacteroidales; f__Prevotellaceae; g__Prevotella; s__Prevotella timonensis |
| ID1263 | 13.123677 | 0.041113 | 7.33 | 1.00 | 1.00 | 7.00 | 75.00 | 24.75 | 90.00 | k__Bacteria; p__Firmicutes; c__Clostridia; o__Clostridiales; f__Ruminococcaceae; g__Ruminococcus; s__(Ruminococcus) |
| ID589537 | 13.105338 | 0.041393 | 0.00 | 0.00 | 0.00 | 1.00 | 5.25 | 3.00 | 1.00 | k__Bacteria; p__Bacteroidetes; c__Bacteroidia; o__Bacteroidales; f__Prevotellaceae; g__Prevotella; s__Prevotella dentasini |
| ID82373 | 13.094595 | 0.041558 | 0.67 | 1.00 | 0.00 | 2.00 | 31.50 | 1.00 | 1.00 | k__Bacteria; p__Firmicutes; c__Negativicutes; o__Selenomonadales; f__Selenomonadaceae; g__Anaerovibrio; s__(Anaerovibrio) |
| ID633701 | 13.077011 | 0.041829 | 0.33 | 0.00 | 0.00 | 1.00 | 26.50 | 10.50 | 2.50 | k__Bacteria; p__Bacteroidetes; c__Bacteroidia; o__Bacteroidales; f__Prevotellaceae; g__Prevotella; s__Prevotella saccharolytica |
| ID84566 | 13.069547 | 0.041945 | 273.00 | 1.00 | 1.00 | 2.00 | 6.50 | 3.75 | 1.00 | k__Bacteria; p__Bacteroidetes; c__Sphingobacteriia; o__Sphingobacteriales; f__Sphingobacteriaceae; g__(Sphingobacteriaceae); s__(Sphingobacteriaceae) |
| ID97139 | 13.068887 | 0.041955 | 0.67 | 0.00 | 0.00 | 1.00 | 8.50 | 2.75 | 8.75 | k__Bacteria; p__Firmicutes; c__Clostridia; o__Clostridiales; f__Clostridiaceae; g__Clostridium; s__Clostridium sp. ASF502 |
| ID85006 | 13.048358 | 0.042274 | 162.67 | 2.50 | 1.00 | 9.00 | 2.75 | 3.00 | 1.50 | k__Bacteria; p__Actinobacteria <phylum>; c__Actinobacteria; o__Micrococcales; f__(Micrococcales); g__(Micrococcales); s__(Micrococcales) |
| ID39483 | 13.041667 | 0.042379 | 0.33 | 0.00 | 0.00 | 1.00 | 4.50 | 1.75 | 7.75 | k__Bacteria; p__Firmicutes; c__Erysipelotrichia; o__Erysipelotrichales; f__Erysipelotrichaceae; g__Faecalitalea; s__Faecalitalea cylindroides |
| ID46506 | 13.035585 | 0.042474 | 0.33 | 0.00 | 0.00 | 1.00 | 17.75 | 7.50 | 133.25 | k__Bacteria; p__Bacteroidetes; c__Bacteroidia; o__Bacteroidales; f__Bacteroidaceae; g__Bacteroides; s__Bacteroides stercoris |
| ID1161942 | 13.017560 | 0.042758 | 0.33 | 0.00 | 0.00 | 1.00 | 200.75 | 9.75 | 20.75 | k__Bacteria; p__Firmicutes; c__Clostridia; o__Clostridiales; f__Ruminococcaceae; g__Ruminococcus; s__Ruminococcus champanellensis |
| ID1283313 | 13.011711 | 0.042850 | 1.67 | 1.00 | 1.00 | 1.00 | 31.50 | 17.25 | 2.50 | k__Bacteria; p__Bacteroidetes; c__Bacteroidia; o__Bacteroidales; f__Prevotellaceae; g__Alloprevotella; s__(Alloprevotella) |
| ID407975 | 13.007534 | 0.042916 | 0.33 | 0.00 | 0.00 | 1.00 | 16.75 | 7.25 | 2.25 | k__Bacteria; p__Bacteroidetes; c__Bacteroidia; o__Bacteroidales; f__Prevotellaceae; g__Prevotella; s__Prevotella pleuritidis |
| ID652721 | 12.991196 | 0.043176 | 0.00 | 0.00 | 0.00 | 1.00 | 9.25 | 9.75 | 2.25 | k__Bacteria; p__Bacteroidetes; c__Bacteroidia; o__Bacteroidales; f__Prevotellaceae; g__Prevotella; s__Prevotella sp. oral taxon 317 |
| ID200643 | 12.979428 | 0.043364 | 8.33 | 1.00 | 1.00 | 1.00 | 30.75 | 13.25 | 5.75 | k__Bacteria; p__Bacteroidetes; c__Bacteroidia; o__(Bacteroidia); f__(Bacteroidia); g__(Bacteroidia); s__(Bacteroidia) |
| ID61170 | 12.963985 | 0.043611 | 0.33 | 0.50 | 0.00 | 1.00 | 3.25 | 1.00 | 8.25 | k__Bacteria; p__Firmicutes; c__Erysipelotrichia; o__Erysipelotrichales; f__Erysipelotrichaceae; g__Holdemania; s__(Holdemania) |
| ID31953 | 12.932609 | 0.044119 | 1.00 | 1.00 | 1.00 | 1.00 | 2.25 | 86.50 | 27.25 | k__Bacteria; p__Actinobacteria <phylum>; c__Actinobacteria; o__Bifidobacteriales; f__Bifidobacteriaceae; g__(Bifidobacteriaceae); s__(Bifidobacteriaceae) |
| ID658087 | 12.914468 | 0.044414 | 0.33 | 0.50 | 0.00 | 1.00 | 22.25 | 6.00 | 32.25 | k__Bacteria; p__Firmicutes; c__Clostridia; o__Clostridiales; f__Lachnospiraceae; g__(Lachnospiraceae bacterium 7_1_58FAA); s__Lachnospiraceae bacterium 7_1_58FAA |
| ID2173 | 12.885000 | 0.044899 | 22.33 | 0.00 | 0.00 | 1.00 | 37.75 | 4.75 | 89.25 | k__Archaea; p__Euryarchaeota; c__Methanobacteria; o__Methanobacteriales; f__Methanobacteriaceae; g__Methanobrevibacter; s__Methanobrevibacter smithii |
| ID28211 | 12.882586 | 0.044939 | 107.00 | 13.00 | 9.00 | 10.00 | 15.00 | 9.00 | 2.25 | k__Bacteria; p__Proteobacteria; c__Alphaproteobacteria; o__(Alphaproteobacteria); f__(Alphaproteobacteria); g__(Alphaproteobacteria); s__(Alphaproteobacteria) |
| ID1263547 | 12.858409 | 0.045340 | 1.67 | 0.00 | 0.00 | 1.00 | 30.00 | 6.75 | 14.00 | k__Bacteria; p__Firmicutes; c__Clostridia; o__Clostridiales; f__Oscillospiraceae; g__Oscillibacter; s__Oscillibacter ruminantium |
| ID1735 | 12.842391 | 0.045608 | 0.33 | 0.50 | 0.00 | 2.00 | 135.50 | 21.50 | 8.25 | k__Bacteria; p__Firmicutes; c__Erysipelotrichia; o__Erysipelotrichales; f__Erysipelotrichaceae; g__Holdemanella; s__Holdemanella biformis |
| ID43668 | 12.842105 | 0.045613 | 20.67 | 0.00 | 0.00 | 0.00 | 0.25 | 0.00 | 0.25 | k__Bacteria; p__Actinobacteria <phylum>; c__Actinobacteria; o__Micrococcales; f__Dermabacteraceae; g__Brachybacterium; s__(Brachybacterium) |
| ID1396 | 12.841263 | 0.045627 | 13.33 | 4.50 | 1.00 | 9.00 | 1.50 | 2.25 | 1.25 | k__Bacteria; p__Firmicutes; c__Bacilli; o__Bacillales; f__Bacillaceae; g__Bacillus; s__Bacillus cereus |
| ID1392836 | 12.834622 | 0.045739 | 2.00 | 0.00 | 0.00 | 1.00 | 10.00 | 2.50 | 8.00 | k__Bacteria; p__Firmicutes; c__Clostridia; o__Clostridiales; f__Lachnospiraceae; g__(Lachnospiraceae bacterium TWA4); s__Lachnospiraceae bacterium TWA4 |

| | | | | | | | | | | |
|---|---|---|---|---|---|---|---|---|---|---|
| ID214851 | 12.834441 | 0.045742 | 1.67 | 0.00 | 0.00 | 1.00 | 93.50 | 50.00 | 466.50 | k__Bacteria; p__Firmicutes; c__Clostridia; o__Clostridiales; f__Ruminococcaceae; g__Subdoligranulum; s__Subdoligranulum variabile |
| ID39491 | 12.827679 | 0.045856 | 0.67 | 1.00 | 0.00 | 15.00 | 1111.75 | 94.75 | 327.75 | k__Bacteria; p__Firmicutes; c__Clostridia; o__Clostridiales; f__Lachnospiraceae; g__([Eubacterium] rectale); s__[Eubacterium] rectale |
| ID552398 | 12.822955 | 0.045935 | 2.33 | 0.00 | 0.00 | 1.00 | 30.00 | 7.50 | 48.00 | k__Bacteria; p__Firmicutes; c__Clostridia; o__Clostridiales; f__Ruminococcaceae; g__(Ruminococcaceae bacterium D16); s__Ruminococcaceae bacterium D16 |
| ID563031 | 12.817241 | 0.046032 | 0.00 | 0.00 | 0.00 | 1.00 | 11.00 | 2.25 | 1.00 | k__Bacteria; p__Bacteroidetes; c__Bacteroidia; o__Bacteroidales; f__Prevotellaceae; g__Prevotella; s__Prevotella sp. C561 |
| ID310297 | 12.816059 | 0.046052 | 2.00 | 0.00 | 0.00 | 1.00 | 26.00 | 12.25 | 420.75 | k__Bacteria; p__Bacteroidetes; c__Bacteroidia; o__Bacteroidales; f__Bacteroidaceae; g__Bacteroides; s__Bacteroides plebeius |
| ID393087 | 12.814238 | 0.046083 | 11.67 | 0.00 | 0.00 | 1.00 | 0.50 | 0.00 | 0.50 | k__Bacteria; p__Firmicutes; c__Bacilli; o__Bacillales; f__Bacillaceae; g__Gracilibacillus; s__Gracilibacillus lacisalsi |
| ID671218 | 12.812212 | 0.046117 | 0.33 | 0.00 | 0.00 | 1.00 | 45.00 | 17.00 | 2.50 | k__Bacteria; p__Bacteroidetes; c__Bacteroidia; o__Bacteroidales; f__Prevotellaceae; g__Alloprevotella; s__Alloprevotella rava |
| ID712469 | 12.806391 | 0.046216 | 0.33 | 0.00 | 1.00 | 1.00 | 32.25 | 5.25 | 2.25 | k__Bacteria; p__Bacteroidetes; c__Bacteroidia; o__Bacteroidales; f__Prevotellaceae; g__Alloprevotella; s__Prevotella sp. oral taxon 473 |
| ID186826 | 12.803144 | 0.046271 | 12.00 | 3.50 | 4.00 | 10.00 | 12.00 | 44.75 | 8.25 | k__Bacteria; p__Firmicutes; c__Bacilli; o__Lactobacillales; f__(Lactobacillales); g__(Lactobacillales); s__(Lactobacillales) |
| ID419005 | 12.773066 | 0.046785 | 0.33 | 0.00 | 0.00 | 1.00 | 12.50 | 3.00 | 1.50 | k__Bacteria; p__Bacteroidetes; c__Bacteroidia; o__Bacteroidales; f__Prevotellaceae; g__Prevotella; s__Prevotella amnii |
| ID1385 | 12.766476 | 0.046898 | 787.00 | 39.50 | 11.00 | 43.00 | 23.75 | 21.25 | 13.50 | k__Bacteria; p__Firmicutes; c__Bacilli; o__Bacillales; f__(Bacillales); g__(Bacillales); s__(Bacillales) |
| ID128827 | 12.738757 | 0.047377 | 10.33 | 1.00 | 1.00 | 3.00 | 48.25 | 27.75 | 33.25 | k__Bacteria; p__Firmicutes; c__Erysipelotrichia; o__Erysipelotrichales; f__Erysipelotrichaceae; g__(Erysipelotrichaceae); s__(Erysipelotrichaceae) |
| ID48479 | 12.736595 | 0.047415 | 11.67 | 1.00 | 1.00 | 1.00 | 29.25 | 17.25 | 4.50 | k__Bacteria; p__(Bacteria); c__(Bacteria); o__(Bacteria); f__(Bacteria); g__(Bacteria); s__(Bacteria) |
| ID438033 | 12.724485 | 0.047626 | 1.67 | 1.00 | 0.00 | 1.00 | 11.25 | 2.75 | 16.00 | k__Bacteria; p__Firmicutes; c__Clostridia; o__Clostridiales; f__Ruminococcaceae; g__Ruminococcus; s__Ruminococcus gauvreauii |
| ID665940 | 12.719704 | 0.047709 | 0.33 | 0.00 | 0.00 | 1.00 | 1.50 | 0.75 | 11.50 | k__Bacteria; p__Firmicutes; c__Clostridia; o__Clostridiales; f__Clostridiaceae; g__Clostridium; s__Clostridium sp. 7_3_54FAA |
| ID397287 | 12.694598 | 0.048150 | 0.33 | 0.00 | 0.00 | 3.00 | 39.50 | 7.00 | 11.00 | k__Bacteria; p__Firmicutes; c__Clostridia; o__Clostridiales; f__Lachnospiraceae; g__(Lachnospiraceae bacterium 28-4); s__Lachnospiraceae bacterium 28-4 |
| ID28136 | 12.690259 | 0.048227 | 0.00 | 0.00 | 0.00 | 1.00 | 7.00 | 3.25 | 1.00 | k__Bacteria; p__Bacteroidetes; c__Bacteroidia; o__Bacteroidales; f__Prevotellaceae; g__Prevotella; s__Prevotella oulorum |
| ID162156 | 12.687773 | 0.048271 | 0.33 | 0.00 | 0.00 | 1.00 | 19.25 | 1.00 | 0.25 | k__Bacteria; p__Bacteroidetes; c__Bacteroidia; o__Bacteroidales; f__Bacteroidaceae; g__Bacteroides; s__uncultured Bacteroides sp. |
| ID2062 | 12.683938 | 0.048339 | 7.67 | 1.00 | 1.00 | 1.00 | 0.75 | 0.50 | 1.00 | k__Bacteria; p__Actinobacteria <phylum>; c__Actinobacteria; o__Streptomycetales; f__Streptomycetaceae; g__(Streptomycetaceae); s__(Streptomycetaceae) |
| ID301092 | 12.683938 | 0.048339 | 16.33 | 0.00 | 0.00 | 0.00 | 0.50 | 0.00 | 0.25 | k__Bacteria; p__Firmicutes; c__Bacilli; o__Bacillales; f__Bacillaceae; g__Marinococcus; s__Marinococcus halotolerans |
| ID1297617 | 12.682326 | 0.048367 | 0.67 | 0.00 | 0.00 | 1.00 | 23.50 | 5.75 | 25.00 | k__Bacteria; p__Firmicutes; c__Clostridia; o__Clostridiales; f__(Intestinimonas); g__Intestinimonas; s__Intestinimonas butyriciproducens |
| ID282402 | 12.681615 | 0.048380 | 0.00 | 0.00 | 0.00 | 1.00 | 7.50 | 1.75 | 0.75 | k__Bacteria; p__Bacteroidetes; c__Bacteroidia; o__Bacteroidales; f__Prevotellaceae; g__Prevotella; s__Prevotella multiformis |
| ID200644 | 12.666667 | 0.048646 | 44.33 | 1.00 | 1.00 | 1.00 | 2.25 | 0.75 | 1.00 | k__Bacteria; p__Bacteroidetes; c__Flavobacteriia; o__Flavobacteriales; f__(Flavobacteriales); g__(Flavobacteriales); s__(Flavobacteriales) |
| ID46206 | 12.659050 | 0.048781 | 1.00 | 1.00 | 0.00 | 1.00 | 9.00 | 1.50 | 6.00 | k__Bacteria; p__Firmicutes; c__Clostridia; o__Clostridiales; f__Lachnospiraceae; g__Pseudobutyrivibrio; s__Pseudobutyrivibrio ruminis |
| ID201174 | 12.651882 | 0.048909 | 6.33 | 1.00 | 1.00 | 1.00 | 4.75 | 45.50 | 7.00 | k__Bacteria; p__Actinobacteria <phylum>; c__(Actinobacteria <phylum>); o__(Actinobacteria <phylum>); f__(Actinobacteria <phylum>); g__(Actinobacteria <phylum>); s__(Actinobacteria <phylu |
| ID1400053 | 12.635748 | 0.049199 | 3.00 | 0.00 | 0.00 | 1.00 | 30.75 | 5.75 | 0.75 | k__Bacteria; p__Bacteroidetes; c__Bacteroidia; o__Bacteroidales; f__(Bacteroidales bacterium CF); g__(Bacteroidales bacterium CF); s__Bacteroidales bacterium CF |
| ID418240 | 12.617166 | 0.049535 | 0.33 | 0.50 | 0.00 | 1.00 | 80.50 | 20.25 | 324.25 | k__Bacteria; p__Firmicutes; c__Clostridia; o__Clostridiales; f__Lachnospiraceae; g__Blautia; s__Blautia wexlerae |
| ID1783272 | 12.610526 | 0.049655 | 87.33 | 19.00 | 8.00 | 34.00 | 159.50 | 207.75 | 167.00 | k__Bacteria; p__(Bacteria); c__(Bacteria); o__(Bacteria); f__(Bacteria); g__(Bacteria); s__(Bacteria) |
| ID84998 | 12.604796 | 0.049759 | 9.67 | 1.00 | 1.00 | 1.00 | 12.50 | 44.75 | 18.50 | k__Bacteria; p__Actinobacteria <phylum>; c__Coriobacteriia; o__(Coriobacteriia); f__(Coriobacteriia); g__(Coriobacteriia); s__(Coriobacteriia) |
| ID52227 | 12.596925 | 0.049903 | 0.33 | 0.00 | 0.00 | 1.00 | 28.00 | 7.25 | 1.25 | k__Bacteria; p__Bacteroidetes; c__Bacteroidia; o__Bacteroidales; f__Prevotellaceae; g__Prevotella; s__Prevotella dentalis |
| ID839 | 12.596575 | 0.049909 | 0.33 | 0.00 | 0.00 | 1.00 | 45.50 | 17.25 | 3.50 | k__Bacteria; p__Bacteroidetes; c__Bacteroidia; o__Bacteroidales; f__Prevotellaceae; g__Prevotella; s__Prevotella ruminicola |
| ID86331 | 12.592280 | 0.049987 | 19.67 | 1.00 | 20.00 | 2.00 | 7.00 | 3.25 | 3.25 | k__Bacteria; p__Firmicutes; c__Clostridia; o__Clostridiales; f__Clostridiales Family XIII. Incertae Sedis; g__Mogibacterium; s__(Mogibacterium) |

**Supplementary Table 5.** Information concerning DNA sequence read-length distribution for taxa identified in this study. DNA sequence read-length distributions were calculated from BWA alignments and BAM files.

| Environmental taxa (species) | BRS1 | BRS2 | BRS3 | BRS4 | BRS5 | Mean |
|---|---|---|---|---|---|---|
| *Babesia bovis* | *0* | *71* | *74* | *74* | *0* | 43.8 |
| *Lactococcus lactis* | *76* | *74* | *76* | *75* | *52* | 70.6 |
| *Ficus sur* | *62* | *73* | *72* | *72* | *56* | 67.0 |
| *Parastrongyloides trichosuri* | *51* | *74* | *76* | *76* | *48* | 65.0 |
| *Podospora anserina* | *59* | *69* | *77* | *74* | *28* | 61.4 |
| *Sorghum bicolor* | *74* | *70* | *73* | *73* | *61* | 70.2 |
| *Triticum aestivum* | *65* | *56* | *68* | *69* | *62* | 64.0 |
| *Zea mays* | *67* | *64* | *71* | *72* | *0* | 54.8 |
| *Bos taurus* | *69* | *62* | *72* | *75* | *36* | 62.8 |
| *Capra hircus* | *70* | *61* | *71* | *76* | *39* | 63.4 |
| *Equus caballus* | *53* | *43* | *61* | *71* | *28* | 51.2 |
| *Canis lupus familiaris* | – | – | – | – | *28* | 28.0 |
| *Homo sapiens* | *72* | *60* | *69* | *74* | *31* | 61.2 |
| *Macaca mulatta* | *67* | *55* | *64* | *74* | *30* | 58.0 |
| *Ovis aries* | *69* | *59* | *71* | *76* | *38* | 62.6 |
| *Pan troglodytes* | *70* | *59* | *69* | *73* | *32* | 60.6 |
| **Commensal taxa (genera)** | **BRS1** | **BRS2** | **BRS3** | **BRS4** | **BRS5** | **Mean** |
| *Methanobrevibacter* | *73* | *72* | *74* | *73* | *61* | 70.6 |
| *Bifidobacterium* | *69* | *73* | *76* | *74* | *71* | 72.6 |
| *Cellulomonas* | *58* | *75* | *77* | *75* | *48* | 66.6 |
| *Slackia* | *50* | *70* | *73* | *73* | *43* | 61.8 |
| *Alistipes* | *81* | *71* | *71* | *72* | *32* | 65.4 |
| *Bacteroides* | *67* | *76* | *77* | *76* | *65* | 72.2 |
| *Flavobacterium* | *51* | *74* | *76* | *75* | *46* | 64.4 |
| *Parabacteroides* | *62* | *72* | *73* | *75* | *68* | 70.0 |
| *Pedobacter* | *48* | *74* | *75* | *75* | *42* | 62.8 |
| *Prevotella* | *56* | *68* | *71* | *73* | *37* | 61.0 |
| *Blautia* | *60* | *69* | *74* | *73* | *65* | 68.2 |
| *Butyrivibrio* | *41* | *70* | *71* | *72* | *37* | 58.2 |
| *Clostridium* | *58* | *73* | *75* | *75* | *48* | 65.8 |

| | | | | | | |
|---|---|---|---|---|---|---|
| *Coprococcus* | *69* | *71* | *72* | *72* | *56* | 68.0 |
| *Dorea* | *52* | *72* | *75* | *73* | *48* | 64.0 |
| *Enterococcus* | *71* | *75* | *77* | *75* | *75* | 74.6 |
| *Eubacterium* | *62* | *70* | *73* | *73* | *51* | 65.8 |
| *Faecalibacterium* | *64* | *72* | *75* | *73* | *45* | 65.8 |
| *Lactobacillus* | *70* | *74* | *76* | *74* | *51* | 69.0 |
| *Lactococcus* | *73* | *74* | *76* | *75* | *50* | 69.6 |
| *Mogibacterium* | *72* | *71* | *72* | *73* | *55* | 68.6 |
| *Mollicutes* | *59* | *72* | *72* | *72* | *34* | 61.8 |
| *Oribacterium* | *65* | *70* | *72* | *72* | *44* | 64.6 |
| *Oscillibacter* | *54* | *71* | *75* | *73* | *42* | 63.0 |
| *Butyrivibrio* | *41* | *70* | *71* | *72* | *37* | 58.2 |
| *Phascolarctobacterium* | *67* | *69* | *75* | *72* | *52* | 67.0 |
| *Romboutsia* | *63* | *69* | *72* | *72* | *60* | 67.2 |
| *Roseburia* | *63* | *69* | *73* | *72* | *65* | 68.4 |
| *Ruminiclostridium* | *61* | *69* | *74* | *72* | *28* | 60.8 |
| *Ruminococcus* | *59* | *73* | *75* | *74* | *56* | 67.4 |
| *Enterobacter* | *69* | *76* | *77* | *76* | *61* | 71.8 |
| *Sarcina* | *40* | *72* | *74* | *77* | *72* | 67.0 |
| *Tissierellia* | *64* | *72* | *75* | *74* | *61* | 69.2 |
| *Enterobacter* | *69* | *76* | *77* | *76* | *61* | 71.8 |
| *Klebsiella* | *63* | *75* | *77* | *76* | *59* | 70.0 |
| *Neisseria* | *46* | *71* | *71* | *72* | *43* | 60.6 |
| *Sphingobium* | *56* | *74* | *77* | *75* | *47* | 65.8 |
| *Variovorax* | *65* | *73* | *76* | *75* | *59* | 69.6 |

| **Pathogenic taxa (genera)** | **BRS1** | **BRS2** | **BRS3** | **BRS4** | **BRS5** | **Mean** |
|---|---|---|---|---|---|---|
| *Aeromonas* | *65* | *73* | *76* | *75* | *47* | 67.2 |
| *Achromobacter* | *69* | *76* | *77* | *76* | *64* | 72.4 |
| *Bordetella* | *66* | *74* | *76* | *75* | *57* | 69.6 |
| *Burkholderia* | *47* | *71* | *75* | *74* | *43* | 62.0 |
| *Comamonas* | *69* | *74* | *76* | *76* | *72* | 73.4 |
| *Delftia* | *76* | *74* | *76* | *75* | *73* | 74.8 |
| *Elizabethkingia* | *71* | *73* | *76* | *74* | *30* | 64.8 |

| | | | | | | |
|---|---|---|---|---|---|---|
| *Listeria* | *68* | *75* | *77* | *75* | *69* | 72.8 |
| *Lysobacter* | *52* | *76* | *78* | *76* | *49* | 66.2 |
| *Mycobacterium* | *69* | *74* | *76* | *75* | *52* | 69.2 |
| *Shigella* | *77* | *76* | *77* | *77* | *77* | 76.8 |
| *Vibrio* | *49* | *72* | *75* | *73* | *45* | 62.8 |
| *Weeksella* | *0* | *75* | *77* | *75* | *0* | 45.4 |

**Supplementary Table 6.** Relative abundance of eighteen significant KEGG pathways detected in BRS and the ancient (Ötzi) and modern (Italian, Hadza and Malawian) comparative IM cohorts.

| KEGG pathways | BRS2 | BRS3 | BRS4 | Hadza_H15 | Hadza_H17 | Hadza_H11 | Hadza_H9 | Italian_IT5 | Italian_IT8 | Italian-IT2 | Italian-IT6 | Malawi _mgm4461146 | Malawi _mgm4461161 | Malawi _mgm4461165 | Malawi _mgm4461162 | Otzi _LPLI | Otzi _UPLI | Otzi _UPLI2 | Otzi _Samll_Int |
|---|---|---|---|---|---|---|---|---|---|---|---|---|---|---|---|---|---|---|---|
| K1000010 Glycolysis / Gluconeogenesis | 261 | 269 | 286 | 192 | 187 | 212 | 216 | 217 | 324 | 290 | 279 | 329 | 513 | 288 | 173 | 385 | 261 | 266 | 300 |
| K1000020 Citrate cycle (TCA cycle) | 342 | 345 | 378 | 116 | 119 | 123 | 122 | 142 | 166 | 146 | 159 | 150 | 189 | 133 | 78 | 98 | 240 | 239 | 294 |
| K1000051 Fructose and mannose metabolism | 94 | 89 | 102 | 111 | 119 | 134 | 135 | 178 | 195 | 147 | 191 | 112 | 176 | 98 | 109 | 297 | 108 | 112 | 118 |
| K1000052 Galactose metabolism | 82 | 77 | 92 | 142 | 145 | 146 | 177 | 187 | 310 | 245 | 283 | 370 | 539 | 319 | 155 | 296 | 77 | 79 | 72 |
| K1000500 Starch and sucrose metabolism | 193 | 187 | 199 | 202 | 208 | 216 | 244 | 243 | 512 | 399 | 356 | 483 | 796 | 405 | 223 | 367 | 164 | 176 | 186 |
| K1000520 Amino sugar and nucleotide sugar metabolism | 206 | 212 | 217 | 205 | 211 | 216 | 241 | 263 | 373 | 313 | 329 | 459 | 706 | 405 | 216 | 535 | 215 | 223 | 237 |
| K1000620 Pyruvate metabolism | 317 | 321 | 353 | 199 | 196 | 219 | 234 | 231 | 342 | 304 | 289 | 285 | 444 | 244 | 167 | 374 | 333 | 339 | 395 |
| K1000630 Glyoxylate and dicarboxylate metabolism | 189 | 180 | 209 | 77 | 85 | 97 | 98 | 109 | 166 | 138 | 137 | 140 | 167 | 108 | 78 | 154 | 165 | 172 | 200 |
| K1000640 Propanoate metabolism | 217 | 221 | 230 | 98 | 91 | 101 | 100 | 113 | 144 | 142 | 135 | 130 | 189 | 114 | 66 | 210 | 246 | 254 | 297 |
| K1000650 Butanoate metabolism | 190 | 192 | 236 | 102 | 114 | 127 | 120 | 123 | 155 | 163 | 151 | 168 | 209 | 138 | 67 | 242 | 227 | 233 | 265 |
| K1000072 Synthesis and degradation of ketone bodies | 27 | 27 | 36 | 8 | 9 | 10 | 10 | 9 | 6 | 9 | 12 | 6 | 8 | 5 | 4 | 28 | 37 | 38 | 48 |
| K1000600 Sphingolipid metabolism | 17 | 18 | 21 | 50 | 55 | 46 | 59 | 72 | 85 | 82 | 109 | 118 | 202 | 111 | 54 | 97 | 21 | 19 | 20 |
| K1001040 Biosynthesis of unsaturated fatty acids | 29 | 30 | 32 | 10 | 10 | 12 | 13 | 14 | 19 | 24 | 16 | 25 | 49 | 21 | 9 | 31 | 49 | 51 | 66 |
| K1000510 N-Glycan biosynthesis | 17 | 15 | 24 | 9 | 11 | 4 | 5 | 6 | 2 | 4 | 3 | 1 | 4 | 1 | 5 | 16 | 22 | 24 | 18 |
| K1000603 Glycosphingolipid biosynthesis - globo series | 6 | 6 | 7 | 21 | 21 | 16 | 19 | 36 | 40 | 42 | 46 | 59 | 84 | 51 | 16 | 29 | 4 | 4 | 3 |
| K1000604 Glycosphingolipid biosynthesis - ganglio series | 2 | 3 | 4 | 6 | 8 | 4 | 6 | 21 | 1 | 9 | 19 | 15 | 4 | 6 | 5 | 9 | 2 | 2 | 1 |
| K1000625 Chloroalkane and chloroalkene degradation | 3 | 2 | 4 | 2 | 2 | 3 | 2 | 3 | 3 | 4 | 4 | 7 | 2 | 4 | 4 | 17 | 6 | 7 | 6 |
| K1000626 Naphthalene degradation | 6 | 6 | 11 | 1 | 3 | 2 | 1 | 0 | 0 | 0 | 1 | 0 | 1 | 0 | 0 | 6 | 7 | 5 | 6 |

**Supplementary Table 7.** Enrichment and depletion of KO metabolic gene categories in the BRS and ancient (Ötzi) and modern (Italian, Hadza and Malawian) comparative IM sample cohorts based on *p* -value (*p* = <0.05) designation. Group significance analyses were performed using Qiime v1.9.1.

| ID | Test-Statistic | *P* | BRS_mean | Otzi-UPLI_mean | Otzi-SI_mean | Otzi-LPLI_mean | Hadza_mean | Malawi_mean | Italian_mean | KEGG categories |
|---|---|---|---|---|---|---|---|---|---|---|
| ID3087 | 18.00000 | 0.00623220 | 3.00 | 4.00 | 4.00 | 1.00 | 1.00 | 0.00 | 1.00 | K03087 RNA polymerase nonessential primary-like sigma factor |
| ID167 | 17.93662 | 0.00639255 | 3.33 | 5.00 | 7.00 | 1.00 | 0.00 | 0.00 | 0.00 | K00167 2-oxoisovalerate dehydrogenase E1 component beta subunit [EC:1.2.4.4] |
| ID322 | 17.93110 | 0.00640670 | 5.33 | 7.50 | 10.00 | 2.00 | 1.00 | 0.00 | 1.00 | K00322 NAD(P) transhydrogenase [EC:1.6.1.1] |
| ID1959 | 17.91396 | 0.00645085 | 4.00 | 12.50 | 17.00 | 2.00 | 1.00 | 0.00 | 1.00 | K01959 pyruvate carboxylase subunit A [EC:6.4.1.1] |
| ID1497 | 17.90448 | 0.00647540 | 4.67 | 3.00 | 4.00 | 1.00 | 1.00 | 0.00 | 1.00 | K01497 GTP cyclohydrolase II [EC:3.5.4.25] |
| ID130 | 17.89080 | 0.00651095 | 3.00 | 8.00 | 11.00 | 2.00 | 1.00 | 0.00 | 1.00 | K00130 betaine-aldehyde dehydrogenase [EC:1.2.1.8] |
| ID11073 | 17.89060 | 0.00651150 | 5.00 | 16.50 | 22.00 | 4.00 | 1.00 | 0.00 | 1.00 | K11073 putrescine transport system substrate-binding protein |
| ID252 | 17.88693 | 0.00652107 | 9.33 | 7.00 | 10.00 | 2.00 | 0.00 | 0.00 | 0.00 | K00252 glutaryl-CoA dehydrogenase [EC:1.3.8.6] |
| ID412 | 17.82775 | 0.00667738 | 12.33 | 24.00 | 13.00 | 2.00 | 1.00 | 0.00 | 0.00 | K00412 ubiquinol-cytochrome c reductase cytochrome b subunit |
| ID257 | 17.82775 | 0.00667738 | 7.33 | 14.50 | 21.00 | 7.00 | 1.00 | 0.00 | 1.00 | K00257 [EC:1.3.99.-] |
| ID2914 | 17.81250 | 0.00671825 | 1.00 | 1.00 | 1.00 | 1.00 | 1.00 | 4.75 | 1.00 | K02914 large subunit ribosomal protein L34 |
| ID1659 | 17.73750 | 0.00692280 | 5.33 | 4.50 | 5.00 | 1.00 | 1.00 | 0.00 | 1.00 | K01659 2-methylcitrate synthase [EC:2.3.3.5] |
| ID2261 | 17.70457 | 0.00701449 | 4.00 | 10.00 | 2.00 | 1.00 | 0.00 | 0.00 | 0.00 | K02261 cytochrome c oxidase subunit 2 |
| ID451 | 17.69081 | 0.00705316 | 4.33 | 4.50 | 6.00 | 1.00 | 0.00 | 0.00 | 0.00 | K00451 homogentisate 1 |
| ID8484 | 17.42281 | 0.00784874 | 4.33 | 11.50 | 17.00 | 2.00 | 1.25 | 0.00 | 1.00 | K08484 phosphotransferase system |
| ID2109 | 17.38441 | 0.00796960 | 1.00 | 3.00 | 5.00 | 3.00 | 1.00 | 7.25 | 2.75 | K02109 F-type H+-transporting ATPase subunit b |
| ID1941 | 17.33138 | 0.00813950 | 9.00 | 10.50 | 15.00 | 2.00 | 1.00 | 0.00 | 1.25 | K01941 urea carboxylase [EC:6.3.4.6] |
| ID1690 | 17.03858 | 0.00914251 | 5.00 | 8.00 | 11.00 | 2.00 | 1.00 | 0.25 | 1.00 | K01690 phosphogluconate dehydratase [EC:4.2.1.12] |
| ID632 | 17.03858 | 0.00914251 | 9.00 | 8.00 | 10.00 | 2.00 | 1.00 | 0.25 | 1.00 | K00632 acetyl-CoA acyltransferase [EC:2.3.1.16] |
| ID2946 | 17.01244 | 0.00923762 | 6.00 | 2.50 | 2.00 | 8.00 | 4.00 | 10.50 | 4.75 | K02946 small subunit ribosomal protein S10 |
| ID799 | 17.00188 | 0.00927634 | 5.00 | 9.50 | 11.00 | 2.00 | 1.00 | 0.00 | 0.75 | K00799 glutathione S-transferase [EC:2.5.1.18] |
| ID2935 | 16.99717 | 0.00929366 | 1.00 | 1.00 | 2.00 | 3.00 | 2.00 | 6.75 | 2.50 | K02935 large subunit ribosomal protein L7/L12 |
| ID991 | 16.99110 | 0.00931602 | 1.00 | 2.00 | 3.00 | 11.00 | 2.00 | 11.00 | 5.25 | K00991 2-C-methyl-D-erythritol 4-phosphate cytidylyltransferase [EC:2.7.7.60] |
| ID651 | 16.97235 | 0.00938542 | 2.00 | 1.00 | 1.00 | 1.00 | 5.00 | 17.00 | 9.75 | K00651 homoserine O-succinyltransferase [EC:2.3.1.46] |
| ID11891 | 16.97143 | 0.00938882 | 3.67 | 11.50 | 14.00 | 2.00 | 1.00 | 0.25 | 1.00 | K11891 type VI secretion system protein ImpL |
| ID940 | 16.93421 | 0.00952810 | 3.00 | 4.00 | 5.00 | 4.00 | 1.00 | 1.25 | 1.00 | K00940 nucleoside-diphosphate kinase [EC:2.7.4.6] |
| ID1637 | 16.91232 | 0.00961096 | 12.00 | 8.50 | 11.00 | 2.00 | 1.00 | 0.25 | 1.00 | K01637 isocitrate lyase [EC:4.1.3.1] |
| ID1493 | 16.90525 | 0.00963783 | 2.33 | 1.00 | 1.00 | 6.00 | 4.00 | 2.50 | 4.25 | K01493 dCMP deaminase [EC:3.5.4.12] |
| ID334 | 16.89087 | 0.00969277 | 4.00 | 4.00 | 4.00 | 1.00 | 1.00 | 0.00 | 1.50 | K00334 NADH-quinone oxidoreductase subunit E [EC:1.6.5.3] |
| ID1776 | 16.89076 | 0.00969323 | 1.67 | 3.00 | 3.00 | 12.00 | 3.00 | 6.00 | 5.50 | K01776 glutamate racemase [EC:5.1.1.3] |
| ID406 | 16.85888 | 0.00981611 | 4.33 | 6.00 | 8.00 | 1.00 | 1.00 | 0.25 | 0.00 | K00406 cytochrome c oxidase cbb3-type subunit III |
| ID404 | 16.84322 | 0.00987704 | 12.67 | 15.50 | 23.00 | 4.00 | 1.00 | 0.25 | 0.00 | K00404 cytochrome c oxidase cbb3-type subunit I [EC:1.9.3.1] |
| ID1818 | 16.83761 | 0.00989896 | 1.00 | 1.00 | 1.00 | 10.00 | 4.50 | 1.75 | 7.75 | K01818 L-fucose isomerase [EC:5.3.1.25] |
| ID7646 | 16.82345 | 0.00995447 | 1.00 | 7.50 | 9.00 | 4.00 | 4.00 | 1.75 | 11.75 | K07646 two-component system |
| ID1803 | 16.79667 | 0.01006030 | 3.33 | 4.00 | 6.00 | 7.00 | 5.75 | 11.00 | 8.25 | K01803 triosephosphate isomerase (TIM) [EC:5.3.1.1] |
| ID12339 | 16.74855 | 0.01025317 | 3.00 | 4.50 | 7.00 | 1.00 | 0.75 | 0.00 | 1.00 | K12339 cysteine synthase B [EC:2.5.1.47] |
| ID568 | 16.74806 | 0.01025515 | 3.33 | 4.00 | 5.00 | 1.00 | 1.00 | 0.00 | 0.75 | K00568 2-polyprenyl-6-hydroxyphenyl methylase / 3-demethylubiquinone-9 3-methyltransferase [EC:2.1.1.222 2.1.1.64] |
| ID873 | 16.73939 | 0.01029029 | 9.33 | 11.00 | 12.00 | 38.00 | 8.75 | 19.50 | 14.50 | K00873 pyruvate kinase [EC:2.7.1.40] |
| ID10941 | 16.73183 | 0.01032101 | 4.33 | 6.00 | 9.00 | 1.00 | 1.00 | 0.00 | 0.75 | K10941 sigma-54 specific transcriptional regulator |
| ID956 | 16.67975 | 0.01053514 | 4.00 | 1.00 | 1.00 | 1.00 | 3.00 | 1.50 | 7.75 | K00956 sulfate adenylyltransferase subunit 1 [EC:2.7.7.4] |
| ID2275 | 16.67892 | 0.01053859 | 9.00 | 4.00 | 6.00 | 1.00 | 0.25 | 0.00 | 0.00 | K02275 cytochrome c oxidase subunit II [EC:1.9.3.1] |
| ID1772 | 16.65636 | 0.01063270 | 7.00 | 5.00 | 6.00 | 1.00 | 1.00 | 0.75 | 1.00 | K01772 ferrochelatase [EC:4.99.1.1] |
| ID1070 | 16.64659 | 0.01067372 | 3.67 | 4.50 | 6.00 | 1.00 | 1.00 | 0.25 | 1.00 | K01070 S-formylglutathione hydrolase [EC:3.1.2.12] |
| ID140 | 16.64624 | 0.01067519 | 9.67 | 21.00 | 28.00 | 5.00 | 1.00 | 0.25 | 1.50 | K00140 malonate-semialdehyde dehydrogenase (acetylating) / methylmalonate-semialdehyde dehydrogenase [EC:1.2.1.18 1.2.1.27] |
| ID1433 | 16.63416 | 0.01072611 | 6.67 | 15.50 | 20.00 | 3.00 | 1.00 | 0.25 | 1.25 | K01433 formyltetrahydrofolate deformylase [EC:3.5.1.10] |
| ID13482 | 16.61111 | 0.01082391 | 3.67 | 7.00 | 10.00 | 2.00 | 0.50 | 0.00 | 0.00 | K13482 xanthine dehydrogenase large subunit [EC:1.17.1.4] |
| ID626 | 16.60924 | 0.01083191 | 14.67 | 20.50 | 26.00 | 21.00 | 4.75 | 2.50 | 5.25 | K00626 acetyl-CoA C-acetyltransferase [EC:2.3.1.9] |
| ID457 | 16.59404 | 0.01089691 | 5.33 | 10.50 | 12.00 | 3.00 | 0.50 | 0.00 | 0.00 | K00457 4-hydroxyphenylpyruvate dioxygenase [EC:1.13.11.27] |
| ID7788 | 16.59404 | 0.01089691 | 4.00 | 10.00 | 13.00 | 2.00 | 1.00 | 0.50 | 1.00 | K07788 multidrug efflux pump |
| ID121 | 16.58648 | 0.01092938 | 9.00 | 9.00 | 9.00 | 2.00 | 1.00 | 0.25 | 1.00 | K00121 S-(hydroxymethyl)glutathione dehydrogenase / alcohol dehydrogenase [EC:1.1.1.284 1.1.1.1] |
| ID302 | 16.58491 | 0.01093616 | 0.67 | 23.00 | 31.00 | 4.00 | 1.00 | 0.00 | 1.00 | K00302 sarcosine oxidase |
| ID1807 | 16.58349 | 0.01094224 | 2.00 | 3.00 | 4.00 | 1.00 | 1.00 | 9.75 | 1.25 | K01807 ribose 5-phosphate isomerase A [EC:5.3.1.6] |
| ID2276 | 16.57933 | 0.01096017 | 6.00 | 4.00 | 6.00 | 1.00 | 0.00 | 0.00 | 0.25 | K02276 cytochrome c oxidase subunit III [EC:1.9.3.1] |
| ID1011 | 16.56643 | 0.01101599 | 7.00 | 4.00 | 5.00 | 9.00 | 1.00 | 0.50 | 1.00 | K01011 thiosulfate/3-mercaptopyruvate sulfurtransferase [EC:2.8.1.1 2.8.1.2] |
| ID117 | 16.55802 | 0.01105247 | 5.00 | 9.50 | 13.00 | 2.00 | 1.25 | 0.25 | 1.00 | K00117 quinoprotein glucose dehydrogenase [EC:1.1.5.2] |
| ID7665 | 16.54832 | 0.01109477 | 1.00 | 11.00 | 13.00 | 2.00 | 1.00 | 0.00 | 2.50 | K07665 two-component system |
| ID3210 | 16.53783 | 0.01114064 | 1.00 | 2.00 | 3.00 | 4.00 | 1.00 | 4.00 | 1.75 | K03210 preprotein translocase subunit YajC |
| ID955 | 16.52645 | 0.01119058 | 4.67 | 8.00 | 10.00 | 2.00 | 1.00 | 0.25 | 1.25 | K00955 bifunctional enzyme CysN/CysC [EC:2.7.7.4 2.7.1.25] |
| ID6445 | 16.52645 | 0.01119058 | 6.33 | 12.00 | 17.00 | 2.00 | 1.25 | 0.25 | 1.00 | K06445 acyl-CoA dehydrogenase [EC:1.3.99.-] |
| ID5365 | 16.52645 | 0.01119058 | 3.33 | 11.00 | 15.00 | 2.00 | 1.25 | 0.25 | 1.00 | K05365 penicillin-binding protein 1B [EC:2.4.1.129 3.4.-.-] |
| ID6596 | 16.50925 | 0.01126655 | 7.67 | 10.00 | 15.00 | 2.00 | 0.50 | 0.00 | 0.00 | K06596 chemosensory pili system protein ChpA (sensor histidine kinase/response regulator) |
| ID3735 | 16.50000 | 0.01130760 | 4.00 | 8.00 | 9.00 | 4.00 | 1.00 | 0.50 | 1.00 | K03735 ethanolamine ammonia-lyase large subunit [EC:4.3.1.7] |
| ID1546 | 16.49783 | 0.01131726 | 1.00 | 7.00 | 7.00 | 2.00 | 1.00 | 0.00 | 3.75 | K01546 K+-transporting ATPase ATPase A chain [EC:3.6.3.12] |
| ID3205 | 16.49732 | 0.01131953 | 1.33 | 1.00 | 2.00 | 1.00 | 30.50 | 14.00 | 60.75 | K03205 type IV secretion system protein VirD4 |
| ID7712 | 16.49072 | 0.01134892 | 6.33 | 8.00 | 11.00 | 2.00 | 1.00 | 0.50 | 1.00 | K07712 two-component system |
| ID1192 | 16.48855 | 0.01135861 | 1.00 | 1.00 | 1.00 | 1.00 | 1.00 | 5.50 | 6.00 | K01192 beta-mannosidase [EC:3.2.1.25] |
| ID3644 | 16.48324 | 0.01138234 | 12.67 | 8.00 | 11.00 | 2.00 | 1.00 | 0.25 | 1.50 | K03644 lipoic acid synthetase [EC:2.8.1.8] |

| | | | | | | | | | | |
|---|---|---|---|---|---|---|---|---|---|---|
| ID286 | 16.47522 | 0.01141826 | 1.00 | 2.00 | 3.00 | 12.00 | 2.75 | 9.00 | 5.50 | K00286 pyrroline-5-carboxylate reductase [EC:1.5.1.2] |
| ID1584 | 16.46926 | 0.01144507 | 11.33 | 7.50 | 10.00 | 2.00 | 1.25 | 0.25 | 1.00 | K01584 arginine decarboxylase [EC:4.1.1.19] |
| ID370 | 16.46413 | 0.01146814 | 15.67 | 8.50 | 11.00 | 2.00 | 1.50 | 1.00 | 1.00 | K00370 nitrate reductase alpha subunit [EC:1.7.99.4] |
| ID3524 | 16.46126 | 0.01148106 | 1.00 | 2.00 | 2.00 | 7.00 | 3.50 | 13.00 | 7.50 | K03524 BirA family transcriptional regulator |
| ID372 | 16.44799 | 0.01154112 | 4.00 | 3.50 | 5.00 | 18.00 | 1.00 | 0.00 | 0.50 | K00372 assimilatory nitrate reductase catalytic subunit [EC:1.7.99.4] |
| ID1893 | 16.44061 | 0.01157459 | 8.00 | 3.00 | 1.00 | 20.00 | 10.00 | 3.75 | 11.75 | K01893 asparaginyl-tRNA synthetase [EC:6.1.1.22] |
| ID1241 | 16.42999 | 0.01162298 | 5.33 | 8.00 | 12.00 | 2.00 | 1.00 | 0.00 | 1.00 | K01241 AMP nucleosidase [EC:3.2.2.4] |
| ID1638 | 16.41476 | 0.01169273 | 12.00 | 13.00 | 18.00 | 3.00 | 1.50 | 0.25 | 1.00 | K01638 malate synthase [EC:2.3.3.9] |
| ID3150 | 16.40550 | 0.01173532 | 1.00 | 3.50 | 2.00 | 23.00 | 8.50 | 4.25 | 12.75 | K03150 2-iminoacetate synthase [EC:4.1.99.19] |
| ID845 | 16.39381 | 0.01178930 | 1.33 | 2.00 | 3.00 | 14.00 | 4.75 | 10.75 | 8.00 | K00845 glucokinase [EC:2.7.1.2] |
| ID10117 | 16.38350 | 0.01183711 | 2.33 | 1.00 | 1.00 | 4.00 | 10.75 | 82.75 | 28.25 | K10117 raffinose/stachyose/melibiose transport system substrate-binding protein |
| ID3381 | 16.37049 | 0.01189767 | 2.00 | 5.00 | 6.00 | 1.00 | 0.25 | 0.00 | 0.00 | K03381 catechol 1 |
| ID3525 | 16.35000 | 0.01199371 | 1.33 | 3.00 | 3.00 | 12.00 | 3.00 | 9.25 | 6.25 | K03525 type III pantothenate kinase [EC:2.7.1.33] |
| ID1712 | 16.33644 | 0.01205769 | 7.67 | 12.00 | 14.00 | 5.00 | 2.50 | 0.25 | 3.50 | K01712 urocanate hydratase [EC:4.2.1.49] |
| ID1961 | 16.33105 | 0.01208318 | 14.00 | 10.50 | 12.00 | 15.00 | 3.00 | 2.50 | 9.25 | K01961 acetyl-CoA carboxylase |
| ID1791 | 16.32917 | 0.01209211 | 6.00 | 4.00 | 1.00 | 15.00 | 5.50 | 3.00 | 3.75 | K01791 UDP-N-acetylglucosamine 2-epimerase (non-hydrolysing) [EC:5.1.3.14] |
| ID647 | 16.31602 | 0.01215461 | 6.33 | 6.00 | 9.00 | 1.00 | 1.00 | 0.75 | 1.00 | K00647 3-oxoacyl-[acyl-carrier-protein] synthase I [EC:2.3.1.41] |
| ID2888 | 16.31107 | 0.01217823 | 2.33 | 2.00 | 3.00 | 3.00 | 1.50 | 7.00 | 3.00 | K02888 large subunit ribosomal protein L21 |
| ID1950 | 16.30406 | 0.01221174 | 3.00 | 4.00 | 1.00 | 19.00 | 6.75 | 15.25 | 14.00 | K01950 NAD+ synthase (glutamine-hydrolysing) [EC:6.3.5.1] |
| ID1929 | 16.29461 | 0.01225705 | 1.67 | 4.00 | 6.00 | 11.00 | 5.50 | 18.75 | 10.75 | K01929 UDP-N-acetylmuramoyl-tripeptide--D-alanyl-D-alanine ligase [EC:6.3.2.10] |
| ID1243 | 16.28435 | 0.01230645 | 1.00 | 1.00 | 1.00 | 11.00 | 2.25 | 12.50 | 4.75 | K01243 adenosylhomocysteine nucleosidase [EC:3.2.2.9] |
| ID13893 | 16.28174 | 0.01231903 | 6.33 | 19.00 | 23.00 | 4.00 | 1.25 | 0.25 | 1.50 | K13893 microcin C transport system substrate-binding protein |
| ID2343 | 16.28125 | 0.01232141 | 12.00 | 9.00 | 11.00 | 23.00 | 7.75 | 29.75 | 12.75 | K02343 DNA polymerase III subunit gamma/tau [EC:2.7.7.7] |
| ID331 | 16.26697 | 0.01239055 | 6.33 | 5.00 | 6.00 | 1.00 | 1.00 | 0.25 | 1.50 | K00331 NADH-quinone oxidoreductase subunit B [EC:1.6.5.3] |
| ID2002 | 16.26036 | 0.01242270 | 3.00 | 21.00 | 26.00 | 4.00 | 1.00 | 0.50 | 1.25 | K02002 glycine betaine/proline transport system substrate-binding protein |
| ID1907 | 16.25336 | 0.01245679 | 3.33 | 4.50 | 6.00 | 1.00 | 0.50 | 0.00 | 0.00 | K01907 acetoacetyl-CoA synthetase [EC:6.2.1.16] |
| ID362 | 16.24897 | 0.01247821 | 9.00 | 13.50 | 19.00 | 3.00 | 1.25 | 0.75 | 1.00 | K00362 nitrite reductase (NADH) large subunit [EC:1.7.1.15] |
| ID1255 | 16.24897 | 0.01247821 | 5.00 | 7.00 | 8.00 | 17.00 | 1.25 | 0.75 | 1.00 | K01255 leucyl aminopeptidase [EC:3.4.11.1] |
| ID3841 | 16.22255 | 0.01260800 | 4.67 | 6.00 | 8.00 | 1.00 | 1.25 | 0.25 | 1.00 | K03841 fructose-1 |
| ID3655 | 16.21463 | 0.01264716 | 9.00 | 11.50 | 13.00 | 19.00 | 17.00 | 63.25 | 26.75 | K03655 ATP-dependent DNA helicase RecG [EC:3.6.4.12] |
| ID3338 | 16.21113 | 0.01266451 | 1.00 | 7.00 | 8.00 | 4.00 | 1.00 | 0.25 | 1.00 | K03338 5-dehydro-2-deoxygluconokinase [EC:2.7.1.92] |
| ID11754 | 16.17800 | 0.01282980 | 2.00 | 4.00 | 5.00 | 11.00 | 4.50 | 16.00 | 10.50 | K11754 dihydrofolate synthase / folylpolyglutamate synthase [EC:6.3.2.12 6.3.2.17] |
| ID3415 | 16.16929 | 0.01287359 | 3.67 | 13.00 | 17.00 | 3.00 | 1.25 | 0.50 | 1.00 | K03415 two-component system |
| ID2864 | 16.16904 | 0.01287487 | 2.00 | 3.00 | 4.00 | 7.00 | 2.75 | 9.75 | 4.00 | K02864 large subunit ribosomal protein L10 |
| ID1682 | 16.16568 | 0.01289179 | 14.67 | 11.00 | 15.00 | 3.00 | 1.25 | 0.50 | 1.00 | K01682 aconitate hydratase 2 / 2-methylisocitrate dehydratase [EC:4.2.1.3 4.2.1.99] |
| ID1077 | 16.16357 | 0.01290243 | 1.00 | 1.00 | 1.00 | 1.00 | 2.00 | 20.75 | 5.50 | K01077 alkaline phosphatase [EC:3.1.3.1] |
| ID10441 | 16.16340 | 0.01290328 | 1.67 | 9.50 | 12.00 | 15.00 | 4.50 | 2.50 | 5.75 | K10441 ribose transport system ATP-binding protein [EC:3.6.3.17] |
| ID1141 | 16.15410 | 0.01295033 | 3.00 | 4.00 | 6.00 | 1.00 | 1.00 | 0.50 | 1.00 | K01141 exodeoxyribonuclease I [EC:3.1.11.1] |
| ID3583 | 16.15339 | 0.01295391 | 3.67 | 11.50 | 16.00 | 3.00 | 1.25 | 0.50 | 1.00 | K03583 exodeoxyribonuclease V gamma subunit [EC:3.1.11.5] |
| ID1969 | 16.12927 | 0.01307671 | 6.67 | 6.50 | 9.00 | 1.00 | 0.50 | 0.00 | 0.00 | K01969 3-methylcrotonyl-CoA carboxylase beta subunit [EC:6.4.1.4] |
| ID10125 | 16.12174 | 0.01311525 | 2.00 | 10.00 | 14.00 | 2.00 | 0.00 | 0.25 | 0.75 | K10125 two-component system |
| ID631 | 16.10417 | 0.01320565 | 4.33 | 11.00 | 15.00 | 2.00 | 1.25 | 0.75 | 1.00 | K00631 glycerol-3-phosphate O-acyltransferase [EC:2.3.1.15] |
| ID1139 | 16.10281 | 0.01321265 | 4.00 | 9.50 | 13.00 | 2.00 | 1.25 | 0.50 | 1.00 | K01139 guanosine-3' |
| ID11963 | 16.10176 | 0.01321809 | 3.00 | 4.00 | 5.00 | 1.00 | 1.00 | 2.75 | 0.00 | K11963 urea transport system ATP-binding protein |
| ID1209 | 16.10169 | 0.01321845 | 7.67 | 1.00 | 1.00 | 1.00 | 5.00 | 23.00 | 11.00 | K01209 alpha-N-arabinofuranosidase [EC:3.2.1.55] |
| ID3582 | 16.08762 | 0.01329135 | 4.00 | 9.00 | 12.00 | 2.00 | 1.25 | 0.75 | 1.00 | K03582 exodeoxyribonuclease V beta subunit [EC:3.1.11.5] |
| ID285 | 16.08621 | 0.01329866 | 4.33 | 9.00 | 13.00 | 2.00 | 1.00 | 0.25 | 0.75 | K00285 D-amino-acid dehydrogenase [EC:1.4.99.1] |
| ID7649 | 16.07786 | 0.01334208 | 1.00 | 5.00 | 8.00 | 1.00 | 0.25 | 0.00 | 0.00 | K07649 two-component system |
| ID11904 | 16.07081 | 0.01337889 | 5.67 | 10.00 | 12.00 | 2.00 | 1.00 | 0.25 | 0.75 | K11904 type VI secretion system secreted protein VgrG |
| ID11907 | 16.07081 | 0.01337889 | 4.33 | 12.50 | 16.00 | 2.00 | 1.00 | 0.25 | 0.75 | K11907 type VI secretion system protein VasG |
| ID8311 | 16.06931 | 0.01338676 | 4.00 | 5.00 | 5.00 | 1.00 | 1.00 | 0.50 | 1.00 | K08311 putative (di)nucleoside polyphosphate hydrolase [EC:3.6.1.-] |
| ID2909 | 16.06713 | 0.01339816 | 4.00 | 2.00 | 2.00 | 5.00 | 2.00 | 7.00 | 2.25 | K02909 large subunit ribosomal protein L31 |
| ID7173 | 16.05550 | 0.01345913 | 1.00 | 1.00 | 1.00 | 4.00 | 2.25 | 7.25 | 4.00 | K07173 S-ribosylhomocysteine lyase [EC:4.4.1.21] |
| ID108 | 16.04410 | 0.01351919 | 3.67 | 11.00 | 15.00 | 3.00 | 1.00 | 0.25 | 0.75 | K00108 choline dehydrogenase |
| ID11076 | 16.04035 | 0.01353905 | 2.67 | 10.00 | 13.00 | 2.00 | 1.00 | 0.25 | 0.75 | K11076 putrescine transport system ATP-binding protein |
| ID3336 | 16.02959 | 0.01359605 | 1.00 | 8.00 | 9.00 | 7.00 | 1.00 | 0.25 | 2.00 | K03336 3D-(3 |
| ID1920 | 16.02072 | 0.01364319 | 4.00 | 4.50 | 6.00 | 1.00 | 1.00 | 0.50 | 1.00 | K01920 glutathione synthase [EC:6.3.2.3] |
| ID13895 | 16.01340 | 0.01368224 | 3.00 | 7.50 | 11.00 | 2.00 | 1.00 | 0.25 | 0.75 | K13895 microcin C transport system permease protein |
| ID2492 | 16.00984 | 0.01370124 | 3.33 | 6.50 | 8.00 | 9.00 | 1.00 | 1.00 | 2.75 | K02492 glutamyl-tRNA reductase [EC:1.2.1.70] |
| ID847 | 16.00938 | 0.01370369 | 1.00 | 3.00 | 3.00 | 8.00 | 5.50 | 12.50 | 10.25 | K00847 fructokinase [EC:2.7.1.4] |
| ID11962 | 16.00115 | 0.01374778 | 3.67 | 5.00 | 7.00 | 1.00 | 1.00 | 3.25 | 0.00 | K11962 urea transport system ATP-binding protein |
| ID2405 | 15.98719 | 0.01382288 | 3.33 | 5.50 | 7.00 | 2.00 | 1.50 | 0.75 | 1.00 | K02405 RNA polymerase sigma factor for flagellar operon FliA |
| ID128 | 15.97043 | 0.01391359 | 12.33 | 18.00 | 24.00 | 16.00 | 1.00 | 6.00 | 2.00 | K00128 aldehyde dehydrogenase (NAD+) [EC:1.2.1.3] |
| ID1012 | 15.96988 | 0.01391652 | 6.33 | 9.50 | 10.00 | 17.00 | 5.25 | 2.75 | 9.50 | K01012 biotin synthase [EC:2.8.1.6] |
| ID8483 | 15.96825 | 0.01392538 | 4.00 | 1.50 | 1.00 | 21.00 | 7.50 | 27.00 | 13.00 | K08483 phosphotransferase system |
| ID335 | 15.94628 | 0.01404522 | 7.00 | 8.00 | 8.00 | 4.00 | 3.25 | 1.50 | 4.75 | K00335 NADH-quinone oxidoreductase subunit F [EC:1.6.5.3] |
| ID10119 | 15.93434 | 0.01411077 | 2.33 | 1.00 | 1.00 | 2.00 | 5.00 | 51.25 | 13.50 | K10119 raffinose/stachyose/melibiose transport system permease protein |
| ID2274 | 15.92806 | 0.01414537 | 28.33 | 9.50 | 13.00 | 2.00 | 0.25 | 0.00 | 0.25 | K02274 cytochrome c oxidase subunit I [EC:1.9.3.1] |
| ID1113 | 15.92092 | 0.01418480 | 2.33 | 5.00 | 5.00 | 1.00 | 0.50 | 0.00 | 0.00 | K01113 alkaline phosphatase D [EC:3.1.3.1] |

| | | | | | | | | | | |
|---|---|---|---|---|---|---|---|---|---|---|
| ID1825 | 15.91387 | 0.01422383 | 4.67 | 9.00 | 13.00 | 2.00 | 1.25 | 0.25 | 0.75 | K01825 3-hydroxyacyl-CoA dehydrogenase / enoyl-CoA hydratase / 3-hydroxybutyryl-CoA epimerase / enoyl-CoA isomerase [EC:1.1.1.35 4.2.1.17 5.1.2.3 5.3.3.8] |
| ID3879 | 15.91083 | 0.01424066 | 2.67 | 4.50 | 2.00 | 1.00 | 0.25 | 0.00 | 0.00 | K03879 NADH-ubiquinone oxidoreductase chain 2 [EC:1.6.5.3] |
| ID1775 | 15.89413 | 0.01433365 | 3.00 | 5.50 | 6.00 | 18.00 | 6.75 | 11.50 | 10.50 | K01775 alanine racemase [EC:5.1.1.1] |
| ID5350 | 15.88308 | 0.01439550 | 2.00 | 1.00 | 1.00 | 1.00 | 2.50 | 13.75 | 4.75 | K05350 beta-glucosidase [EC:3.2.1.21] |
| ID2939 | 15.85929 | 0.01452950 | 1.67 | 2.00 | 3.00 | 5.00 | 2.00 | 8.75 | 4.00 | K02939 large subunit ribosomal protein L9 |
| ID928 | 15.85267 | 0.01456700 | 10.33 | 9.00 | 11.00 | 12.00 | 7.50 | 22.75 | 14.75 | K00928 aspartate kinase [EC:2.7.2.4] |
| ID7667 | 15.84885 | 0.01458864 | 1.00 | 2.50 | 3.00 | 1.00 | 3.50 | 1.25 | 4.75 | K07667 two-component system |
| ID12308 | 15.83407 | 0.01467281 | 1.67 | 1.00 | 1.00 | 17.00 | 6.00 | 52.75 | 14.25 | K12308 beta-galactosidase [EC:3.2.1.23] |
| ID995 | 15.82722 | 0.01471204 | 2.33 | 3.00 | 3.00 | 7.00 | 1.50 | 12.25 | 7.00 | K00995 CDP-diacylglycerol--glycerol-3-phosphate 3-phosphatidyltransferase [EC:2.7.8.5] |
| ID1676 | 15.82593 | 0.01471939 | 13.00 | 7.00 | 10.00 | 2.00 | 4.00 | 0.50 | 3.50 | K01676 fumarate hydratase |
| ID290 | 15.82345 | 0.01473361 | 4.00 | 1.00 | 1.00 | 1.00 | 7.25 | 2.25 | 10.00 | K00290 saccharopine dehydrogenase (NAD+ |
| ID163 | 15.81281 | 0.01479476 | 20.33 | 18.50 | 23.00 | 4.00 | 1.50 | 0.50 | 1.00 | K00163 pyruvate dehydrogenase E1 component [EC:1.2.4.1] |
| ID705 | 15.80343 | 0.01484884 | 3.67 | 4.00 | 5.00 | 19.00 | 11.75 | 48.00 | 23.75 | K00705 4-alpha-glucanotransferase [EC:2.4.1.25] |
| ID3100 | 15.80077 | 0.01486423 | 6.67 | 6.00 | 7.00 | 14.00 | 5.00 | 14.00 | 9.25 | K03100 signal peptidase I [EC:3.4.21.89] |
| ID641 | 15.79412 | 0.01490273 | 6.33 | 6.50 | 8.00 | 1.00 | 1.00 | 0.50 | 1.00 | K00641 homoserine O-acetyltransferase [EC:2.3.1.31] |
| ID1190 | 15.77768 | 0.01499832 | 3.33 | 2.00 | 1.00 | 48.00 | 32.75 | 60.50 | 51.00 | K01190 beta-galactosidase [EC:3.2.1.23] |
| ID248 | 15.77298 | 0.01502576 | 3.33 | 9.50 | 10.00 | 11.00 | 1.50 | 0.50 | 1.25 | K00248 butyryl-CoA dehydrogenase [EC:1.3.8.1] |
| ID1442 | 15.77246 | 0.01502880 | 1.00 | 1.00 | 1.00 | 15.00 | 3.00 | 15.75 | 12.50 | K01442 choloylglycine hydrolase [EC:3.5.1.24] |
| ID3101 | 15.76944 | 0.01504643 | 2.33 | 3.00 | 3.00 | 8.00 | 1.25 | 5.75 | 2.50 | K03101 signal peptidase II [EC:3.4.23.36] |
| ID336 | 15.75874 | 0.01510916 | 4.67 | 8.00 | 9.00 | 4.00 | 2.75 | 1.00 | 4.75 | K00336 NADH-quinone oxidoreductase subunit G [EC:1.6.5.3] |
| ID116 | 15.75668 | 0.01512131 | 8.00 | 10.00 | 14.00 | 2.00 | 1.00 | 0.50 | 0.75 | K00116 malate dehydrogenase (quinone) [EC:1.1.5.4] |
| ID648 | 15.75272 | 0.01514458 | 7.33 | 5.50 | 7.00 | 10.00 | 3.25 | 2.75 | 5.00 | K00648 3-oxoacyl-[acyl-carrier-protein] synthase III [EC:2.3.1.180] |
| ID3394 | 15.75000 | 0.01516061 | 1.00 | 4.00 | 4.00 | 4.00 | 1.00 | 1.50 | 1.00 | K03394 precorrin-2/cobalt-factor-2 C20-methyltransferase [EC:2.1.1.130 2.1.1.151] |
| ID2038 | 15.73997 | 0.01521982 | 4.67 | 8.00 | 10.00 | 4.00 | 3.00 | 12.75 | 6.25 | K02038 phosphate transport system permease protein |
| ID1478 | 15.72039 | 0.01533606 | 1.00 | 6.50 | 7.00 | 9.00 | 1.00 | 1.50 | 1.00 | K01478 arginine deiminase [EC:3.5.3.6] |
| ID3816 | 15.71322 | 0.01537890 | 1.00 | 4.00 | 5.00 | 9.00 | 3.00 | 7.00 | 4.75 | K03816 xanthine phosphoribosyltransferase [EC:2.4.2.22] |
| ID1692 | 15.71091 | 0.01539269 | 3.67 | 6.50 | 8.00 | 1.00 | 0.75 | 0.00 | 0.25 | K01692 enoyl-CoA hydratase [EC:4.2.1.17] |
| ID2454 | 15.71006 | 0.01539776 | 7.33 | 10.00 | 13.00 | 3.00 | 1.25 | 1.50 | 1.00 | K02454 general secretion pathway protein E |
| ID1679 | 15.70496 | 0.01542831 | 10.33 | 7.00 | 8.00 | 2.00 | 2.25 | 1.25 | 1.00 | K01679 fumarate hydratase |
| ID249 | 15.69458 | 0.01549062 | 10.67 | 13.50 | 18.00 | 3.00 | 0.50 | 0.00 | 0.25 | K00249 acyl-CoA dehydrogenase [EC:1.3.8.7] |
| ID965 | 15.69430 | 0.01549233 | 2.33 | 1.00 | 1.00 | 9.00 | 6.50 | 21.00 | 12.00 | K00965 UDPglucose--hexose-1-phosphate uridylyltransferase [EC:2.7.7.12] |
| ID381 | 15.69107 | 0.01551178 | 9.33 | 9.50 | 13.00 | 2.00 | 0.75 | 0.25 | 1.25 | K00381 sulfite reductase (NADPH) hemoprotein beta-component [EC:1.8.1.2] |
| ID957 | 15.69019 | 0.01551707 | 9.67 | 6.00 | 7.00 | 2.00 | 2.50 | 1.50 | 6.50 | K00957 sulfate adenylyltransferase subunit 2 [EC:2.7.7.4] |
| ID1591 | 15.67643 | 0.01560019 | 3.00 | 3.00 | 3.00 | 8.00 | 5.75 | 15.50 | 10.00 | K01591 orotidine-5'-phosphate decarboxylase [EC:4.1.1.23] |
| ID164 | 15.65064 | 0.01575718 | 23.67 | 19.50 | 25.00 | 4.00 | 1.25 | 0.75 | 1.50 | K00164 2-oxoglutarate dehydrogenase E1 component [EC:1.2.4.2] |
| ID405 | 15.64638 | 0.01578326 | 4.00 | 5.50 | 9.00 | 1.00 | 0.75 | 0.25 | 0.00 | K00405 cytochrome c oxidase cbb3-type subunit II |
| ID10118 | 15.64000 | 0.01582237 | 1.67 | 1.00 | 1.00 | 3.00 | 4.50 | 53.75 | 13.25 | K10118 raffinose/stachyose/melibiose transport system permease protein |
| ID3584 | 15.63205 | 0.01587128 | 1.67 | 2.50 | 3.00 | 6.00 | 1.75 | 8.75 | 5.00 | K03584 DNA repair protein RecO (recombination protein O) |
| ID850 | 15.62544 | 0.01591200 | 5.00 | 2.50 | 1.00 | 34.00 | 15.50 | 9.00 | 28.75 | K00850 6-phosphofructokinase 1 [EC:2.7.1.11] |
| ID341 | 15.61820 | 0.01595678 | 5.67 | 8.00 | 10.00 | 2.00 | 3.50 | 0.50 | 2.50 | K00341 NADH-quinone oxidoreductase subunit L [EC:1.6.5.3] |
| ID864 | 15.60879 | 0.01601510 | 10.33 | 13.00 | 15.00 | 26.00 | 7.50 | 3.00 | 8.25 | K00864 glycerol kinase [EC:2.7.1.30] |
| ID1737 | 15.60268 | 0.01605312 | 3.33 | 4.00 | 4.00 | 1.00 | 1.00 | 1.00 | 1.50 | K01737 6-pyruvoyltetrahydropterin/6-carboxytetrahydropterin synthase [EC:4.2.3.12 4.1.2.50] |
| ID253 | 15.58978 | 0.01613360 | 3.67 | 6.00 | 9.00 | 1.00 | 0.25 | 0.00 | 0.25 | K00253 isovaleryl-CoA dehydrogenase [EC:1.3.8.4] |
| ID859 | 15.58683 | 0.01615207 | 1.00 | 1.50 | 2.00 | 5.00 | 1.25 | 6.25 | 2.25 | K00859 dephospho-CoA kinase [EC:2.7.1.24] |
| ID2434 | 15.58644 | 0.01615455 | 13.33 | 8.00 | 10.00 | 3.00 | 5.25 | 22.00 | 8.00 | K02434 aspartyl-tRNA(Asn)/glutamyl-tRNA(Gln) amidotransferase subunit B [EC:6.3.5.6 6.3.5.7] |
| ID604 | 15.57059 | 0.01625407 | 3.00 | 4.00 | 5.00 | 10.00 | 4.00 | 10.50 | 7.25 | K00604 methionyl-tRNA formyltransferase [EC:2.1.2.9] |
| ID1520 | 15.56113 | 0.01631380 | 2.00 | 2.50 | 3.00 | 8.00 | 3.00 | 6.50 | 4.50 | K01520 dUTP pyrophosphatase [EC:3.6.1.23] |
| ID849 | 15.55348 | 0.01636227 | 1.67 | 1.00 | 1.00 | 10.00 | 8.75 | 16.50 | 11.50 | K00849 galactokinase [EC:2.7.1.6] |
| ID1814 | 15.55095 | 0.01637829 | 5.33 | 5.00 | 6.00 | 1.00 | 2.50 | 1.25 | 5.00 | K01814 phosphoribosylformimino-5-aminoimidazole carboxamide ribotide isomerase [EC:5.3.1.16] |
| ID2341 | 15.54527 | 0.01641437 | 2.00 | 4.00 | 5.00 | 9.00 | 2.75 | 13.25 | 7.25 | K02341 DNA polymerase III subunit delta' [EC:2.7.7.7] |
| ID2887 | 15.54121 | 0.01644019 | 3.00 | 2.00 | 3.00 | 5.00 | 3.50 | 6.50 | 3.25 | K02887 large subunit ribosomal protein L20 |
| ID854 | 15.53254 | 0.01649553 | 4.67 | 5.00 | 6.00 | 2.00 | 5.25 | 15.00 | 10.25 | K00854 xylulokinase [EC:2.7.1.17] |
| ID1658 | 15.51146 | 0.01663075 | 3.67 | 4.00 | 4.00 | 1.00 | 1.75 | 1.50 | 3.75 | K01658 anthranilate synthase component II [EC:4.1.3.27] |
| ID12251 | 15.49702 | 0.01672401 | 4.67 | 3.00 | 5.00 | 1.00 | 2.75 | 0.50 | 2.75 | K12251 N-carbamoylputrescine amidase [EC:3.5.1.53] |
| ID303 | 15.49180 | 0.01675783 | 1.00 | 20.00 | 28.00 | 4.00 | 1.00 | 0.25 | 1.25 | K00303 sarcosine oxidase |
| ID11959 | 15.48179 | 0.01682290 | 9.33 | 8.50 | 11.00 | 2.00 | 1.00 | 6.00 | 0.50 | K11959 urea transport system substrate-binding protein |
| ID602 | 15.46815 | 0.01691194 | 10.67 | 11.00 | 12.00 | 14.00 | 9.75 | 16.25 | 13.25 | K00602 phosphoribosylaminoimidazolecarboxamide formyltransferase / IMP cyclohydrolase [EC:2.1.2.3 3.5.4.10] |
| ID11752 | 15.45432 | 0.01700270 | 2.00 | 4.50 | 5.00 | 7.00 | 3.50 | 10.00 | 6.75 | K11752 diaminohydroxyphosphoribosylaminopyrimidine deaminase / 5-amino-6-(5-phosphoribosylamino)uracil reductase [EC:3.5.4.26 1.1.1.193] |
| ID161 | 15.45311 | 0.01701068 | 6.33 | 5.00 | 8.00 | 1.00 | 1.00 | 1.25 | 1.25 | K00161 pyruvate dehydrogenase E1 component alpha subunit [EC:1.2.4.1] |
| ID1960 | 15.44569 | 0.01705956 | 5.33 | 10.00 | 14.00 | 2.00 | 3.25 | 0.50 | 2.50 | K01960 pyruvate carboxylase subunit B [EC:6.4.1.1] |
| ID975 | 15.43877 | 0.01710531 | 6.00 | 2.50 | 2.00 | 32.00 | 12.00 | 14.75 | 21.00 | K00975 glucose-1-phosphate adenylyltransferase [EC:2.7.7.27] |
| ID658 | 15.43627 | 0.01712181 | 10.33 | 4.50 | 6.00 | 1.00 | 0.75 | 0.75 | 1.00 | K00658 2-oxoglutarate dehydrogenase E2 component (dihydrolipoamide succinyltransferase) [EC:2.3.1.61] |
| ID5601 | 15.43553 | 0.01712676 | 1.00 | 9.00 | 2.00 | 34.00 | 7.25 | 4.00 | 9.75 | K05601 hydroxylamine reductase [EC:1.7.99.1] |
| ID48 | 15.43267 | 0.01714572 | 1.00 | 1.00 | 1.00 | 6.00 | 5.25 | 12.50 | 9.75 | K00048 lactaldehyde reductase [EC:1.1.1.77] |
| ID11717 | 15.42175 | 0.01721827 | 7.33 | 2.00 | 4.00 | 1.00 | 3.25 | 12.00 | 4.50 | K11717 cysteine desulfurase / selenocysteine lyase [EC:2.8.1.7 4.4.1.16] |
| ID2563 | 15.42054 | 0.01722634 | 2.67 | 4.50 | 6.00 | 13.00 | 5.00 | 12.50 | 8.75 | K02563 UDP-N-acetylglucosamine--N-acetylmuramyl-(pentapeptide) pyrophosphoryl-undecaprenol N-acetylglucosamine transferase [EC:2.4.1.227] |
| ID1938 | 15.41836 | 0.01724085 | 2.67 | 3.50 | 1.00 | 23.00 | 9.50 | 23.25 | 16.25 | K01938 formate--tetrahydrofolate ligase [EC:6.3.4.3] |
| ID1928 | 15.41446 | 0.01726689 | 4.33 | 5.00 | 6.00 | 12.00 | 8.00 | 15.50 | 11.50 | K01928 UDP-N-acetylmuramoyl-L-alanyl-D-glutamate--2 |
| ID14 | 15.40524 | 0.01732851 | 1.67 | 4.00 | 5.00 | 8.00 | 4.50 | 9.25 | 10.50 | K00014 shikimate dehydrogenase [EC:1.1.1.25] |

| | | | | | | | | | | |
|---|---|---|---|---|---|---|---|---|---|---|
| ID3550 | 15.40390 | 0.01733751 | 1.33 | 4.00 | 4.00 | 5.00 | 1.75 | 6.75 | 3.75 | K03550 holliday junction DNA helicase RuvA [EC:3.6.4.12] |
| ID2412 | 15.39397 | 0.01740415 | 8.00 | 7.00 | 7.00 | 3.00 | 2.25 | 3.25 | 1.00 | K02412 flagellum-specific ATP synthase [EC:3.6.3.14] |
| ID1159 | 15.39118 | 0.01742298 | 1.33 | 2.00 | 3.00 | 1.00 | 1.00 | 5.00 | 1.75 | K01159 crossover junction endodeoxyribonuclease RuvC [EC:3.1.22.4] |
| ID13821 | 15.36240 | 0.01761781 | 7.33 | 17.00 | 23.00 | 4.00 | 1.25 | 0.50 | 3.75 | K13821 RHH-type transcriptional regulator |
| ID3470 | 15.34344 | 0.01774730 | 1.67 | 3.00 | 3.00 | 6.00 | 3.00 | 10.25 | 4.50 | K03470 ribonuclease HII [EC:3.1.26.4] |
| ID338 | 15.34155 | 0.01776028 | 5.00 | 5.00 | 5.00 | 1.00 | 1.00 | 0.50 | 1.25 | K00338 NADH-quinone oxidoreductase subunit I [EC:1.6.5.3] |
| ID57 | 15.33356 | 0.01781515 | 3.33 | 5.00 | 6.00 | 9.00 | 5.75 | 16.50 | 8.00 | K00057 glycerol-3-phosphate dehydrogenase (NAD(P)+) [EC:1.1.1.94] |
| ID11358 | 15.32093 | 0.01790221 | 1.00 | 1.00 | 1.00 | 1.00 | 3.50 | 13.75 | 6.75 | K11358 aspartate aminotransferase [EC:2.6.1.1] |
| ID75 | 15.31878 | 0.01791705 | 3.33 | 2.50 | 3.00 | 11.00 | 3.50 | 14.25 | 5.75 | K00075 UDP-N-acetylmuramate dehydrogenase [EC:1.3.1.98] |
| ID2011 | 15.29837 | 0.01805882 | 3.33 | 8.00 | 11.00 | 4.00 | 3.50 | 1.25 | 1.50 | K02011 iron(III) transport system permease protein |
| ID3885 | 15.28381 | 0.01816055 | 7.33 | 4.00 | 5.00 | 2.00 | 1.00 | 2.00 | 1.00 | K03885 NADH dehydrogenase [EC:1.6.99.3] |
| ID432 | 15.28367 | 0.01816152 | 4.00 | 8.50 | 8.00 | 10.00 | 1.00 | 2.50 | 1.25 | K00432 glutathione peroxidase [EC:1.11.1.9] |
| ID575 | 15.27783 | 0.01820245 | 5.00 | 13.00 | 15.00 | 3.00 | 1.50 | 1.75 | 1.00 | K00575 chemotaxis protein methyltransferase CheR [EC:2.1.1.80] |
| ID2757 | 15.27772 | 0.01820322 | 1.00 | 1.00 | 1.00 | 1.00 | 1.75 | 23.00 | 4.50 | K02757 PTS system |
| ID3781 | 15.27588 | 0.01821618 | 46.00 | 26.00 | 33.00 | 7.00 | 1.00 | 1.25 | 2.00 | K03781 catalase [EC:1.11.1.6] |
| ID627 | 15.26901 | 0.01826448 | 10.67 | 9.50 | 13.00 | 2.00 | 1.00 | 1.00 | 1.00 | K00627 pyruvate dehydrogenase E2 component (dihydrolipoamide acetyltransferase) [EC:2.3.1.12] |
| ID791 | 15.26888 | 0.01826542 | 3.00 | 4.00 | 5.00 | 7.00 | 4.00 | 11.25 | 5.50 | K00791 tRNA dimethylallyltransferase [EC:2.5.1.75] |
| ID2049 | 15.26447 | 0.01829654 | 6.00 | 8.50 | 9.00 | 8.00 | 1.00 | 5.75 | 2.25 | K02049 |
| ID1599 | 15.24977 | 0.01840053 | 11.00 | 5.00 | 7.00 | 2.00 | 1.00 | 7.75 | 1.50 | K01599 uroporphyrinogen decarboxylase [EC:4.1.1.37] |
| ID12410 | 15.24643 | 0.01842426 | 1.67 | 2.50 | 3.00 | 11.00 | 3.75 | 13.50 | 5.50 | K12410 |
| ID215 | 15.24642 | 0.01842433 | 3.33 | 4.50 | 6.00 | 9.00 | 3.75 | 13.50 | 6.50 | K00215 4-hydroxy-tetrahydrodipicolinate reductase [EC:1.17.1.8] |
| ID1029 | 15.23559 | 0.01850140 | 4.67 | 4.00 | 5.00 | 1.00 | 0.75 | 0.00 | 0.50 | K01029 3-oxoacid CoA-transferase subunit B [EC:2.8.3.5] |
| ID1886 | 15.23184 | 0.01852816 | 15.67 | 13.50 | 14.00 | 20.00 | 9.75 | 2.75 | 16.75 | K01886 glutaminyl-tRNA synthetase [EC:6.1.1.18] |
| ID3182 | 15.23044 | 0.01853818 | 10.00 | 9.50 | 13.00 | 3.00 | 1.25 | 0.75 | 2.00 | K03182 4-hydroxy-3-polyprenylbenzoate decarboxylase [EC:4.1.1.98] |
| ID3588 | 15.22118 | 0.01860446 | 4.00 | 6.00 | 8.00 | 10.00 | 3.00 | 12.00 | 6.25 | K03588 cell division protein FtsW |
| ID9811 | 15.20874 | 0.01869385 | 1.67 | 4.00 | 5.00 | 7.00 | 2.50 | 10.00 | 6.00 | K09811 cell division transport system permease protein |
| ID3589 | 15.20578 | 0.01871522 | 1.00 | 4.50 | 6.00 | 4.00 | 1.50 | 8.75 | 3.00 | K03589 cell division protein FtsQ |
| ID2409 | 15.19929 | 0.01876208 | 4.33 | 6.50 | 9.00 | 1.00 | 1.75 | 2.25 | 1.00 | K02409 flagellar M-ring protein FliF |
| ID2072 | 15.18696 | 0.01885135 | 1.00 | 4.50 | 6.00 | 1.00 | 1.25 | 13.75 | 2.00 | K02072 D-methionine transport system permease protein |
| ID2301 | 15.18170 | 0.01888960 | 3.67 | 5.50 | 8.00 | 1.00 | 0.75 | 0.50 | 1.00 | K02301 protoheme IX farnesyltransferase [EC:2.5.1.-] |
| ID2340 | 15.17631 | 0.01892886 | 2.33 | 5.00 | 7.00 | 8.00 | 2.75 | 9.75 | 5.25 | K02340 DNA polymerase III subunit delta [EC:2.7.7.7] |
| ID2963 | 15.17308 | 0.01895245 | 3.67 | 2.00 | 2.00 | 5.00 | 2.25 | 5.25 | 2.75 | K02963 small subunit ribosomal protein S18 |
| ID1455 | 15.16368 | 0.01902118 | 7.33 | 7.50 | 10.00 | 1.00 | 1.00 | 0.25 | 0.50 | K01455 formamidase [EC:3.5.1.49] |
| ID2115 | 15.13692 | 0.01921814 | 4.00 | 8.00 | 11.00 | 9.00 | 4.50 | 12.50 | 8.25 | K02115 F-type H+-transporting ATPase subunit gamma |
| ID3621 | 15.13462 | 0.01923516 | 2.33 | 3.50 | 4.00 | 10.00 | 2.75 | 4.75 | 5.25 | K03621 glycerol-3-phosphate acyltransferase PlsX [EC:2.3.1.15] |
| ID1835 | 15.11620 | 0.01937196 | 6.67 | 9.50 | 9.00 | 19.00 | 12.75 | 21.50 | 21.00 | K01835 phosphoglucomutase [EC:5.4.2.2] |
| ID2428 | 15.09673 | 0.01951758 | 3.33 | 3.00 | 4.00 | 5.00 | 3.25 | 14.50 | 6.25 | K02428 XTP/dITP diphosphohydrolase [EC:3.6.1.66] |
| ID525 | 15.09367 | 0.01954057 | 44.00 | 41.50 | 21.00 | 37.00 | 5.00 | 19.50 | 12.50 | K00525 ribonucleoside-diphosphate reductase alpha chain [EC:1.17.4.1] |
| ID1720 | 15.08991 | 0.01956881 | 2.67 | 4.50 | 6.00 | 1.00 | 1.00 | 1.00 | 1.00 | K01720 2-methylcitrate dehydratase [EC:4.2.1.79] |
| ID2535 | 15.08864 | 0.01957836 | 4.67 | 6.50 | 8.00 | 1.00 | 1.25 | 0.75 | 0.75 | K02535 UDP-3-O-[3-hydroxymyristoyl] N-acetylglucosamine deacetylase [EC:3.5.1.108] |
| ID1878 | 15.08506 | 0.01960536 | 9.67 | 7.00 | 9.00 | 2.00 | 1.25 | 2.75 | 1.00 | K01878 glycyl-tRNA synthetase alpha chain [EC:6.1.1.14] |
| ID5349 | 15.06866 | 0.01972935 | 11.33 | 11.50 | 14.00 | 4.00 | 28.50 | 87.25 | 43.25 | K05349 beta-glucosidase [EC:3.2.1.21] |
| ID931 | 15.06356 | 0.01976809 | 3.00 | 4.00 | 6.00 | 11.00 | 4.00 | 15.75 | 7.75 | K00931 glutamate 5-kinase [EC:2.7.2.11] |
| ID2114 | 15.04437 | 0.01991444 | 2.00 | 3.00 | 4.00 | 4.00 | 1.75 | 4.75 | 3.50 | K02114 F-type H+-transporting ATPase subunit epsilon |
| ID1491 | 15.03173 | 0.02001137 | 7.33 | 6.00 | 8.00 | 9.00 | 5.50 | 12.25 | 6.75 | K01491 methylenetetrahydrofolate dehydrogenase (NADP+) / methenyltetrahydrofolate cyclohydrolase [EC:1.5.1.5 3.5.4.9] |
| ID3183 | 15.03028 | 0.02002251 | 6.67 | 6.00 | 7.00 | 1.00 | 1.00 | 1.25 | 1.75 | K03183 demethylmenaquinone methyltransferase / 2-methoxy-6-polyprenyl-1 |
| ID4517 | 15.02584 | 0.02005670 | 1.67 | 1.00 | 1.00 | 1.00 | 2.75 | 8.75 | 4.25 | K04517 prephenate dehydrogenase [EC:1.3.1.12] |
| ID3527 | 15.01784 | 0.02011841 | 5.33 | 6.00 | 9.00 | 8.00 | 5.50 | 12.50 | 9.25 | K03527 4-hydroxy-3-methylbut-2-enyl diphosphate reductase [EC:1.17.1.2] |
| ID973 | 15.01570 | 0.02013499 | 10.67 | 11.00 | 11.00 | 11.00 | 13.25 | 15.75 | 9.50 | K00973 glucose-1-phosphate thymidylyltransferase [EC:2.7.7.24] |
| ID281 | 15.01176 | 0.02016541 | 14.33 | 15.50 | 23.00 | 4.00 | 1.25 | 1.00 | 3.75 | K00281 glycine dehydrogenase [EC:1.4.4.2] |
| ID1740 | 14.99336 | 0.02030845 | 12.00 | 7.00 | 8.00 | 4.00 | 16.00 | 28.25 | 27.50 | K01740 O-acetylhomoserine (thiol)-lyase [EC:2.5.1.49] |
| ID16 | 14.98794 | 0.02035076 | 1.00 | 2.00 | 1.00 | 16.00 | 4.75 | 22.00 | 7.75 | K00016 L-lactate dehydrogenase [EC:1.1.1.27] |
| ID7718 | 14.98569 | 0.02036833 | 1.00 | 1.00 | 1.00 | 12.00 | 3.50 | 3.00 | 10.25 | K07718 two-component system |
| ID1524 | 14.98542 | 0.02037039 | 3.33 | 8.50 | 10.00 | 11.00 | 2.00 | 16.25 | 6.00 | K01524 exopolyphosphatase / guanosine-5'-triphosphate |
| ID833 | 14.95940 | 0.02057484 | 4.67 | 9.50 | 11.00 | 3.00 | 1.25 | 1.00 | 3.00 | K00833 adenosylmethionine-8-amino-7-oxononanoate aminotransferase [EC:2.6.1.62] |
| ID2338 | 14.95420 | 0.02061591 | 6.33 | 10.00 | 13.00 | 21.00 | 4.25 | 22.50 | 10.75 | K02338 DNA polymerase III subunit beta [EC:2.7.7.7] |
| ID527 | 14.94743 | 0.02066958 | 9.00 | 8.00 | 8.00 | 37.00 | 22.25 | 29.25 | 31.00 | K00527 ribonucleoside-triphosphate reductase [EC:1.17.4.2] |
| ID656 | 14.93074 | 0.02080231 | 5.67 | 10.50 | 3.00 | 34.00 | 20.00 | 25.00 | 30.75 | K00656 formate C-acetyltransferase [EC:2.3.1.54] |
| ID954 | 14.92466 | 0.02085087 | 2.67 | 4.00 | 5.00 | 5.00 | 2.00 | 7.75 | 4.25 | K00954 pantetheine-phosphate adenylyltransferase [EC:2.7.7.3] |
| ID1685 | 14.92394 | 0.02085658 | 2.00 | 1.00 | 1.00 | 1.00 | 3.75 | 1.00 | 4.50 | K01685 altronate hydrolase [EC:4.2.1.7] |
| ID4561 | 14.91378 | 0.02093805 | 11.67 | 3.00 | 5.00 | 1.00 | 1.00 | 0.50 | 0.50 | K04561 nitric oxide reductase subunit B [EC:1.7.2.5] |
| ID1182 | 14.87825 | 0.02122508 | 2.33 | 1.00 | 1.00 | 1.00 | 4.00 | 29.00 | 10.50 | K01182 oligo-1 |
| ID798 | 14.86513 | 0.02133203 | 1.33 | 5.00 | 6.00 | 5.00 | 1.00 | 1.00 | 2.00 | K00798 cob(I)alamin adenosyltransferase [EC:2.5.1.17] |
| ID337 | 14.84381 | 0.02150694 | 5.67 | 6.00 | 7.00 | 1.00 | 1.75 | 1.00 | 2.25 | K00337 NADH-quinone oxidoreductase subunit H [EC:1.6.5.3] |
| ID3883 | 14.82280 | 0.02168064 | 4.33 | 19.50 | 4.00 | 1.00 | 0.25 | 0.25 | 0.00 | K03883 NADH-ubiquinone oxidoreductase chain 5 [EC:1.6.5.3] |
| ID2316 | 14.81001 | 0.02178702 | 9.00 | 12.00 | 14.00 | 18.00 | 7.25 | 22.25 | 13.25 | K02316 DNA primase [EC:2.7.7.-] |
| ID2113 | 14.80726 | 0.02180993 | 1.00 | 4.00 | 5.00 | 5.00 | 1.00 | 10.75 | 4.00 | K02113 F-type H+-transporting ATPase subunit delta |
| ID997 | 14.78810 | 0.02197041 | 0.67 | 1.00 | 1.00 | 4.00 | 1.00 | 7.00 | 1.25 | K00997 holo-[acyl-carrier protein] synthase [EC:2.7.8.7] |
| ID4564 | 14.77143 | 0.02211096 | 9.00 | 7.50 | 8.00 | 5.00 | 1.00 | 0.75 | 1.25 | K04564 superoxide dismutase |

| | | | | | | | | | | |
|---|---|---|---|---|---|---|---|---|---|---|
| ID1895 | 14.76729 | 0.02214601 | 27.67 | 26.00 | 37.00 | 6.00 | 6.25 | 2.25 | 8.25 | K01895 acetyl-CoA synthetase [EC:6.2.1.1] |
| ID1845 | 14.75789 | 0.02222572 | 12.33 | 12.00 | 15.00 | 13.00 | 2.00 | 1.50 | 4.75 | K01845 glutamate-1-semialdehyde 2 |
| ID11903 | 14.72768 | 0.02248386 | 3.67 | 4.50 | 6.00 | 1.00 | 1.00 | 0.50 | 0.50 | K11903 type VI secretion system secreted protein Hcp |
| ID41 | 14.72687 | 0.02249081 | 1.00 | 1.00 | 1.00 | 1.00 | 4.50 | 1.00 | 6.25 | K00041 tagaturonate reductase [EC:1.1.1.58] |
| ID12574 | 14.70372 | 0.02269061 | 10.33 | 2.00 | 1.00 | 18.00 | 4.50 | 17.25 | 6.75 | K12574 ribonuclease J [EC:3.1.-.-] |
| ID819 | 14.70157 | 0.02270924 | 5.00 | 1.00 | 1.00 | 1.00 | 1.00 | 0.00 | 1.25 | K00819 ornithine--oxo-acid transaminase [EC:2.6.1.13] |
| ID655 | 14.69867 | 0.02273444 | 4.00 | 8.00 | 10.00 | 14.00 | 3.75 | 7.75 | 7.25 | K00655 1-acyl-sn-glycerol-3-phosphate acyltransferase [EC:2.3.1.51] |
| ID800 | 14.69367 | 0.02277786 | 3.33 | 6.50 | 8.00 | 8.00 | 3.75 | 13.25 | 7.25 | K00800 3-phosphoshikimate 1-carboxyvinyltransferase [EC:2.5.1.19] |
| ID951 | 14.67544 | 0.02293695 | 19.00 | 16.50 | 19.00 | 23.00 | 13.25 | 27.75 | 24.00 | K00951 GTP pyrophosphokinase [EC:2.7.6.5] |
| ID1191 | 14.67336 | 0.02295515 | 1.00 | 2.00 | 2.00 | 41.00 | 1.25 | 39.00 | 7.75 | K01191 alpha-mannosidase [EC:3.2.1.24] |
| ID2108 | 14.67049 | 0.02298040 | 5.00 | 7.00 | 10.00 | 8.00 | 4.50 | 14.25 | 8.25 | K02108 F-type H+-transporting ATPase subunit a |
| ID2040 | 14.66566 | 0.02302274 | 5.00 | 7.00 | 8.00 | 5.00 | 4.25 | 15.50 | 7.75 | K02040 phosphate transport system substrate-binding protein |
| ID762 | 14.66168 | 0.02305772 | 3.00 | 5.00 | 7.00 | 8.00 | 5.50 | 9.00 | 8.00 | K00762 orotate phosphoribosyltransferase [EC:2.4.2.10] |
| ID5847 | 14.66124 | 0.02306161 | 2.33 | 10.00 | 8.00 | 9.00 | 1.00 | 0.75 | 1.00 | K05847 osmoprotectant transport system ATP-binding protein |
| ID2892 | 14.65642 | 0.02310406 | 2.00 | 3.00 | 4.00 | 4.00 | 2.75 | 6.25 | 3.25 | K02892 large subunit ribosomal protein L23 |
| ID3856 | 14.65338 | 0.02313093 | 1.00 | 2.00 | 1.00 | 9.00 | 4.00 | 3.25 | 6.00 | K03856 3-deoxy-7-phosphoheptulonate synthase [EC:2.5.1.54] |
| ID77 | 14.65200 | 0.02314310 | 1.00 | 4.00 | 5.00 | 9.00 | 3.00 | 6.00 | 7.75 | K00077 2-dehydropantoate 2-reductase [EC:1.1.1.169] |
| ID7456 | 14.64397 | 0.02321415 | 3.67 | 2.00 | 1.00 | 18.00 | 8.50 | 7.00 | 15.50 | K07456 DNA mismatch repair protein MutS2 |
| ID1585 | 14.64016 | 0.02324788 | 10.00 | 11.00 | 13.00 | 3.00 | 4.25 | 1.00 | 5.00 | K01585 arginine decarboxylase [EC:4.1.1.19] |
| ID123 | 14.61727 | 0.02345174 | 21.33 | 8.00 | 10.00 | 4.00 | 2.00 | 1.50 | 1.25 | K00123 formate dehydrogenase major subunit [EC:1.2.1.2] |
| ID11069 | 14.61574 | 0.02346548 | 1.00 | 1.00 | 1.00 | 10.00 | 4.00 | 5.25 | 7.25 | K11069 spermidine/putrescine transport system substrate-binding protein |
| ID12368 | 14.59067 | 0.02369082 | 1.00 | 21.00 | 27.00 | 4.00 | 1.25 | 0.50 | 1.00 | K12368 dipeptide transport system substrate-binding protein |
| ID645 | 14.58943 | 0.02370201 | 2.00 | 4.00 | 5.00 | 8.00 | 4.75 | 6.00 | 6.75 | K00645 [acyl-carrier-protein] S-malonyltransferase [EC:2.3.1.39] |
| ID2016 | 14.57394 | 0.02384235 | 5.67 | 3.50 | 3.00 | 23.00 | 4.50 | 5.75 | 13.75 | K02016 iron complex transport system substrate-binding protein |
| ID1817 | 14.56981 | 0.02387989 | 1.33 | 3.00 | 2.00 | 1.00 | 1.00 | 10.00 | 2.75 | K01817 phosphoribosylanthranilate isomerase [EC:5.3.1.24] |
| ID3629 | 14.55882 | 0.02398007 | 3.67 | 9.00 | 11.00 | 16.00 | 3.50 | 17.25 | 6.50 | K03629 DNA replication and repair protein RecF |
| ID3 | 14.55824 | 0.02398542 | 5.33 | 9.00 | 11.00 | 3.00 | 4.00 | 17.25 | 11.25 | K00003 homoserine dehydrogenase [EC:1.1.1.3] |
| ID1998 | 14.55176 | 0.02404472 | 3.67 | 13.00 | 18.00 | 3.00 | 2.50 | 23.75 | 3.75 | K01998 branched-chain amino acid transport system permease protein |
| ID1007 | 14.55028 | 0.02405819 | 11.33 | 15.00 | 18.00 | 4.00 | 1.25 | 1.25 | 1.25 | K01007 pyruvate |
| ID2342 | 14.54409 | 0.02411501 | 7.67 | 8.50 | 11.00 | 11.00 | 2.00 | 9.25 | 3.50 | K02342 DNA polymerase III subunit epsilon [EC:2.7.7.7] |
| ID2416 | 14.52228 | 0.02431609 | 6.33 | 6.50 | 7.00 | 2.00 | 1.75 | 2.50 | 1.00 | K02416 flagellar motor switch protein FliM |
| ID925 | 14.51838 | 0.02435226 | 4.33 | 2.00 | 2.00 | 21.00 | 9.50 | 15.50 | 12.00 | K00925 acetate kinase [EC:2.7.2.1] |
| ID228 | 14.51827 | 0.02435323 | 5.33 | 5.00 | 7.00 | 1.00 | 0.75 | 0.25 | 0.75 | K00228 coproporphyrinogen III oxidase [EC:1.3.3.3] |
| ID3111 | 14.51622 | 0.02437232 | 5.67 | 5.00 | 4.00 | 12.00 | 3.50 | 18.00 | 7.00 | K03111 single-strand DNA-binding protein |
| ID135 | 14.51302 | 0.02440203 | 8.67 | 14.00 | 18.00 | 3.00 | 1.00 | 2.25 | 0.75 | K00135 succinate-semialdehyde dehydrogenase / glutarate-semialdehyde dehydrogenase [EC:1.2.1.16 1.2.1.79 1.2.1.20] |
| ID919 | 14.50796 | 0.02444902 | 2.67 | 6.00 | 8.00 | 9.00 | 2.75 | 15.50 | 5.50 | K00919 4-diphosphocytidyl-2-C-methyl-D-erythritol kinase [EC:2.7.1.148] |
| ID6001 | 14.50683 | 0.02445954 | 2.67 | 1.00 | 1.00 | 1.00 | 4.75 | 1.00 | 4.50 | K06001 tryptophan synthase beta chain [EC:4.2.1.20] |
| ID3763 | 14.47705 | 0.02473825 | 9.00 | 4.00 | 2.00 | 35.00 | 12.50 | 7.25 | 22.00 | K03763 DNA polymerase III subunit alpha |
| ID1193 | 14.46839 | 0.02481991 | 1.67 | 1.00 | 1.00 | 10.00 | 4.50 | 12.75 | 11.50 | K01193 beta-fructofuranosidase [EC:3.2.1.26] |
| ID13789 | 14.46711 | 0.02483202 | 1.67 | 3.00 | 4.00 | 7.00 | 3.25 | 2.25 | 3.50 | K13789 geranylgeranyl diphosphate synthase |
| ID1778 | 14.46234 | 0.02487705 | 3.33 | 5.50 | 6.00 | 7.00 | 3.25 | 10.00 | 5.25 | K01778 diaminopimelate epimerase [EC:5.1.1.7] |
| ID1808 | 14.45722 | 0.02492556 | 2.00 | 2.00 | 1.00 | 5.00 | 3.00 | 2.25 | 4.00 | K01808 ribose 5-phosphate isomerase B [EC:5.3.1.6] |
| ID1647 | 14.45358 | 0.02496006 | 15.67 | 9.00 | 10.00 | 3.00 | 7.75 | 16.75 | 23.00 | K01647 citrate synthase [EC:2.3.3.1] |
| ID13378 | 14.44467 | 0.02504474 | 6.00 | 8.00 | 11.00 | 2.00 | 1.75 | 0.50 | 2.50 | K13378 NADH-quinone oxidoreductase subunit C/D [EC:1.6.5.3] |
| ID1206 | 14.44381 | 0.02505299 | 2.00 | 1.00 | 1.00 | 17.00 | 4.75 | 30.00 | 12.25 | K01206 alpha-L-fucosidase [EC:3.2.1.51] |
| ID2232 | 14.41984 | 0.02528226 | 1.33 | 6.50 | 7.00 | 9.00 | 3.25 | 2.25 | 4.25 | K02232 adenosylcobyric acid synthase [EC:6.3.5.10] |
| ID7699 | 14.40322 | 0.02544239 | 1.33 | 3.00 | 1.00 | 11.00 | 2.00 | 0.75 | 4.75 | K07699 two-component system |
| ID2117 | 14.39542 | 0.02551784 | 1.67 | 2.00 | 1.00 | 15.00 | 6.75 | 4.00 | 13.50 | K02117 V/A-type H+-transporting ATPase subunit A [EC:3.6.3.14] |
| ID2400 | 14.39334 | 0.02553800 | 8.33 | 9.00 | 11.00 | 3.00 | 3.50 | 2.25 | 1.25 | K02400 flagellar biosynthesis protein FlhA |
| ID3217 | 14.39299 | 0.02554146 | 6.67 | 8.50 | 11.00 | 10.00 | 5.25 | 20.00 | 9.75 | K03217 YidC/Oxa1 family membrane protein insertase |
| ID1613 | 14.38505 | 0.02561850 | 3.67 | 6.00 | 7.00 | 12.00 | 1.75 | 1.25 | 2.50 | K01613 phosphatidylserine decarboxylase [EC:4.1.1.65] |
| ID8 | 14.37677 | 0.02569918 | 1.00 | 1.00 | 1.00 | 1.00 | 1.50 | 0.75 | 7.25 | K00008 L-iditol 2-dehydrogenase [EC:1.1.1.14] |
| ID1681 | 14.37103 | 0.02575527 | 36.67 | 24.50 | 31.00 | 7.00 | 11.50 | 20.00 | 19.75 | K01681 aconitate hydratase [EC:4.2.1.3] |
| ID111 | 14.36429 | 0.02582124 | 5.33 | 8.00 | 8.00 | 20.00 | 4.00 | 1.50 | 4.75 | K00111 glycerol-3-phosphate dehydrogenase [EC:1.1.5.3] |
| ID394 | 14.36155 | 0.02584811 | 0.33 | 1.00 | 1.00 | 1.00 | 2.75 | 1.00 | 6.75 | K00394 adenylylsulfate reductase |
| ID1443 | 14.35676 | 0.02589515 | 1.33 | 2.00 | 2.00 | 12.00 | 5.00 | 15.25 | 7.75 | K01443 N-acetylglucosamine-6-phosphate deacetylase [EC:3.5.1.25] |
| ID1428 | 14.35345 | 0.02592774 | 4.00 | 8.50 | 11.00 | 2.00 | 1.00 | 6.75 | 1.00 | K01428 urease subunit alpha [EC:3.5.1.5] |
| ID7407 | 14.35239 | 0.02593812 | 2.33 | 2.00 | 1.00 | 19.00 | 12.50 | 44.25 | 26.00 | K07407 alpha-galactosidase [EC:3.2.1.22] |
| ID1903 | 14.32258 | 0.02623316 | 12.00 | 7.50 | 10.00 | 2.00 | 1.00 | 7.25 | 2.00 | K01903 succinyl-CoA synthetase beta subunit [EC:6.2.1.5] |
| ID371 | 14.32115 | 0.02624744 | 8.33 | 6.50 | 7.00 | 1.00 | 1.00 | 0.50 | 1.00 | K00371 nitrate reductase beta subunit [EC:1.7.99.4] |
| ID681 | 14.31133 | 0.02634533 | 6.00 | 12.00 | 14.00 | 2.00 | 1.25 | 1.50 | 1.25 | K00681 gamma-glutamyltranspeptidase / glutathione hydrolase [EC:2.3.2.2 3.4.19.13] |
| ID7516 | 14.30769 | 0.02638169 | 7.00 | 1.00 | 1.00 | 1.00 | 0.25 | 0.00 | 0.25 | K07516 3-hydroxyacyl-CoA dehydrogenase [EC:1.1.1.35] |
| ID11071 | 14.30579 | 0.02640078 | 1.00 | 1.00 | 1.00 | 7.00 | 3.25 | 3.25 | 5.00 | K11071 spermidine/putrescine transport system permease protein |
| ID1897 | 14.27181 | 0.02674300 | 20.33 | 18.00 | 22.00 | 4.00 | 11.25 | 65.75 | 18.75 | K01897 long-chain acyl-CoA synthetase [EC:6.2.1.3] |
| ID949 | 14.24186 | 0.02704806 | 0.67 | 1.00 | 1.00 | 4.00 | 1.25 | 8.50 | 2.75 | K00949 thiamine pyrophosphokinase [EC:2.7.6.2] |
| ID1610 | 14.23940 | 0.02707332 | 13.67 | 10.50 | 14.00 | 3.00 | 16.50 | 4.00 | 17.75 | K01610 phosphoenolpyruvate carboxykinase (ATP) [EC:4.1.1.49] |
| ID2556 | 14.23893 | 0.02707813 | 5.00 | 8.00 | 11.00 | 2.00 | 1.50 | 1.75 | 1.00 | K02556 chemotaxis protein MotA |
| ID3531 | 14.22770 | 0.02719356 | 7.00 | 7.00 | 9.00 | 11.00 | 5.75 | 16.00 | 9.75 | K03531 cell division protein FtsZ |
| ID10536 | 14.21001 | 0.02737623 | 2.67 | 6.00 | 7.00 | 2.00 | 1.25 | 0.75 | 2.50 | K10536 agmatine deiminase [EC:3.5.3.12] |

| | | | | | | | | | | |
|---|---|---|---|---|---|---|---|---|---|---|
| ID2051 | 14.20935 | 0.02738306 | 7.33 | 8.00 | 9.00 | 11.00 | 2.75 | 2.75 | 4.25 | K02051 |
| ID13747 | 14.19361 | 0.02754660 | 4.67 | 2.50 | 3.00 | 1.00 | 4.00 | 1.00 | 5.50 | K13747 carboxynorspermidine decarboxylase [EC:4.1.1.96] |
| ID1962 | 14.17479 | 0.02774340 | 7.33 | 7.50 | 8.00 | 7.00 | 1.25 | 2.25 | 3.50 | K01962 acetyl-CoA carboxylase carboxyl transferase subunit alpha [EC:6.4.1.2] |
| ID12573 | 14.14513 | 0.02805631 | 10.67 | 16.50 | 22.00 | 19.00 | 7.75 | 6.25 | 11.00 | K12573 ribonuclease R [EC:3.1.-.-] |
| ID36 | 14.13876 | 0.02812394 | 12.00 | 17.00 | 21.00 | 4.00 | 0.75 | 11.00 | 2.00 | K00036 glucose-6-phosphate 1-dehydrogenase [EC:1.1.1.49] |
| ID2410 | 14.12919 | 0.02822587 | 4.67 | 8.50 | 9.00 | 3.00 | 2.25 | 2.00 | 1.00 | K02410 flagellar motor switch protein FliG |
| ID4041 | 14.12856 | 0.02823260 | 2.00 | 2.50 | 1.00 | 4.00 | 7.25 | 2.75 | 11.25 | K04041 fructose-1 |
| ID768 | 14.06303 | 0.02893994 | 1.00 | 3.50 | 4.00 | 8.00 | 1.50 | 1.00 | 2.75 | K00768 nicotinate-nucleotide--dimethylbenzimidazole phosphoribosyltransferase [EC:2.4.2.21] |
| ID3074 | 14.05742 | 0.02900129 | 1.00 | 3.50 | 5.00 | 7.00 | 1.25 | 1.25 | 1.00 | K03074 preprotein translocase subunit SecF |
| ID1486 | 14.05479 | 0.02903003 | 2.00 | 1.00 | 1.00 | 2.00 | 4.50 | 3.50 | 8.00 | K01486 adenine deaminase [EC:3.5.4.2] |
| ID1091 | 14.04527 | 0.02913454 | 3.00 | 7.00 | 10.00 | 2.00 | 4.50 | 9.25 | 9.50 | K01091 phosphoglycolate phosphatase [EC:3.1.3.18] |
| ID2050 | 14.03954 | 0.02919754 | 4.00 | 7.50 | 8.00 | 8.00 | 1.50 | 6.50 | 2.50 | K02050 |
| ID761 | 14.03128 | 0.02928870 | 4.33 | 4.00 | 5.00 | 8.00 | 4.25 | 12.75 | 6.50 | K00761 uracil phosphoribosyltransferase [EC:2.4.2.9] |
| ID29 | 14.03028 | 0.02929972 | 8.00 | 8.00 | 10.00 | 2.00 | 2.50 | 1.00 | 3.75 | K00029 malate dehydrogenase (oxaloacetate-decarboxylating)(NADP+) [EC:1.1.1.40] |
| ID13038 | 14.01542 | 0.02946444 | 3.33 | 6.00 | 8.00 | 11.00 | 4.50 | 13.00 | 6.25 | K13038 phosphopantothenoylcysteine decarboxylase / phosphopantothenate--cysteine ligase [EC:4.1.1.36 6.3.2.5] |
| ID937 | 14.00902 | 0.02953554 | 11.00 | 11.50 | 16.00 | 4.00 | 6.50 | 30.00 | 21.50 | K00937 polyphosphate kinase [EC:2.7.4.1] |
| ID10546 | 13.99355 | 0.02970832 | 1.00 | 1.00 | 1.00 | 1.00 | 1.00 | 5.50 | 2.25 | K10546 putative multiple sugar transport system substrate-binding protein |
| ID836 | 13.97391 | 0.02992898 | 6.00 | 2.00 | 2.00 | 1.00 | 1.25 | 0.25 | 1.00 | K00836 diaminobutyrate-2-oxoglutarate transaminase [EC:2.6.1.76] |
| ID2502 | 13.96970 | 0.02997650 | 2.33 | 5.50 | 8.00 | 1.00 | 1.50 | 2.25 | 4.00 | K02502 ATP phosphoribosyltransferase regulatory subunit |
| ID878 | 13.96677 | 0.03000953 | 0.67 | 1.00 | 1.00 | 5.00 | 2.00 | 9.25 | 3.50 | K00878 hydroxyethylthiazole kinase [EC:2.7.1.50] |
| ID605 | 13.96522 | 0.03002715 | 5.33 | 20.00 | 26.00 | 4.00 | 2.50 | 1.00 | 3.25 | K00605 aminomethyltransferase [EC:2.1.2.10] |
| ID3732 | 13.95915 | 0.03009578 | 5.00 | 5.50 | 7.00 | 1.00 | 1.25 | 0.75 | 1.00 | K03732 ATP-dependent RNA helicase RhlB [EC:3.6.4.13] |
| ID1749 | 13.90929 | 0.03066600 | 4.00 | 4.00 | 5.00 | 7.00 | 1.25 | 1.25 | 2.00 | K01749 hydroxymethylbilane synthase [EC:2.5.1.61] |
| ID1081 | 13.90417 | 0.03072519 | 2.00 | 4.50 | 6.00 | 1.00 | 1.00 | 5.00 | 2.25 | K01081 5'-nucleotidase [EC:3.1.3.5] |
| ID2037 | 13.89671 | 0.03081155 | 5.33 | 9.00 | 12.00 | 5.00 | 3.00 | 12.50 | 8.00 | K02037 phosphate transport system permease protein |
| ID162 | 13.88462 | 0.03095197 | 9.00 | 6.00 | 9.00 | 1.00 | 1.00 | 1.25 | 1.25 | K00162 pyruvate dehydrogenase E1 component beta subunit [EC:1.2.4.1] |
| ID1840 | 13.85991 | 0.03124083 | 6.00 | 1.00 | 1.00 | 1.00 | 2.50 | 1.00 | 2.50 | K01840 phosphomannomutase [EC:5.4.2.8] |
| ID1581 | 13.83929 | 0.03148387 | 5.33 | 11.50 | 16.00 | 3.00 | 1.75 | 2.00 | 1.25 | K01581 ornithine decarboxylase [EC:4.1.1.17] |
| ID697 | 13.83630 | 0.03151922 | 5.33 | 1.00 | 1.00 | 1.00 | 0.75 | 0.25 | 1.00 | K00697 trehalose 6-phosphate synthase [EC:2.4.1.15] |
| ID1739 | 13.83058 | 0.03158703 | 2.67 | 3.00 | 4.00 | 1.00 | 1.25 | 7.75 | 2.00 | K01739 cystathionine gamma-synthase [EC:2.5.1.48] |
| ID1912 | 13.82414 | 0.03166350 | 4.67 | 1.00 | 1.00 | 1.00 | 5.75 | 1.00 | 6.50 | K01912 phenylacetate-CoA ligase [EC:6.2.1.30] |
| ID6131 | 13.80346 | 0.03191035 | 4.33 | 7.50 | 8.00 | 43.00 | 8.50 | 5.50 | 20.00 | K06131 cardiolipin synthase [EC:2.7.8.-] |
| ID11927 | 13.80206 | 0.03192709 | 4.67 | 8.50 | 11.00 | 2.00 | 1.25 | 1.75 | 2.75 | K11927 ATP-dependent RNA helicase RhlE [EC:3.6.4.13] |
| ID1966 | 13.77465 | 0.03225731 | 8.67 | 1.00 | 1.00 | 1.00 | 1.00 | 0.50 | 2.50 | K01966 propionyl-CoA carboxylase beta chain [EC:6.4.1.3] |
| ID2013 | 13.77279 | 0.03227984 | 8.33 | 4.00 | 4.00 | 19.00 | 4.25 | 9.75 | 9.25 | K02013 iron complex transport system ATP-binding protein [EC:3.6.3.34] |
| ID1439 | 13.76928 | 0.03232232 | 6.33 | 5.00 | 7.00 | 10.00 | 3.00 | 13.25 | 7.50 | K01439 succinyl-diaminopimelate desuccinylase [EC:3.5.1.18] |
| ID876 | 13.75997 | 0.03243539 | 2.00 | 2.00 | 1.00 | 14.00 | 8.50 | 5.75 | 10.00 | K00876 uridine kinase [EC:2.7.1.48] |
| ID426 | 13.75144 | 0.03253936 | 3.33 | 9.00 | 10.00 | 2.00 | 1.25 | 1.25 | 1.75 | K00426 cytochrome d ubiquinol oxidase subunit II [EC:1.10.3.-] |
| ID2959 | 13.74670 | 0.03259726 | 3.00 | 2.00 | 2.00 | 3.00 | 2.00 | 7.25 | 2.75 | K02959 small subunit ribosomal protein S16 |
| ID1488 | 13.72038 | 0.03292048 | 5.00 | 6.00 | 6.00 | 12.00 | 1.75 | 1.50 | 1.00 | K01488 adenosine deaminase [EC:3.5.4.4] |
| ID1006 | 13.70671 | 0.03308954 | 4.33 | 3.50 | 1.00 | 28.00 | 28.25 | 6.50 | 29.50 | K01006 pyruvate |
| ID817 | 13.70283 | 0.03313773 | 6.33 | 8.00 | 11.00 | 2.00 | 4.50 | 13.25 | 7.50 | K00817 histidinol-phosphate aminotransferase [EC:2.6.1.9] |
| ID1958 | 13.69981 | 0.03317518 | 10.33 | 5.50 | 3.00 | 4.00 | 1.00 | 4.00 | 2.00 | K01958 pyruvate carboxylase [EC:6.4.1.1] |
| ID2913 | 13.69029 | 0.03329380 | 3.67 | 2.00 | 2.00 | 2.00 | 1.75 | 7.50 | 2.00 | K02913 large subunit ribosomal protein L33 |
| ID11070 | 13.68956 | 0.03330287 | 1.00 | 1.50 | 1.00 | 6.00 | 4.50 | 5.00 | 6.00 | K11070 spermidine/putrescine transport system permease protein |
| ID1698 | 13.68750 | 0.03332858 | 6.00 | 8.00 | 11.00 | 9.00 | 1.75 | 1.75 | 4.75 | K01698 porphobilinogen synthase [EC:4.2.1.24] |
| ID11103 | 13.67442 | 0.03349225 | 1.00 | 6.00 | 8.00 | 1.00 | 1.25 | 0.50 | 1.00 | K11103 aerobic C4-dicarboxylate transport protein |
| ID2895 | 13.66344 | 0.03363016 | 2.33 | 2.50 | 3.00 | 4.00 | 2.25 | 6.00 | 3.75 | K02895 large subunit ribosomal protein L24 |
| ID1734 | 13.65660 | 0.03371641 | 1.33 | 1.00 | 1.00 | 4.00 | 2.50 | 1.25 | 4.25 | K01734 methylglyoxal synthase [EC:4.2.3.3] |
| ID1914 | 13.65562 | 0.03372880 | 1.00 | 1.00 | 1.00 | 9.00 | 5.75 | 2.50 | 5.75 | K01914 aspartate--ammonia ligase [EC:6.3.1.1] |
| ID756 | 13.65115 | 0.03378525 | 1.33 | 2.00 | 1.00 | 11.00 | 2.25 | 7.00 | 2.00 | K00756 pyrimidine-nucleoside phosphorylase [EC:2.4.2.2] |
| ID1270 | 13.64701 | 0.03383772 | 1.67 | 2.00 | 1.00 | 31.00 | 5.00 | 2.00 | 7.00 | K01270 dipeptidase D [EC:3.4.13.-] |
| ID1714 | 13.62522 | 0.03411461 | 5.00 | 8.00 | 10.00 | 10.00 | 6.00 | 28.25 | 9.50 | K01714 4-hydroxy-tetrahydrodipicolinate synthase [EC:4.3.3.7] |
| ID1572 | 13.62280 | 0.03414551 | 1.33 | 1.00 | 0.00 | 1.00 | 9.50 | 3.75 | 12.50 | K01572 oxaloacetate decarboxylase |
| ID3581 | 13.61190 | 0.03428503 | 3.33 | 12.00 | 13.00 | 24.00 | 4.50 | 8.75 | 12.00 | K03581 exodeoxyribonuclease V alpha subunit [EC:3.1.11.5] |
| ID1999 | 13.59458 | 0.03450774 | 8.00 | 18.00 | 24.00 | 5.00 | 4.00 | 24.75 | 5.75 | K01999 branched-chain amino acid transport system substrate-binding protein |
| ID526 | 13.59143 | 0.03454842 | 17.33 | 21.00 | 16.00 | 15.00 | 2.00 | 13.50 | 5.50 | K00526 ribonucleoside-diphosphate reductase beta chain [EC:1.17.4.1] |
| ID3076 | 13.58459 | 0.03463686 | 13.33 | 11.00 | 15.00 | 16.00 | 6.50 | 21.00 | 8.75 | K03076 preprotein translocase subunit SecY |
| ID1925 | 13.57063 | 0.03481807 | 5.00 | 6.00 | 7.00 | 17.00 | 6.25 | 14.50 | 12.00 | K01925 UDP-N-acetylmuramoylalanine--D-glutamate ligase [EC:6.3.2.9] |
| ID868 | 13.56959 | 0.03483151 | 2.33 | 3.50 | 3.00 | 8.00 | 2.50 | 10.25 | 5.25 | K00868 pyridoxine kinase [EC:2.7.1.35] |
| ID1733 | 13.56519 | 0.03488890 | 12.33 | 12.50 | 14.00 | 3.00 | 7.75 | 16.50 | 11.50 | K01733 threonine synthase [EC:4.2.3.1] |
| ID874 | 13.56278 | 0.03492033 | 1.33 | 3.00 | 2.00 | 8.00 | 9.00 | 4.50 | 9.00 | K00874 2-dehydro-3-deoxygluconokinase [EC:2.7.1.45] |
| ID1879 | 13.55042 | 0.03508189 | 4.00 | 10.00 | 14.00 | 3.00 | 1.50 | 3.75 | 1.25 | K01879 glycyl-tRNA synthetase beta chain [EC:6.1.1.14] |
| ID981 | 13.54163 | 0.03519726 | 3.67 | 6.00 | 6.00 | 7.00 | 3.25 | 11.00 | 5.50 | K00981 phosphatidate cytidylyltransferase [EC:2.7.7.41] |
| ID6148 | 13.53900 | 0.03523183 | 1.67 | 5.50 | 7.00 | 34.00 | 1.00 | 1.25 | 2.00 | K06148 ATP-binding cassette |
| ID941 | 13.53520 | 0.03528187 | 1.67 | 3.50 | 5.00 | 5.00 | 2.75 | 9.50 | 5.50 | K00941 hydroxymethylpyrimidine/phosphomethylpyrimidine kinase [EC:2.7.1.49 2.7.4.7] |
| ID2931 | 13.52851 | 0.03537012 | 8.67 | 7.50 | 9.00 | 10.00 | 6.00 | 10.75 | 6.00 | K02931 large subunit ribosomal protein L5 |
| ID1547 | 13.52436 | 0.03542491 | 1.00 | 8.50 | 9.00 | 3.00 | 1.00 | 0.75 | 4.00 | K01547 K+-transporting ATPase ATPase B chain [EC:3.6.3.12] |
| ID848 | 13.51868 | 0.03550009 | 1.00 | 1.50 | 1.00 | 3.00 | 3.00 | 2.25 | 5.25 | K00848 rhamnulokinase [EC:2.7.1.5] |

| | | | | | | | | | | |
|---|---|---|---|---|---|---|---|---|---|---|
| ID3092 | 13.49663 | 0.03579338 | 5.00 | 9.00 | 12.00 | 11.00 | 2.00 | 4.25 | 3.50 | K03092 RNA polymerase sigma-54 factor |
| ID688 | 13.49604 | 0.03580129 | 7.00 | 18.50 | 22.00 | 53.00 | 23.75 | 37.75 | 43.75 | K00688 starch phosphorylase [EC:2.4.1.1] |
| ID40 | 13.48788 | 0.03591039 | 2.00 | 1.50 | 1.00 | 1.00 | 5.00 | 4.50 | 5.25 | K00040 fructuronate reductase [EC:1.1.1.57] |
| ID806 | 13.48037 | 0.03601113 | 5.00 | 6.50 | 7.00 | 14.00 | 4.25 | 10.25 | 6.25 | K00806 undecaprenyl diphosphate synthase [EC:2.5.1.31] |
| ID548 | 13.47673 | 0.03606006 | 21.00 | 20.50 | 24.00 | 11.00 | 9.50 | 4.75 | 9.50 | K00548 5-methyltetrahydrofolate--homocysteine methyltransferase [EC:2.1.1.13] |
| ID11961 | 13.47629 | 0.03606592 | 4.00 | 4.00 | 6.00 | 1.00 | 1.00 | 4.50 | 0.50 | K11961 urea transport system permease protein |
| ID2557 | 13.47406 | 0.03609592 | 3.00 | 6.00 | 7.00 | 1.00 | 1.75 | 2.25 | 1.00 | K02557 chemotaxis protein MotB |
| ID1945 | 13.47038 | 0.03614557 | 8.00 | 7.50 | 8.00 | 11.00 | 8.25 | 14.50 | 13.25 | K01945 phosphoribosylamine--glycine ligase [EC:6.3.4.13] |
| ID4072 | 13.46872 | 0.03616787 | 1.33 | 3.00 | 1.00 | 34.00 | 7.00 | 29.00 | 14.25 | K04072 acetaldehyde dehydrogenase / alcohol dehydrogenase [EC:1.2.1.10 1.1.1.1] |
| ID1921 | 13.45371 | 0.03637082 | 5.00 | 6.50 | 8.00 | 9.00 | 5.25 | 18.25 | 9.25 | K01921 D-alanine-D-alanine ligase [EC:6.3.2.4] |
| ID2810 | 13.43756 | 0.03659034 | 1.00 | 1.50 | 1.00 | 7.00 | 1.00 | 5.25 | 4.75 | K02810 PTS system |
| ID1703 | 13.42077 | 0.03681990 | 14.67 | 6.50 | 8.00 | 2.00 | 9.50 | 17.75 | 16.75 | K01703 3-isopropylmalate/(R)-2-methylmalate dehydratase large subunit [EC:4.2.1.33 4.2.1.35] |
| ID1918 | 13.40642 | 0.03701719 | 2.33 | 2.50 | 4.00 | 1.00 | 2.50 | 1.25 | 4.75 | K01918 pantoate--beta-alanine ligase [EC:6.3.2.1] |
| ID5846 | 13.39548 | 0.03716827 | 1.00 | 7.50 | 7.00 | 13.00 | 1.00 | 1.50 | 1.00 | K05846 osmoprotectant transport system permease protein |
| ID3657 | 13.38588 | 0.03730132 | 19.67 | 17.50 | 18.00 | 86.00 | 17.50 | 77.25 | 33.00 | K03657 DNA helicase II / ATP-dependent DNA helicase PcrA [EC:3.6.4.12] |
| ID3783 | 13.37379 | 0.03746956 | 2.00 | 2.00 | 1.00 | 13.00 | 3.75 | 2.00 | 3.75 | K03783 purine-nucleoside phosphorylase [EC:2.4.2.1] |
| ID2879 | 13.36446 | 0.03759989 | 3.00 | 3.00 | 4.00 | 6.00 | 3.25 | 7.50 | 4.25 | K02879 large subunit ribosomal protein L17 |
| ID1130 | 13.36068 | 0.03765275 | 1.33 | 5.00 | 7.00 | 1.00 | 1.25 | 0.25 | 1.50 | K01130 arylsulfatase [EC:3.1.6.1] |
| ID10440 | 13.35817 | 0.03768799 | 1.00 | 4.50 | 6.00 | 6.00 | 1.50 | 1.75 | 2.25 | K10440 ribose transport system permease protein |
| ID425 | 13.33713 | 0.03798394 | 9.33 | 19.00 | 22.00 | 4.00 | 2.25 | 3.00 | 2.00 | K00425 cytochrome d ubiquinol oxidase subunit I [EC:1.10.3.-] |
| ID5341 | 13.32429 | 0.03816575 | 1.00 | 1.00 | 0.00 | 1.00 | 3.00 | 2.50 | 5.75 | K05341 amylosucrase [EC:2.4.1.4] |
| ID11072 | 13.32242 | 0.03819220 | 2.00 | 1.00 | 1.00 | 10.00 | 6.25 | 3.75 | 10.25 | K11072 spermidine/putrescine transport system ATP-binding protein [EC:3.6.3.31] |
| ID1997 | 13.32205 | 0.03819754 | 3.33 | 13.00 | 16.00 | 3.00 | 3.50 | 16.50 | 4.00 | K01997 branched-chain amino acid transport system permease protein |
| ID1338 | 13.27856 | 0.03881961 | 21.67 | 35.00 | 37.00 | 54.00 | 15.25 | 10.50 | 21.75 | K01338 ATP-dependent Lon protease [EC:3.4.21.53] |
| ID390 | 13.26639 | 0.03899533 | 4.67 | 5.00 | 6.00 | 1.00 | 2.50 | 1.50 | 1.75 | K00390 phosphoadenosine phosphosulfate reductase [EC:1.8.4.8] |
| ID1627 | 13.26264 | 0.03904969 | 5.33 | 5.00 | 7.00 | 1.00 | 1.75 | 1.50 | 1.75 | K01627 2-dehydro-3-deoxyphosphooctonate aldolase (KDO 8-P synthase) [EC:2.5.1.55] |
| ID3110 | 13.26121 | 0.03907047 | 4.33 | 5.00 | 7.00 | 8.00 | 6.00 | 18.25 | 7.00 | K03110 fused signal recognition particle receptor |
| ID31 | 13.25531 | 0.03915615 | 24.67 | 11.00 | 13.00 | 3.00 | 7.00 | 12.50 | 10.50 | K00031 isocitrate dehydrogenase [EC:1.1.1.42] |
| ID1812 | 13.24301 | 0.03933517 | 5.67 | 1.50 | 1.00 | 9.00 | 5.00 | 3.00 | 5.50 | K01812 glucuronate isomerase [EC:5.3.1.12] |
| ID4066 | 13.23011 | 0.03952394 | 6.67 | 12.50 | 17.00 | 20.00 | 8.50 | 22.25 | 15.25 | K04066 primosomal protein N' (replication factor Y) (superfamily II helicase) [EC:3.6.4.-] |
| ID1571 | 13.21009 | 0.03981840 | 2.67 | 1.00 | 1.00 | 1.00 | 7.00 | 3.00 | 11.50 | K01571 oxaloacetate decarboxylase |
| ID2047 | 13.17672 | 0.04031383 | 2.67 | 4.00 | 6.00 | 1.00 | 1.00 | 1.00 | 2.50 | K02047 sulfate transport system permease protein |
| ID3654 | 13.11111 | 0.04130504 | 15.33 | 23.00 | 29.00 | 32.00 | 10.00 | 20.00 | 16.50 | K03654 ATP-dependent DNA helicase RecQ [EC:3.6.4.12] |
| ID2564 | 13.10997 | 0.04132243 | 3.33 | 1.00 | 1.00 | 10.00 | 6.75 | 13.25 | 8.75 | K02564 glucosamine-6-phosphate deaminase [EC:3.5.99.6] |
| ID7636 | 13.10606 | 0.04138227 | 3.33 | 6.00 | 6.00 | 13.00 | 5.75 | 6.00 | 9.50 | K07636 two-component system |
| ID2314 | 13.08561 | 0.04169637 | 12.00 | 13.50 | 15.00 | 29.00 | 7.50 | 23.25 | 14.50 | K02314 replicative DNA helicase [EC:3.6.4.12] |
| ID3590 | 13.06922 | 0.04194980 | 10.33 | 8.00 | 11.00 | 9.00 | 2.00 | 4.00 | 2.75 | K03590 cell division protein FtsA |
| ID1424 | 13.05987 | 0.04209497 | 2.00 | 3.00 | 3.00 | 9.00 | 2.75 | 11.25 | 5.00 | K01424 L-asparaginase [EC:3.5.1.1] |
| ID767 | 13.04208 | 0.04237250 | 2.67 | 4.00 | 4.00 | 6.00 | 2.75 | 10.25 | 6.25 | K00767 nicotinate-nucleotide pyrophosphorylase (carboxylating) [EC:2.4.2.19] |
| ID5343 | 13.03857 | 0.04242758 | 16.67 | 12.00 | 15.00 | 3.00 | 1.00 | 11.00 | 4.75 | K05343 maltose alpha-D-glucosyltransferase/ alpha-amylase [EC:5.4.99.16 3.2.1.1] |
| ID3408 | 13.03738 | 0.04244613 | 4.67 | 6.00 | 7.00 | 1.00 | 1.50 | 2.25 | 1.00 | K03408 purine-binding chemotaxis protein CheW |
| ID1915 | 13.02579 | 0.04262817 | 41.67 | 58.50 | 75.00 | 41.00 | 24.25 | 34.75 | 37.00 | K01915 glutamine synthetase [EC:6.3.1.2] |
| ID970 | 13.01829 | 0.04274638 | 4.67 | 8.00 | 12.00 | 2.00 | 1.25 | 13.50 | 2.00 | K00970 poly(A) polymerase [EC:2.7.7.19] |
| ID383 | 12.99306 | 0.04314636 | 2.67 | 6.00 | 7.00 | 1.00 | 0.75 | 1.50 | 1.00 | K00383 glutathione reductase (NADPH) [EC:1.8.1.7] |
| ID2867 | 12.99295 | 0.04314807 | 5.33 | 3.00 | 4.00 | 8.00 | 4.25 | 9.00 | 5.75 | K02867 large subunit ribosomal protein L11 |
| ID4487 | 12.98722 | 0.04323927 | 6.33 | 12.00 | 13.00 | 22.00 | 9.25 | 18.00 | 15.25 | K04487 cysteine desulfurase [EC:2.8.1.7] |
| ID1996 | 12.98471 | 0.04327936 | 3.67 | 12.50 | 16.00 | 3.00 | 3.75 | 10.00 | 5.00 | K01996 branched-chain amino acid transport system ATP-binding protein |
| ID1187 | 12.97579 | 0.04342198 | 14.33 | 2.00 | 1.00 | 33.00 | 9.25 | 24.75 | 24.75 | K01187 alpha-glucosidase [EC:3.2.1.20] |
| ID2945 | 12.96123 | 0.04365577 | 22.00 | 11.50 | 15.00 | 11.00 | 9.25 | 15.00 | 11.50 | K02945 small subunit ribosomal protein S1 |
| ID3587 | 12.95632 | 0.04373490 | 3.67 | 9.00 | 12.00 | 2.00 | 3.00 | 15.00 | 6.00 | K03587 cell division protein FtsI (penicillin-binding protein 3) |
| ID2073 | 12.93911 | 0.04401321 | 3.00 | 10.00 | 13.00 | 3.00 | 3.75 | 18.25 | 4.00 | K02073 D-methionine transport system substrate-binding protein |
| ID1008 | 12.92623 | 0.04422256 | 2.67 | 4.00 | 4.00 | 8.00 | 2.25 | 1.25 | 3.25 | K01008 selenide |
| ID4077 | 12.92385 | 0.04426121 | 13.33 | 12.00 | 13.00 | 20.00 | 13.50 | 19.50 | 16.50 | K04077 chaperonin GroEL |
| ID702 | 12.92027 | 0.04431968 | 2.00 | 1.50 | 1.00 | 2.00 | 9.75 | 4.50 | 9.50 | K00702 cellobiose phosphorylase [EC:2.4.1.20] |
| ID1653 | 12.91875 | 0.04434450 | 3.00 | 3.00 | 4.00 | 1.00 | 2.75 | 8.25 | 5.50 | K01653 acetolactate synthase I/III small subunit [EC:2.2.1.6] |
| ID8303 | 12.90226 | 0.04461458 | 14.33 | 13.00 | 14.00 | 29.00 | 13.00 | 23.50 | 20.75 | K08303 putative protease [EC:3.4.-.-] |
| ID5 | 12.88718 | 0.04486298 | 1.00 | 1.00 | 1.00 | 6.00 | 1.75 | 3.25 | 3.75 | K00005 glycerol dehydrogenase [EC:1.1.1.6] |
| ID677 | 12.87805 | 0.04501394 | 5.00 | 5.00 | 6.00 | 1.00 | 2.00 | 1.00 | 2.75 | K00677 UDP-N-acetylglucosamine acyltransferase [EC:2.3.1.129] |
| ID1426 | 12.87179 | 0.04511764 | 6.33 | 9.00 | 11.00 | 2.00 | 1.00 | 4.50 | 1.00 | K01426 amidase [EC:3.5.1.4] |
| ID748 | 12.85759 | 0.04535398 | 2.67 | 4.00 | 5.00 | 1.00 | 1.25 | 1.00 | 2.00 | K00748 lipid-A-disaccharide synthase [EC:2.4.1.182] |
| ID2948 | 12.85714 | 0.04536146 | 6.67 | 2.00 | 3.00 | 4.00 | 4.75 | 7.50 | 4.75 | K02948 small subunit ribosomal protein S11 |
| ID12257 | 12.84375 | 0.04558541 | 1.00 | 1.00 | 0.00 | 1.00 | 7.00 | 3.00 | 12.00 | K12257 SecD/SecF fusion protein |
| ID6153 | 12.82660 | 0.04587372 | 2.67 | 6.50 | 8.00 | 9.00 | 4.50 | 8.50 | 7.75 | K06153 undecaprenyl-diphosphatase [EC:3.6.1.27] |
| ID1940 | 12.82426 | 0.04591321 | 15.67 | 10.50 | 13.00 | 5.00 | 11.75 | 13.50 | 17.75 | K01940 argininosuccinate synthase [EC:6.3.4.5] |
| ID963 | 12.81510 | 0.04606793 | 10.00 | 11.00 | 12.00 | 19.00 | 1.75 | 14.75 | 4.50 | K00963 UTP--glucose-1-phosphate uridylyltransferase [EC:2.7.7.9] |
| ID2523 | 12.80158 | 0.04629741 | 3.67 | 5.00 | 6.00 | 1.00 | 1.25 | 1.25 | 2.25 | K02523 octaprenyl-diphosphate synthase [EC:2.5.1.90] |
| ID2203 | 12.78151 | 0.04663987 | 3.67 | 4.00 | 5.00 | 1.00 | 1.75 | 1.00 | 2.00 | K02203 phosphoserine / homoserine phosphotransferase [EC:3.1.3.3 2.7.1.39] |
| ID1890 | 12.76857 | 0.04686192 | 12.00 | 12.50 | 15.00 | 24.00 | 12.50 | 23.00 | 17.75 | K01890 phenylalanyl-tRNA synthetase beta chain [EC:6.1.1.20] |
| ID1783 | 12.74298 | 0.04730387 | 5.33 | 5.00 | 6.00 | 6.00 | 3.25 | 7.00 | 5.75 | K01783 ribulose-phosphate 3-epimerase [EC:5.1.3.1] |

| | | | | | | | | | |
|---|---|---|---|---|---|---|---|---|---|
| ID764 | 12.72937 | 0.04754063 | 16.33 | 12.00 | 16.00 | 13.00 | 10.25 | 16.00 | 17.25 | K00764 amidophosphoribosyltransferase [EC:2.4.2.14] |
| ID2967 | 12.72418 | 0.04763110 | 12.00 | 6.00 | 7.00 | 7.00 | 6.75 | 11.00 | 8.50 | K02967 small subunit ribosomal protein S2 |
| ID1494 | 12.70796 | 0.04791515 | 7.33 | 5.50 | 5.00 | 1.00 | 1.00 | 10.25 | 1.50 | K01494 dCTP deaminase [EC:3.5.4.13] |
| ID11690 | 12.69246 | 0.04818807 | 1.67 | 3.00 | 4.00 | 1.00 | 2.25 | 1.75 | 6.75 | K11690 C4-dicarboxylate transporter |
| ID930 | 12.66473 | 0.04868006 | 5.33 | 6.00 | 9.00 | 2.00 | 4.75 | 10.25 | 8.75 | K00930 acetylglutamate kinase [EC:2.7.2.8] |
| ID2501 | 12.66033 | 0.04875855 | 3.67 | 4.50 | 6.00 | 1.00 | 2.50 | 7.25 | 4.50 | K02501 glutamine amidotransferase [EC:2.4.2.-] |
| ID13 | 12.65692 | 0.04881939 | 5.33 | 6.00 | 8.00 | 2.00 | 5.50 | 13.75 | 10.50 | K00013 histidinol dehydrogenase [EC:1.1.1.23] |
| ID13954 | 12.65278 | 0.04889346 | 1.00 | 4.00 | 5.00 | 10.00 | 1.75 | 5.00 | 1.00 | K13954 alcohol dehydrogenase [EC:1.1.1.1] |
| ID788 | 12.64315 | 0.04906602 | 1.00 | 2.00 | 3.00 | 7.00 | 2.50 | 3.25 | 4.75 | K00788 thiamine-phosphate pyrophosphorylase [EC:2.5.1.3] |
| ID2335 | 12.64002 | 0.04912230 | 14.67 | 115.00 | 22.00 | 31.00 | 13.50 | 26.50 | 18.00 | K02335 DNA polymerase I [EC:2.7.7.7] |
| ID4486 | 12.63000 | 0.04930266 | 1.00 | 1.00 | 1.00 | 26.00 | 3.00 | 3.25 | 4.25 | K04486 histidinol-phosphatase (PHP family) [EC:3.1.3.15] |
| ID2048 | 12.62432 | 0.04940519 | 5.33 | 12.00 | 15.00 | 3.00 | 2.00 | 1.50 | 4.50 | K02048 sulfate transport system substrate-binding protein |
| ID796 | 12.61521 | 0.04957012 | 1.67 | 2.50 | 3.00 | 6.00 | 1.25 | 8.50 | 3.00 | K00796 dihydropteroate synthase [EC:2.5.1.15] |
| ID891 | 12.60507 | 0.04975427 | 1.00 | 4.00 | 5.00 | 4.00 | 2.75 | 3.25 | 4.75 | K00891 shikimate kinase [EC:2.7.1.71] |
| ID3628 | 12.59840 | 0.04987558 | 16.67 | 15.00 | 18.00 | 17.00 | 7.50 | 20.25 | 8.00 | K03628 transcription termination factor Rho |
| ID2313 | 12.59565 | 0.04992588 | 17.00 | 14.00 | 15.00 | 28.00 | 7.50 | 28.50 | 14.75 | K02313 chromosomal replication initiator protein |

**Supplementary Table 8.** Enrichment and depletion of KO metabolic gene categories in the ancient (BRS and Ötzi) and modern (Italian, Hadza and Malawian) comparative IM sample cohorts based on false discovery rate (FDR) corrected *p* -values (*q* = <0.05).

| ID | Test-Statistic | FDR_P | Ancient_mean | Modern_mean | KEGG categories |
|---|---|---|---|---|---|
| ID11358 | 13.3618605 | 0.00288992 | 1.00 | 8.00 | K11358 aspartate aminotransferase [EC:2.6.1.1] |
| ID128 | 13.2754159 | 0.00288992 | 16.14 | 3.00 | K00128 aldehyde dehydrogenase (NAD+) [EC:1.2.1.3] |
| ID123 | 13.4329174 | 0.00288992 | 13.43 | 1.58 | K00123 formate dehydrogenase major subunit [EC:1.2.1.2] |
| ID3781 | 13.2754159 | 0.00288992 | 32.86 | 1.42 | K03781 catalase [EC:1.11.1.6] |
| ID575 | 13.4858550 | 0.00288992 | 8.43 | 1.42 | K00575 chemotaxis protein methyltransferase CheR [EC:2.1.1.80] |
| ID1007 | 13.6280835 | 0.00288992 | 12.29 | 1.25 | K01007 pyruvate |
| ID2454 | 14.1656805 | 0.00288992 | 8.29 | 1.25 | K02454 general secretion pathway protein E |
| ID164 | 14.0961727 | 0.00288992 | 19.86 | 1.17 | K00164 2-oxoglutarate dehydrogenase E1 component [EC:1.2.4.2] |
| ID370 | 14.8784161 | 0.00288992 | 11.00 | 1.17 | K00370 nitrate reductase alpha subunit [EC:1.7.99.4] |
| ID248 | 13.3370474 | 0.00288992 | 7.14 | 1.08 | K00248 butyryl-CoA dehydrogenase [EC:1.3.8.1] |
| ID2405 | 13.4858550 | 0.00288992 | 4.29 | 1.08 | K02405 RNA polymerase sigma factor for flagellar operon FliA |
| ID940 | 15.7500000 | 0.00288992 | 3.71 | 1.08 | K00940 nucleoside-diphosphate kinase [EC:2.7.4.6] |
| ID163 | 14.1517241 | 0.00288992 | 17.86 | 1.00 | K00163 pyruvate dehydrogenase E1 component [EC:1.2.4.1] |
| ID13893 | 13.3123262 | 0.00288992 | 12.00 | 1.00 | K13893 microcin C transport system substrate-binding protein |
| ID362 | 14.7474333 | 0.00288992 | 10.86 | 1.00 | K00362 nitrite reductase (NADH) large subunit [EC:1.7.1.15] |
| ID627 | 14.4131919 | 0.00288992 | 9.43 | 1.00 | K00627 pyruvate dehydrogenase E2 component (dihydrolipoamide acetyltransferase) [EC:2.3.1.12] |
| ID4564 | 13.6800000 | 0.00288992 | 7.86 | 1.00 | K04564 superoxide dismutase |
| ID1255 | 14.7474333 | 0.00288992 | 7.71 | 1.00 | K01255 leucyl aminopeptidase [EC:3.4.11.1] |
| ID631 | 14.4280203 | 0.00288992 | 7.43 | 1.00 | K00631 glycerol-3-phosphate O-acyltransferase [EC:2.3.1.15] |
| ID3582 | 14.4131919 | 0.00288992 | 6.29 | 1.00 | K03582 exodeoxyribonuclease V beta subunit [EC:3.1.11.5] |
| ID140 | 13.3370474 | 0.00288992 | 14.86 | 0.92 | K00140 malonate-semialdehyde dehydrogenase (acetylating) / methylmalonate-semialdehyde dehydrogenase [EC:1.2.1.18 1.2.1.27] |
| ID1682 | 14.1656805 | 0.00288992 | 12.00 | 0.92 | K01682 aconitate hydratase 2 / 2-methylisocitrate dehydratase [EC:4.2.1.3 4.2.1.99] |
| ID1638 | 13.3944945 | 0.00288992 | 11.86 | 0.92 | K01638 malate synthase [EC:2.3.3.9] |
| ID2002 | 14.1656805 | 0.00288992 | 11.57 | 0.92 | K02002 glycine betaine/proline transport system substrate-binding protein |
| ID3415 | 14.1377953 | 0.00288992 | 8.14 | 0.92 | K03415 two-component system |
| ID3583 | 14.1238938 | 0.00288992 | 7.57 | 0.92 | K03583 exodeoxyribonuclease V gamma subunit [EC:3.1.11.5] |
| ID1139 | 13.8304100 | 0.00288992 | 6.57 | 0.92 | K01139 guanosine-3' |
| ID5847 | 13.4991547 | 0.00288992 | 6.29 | 0.92 | K05847 osmoprotectant transport system ATP-binding protein |
| ID1433 | 13.6669838 | 0.00288992 | 10.57 | 0.83 | K01433 formyltetrahydrofolate deformylase [EC:3.5.1.10] |
| ID6445 | 13.3689568 | 0.00288992 | 8.86 | 0.83 | K06445 acyl-CoA dehydrogenase [EC:1.3.99.-] |
| ID1584 | 13.3689568 | 0.00288992 | 8.71 | 0.83 | K01584 arginine decarboxylase [EC:4.1.1.19] |
| ID117 | 13.3944945 | 0.00288992 | 7.00 | 0.83 | K00117 quinoprotein glucose dehydrogenase [EC:1.1.5.2] |
| ID5365 | 13.3689568 | 0.00288992 | 7.00 | 0.83 | K05365 penicillin-binding protein 1B [EC:2.4.1.129 3.4.-.-] |
| ID7712 | 14.8082474 | 0.00288992 | 6.86 | 0.83 | K07712 two-component system |
| ID7788 | 14.7625899 | 0.00288992 | 6.71 | 0.83 | K07788 multidrug efflux pump |
| ID1011 | 14.7929969 | 0.00288992 | 6.14 | 0.83 | K01011 thiosulfate/3-mercaptopyruvate sulfurtransferase [EC:2.8.1.1 2.8.1.2] |
| ID955 | 13.3689568 | 0.00288992 | 6.00 | 0.83 | K00955 bifunctional enzyme CysN/CysC [EC:2.7.7.4 2.7.1.25] |
| ID3735 | 14.7777778 | 0.00288992 | 5.86 | 0.83 | K03735 ethanolamine ammonia-lyase large subunit [EC:4.3.1.7] |
| ID1637 | 14.1517241 | 0.00288992 | 9.43 | 0.75 | K01637 isocitrate lyase [EC:4.1.3.1] |
| ID116 | 14.2077151 | 0.00288992 | 8.57 | 0.75 | K00116 malate dehydrogenase (quinone) [EC:1.1.5.4] |
| ID121 | 14.2783300 | 0.00288992 | 8.00 | 0.75 | K00121 S-(hydroxymethyl)glutathione dehydrogenase / alcohol dehydrogenase [EC:1.1.1.284 1.1.1.1] |
| ID632 | 14.2077151 | 0.00288992 | 7.86 | 0.75 | K00632 acetyl-CoA acyltransferase [EC:2.3.1.16] |
| ID11891 | 14.1517241 | 0.00288992 | 7.14 | 0.75 | K11891 type VI secretion system protein ImpL |
| ID1690 | 14.2077151 | 0.00288992 | 6.29 | 0.75 | K01690 phosphogluconate dehydratase [EC:4.2.1.12] |
| ID257 | 13.7454545 | 0.00288992 | 11.29 | 0.67 | K00257 [EC:1.3.99.-] |
| ID11073 | 13.7850288 | 0.00288992 | 10.57 | 0.67 | K11073 putrescine transport system substrate-binding protein |
| ID11907 | 13.7454545 | 0.00288992 | 8.00 | 0.67 | K11907 type VI secretion system protein VasG |
| ID1959 | 13.7323136 | 0.00288992 | 8.00 | 0.67 | K01959 pyruvate carboxylase subunit A [EC:6.4.1.1] |
| ID108 | 13.7718121 | 0.00288992 | 7.29 | 0.67 | K00108 choline dehydrogenase |

| | | | | | |
|---|---|---|---|---|---|
| ID11904 | 13.7454545 | 0.00288992 | 7.29 | 0.67 | K11904 type VI secretion system secreted protein VgrG |
| ID285 | 13.7586207 | 0.00288992 | 6.57 | 0.67 | K00285 D-amino-acid dehydrogenase [EC:1.4.99.1] |
| ID11076 | 13.7982709 | 0.00288992 | 6.14 | 0.67 | K11076 putrescine transport system ATP-binding protein |
| ID322 | 13.7454545 | 0.00288992 | 6.14 | 0.67 | K00322 NAD(P) transhydrogenase [EC:1.6.1.1] |
| ID130 | 13.7586207 | 0.00288992 | 5.43 | 0.67 | K00130 betaine-aldehyde dehydrogenase [EC:1.2.1.8] |
| ID13895 | 13.7454545 | 0.00288992 | 5.29 | 0.67 | K13895 microcin C transport system permease protein |
| ID799 | 13.5000000 | 0.00288992 | 6.71 | 0.58 | K00799 glutathione S-transferase [EC:2.5.1.18] |
| ID372 | 13.4620431 | 0.00288992 | 6.00 | 0.50 | K00372 assimilatory nitrate reductase catalytic subunit [EC:1.7.99.4] |
| ID404 | 13.5254237 | 0.00288992 | 13.71 | 0.42 | K00404 cytochrome c oxidase cbb3-type subunit I [EC:1.9.3.1] |
| ID412 | 13.7454545 | 0.00288992 | 14.29 | 0.33 | K00412 ubiquinol-cytochrome c reductase cytochrome b subunit |
| ID10125 | 13.8782609 | 0.00288992 | 6.00 | 0.33 | K10125 two-component system |
| ID249 | 14.1517241 | 0.00288992 | 11.43 | 0.25 | K00249 acyl-CoA dehydrogenase [EC:1.3.8.7] |
| ID2274 | 14.7625899 | 0.00288992 | 17.00 | 0.17 | K02274 cytochrome c oxidase subunit I [EC:1.9.3.1] |
| ID6596 | 14.7625899 | 0.00288992 | 8.57 | 0.17 | K06596 chemosensory pili system protein ChpA (sensor histidine kinase/response regulator) |
| ID3883 | 13.5011103 | 0.00288992 | 8.14 | 0.17 | K03883 NADH-ubiquinone oxidoreductase chain 5 [EC:1.6.5.3] |
| ID457 | 14.7625899 | 0.00288992 | 7.43 | 0.17 | K00457 4-hydroxyphenylpyruvate dioxygenase [EC:1.13.11.27] |
| ID1969 | 14.1552667 | 0.00288992 | 6.14 | 0.17 | K01969 3-methylcrotonyl-CoA carboxylase beta subunit [EC:6.4.1.4] |
| ID13482 | 14.7777778 | 0.00288992 | 5.29 | 0.17 | K13482 xanthine dehydrogenase large subunit [EC:1.17.1.4] |
| ID253 | 14.1260504 | 0.00288992 | 4.71 | 0.17 | K00253 isovaleryl-CoA dehydrogenase [EC:1.3.8.4] |
| ID1907 | 14.1552667 | 0.00288992 | 3.71 | 0.17 | K01907 acetoacetyl-CoA synthetase [EC:6.2.1.16] |
| ID1113 | 13.5573064 | 0.00288992 | 3.29 | 0.17 | K01113 alkaline phosphatase D [EC:3.1.3.1] |
| ID2275 | 15.2767274 | 0.00288992 | 6.00 | 0.08 | K02275 cytochrome c oxidase subunit II [EC:1.9.3.1] |
| ID2276 | 15.2933868 | 0.00288992 | 4.71 | 0.08 | K02276 cytochrome c oxidase subunit III [EC:1.9.3.1] |
| ID3381 | 14.9597190 | 0.00288992 | 3.29 | 0.08 | K03381 catechol 1 |
| ID7649 | 14.4922930 | 0.00288992 | 3.14 | 0.08 | K07649 two-component system |
| ID3879 | 14.9760863 | 0.00288992 | 2.86 | 0.08 | K03879 NADH-ubiquinone oxidoreductase chain 2 [EC:1.6.5.3] |
| ID252 | 16.9187279 | 0.00288992 | 7.71 | 0.00 | K00252 glutaryl-CoA dehydrogenase [EC:1.3.8.6] |
| ID2261 | 16.8393904 | 0.00288992 | 5.00 | 0.00 | K02261 cytochrome c oxidase subunit 2 |
| ID451 | 16.9187279 | 0.00288992 | 4.14 | 0.00 | K00451 homogentisate 1 |
| ID167 | 16.8591549 | 0.00288992 | 4.00 | 0.00 | K00167 2-oxoisovalerate dehydrogenase E1 component beta subunit [EC:1.2.4.4] |
| ID135 | 13.1201586 | 0.00297619 | 10.71 | 1.33 | K00135 succinate-semialdehyde dehydrogenase / glutarate-semialdehyde dehydrogenase [EC:1.2.1.16 1.2.1.79 1.2.1.20] |
| ID3182 | 13.0612050 | 0.00297619 | 9.29 | 1.33 | K03182 4-hydroxy-3-polyprenylbenzoate decarboxylase [EC:4.1.1.98] |
| ID3885 | 13.1622137 | 0.00297619 | 5.29 | 1.33 | K03885 NADH dehydrogenase [EC:1.6.99.3] |
| ID1941 | 13.1188360 | 0.00297619 | 9.29 | 0.75 | K01941 urea carboxylase [EC:6.3.4.6] |
| ID381 | 13.1188360 | 0.00297619 | 8.86 | 0.75 | K00381 sulfite reductase (NADPH) hemoprotein beta-component [EC:1.8.1.2] |
| ID8484 | 13.0821229 | 0.00297619 | 7.86 | 0.75 | K08484 phosphotransferase system |
| ID1825 | 13.0943377 | 0.00297619 | 6.71 | 0.75 | K01825 3-hydroxyacyl-CoA dehydrogenase / enoyl-CoA hydratase / 3-hydroxybutyryl-CoA epimerase / enoyl-CoA isomerase [EC:1.1.1.35 4.2.1.17 5.1.2.3 5.3.3.8] |
| ID3205 | 12.8479428 | 0.0030813 | 1.29 | 35.08 | K03205 type IV secretion system protein VirD4 |
| ID10119 | 12.7453416 | 0.0030813 | 1.71 | 23.25 | K10119 raffinose/stachyose/melibiose transport system permease protein |
| ID651 | 12.8135593 | 0.0030813 | 1.43 | 10.58 | K00651 homoserine O-succinyltransferase [EC:2.3.1.46] |
| ID40 | 12.8940754 | 0.0030813 | 1.57 | 4.92 | K00040 fructuronate reductase [EC:1.1.1.57] |
| ID626 | 12.7566607 | 0.0030813 | 18.86 | 4.17 | K00626 acetyl-CoA C-acetyltransferase [EC:2.3.1.9] |
| ID1845 | 12.9522092 | 0.0030813 | 12.71 | 2.75 | K01845 glutamate-1-semialdehyde 2 |
| ID1962 | 12.7839888 | 0.0030813 | 7.43 | 2.33 | K01962 acetyl-CoA carboxylase carboxyl transferase subunit alpha [EC:6.4.1.2] |
| ID281 | 12.7731092 | 0.0030813 | 14.43 | 2.00 | K00281 glycine dehydrogenase [EC:1.4.4.2] |
| ID681 | 12.8360310 | 0.0030813 | 8.29 | 1.33 | K00681 gamma-glutamyltranspeptidase / glutathione hydrolase [EC:2.3.2.2 3.4.19.13] |
| ID3644 | 12.7450120 | 0.0030813 | 9.57 | 0.92 | K03644 lipoic acid synthetase [EC:2.8.1.8] |
| ID647 | 12.7327716 | 0.0030813 | 5.86 | 0.92 | K00647 3-oxoacyl-[acyl-carrier-protein] synthase I [EC:2.3.1.41] |
| ID1772 | 12.7777638 | 0.0030813 | 5.43 | 0.92 | K01772 ferrochelatase [EC:4.99.1.1] |
| ID1241 | 12.9612160 | 0.0030813 | 6.57 | 0.67 | K01241 AMP nucleosidase [EC:3.2.2.4] |
| ID10117 | 12.6778464 | 0.00312589 | 2.00 | 40.58 | K10117 raffinose/stachyose/melibiose transport system substrate-binding protein |
| ID2051 | 12.6685056 | 0.00312589 | 8.29 | 3.25 | K02051 |

| | | | | | |
|---|---|---|---|---|---|
| ID5349 | 12.6443662 | 0.00313547 | 10.71 | 53.00 | K05349 beta-glucosidase [EC:3.2.1.21] |
| ID405 | 12.5879917 | 0.00318264 | 4.71 | 0.33 | K00405 cytochrome c oxidase cbb3-type subunit II |
| ID1692 | 12.5758411 | 0.00318264 | 4.71 | 0.33 | K01692 enoyl-CoA hydratase [EC:4.2.1.17] |
| ID7516 | 12.5622711 | 0.00318264 | 3.57 | 0.17 | K07516 3-hydroxyacyl-CoA dehydrogenase [EC:1.1.1.35] |
| ID1878 | 12.5294801 | 0.00320843 | 7.71 | 1.67 | K01878 glycyl-tRNA synthetase alpha chain [EC:6.1.1.14] |
| ID1749 | 12.4925200 | 0.00324195 | 4.57 | 1.50 | K01749 hydroxymethylbilane synthase [EC:2.5.1.61] |
| ID1488 | 12.4592551 | 0.00326964 | 6.43 | 1.42 | K01488 adenosine deaminase [EC:3.5.4.4] |
| ID1740 | 12.1241301 | 0.00358116 | 8.86 | 23.92 | K01740 O-acetylhomoserine (thiol)-lyase [EC:2.5.1.49] |
| ID10118 | 12.1672381 | 0.00358116 | 1.57 | 23.83 | K10118 raffinose/stachyose/melibiose transport system permease protein |
| ID1712 | 12.1348784 | 0.00358116 | 9.43 | 2.08 | K01712 urocanate hydratase [EC:4.2.1.49] |
| ID658 | 12.1433240 | 0.00358116 | 6.71 | 0.83 | K00658 2-oxoglutarate dehydrogenase E2 component (dihydrolipoamide succinyltransferase) [EC:2.3.1.61] |
| ID641 | 12.1433240 | 0.00358116 | 5.86 | 0.83 | K00641 homoserine O-acetyltransferase [EC:2.3.1.31] |
| ID8311 | 12.2635549 | 0.00358116 | 4.00 | 0.83 | K08311 putative (di)nucleoside polyphosphate hydrolase [EC:3.6.1.-] |
| ID1920 | 12.1565664 | 0.00358116 | 4.00 | 0.83 | K01920 glutathione synthase [EC:6.3.2.3] |
| ID1141 | 12.1831382 | 0.00358116 | 3.43 | 0.83 | K01141 exodeoxyribonuclease I [EC:3.1.11.1] |
| ID406 | 12.1345682 | 0.00358116 | 4.86 | 0.42 | K00406 cytochrome c oxidase cbb3-type subunit III |
| ID1029 | 12.2045080 | 0.00358116 | 4.00 | 0.42 | K01029 3-oxoacid CoA-transferase subunit B [EC:2.8.3.5] |
| ID1581 | 12.0998980 | 0.00359752 | 8.29 | 1.67 | K01581 ornithine decarboxylase [EC:4.1.1.17] |
| ID1182 | 11.9787636 | 0.00379742 | 1.57 | 14.50 | K01182 oligo-1 |
| ID3590 | 11.9680300 | 0.00379742 | 9.57 | 2.92 | K03590 cell division protein FtsA |
| ID425 | 11.9359441 | 0.00383172 | 13.14 | 2.42 | K00425 cytochrome d ubiquinol oxidase subunit I [EC:1.10.3.-] |
| ID4517 | 11.8119415 | 0.00393105 | 1.29 | 5.25 | K04517 prephenate dehydrogenase [EC:1.3.1.12] |
| ID1961 | 11.7988705 | 0.00393105 | 12.86 | 4.92 | K01961 acetyl-CoA carboxylase |
| ID432 | 11.8293082 | 0.00393105 | 6.71 | 1.58 | K00432 glutathione peroxidase [EC:1.11.1.9] |
| ID1679 | 11.8067547 | 0.00393105 | 7.86 | 1.50 | K01679 fumarate hydratase |
| ID2301 | 11.8050442 | 0.00393105 | 4.43 | 0.75 | K02301 protoheme IX farnesyltransferase [EC:2.5.1.-] |
| ID1070 | 11.8416679 | 0.00393105 | 3.86 | 0.75 | K01070 S-formylglutathione hydrolase [EC:3.1.2.12] |
| ID1659 | 11.7582857 | 0.00394009 | 4.43 | 0.67 | K01659 2-methylcitrate synthase [EC:2.3.3.5] |
| ID1497 | 11.6997868 | 0.00394009 | 3.57 | 0.67 | K01497 GTP cyclohydrolase II [EC:3.5.4.25] |
| ID3087 | 11.7348161 | 0.00394009 | 3.14 | 0.67 | K03087 RNA polymerase nonessential primary-like sigma factor |
| ID1455 | 11.7068380 | 0.00394009 | 6.86 | 0.58 | K01455 formamidase [EC:3.5.1.49] |
| ID10941 | 11.6954942 | 0.00394009 | 5.00 | 0.58 | K10941 sigma-54 specific transcriptional regulator |
| ID228 | 11.7638889 | 0.00394009 | 4.86 | 0.58 | K00228 coproporphyrinogen III oxidase [EC:1.3.3.3] |
| ID568 | 11.7068380 | 0.00394009 | 3.43 | 0.58 | K00568 2-polyprenyl-6-hydroxyphenyl methylase / 3-demethylubiquinone-9 3-methyltransferase [EC:2.1.1.222 2.1.1.64] |
| ID4561 | 11.6534051 | 0.00394263 | 6.71 | 0.67 | K04561 nitric oxide reductase subunit B [EC:1.7.2.5] |
| ID11903 | 11.6649660 | 0.00394263 | 3.86 | 0.67 | K11903 type VI secretion system secreted protein Hcp |
| ID12339 | 11.6615942 | 0.00394263 | 3.71 | 0.58 | K12339 cysteine synthase B [EC:2.5.1.47] |
| ID1613 | 11.6132476 | 0.00399969 | 6.00 | 1.83 | K01613 phosphatidylserine decarboxylase [EC:4.1.1.65] |
| ID426 | 11.5943804 | 0.00401161 | 5.71 | 1.42 | K00426 cytochrome d ubiquinol oxidase subunit II [EC:1.10.3.-] |
| ID2492 | 11.5731159 | 0.00402897 | 5.71 | 1.58 | K02492 glutamyl-tRNA reductase [EC:1.2.1.70] |
| ID1077 | 11.4950000 | 0.00417228 | 1.00 | 9.42 | K01077 alkaline phosphatase [EC:3.1.3.1] |
| ID525 | 11.4587260 | 0.00422476 | 39.00 | 12.33 | K00525 ribonucleoside-diphosphate reductase alpha chain [EC:1.17.4.1] |
| ID2556 | 11.4255403 | 0.00427102 | 6.29 | 1.42 | K02556 chemotaxis protein MotA |
| ID600 | 11.2432680 | 0.00450554 | 17.00 | 10.42 | K00600 glycine hydroxymethyltransferase [EC:2.1.2.1] |
| ID48 | 11.2695957 | 0.00450554 | 1.71 | 9.17 | K00048 lactaldehyde reductase [EC:1.1.1.77] |
| ID548 | 11.2632029 | 0.00450554 | 19.86 | 7.92 | K00548 5-methyltetrahydrofolate--homocysteine methyltransferase [EC:2.1.1.13] |
| ID5350 | 11.2695957 | 0.00450554 | 1.43 | 7.00 | K05350 beta-glucosidase [EC:3.2.1.21] |
| ID648 | 11.2389163 | 0.00450554 | 7.14 | 3.67 | K00648 3-oxoacyl-[acyl-carrier-protein] synthase III [EC:2.3.1.180] |
| ID2410 | 11.2381285 | 0.00450554 | 6.14 | 1.75 | K02410 flagellar motor switch protein FliG |
| ID338 | 11.2534455 | 0.00450554 | 4.43 | 0.92 | K00338 NADH-quinone oxidoreductase subunit I [EC:1.6.5.3] |
| ID2535 | 11.1493540 | 0.00469525 | 5.14 | 0.92 | K02535 UDP-3-O-[3-hydroxymyristoyl] N-acetylglucosamine deacetylase [EC:3.5.1.108] |
| ID2412 | 11.1037259 | 0.00478073 | 6.86 | 2.17 | K02412 flagellum-specific ATP synthase [EC:3.6.3.14] |

| | | | | | |
|---|---|---|---|---|---|
| ID371 | 11.0153868 | 0.00494932 | 6.57 | 0.83 | K00371 nitrate reductase beta subunit [EC:1.7.99.4] |
| ID3841 | 11.0262823 | 0.00494932 | 5.00 | 0.83 | K03841 fructose-1 |
| ID864 | 10.9993656 | 0.00496029 | 14.00 | 6.25 | K00864 glycerol kinase [EC:2.7.1.30] |
| ID2400 | 10.9327446 | 0.00510911 | 8.14 | 2.33 | K02400 flagellar biosynthesis protein FlhA |
| ID2416 | 10.9176283 | 0.00511837 | 5.86 | 1.75 | K02416 flagellar motor switch protein FliM |
| ID833 | 10.8745421 | 0.00517334 | 6.71 | 1.75 | K00833 adenosylmethionine-8-amino-7-oxononanoate aminotransferase [EC:2.6.1.62] |
| ID161 | 10.8778626 | 0.00517334 | 5.43 | 1.17 | K00161 pyruvate dehydrogenase E1 component alpha subunit [EC:1.2.4.1] |
| ID605 | 10.8346050 | 0.00525328 | 12.29 | 2.25 | K00605 aminomethyltransferase [EC:2.1.2.10] |
| ID1486 | 10.7381605 | 0.00550008 | 1.57 | 5.33 | K01486 adenine deaminase [EC:3.5.4.2] |
| ID1720 | 10.7142857 | 0.00553733 | 3.43 | 1.00 | K01720 2-methylcitrate dehydratase [EC:4.2.1.79] |
| ID13821 | 10.6796419 | 0.00560759 | 11.86 | 1.83 | K13821 RHH-type transcriptional regulator |
| ID3655 | 10.6061072 | 0.00579969 | 11.71 | 35.67 | K03655 ATP-dependent DNA helicase RecG [EC:3.6.4.12] |
| ID1737 | 10.5057398 | 0.00608642 | 3.29 | 1.17 | K01737 6-pyruvoyltetrahydropterin/6-carboxytetrahydropterin synthase [EC:4.2.3.12 4.1.2.50] |
| ID334 | 10.4868969 | 0.00611199 | 3.57 | 0.83 | K00334 NADH-quinone oxidoreductase subunit E [EC:1.6.5.3] |
| ID13378 | 10.4288642 | 0.00626951 | 6.71 | 1.58 | K13378 NADH-quinone oxidoreductase subunit C/D [EC:1.6.5.3] |
| ID11959 | 10.4085634 | 0.0063013 | 8.29 | 2.50 | K11959 urea transport system substrate-binding protein |
| ID3732 | 10.3869516 | 0.00633798 | 4.86 | 1.00 | K03732 ATP-dependent RNA helicase RhlB [EC:3.6.4.13] |
| ID331 | 10.3712442 | 0.00635475 | 5.14 | 0.92 | K00331 NADH-quinone oxidoreductase subunit B [EC:1.6.5.3] |
| ID1698 | 10.2606567 | 0.00670795 | 7.71 | 2.75 | K01698 porphobilinogen synthase [EC:4.2.1.24] |
| ID335 | 10.2365211 | 0.00675701 | 7.00 | 3.17 | K00335 NADH-quinone oxidoreductase subunit F [EC:1.6.5.3] |
| ID5341 | 10.0506122 | 0.0073886 | 0.86 | 3.75 | K05341 amylosucrase [EC:2.4.1.4] |
| ID244 | 10.0601934 | 0.0073886 | 0.86 | 3.33 | K00244 fumarate reductase flavoprotein subunit [EC:1.3.5.4] |
| ID3183 | 10.0331633 | 0.00741653 | 5.71 | 1.33 | K03183 demethylmenaquinone methyltransferase / 2-methoxy-6-polyprenyl-1 |
| ID847 | 9.9620835 | 0.00766119 | 2.86 | 9.42 | K00847 fructokinase [EC:2.7.1.4] |
| ID11927 | 9.9525885 | 0.00766119 | 6.29 | 1.92 | K11927 ATP-dependent RNA helicase RhlE [EC:3.6.4.13] |
| ID849 | 9.9354469 | 0.00768967 | 2.57 | 12.25 | K00849 galactokinase [EC:2.7.1.6] |
| ID2757 | 9.8762376 | 0.00789515 | 1.00 | 9.75 | K02757 PTS system |
| ID11755 | 9.8664688 | 0.00789515 | 1.00 | 3.33 | K11755 phosphoribosyl-ATP pyrophosphohydrolase / phosphoribosyl-AMP cyclohydrolase [EC:3.6.1.31 3.5.4.19] |
| ID1591 | 9.8531530 | 0.00790881 | 3.71 | 10.42 | K01591 orotidine-5'-phosphate decarboxylase [EC:4.1.1.23] |
| ID162 | 9.8372781 | 0.00793376 | 7.00 | 1.17 | K00162 pyruvate dehydrogenase E1 component beta subunit [EC:1.2.4.1] |
| ID705 | 9.8043724 | 0.00803308 | 6.14 | 27.83 | K00705 4-alpha-glucanotransferase [EC:2.4.1.25] |
| ID558 | 9.5833228 | 0.00896255 | 8.14 | 14.75 | K00558 DNA (cytosine-5)-methyltransferase 1 [EC:2.1.1.37] |
| ID965 | 9.5917961 | 0.00896255 | 2.71 | 13.17 | K00965 UDPglucose--hexose-1-phosphate uridylyltransferase [EC:2.7.7.12] |
| ID702 | 9.5073359 | 0.009242 | 1.71 | 7.92 | K00702 cellobiose phosphorylase [EC:2.4.1.20] |
| ID111 | 9.5159088 | 0.009242 | 8.57 | 3.42 | K00111 glycerol-3-phosphate dehydrogenase [EC:1.1.5.3] |
| ID3524 | 9.3972777 | 0.00965939 | 2.29 | 8.00 | K03524 BirA family transcriptional regulator |
| ID341 | 9.4140436 | 0.00965939 | 6.43 | 2.17 | K00341 NADH-quinone oxidoreductase subunit L [EC:1.6.5.3] |
| ID383 | 9.4015653 | 0.00965939 | 4.00 | 1.08 | K00383 glutathione reductase (NADPH) [EC:1.8.1.7] |
| ID7407 | 9.3556231 | 0.00972859 | 4.43 | 27.58 | K07407 alpha-galactosidase [EC:3.2.1.22] |
| ID1895 | 9.3556231 | 0.00972859 | 25.43 | 5.58 | K01895 acetyl-CoA synthetase [EC:6.2.1.1] |
| ID1676 | 9.3722405 | 0.00972859 | 9.29 | 2.67 | K01676 fumarate hydratase |
| ID12573 | 9.3390645 | 0.00976655 | 15.14 | 8.33 | K12573 ribonuclease R [EC:3.1.-.-] |
| ID1879 | 9.3137255 | 0.00985202 | 7.00 | 2.17 | K01879 glycyl-tRNA synthetase beta chain [EC:6.1.1.14] |
| ID2048 | 9.2953222 | 0.00990097 | 8.29 | 2.67 | K02048 sulfate transport system substrate-binding protein |
| ID3092 | 9.2201579 | 0.01026368 | 8.00 | 3.25 | K03092 RNA polymerase sigma-54 factor |
| ID7173 | 9.1513761 | 0.01060316 | 1.43 | 4.50 | K07173 S-ribosylhomocysteine lyase [EC:4.4.1.21] |
| ID1803 | 9.1218095 | 0.01072202 | 4.43 | 8.33 | K01803 triosephosphate isomerase (TIM) [EC:5.3.1.1] |
| ID29 | 9.1005030 | 0.01079367 | 7.43 | 2.42 | K00029 malate dehydrogenase (oxaloacetate-decarboxylating)(NADP+) [EC:1.1.1.40] |
| ID3072 | 9.0410781 | 0.011095 | 6.71 | 2.08 | K03072 preprotein translocase subunit SecD |
| ID2049 | 9.0189873 | 0.01117457 | 7.43 | 3.00 | K02049 |
| ID1572 | 8.9658055 | 0.01140914 | 1.00 | 8.58 | K01572 oxaloacetate decarboxylase |
| ID1585 | 8.9622642 | 0.01140914 | 9.71 | 3.42 | K01585 arginine decarboxylase [EC:4.1.1.19] |

| | | | | | |
|---|---|---|---|---|---|
| ID1960 | 8.9542190 | 0.01140914 | 7.43 | 2.08 | K01960 pyruvate carboxylase subunit B [EC:6.4.1.1] |
| ID1426 | 8.9393569 | 0.01144476 | 7.14 | 2.17 | K01426 amidase [EC:3.5.1.4] |
| ID836 | 8.9308756 | 0.01144476 | 3.57 | 0.83 | K00836 diaminobutyrate-2-oxoglutarate transaminase [EC:2.6.1.76] |
| ID7192 | 8.8146833 | 0.01208071 | 0.86 | 3.00 | K07192 flotillin |
| ID1745 | 8.8189588 | 0.01208071 | 10.43 | 2.50 | K01745 histidine ammonia-lyase [EC:4.3.1.3] |
| ID1190 | 8.7653779 | 0.01229458 | 9.00 | 48.08 | K01190 beta-galactosidase [EC:3.2.1.23] |
| ID337 | 8.7699822 | 0.01229458 | 5.29 | 1.67 | K00337 NADH-quinone oxidoreductase subunit H [EC:1.6.5.3] |
| ID1958 | 8.7078102 | 0.01260243 | 7.00 | 2.33 | K01958 pyruvate carboxylase [EC:6.4.1.1] |
| ID2409 | 8.7031633 | 0.01260243 | 5.14 | 1.67 | K02409 flagellar M-ring protein FliF |
| ID1928 | 8.5694392 | 0.0134992 | 5.86 | 11.67 | K01928 UDP-N-acetylmuramoyl-L-alanyl-D-glutamate--2 |
| ID2011 | 8.5496302 | 0.0135837 | 5.86 | 2.08 | K02011 iron(III) transport system permease protein |
| ID131 | 8.4960568 | 0.01392505 | 4.71 | 0.58 | K00131 glyceraldehyde-3-phosphate dehydrogenase (NADP+) [EC:1.2.1.9] |
| ID5592 | 8.4498589 | 0.0142177 | 11.43 | 6.25 | K05592 ATP-dependent RNA helicase DeaD [EC:3.6.4.13] |
| ID12308 | 8.3746823 | 0.01461679 | 3.57 | 24.33 | K12308 beta-galactosidase [EC:3.2.1.23] |
| ID2564 | 8.3821397 | 0.01461679 | 3.29 | 9.58 | K02564 glucosamine-6-phosphate deaminase [EC:3.5.99.6] |
| ID1193 | 8.3821397 | 0.01461679 | 2.57 | 9.58 | K01193 beta-fructofuranosidase [EC:3.2.1.26] |
| ID1206 | 8.3302149 | 0.01484457 | 3.71 | 15.67 | K01206 alpha-L-fucosidase [EC:3.2.1.51] |
| ID2012 | 8.3382857 | 0.01484457 | 9.14 | 1.17 | K02012 iron(III) transport system substrate-binding protein |
| ID1627 | 8.3029695 | 0.01499898 | 4.86 | 1.67 | K01627 2-dehydro-3-deoxyphosphooctonate aldolase (KDO 8-P synthase) [EC:2.5.1.55] |
| ID10536 | 8.2952025 | 0.01499898 | 4.14 | 1.50 | K10536 agmatine deiminase [EC:3.5.3.12] |
| ID1835 | 8.2862173 | 0.01500668 | 9.57 | 18.42 | K01835 phosphoglucomutase [EC:5.4.2.2] |
| ID11070 | 8.1204867 | 0.01629775 | 1.86 | 5.17 | K11070 spermidine/putrescine transport system permease protein |
| ID3408 | 8.1259457 | 0.01629775 | 4.86 | 1.58 | K03408 purine-binding chemotaxis protein CheW |
| ID1915 | 8.0584845 | 0.01673834 | 51.14 | 32.00 | K01915 glutamine synthetase [EC:6.3.1.2] |
| ID1629 | 8.0563147 | 0.01673834 | 1.14 | 3.17 | K01629 rhamnulose-1-phosphate aldolase [EC:4.1.2.19] |
| ID12257 | 7.9637897 | 0.01753953 | 0.86 | 7.33 | K12257 SecD/SecF fusion protein |
| ID16 | 7.9245500 | 0.01784674 | 3.43 | 11.50 | K00016 L-lactate dehydrogenase [EC:1.1.1.27] |
| ID1443 | 7.8892984 | 0.01804239 | 3.14 | 9.33 | K01443 N-acetylglucosamine-6-phosphate deacetylase [EC:3.5.1.25] |
| ID874 | 7.8963236 | 0.01804239 | 2.86 | 7.50 | K00874 2-dehydro-3-deoxygluconokinase [EC:2.7.1.45] |
| ID1142 | 7.8752855 | 0.01804411 | 13.00 | 7.08 | K01142 exodeoxyribonuclease III [EC:3.1.11.2] |
| ID2935 | 7.8737353 | 0.01804411 | 1.43 | 3.75 | K02935 large subunit ribosomal protein L7/L12 |
| ID2203 | 7.8546967 | 0.01815816 | 3.57 | 1.58 | K02203 phosphoserine / homoserine phosphotransferase [EC:3.1.3.3 2.7.1.39] |
| ID8483 | 7.8197279 | 0.01841548 | 5.29 | 15.83 | K08483 phosphotransferase system |
| ID3412 | 7.8140716 | 0.01841548 | 4.43 | 1.67 | K03412 two-component system |
| ID1338 | 7.7922420 | 0.01850363 | 32.29 | 15.83 | K01338 ATP-dependent Lon protease [EC:3.4.21.53] |
| ID3338 | 7.7903836 | 0.01850363 | 4.14 | 0.75 | K03338 5-dehydro-2-deoxygluconokinase [EC:2.7.1.92] |
| ID6215 | 7.7485521 | 0.01885889 | 1.86 | 6.25 | K06215 pyridoxal 5'-phosphate synthase pdxS subunit [EC:4.3.3.6] |
| ID854 | 7.7207297 | 0.01903352 | 4.57 | 10.17 | K00854 xylulokinase [EC:2.7.1.17] |
| ID2523 | 7.7170487 | 0.01903352 | 4.00 | 1.58 | K02523 octaprenyl-diphosphate synthase [EC:2.5.1.90] |
| ID640 | 7.6862313 | 0.01928225 | 7.86 | 4.67 | K00640 serine O-acetyltransferase [EC:2.3.1.30] |
| ID10206 | 7.6730769 | 0.01934436 | 1.00 | 6.08 | K10206 LL-diaminopimelate aminotransferase [EC:2.6.1.83] |
| ID1243 | 7.5871572 | 0.02004376 | 2.43 | 6.50 | K01243 adenosylhomocysteine nucleosidase [EC:3.2.2.9] |
| ID36 | 7.5871572 | 0.02004376 | 13.57 | 4.58 | K00036 glucose-6-phosphate 1-dehydrogenase [EC:1.1.1.49] |
| ID390 | 7.5940671 | 0.02004376 | 4.43 | 1.92 | K00390 phosphoadenosine phosphosulfate reductase [EC:1.8.4.8] |
| ID2000 | 7.5319336 | 0.02058479 | 5.43 | 1.33 | K02000 glycine betaine/proline transport system ATP-binding protein [EC:3.6.3.32] |
| ID848 | 7.4536439 | 0.02141379 | 1.43 | 3.50 | K00848 rhamnulokinase [EC:2.7.1.5] |
| ID1903 | 7.4243697 | 0.02166462 | 9.00 | 3.42 | K01903 succinyl-CoA synthetase beta subunit [EC:6.2.1.5] |
| ID104 | 7.4184037 | 0.02166462 | 6.71 | 2.33 | K00104 glycolate oxidase [EC:1.1.3.15] |
| ID878 | 7.4058203 | 0.02173079 | 1.43 | 4.92 | K00878 hydroxyethylthiazole kinase [EC:2.7.1.50] |
| ID1938 | 7.3594755 | 0.02195233 | 5.57 | 16.33 | K01938 formate--tetrahydrofolate ligase [EC:6.3.4.3] |
| ID1929 | 7.3659768 | 0.02195233 | 4.29 | 11.67 | K01929 UDP-N-acetylmuramoyl-tripeptide--D-alanyl-D-alanine ligase [EC:6.3.2.10] |
| ID526 | 7.3659768 | 0.02195233 | 17.86 | 7.00 | K00526 ribonucleoside-diphosphate reductase beta chain [EC:1.17.4.1] |

| | | | | | |
|---|---|---|---|---|---|
| ID1786 | 7.3684423 | 0.02195233 | 1.43 | 4.50 | K01786 L-ribulose-5-phosphate 4-epimerase [EC:5.1.3.4] |
| ID1442 | 7.3318966 | 0.02220562 | 3.00 | 10.42 | K01442 choloylglycine hydrolase [EC:3.5.1.24] |
| ID394 | 7.3201924 | 0.02226473 | 0.71 | 3.50 | K00394 adenylylsulfate reductase |
| ID2372 | 7.2929037 | 0.02251883 | 4.57 | 2.83 | K02372 3-hydroxyacyl-[acyl-carrier-protein] dehydratase [EC:4.2.1.59] |
| ID11071 | 7.1857536 | 0.0238124 | 1.86 | 3.83 | K11071 spermidine/putrescine transport system permease protein |
| ID527 | 7.1692547 | 0.02385033 | 12.57 | 27.50 | K00527 ribonucleoside-triphosphate reductase [EC:1.17.4.2] |
| ID748 | 7.1708302 | 0.02385033 | 3.14 | 1.42 | K00748 lipid-A-disaccharide synthase [EC:2.4.1.182] |
| ID11069 | 7.1594563 | 0.02389047 | 2.29 | 5.50 | K11069 spermidine/putrescine transport system substrate-binding protein |
| ID656 | 7.1062005 | 0.02451815 | 10.71 | 25.25 | K00656 formate C-acetyltransferase [EC:2.3.1.54] |
| ID336 | 7.0181935 | 0.02565621 | 6.14 | 2.83 | K00336 NADH-quinone oxidoreductase subunit G [EC:1.6.5.3] |
| ID1804 | 6.9931061 | 0.02592119 | 3.29 | 9.42 | K01804 L-arabinose isomerase [EC:5.3.1.4] |
| ID1192 | 6.9618321 | 0.02628009 | 1.00 | 4.17 | K01192 beta-mannosidase [EC:3.2.1.25] |
| ID845 | 6.9250316 | 0.02672699 | 3.57 | 7.83 | K00845 glucokinase [EC:2.7.1.2] |
| ID1950 | 6.9066952 | 0.02690283 | 5.29 | 12.00 | K01950 NAD+ synthase (glutamine-hydrolysing) [EC:6.3.5.1] |
| ID3407 | 6.8489086 | 0.02759513 | 7.00 | 3.08 | K03407 two-component system |
| ID6148 | 6.8481429 | 0.02759513 | 8.14 | 1.42 | K06148 ATP-binding cassette |
| ID226 | 6.8400786 | 0.02761881 | 0.43 | 5.00 | K00226 dihydroorotate dehydrogenase (fumarate) [EC:1.3.98.1] |
| ID2881 | 6.8236233 | 0.02777097 | 2.43 | 4.50 | K02881 large subunit ribosomal protein L18 |
| ID342 | 6.8172757 | 0.02777097 | 4.86 | 2.00 | K00342 NADH-quinone oxidoreductase subunit M [EC:1.6.5.3] |
| ID14 | 6.7875384 | 0.02806081 | 3.71 | 8.08 | K00014 shikimate dehydrogenase [EC:1.1.1.25] |
| ID2517 | 6.7794370 | 0.02806081 | 6.14 | 2.00 | K02517 lipid A biosynthesis lauroyl acyltransferase [EC:2.3.1.-] |
| ID11963 | 6.7794370 | 0.02806081 | 3.29 | 1.25 | K11963 urea transport system ATP-binding protein |
| ID4072 | 6.7212891 | 0.02888684 | 6.43 | 16.75 | K04072 acetaldehyde dehydrogenase / alcohol dehydrogenase [EC:1.2.1.10 1.1.1.1] |
| ID925 | 6.7093825 | 0.02897686 | 5.71 | 12.33 | K00925 acetate kinase [EC:2.7.2.1] |
| ID2557 | 6.5784273 | 0.03107643 | 4.14 | 1.67 | K02557 chemotaxis protein MotB |
| ID12410 | 6.5433673 | 0.03147156 | 3.43 | 7.58 | K12410 |
| ID1785 | 6.5492149 | 0.03147156 | 2.43 | 6.75 | K01785 aldose 1-epimerase [EC:5.1.3.3] |
| ID697 | 6.5297407 | 0.03160229 | 2.86 | 0.67 | K00697 trehalose 6-phosphate synthase [EC:2.4.1.15] |
| ID1209 | 6.5085004 | 0.03187016 | 3.86 | 13.00 | K01209 alpha-N-arabinofuranosidase [EC:3.2.1.55] |
| ID303 | 6.4875878 | 0.03213488 | 10.71 | 0.83 | K00303 sarcosine oxidase |
| ID975 | 6.4568911 | 0.03258121 | 8.14 | 15.92 | K00975 glucose-1-phosphate adenylyltransferase [EC:2.7.7.27] |
| ID3856 | 6.3840734 | 0.03382747 | 2.43 | 4.42 | K03856 3-deoxy-7-phosphoheptulonate synthase [EC:2.5.1.54] |
| ID57 | 6.3668656 | 0.0339223 | 5.00 | 10.08 | K00057 glycerol-3-phosphate dehydrogenase (NAD(P)+) [EC:1.1.1.94] |
| ID1686 | 6.3668656 | 0.0339223 | 2.71 | 6.00 | K01686 mannonate dehydratase [EC:4.2.1.8] |
| ID11754 | 6.2989524 | 0.03512573 | 4.29 | 10.33 | K11754 dihydrofolate synthase / folylpolyglutamate synthase [EC:6.3.2.12 6.3.2.17] |
| ID262 | 6.2766355 | 0.03520908 | 10.00 | 17.92 | K00262 glutamate dehydrogenase (NADP+) [EC:1.4.1.4] |
| ID876 | 6.2821998 | 0.03520908 | 3.57 | 8.08 | K00876 uridine kinase [EC:2.7.1.48] |
| ID41 | 6.2834646 | 0.03520908 | 1.00 | 3.92 | K00041 tagaturonate reductase [EC:1.1.1.58] |
| ID6861 | 6.2597284 | 0.03542657 | 4.00 | 2.00 | K06861 lipopolysaccharide export system ATP-binding protein [EC:3.6.3.-] |
| ID5343 | 6.2142857 | 0.03610343 | 13.14 | 5.58 | K05343 maltose alpha-D-glucosyltransferase/ alpha-amylase [EC:5.4.99.16 3.2.1.1] |
| ID12368 | 6.2146630 | 0.03610343 | 10.86 | 0.92 | K12368 dipeptide transport system substrate-binding protein |
| ID1251 | 6.1639450 | 0.03702097 | 11.57 | 4.42 | K01251 adenosylhomocysteinase [EC:3.3.1.1] |
| ID290 | 6.1528687 | 0.03712935 | 2.29 | 6.50 | K00290 saccharopine dehydrogenase (NAD+ |
| ID145 | 6.1363287 | 0.03735374 | 5.29 | 9.17 | K00145 N-acetyl-gamma-glutamyl-phosphate reductase [EC:1.2.1.38] |
| ID2047 | 6.1149425 | 0.03768317 | 3.29 | 1.50 | K02047 sulfate transport system permease protein |
| ID1834 | 6.0970651 | 0.03769187 | 1.29 | 10.08 | K01834 2 |
| ID11962 | 6.1015090 | 0.03769187 | 4.14 | 1.42 | K11962 urea transport system ATP-binding protein |
| ID3394 | 6.1071429 | 0.03769187 | 2.71 | 1.17 | K03394 precorrin-2/cobalt-factor-2 C20-methyltransferase [EC:2.1.1.130 2.1.1.151] |
| ID5846 | 6.0519816 | 0.03853988 | 5.43 | 1.17 | K05846 osmoprotectant transport system permease protein |
| ID1478 | 6.0267857 | 0.03896653 | 4.57 | 1.17 | K01478 arginine deiminase [EC:3.5.3.6] |
| ID677 | 5.9887430 | 0.0396865 | 4.57 | 1.92 | K00677 UDP-N-acetylglucosamine acyltransferase [EC:2.3.1.129] |
| ID831 | 5.8948625 | 0.04172221 | 5.71 | 10.42 | K00831 phosphoserine aminotransferase [EC:2.6.1.52] |

| | | | | | |
|---|---|---|---|---|---|
| ID1902 | 5.8592157 | 0.04243822 | 7.86 | 2.58 | K01902 succinyl-CoA synthetase alpha subunit [EC:6.2.1.5] |
| ID2876 | 5.8301370 | 0.04300662 | 3.57 | 5.83 | K02876 large subunit ribosomal protein L15 |
| ID819 | 5.7639491 | 0.04451336 | 2.71 | 0.75 | K00819 ornithine--oxo-acid transaminase [EC:2.6.1.13] |
| ID1775 | 5.6797153 | 0.04655165 | 6.29 | 9.58 | K01775 alanine racemase [EC:5.1.1.1] |
| ID1428 | 5.6338142 | 0.04763392 | 6.00 | 2.92 | K01428 urease subunit alpha [EC:3.5.1.5] |
| ID688 | 5.6197183 | 0.04786609 | 19.00 | 35.08 | K00688 starch phosphorylase [EC:2.4.1.1] |
| ID798 | 5.5803571 | 0.04879874 | 3.57 | 1.33 | K00798 cob(I)alamin adenosyltransferase [EC:2.5.1.17] |

**Supplementary Table 9.** Enrichment and depletion of KO metabolic gene categories in the ancient and modern comparative IM sample cohort as calculated for the twenty-four authenticated ancient IM taxa.

| Taxa KO genes | BRS1 | BRS2 | BRS3 | BRS4 | BRS5 | Otzi_LPLI | Otzi_Small | Otzi_UPLI2 | Otzi_UPLI | Hadza_H11 | Hadza_H15 | Hadza_H17 | Hadza_H9 | Italian_IT2 | Italian_IT5 | Italian_IT6 | Italian_IT8 | Malawi_4461146 | Malawi_4461161 | Malawi_4461162 | Malawi_4461165 |
|---|---|---|---|---|---|---|---|---|---|---|---|---|---|---|---|---|---|---|---|---|---|
| Alistipes:K00029 | 0 | 0 | 0 | 0.0016 | 0 | 0 | 0 | 0 | 0 | 0 | 0.000477 | 0.0003415 | 0.002235 | 0.004593 | 0.001453 | 0.009139 | 0.000987 | 0 | 0 | 0 | 0 |
| Alistipes:K00111 | 0 | 0 | 0 | 0.0016 | 0 | 0 | 0 | 0 | 0 | 0 | 0 | 0 | 0 | 0.002923 | 0.003529 | 0.010553 | 0 | 0 | 0 | 0 | 0 |
| Alistipes:K00128 | 0 | 0.0205 | 0 | 0.0113 | 0 | 0 | 0 | 0 | 0 | 0.0149858 | 0.01575 | 0.0046673 | 0.001118 | 0.000835 | 0.000311 | 0.002067 | 0 | 0 | 0 | 0 | 0 |
| Alistipes:K00248 | 0 | 0 | 0 | 0 | 0 | 0 | 0 | 0 | 0 | 0 | 0 | 0 | 0 | 0.000418 | 0.00083 | 0.002829 | 0 | 0 | 0 | 0 | 0 |
| Alistipes:K00257 | 0 | 0 | 0 | 0 | 0 | 0 | 0 | 0 | 0 | 0 | 0 | 0 | 0 | 0.000418 | 0 | 0.000109 | 0 | 0 | 0 | 0 | 0 |
| Alistipes:K00281 | 0 | 0 | 0 | 0.0226 | 0 | 0 | 0 | 0 | 0 | 0 | 0.001432 | 0.0003415 | 0 | 0.011692 | 0.004463 | 0.026002 | 0 | 0 | 0 | 0 | 0 |
| Alistipes:K00331 | 0 | 0 | 0 | 0 | 0 | 0 | 0 | 0 | 0 | 0 | 0 | 0.0004553 | 0.001118 | 0.002923 | 0.007059 | 0.011097 | 0 | 0 | 0 | 0 | 0 |
| Alistipes:K00334 | 0 | 0 | 0 | 0 | 0 | 0 | 0 | 0 | 0 | 0 | 0 | 0 | 0 | 0 | 0 | 0.000109 | 0 | 0 | 0 | 0 | 0 |
| Alistipes:K00335 | 0 | 0 | 0 | 0 | 0 | 0 | 0 | 0 | 0 | 0 | 0 | 0 | 0 | 0.000418 | 0.000415 | 0.001523 | 0 | 0 | 0 | 0 | 0 |
| Alistipes:K00336 | 0 | 0 | 0 | 0 | 0 | 0 | 0 | 0 | 0 | 0 | 0 | 0 | 0 | 0 | 0.000208 | 0.000109 | 0 | 0 | 0 | 0 | 0 |
| Alistipes:K00337 | 0 | 0 | 0 | 0.0032 | 0 | 0 | 0 | 0 | 0 | 0.0010704 | 0.000477 | 0 | 0.001118 | 0.007099 | 0.007993 | 0.019039 | 0.000987 | 0 | 0 | 0 | 0 |
| Alistipes:K00338 | 0 | 0 | 0 | 0 | 0 | 0 | 0 | 0 | 0 | 0 | 0 | 0 | 0 | 0.002506 | 0.002076 | 0.006093 | 0 | 0 | 0 | 0 | 0 |
| Alistipes:K00341 | 0 | 0.0205 | 0 | 0 | 0 | 0 | 0 | 0 | 0 | 0.0021408 | 0 | 0.0004553 | 0 | 0.007934 | 0.00436 | 0.014796 | 0.001974 | 0 | 0 | 0 | 0 |
| Alistipes:K00342 | 0 | 0 | 0 | 0 | 0 | 0 | 0 | 0 | 0 | 0 | 0.000477 | 0.0005692 | 0.002235 | 0.005011 | 0.006124 | 0.019257 | 0.001974 | 0 | 0 | 0 | 0 |
| Alistipes:K00390 | 0 | 0 | 0 | 0 | 0 | 0 | 0 | 0 | 0 | 0 | 0 | 0 | 0 | 0.00167 | 0.006124 | 0.005331 | 0 | 0 | 0 | 0 | 0 |
| Alistipes:K00425 | 0 | 0 | 0 | 0 | 0 | 0 | 0 | 0 | 0 | 0.0010704 | 0 | 0 | 0 | 0.007099 | 0.003322 | 0.009792 | 0 | 0 | 0 | 0 | 0 |
| Alistipes:K00426 | 0 | 0 | 0 | 0 | 0 | 0 | 0 | 0 | 0 | 0.0010704 | 0.000955 | 0 | 0.001118 | 0.001253 | 0.001453 | 0.006201 | 0 | 0 | 0 | 0 | 0 |
| Alistipes:K00525 | 0 | 0.0205 | 0 | 0.0226 | 0 | 0 | 0 | 0 | 0 | 0 | 0 | 0 | 0.001118 | 0.022132 | 0.018477 | 0.039384 | 0 | 0 | 0 | 0 | 0 |
| Alistipes:K00548 | 0 | 0 | 0 | 0.0016 | 0 | 4.15E-05 | 0 | 0 | 0 | 0.0053521 | 0.007159 | 0.0001138 | 0 | 0.009187 | 0.003114 | 0.02796 | 0.000987 | 0 | 0 | 0.021473051 | 0 |
| Alistipes:K00600 | 0 | 0.0205 | 0.0356 | 0.0226 | 0 | 0 | 0 | 0 | 0 | 0.0096337 | 0.009545 | 0.0036428 | 0 | 0.012528 | 0.008304 | 0.020018 | 0.001974 | 0 | 0 | 0 | 0 |
| Alistipes:K00627 | 0 | 0 | 0 | 0 | 0 | 0 | 0 | 0 | 0 | 0 | 0.000955 | 0 | 0 | 0.000418 | 0 | 0 | 0 | 0 | 0 | 0 | 0 |
| Alistipes:K00640 | 0 | 0 | 0 | 0 | 0 | 0 | 0 | 0 | 0 | 0 | 0 | 0 | 0 | 0.000835 | 0.000104 | 0.000109 | 0 | 0 | 0 | 0 | 0 |
| Alistipes:K00641 | 0 | 0 | 0 | 0.0016 | 0 | 0 | 0 | 0 | 0 | 0.0053521 | 0.003341 | 0.0001138 | 0.002235 | 0.003341 | 0.001868 | 0.010771 | 0.000987 | 0 | 0 | 0.021473051 | 0 |
| Alistipes:K00647 | 0 | 0 | 0 | 0 | 0 | 0 | 0 | 0 | 0 | 0.0021408 | 0 | 0.0001138 | 0.001118 | 0.003341 | 0.001661 | 0.014579 | 0 | 0 | 0 | 0 | 0 |
| Alistipes:K00648 | 0 | 0 | 0 | 0 | 0 | 0 | 0 | 0 | 0 | 0 | 0.000477 | 0 | 0 | 0.006264 | 0.00436 | 0.010009 | 0 | 0 | 0 | 0 | 0 |
| Alistipes:K00677 | 0 | 0 | 0 | 0.0048 | 0 | 0 | 0 | 0 | 0 | 0 | 0 | 0 | 0 | 0.004176 | 0.002491 | 0.010118 | 0 | 0 | 0 | 0 | 0 |
| Alistipes:K00748 | 0 | 0 | 0 | 0.0032 | 0 | 0 | 0 | 0 | 0 | 0 | 0 | 0 | 0 | 0.002506 | 0.002076 | 0.009248 | 0.000987 | 0 | 0 | 0 | 0 |
| Alistipes:K00798 | 0 | 0 | 0 | 0 | 0 | 0 | 0 | 0 | 0 | 0 | 0.002864 | 0.0001138 | 0.001118 | 0.000418 | 0.001868 | 0.003699 | 0 | 0 | 0 | 0 | 0 |
| Alistipes:K00833 | 0 | 0 | 0 | 0 | 0 | 0 | 0 | 0 | 0 | 0 | 0 | 0 | 0 | 0.00167 | 0 | 0 | 0 | 0 | 0 | 0 | 0 |
| Alistipes:K00864 | 0 | 0 | 0 | 0 | 0 | 0 | 0 | 0 | 0 | 0 | 0.000477 | 0.0001138 | 0 | 0.000835 | 0.000104 | 0.000544 | 0 | 0 | 0 | 0 | 0 |
| Alistipes:K01142 | 0 | 0 | 0 | 0 | 0 | 0 | 0 | 0 | 0 | 0 | 0 | 0 | 0 | 0.002506 | 0.000415 | 0.00272 | 0 | 0 | 0 | 0 | 0 |
| Alistipes:K01251 | 0 | 0 | 0.0356 | 0.0016 | 0 | 0 | 0 | 0 | 0 | 0.0010704 | 0.00525 | 0.0004553 | 0 | 0.008769 | 0.006747 | 0.022629 | 0.000987 | 0 | 0 | 0 | 0 |
| Alistipes:K01338 | 0 | 0.0205 | 0 | 0.0129 | 0 | 0 | 0 | 0 | 0 | 0 | 0.000955 | 0.0001138 | 0.001118 | 0.019209 | 0.007059 | 0.028613 | 0.000987 | 0 | 0 | 0 | 0 |
| Alistipes:K01428 | 0 | 0 | 0 | 0.0016 | 0 | 0 | 0 | 0 | 0 | 0 | 0 | 0 | 0 | 0.000418 | 0.001453 | 0.004678 | 0 | 0 | 0 | 0 | 0 |
| Alistipes:K01585 | 0 | 0 | 0 | 0.0048 | 0 | 4.15E-05 | 0 | 0 | 0 | 0.0010704 | 0.004773 | 0.0012522 | 0 | 0.021297 | 0.025224 | 0.025893 | 0 | 0 | 0 | 0 | 0 |
| Alistipes:K01613 | 0 | 0.0205 | 0 | 0.0032 | 0 | 0 | 0 | 0 | 0 | 0 | 0 | 0 | 0.001118 | 0.005429 | 0.002803 | 0.008377 | 0.000987 | 0 | 0 | 0 | 0 |
| Alistipes:K01627 | 0 | 0 | 0 | 0.0016 | 0 | 0 | 0 | 0 | 0 | 0 | 0 | 0.0004553 | 0 | 0.002506 | 0.001453 | 0.006963 | 0 | 0 | 0 | 0 | 0 |
| Alistipes:K01676 | 0 | 0 | 0.0356 | 0.021 | 0 | 0 | 0 | 0 | 0 | 0 | 0 | 0 | 0 | 0.01211 | 0.005917 | 0.020998 | 0 | 0 | 0 | 0 | 0 |
| Alistipes:K01712 | 0 | 0.0205 | 0 | 0.0032 | 0 | 0 | 0 | 0 | 0 | 0.0171266 | 0.009545 | 0.0070579 | 0.006706 | 0.002923 | 0.002595 | 0.011532 | 0.001974 | 0 | 0 | 0 | 0 |
| Alistipes:K01737 | 0 | 0 | 0 | 0.0032 | 0 | 0 | 0 | 0 | 0 | 0 | 0 | 0 | 0 | 0.001253 | 0.000623 | 0.002285 | 0 | 0 | 0 | 0 | 0 |
| Alistipes:K01745 | 0 | 0.0409 | 0.0356 | 0.0016 | 0 | 0 | 0 | 0 | 0 | 0.0321124 | 0.011932 | 0.0060333 | 0.014529 | 0.010857 | 0.005398 | 0.022412 | 0.000987 | 0 | 0 | 0 | 0 |
| Alistipes:K01895 | 0 | 0 | 0 | 0 | 0 | 4.15E-05 | 0 | 0 | 0 | 0 | 0 | 0.0001138 | 0.002235 | 0.00167 | 0.00083 | 0.005113 | 0 | 0 | 0 | 0 | 0 |
| Alistipes:K01915 | 0 | 0.0409 | 0 | 0.0065 | 0 | 0 | 0 | 0 | 0 | 0.0021408 | 0.000477 | 0.0002277 | 0.001118 | 0.016703 | 0.009757 | 0.030463 | 0.001974 | 0 | 0 | 0 | 0 |
| Alistipes:K01960 | 0 | 0 | 0 | 0.0129 | 0 | 0 | 0 | 0 | 0 | 0 | 0.000477 | 0.0001138 | 0 | 0.014615 | 0.00737 | 0.015667 | 0.000987 | 0 | 0 | 0 | 0 |
| Alistipes:K02372 | 0 | 0 | 0 | 0 | 0 | 0 | 0 | 0 | 0 | 0 | 0 | 0 | 0 | 0 | 0.000104 | 0.001306 | 0 | 0 | 0 | 0 | 0 |
| Alistipes:K02454 | 0 | 0 | 0 | 0 | 0 | 0 | 0 | 0 | 0 | 0 | 0 | 0 | 0 | 0 | 0 | 0.000109 | 0 | 0 | 0 | 0 | 0 |
| Alistipes:K02517 | 0 | 0 | 0 | 0.0016 | 0 | 0 | 0 | 0 | 0 | 0 | 0 | 0 | 0 | 0.005011 | 0.002284 | 0.008051 | 0 | 0 | 0 | 0 | 0 |
| Alistipes:K02523 | 0 | 0 | 0 | 0.0016 | 0 | 0 | 0 | 0 | 0 | 0 | 0 | 0 | 0 | 0.00167 | 0.001246 | 0.008051 | 0 | 0 | 0 | 0 | 0 |
| Alistipes:K03092 | 0 | 0 | 0 | 0.0065 | 0 | 0 | 0 | 0 | 0 | 0 | 0 | 0 | 0 | 0.005429 | 0.005709 | 0.011206 | 0 | 0 | 0 | 0 | 0 |
| Alistipes:K03182 | 0 | 0 | 0 | 0.0048 | 0 | 0 | 0 | 0 | 0 | 0 | 0 | 0 | 0 | 0.003341 | 0.001038 | 0.010118 | 0 | 0 | 0 | 0 | 0 |
| Alistipes:K03183 | 0 | 0 | 0 | 0 | 0 | 0 | 0 | 0 | 0 | 0 | 0 | 0 | 0 | 0.000835 | 0.00083 | 0.006637 | 0.000987 | 0 | 0 | 0 | 0 |
| Alistipes:K03590 | 0 | 0 | 0 | 0 | 0 | 0 | 0 | 0 | 0 | 0 | 0 | 0 | 0 | 0.00167 | 0.001661 | 0.008595 | 0.000987 | 0 | 0 | 0 | 0 |
| Alistipes:K03644 | 0 | 0 | 0 | 0.0065 | 0 | 0 | 0 | 0 | 0 | 0 | 0 | 0 | 0 | 0.005846 | 0.002595 | 0.013599 | 0.000987 | 0 | 0 | 0 | 0 |
| Alistipes:K03781 | 0 | 0 | 0 | 0.0097 | 0 | 0 | 0 | 0 | 0 | 0 | 0 | 0 | 0 | 0.004176 | 0.002906 | 0.010444 | 0.000987 | 0 | 0 | 0 | 0 |
| Alistipes:K04564 | 0 | 0 | 0 | 0.0016 | 0 | 0 | 0 | 0 | 0 | 0 | 0 | 0.0002277 | 0.001118 | 0.005011 | 0.001765 | 0.00544 | 0 | 0 | 0 | 0 | 0 |
| Alistipes:K05343 | 0 | 0 | 0 | 0 | 0 | 0 | 0 | 0 | 0 | 0 | 0 | 0 | 0 | 0.00167 | 0 | 0.003264 | 0 | 0 | 0 | 0 | 0 |
| Alistipes:K05592 | 0 | 0 | 0 | 0 | 0 | 0.00017 | 0 | 0.0030776 | 0.001025 | 0 | 0 | 0 | 0.001118 | 0.007099 | 0.003322 | 0.014035 | 0.000987 | 0 | 0 | 0 | 0 |

| | | | | | | | | | | | | | | | | | | | | | |
|---|---|---|---|---|---|---|---|---|---|---|---|---|---|---|---|---|---|---|---|---|---|
| Alistipes:K06861 | 0 | 0.0205 | 0 | 0.0081 | 0 | 0 | 0 | 0 | 0 | 0 | 0 | 0 | 0 | 0.008352 | 0.003322 | 0.010009 | 0.000987 | 0 | 0 | 0 | 0 |
| Alistipes:K11927 | 0 | 0 | 0 | 0 | 0 | 0 | 0 | 0 | 0 | 0 | 0 | 0 | 0 | 0.004176 | 0.00083 | 0.007072 | 0 | 0 | 0 | 0 | 0 |
| Alistipes:K12339 | 0 | 0 | 0.0356 | 0.0032 | 0 | 0 | 0 | 0 | 0 | 0.0053521 | 0.000955 | 0 | 0 | 0.002088 | 0.002076 | 0.006528 | 0 | 0 | 0 | 0 | 0 |
| Alistipes:K12573 | 0 | 0 | 0 | 0.0032 | 0 | 0 | 0 | 0 | 0 | 0 | 0.000477 | 0.0004553 | 0.001118 | 0.008769 | 0.003425 | 0.021106 | 0 | 0 | 0 | 0 | 0 |
| Alistipes:K13378 | 0 | 0 | 0 | 0 | 0 | 0 | 0 | 0 | 0 | 0 | 0.000955 | 0 | 0 | 0.004176 | 0.007578 | 0.018604 | 0.000987 | 0 | 0 | 0 | 0 |
| Bacteroides:K00029 | 0 | 0.0614 | 0.0356 | 0.0275 | 0 | 4.15E-05 | 0 | 0 | 0 | 0.0160562 | 0.019568 | 0.0165063 | 0.031293 | 0.016703 | 0.150513 | 0.102159 | 0.009872 | 0 | 0 | 0 | 0 |
| Bacteroides:K00036 | 0 | 0 | 0 | 0 | 0 | 0 | 0 | 0 | 0 | 0 | 0 | 0 | 0.002235 | 0.005011 | 0.048891 | 0.023065 | 0.000987 | 0 | 0 | 0 | 0 |
| Bacteroides:K00111 | 0 | 0 | 0 | 0 | 0 | 0 | 0 | 0 | 0 | 0.0010704 | 0 | 0 | 0 | 0 | 0.000208 | 0 | 0 | 0 | 0 | 0 | 0 |
| Bacteroides:K00257 | 0 | 0 | 0 | 0 | 0 | 8.29E-05 | 0 | 0 | 0 | 0.0107041 | 0.004295 | 0.0023906 | 0.011176 | 0.004176 | 0.005813 | 0.007072 | 0 | 0 | 0 | 0 | 0 |
| Bacteroides:K00281 | 0 | 0 | 0 | 0.0275 | 0 | 0 | 0 | 0 | 0 | 0.0010704 | 0.001432 | 0.00979 | 0.029058 | 0.007517 | 0.127573 | 0.081488 | 0.005923 | 0.0373413 | 0 | 0 | 0 |
| Bacteroides:K00331 | 0 | 0 | 0 | 0.0048 | 0 | 4.15E-05 | 0 | 0 | 0 | 0.0021408 | 0.001909 | 0.0027321 | 0.016764 | 0.004176 | 0.025951 | 0.019583 | 0.000987 | 0 | 0 | 0 | 0 |
| Bacteroides:K00334 | 0 | 0 | 0 | 0 | 0 | 0 | 0 | 0 | 0 | 0 | 0 | 0 | 0 | 0.000418 | 0.001972 | 0.000979 | 0.000987 | 0 | 0 | 0 | 0 |
| Bacteroides:K00335 | 0 | 0 | 0 | 0 | 0 | 0 | 0 | 0 | 0 | 0 | 0 | 0 | 0 | 0.004593 | 0.005086 | 0.00544 | 0 | 0 | 0 | 0 | 0 |
| Bacteroides:K00336 | 0 | 0 | 0 | 0 | 0 | 0 | 0 | 0 | 0 | 0 | 0.000477 | 0 | 0.002235 | 0.006264 | 0.005086 | 0.011315 | 0 | 0 | 0 | 0 | 0 |
| Bacteroides:K00337 | 0 | 0.0409 | 0 | 0.0065 | 0 | 0 | 0 | 0 | 0 | 0.0021408 | 0.002864 | 0.0019352 | 0.011176 | 0.005011 | 0.063319 | 0.028613 | 0.003949 | 0 | 0 | 0 | 0 |
| Bacteroides:K00338 | 0 | 0 | 0 | 0.0113 | 0 | 0 | 0 | 0 | 0 | 0.0021408 | 0.000955 | 0.0015937 | 0.003353 | 0.002506 | 0.028234 | 0.017407 | 0.001974 | 0 | 0 | 0 | 0 |
| Bacteroides:K00341 | 0 | 0 | 0 | 0.0129 | 0 | 0 | 0 | 0 | 0 | 0.0139154 | 0.004295 | 0.0124082 | 0.014529 | 0.007517 | 0.05896 | 0.059076 | 0.001974 | 0 | 0 | 0 | 0 |
| Bacteroides:K00342 | 0 | 0 | 0 | 0.0129 | 0 | 0 | 0 | 0 | 0 | 0.0064225 | 0.007636 | 0.0130912 | 0.018999 | 0.007099 | 0.102557 | 0.060164 | 0.004936 | 0 | 0 | 0 | 0 |
| Bacteroides:K00390 | 0 | 0 | 0 | 0 | 0 | 0 | 0 | 0 | 0 | 0 | 0 | 0.0001138 | 0.001118 | 0 | 0.000311 | 0.001958 | 0 | 0 | 0 | 0 | 0 |
| Bacteroides:K00425 | 0 | 0 | 0 | 0.0016 | 0 | 0 | 0 | 0 | 0 | 0.0032112 | 0.002386 | 0.006261 | 0.007823 | 0.009604 | 0.035812 | 0.034815 | 0 | 0 | 0 | 0 | 0 |
| Bacteroides:K00426 | 0 | 0 | 0 | 0 | 0 | 4.15E-05 | 0 | 0 | 0 | 0.0042816 | 0.016705 | 0.005578 | 0.002235 | 0.003341 | 0.043805 | 0.025567 | 0.000987 | 0 | 0 | 0 | 0 |
| Bacteroides:K00432 | 0 | 0 | 0 | 0 | 0 | 0 | 0.002207 | 0 | 0 | 0 | 0 | 0 | 0.001118 | 0.001253 | 0.033424 | 0.003373 | 0 | 0 | 0 | 0 | 0.007893906 |
| Bacteroides:K00525 | 0 | 0.0409 | 0.1426 | 0.0307 | 0 | 4.15E-05 | 0 | 0 | 0 | 0.0160562 | 0.008591 | 0.0159371 | 0.020117 | 0.01211 | 0.163177 | 0.064842 | 0.004936 | 0 | 0 | 0.021473051 | 0 |
| Bacteroides:K00526 | 0 | 0 | 0 | 0 | 0 | 0 | 0 | 0 | 0 | 0.0010704 | 0.000955 | 0.0002277 | 0.001118 | 0.004176 | 0.044843 | 0.010771 | 0.000987 | 0 | 0 | 0 | 0 |
| Bacteroides:K00548 | 0 | 0 | 0 | 0.0129 | 0 | 0 | 0 | 0 | 0 | 0.0010704 | 0.018136 | 0.011839 | 0.005588 | 0.00167 | 0.011107 | 0.013273 | 0.002962 | 0 | 0 | 0 | 0 |
| Bacteroides:K00600 | 0 | 0.0205 | 0.0356 | 0.0113 | 0 | 4.15E-05 | 0 | 0 | 0 | 0.0010704 | 0.006682 | 0.0079686 | 0.010058 | 0.012945 | 0.062697 | 0.038405 | 0.00691 | 0.0373413 | 0 | 0.021473051 | 0 |
| Bacteroides:K00640 | 0 | 0 | 0 | 0.0048 | 0 | 0 | 0 | 0 | 0 | 0.0032112 | 0.001909 | 0.0081962 | 0.003353 | 0.007099 | 0.056884 | 0.027852 | 0.000987 | 0.0373413 | 0 | 0 | 0.015787812 |
| Bacteroides:K00647 | 0 | 0 | 0 | 0.0016 | 0 | 0 | 0 | 0 | 0 | 0.0021408 | 0.000477 | 0.0005692 | 0.00447 | 0.003758 | 0.032179 | 0.023391 | 0 | 0 | 0 | 0 | 0 |
| Bacteroides:K00648 | 0 | 0 | 0.0713 | 0.0081 | 0 | 0 | 0 | 0 | 0 | 0 | 0.001909 | 0.0018214 | 0.006706 | 0.005011 | 0.082419 | 0.030245 | 0.003949 | 0 | 0 | 0 | 0.007893906 |
| Bacteroides:K00658 | 0 | 0 | 0 | 0 | 0 | 0 | 0 | 0 | 0 | 0 | 0 | 0 | 0 | 0.000835 | 0.002595 | 0.000762 | 0 | 0 | 0 | 0 | 0 |
| Bacteroides:K00677 | 0 | 0 | 0 | 0.0065 | 0 | 0 | 0 | 0 | 0 | 0.0074929 | 0.001909 | 0.0053503 | 0.013411 | 0.008769 | 0.095913 | 0.04428 | 0.00691 | 0 | 0 | 0.042946103 | 0 |
| Bacteroides:K00748 | 0 | 0 | 0 | 0.0016 | 0 | 0 | 0 | 0 | 0 | 0.0042816 | 0.002386 | 0.006261 | 0.008941 | 0.004176 | 0.063942 | 0.029484 | 0.004936 | 0 | 0 | 0 | 0 |
| Bacteroides:K00798 | 0 | 0 | 0 | 0 | 0 | 0 | 0 | 0 | 0 | 0 | 0 | 0 | 0 | 0.000418 | 0 | 0 | 0 | 0 | 0 | 0 | 0 |
| Bacteroides:K00819 | 0 | 0.0205 | 0 | 0 | 0 | 0 | 0 | 0 | 0 | 0 | 0 | 0 | 0.002235 | 0.004176 | 0.07671 | 0.024914 | 0 | 0 | 0 | 0 | 0 |
| Bacteroides:K00833 | 0 | 0 | 0 | 0 | 0 | 4.15E-05 | 0 | 0 | 0 | 0.0021408 | 0.001909 | 0.0003415 | 0.003353 | 0.007099 | 0.039341 | 0.026981 | 0.003949 | 0 | 0 | 0.021473051 | 0 |
| Bacteroides:K00864 | 0 | 0 | 0 | 0 | 0 | 0 | 0 | 0 | 0 | 0.0021408 | 0.014318 | 0.004212 | 0.002235 | 0.002506 | 0.043805 | 0.016428 | 0.000987 | 0 | 0 | 0 | 0 |
| Bacteroides:K00940 | 0 | 0.0614 | 0.0356 | 0.0081 | 0 | 0 | 0 | 0 | 0 | 0.0053521 | 0.005727 | 0.004212 | 0.017882 | 0.000418 | 0.028649 | 0.009356 | 0.000987 | 0 | 0 | 0.021473051 | 0 |
| Bacteroides:K00955 | 0 | 0 | 0 | 0 | 0 | 0 | 0 | 0 | 0 | 0 | 0 | 0.0004553 | 0.001118 | 0 | 0.000727 | 0.000653 | 0 | 0 | 0 | 0 | 0 |
| Bacteroides:K01113 | 0 | 0 | 0 | 0 | 0 | 0 | 0 | 0 | 0 | 0 | 0 | 0.0001138 | 0 | 0 | 0 | 0 | 0 | 0 | 0 | 0 | 0 |
| Bacteroides:K01142 | 0 | 0 | 0.0713 | 0.0032 | 0 | 4.15E-05 | 0 | 0 | 0 | 0.0010704 | 0.001909 | 0.0027321 | 0.001118 | 0.011275 | 0.008097 | 0.008595 | 0.001974 | 0 | 0 | 0 | 0 |
| Bacteroides:K01241 | 0 | 0.0205 | 0 | 0.0032 | 0 | 0 | 0 | 0 | 0 | 0.0010704 | 0.000477 | 0.0020491 | 0.005588 | 0.002923 | 0.037057 | 0.01719 | 0 | 0 | 0 | 0 | 0 |
| Bacteroides:K01251 | 0 | 0.0205 | 0 | 0 | 0 | 0 | 0 | 0 | 0 | 0 | 0 | 0 | 0.002235 | 0.001253 | 0.004048 | 0.002394 | 0 | 0 | 0 | 0.021473051 | 0 |
| Bacteroides:K01338 | 0 | 0.0205 | 0.0356 | 0.0162 | 0 | 4.15E-05 | 0 | 0 | 0 | 0.0096337 | 0.004295 | 0.0359724 | 0.015646 | 0.008352 | 0.143351 | 0.052113 | 0.005923 | 0 | 0 | 0.021473051 | 0 |
| Bacteroides:K01433 | 0 | 0.0205 | 0 | 0 | 0 | 0 | 0 | 0 | 0 | 0 | 0.000477 | 0 | 0.002235 | 0.002088 | 0.055742 | 0.010553 | 0.002962 | 0 | 0 | 0 | 0 |
| Bacteroides:K01585 | 0 | 0 | 0 | 0 | 0 | 4.15E-05 | 0 | 0 | 0 | 0.0021408 | 0.004295 | 0.0047811 | 0.015646 | 0.019209 | 0.140029 | 0.086275 | 0.007897 | 0 | 0 | 0 | 0 |
| Bacteroides:K01613 | 0 | 0 | 0 | 0.0048 | 0 | 0 | 0 | 0 | 0 | 0.0010704 | 0 | 0.0025044 | 0.002235 | 0.004593 | 0.034151 | 0.016646 | 0.000987 | 0 | 0 | 0 | 0 |
| Bacteroides:K01627 | 0 | 0 | 0.0356 | 0.0032 | 0 | 0 | 0 | 0 | 0 | 0.0021408 | 0.001432 | 0.0014799 | 0.005588 | 0.004176 | 0.043286 | 0.018169 | 0 | 0 | 0 | 0 | 0 |
| Bacteroides:K01629 | 0 | 0.0205 | 0 | 0.0065 | 0 | 4.15E-05 | 0 | 0 | 0 | 0.0032112 | 0.007159 | 0.0026182 | 0.011176 | 0.003341 | 0.060828 | 0.029701 | 0 | 0 | 0 | 0 | 0 |
| Bacteroides:K01676 | 0 | 0.0409 | 0 | 0.0323 | 0 | 4.15E-05 | 0 | 0 | 0.001025 | 0.0235491 | 0.015273 | 0.006944 | 0.025705 | 0.01378 | 0.089374 | 0.063428 | 0.000987 | 0 | 0 | 0 | 0 |
| Bacteroides:K01712 | 0 | 0 | 0 | 0.0097 | 0 | 0.00012 | 0.002207 | 0.0023082 | 0.001025 | 0.0010704 | 0.000477 | 0.0003415 | 0.007823 | 0.007517 | 0.090308 | 0.060055 | 0.005923 | 0 | 0 | 0 | 0 |
| Bacteroides:K01737 | 0 | 0 | 0 | 0 | 0 | 0 | 0 | 0 | 0 | 0.0021408 | 0.001909 | 0.0003415 | 0.002235 | 0.000835 | 0.019619 | 0.008486 | 0 | 0 | 0 | 0 | 0 |
| Bacteroides:K01745 | 0 | 0.0205 | 0 | 0.0048 | 0 | 0 | 0 | 0 | 0 | 0.0042816 | 0.003818 | 0.0003415 | 0.010058 | 0.01378 | 0.093733 | 0.057988 | 0.001974 | 0 | 0 | 0 | 0 |
| Bacteroides:K01845 | 0 | 0 | 0 | 0 | 0 | 4.15E-05 | 0 | 0 | 0 | 0 | 0 | 0 | 0 | 0.000835 | 0 | 0 | 0 | 0 | 0 | 0 | 0.007893906 |
| Bacteroides:K01895 | 0 | 0 | 0 | 0.0081 | 0 | 0 | 0 | 0 | 0 | 0.0032112 | 0.00525 | 0.0025044 | 0.016764 | 0.008769 | 0.128507 | 0.058097 | 0.002962 | 0 | 0 | 0.021473051 | 0 |
| Bacteroides:K01902 | 0 | 0 | 0 | 0 | 0 | 0 | 0 | 0 | 0 | 0 | 0 | 0 | 0 | 0.000418 | 0.002076 | 0.000326 | 0 | 0 | 0 | 0 | 0 |
| Bacteroides:K01903 | 0 | 0 | 0 | 0 | 0 | 0 | 0 | 0 | 0 | 0 | 0 | 0 | 0.001118 | 0 | 0.002906 | 0.000435 | 0 | 0 | 0 | 0 | 0 |
| Bacteroides:K01915 | 0 | 0.1228 | 0.0356 | 0.0549 | 0 | 0 | 0 | 0 | 0 | 0.0278307 | 0.029591 | 0.0392737 | 0.05141 | 0.034659 | 0.252758 | 0.171353 | 0.014808 | 0 | 0 | 0 | 0 |
| Bacteroides:K01960 | 0 | 0 | 0.0713 | 0.0145 | 0 | 4.15E-05 | 0 | 0 | 0 | 0.0064225 | 0.002386 | 0.0067164 | 0.013411 | 0.014198 | 0.096536 | 0.058532 | 0.001974 | 0 | 0 | 0 | 0.007893906 |
| Bacteroides:K01961 | 0 | 0 | 0 | 0 | 0 | 0.00012 | 0 | 0 | 0 | 0.0021408 | 0.004773 | 0.0063749 | 0.017882 | 0.015451 | 0.119995 | 0.086057 | 0.000987 | 0 | 0 | 0 | 0 |
| Bacteroides:K02000 | 0 | 0 | 0 | 0 | 0 | 0 | 0 | 0 | 0 | 0 | 0 | 0 | 0.003353 | 0.007517 | 0.064461 | 0.02437 | 0 | 0 | 0 | 0.021473051 | 0.007893906 |

| | | | | | | | | | | | | | | | | | | | | | |
|---|---|---|---|---|---|---|---|---|---|---|---|---|---|---|---|---|---|---|---|---|---|
| Bacteroides:K02002 | 0 | 0 | 0 | 0 | 0 | 0 | 0 | 0 | 0 | 0 | 0 | 0 | 0 | 0.000418 | 0.017543 | 0.006854 | 0 | 0 | 0 | 0 | 0 |
| Bacteroides:K02049 | 0 | 0 | 0 | 0 | 0 | 0 | 0 | 0 | 0 | 0 | 0 | 0 | 0 | 0 | 0.000104 | 0 | 0 | 0 | 0 | 0 | 0 |
| Bacteroides:K02051 | 0 | 0 | 0 | 0 | 0 | 0 | 0 | 0 | 0 | 0 | 0 | 0 | 0 | 0.000418 | 0.001765 | 0.000762 | 0 | 0 | 0 | 0 | 0 |
| Bacteroides:K02372 | 0 | 0 | 0 | 0 | 0 | 0 | 0 | 0 | 0 | 0 | 0 | 0 | 0 | 0.000835 | 0.005086 | 0.00087 | 0 | 0 | 0 | 0 | 0 |
| Bacteroides:K02517 | 0 | 0 | 0 | 0 | 0 | 0 | 0 | 0 | 0 | 0.0021408 | 0.01575 | 0.0083101 | 0.016764 | 0.003341 | 0.062904 | 0.025349 | 0.00691 | 0.112023898 | 0.003471861 | 0.042946103 | 0 |
| Bacteroides:K02523 | 0 | 0.0409 | 0 | 0.0032 | 0 | 0 | 0 | 0 | 0 | 0 | 0 | 0.001366 | 0 | 0.005011 | 0.074738 | 0.034706 | 0.002962 | 0 | 0 | 0 | 0 |
| Bacteroides:K02557 | 0 | 0 | 0 | 0 | 0 | 0 | 0 | 0 | 0 | 0 | 0 | 0.0017076 | 0 | 0.000418 | 0.002076 | 0.001414 | 0 | 0 | 0 | 0 | 0 |
| Bacteroides:K03092 | 0 | 0.0205 | 0 | 0 | 0 | 4.15E-05 | 0 | 0 | 0 | 0 | 0.000477 | 0.0066025 | 0.014529 | 0.004593 | 0.090723 | 0.043083 | 0.008885 | 0 | 0 | 0 | 0.007893906 |
| Bacteroides:K03183 | 0 | 0.0205 | 0 | 0.0097 | 0 | 0 | 0 | 0 | 0 | 0.0021408 | 0.001432 | 0.0026182 | 0.011176 | 0.005011 | 0.052524 | 0.030898 | 0.002962 | 0 | 0 | 0 | 0 |
| Bacteroides:K03394 | 0 | 0 | 0 | 0 | 0 | 0 | 0 | 0 | 0 | 0 | 0 | 0 | 0.003353 | 0 | 0.008927 | 0.001741 | 0 | 0 | 0 | 0 | 0 |
| Bacteroides:K03590 | 0 | 0 | 0 | 0 | 0 | 0 | 0 | 0 | 0 | 0 | 0 | 0.0004553 | 0.006706 | 0.01044 | 0.094771 | 0.056791 | 0 | 0 | 0 | 0 | 0 |
| Bacteroides:K03644 | 0 | 0 | 0 | 0 | 0 | 0 | 0 | 0 | 0 | 0 | 0.000477 | 0.0021629 | 0.006706 | 0.003758 | 0.033632 | 0.021215 | 0.001974 | 0 | 0 | 0 | 0 |
| Bacteroides:K03781 | 0 | 0 | 0.0356 | 0.0081 | 0 | 0 | 0 | 0 | 0 | 0 | 0 | 0 | 0 | 0 | 0.003114 | 0.00544 | 0.000987 | 0 | 0 | 0 | 0 |
| Bacteroides:K03885 | 0 | 0 | 0 | 0 | 0 | 0 | 0 | 0 | 0 | 0.0064225 | 0.000955 | 0.0031874 | 0.008941 | 0.00167 | 0.003425 | 0.002611 | 0.001974 | 0 | 0 | 0 | 0 |
| Bacteroides:K04564 | 0 | 0 | 0 | 0 | 0 | 0 | 0 | 0 | 0 | 0 | 0.000477 | 0 | 0.001118 | 0.003341 | 0.024497 | 0.01175 | 0.001974 | 0 | 0 | 0 | 0 |
| Bacteroides:K05343 | 0 | 0 | 0 | 0 | 0 | 0 | 0 | 0 | 0 | 0 | 0 | 0.0002277 | 0 | 0 | 0.001246 | 0 | 0 | 0 | 0 | 0 | 0 |
| Bacteroides:K05592 | 0 | 0 | 0 | 0.0145 | 0 | 0 | 0 | 0 | 0 | 0.0032112 | 0.002386 | 0.0029598 | 0.011176 | 0.008769 | 0.091242 | 0.063319 | 0.00691 | 0 | 0 | 0 | 0.007893906 |
| Bacteroides:K06861 | 0 | 0 | 0 | 0.0032 | 0 | 0.00012 | 0 | 0 | 0 | 0.0010704 | 0.003818 | 0.0044396 | 0.00447 | 0.003341 | 0.043182 | 0.022412 | 0.002962 | 0 | 0 | 0 | 0 |
| Bacteroides:K07712 | 0 | 0 | 0 | 0 | 0 | 0 | 0 | 0 | 0 | 0 | 0 | 0.0001138 | 0 | 0 | 0 | 0 | 0 | 0 | 0 | 0 | 0 |
| Bacteroides:K10536 | 0 | 0 | 0 | 0 | 0 | 0 | 0 | 0 | 0 | 0 | 0 | 0.0001138 | 0 | 0 | 0 | 0 | 0 | 0 | 0 | 0 | 0 |
| Bacteroides:K11076 | 0 | 0 | 0 | 0 | 0 | 0 | 0 | 0 | 0 | 0 | 0 | 0.0001138 | 0 | 0 | 0 | 0 | 0 | 0 | 0 | 0 | 0 |
| Bacteroides:K11927 | 0 | 0 | 0 | 0 | 0 | 4.15E-05 | 0 | 0 | 0 | 0.0010704 | 0.000477 | 0.000683 | 0.005588 | 0.005011 | 0.086779 | 0.038296 | 0 | 0 | 0 | 0 | 0 |
| Bacteroides:K12573 | 0 | 0.0205 | 0 | 0.0129 | 0 | 4.15E-05 | 0 | 0 | 0 | 0.0160562 | 0.002864 | 0.0142296 | 0.029058 | 0.017956 | 0.139199 | 0.078442 | 0.009872 | 0 | 0 | 0 | 0.015787812 |
| Bacteroides:K13378 | 0 | 0 | 0.0713 | 0.0097 | 0 | 0 | 0 | 0 | 0 | 0.0010704 | 0.002386 | 0.0014799 | 0.010058 | 0.008352 | 0.110134 | 0.044062 | 0.002962 | 0 | 0 | 0 | 0 |
| Blautia:K00104 | 0 | 0 | 0 | 0 | 0 | 4.15E-05 | 0 | 0 | 0 | 0.0064225 | 0.000955 | 0.0088793 | 0.00447 | 0.043846 | 0.002906 | 0.003264 | 0.030602 | 0 | 0 | 0 | 0 |
| Blautia:K00111 | 0 | 0 | 0 | 0.0016 | 0 | 0 | 0 | 0 | 0.001025 | 0 | 0.001432 | 0.0015937 | 0.00447 | 0.077671 | 0.0082 | 0.007398 | 0.059231 | 0 | 0 | 0.021473051 | 0 |
| Blautia:K00123 | 0 | 0 | 0 | 0 | 0 | 0 | 0 | 0 | 0 | 0 | 0 | 0.0038704 | 0 | 0 | 0 | 0 | 0 | 0 | 0 | 0 | 0 |
| Blautia:K00128 | 0 | 0.0205 | 0 | 0 | 0 | 0 | 0 | 0 | 0 | 0.0021408 | 0.00525 | 0.0054642 | 0.002235 | 0.065143 | 0.00737 | 0.006745 | 0.04541 | 0 | 0 | 0.021473051 | 0 |
| Blautia:K00140 | 0 | 0 | 0 | 0 | 0 | 0 | 0 | 0 | 0 | 0.0021408 | 0 | 0.0007969 | 0.001118 | 0.045099 | 0.006955 | 0.004678 | 0.033564 | 0.0373413 | 0 | 0 | 0 |
| Blautia:K00303 | 0 | 0 | 0 | 0 | 0 | 0 | 0 | 0 | 0 | 0 | 0 | 0 | 0.002235 | 0 | 0 | 0 | 0 | 0 | 0 | 0 | 0 |
| Blautia:K00334 | 0 | 0 | 0 | 0 | 0 | 0.00012 | 0 | 0 | 0 | 0.0032112 | 0.00525 | 0.0071717 | 0.012294 | 0.016703 | 0.0082 | 0.007507 | 0.021718 | 0 | 0 | 0 | 0 |
| Blautia:K00335 | 0 | 0 | 0 | 0.0032 | 0 | 0.00021 | 0 | 0 | 0 | 0.0107041 | 0.006682 | 0.0224258 | 0.011176 | 0.071407 | 0.018892 | 0.026329 | 0.112538 | 0 | 0 | 0.042946103 | 0 |
| Blautia:K00336 | 0 | 0 | 0 | 0.0032 | 0 | 0.00017 | 0 | 0 | 0 | 0.0203378 | 0.014318 | 0.0241334 | 0.016764 | 0.039253 | 0.019826 | 0.032639 | 0.113525 | 0 | 0 | 0.042946103 | 0 |
| Blautia:K00525 | 0 | 0 | 0 | 0 | 0 | 0 | 0 | 0 | 0 | 0 | 0 | 0 | 0 | 0 | 0 | 0.000435 | 0 | 0 | 0 | 0 | 0 |
| Blautia:K00526 | 0 | 0 | 0 | 0 | 0 | 0 | 0 | 0 | 0 | 0 | 0 | 0 | 0 | 0.00167 | 0.000519 | 0.004352 | 0 | 0 | 0 | 0 | 0 |
| Blautia:K00548 | 0 | 0 | 0 | 0.0048 | 0 | 8.29E-05 | 0 | 0 | 0 | 0.0096337 | 0.006682 | 0.0140019 | 0.006706 | 0.071407 | 0.008616 | 0.028613 | 0.043436 | 0 | 0 | 0.128838308 | 0 |
| Blautia:K00600 | 0 | 0 | 0 | 0 | 0 | 0 | 0 | 0 | 0 | 0 | 0.000477 | 0.0011384 | 0.001118 | 0.000835 | 0.000311 | 0.001306 | 0 | 0 | 0 | 0 | 0 |
| Blautia:K00626 | 0 | 0 | 0 | 0 | 0 | 0 | 0 | 0 | 0 | 0 | 0 | 0 | 0 | 0.000418 | 0.000104 | 0 | 0 | 0 | 0 | 0 | 0 |
| Blautia:K00640 | 0 | 0 | 0 | 0 | 0 | 0 | 0 | 0 | 0 | 0.0042816 | 0.004295 | 0.006944 | 0.005588 | 0.031736 | 0.008823 | 0.010771 | 0.03159 | 0 | 0 | 0 | 0 |
| Blautia:K00648 | 0 | 0 | 0 | 0 | 0 | 0 | 0 | 0 | 0 | 0.0074929 | 0.004295 | 0.0054642 | 0.005588 | 0.068066 | 0.005086 | 0.008377 | 0.039487 | 0 | 0 | 0.021473051 | 0 |
| Blautia:K00798 | 0 | 0 | 0 | 0 | 0 | 0 | 0 | 0 | 0 | 0.0021408 | 0.000955 | 0.0019352 | 0.003353 | 0.022967 | 0.004048 | 0.00359 | 0.021718 | 0 | 0 | 0.021473051 | 0 |
| Blautia:K00833 | 0 | 0 | 0 | 0 | 0 | 0 | 0 | 0 | 0 | 0.0053521 | 0.000955 | 0.0004553 | 0 | 0 | 0.000311 | 0.000326 | 0 | 0 | 0 | 0 | 0 |
| Blautia:K00864 | 0 | 0 | 0 | 0.0016 | 0 | 0 | 0 | 0 | 0 | 0.0032112 | 0.001432 | 0.0073994 | 0.00447 | 0.057209 | 0.00737 | 0.018278 | 0.043436 | 0 | 0 | 0 | 0 |
| Blautia:K01142 | 0 | 0 | 0 | 0.0016 | 0 | 4.15E-05 | 0 | 0 | 0 | 0.0032112 | 0.000955 | 0.0023906 | 0.012294 | 0.031319 | 0.004152 | 0.004787 | 0.032577 | 0 | 0 | 0 | 0 |
| Blautia:K01338 | 0 | 0 | 0 | 0 | 0 | 8.29E-05 | 0 | 0 | 0 | 0.0171266 | 0.01575 | 0.0217428 | 0.016764 | 0.175803 | 0.022421 | 0.042757 | 0.132282 | 0.0373413 | 0 | 0.107365257 | 0 |
| Blautia:K01428 | 0 | 0 | 0 | 0 | 0 | 4.15E-05 | 0 | 0 | 0 | 0.0149858 | 0.001432 | 0.0095623 | 0.001118 | 0 | 0.001246 | 0.002067 | 0.003949 | 0 | 0 | 0 | 0 |
| Blautia:K01488 | 0 | 0 | 0 | 0 | 0 | 0 | 0 | 0 | 0 | 0 | 0.000477 | 0.0001138 | 0 | 0.022132 | 0.003218 | 0.000762 | 0.009872 | 0 | 0 | 0 | 0 |
| Blautia:K01497 | 0 | 0 | 0 | 0 | 0 | 0 | 0 | 0 | 0 | 0 | 0 | 0.0001138 | 0 | 0 | 0 | 0 | 0 | 0 | 0 | 0 | 0 |
| Blautia:K01613 | 0 | 0 | 0 | 0 | 0 | 0 | 0 | 0 | 0 | 0.0021408 | 0 | 0.0002277 | 0.001118 | 0.032989 | 0.003633 | 0.002285 | 0.017769 | 0 | 0 | 0.042946103 | 0 |
| Blautia:K01629 | 0 | 0 | 0 | 0 | 0 | 0 | 0 | 0 | 0 | 0.0064225 | 0.001432 | 0.0044396 | 0.00447 | 0.013363 | 0.004048 | 0.006854 | 0.03159 | 0 | 0 | 0 | 0 |
| Blautia:K01698 | 0 | 0 | 0 | 0.0016 | 0 | 8.29E-05 | 0 | 0 | 0 | 0.0021408 | 0.003341 | 0.0083101 | 0.001118 | 0.041758 | 0.007474 | 0.00903 | 0.037513 | 0 | 0 | 0.021473051 | 0 |
| Blautia:K01712 | 0 | 0 | 0 | 0 | 0 | 0 | 0 | 0 | 0 | 0 | 0 | 0.0001138 | 0 | 0 | 0 | 0 | 0 | 0 | 0 | 0 | 0 |
| Blautia:K01737 | 0 | 0 | 0 | 0 | 0 | 4.15E-05 | 0 | 0 | 0 | 0 | 0 | 0.0001138 | 0 | 0.002506 | 0 | 0.000109 | 0.000987 | 0 | 0 | 0 | 0 |
| Blautia:K01745 | 0 | 0 | 0 | 0 | 0 | 0 | 0 | 0 | 0 | 0 | 0 | 0.0002277 | 0 | 0 | 0 | 0 | 0 | 0 | 0 | 0 | 0 |
| Blautia:K01749 | 0 | 0 | 0 | 0 | 0 | 4.15E-05 | 0 | 0 | 0 | 0.0021408 | 0.001909 | 0.0025044 | 0.002235 | 0.032572 | 0.002803 | 0.005331 | 0.009872 | 0 | 0 | 0.021473051 | 0 |
| Blautia:K01845 | 0 | 0 | 0 | 0 | 0 | 4.15E-05 | 0 | 0 | 0 | 0.0032112 | 0.004295 | 0.0060333 | 0.003353 | 0.051363 | 0.005086 | 0.008921 | 0.04541 | 0 | 0 | 0.021473051 | 0 |
| Blautia:K01902 | 0 | 0 | 0 | 0 | 0 | 0 | 0 | 0 | 0 | 0 | 0.000477 | 0.0001138 | 0 | 0 | 0 | 0.000109 | 0 | 0 | 0 | 0 | 0 |
| Blautia:K01915 | 0 | 0 | 0 | 0.0081 | 0 | 0.00012 | 0 | 0 | 0 | 0.0203378 | 0.032932 | 0.0469007 | 0.05588 | 0.245122 | 0.044324 | 0.06778 | 0.308986 | 0 | 0 | 0.064419154 | 0.015787812 |
| Blautia:K01959 | 0 | 0 | 0 | 0 | 0 | 0 | 0 | 0 | 0 | 0 | 0 | 0.0001138 | 0 | 0 | 0 | 0 | 0 | 0 | 0 | 0 | 0 |
| Blautia:K01961 | 0 | 0 | 0 | 0 | 0 | 0 | 0 | 0 | 0 | 0.0096337 | 0.002386 | 0.0099038 | 0.005588 | 0.07433 | 0.013598 | 0.019801 | 0.074038 | 0 | 0 | 0 | 0 |

| | | | | | | | | | | | | | | | | | | | | | |
|---|---|---|---|---|---|---|---|---|---|---|---|---|---|---|---|---|---|---|---|---|---|
| Blautia:K01962 | 0 | 0 | 0 | 0 | 0 | 0 | 0 | 0 | 0 | 0 | 0 | 0.0002277 | 0 | 0 | 0 | 0.000218 | 0 | 0 | 0 | 0 | 0 |
| Blautia:K02047 | 0 | 0 | 0 | 0 | 0 | 0 | 0 | 0 | 0 | 0 | 0 | 0 | 0 | 0.00167 | 0.000415 | 0.000326 | 0.003949 | 0 | 0 | 0 | 0 |
| Blautia:K02048 | 0 | 0 | 0 | 0 | 0 | 4.15E-05 | 0 | 0 | 0 | 0.0128449 | 0.0105 | 0.0112698 | 0.005588 | 0.099385 | 0.009135 | 0.013273 | 0.137218 | 0 | 0 | 0.064419154 | 0 |
| Blautia:K02049 | 0 | 0 | 0 | 0 | 0 | 0 | 0 | 0 | 0 | 0.0042816 | 0.001909 | 0.0047811 | 0.002235 | 0.041341 | 0.004671 | 0.009139 | 0.019744 | 0 | 0 | 0 | 0 |
| Blautia:K02051 | 0 | 0 | 0 | 0 | 0 | 4.15E-05 | 0 | 0 | 0 | 0.0042816 | 0.000955 | 0.0085378 | 0.006706 | 0.036747 | 0.007889 | 0.011859 | 0.066141 | 0 | 0 | 0.107365257 | 0 |
| Blautia:K02203 | 0 | 0 | 0 | 0 | 0 | 4.15E-05 | 0 | 0 | 0 | 0.0032112 | 0.010977 | 0.0031874 | 0.003353 | 0.005429 | 0.001349 | 0.001741 | 0.020731 | 0 | 0 | 0 | 0 |
| Blautia:K02372 | 0 | 0 | 0 | 0 | 0 | 0 | 0 | 0 | 0 | 0.0032112 | 0.001432 | 0.0033013 | 0.002235 | 0.017539 | 0.001868 | 0.005005 | 0.020731 | 0.0373413 | 0 | 0.064419154 | 0 |
| Blautia:K02492 | 0 | 0 | 0 | 0 | 0 | 0 | 0 | 0 | 0 | 0.0032112 | 0.002386 | 0.0028459 | 0.005588 | 0.058879 | 0.005917 | 0.006854 | 0.042449 | 0 | 0 | 0.042946103 | 0 |
| Blautia:K02557 | 0 | 0 | 0 | 0 | 0 | 0 | 0 | 0 | 0 | 0 | 0.001909 | 0.0025044 | 0.001118 | 0.009604 | 0.001868 | 0.004025 | 0.014808 | 0.0373413 | 0 | 0.021473051 | 0 |
| Blautia:K03092 | 0 | 0 | 0 | 0 | 0 | 0 | 0 | 0 | 0 | 0 | 0 | 0 | 0 | 0.001253 | 0.002076 | 0.000979 | 0 | 0 | 0 | 0 | 0 |
| Blautia:K03182 | 0 | 0 | 0 | 0 | 0 | 4.15E-05 | 0 | 0 | 0 | 0 | 0.000477 | 0.0003415 | 0 | 0.019626 | 0.00737 | 0.005113 | 0.056269 | 0 | 0 | 0 | 0 |
| Blautia:K03338 | 0 | 0 | 0 | 0 | 0 | 0 | 0 | 0 | 0 | 0 | 0 | 0.0002277 | 0.001118 | 0.009604 | 0.002803 | 0.000762 | 0.009872 | 0 | 0 | 0 | 0 |
| Blautia:K03394 | 0 | 0 | 0 | 0 | 0 | 0 | 0 | 0 | 0 | 0.0085633 | 0.001909 | 0.0067164 | 0.002235 | 0.022967 | 0.00301 | 0.003808 | 0.009872 | 0 | 0 | 0.021473051 | 0 |
| Blautia:K05592 | 0 | 0 | 0 | 0 | 0 | 0 | 0 | 0 | 0 | 0 | 0 | 0 | 0 | 0 | 0.000311 | 0 | 0 | 0 | 0 | 0 | 0 |
| Blautia:K05847 | 0 | 0 | 0 | 0 | 0 | 0 | 0 | 0 | 0 | 0 | 0 | 0 | 0 | 0.000835 | 0 | 0.000653 | 0 | 0 | 0 | 0 | 0 |
| Blautia:K06148 | 0 | 0 | 0 | 0 | 0 | 0 | 0 | 0 | 0 | 0 | 0 | 0 | 0 | 0 | 0.000208 | 0.000326 | 0 | 0 | 0 | 0 | 0 |
| Blautia:K10536 | 0 | 0 | 0 | 0 | 0 | 0 | 0 | 0 | 0 | 0 | 0 | 0 | 0 | 0 | 0.000104 | 0.002502 | 0 | 0 | 0 | 0 | 0 |
| Blautia:K12573 | 0 | 0 | 0 | 0 | 0 | 0 | 0 | 0 | 0 | 0.0085633 | 0.005727 | 0.0079686 | 0.014529 | 0.099802 | 0.013079 | 0.016755 | 0.092795 | 0 | 0 | 0.021473051 | 0 |
| Butyrivibrio:K00111 | 0 | 0 | 0 | 0.0048 | 0 | 0 | 0 | 0 | 0 | 0 | 0 | 0.0005692 | 0.001118 | 0.000418 | 0.000104 | 0.000326 | 0.000987 | 0 | 0 | 0 | 0 |
| Butyrivibrio:K00128 | 0 | 0.0205 | 0 | 0.021 | 0 | 0 | 0 | 0 | 0 | 0 | 0 | 0 | 0 | 0 | 0 | 0 | 0 | 0 | 0 | 0 | 0 |
| Butyrivibrio:K00248 | 0 | 0 | 0 | 0.0016 | 0 | 4.15E-05 | 0 | 0 | 0 | 0 | 0 | 0.0010245 | 0.002235 | 0.000418 | 0 | 0.000109 | 0 | 0 | 0 | 0 | 0 |
| Butyrivibrio:K00372 | 0 | 0 | 0 | 0.0081 | 0 | 0 | 0 | 0 | 0 | 0 | 0 | 0 | 0 | 0 | 0 | 0 | 0 | 0 | 0 | 0 | 0 |
| Butyrivibrio:K00390 | 0 | 0 | 0 | 0 | 0 | 0 | 0 | 0 | 0 | 0 | 0 | 0.0001138 | 0 | 0 | 0 | 0 | 0 | 0 | 0 | 0 | 0 |
| Butyrivibrio:K00432 | 0 | 0 | 0 | 0.0048 | 0 | 4.15E-05 | 0 | 0 | 0 | 0 | 0 | 0 | 0 | 0 | 0 | 0 | 0 | 0 | 0 | 0 | 0 |
| Butyrivibrio:K00548 | 0 | 0 | 0 | 0 | 0 | 0 | 0 | 0 | 0 | 0.0010704 | 0.001909 | 0.0012522 | 0.002235 | 0 | 0.000311 | 0.000326 | 0 | 0 | 0 | 0 | 0 |
| Butyrivibrio:K00575 | 0 | 0 | 0 | 0 | 0 | 0 | 0 | 0 | 0 | 0 | 0.000477 | 0 | 0 | 0 | 0 | 0.000109 | 0 | 0 | 0 | 0 | 0 |
| Butyrivibrio:K00600 | 0 | 0 | 0 | 0.0065 | 0 | 0 | 0 | 0 | 0 | 0.0010704 | 0.000955 | 0.000683 | 0 | 0.00167 | 0.000104 | 0.000435 | 0.000987 | 0 | 0 | 0 | 0 |
| Butyrivibrio:K00626 | 0 | 0 | 0 | 0.0048 | 0 | 0.00091 | 0 | 0 | 0 | 0.0021408 | 0.004295 | 0.001366 | 0.003353 | 0.001253 | 0 | 0.000326 | 0 | 0 | 0 | 0 | 0 |
| Butyrivibrio:K00640 | 0 | 0 | 0 | 0 | 0 | 0 | 0 | 0 | 0 | 0 | 0 | 0.0001138 | 0 | 0.001253 | 0 | 0 | 0 | 0 | 0 | 0 | 0 |
| Butyrivibrio:K00648 | 0 | 0 | 0 | 0 | 0 | 0 | 0 | 0 | 0 | 0 | 0 | 0.0003415 | 0 | 0 | 0 | 0 | 0 | 0 | 0 | 0 | 0 |
| Butyrivibrio:K00798 | 0 | 0 | 0 | 0 | 0 | 0 | 0 | 0 | 0 | 0 | 0.000955 | 0 | 0 | 0 | 0 | 0.000326 | 0 | 0 | 0 | 0 | 0 |
| Butyrivibrio:K00864 | 0 | 0 | 0.0356 | 0.0162 | 0 | 0 | 0 | 0 | 0 | 0.0010704 | 0.000955 | 0.000683 | 0.002235 | 0.00167 | 0 | 0.000109 | 0 | 0 | 0 | 0 | 0 |
| Butyrivibrio:K01142 | 0 | 0 | 0 | 0.0016 | 0 | 0 | 0 | 0 | 0 | 0 | 0.000477 | 0.0004553 | 0.002235 | 0.003758 | 0.000415 | 0.000218 | 0 | 0 | 0 | 0 | 0 |
| Butyrivibrio:K01338 | 0 | 0 | 0 | 0.0016 | 0 | 0 | 0 | 0 | 0 | 0 | 0.000477 | 0.0003415 | 0 | 0.000418 | 0.000208 | 0 | 0 | 0 | 0 | 0 | 0 |
| Butyrivibrio:K01497 | 0 | 0 | 0 | 0 | 0 | 0 | 0 | 0 | 0 | 0 | 0 | 0.0001138 | 0 | 0 | 0 | 0 | 0 | 0 | 0 | 0 | 0 |
| Butyrivibrio:K01629 | 0 | 0 | 0 | 0.0081 | 0 | 0 | 0 | 0 | 0 | 0.0042816 | 0.000477 | 0.0005692 | 0 | 0.000418 | 0.000104 | 0.000109 | 0 | 0 | 0 | 0 | 0 |
| Butyrivibrio:K01679 | 0 | 0 | 0 | 0 | 0 | 0 | 0 | 0 | 0 | 0.0010704 | 0 | 0.0001138 | 0.002235 | 0.000418 | 0 | 0 | 0 | 0 | 0 | 0 | 0 |
| Butyrivibrio:K01845 | 0 | 0 | 0 | 0 | 0 | 0 | 0 | 0 | 0 | 0.0010704 | 0.000477 | 0.0011384 | 0.001118 | 0.000835 | 0 | 0 | 0 | 0 | 0 | 0 | 0 |
| Butyrivibrio:K01895 | 0 | 0 | 0 | 0 | 0 | 0 | 0 | 0 | 0 | 0.0021408 | 0.000477 | 0 | 0 | 0 | 0 | 0 | 0 | 0 | 0 | 0 | 0 |
| Butyrivibrio:K01915 | 0 | 0 | 0 | 0.0162 | 0 | 8.29E-05 | 0 | 0 | 0 | 0.0042816 | 0.00525 | 0.0047811 | 0.02347 | 0.002088 | 0.000208 | 0.004243 | 0.004936 | 0 | 0 | 0 | 0 |
| Butyrivibrio:K01961 | 0 | 0 | 0 | 0 | 0 | 0 | 0 | 0 | 0 | 0 | 0 | 0 | 0 | 0.000418 | 0 | 0.000218 | 0 | 0 | 0 | 0 | 0 |
| Butyrivibrio:K01962 | 0 | 0 | 0 | 0 | 0 | 4.15E-05 | 0 | 0 | 0 | 0 | 0.003341 | 0.0011384 | 0.002235 | 0.021297 | 0 | 0.005657 | 0.021718 | 0 | 0 | 0 | 0 |
| Butyrivibrio:K02011 | 0 | 0 | 0 | 0 | 0 | 0 | 0 | 0 | 0 | 0.0010704 | 0.000477 | 0.0005692 | 0.011176 | 0.002088 | 0 | 0 | 0 | 0 | 0 | 0 | 0 |
| Butyrivibrio:K02012 | 0 | 0 | 0 | 0 | 0 | 0 | 0 | 0 | 0 | 0 | 0.000955 | 0.0001138 | 0.002235 | 0 | 0 | 0 | 0 | 0 | 0 | 0 | 0 |
| Butyrivibrio:K02049 | 0 | 0 | 0 | 0 | 0 | 0 | 0 | 0 | 0 | 0 | 0 | 0.0005692 | 0 | 0 | 0 | 0 | 0 | 0 | 0 | 0 | 0 |
| Butyrivibrio:K02051 | 0 | 0 | 0 | 0 | 0 | 0 | 0 | 0 | 0 | 0 | 0 | 0.0005692 | 0 | 0 | 0 | 0 | 0 | 0 | 0 | 0 | 0 |
| Butyrivibrio:K02203 | 0 | 0 | 0 | 0.0032 | 0 | 4.15E-05 | 0 | 0 | 0 | 0 | 0.000955 | 0.000683 | 0.002235 | 0.00167 | 0 | 0.000109 | 0 | 0 | 0 | 0 | 0 |
| Butyrivibrio:K02372 | 0 | 0 | 0 | 0 | 0 | 0 | 0 | 0 | 0 | 0 | 0 | 0.0002277 | 0.001118 | 0.002088 | 0.000311 | 0.000218 | 0.000987 | 0 | 0 | 0 | 0 |
| Butyrivibrio:K02400 | 0 | 0 | 0 | 0 | 0 | 0.00021 | 0 | 0 | 0 | 0.0032112 | 0.002864 | 0.0015937 | 0.010058 | 0.000418 | 0.000104 | 0.001632 | 0 | 0 | 0 | 0 | 0 |
| Butyrivibrio:K02405 | 0 | 0 | 0 | 0 | 0 | 0 | 0 | 0 | 0 | 0.0010704 | 0.002386 | 0.0001138 | 0 | 0 | 0 | 0.000435 | 0 | 0 | 0 | 0 | 0 |
| Butyrivibrio:K02410 | 0 | 0 | 0 | 0 | 0 | 4.15E-05 | 0 | 0 | 0 | 0 | 0.001432 | 0.0005692 | 0 | 0 | 0 | 0 | 0 | 0 | 0 | 0 | 0 |
| Butyrivibrio:K02412 | 0 | 0 | 0 | 0 | 0 | 4.15E-05 | 0 | 0 | 0 | 0.0010704 | 0.003341 | 0.0012522 | 0.00447 | 0.00167 | 0 | 0.000435 | 0.000987 | 0 | 0 | 0 | 0 |
| Butyrivibrio:K02416 | 0 | 0 | 0 | 0 | 0 | 0 | 0 | 0 | 0 | 0 | 0.000477 | 0.0003415 | 0 | 0 | 0 | 0 | 0 | 0 | 0 | 0 | 0 |
| Butyrivibrio:K02556 | 0 | 0 | 0 | 0 | 0 | 0 | 0 | 0 | 0 | 0 | 0 | 0.0001138 | 0 | 0 | 0 | 0 | 0 | 0 | 0 | 0 | 0 |
| Butyrivibrio:K03407 | 0 | 0 | 0 | 0 | 0 | 0 | 0 | 0 | 0 | 0.0021408 | 0.000477 | 0.0003415 | 0.008941 | 0.000418 | 0 | 0.000218 | 0 | 0 | 0 | 0 | 0 |
| Butyrivibrio:K03412 | 0 | 0 | 0 | 0 | 0 | 0 | 0 | 0 | 0 | 0 | 0 | 0.0001138 | 0 | 0 | 0 | 0 | 0 | 0 | 0 | 0 | 0 |
| Butyrivibrio:K03415 | 0 | 0 | 0 | 0 | 0 | 0 | 0 | 0 | 0 | 0.0010704 | 0.001432 | 0.0007969 | 0.003353 | 0 | 0 | 0.000544 | 0 | 0 | 0 | 0 | 0 |
| Butyrivibrio:K05343 | 0 | 0 | 0 | 0.0065 | 0 | 0 | 0 | 0 | 0 | 0.0010704 | 0 | 0.001366 | 0.001118 | 0.033407 | 0 | 0.002285 | 0.004936 | 0 | 0 | 0 | 0 |
| Butyrivibrio:K05592 | 0 | 0 | 0 | 0 | 0 | 0 | 0 | 0 | 0 | 0 | 0.000477 | 0.0005692 | 0 | 0 | 0 | 0.000109 | 0 | 0 | 0 | 0 | 0 |
| Butyrivibrio:K06148 | 0 | 0 | 0 | 0 | 0 | 0 | 0 | 0 | 0 | 0 | 0 | 0.0001138 | 0 | 0 | 0 | 0 | 0 | 0 | 0 | 0 | 0 |

| | | | | | | | | | | | | | | | | | | | | | |
|---|---|---|---|---|---|---|---|---|---|---|---|---|---|---|---|---|---|---|---|---|---|
| Butyrivibrio:K10536 | 0 | 0 | 0 | 0 | 0 | 4.15E-05 | 0 | 0 | 0 | 0 | 0.000955 | 0.001366 | 0 | 0.00167 | 0 | 0.002937 | 0.007897 | 0 | 0 | 0 | 0 |
| Butyrivibrio:K12573 | 0 | 0 | 0 | 0 | 0 | 0 | 0 | 0 | 0 | 0.0021408 | 0.002864 | 0.001366 | 0.00447 | 0.000835 | 0.000104 | 0.000326 | 0.000987 | 0 | 0 | 0 | 0 |
| Cellulomonas:K00036 | 0 | 0 | 0.0713 | 0.0194 | 0 | 0 | 0 | 0 | 0 | 0 | 0 | 0 | 0 | 0 | 0 | 0 | 0 | 0 | 0 | 0 | 0 |
| Cellulomonas:K00111 | 0 | 0.0205 | 0.0356 | 0.0242 | 0 | 0 | 0 | 0 | 0 | 0 | 0 | 0 | 0 | 0 | 0 | 0 | 0 | 0 | 0 | 0 | 0 |
| Cellulomonas:K00116 | 0 | 0 | 0.0356 | 0.0032 | 0 | 0 | 0 | 0 | 0 | 0 | 0 | 0 | 0 | 0 | 0 | 0 | 0 | 0 | 0 | 0 | 0 |
| Cellulomonas:K00130 | 0 | 0 | 0 | 0.0016 | 0 | 0 | 0 | 0 | 0 | 0 | 0 | 0 | 0 | 0 | 0 | 0 | 0 | 0 | 0 | 0 | 0 |
| Cellulomonas:K00135 | 0 | 0.0205 | 0 | 0.0065 | 0 | 0 | 0 | 0 | 0 | 0 | 0 | 0 | 0 | 0 | 0 | 0 | 0 | 0 | 0 | 0 | 0 |
| Cellulomonas:K00161 | 0 | 0.0205 | 0 | 0.0275 | 0 | 0 | 0 | 0 | 0 | 0 | 0 | 0 | 0 | 0 | 0 | 0 | 0 | 0 | 0 | 0 | 0 |
| Cellulomonas:K00162 | 0 | 0.0205 | 0 | 0.0081 | 0 | 0 | 0 | 0 | 0 | 0 | 0 | 0 | 0 | 0 | 0 | 0 | 0 | 0 | 0 | 0 | 0 |
| Cellulomonas:K00163 | 0 | 0.0819 | 0 | 0.0468 | 0 | 0 | 0 | 0 | 0 | 0 | 0 | 0 | 0 | 0 | 0 | 0 | 0 | 0 | 0 | 0 | 0 |
| Cellulomonas:K00164 | 0 | 0.0409 | 0.0356 | 0.042 | 0 | 0 | 0 | 0 | 0 | 0 | 0 | 0 | 0 | 0 | 0 | 0 | 0 | 0 | 0 | 0 | 0 |
| Cellulomonas:K00252 | 0 | 0 | 0 | 0.0016 | 0 | 0 | 0 | 0 | 0 | 0 | 0 | 0 | 0 | 0 | 0 | 0 | 0 | 0 | 0 | 0 | 0 |
| Cellulomonas:K00281 | 0 | 0 | 0 | 0.0065 | 0 | 0 | 0 | 0 | 0 | 0 | 0 | 0 | 0 | 0 | 0 | 0 | 0 | 0 | 0 | 0 | 0 |
| Cellulomonas:K00331 | 0 | 0 | 0 | 0.0032 | 0 | 0 | 0 | 0 | 0 | 0 | 0 | 0 | 0 | 0 | 0 | 0 | 0 | 0 | 0 | 0 | 0 |
| Cellulomonas:K00334 | 0 | 0 | 0 | 0.0065 | 0 | 0 | 0 | 0 | 0 | 0 | 0 | 0 | 0 | 0 | 0 | 0 | 0 | 0 | 0 | 0 | 0 |
| Cellulomonas:K00335 | 0 | 0.0205 | 0 | 0.0145 | 0 | 4.15E-05 | 0 | 0 | 0 | 0 | 0 | 0 | 0 | 0 | 0 | 0 | 0 | 0 | 0 | 0 | 0 |
| Cellulomonas:K00336 | 0 | 0 | 0 | 0.0097 | 0 | 0 | 0 | 0 | 0 | 0 | 0 | 0 | 0 | 0 | 0 | 0 | 0 | 0 | 0 | 0 | 0 |
| Cellulomonas:K00337 | 0 | 0 | 0 | 0.0065 | 0 | 0 | 0 | 0 | 0 | 0 | 0 | 0 | 0 | 0 | 0 | 0 | 0 | 0 | 0 | 0 | 0 |
| Cellulomonas:K00338 | 0 | 0 | 0 | 0.0081 | 0 | 0 | 0 | 0 | 0 | 0 | 0 | 0 | 0 | 0 | 0 | 0 | 0 | 0 | 0 | 0 | 0 |
| Cellulomonas:K00341 | 0 | 0.0205 | 0.0356 | 0.0097 | 0 | 0 | 0 | 0 | 0 | 0 | 0 | 0 | 0 | 0 | 0 | 0 | 0 | 0 | 0 | 0 | 0 |
| Cellulomonas:K00342 | 0 | 0.0205 | 0 | 0.0113 | 0 | 0 | 0 | 0 | 0 | 0 | 0 | 0 | 0 | 0 | 0 | 0 | 0 | 0 | 0 | 0 | 0 |
| Cellulomonas:K00362 | 0 | 0 | 0 | 0.021 | 0 | 0 | 0 | 0 | 0 | 0 | 0 | 0 | 0 | 0 | 0 | 0 | 0 | 0 | 0 | 0 | 0 |
| Cellulomonas:K00370 | 0 | 0.0205 | 0 | 0.0016 | 0 | 0 | 0 | 0 | 0 | 0 | 0 | 0 | 0 | 0 | 0 | 0 | 0 | 0 | 0 | 0 | 0 |
| Cellulomonas:K00371 | 0 | 0 | 0 | 0.0016 | 0 | 0 | 0 | 0 | 0 | 0 | 0 | 0 | 0 | 0 | 0 | 0 | 0 | 0 | 0 | 0 | 0 |
| Cellulomonas:K00372 | 0 | 0 | 0 | 0.0016 | 0 | 0 | 0 | 0 | 0 | 0 | 0 | 0 | 0 | 0 | 0 | 0 | 0 | 0 | 0 | 0 | 0 |
| Cellulomonas:K00390 | 0 | 0 | 0 | 0.0097 | 0 | 0 | 0 | 0 | 0 | 0 | 0 | 0 | 0 | 0 | 0 | 0 | 0 | 0 | 0 | 0 | 0 |
| Cellulomonas:K00425 | 0 | 0.0205 | 0 | 0.0048 | 0 | 0 | 0 | 0 | 0 | 0 | 0 | 0 | 0 | 0 | 0 | 0 | 0 | 0 | 0 | 0 | 0 |
| Cellulomonas:K00426 | 0 | 0.0205 | 0 | 0.0081 | 0 | 0 | 0 | 0 | 0 | 0 | 0 | 0 | 0 | 0 | 0 | 0 | 0 | 0 | 0 | 0 | 0 |
| Cellulomonas:K00525 | 0 | 0.0614 | 0.0356 | 0.0275 | 0 | 0 | 0 | 0 | 0 | 0 | 0 | 0 | 0 | 0 | 0 | 0 | 0 | 0 | 0 | 0 | 0 |
| Cellulomonas:K00526 | 0 | 0 | 0 | 0.0081 | 0 | 0 | 0 | 0 | 0 | 0 | 0 | 0 | 0 | 0 | 0 | 0 | 0 | 0 | 0 | 0 | 0 |
| Cellulomonas:K00575 | 0 | 0 | 0 | 0.0016 | 0 | 0 | 0 | 0 | 0 | 0 | 0 | 0 | 0 | 0 | 0 | 0 | 0 | 0 | 0 | 0 | 0 |
| Cellulomonas:K00600 | 0 | 0 | 0 | 0.0065 | 0 | 0 | 0 | 0 | 0 | 0 | 0 | 0 | 0 | 0 | 0 | 0 | 0 | 0 | 0 | 0 | 0 |
| Cellulomonas:K00626 | 0 | 0.0205 | 0 | 0.0032 | 0 | 4.15E-05 | 0 | 0 | 0 | 0 | 0 | 0 | 0 | 0 | 0 | 0 | 0 | 0 | 0 | 0 | 0 |
| Cellulomonas:K00627 | 0 | 0 | 0 | 0.0048 | 0 | 0 | 0 | 0 | 0 | 0 | 0 | 0 | 0 | 0 | 0 | 0 | 0 | 0 | 0 | 0 | 0 |
| Cellulomonas:K00632 | 0 | 0 | 0 | 0.0065 | 0 | 0 | 0 | 0 | 0 | 0 | 0 | 0 | 0 | 0 | 0 | 0 | 0 | 0 | 0 | 0 | 0 |
| Cellulomonas:K00640 | 0 | 0 | 0 | 0.0016 | 0 | 0 | 0 | 0 | 0 | 0 | 0 | 0 | 0 | 0 | 0 | 0 | 0 | 0 | 0 | 0 | 0 |
| Cellulomonas:K00641 | 0 | 0.0614 | 0 | 0.0275 | 0 | 4.15E-05 | 0 | 0 | 0 | 0 | 0 | 0 | 0 | 0 | 0 | 0 | 0 | 0 | 0 | 0 | 0 |
| Cellulomonas:K00648 | 0 | 0.0409 | 0 | 0.0081 | 0 | 0 | 0 | 0 | 0 | 0 | 0 | 0 | 0 | 0 | 0 | 0 | 0 | 0 | 0 | 0 | 0 |
| Cellulomonas:K00658 | 0 | 0.0205 | 0.0356 | 0.0048 | 0 | 0 | 0 | 0 | 0 | 0 | 0 | 0 | 0 | 0 | 0 | 0 | 0 | 0 | 0 | 0 | 0 |
| Cellulomonas:K00681 | 0 | 0 | 0 | 0.0081 | 0 | 0 | 0 | 0 | 0 | 0 | 0 | 0 | 0 | 0 | 0 | 0 | 0 | 0 | 0 | 0 | 0 |
| Cellulomonas:K00697 | 0 | 0.0409 | 0 | 0.0145 | 0 | 0 | 0 | 0 | 0 | 0 | 0 | 0 | 0 | 0 | 0 | 0 | 0 | 0 | 0 | 0 | 0 |
| Cellulomonas:K00819 | 0 | 0 | 0 | 0.0016 | 0 | 0 | 0 | 0 | 0 | 0 | 0 | 0 | 0 | 0 | 0 | 0 | 0 | 0 | 0 | 0 | 0 |
| Cellulomonas:K00864 | 0 | 0 | 0.0356 | 0.0129 | 0 | 0 | 0 | 0 | 0 | 0 | 0 | 0 | 0 | 0 | 0 | 0 | 0 | 0 | 0 | 0 | 0 |
| Cellulomonas:K00940 | 0 | 0.0205 | 0 | 0.0016 | 0 | 0 | 0 | 0 | 0 | 0 | 0 | 0 | 0 | 0 | 0 | 0 | 0 | 0 | 0 | 0 | 0 |
| Cellulomonas:K01011 | 0 | 0 | 0 | 0.0097 | 0 | 0 | 0 | 0 | 0 | 0 | 0 | 0 | 0 | 0 | 0 | 0 | 0 | 0 | 0 | 0 | 0 |
| Cellulomonas:K01142 | 0 | 0.0409 | 0 | 0.0081 | 0 | 0 | 0 | 0 | 0 | 0 | 0 | 0 | 0 | 0 | 0 | 0 | 0 | 0 | 0 | 0 | 0 |
| Cellulomonas:K01255 | 0 | 0 | 0.0713 | 0.0162 | 0 | 0 | 0 | 0 | 0 | 0 | 0 | 0 | 0 | 0 | 0 | 0 | 0 | 0 | 0 | 0 | 0 |
| Cellulomonas:K01428 | 0 | 0 | 0 | 0.0016 | 0 | 0 | 0 | 0 | 0 | 0 | 0 | 0 | 0 | 0 | 0 | 0 | 0 | 0 | 0 | 0 | 0 |
| Cellulomonas:K01433 | 0 | 0.0205 | 0 | 0.0129 | 0 | 0 | 0 | 0 | 0 | 0 | 0 | 0 | 0 | 0 | 0 | 0 | 0 | 0 | 0 | 0 | 0 |
| Cellulomonas:K01488 | 0 | 0 | 0 | 0.0081 | 0 | 4.15E-05 | 0 | 0 | 0 | 0 | 0 | 0 | 0 | 0 | 0 | 0 | 0 | 0 | 0 | 0 | 0 |
| Cellulomonas:K01679 | 0 | 0 | 0.0356 | 0 | 0 | 0 | 0 | 0 | 0 | 0 | 0 | 0 | 0 | 0 | 0 | 0 | 0 | 0 | 0 | 0 | 0 |
| Cellulomonas:K01698 | 0 | 0 | 0 | 0.0065 | 0 | 0 | 0 | 0 | 0 | 0 | 0 | 0 | 0 | 0 | 0 | 0 | 0 | 0 | 0 | 0 | 0 |
| Cellulomonas:K01712 | 0 | 0 | 0 | 0.0032 | 0 | 0 | 0 | 0 | 0 | 0 | 0 | 0 | 0 | 0 | 0 | 0 | 0 | 0 | 0 | 0 | 0 |
| Cellulomonas:K01749 | 0 | 0 | 0.0713 | 0.0016 | 0 | 0 | 0 | 0 | 0 | 0 | 0 | 0 | 0 | 0 | 0 | 0 | 0 | 0 | 0 | 0 | 0 |
| Cellulomonas:K01772 | 0 | 0 | 0 | 0.0032 | 0 | 4.15E-05 | 0 | 0 | 0 | 0 | 0 | 0 | 0 | 0 | 0 | 0 | 0 | 0 | 0 | 0 | 0 |
| Cellulomonas:K01845 | 0 | 0 | 0 | 0.0032 | 0 | 0 | 0 | 0 | 0 | 0 | 0 | 0 | 0 | 0 | 0 | 0 | 0 | 0 | 0 | 0 | 0 |
| Cellulomonas:K01895 | 0 | 0.0614 | 0.0356 | 0.0355 | 0 | 0 | 0 | 0 | 0 | 0 | 0 | 0 | 0 | 0 | 0 | 0 | 0 | 0 | 0 | 0 | 0 |
| Cellulomonas:K01902 | 0 | 0 | 0 | 0.0097 | 0 | 0 | 0 | 0 | 0 | 0 | 0 | 0 | 0 | 0 | 0 | 0 | 0 | 0 | 0 | 0 | 0 |
| Cellulomonas:K01903 | 0 | 0 | 0 | 0.0113 | 0 | 0 | 0 | 0 | 0 | 0 | 0 | 0 | 0 | 0 | 0 | 0 | 0 | 0 | 0 | 0 | 0 |
| Cellulomonas:K01915 | 0 | 0.0819 | 0.0356 | 0.0533 | 0 | 4.15E-05 | 0 | 0 | 0 | 0 | 0 | 0 | 0 | 0 | 0 | 0 | 0 | 0 | 0 | 0 | 0 |

| | | | | | | | | | | | | | | | | | | | | | |
|---|---|---|---|---|---|---|---|---|---|---|---|---|---|---|---|---|---|---|---|---|---|
| Cellulomonas:K01941 | 0 | 0 | 0 | 0.0048 | 0 | 0 | 0 | 0 | 0 | 0 | 0 | 0 | 0 | 0 | 0 | 0 | 0 | 0 | 0 | 0 | 0 |
| Cellulomonas:K02049 | 0 | 0 | 0 | 0.0048 | 0 | 0 | 0 | 0 | 0 | 0 | 0 | 0 | 0 | 0 | 0 | 0 | 0 | 0 | 0 | 0 | 0 |
| Cellulomonas:K02051 | 0 | 0 | 0 | 0.0129 | 0 | 0 | 0 | 0 | 0 | 0 | 0 | 0 | 0 | 0 | 0 | 0 | 0 | 0 | 0 | 0 | 0 |
| Cellulomonas:K02274 | 0 | 0 | 0 | 0.0113 | 0 | 4.15E-05 | 0 | 0 | 0 | 0 | 0 | 0 | 0 | 0 | 0 | 0 | 0 | 0 | 0 | 0 | 0 |
| Cellulomonas:K02275 | 0 | 0.0205 | 0 | 0.0113 | 0 | 0 | 0 | 0 | 0 | 0 | 0 | 0 | 0 | 0 | 0 | 0 | 0 | 0 | 0 | 0 | 0 |
| Cellulomonas:K02276 | 0 | 0 | 0 | 0.0113 | 0 | 0 | 0 | 0 | 0 | 0 | 0 | 0 | 0 | 0 | 0 | 0 | 0 | 0 | 0 | 0 | 0 |
| Cellulomonas:K02301 | 0 | 0 | 0 | 0.0081 | 0 | 0 | 0 | 0 | 0 | 0 | 0 | 0 | 0 | 0 | 0 | 0 | 0 | 0 | 0 | 0 | 0 |
| Cellulomonas:K02400 | 0 | 0 | 0 | 0.0048 | 0 | 0 | 0 | 0 | 0.001025 | 0 | 0 | 0 | 0 | 0 | 0 | 0 | 0 | 0 | 0 | 0 | 0 |
| Cellulomonas:K02405 | 0 | 0 | 0 | 0.0032 | 0 | 0 | 0 | 0 | 0 | 0 | 0 | 0 | 0 | 0 | 0 | 0 | 0 | 0 | 0 | 0 | 0 |
| Cellulomonas:K02410 | 0 | 0 | 0 | 0 | 0 | 4.15E-05 | 0 | 0 | 0 | 0 | 0 | 0 | 0 | 0 | 0 | 0 | 0 | 0 | 0 | 0 | 0 |
| Cellulomonas:K02412 | 0 | 0 | 0 | 0.0048 | 0 | 0 | 0 | 0 | 0 | 0 | 0 | 0 | 0 | 0 | 0 | 0 | 0 | 0 | 0 | 0 | 0 |
| Cellulomonas:K02557 | 0 | 0.0205 | 0 | 0.0016 | 0 | 0 | 0 | 0 | 0.001025 | 0 | 0 | 0 | 0 | 0 | 0 | 0 | 0 | 0 | 0 | 0 | 0 |
| Cellulomonas:K03072 | 0 | 0.0205 | 0 | 0.0032 | 0 | 0 | 0 | 0 | 0 | 0 | 0 | 0 | 0 | 0 | 0 | 0 | 0 | 0 | 0 | 0 | 0 |
| Cellulomonas:K03183 | 0 | 0 | 0 | 0.0016 | 0 | 0 | 0 | 0 | 0 | 0 | 0 | 0 | 0 | 0 | 0 | 0 | 0 | 0 | 0 | 0 | 0 |
| Cellulomonas:K03407 | 0 | 0 | 0 | 0.0032 | 0 | 4.15E-05 | 0 | 0 | 0 | 0 | 0 | 0 | 0 | 0 | 0 | 0 | 0 | 0 | 0 | 0 | 0 |
| Cellulomonas:K03582 | 0 | 0 | 0.0356 | 0.0129 | 0 | 0 | 0 | 0 | 0 | 0 | 0 | 0 | 0 | 0 | 0 | 0 | 0 | 0 | 0 | 0 | 0 |
| Cellulomonas:K03583 | 0 | 0.0614 | 0 | 0.0048 | 0 | 0 | 0 | 0 | 0 | 0 | 0 | 0 | 0 | 0 | 0 | 0 | 0 | 0 | 0 | 0 | 0 |
| Cellulomonas:K03644 | 0 | 0 | 0 | 0.0129 | 0 | 0 | 0 | 0 | 0 | 0 | 0 | 0 | 0 | 0 | 0 | 0 | 0 | 0 | 0 | 0 | 0 |
| Cellulomonas:K03781 | 0 | 0 | 0 | 0.0162 | 0 | 0 | 0 | 0 | 0 | 0 | 0 | 0 | 0 | 0 | 0 | 0 | 0 | 0 | 0 | 0 | 0 |
| Cellulomonas:K03885 | 0 | 0.0205 | 0 | 0.0162 | 0 | 0 | 0 | 0.0007694 | 0 | 0 | 0 | 0 | 0 | 0 | 0 | 0 | 0 | 0 | 0 | 0 | 0 |
| Cellulomonas:K04564 | 0 | 0 | 0.0713 | 0.0129 | 0 | 0 | 0 | 0 | 0.001025 | 0 | 0 | 0 | 0 | 0 | 0 | 0 | 0 | 0 | 0 | 0 | 0 |
| Cellulomonas:K05343 | 0 | 0.0205 | 0 | 0.0178 | 0 | 0 | 0 | 0 | 0 | 0 | 0 | 0 | 0 | 0 | 0 | 0 | 0 | 0 | 0 | 0 | 0 |
| Cellulomonas:K05592 | 0 | 0 | 0 | 0.0048 | 0 | 0 | 0 | 0 | 0 | 0 | 0 | 0 | 0 | 0 | 0 | 0 | 0 | 0 | 0 | 0 | 0 |
| Cellulomonas:K05846 | 0 | 0 | 0 | 0.0048 | 0 | 0 | 0 | 0 | 0 | 0 | 0 | 0 | 0 | 0 | 0 | 0 | 0 | 0 | 0 | 0 | 0 |
| Cellulomonas:K05847 | 0 | 0 | 0.0356 | 0.0032 | 0 | 0 | 0 | 0 | 0 | 0 | 0 | 0 | 0 | 0 | 0 | 0 | 0 | 0 | 0 | 0 | 0 |
| Cellulomonas:K11959 | 0 | 0.0205 | 0 | 0.0291 | 0 | 0 | 0 | 0 | 0 | 0 | 0 | 0 | 0 | 0 | 0 | 0 | 0 | 0 | 0 | 0 | 0 |
| Cellulomonas:K11962 | 0 | 0 | 0 | 0.0016 | 0 | 0 | 0 | 0 | 0 | 0 | 0 | 0 | 0 | 0 | 0 | 0 | 0 | 0 | 0 | 0 | 0 |
| Cellulomonas:K11963 | 0 | 0 | 0 | 0.0016 | 0 | 0 | 0 | 0 | 0 | 0 | 0 | 0 | 0 | 0 | 0 | 0 | 0 | 0 | 0 | 0 | 0 |
| Clostridium:K00029 | 0 | 0 | 0 | 0 | 0 | 4.15E-05 | 0 | 0.0023082 | 0 | 0 | 0 | 0 | 0 | 0 | 0 | 0 | 0 | 0 | 0 | 0 | 0 |
| Clostridium:K00036 | 0 | 0 | 0 | 0 | 0 | 8.29E-05 | 0 | 0.0023082 | 0.003076 | 0 | 0 | 0 | 0 | 0 | 0 | 0 | 0 | 0 | 0 | 0 | 0 |
| Clostridium:K00104 | 0 | 0 | 0 | 0.0065 | 0 | 0.09704 | 0.216254 | 0.2815969 | 0.268679 | 0.0042816 | 0.002386 | 0.0015937 | 0.00447 | 0.003758 | 0.001246 | 0.000109 | 0 | 0 | 0 | 0.021473051 | 0 |
| Clostridium:K00111 | 0 | 0 | 0 | 0 | 0 | 0.15679 | 0.147847 | 0.1646496 | 0.188691 | 0.0096337 | 0.007159 | 0.0036428 | 0.013411 | 0.002506 | 0.002491 | 0.000544 | 0 | 0 | 0 | 0 | 0 |
| Clostridium:K00121 | 0 | 0 | 0 | 0 | 0 | 0.00012 | 0 | 0.0023082 | 0.003076 | 0 | 0 | 0 | 0 | 0 | 0 | 0 | 0 | 0 | 0 | 0 | 0 |
| Clostridium:K00123 | 0 | 0 | 0 | 0.0016 | 0 | 0.0063 | 0 | 0.0023082 | 0.003076 | 0 | 0.000477 | 0.0001138 | 0 | 0 | 0 | 0 | 0 | 0 | 0 | 0 | 0 |
| Clostridium:K00128 | 0 | 0 | 0 | 0.0048 | 0 | 0.09787 | 0.011033 | 0.0392389 | 0.056402 | 0.0021408 | 0.001432 | 0.0002277 | 0.001118 | 0 | 0 | 0.000109 | 0 | 0 | 0 | 0 | 0 |
| Clostridium:K00130 | 0 | 0 | 0 | 0.0016 | 0 | 0.00066 | 0 | 0 | 0 | 0.0032112 | 0.003341 | 0.0028459 | 0.00447 | 0.005429 | 0 | 0.001306 | 0.002962 | 0 | 0 | 0 | 0 |
| Clostridium:K00131 | 0 | 0 | 0 | 0 | 0 | 0.14655 | 0.004413 | 0.0723227 | 0.08409 | 0 | 0 | 0.0002277 | 0 | 0 | 0 | 0 | 0 | 0 | 0 | 0 | 0 |
| Clostridium:K00140 | 0 | 0 | 0 | 0 | 0 | 0.00261 | 0 | 0.0153878 | 0.006153 | 0 | 0 | 0 | 0 | 0.000418 | 0 | 0 | 0 | 0 | 0 | 0 | 0 |
| Clostridium:K00161 | 0 | 0 | 0 | 0 | 0 | 0.00066 | 0 | 0.0007694 | 0 | 0.0064225 | 0.001432 | 0.0007969 | 0.001118 | 0.010022 | 0 | 0.001632 | 0 | 0 | 0 | 0 | 0 |
| Clostridium:K00162 | 0 | 0 | 0 | 0 | 0 | 0.00025 | 0 | 0 | 0 | 0 | 0 | 0.0002277 | 0 | 0.00167 | 0 | 0.000326 | 0 | 0 | 0 | 0 | 0 |
| Clostridium:K00248 | 0 | 0.0205 | 0 | 0.0097 | 0 | 0.08103 | 0.278041 | 0.263901 | 0.226634 | 0.0032112 | 0.00525 | 0.0026182 | 0.003353 | 0.007934 | 0.000519 | 0.000326 | 0.000987 | 0 | 0 | 0 | 0 |
| Clostridium:K00257 | 0 | 0 | 0 | 0 | 0 | 0.02762 | 0 | 0.017696 | 0.012306 | 0.0010704 | 0 | 0.0001138 | 0 | 0 | 0 | 0 | 0 | 0 | 0 | 0 | 0 |
| Clostridium:K00303 | 0 | 0 | 0 | 0 | 0 | 0 | 0 | 0 | 0 | 0.0021408 | 0.001909 | 0.0020491 | 0.012294 | 0.000835 | 0 | 0.005005 | 0 | 0 | 0 | 0.021473051 | 0 |
| Clostridium:K00331 | 0 | 0 | 0 | 0 | 0 | 0.00037 | 0 | 0 | 0 | 0 | 0 | 0 | 0 | 0 | 0 | 0 | 0 | 0 | 0 | 0 | 0 |
| Clostridium:K00334 | 0 | 0 | 0 | 0 | 0 | 0.0039 | 0 | 0.0338532 | 0.027688 | 0.0053521 | 0.000955 | 0.0019352 | 0.003353 | 0.000835 | 0.000104 | 0.00087 | 0 | 0 | 0 | 0 | 0 |
| Clostridium:K00335 | 0 | 0.0409 | 0.0356 | 0.0258 | 0 | 0.0102 | 0.004413 | 0.0915575 | 0.081014 | 0.0214082 | 0.018136 | 0.0169617 | 0.005588 | 0.017539 | 0.000208 | 0.000762 | 0.001974 | 0 | 0 | 0 | 0 |
| Clostridium:K00336 | 0 | 0 | 0 | 0.0032 | 0 | 0.01053 | 0.015447 | 0.0700145 | 0.037943 | 0.0064225 | 0.008114 | 0.0058057 | 0.002235 | 0.017539 | 0.001868 | 0.000979 | 0.002962 | 0 | 0 | 0 | 0 |
| Clostridium:K00337 | 0 | 0 | 0 | 0 | 0 | 0.0012 | 0 | 0 | 0 | 0 | 0 | 0 | 0 | 0 | 0 | 0 | 0 | 0 | 0 | 0 | 0 |
| Clostridium:K00338 | 0 | 0 | 0 | 0 | 0 | 0.00041 | 0 | 0 | 0 | 0 | 0 | 0 | 0 | 0 | 0 | 0 | 0 | 0 | 0 | 0 | 0 |
| Clostridium:K00341 | 0 | 0 | 0 | 0 | 0 | 0.00137 | 0 | 0 | 0 | 0 | 0 | 0 | 0 | 0 | 0 | 0 | 0 | 0 | 0 | 0 | 0 |
| Clostridium:K00342 | 0 | 0 | 0 | 0 | 0 | 0.00087 | 0 | 0.0015388 | 0 | 0 | 0 | 0 | 0 | 0 | 0 | 0 | 0 | 0 | 0 | 0 | 0 |
| Clostridium:K00362 | 0 | 0 | 0 | 0 | 0 | 8.29E-05 | 0 | 0 | 0.003076 | 0 | 0 | 0 | 0 | 0 | 0 | 0 | 0 | 0 | 0 | 0 | 0 |
| Clostridium:K00372 | 0 | 0 | 0.0356 | 0 | 0 | 0.13925 | 0.00662 | 0.0169266 | 0.024612 | 0 | 0.000477 | 0.0002277 | 0.003353 | 0.000418 | 0 | 0 | 0 | 0 | 0 | 0 | 0 |
| Clostridium:K00390 | 0 | 0 | 0 | 0 | 0 | 0.00041 | 0 | 0 | 0 | 0 | 0 | 0 | 0 | 0 | 0 | 0 | 0 | 0 | 0 | 0 | 0 |
| Clostridium:K00425 | 0 | 0 | 0 | 0 | 0 | 0.00029 | 0 | 0.0007694 | 0 | 0 | 0 | 0 | 0 | 0 | 0 | 0 | 0 | 0 | 0 | 0 | 0 |
| Clostridium:K00426 | 0 | 0 | 0 | 0 | 0 | 4.15E-05 | 0 | 0 | 0.001025 | 0 | 0 | 0 | 0 | 0 | 0 | 0 | 0 | 0 | 0 | 0 | 0 |
| Clostridium:K00432 | 0 | 0 | 0 | 0.0016 | 0 | 0.06672 | 0.063994 | 0.0954044 | 0.094345 | 0 | 0 | 0.0003415 | 0 | 0 | 0 | 0 | 0 | 0 | 0 | 0 | 0 |
| Clostridium:K00525 | 0 | 0 | 0 | 0.0016 | 0 | 0.22497 | 0.233908 | 0.2685173 | 0.318929 | 0 | 0.001432 | 0.0022767 | 0.001118 | 0 | 0 | 0.00087 | 0 | 0 | 0 | 0 | 0 |
| Clostridium:K00526 | 0 | 0 | 0 | 0.0016 | 0 | 0.08281 | 0.041927 | 0.0446247 | 0.060504 | 0.0010704 | 0.000955 | 0.000683 | 0.001118 | 0.000418 | 0.000727 | 0 | 0 | 0 | 0 | 0 | 0 |
| Clostridium:K00548 | 0 | 0 | 0 | 0.0032 | 0 | 0.03898 | 0.244941 | 0.1892701 | 0.184589 | 0.0042816 | 0.008591 | 0.0050088 | 0.003353 | 0.015451 | 0.002906 | 0.00272 | 0.001974 | 0 | 0 | 0 | 0 |

| | | | | | | | | | | | | | | | | | | | | |
|---|---|---|---|---|---|---|---|---|---|---|---|---|---|---|---|---|---|---|---|---|
| Clostridium:K00575 | 0 | 0 | 0 | 0 | 0 | 0.00236 | 0.0662 | 0.0823248 | 0.076912 | 0 | 0.000477 | 0.0001138 | 0 | 0 | 0 | 0 | 0 | 0 | 0 | 0 | 0 |
| Clostridium:K00600 | 0 | 0 | 0 | 0.0145 | 0 | 0.09011 | 0.07944 | 0.1046371 | 0.090243 | 0.0139154 | 0.010977 | 0.0052365 | 0.029058 | 0.055121 | 0.000415 | 0.017081 | 0.059231 | 0 | 0 | 0 | 0.007893906 |
| Clostridium:K00626 | 0 | 0 | 0 | 0.0016 | 0 | 0.12963 | 0.163294 | 0.1038677 | 0.13434 | 0.0139154 | 0.007159 | 0.0070579 | 0.015646 | 0.005429 | 0.01256 | 0.002176 | 0.002962 | 0 | 0 | 0.042946103 | 0 |
| Clostridium:K00627 | 0 | 0 | 0 | 0 | 0 | 4.15E-05 | 0 | 0 | 0 | 0.0010704 | 0 | 0 | 0 | 0.00167 | 0 | 0 | 0 | 0 | 0 | 0 | 0 |
| Clostridium:K00640 | 0 | 0.0205 | 0 | 0.0016 | 0 | 0.03131 | 0.068407 | 0.0284675 | 0.02051 | 0.0021408 | 0.000955 | 0.0015937 | 0.005588 | 0 | 0.003737 | 0.002285 | 0.00691 | 0 | 0 | 0 | 0 |
| Clostridium:K00648 | 0 | 0 | 0 | 0 | 0 | 0.06527 | 0.189774 | 0.0530879 | 0.06153 | 0 | 0.000477 | 0.0002277 | 0.002235 | 0.002923 | 0.000623 | 0.001523 | 0 | 0.0373413 | 0 | 0 | 0 |
| Clostridium:K00681 | 0 | 0 | 0 | 0 | 0 | 0.00017 | 0 | 0 | 0 | 0.0010704 | 0.000477 | 0.0035289 | 0.003353 | 0.000835 | 0 | 0.001958 | 0 | 0 | 0 | 0 | 0 |
| Clostridium:K00798 | 0 | 0 | 0 | 0 | 0 | 0.03069 | 0.0331 | 0.0092327 | 0.012306 | 0.0010704 | 0 | 0.0004553 | 0 | 0.001253 | 0.000104 | 0.000218 | 0 | 0 | 0 | 0 | 0 |
| Clostridium:K00819 | 0 | 0 | 0 | 0 | 0 | 0 | 0 | 0.0007694 | 0 | 0 | 0 | 0 | 0 | 0 | 0 | 0 | 0 | 0 | 0 | 0 | 0 |
| Clostridium:K00833 | 0 | 0 | 0 | 0 | 0 | 0.00966 | 0.05296 | 0.1646496 | 0.133314 | 0 | 0 | 0.0005692 | 0 | 0.001253 | 0.000104 | 0.000435 | 0 | 0 | 0 | 0 | 0 |
| Clostridium:K00836 | 0 | 0 | 0 | 0 | 0 | 0 | 0 | 0 | 0 | 0 | 0.000477 | 0.0002277 | 0 | 0.002923 | 0 | 0.000653 | 0.002962 | 0 | 0 | 0 | 0 |
| Clostridium:K00864 | 0 | 0.0205 | 0 | 0.0113 | 0 | 0.18209 | 0.247148 | 0.1738823 | 0.190742 | 0.0139154 | 0.006205 | 0.0080824 | 0.00447 | 0.007099 | 0.001349 | 0.004352 | 0.002962 | 0 | 0 | 0.042946103 | 0 |
| Clostridium:K00940 | 0 | 0 | 0 | 0 | 0 | 0.02397 | 0.022067 | 0.0123102 | 0.008204 | 0 | 0 | 0 | 0 | 0 | 0 | 0 | 0 | 0 | 0 | 0 | 0 |
| Clostridium:K00955 | 0 | 0 | 0 | 0.0016 | 0 | 0.00012 | 0.002207 | 0.0061551 | 0.009229 | 0 | 0 | 0 | 0 | 0 | 0 | 0.000109 | 0 | 0 | 0 | 0.021473051 | 0 |
| Clostridium:K01011 | 0 | 0 | 0 | 0 | 0 | 0.06614 | 0 | 0.0253899 | 0.021535 | 0 | 0 | 0 | 0 | 0 | 0 | 0 | 0 | 0 | 0 | 0 | 0 |
| Clostridium:K01029 | 0 | 0 | 0 | 0 | 0 | 0.00029 | 0.011033 | 0.0115409 | 0.013331 | 0 | 0 | 0 | 0 | 0 | 0 | 0 | 0 | 0 | 0 | 0 | 0 |
| Clostridium:K01142 | 0 | 0.0205 | 0.0356 | 0.0048 | 0 | 0.04848 | 0.024273 | 0.0854024 | 0.090243 | 0.0064225 | 0.002864 | 0.0014799 | 0.013411 | 0.005011 | 0.001349 | 0.001088 | 0 | 0 | 0 | 0 | 0 |
| Clostridium:K01251 | 0 | 0 | 0 | 0.0065 | 0 | 0 | 0 | 0 | 0 | 0.0010704 | 0 | 0 | 0 | 0 | 0 | 0 | 0 | 0 | 0 | 0 | 0 |
| Clostridium:K01255 | 0 | 0 | 0 | 0 | 0 | 0.12714 | 0.01324 | 0.0615512 | 0.069734 | 0.0010704 | 0.000477 | 0.0005692 | 0 | 0.000418 | 0 | 0.000109 | 0 | 0 | 0 | 0.021473051 | 0 |
| Clostridium:K01338 | 0 | 0.0409 | 0 | 0.0129 | 0 | 0.37135 | 0.436922 | 0.5131835 | 0.528129 | 0.0107041 | 0.006682 | 0.0107007 | 0.02347 | 0.018791 | 0.035812 | 0.012511 | 0.017769 | 0 | 0 | 0.021473051 | 0 |
| Clostridium:K01428 | 0 | 0 | 0 | 0 | 0 | 0.00017 | 0 | 0 | 0 | 0 | 0.001909 | 0.0063749 | 0 | 0.000418 | 0 | 0.000762 | 0 | 0.0373413 | 0 | 0.021473051 | 0 |
| Clostridium:K01478 | 0 | 0 | 0 | 0.0048 | 0 | 0.05963 | 0.008827 | 0.0361614 | 0.034867 | 0.0010704 | 0.001432 | 0.0003415 | 0 | 0 | 0 | 0 | 0 | 0 | 0 | 0 | 0 |
| Clostridium:K01488 | 0 | 0 | 0 | 0.0016 | 0 | 0.08592 | 0.011033 | 0.045394 | 0.060504 | 0 | 0.000477 | 0 | 0 | 0 | 0 | 0.000218 | 0 | 0 | 0 | 0 | 0 |
| Clostridium:K01585 | 0 | 0 | 0 | 0.0016 | 0 | 0.00153 | 0.02648 | 0.0361614 | 0.030765 | 0 | 0 | 0 | 0 | 0 | 0 | 0 | 0 | 0 | 0 | 0 | 0 |
| Clostridium:K01613 | 0 | 0 | 0 | 0 | 0 | 0.09111 | 0.05958 | 0.0838636 | 0.095371 | 0 | 0 | 0 | 0 | 0.000835 | 0.000104 | 0 | 0 | 0 | 0 | 0 | 0 |
| Clostridium:K01629 | 0 | 0 | 0 | 0 | 0 | 0.01016 | 0.077234 | 0.1138698 | 0.087167 | 0.0010704 | 0.000955 | 0.0019352 | 0 | 0.002088 | 0.000104 | 0.001523 | 0.002962 | 0 | 0 | 0 | 0 |
| Clostridium:K01679 | 0 | 0 | 0 | 0 | 0 | 0 | 0 | 0 | 0 | 0 | 0 | 0.0001138 | 0 | 0 | 0 | 0 | 0 | 0 | 0 | 0 | 0 |
| Clostridium:K01698 | 0 | 0.0205 | 0 | 0.0032 | 0 | 0.05698 | 0.075027 | 0.0769391 | 0.083065 | 0 | 0.001432 | 0.0014799 | 0 | 0.002088 | 0 | 0 | 0.000987 | 0 | 0 | 0 | 0 |
| Clostridium:K01712 | 0 | 0 | 0 | 0 | 0 | 0.00365 | 0.05296 | 0.0800166 | 0.077938 | 0 | 0.002864 | 0.0011384 | 0.003353 | 0.00167 | 0 | 0.000326 | 0.000987 | 0 | 0 | 0 | 0 |
| Clostridium:K01737 | 0 | 0 | 0 | 0 | 0 | 0.00211 | 0.030893 | 0.0330838 | 0.023586 | 0 | 0 | 0 | 0 | 0.000835 | 0 | 0 | 0 | 0 | 0 | 0 | 0 |
| Clostridium:K01745 | 0 | 0 | 0 | 0 | 0 | 0.00464 | 0.055167 | 0.1030983 | 0.088192 | 0.0010704 | 0 | 0.0002277 | 0 | 0.000418 | 0.000104 | 0 | 0 | 0 | 0 | 0 | 0 |
| Clostridium:K01749 | 0 | 0 | 0 | 0 | 0 | 0.04926 | 0.057374 | 0.0284675 | 0.025637 | 0 | 0 | 0.0011384 | 0 | 0 | 0.000104 | 0.000218 | 0 | 0 | 0 | 0 | 0 |
| Clostridium:K01845 | 0 | 0 | 0 | 0.0355 | 0 | 0.08124 | 0.110334 | 0.1184861 | 0.136391 | 0.0010704 | 0.002386 | 0.0023906 | 0.002235 | 0.005011 | 0.000104 | 0.007942 | 0.024679 | 0 | 0 | 0 | 0 |
| Clostridium:K01879 | 0 | 0 | 0 | 0 | 0 | 4.15E-05 | 0 | 0 | 0 | 0 | 0 | 0 | 0 | 0 | 0 | 0 | 0 | 0 | 0 | 0 | 0 |
| Clostridium:K01895 | 0 | 0 | 0 | 0 | 0 | 4.15E-05 | 0 | 0 | 0 | 0.0032112 | 0.003818 | 0.0007969 | 0.010058 | 0.020044 | 0 | 0.005984 | 0.022705 | 0 | 0 | 0 | 0 |
| Clostridium:K01915 | 0 | 0.0205 | 0 | 0.0097 | 0 | 0.23222 | 0.158881 | 0.1938864 | 0.189716 | 0.0310419 | 0.021 | 0.0177585 | 0.046939 | 0.050945 | 0.002387 | 0.016972 | 0.044423 | 0 | 0 | 0.021473051 | 0 |
| Clostridium:K01941 | 0 | 0 | 0 | 0 | 0 | 0 | 0 | 0 | 0 | 0 | 0.000955 | 0.0001138 | 0 | 0.000418 | 0 | 0.000218 | 0 | 0 | 0 | 0 | 0 |
| Clostridium:K01958 | 0 | 0 | 0 | 0.0162 | 0 | 0.01708 | 0.260388 | 0.3185277 | 0.223558 | 0.0021408 | 0.002386 | 0.005578 | 0 | 0.006264 | 0.000623 | 0.00185 | 0.001974 | 0 | 0 | 0 | 0 |
| Clostridium:K01960 | 0 | 0 | 0 | 0 | 0 | 0 | 0 | 0 | 0 | 0 | 0 | 0.0001138 | 0 | 0 | 0 | 0 | 0 | 0 | 0 | 0 | 0 |
| Clostridium:K01961 | 0 | 0.0205 | 0 | 0 | 0 | 0.10064 | 0.108127 | 0.1000208 | 0.121008 | 0.0042816 | 0.010023 | 0.0020491 | 0.018999 | 0.029648 | 0.000415 | 0.011968 | 0.037513 | 0 | 0 | 0 | 0 |
| Clostridium:K01962 | 0 | 0 | 0 | 0.0016 | 0 | 0.03861 | 0.050754 | 0.0684758 | 0.048198 | 0.0010704 | 0.002386 | 0.0011384 | 0.003353 | 0.011275 | 0.000104 | 0.006093 | 0.01382 | 0 | 0 | 0.042946103 | 0 |
| Clostridium:K02011 | 0 | 0 | 0 | 0 | 0 | 0.01887 | 0.022067 | 0.0115409 | 0.018459 | 0 | 0.000477 | 0 | 0.002235 | 0 | 0.000727 | 0.000218 | 0 | 0 | 0 | 0 | 0 |
| Clostridium:K02012 | 0 | 0 | 0 | 0 | 0 | 0.16596 | 0.116954 | 0.0769391 | 0.070759 | 0 | 0 | 0.0004553 | 0.001118 | 0.000418 | 0.001038 | 0.000435 | 0.001974 | 0 | 0 | 0 | 0 |
| Clostridium:K02047 | 0 | 0 | 0 | 0.0032 | 0 | 0 | 0 | 0 | 0 | 0.0010704 | 0 | 0.0003415 | 0 | 0 | 0 | 0.000218 | 0.00691 | 0 | 0 | 0 | 0 |
| Clostridium:K02048 | 0 | 0 | 0 | 0 | 0 | 0 | 0 | 0 | 0 | 0 | 0.000477 | 0.0001138 | 0.002235 | 0.000835 | 0.000104 | 0.000544 | 0.005923 | 0 | 0 | 0 | 0 |
| Clostridium:K02049 | 0 | 0 | 0 | 0 | 0 | 0.04711 | 0.08606 | 0.06309 | 0.066657 | 0.0032112 | 0.000955 | 0.0027321 | 0.003353 | 0.007517 | 0.001349 | 0.001414 | 0.000987 | 0 | 0 | 0.021473051 | 0 |
| Clostridium:K02051 | 0 | 0 | 0 | 0 | 0 | 0.06934 | 0.05296 | 0.0907881 | 0.09332 | 0.0074929 | 0.003341 | 0.0030736 | 0.018999 | 0.002506 | 0.000415 | 0.003046 | 0.000987 | 0.074682599 | 0 | 0 | 0 |
| Clostridium:K02203 | 0 | 0 | 0 | 0.0016 | 0 | 0 | 0 | 0 | 0 | 0.0010704 | 0.000477 | 0.0004553 | 0 | 0.000418 | 0.000104 | 0.000109 | 0 | 0 | 0 | 0 | 0 |
| Clostridium:K02372 | 0 | 0 | 0 | 0 | 0 | 0.02538 | 0.041927 | 0.0792472 | 0.066657 | 0.0042816 | 0.000955 | 0.0003415 | 0.002235 | 0.001253 | 0.000934 | 0.003373 | 0.000987 | 0 | 0 | 0 | 0 |
| Clostridium:K02400 | 0 | 0 | 0 | 0 | 0 | 0.00717 | 0.165501 | 0.1446454 | 0.118957 | 0.0021408 | 0.001432 | 0.0007969 | 0 | 0 | 0 | 0.000326 | 0.000987 | 0 | 0 | 0 | 0 |
| Clostridium:K02405 | 0 | 0 | 0 | 0 | 0 | 0.00352 | 0.05958 | 0.0500104 | 0.050249 | 0 | 0 | 0.0001138 | 0 | 0 | 0 | 0 | 0 | 0 | 0 | 0 | 0 |
| Clostridium:K02409 | 0 | 0 | 0 | 0 | 0 | 0.00058 | 0.055167 | 0.0130796 | 0.009229 | 0 | 0 | 0.0001138 | 0 | 0 | 0 | 0 | 0 | 0 | 0 | 0 | 0.007893906 |
| Clostridium:K02410 | 0 | 0 | 0 | 0 | 0 | 0.0051 | 0.088267 | 0.1131004 | 0.114855 | 0 | 0.000477 | 0.0002277 | 0 | 0 | 0 | 0.000109 | 0 | 0 | 0 | 0 | 0 |
| Clostridium:K02412 | 0 | 0 | 0 | 0.0016 | 0 | 0.00742 | 0.139021 | 0.1323352 | 0.143569 | 0 | 0.000477 | 0.0005692 | 0.003353 | 0 | 0 | 0 | 0 | 0 | 0 | 0 | 0 |
| Clostridium:K02416 | 0 | 0 | 0 | 0 | 0 | 0.00614 | 0.083854 | 0.0969432 | 0.105626 | 0 | 0.000477 | 0.0001138 | 0.003353 | 0.000418 | 0 | 0.000109 | 0 | 0 | 0 | 0 | 0 |
| Clostridium:K02454 | 0 | 0 | 0 | 0 | 0 | 0.00095 | 0 | 0 | 0 | 0 | 0 | 0 | 0 | 0 | 0 | 0 | 0 | 0 | 0 | 0 | 0 |
| Clostridium:K02492 | 0 | 0 | 0 | 0 | 0 | 0.06262 | 0.05296 | 0.0161572 | 0.022561 | 0 | 0 | 0 | 0 | 0.00167 | 0 | 0 | 0.000987 | 0 | 0 | 0 | 0 |
| Clostridium:K02523 | 0 | 0 | 0 | 0 | 0 | 0.00033 | 0.00662 | 0.0146184 | 0.016408 | 0 | 0 | 0 | 0 | 0 | 0 | 0 | 0 | 0 | 0 | 0 | 0 |
| Clostridium:K02556 | 0 | 0 | 0 | 0 | 0 | 0.00249 | 0.0662 | 0.0338532 | 0.028714 | 0 | 0 | 0 | 0 | 0 | 0 | 0 | 0 | 0 | 0 | 0 | 0 |
| Clostridium:K02557 | 0 | 0 | 0 | 0 | 0 | 0.00066 | 0.035307 | 0.0130796 | 0.014357 | 0 | 0 | 0 | 0 | 0 | 0 | 0 | 0 | 0 | 0 | 0 | 0 |

| | | | | | | | | | | | | | | | | | | | | | |
|---|---|---|---|---|---|---|---|---|---|---|---|---|---|---|---|---|---|---|---|---|---|
| Clostridium:K03072 | 0 | 0 | 0 | 0 | 0 | 0.08194 | 0.103714 | 0.049241 | 0.0523 | 0 | 0.000955 | 0.0003415 | 0.001118 | 0.002088 | 0 | 0.002176 | 0.014808 | 0 | 0 | 0.021473051 | 0 |
| Clostridium:K03092 | 0 | 0 | 0 | 0 | 0 | 0.06892 | 0.04634 | 0.0523186 | 0.055377 | 0 | 0.000477 | 0.0001138 | 0 | 0 | 0.004983 | 0.001306 | 0.030602 | 0 | 0 | 0 | 0 |
| Clostridium:K03182 | 0 | 0 | 0 | 0.0016 | 0 | 0.00017 | 0 | 0 | 0 | 0.0010704 | 0.003341 | 0.0009107 | 0 | 0.002506 | 0.000415 | 0.000544 | 0.003949 | 0 | 0 | 0 | 0 |
| Clostridium:K03183 | 0 | 0 | 0 | 0 | 0 | 0.00182 | 0.03972 | 0.0646288 | 0.051275 | 0 | 0 | 0 | 0 | 0 | 0 | 0 | 0 | 0 | 0 | 0 | 0 |
| Clostridium:K03338 | 0 | 0 | 0 | 0 | 0 | 0.01791 | 0.004413 | 0.0076939 | 0.009229 | 0.0064225 | 0.002386 | 0.0014799 | 0.001118 | 0 | 0 | 0 | 0.000987 | 0 | 0 | 0 | 0 |
| Clostridium:K03394 | 0 | 0 | 0 | 0 | 0 | 0.03069 | 0.048547 | 0.0646288 | 0.069734 | 0 | 0 | 0.0003415 | 0 | 0.000418 | 0 | 0 | 0 | 0 | 0 | 0 | 0 |
| Clostridium:K03407 | 0 | 0 | 0 | 0 | 0 | 0.00701 | 0.152261 | 0.143876 | 0.127161 | 0 | 0 | 0.000683 | 0.001118 | 0 | 0 | 0.000109 | 0 | 0 | 0 | 0 | 0 |
| Clostridium:K03408 | 0 | 0 | 0 | 0 | 0 | 0.00041 | 0.02648 | 0.0238511 | 0.033841 | 0 | 0 | 0.0002277 | 0 | 0 | 0 | 0 | 0 | 0 | 0 | 0 | 0 |
| Clostridium:K03412 | 0 | 0 | 0 | 0 | 0 | 0.00199 | 0.101507 | 0.0707839 | 0.063581 | 0 | 0 | 0 | 0 | 0 | 0 | 0.000109 | 0 | 0 | 0 | 0 | 0 |
| Clostridium:K03415 | 0 | 0 | 0 | 0 | 0 | 0.00361 | 0.05958 | 0.0323144 | 0.036918 | 0 | 0 | 0 | 0 | 0 | 0 | 0 | 0 | 0 | 0 | 0 | 0 |
| Clostridium:K03590 | 0 | 0 | 0 | 0 | 0 | 0.0622 | 0.061787 | 0.0415471 | 0.046147 | 0 | 0 | 0 | 0 | 0 | 0 | 0 | 0 | 0 | 0 | 0 | 0 |
| Clostridium:K03644 | 0 | 0 | 0 | 0 | 0 | 0.00012 | 0 | 0.0400083 | 0.026663 | 0 | 0 | 0 | 0 | 0 | 0 | 0 | 0 | 0 | 0 | 0 | 0 |
| Clostridium:K03735 | 0 | 0 | 0 | 0 | 0 | 0.01004 | 0.004413 | 0.0677064 | 0.092294 | 0.0010704 | 0 | 0.0002277 | 0.001118 | 0.000418 | 0 | 0 | 0 | 0 | 0 | 0.021473051 | 0 |
| Clostridium:K03781 | 0 | 0 | 0 | 0 | 0 | 0.01767 | 0.008827 | 0.0230817 | 0.021535 | 0 | 0 | 0 | 0.001118 | 0 | 0 | 0 | 0 | 0 | 0 | 0 | 0 |
| Clostridium:K03885 | 0 | 0 | 0 | 0 | 0 | 0.00373 | 0 | 0 | 0 | 0 | 0 | 0 | 0 | 0 | 0 | 0 | 0 | 0 | 0 | 0 | 0 |
| Clostridium:K04564 | 0 | 0 | 0 | 0 | 0 | 0.0299 | 0.125781 | 0.080786 | 0.09332 | 0 | 0.000477 | 0.0001138 | 0 | 0 | 0 | 0.000762 | 0.001974 | 0 | 0 | 0 | 0.007893906 |
| Clostridium:K05343 | 0 | 0 | 0 | 0 | 0 | 0 | 0 | 0 | 0 | 0.0021408 | 0.001909 | 0.0027321 | 0.003353 | 0.00167 | 0.000311 | 0.000979 | 0.000987 | 0 | 0 | 0 | 0 |
| Clostridium:K05592 | 0 | 0.0409 | 0 | 0 | 0 | 0.12105 | 0.136814 | 0.1169474 | 0.111779 | 0.0042816 | 0.002864 | 0.0028459 | 0.005588 | 0.00167 | 0.000311 | 0.002937 | 0 | 0 | 0 | 0 | 0.007893906 |
| Clostridium:K05846 | 0 | 0 | 0 | 0 | 0 | 0.09496 | 0.194188 | 0.2200457 | 0.200997 | 0.0010704 | 0 | 0.0002277 | 0 | 0.00167 | 0 | 0 | 0 | 0 | 0 | 0 | 0.007893906 |
| Clostridium:K05847 | 0 | 0 | 0 | 0 | 0 | 0.0656 | 0.163294 | 0.291599 | 0.246119 | 0 | 0.000477 | 0.0005692 | 0 | 0.000835 | 0.000104 | 0.000109 | 0 | 0 | 0 | 0 | 0 |
| Clostridium:K06148 | 0 | 0 | 0 | 0 | 0 | 0.27672 | 0.05958 | 0.0523186 | 0.047173 | 0.0010704 | 0.001909 | 0.000683 | 0 | 0.001253 | 0 | 0 | 0 | 0 | 0 | 0 | 0 |
| Clostridium:K10536 | 0 | 0 | 0 | 0.0032 | 0 | 4.15E-05 | 0 | 0 | 0 | 0 | 0.000477 | 0.0001138 | 0 | 0.007099 | 0.000104 | 0.001306 | 0.000987 | 0 | 0 | 0 | 0 |
| Clostridium:K11927 | 0 | 0 | 0 | 0.0016 | 0 | 0.00124 | 0.017653 | 0.0423165 | 0.057428 | 0 | 0 | 0.000683 | 0 | 0 | 0.000104 | 0 | 0 | 0 | 0 | 0 | 0 |
| Clostridium:K11963 | 0 | 0 | 0 | 0 | 0 | 0 | 0 | 0 | 0 | 0 | 0 | 0.0001138 | 0 | 0 | 0 | 0 | 0 | 0 | 0 | 0 | 0 |
| Clostridium:K12573 | 0 | 0 | 0 | 0.0016 | 0 | 0.12258 | 0.167707 | 0.1600332 | 0.168181 | 0.0096337 | 0.006682 | 0.0044396 | 0.010058 | 0.008769 | 0.000623 | 0.001741 | 0 | 0 | 0 | 0 | 0 |
| Enterobacter:K00029 | 0 | 0 | 0 | 0 | 0 | 0 | 0 | 0 | 0 | 0 | 0 | 0.0002277 | 0 | 0 | 0 | 0 | 0 | 0 | 0 | 0 | 0 |
| Enterobacter:K00108 | 0 | 0 | 0 | 0 | 0 | 0.00021 | 0.008827 | 0.0076939 | 0.004102 | 0 | 0 | 0 | 0 | 0 | 0 | 0 | 0 | 0 | 0 | 0 | 0 |
| Enterobacter:K00111 | 0 | 0 | 0 | 0 | 0 | 0 | 0 | 0 | 0 | 0.0021408 | 0 | 0.0003415 | 0.001118 | 0 | 0 | 0 | 0 | 0 | 0 | 0 | 0 |
| Enterobacter:K00116 | 0 | 0 | 0 | 0 | 0 | 0 | 0 | 0 | 0 | 0 | 0 | 0.0001138 | 0 | 0 | 0 | 0 | 0 | 0 | 0 | 0 | 0 |
| Enterobacter:K00117 | 0 | 0 | 0 | 0 | 0 | 0 | 0 | 0 | 0 | 0 | 0 | 0.0001138 | 0 | 0 | 0 | 0 | 0 | 0 | 0 | 0 | 0 |
| Enterobacter:K00121 | 0 | 0 | 0 | 0.0016 | 0 | 0 | 0 | 0 | 0 | 0.0010704 | 0 | 0 | 0 | 0 | 0 | 0 | 0 | 0 | 0 | 0 | 0 |
| Enterobacter:K00123 | 0 | 0 | 0 | 0 | 0 | 0 | 0 | 0 | 0 | 0.0010704 | 0.000477 | 0.0003415 | 0 | 0 | 0 | 0 | 0 | 0 | 0 | 0 | 0 |
| Enterobacter:K00135 | 0 | 0 | 0 | 0 | 0 | 0 | 0 | 0 | 0 | 0.0010704 | 0 | 0.0002277 | 0 | 0 | 0 | 0 | 0 | 0 | 0 | 0 | 0 |
| Enterobacter:K00163 | 0 | 0 | 0 | 0 | 0 | 0 | 0 | 0 | 0.001025 | 0 | 0 | 0.0002277 | 0 | 0 | 0 | 0 | 0 | 0 | 0 | 0 | 0 |
| Enterobacter:K00164 | 0 | 0 | 0 | 0 | 0 | 0 | 0 | 0 | 0 | 0.0010704 | 0 | 0.0002277 | 0 | 0 | 0 | 0 | 0 | 0 | 0 | 0 | 0 |
| Enterobacter:K00281 | 0 | 0 | 0 | 0 | 0 | 0 | 0 | 0 | 0 | 0 | 0 | 0.0003415 | 0 | 0 | 0 | 0 | 0.000987 | 0 | 0 | 0 | 0 |
| Enterobacter:K00285 | 0 | 0 | 0 | 0.0081 | 0 | 0 | 0 | 0 | 0 | 0 | 0 | 0.0002277 | 0 | 0 | 0 | 0 | 0 | 0 | 0 | 0 | 0 |
| Enterobacter:K00322 | 0 | 0 | 0 | 0 | 0 | 0 | 0 | 0 | 0 | 0.0010704 | 0 | 0 | 0 | 0 | 0 | 0 | 0 | 0 | 0 | 0 | 0 |
| Enterobacter:K00331 | 0 | 0 | 0 | 0 | 0 | 0 | 0 | 0 | 0 | 0.0010704 | 0 | 0 | 0 | 0 | 0 | 0.000109 | 0 | 0 | 0 | 0 | 0 |
| Enterobacter:K00334 | 0 | 0 | 0 | 0 | 0 | 4.15E-05 | 0 | 0 | 0 | 0.0010704 | 0 | 0 | 0 | 0 | 0 | 0 | 0 | 0 | 0 | 0 | 0 |
| Enterobacter:K00336 | 0 | 0 | 0 | 0 | 0 | 0 | 0 | 0 | 0 | 0 | 0 | 0.0001138 | 0 | 0 | 0 | 0 | 0 | 0 | 0 | 0 | 0 |
| Enterobacter:K00341 | 0 | 0 | 0 | 0 | 0 | 0 | 0 | 0 | 0 | 0 | 0 | 0.0001138 | 0 | 0 | 0 | 0 | 0 | 0 | 0 | 0 | 0 |
| Enterobacter:K00342 | 0 | 0 | 0 | 0 | 0 | 0 | 0 | 0 | 0 | 0 | 0 | 0.0001138 | 0 | 0 | 0 | 0 | 0 | 0 | 0 | 0 | 0 |
| Enterobacter:K00362 | 0 | 0 | 0 | 0 | 0 | 0 | 0 | 0 | 0 | 0 | 0.000477 | 0.0002277 | 0 | 0 | 0 | 0 | 0 | 0 | 0 | 0 | 0 |
| Enterobacter:K00370 | 0 | 0 | 0 | 0 | 0 | 0 | 0 | 0 | 0.001025 | 0 | 0.000477 | 0.0004553 | 0 | 0 | 0 | 0 | 0.001974 | 0 | 0 | 0 | 0 |
| Enterobacter:K00371 | 0 | 0 | 0 | 0 | 0 | 0 | 0 | 0 | 0 | 0 | 0 | 0.0001138 | 0 | 0 | 0 | 0 | 0.000987 | 0 | 0 | 0 | 0 |
| Enterobacter:K00372 | 0 | 0 | 0 | 0 | 0 | 0 | 0 | 0 | 0 | 0 | 0 | 0.0002277 | 0 | 0 | 0 | 0 | 0 | 0 | 0 | 0 | 0 |
| Enterobacter:K00390 | 0 | 0 | 0 | 0 | 0 | 0 | 0 | 0 | 0 | 0.0010704 | 0 | 0 | 0 | 0 | 0 | 0 | 0 | 0 | 0 | 0 | 0 |
| Enterobacter:K00425 | 0 | 0 | 0 | 0.0032 | 0 | 0 | 0 | 0 | 0.001025 | 0 | 0.000955 | 0.0003415 | 0 | 0 | 0 | 0 | 0.000987 | 0 | 0 | 0 | 0 |
| Enterobacter:K00426 | 0 | 0 | 0 | 0 | 0 | 0 | 0 | 0 | 0 | 0 | 0 | 0.0005692 | 0 | 0 | 0 | 0 | 0 | 0 | 0 | 0 | 0 |
| Enterobacter:K00432 | 0 | 0 | 0 | 0 | 0 | 0 | 0 | 0 | 0 | 0 | 0 | 0.0001138 | 0 | 0 | 0 | 0 | 0 | 0 | 0 | 0 | 0 |
| Enterobacter:K00525 | 0 | 0 | 0 | 0 | 0 | 0 | 0 | 0 | 0 | 0 | 0.000477 | 0.0001138 | 0 | 0 | 0 | 0 | 0 | 0 | 0 | 0 | 0 |
| Enterobacter:K00526 | 0 | 0 | 0 | 0 | 0 | 0 | 0 | 0 | 0 | 0.0010704 | 0 | 0 | 0 | 0 | 0 | 0 | 0 | 0 | 0 | 0 | 0 |
| Enterobacter:K00568 | 0 | 0.0205 | 0.0356 | 0.0178 | 0 | 0 | 0 | 0 | 0 | 0 | 0 | 0 | 0 | 0 | 0 | 0 | 0 | 0 | 0 | 0 | 0 |
| Enterobacter:K00575 | 0 | 0 | 0 | 0 | 0 | 0 | 0 | 0 | 0 | 0 | 0 | 0 | 0.001118 | 0 | 0 | 0 | 0 | 0 | 0 | 0 | 0 |
| Enterobacter:K00626 | 0 | 0 | 0 | 0.0065 | 0 | 4.15E-05 | 0.00662 | 0.0046163 | 0 | 0 | 0 | 0.0001138 | 0.001118 | 0 | 0 | 0 | 0 | 0 | 0 | 0 | 0 |
| Enterobacter:K00631 | 0 | 0 | 0 | 0 | 0 | 0 | 0 | 0 | 0 | 0.0010704 | 0 | 0.0002277 | 0 | 0 | 0 | 0 | 0.000987 | 0 | 0 | 0 | 0 |
| Enterobacter:K00632 | 0 | 0 | 0 | 0 | 0 | 0 | 0 | 0 | 0 | 0 | 0 | 0 | 0 | 0 | 0 | 0 | 0.000987 | 0 | 0 | 0 | 0 |
| Enterobacter:K00640 | 0 | 0 | 0 | 0 | 0 | 0 | 0 | 0 | 0 | 0 | 0 | 0 | 0.001118 | 0 | 0 | 0 | 0 | 0 | 0 | 0 | 0 |
| Enterobacter:K00647 | 0 | 0 | 0 | 0 | 0 | 0 | 0 | 0 | 0 | 0 | 0.000477 | 0.0001138 | 0 | 0 | 0 | 0 | 0 | 0 | 0 | 0 | 0 |

| | | | | | | | | | | | | | | | | | | | | | |
|---|---|---|---|---|---|---|---|---|---|---|---|---|---|---|---|---|---|---|---|---|---|
| Enterobacter:K00648 | 0 | 0 | 0 | 0 | 0 | 0 | 0 | 0 | 0 | 0 | 0 | 0.0001138 | 0 | 0 | 0 | 0 | 0 | 0 | 0 | 0 | 0 |
| Enterobacter:K00681 | 0 | 0 | 0 | 0 | 0 | 0 | 0 | 0 | 0 | 0 | 0 | 0.0004553 | 0 | 0 | 0 | 0 | 0.000987 | 0 | 0 | 0 | 0 |
| Enterobacter:K00748 | 0 | 0 | 0 | 0.0048 | 0 | 0 | 0 | 0 | 0 | 0.0010704 | 0.000477 | 0 | 0 | 0 | 0 | 0 | 0 | 0 | 0 | 0 | 0 |
| Enterobacter:K00798 | 0 | 0 | 0 | 0 | 0 | 0 | 0 | 0 | 0 | 0 | 0 | 0.0001138 | 0 | 0 | 0 | 0 | 0 | 0 | 0 | 0 | 0 |
| Enterobacter:K00799 | 0 | 0 | 0 | 0 | 0 | 0 | 0 | 0 | 0 | 0 | 0 | 0.0004553 | 0 | 0 | 0 | 0 | 0 | 0 | 0 | 0 | 0 |
| Enterobacter:K00833 | 0 | 0 | 0.0356 | 0 | 0 | 0 | 0 | 0 | 0 | 0 | 0 | 0 | 0 | 0 | 0 | 0 | 0 | 0 | 0 | 0 | 0 |
| Enterobacter:K00864 | 0 | 0 | 0 | 0 | 0 | 0 | 0 | 0 | 0 | 0.0010704 | 0 | 0.0001138 | 0 | 0 | 0 | 0 | 0 | 0 | 0 | 0 | 0 |
| Enterobacter:K01029 | 0 | 0 | 0 | 0.0032 | 0 | 0 | 0 | 0 | 0 | 0 | 0 | 0 | 0 | 0 | 0 | 0 | 0 | 0 | 0 | 0 | 0 |
| Enterobacter:K01070 | 0 | 0 | 0 | 0.0032 | 0 | 0 | 0 | 0 | 0 | 0 | 0 | 0.0001138 | 0 | 0 | 0 | 0 | 0 | 0 | 0 | 0 | 0 |
| Enterobacter:K01139 | 0 | 0 | 0 | 0 | 0 | 0 | 0 | 0 | 0 | 0.0021408 | 0 | 0 | 0 | 0 | 0 | 0 | 0 | 0 | 0 | 0 | 0 |
| Enterobacter:K01142 | 0 | 0 | 0 | 0 | 0 | 0 | 0 | 0 | 0.001025 | 0 | 0 | 0 | 0 | 0 | 0 | 0 | 0 | 0 | 0 | 0 | 0 |
| Enterobacter:K01241 | 0 | 0 | 0 | 0 | 0 | 0 | 0 | 0 | 0 | 0.0010704 | 0 | 0 | 0 | 0 | 0 | 0 | 0 | 0 | 0 | 0 | 0 |
| Enterobacter:K01255 | 0 | 0 | 0 | 0.0032 | 0 | 0 | 0 | 0 | 0 | 0.0010704 | 0 | 0 | 0 | 0 | 0 | 0 | 0.001974 | 0 | 0 | 0 | 0 |
| Enterobacter:K01338 | 0 | 0 | 0 | 0 | 0 | 0 | 0 | 0.0007694 | 0 | 0 | 0 | 0 | 0 | 0 | 0 | 0 | 0 | 0 | 0 | 0 | 0 |
| Enterobacter:K01428 | 0 | 0 | 0 | 0.0065 | 0 | 0 | 0 | 0 | 0 | 0 | 0 | 0.0002277 | 0 | 0 | 0 | 0 | 0 | 0 | 0 | 0 | 0 |
| Enterobacter:K01478 | 0 | 0 | 0 | 0.0032 | 0 | 0 | 0 | 0 | 0 | 0 | 0 | 0.0003415 | 0 | 0 | 0 | 0 | 0 | 0 | 0 | 0 | 0 |
| Enterobacter:K01497 | 0 | 0 | 0 | 0 | 0 | 0 | 0 | 0 | 0 | 0 | 0 | 0.0001138 | 0 | 0 | 0 | 0 | 0 | 0 | 0 | 0 | 0 |
| Enterobacter:K01581 | 0 | 0 | 0 | 0 | 0 | 0 | 0.002207 | 0 | 0.001025 | 0 | 0.000955 | 0.0005692 | 0 | 0 | 0 | 0 | 0 | 0 | 0 | 0 | 0 |
| Enterobacter:K01585 | 0 | 0 | 0 | 0.0016 | 0 | 0 | 0 | 0 | 0 | 0.0032112 | 0 | 0.0003415 | 0 | 0 | 0 | 0 | 0 | 0 | 0 | 0 | 0 |
| Enterobacter:K01613 | 0 | 0 | 0 | 0.0016 | 0 | 0 | 0 | 0 | 0 | 0.0042816 | 0 | 0.0001138 | 0 | 0 | 0 | 0 | 0 | 0 | 0 | 0 | 0 |
| Enterobacter:K01629 | 0 | 0 | 0 | 0 | 0 | 0 | 0 | 0 | 0 | 0 | 0.000477 | 0.0001138 | 0 | 0 | 0 | 0 | 0 | 0 | 0 | 0 | 0 |
| Enterobacter:K01676 | 0 | 0 | 0 | 0 | 0 | 0 | 0 | 0 | 0 | 0 | 0 | 0.0002277 | 0 | 0 | 0 | 0 | 0 | 0 | 0 | 0 | 0 |
| Enterobacter:K01690 | 0 | 0 | 0 | 0 | 0 | 0 | 0 | 0 | 0 | 0 | 0 | 0.0001138 | 0 | 0 | 0 | 0 | 0.000987 | 0 | 0 | 0 | 0 |
| Enterobacter:K01698 | 0 | 0 | 0 | 0 | 0 | 0 | 0 | 0 | 0 | 0 | 0 | 0 | 0 | 0 | 0 | 0 | 0.001974 | 0 | 0 | 0 | 0 |
| Enterobacter:K01712 | 0 | 0 | 0 | 0 | 0 | 0 | 0 | 0 | 0 | 0 | 0 | 0.0001138 | 0 | 0 | 0 | 0 | 0 | 0 | 0 | 0 | 0 |
| Enterobacter:K01737 | 0 | 0 | 0 | 0 | 0 | 0 | 0.002207 | 0.0007694 | 0.001025 | 0 | 0 | 0 | 0 | 0 | 0 | 0 | 0 | 0 | 0 | 0 | 0 |
| Enterobacter:K01825 | 0 | 0 | 0 | 0.0016 | 0 | 0 | 0 | 0 | 0 | 0.0010704 | 0 | 0 | 0.001118 | 0 | 0 | 0 | 0 | 0 | 0 | 0 | 0 |
| Enterobacter:K01879 | 0 | 0 | 0 | 0 | 0 | 0 | 0 | 0 | 0 | 0 | 0.000477 | 0 | 0 | 0 | 0 | 0 | 0 | 0 | 0 | 0 | 0 |
| Enterobacter:K01895 | 0 | 0 | 0 | 0 | 0 | 0 | 0 | 0 | 0.001025 | 0 | 0 | 0.0001138 | 0 | 0 | 0 | 0 | 0 | 0 | 0 | 0 | 0 |
| Enterobacter:K01903 | 0 | 0 | 0 | 0 | 0 | 0 | 0 | 0 | 0 | 0.0010704 | 0 | 0 | 0 | 0 | 0 | 0 | 0 | 0 | 0 | 0 | 0 |
| Enterobacter:K01920 | 0 | 0 | 0 | 0 | 0 | 0 | 0 | 0 | 0 | 0 | 0 | 0.0001138 | 0 | 0 | 0 | 0 | 0 | 0 | 0 | 0 | 0 |
| Enterobacter:K01941 | 0 | 0 | 0 | 0.0016 | 0 | 0 | 0 | 0 | 0 | 0 | 0 | 0 | 0 | 0 | 0 | 0 | 0 | 0 | 0 | 0 | 0 |
| Enterobacter:K02000 | 0 | 0 | 0 | 0 | 0 | 0 | 0 | 0 | 0 | 0.0010704 | 0 | 0 | 0 | 0 | 0 | 0 | 0 | 0 | 0 | 0 | 0 |
| Enterobacter:K02011 | 0 | 0 | 0 | 0 | 0 | 0 | 0 | 0 | 0 | 0 | 0 | 0.0001138 | 0 | 0 | 0 | 0 | 0 | 0 | 0 | 0 | 0 |
| Enterobacter:K02012 | 0 | 0 | 0 | 0 | 0 | 0 | 0 | 0.0023082 | 0 | 0 | 0 | 0 | 0 | 0 | 0 | 0 | 0 | 0 | 0 | 0 | 0 |
| Enterobacter:K02047 | 0 | 0 | 0 | 0 | 0 | 0 | 0 | 0 | 0 | 0 | 0 | 0.0001138 | 0 | 0 | 0 | 0 | 0 | 0 | 0 | 0 | 0 |
| Enterobacter:K02048 | 0 | 0 | 0 | 0 | 0 | 0 | 0 | 0 | 0 | 0 | 0 | 0.0001138 | 0 | 0 | 0 | 0 | 0 | 0 | 0 | 0 | 0 |
| Enterobacter:K02049 | 0 | 0 | 0 | 0 | 0 | 4.15E-05 | 0 | 0.0007694 | 0.001025 | 0 | 0 | 0 | 0 | 0 | 0 | 0 | 0 | 0 | 0 | 0 | 0 |
| Enterobacter:K02301 | 0 | 0 | 0 | 0 | 0 | 0 | 0 | 0 | 0 | 0 | 0 | 0.0001138 | 0 | 0 | 0 | 0 | 0 | 0 | 0 | 0 | 0 |
| Enterobacter:K02372 | 0 | 0 | 0 | 0 | 0 | 0 | 0 | 0 | 0 | 0.0010704 | 0 | 0 | 0 | 0 | 0 | 0 | 0 | 0 | 0 | 0 | 0 |
| Enterobacter:K02405 | 0 | 0 | 0 | 0 | 0 | 0 | 0 | 0 | 0 | 0 | 0 | 0.0001138 | 0 | 0 | 0 | 0 | 0 | 0 | 0 | 0 | 0 |
| Enterobacter:K02409 | 0 | 0 | 0 | 0 | 0 | 0 | 0 | 0 | 0 | 0 | 0 | 0.0002277 | 0 | 0 | 0 | 0 | 0 | 0 | 0 | 0 | 0 |
| Enterobacter:K02410 | 0 | 0 | 0 | 0 | 0 | 0 | 0 | 0 | 0 | 0 | 0 | 0.0001138 | 0 | 0 | 0 | 0 | 0 | 0 | 0 | 0 | 0 |
| Enterobacter:K02412 | 0 | 0 | 0 | 0 | 0 | 0 | 0 | 0 | 0 | 0 | 0 | 0.0001138 | 0 | 0 | 0 | 0 | 0 | 0 | 0 | 0 | 0 |
| Enterobacter:K02416 | 0 | 0 | 0 | 0 | 0 | 0 | 0 | 0 | 0 | 0 | 0 | 0.0001138 | 0 | 0 | 0 | 0 | 0 | 0 | 0 | 0 | 0 |
| Enterobacter:K02492 | 0 | 0 | 0 | 0 | 0 | 0 | 0 | 0 | 0 | 0 | 0 | 0 | 0.001118 | 0 | 0 | 0 | 0.000987 | 0 | 0 | 0 | 0 |
| Enterobacter:K02517 | 0 | 0 | 0 | 0 | 0 | 0 | 0 | 0 | 0 | 0 | 0 | 0.0001138 | 0 | 0 | 0 | 0 | 0.000987 | 0 | 0 | 0 | 0 |
| Enterobacter:K02523 | 0 | 0 | 0 | 0 | 0 | 0 | 0 | 0 | 0 | 0.0010704 | 0.000477 | 0 | 0 | 0 | 0 | 0 | 0 | 0 | 0 | 0 | 0 |
| Enterobacter:K02556 | 0 | 0 | 0 | 0 | 0 | 0 | 0 | 0 | 0 | 0 | 0.000477 | 0 | 0 | 0 | 0 | 0 | 0 | 0 | 0 | 0 | 0 |
| Enterobacter:K02557 | 0 | 0 | 0 | 0 | 0 | 0 | 0 | 0 | 0 | 0.0010704 | 0.000477 | 0 | 0 | 0 | 0 | 0 | 0 | 0 | 0 | 0 | 0 |
| Enterobacter:K03072 | 0 | 0 | 0 | 0 | 0 | 0 | 0 | 0 | 0 | 0 | 0 | 0.0002277 | 0.001118 | 0 | 0 | 0 | 0 | 0 | 0 | 0 | 0 |
| Enterobacter:K03092 | 0 | 0 | 0 | 0 | 0 | 0 | 0 | 0 | 0 | 0 | 0 | 0.0004553 | 0 | 0 | 0 | 0 | 0 | 0 | 0 | 0 | 0 |
| Enterobacter:K03182 | 0 | 0 | 0 | 0 | 0 | 0 | 0 | 0 | 0 | 0 | 0 | 0.0001138 | 0 | 0 | 0 | 0 | 0 | 0 | 0 | 0 | 0 |
| Enterobacter:K03183 | 0 | 0.0205 | 0 | 0 | 0 | 0 | 0 | 0 | 0 | 0 | 0 | 0.0002277 | 0 | 0 | 0 | 0 | 0.000987 | 0 | 0 | 0 | 0 |
| Enterobacter:K03407 | 0 | 0 | 0 | 0 | 0 | 0 | 0 | 0 | 0 | 0 | 0 | 0.0001138 | 0 | 0 | 0 | 0 | 0 | 0 | 0 | 0 | 0 |
| Enterobacter:K03408 | 0 | 0 | 0 | 0 | 0 | 0 | 0 | 0 | 0 | 0 | 0 | 0.0001138 | 0 | 0 | 0 | 0 | 0 | 0 | 0 | 0 | 0 |
| Enterobacter:K03412 | 0 | 0 | 0 | 0 | 0 | 0 | 0 | 0 | 0 | 0 | 0 | 0.0001138 | 0.001118 | 0 | 0 | 0 | 0 | 0 | 0 | 0 | 0 |
| Enterobacter:K03415 | 0 | 0 | 0 | 0 | 0 | 0 | 0 | 0 | 0 | 0 | 0 | 0.0001138 | 0 | 0 | 0 | 0 | 0 | 0 | 0 | 0 | 0 |
| Enterobacter:K03582 | 0 | 0 | 0 | 0 | 0 | 0 | 0 | 0 | 0 | 0 | 0.000477 | 0 | 0 | 0 | 0 | 0 | 0 | 0 | 0 | 0 | 0 |
| Enterobacter:K03583 | 0 | 0 | 0 | 0 | 0 | 0 | 0 | 0 | 0 | 0 | 0 | 0.0005692 | 0 | 0 | 0 | 0 | 0 | 0 | 0 | 0 | 0 |

| | | | | | | | | | | | | | | | | | | | | | |
|---|---|---|---|---|---|---|---|---|---|---|---|---|---|---|---|---|---|---|---|---|---|
| Enterobacter:K03732 | 0 | 0 | 0 | 0 | 0 | 0 | 0 | 0 | 0 | 0.0010704 | 0 | 0 | 0 | 0 | 0 | 0 | 0 | 0 | 0 | 0 | 0 |
| Enterobacter:K03781 | 0 | 0 | 0 | 0 | 0 | 0 | 0 | 0 | 0 | 0 | 0 | 0.0003415 | 0 | 0 | 0 | 0 | 0 | 0 | 0 | 0 | 0 |
| Enterobacter:K03841 | 0 | 0 | 0 | 0 | 0 | 0 | 0 | 0 | 0 | 0.0021408 | 0 | 0 | 0 | 0 | 0 | 0 | 0 | 0 | 0 | 0 | 0 |
| Enterobacter:K03885 | 0 | 0 | 0.0713 | 0 | 0 | 0 | 0 | 0 | 0 | 0 | 0 | 0.0001138 | 0 | 0 | 0 | 0 | 0 | 0 | 0 | 0 | 0 |
| Enterobacter:K04564 | 0 | 0 | 0.0356 | 0 | 0 | 0 | 0 | 0 | 0 | 0 | 0 | 0 | 0 | 0 | 0 | 0 | 0 | 0 | 0 | 0 | 0 |
| Enterobacter:K05343 | 0 | 0 | 0 | 0 | 0 | 0 | 0 | 0 | 0 | 0 | 0.000477 | 0.0005692 | 0 | 0 | 0 | 0 | 0 | 0 | 0 | 0 | 0 |
| Enterobacter:K05365 | 0 | 0 | 0 | 0 | 0 | 0 | 0 | 0 | 0 | 0 | 0 | 0.0001138 | 0 | 0 | 0 | 0 | 0 | 0 | 0 | 0 | 0 |
| Enterobacter:K05592 | 0 | 0 | 0 | 0 | 0 | 0 | 0 | 0 | 0 | 0 | 0.000477 | 0 | 0 | 0 | 0 | 0 | 0 | 0 | 0 | 0 | 0 |
| Enterobacter:K06445 | 0 | 0 | 0 | 0 | 0 | 0 | 0 | 0 | 0 | 0.0021408 | 0 | 0.0002277 | 0 | 0 | 0 | 0 | 0 | 0 | 0 | 0 | 0 |
| Enterobacter:K06861 | 0 | 0 | 0 | 0 | 0 | 0 | 0 | 0 | 0 | 0.0010704 | 0 | 0 | 0 | 0 | 0 | 0 | 0 | 0 | 0 | 0 | 0 |
| Enterobacter:K07649 | 0 | 0.0205 | 0 | 0 | 0 | 0 | 0 | 0 | 0 | 0 | 0 | 0 | 0 | 0 | 0 | 0 | 0 | 0 | 0 | 0 | 0 |
| Enterobacter:K07712 | 0 | 0 | 0 | 0 | 0 | 0 | 0 | 0 | 0 | 0 | 0 | 0.0001138 | 0 | 0 | 0 | 0 | 0 | 0 | 0 | 0 | 0 |
| Enterobacter:K07788 | 0 | 0 | 0 | 0 | 0 | 0 | 0 | 0 | 0.001025 | 0 | 0 | 0 | 0 | 0 | 0 | 0 | 0 | 0 | 0 | 0 | 0 |
| Enterobacter:K08311 | 0 | 0 | 0 | 0 | 0 | 0 | 0 | 0 | 0 | 0.0010704 | 0 | 0.0002277 | 0 | 0 | 0 | 0 | 0 | 0 | 0 | 0 | 0 |
| Enterobacter:K08484 | 0 | 0 | 0 | 0 | 0 | 0 | 0 | 0 | 0 | 0 | 0 | 0.0002277 | 0 | 0 | 0 | 0 | 0.000987 | 0 | 0 | 0 | 0 |
| Enterobacter:K11891 | 0 | 0 | 0 | 0 | 0 | 0 | 0 | 0 | 0 | 0 | 0 | 0.0003415 | 0.001118 | 0 | 0 | 0 | 0.000987 | 0 | 0 | 0 | 0 |
| Enterobacter:K11903 | 0 | 0 | 0 | 0 | 0 | 0 | 0 | 0 | 0 | 0 | 0 | 0.0003415 | 0 | 0 | 0 | 0 | 0 | 0 | 0 | 0 | 0 |
| Enterobacter:K11904 | 0 | 0 | 0 | 0 | 0 | 0 | 0 | 0 | 0 | 0 | 0 | 0.0003415 | 0 | 0 | 0 | 0 | 0 | 0 | 0 | 0 | 0 |
| Enterobacter:K11959 | 0 | 0 | 0 | 0.0032 | 0 | 0 | 0.002207 | 0.0015388 | 0 | 0 | 0 | 0 | 0 | 0 | 0 | 0 | 0 | 0 | 0 | 0 | 0 |
| Enterobacter:K11962 | 0 | 0 | 0 | 0 | 0 | 0 | 0 | 0 | 0.001025 | 0 | 0 | 0 | 0 | 0 | 0 | 0 | 0 | 0 | 0 | 0 | 0 |
| Enterobacter:K11963 | 0 | 0 | 0 | 0 | 0 | 0 | 0 | 0 | 0 | 0 | 0 | 0.0001138 | 0 | 0 | 0 | 0 | 0 | 0 | 0 | 0 | 0 |
| Enterobacter:K12368 | 0 | 0 | 0 | 0 | 0 | 0 | 0 | 0 | 0 | 0 | 0 | 0.0005692 | 0 | 0 | 0 | 0 | 0.001974 | 0 | 0 | 0 | 0 |
| Enterobacter:K12573 | 0 | 0 | 0 | 0 | 0 | 0 | 0 | 0 | 0 | 0 | 0 | 0.0001138 | 0.001118 | 0 | 0 | 0 | 0 | 0 | 0 | 0 | 0 |
| Enterobacter:K13378 | 0 | 0 | 0 | 0 | 0 | 0 | 0 | 0 | 0 | 0 | 0.000477 | 0.0001138 | 0 | 0 | 0 | 0 | 0 | 0 | 0 | 0 | 0 |
| Enterococcus:K00036 | 0 | 0 | 0 | 0 | 0 | 0 | 0 | 0 | 0 | 0 | 0 | 0 | 0 | 0.000835 | 0 | 0 | 0 | 0 | 0.052077909 | 0 | 0 |
| Enterococcus:K00104 | 0 | 0 | 0 | 0 | 0 | 0 | 0 | 0 | 0 | 0 | 0 | 0.0001138 | 0 | 0 | 0 | 0 | 0 | 0 | 0 | 0 | 0 |
| Enterococcus:K00111 | 0 | 0 | 0 | 0 | 0 | 0 | 0 | 0 | 0 | 0 | 0.000477 | 0 | 0 | 0 | 0 | 0.000109 | 0 | 0 | 0 | 0 | 0 |
| Enterococcus:K00121 | 0 | 0 | 0 | 0 | 0 | 0 | 0 | 0 | 0 | 0 | 0 | 0 | 0 | 0.000418 | 0 | 0 | 0 | 0 | 0.003471861 | 0 | 0 |
| Enterococcus:K00123 | 0 | 0 | 0 | 0 | 0 | 8.29E-05 | 0 | 0 | 0 | 0 | 0 | 0.0001138 | 0 | 0 | 0 | 0 | 0 | 0 | 0 | 0 | 0 |
| Enterococcus:K00135 | 0 | 0 | 0 | 0 | 0 | 0 | 0 | 0 | 0 | 0 | 0 | 0 | 0 | 0 | 0 | 0 | 0.000987 | 0 | 0.076380933 | 0 | 0 |
| Enterococcus:K00161 | 0 | 0 | 0 | 0.0016 | 0 | 0 | 0 | 0 | 0 | 0 | 0 | 0.0001138 | 0 | 0.00167 | 0 | 0 | 0 | 0 | 0.027774885 | 0 | 0 |
| Enterococcus:K00162 | 0 | 0 | 0 | 0 | 0 | 0 | 0 | 0 | 0 | 0 | 0 | 0 | 0 | 0.001253 | 0 | 0 | 0 | 0 | 0.027774885 | 0 | 0 |
| Enterococcus:K00383 | 0 | 0 | 0 | 0.0016 | 0 | 0 | 0 | 0 | 0 | 0 | 0 | 0 | 0 | 0.001253 | 0 | 0 | 0 | 0 | 0.083324654 | 0 | 0 |
| Enterococcus:K00432 | 0 | 0 | 0 | 0 | 0 | 0 | 0 | 0 | 0 | 0 | 0 | 0 | 0 | 0.001253 | 0 | 0 | 0 | 0 | 0.038190466 | 0 | 0 |
| Enterococcus:K00525 | 0 | 0 | 0 | 0.0016 | 0 | 0 | 0 | 0 | 0 | 0 | 0 | 0 | 0 | 0.003758 | 0 | 0 | 0.000987 | 0 | 0.173593029 | 0 | 0 |
| Enterococcus:K00526 | 0 | 0.0205 | 0 | 0 | 0 | 0 | 0 | 0 | 0 | 0 | 0 | 0 | 0 | 0 | 0 | 0 | 0 | 0 | 0.031246745 | 0 | 0 |
| Enterococcus:K00600 | 0 | 0 | 0 | 0.0016 | 0 | 4.15E-05 | 0 | 0 | 0 | 0.0021408 | 0.000955 | 0.0001138 | 0 | 0.002088 | 0 | 0 | 0.000987 | 0 | 0.055549769 | 0 | 0 |
| Enterococcus:K00626 | 0 | 0 | 0 | 0 | 0 | 0 | 0 | 0 | 0 | 0 | 0 | 0 | 0 | 0 | 0 | 0 | 0 | 0 | 0.045134187 | 0 | 0 |
| Enterococcus:K00627 | 0 | 0 | 0 | 0 | 0 | 0 | 0 | 0 | 0 | 0 | 0 | 0 | 0 | 0.002088 | 0 | 0 | 0 | 0 | 0.045134187 | 0 | 0 |
| Enterococcus:K00640 | 0 | 0 | 0 | 0 | 0 | 0 | 0 | 0 | 0 | 0 | 0 | 0 | 0 | 0.002506 | 0 | 0 | 0 | 0 | 0.017359303 | 0 | 0 |
| Enterococcus:K00648 | 0 | 0 | 0 | 0 | 0 | 0 | 0 | 0 | 0 | 0 | 0 | 0.0001138 | 0 | 0.001253 | 0 | 0 | 0 | 0 | 0.017359303 | 0 | 0 |
| Enterococcus:K00864 | 0 | 0 | 0 | 0 | 0 | 0 | 0 | 0 | 0 | 0 | 0 | 0.0001138 | 0 | 0.000835 | 0 | 0 | 0 | 0 | 0.020831163 | 0 | 0 |
| Enterococcus:K00940 | 0 | 0 | 0 | 0 | 0 | 0 | 0 | 0 | 0 | 0 | 0 | 0 | 0 | 0.002506 | 0 | 0 | 0 | 0 | 0.038190466 | 0 | 0 |
| Enterococcus:K01142 | 0 | 0 | 0 | 0.0048 | 0 | 0 | 0 | 0 | 0 | 0.0010704 | 0.002864 | 0.0018214 | 0.002235 | 0.001253 | 0 | 0.000218 | 0.000987 | 0 | 0.048606048 | 0 | 0 |
| Enterococcus:K01426 | 0 | 0 | 0 | 0.0016 | 0 | 4.15E-05 | 0 | 0 | 0 | 0 | 0 | 0 | 0 | 0.00167 | 0.000104 | 0 | 0 | 0 | 0.05902163 | 0 | 0 |
| Enterococcus:K01478 | 0 | 0 | 0 | 0 | 0 | 0 | 0 | 0 | 0 | 0 | 0.000477 | 0.0005692 | 0 | 0.002506 | 0.000104 | 0.000109 | 0 | 0 | 0.041662327 | 0 | 0 |
| Enterococcus:K01629 | 0 | 0 | 0 | 0 | 0 | 0 | 0 | 0 | 0 | 0 | 0 | 0.0001138 | 0 | 0 | 0 | 0 | 0 | 0 | 0 | 0 | 0 |
| Enterococcus:K01878 | 0 | 0 | 0 | 0 | 0 | 0 | 0 | 0 | 0 | 0 | 0 | 0 | 0 | 0.000418 | 0 | 0 | 0 | 0 | 0.027774885 | 0 | 0 |
| Enterococcus:K01879 | 0 | 0 | 0 | 0 | 0 | 4.15E-05 | 0 | 0 | 0 | 0 | 0.000477 | 0 | 0 | 0.000835 | 0 | 0 | 0 | 0 | 0.086796514 | 0 | 0 |
| Enterococcus:K01915 | 0 | 0 | 0 | 0.0081 | 0 | 4.15E-05 | 0 | 0 | 0 | 0 | 0 | 0 | 0 | 0.004593 | 0 | 0 | 0 | 0 | 0.052077909 | 0 | 0 |
| Enterococcus:K01958 | 0 | 0.0205 | 0 | 0.0016 | 0 | 0 | 0 | 0.0007694 | 0 | 0 | 0 | 0 | 0 | 0.01044 | 0 | 0.000218 | 0.001974 | 0 | 0.145818144 | 0 | 0 |
| Enterococcus:K01961 | 0 | 0 | 0 | 0.0016 | 0 | 0 | 0 | 0 | 0 | 0 | 0 | 0.0001138 | 0 | 0.002088 | 0.000104 | 0 | 0.000987 | 0 | 0.003471861 | 0 | 0 |
| Enterococcus:K01962 | 0 | 0 | 0.0356 | 0 | 0 | 0 | 0 | 0 | 0 | 0 | 0 | 0.0002277 | 0.001118 | 0.001253 | 0 | 0.000435 | 0.003949 | 0 | 0.013887442 | 0 | 0 |
| Enterococcus:K02000 | 0 | 0 | 0 | 0 | 0 | 0 | 0 | 0 | 0 | 0 | 0 | 0 | 0 | 0.001253 | 0 | 0 | 0 | 0 | 0.05902163 | 0 | 0 |
| Enterococcus:K02002 | 0 | 0 | 0 | 0 | 0 | 0 | 0 | 0 | 0 | 0 | 0 | 0 | 0 | 0.000835 | 0 | 0 | 0 | 0 | 0 | 0 | 0 |
| Enterococcus:K02012 | 0 | 0 | 0 | 0 | 0 | 0 | 0 | 0 | 0 | 0 | 0 | 0 | 0 | 0 | 0 | 0.000218 | 0 | 0 | 0 | 0 | 0 |
| Enterococcus:K02049 | 0 | 0 | 0 | 0 | 0 | 0 | 0 | 0 | 0 | 0 | 0 | 0 | 0 | 0.000418 | 0 | 0 | 0 | 0 | 0 | 0 | 0 |
| Enterococcus:K02372 | 0 | 0 | 0 | 0 | 0 | 0 | 0 | 0 | 0 | 0.0010704 | 0 | 0 | 0 | 0.000418 | 0 | 0 | 0 | 0 | 0.034718606 | 0 | 0 |
| Enterococcus:K02557 | 0 | 0 | 0 | 0 | 0 | 4.15E-05 | 0 | 0 | 0 | 0 | 0 | 0 | 0 | 0 | 0 | 0 | 0 | 0 | 0 | 0 | 0 |
| Enterococcus:K03092 | 0 | 0 | 0 | 0.0016 | 0 | 8.29E-05 | 0 | 0 | 0 | 0.0010704 | 0 | 0 | 0 | 0.003758 | 0 | 0 | 0 | 0 | 0.048606048 | 0 | 0 |

| | | | | | | | | | | | | | | | | | | | | | |
|---|---|---|---|---|---|---|---|---|---|---|---|---|---|---|---|---|---|---|---|---|---|
| Enterococcus:K03182 | 0 | 0 | 0 | 0 | 0 | 0 | 0 | 0 | 0 | 0 | 0 | 0 | 0 | 0.002506 | 0 | 0 | 0 | 0 | 0 | 0 | 0 |
| Enterococcus:K03338 | 0 | 0 | 0 | 0.0048 | 0 | 0 | 0 | 0 | 0 | 0 | 0.000477 | 0.0003415 | 0 | 0 | 0 | 0 | 0 | 0 | 0 | 0 | 0 |
| Enterococcus:K03590 | 0 | 0 | 0 | 0 | 0 | 0 | 0 | 0 | 0 | 0 | 0 | 0 | 0 | 0.004176 | 0 | 0 | 0 | 0 | 0.05902163 | 0 | 0 |
| Enterococcus:K03885 | 0 | 0 | 0 | 0.0016 | 0 | 4.15E-05 | 0 | 0 | 0 | 0 | 0 | 0 | 0 | 0.007099 | 0 | 0.000109 | 0 | 0 | 0.111099538 | 0 | 0 |
| Enterococcus:K04564 | 0 | 0 | 0 | 0 | 0 | 0 | 0 | 0 | 0 | 0 | 0 | 0 | 0 | 0 | 0 | 0 | 0.000987 | 0 | 0.013887442 | 0 | 0 |
| Enterococcus:K05592 | 0 | 0 | 0 | 0 | 0 | 0 | 0 | 0 | 0 | 0 | 0 | 0 | 0 | 0.003758 | 0 | 0 | 0 | 0 | 0.104155817 | 0 | 0 |
| Enterococcus:K05846 | 0 | 0 | 0 | 0 | 0 | 0 | 0 | 0 | 0 | 0 | 0 | 0 | 0 | 0.000835 | 0 | 0 | 0 | 0 | 0.055549769 | 0 | 0 |
| Enterococcus:K05847 | 0 | 0 | 0 | 0 | 0 | 0 | 0 | 0 | 0 | 0 | 0 | 0 | 0 | 0.00167 | 0 | 0 | 0 | 0 | 0.017359303 | 0 | 0 |
| Enterococcus:K10536 | 0 | 0 | 0 | 0.0032 | 0 | 4.15E-05 | 0 | 0 | 0 | 0.0021408 | 0 | 0.0005692 | 0.003353 | 0.00167 | 0 | 0.001088 | 0.002962 | 0 | 0.031246745 | 0 | 0 |
| Enterococcus:K10941 | 0 | 0 | 0 | 0 | 0 | 0 | 0 | 0 | 0 | 0 | 0 | 0.0001138 | 0 | 0 | 0 | 0 | 0 | 0 | 0 | 0 | 0 |
| Enterococcus:K12573 | 0 | 0 | 0 | 0.0048 | 0 | 4.15E-05 | 0 | 0 | 0 | 0 | 0 | 0 | 0 | 0.001253 | 0 | 0 | 0 | 0 | 0.072909072 | 0 | 0 |
| Eubacterium:K00029 | 0 | 0 | 0 | 0 | 0 | 0 | 0 | 0 | 0 | 0 | 0 | 0 | 0 | 0 | 0.000104 | 0 | 0 | 0 | 0 | 0 | 0 |
| Eubacterium:K00104 | 0 | 0 | 0 | 0 | 0 | 0 | 0 | 0 | 0 | 0.0021408 | 0.001432 | 0.0019352 | 0.002235 | 0 | 0.002699 | 0.002285 | 0.000987 | 0 | 0 | 0 | 0 |
| Eubacterium:K00111 | 0 | 0 | 0 | 0 | 0 | 0 | 0 | 0 | 0 | 0.0074929 | 0.002864 | 0.0066025 | 0.002235 | 0.018374 | 0.004048 | 0.006419 | 0.017769 | 0 | 0 | 0.021473051 | 0 |
| Eubacterium:K00123 | 0 | 0 | 0 | 0.0178 | 0 | 0 | 0 | 0 | 0 | 0.0139154 | 0.001432 | 0.0054642 | 0.003353 | 0.00167 | 0.000623 | 0.003155 | 0.000987 | 0 | 0 | 0 | 0 |
| Eubacterium:K00128 | 0 | 0 | 0 | 0 | 0 | 0 | 0 | 0 | 0 | 0.0010704 | 0 | 0.0033013 | 0.001118 | 0.001253 | 0.000104 | 0.001958 | 0 | 0 | 0 | 0 | 0 |
| Eubacterium:K00248 | 0 | 0 | 0 | 0.0016 | 0 | 0 | 0 | 0 | 0 | 0.0010704 | 0 | 0.0004553 | 0 | 0 | 0.000311 | 0.000653 | 0 | 0 | 0 | 0 | 0 |
| Eubacterium:K00334 | 0 | 0 | 0 | 0.0097 | 0 | 0 | 0 | 0 | 0 | 0 | 0 | 0 | 0 | 0 | 0 | 0 | 0 | 0 | 0 | 0 | 0 |
| Eubacterium:K00335 | 0 | 0 | 0 | 0 | 0 | 0 | 0 | 0 | 0 | 0 | 0 | 0.0004553 | 0.002235 | 0.000418 | 0.000623 | 0.000653 | 0 | 0 | 0 | 0 | 0 |
| Eubacterium:K00336 | 0 | 0 | 0 | 0 | 0 | 0 | 0 | 0 | 0 | 0.0032112 | 0.000955 | 0.0076271 | 0.001118 | 0.014198 | 0.00955 | 0.011641 | 0.000987 | 0 | 0 | 0 | 0 |
| Eubacterium:K00390 | 0 | 0 | 0 | 0 | 0 | 4.15E-05 | 0 | 0 | 0 | 0 | 0.000477 | 0.0004553 | 0 | 0 | 0.000104 | 0 | 0 | 0 | 0 | 0 | 0 |
| Eubacterium:K00548 | 0 | 0 | 0 | 0 | 0 | 0.00021 | 0 | 0 | 0 | 0.0117745 | 0.002864 | 0.005578 | 0.003353 | 0.009604 | 0.002906 | 0.010444 | 0.001974 | 0 | 0 | 0 | 0 |
| Eubacterium:K00575 | 0 | 0 | 0 | 0 | 0 | 0 | 0 | 0 | 0 | 0.0064225 | 0.000955 | 0.0014799 | 0 | 0.003758 | 0.000934 | 0.001523 | 0 | 0 | 0 | 0 | 0 |
| Eubacterium:K00600 | 0 | 0 | 0 | 0.0016 | 0 | 0 | 0 | 0 | 0 | 0.0064225 | 0.000955 | 0.0030736 | 0.005588 | 0.001253 | 0.00737 | 0.0099 | 0.000987 | 0 | 0 | 0.021473051 | 0 |
| Eubacterium:K00626 | 0 | 0 | 0 | 0.0291 | 0 | 0 | 0 | 0 | 0 | 0.0032112 | 0.000955 | 0.0027321 | 0.001118 | 0.001253 | 0.007059 | 0.004678 | 0.001974 | 0 | 0 | 0 | 0 |
| Eubacterium:K00640 | 0 | 0 | 0 | 0 | 0 | 8.29E-05 | 0 | 0 | 0 | 0.0096337 | 0.003341 | 0.0037566 | 0.003353 | 0.003341 | 0.00083 | 0.00544 | 0.033564 | 0 | 0 | 0.021473051 | 0 |
| Eubacterium:K00648 | 0 | 0 | 0 | 0 | 0 | 0 | 0 | 0 | 0 | 0.0042816 | 0 | 0.0022767 | 0 | 0.006681 | 0.000415 | 0.001306 | 0.000987 | 0 | 0 | 0 | 0 |
| Eubacterium:K00798 | 0 | 0 | 0 | 0 | 0 | 0 | 0 | 0 | 0 | 0 | 0 | 0 | 0 | 0 | 0.000104 | 0.000979 | 0 | 0 | 0 | 0 | 0 |
| Eubacterium:K00864 | 0 | 0 | 0 | 0 | 0 | 0 | 0 | 0 | 0 | 0.0032112 | 0.000477 | 0.0059195 | 0.001118 | 0.020462 | 0.007785 | 0.013491 | 0.018756 | 0 | 0 | 0 | 0 |
| Eubacterium:K00955 | 0 | 0 | 0 | 0 | 0 | 0 | 0 | 0 | 0 | 0 | 0 | 0.0017076 | 0 | 0 | 0 | 0 | 0.000987 | 0 | 0 | 0 | 0 |
| Eubacterium:K01142 | 0 | 0 | 0 | 0 | 0 | 4.15E-05 | 0 | 0 | 0 | 0.0021408 | 0.000477 | 0.0009107 | 0.001118 | 0.01044 | 0.000519 | 0.001197 | 0.000987 | 0 | 0 | 0 | 0 |
| Eubacterium:K01251 | 0 | 0 | 0 | 0 | 0 | 4.15E-05 | 0 | 0 | 0 | 0.0224786 | 0.001432 | 0.0064887 | 0.005588 | 0.004176 | 0.003633 | 0.003917 | 0 | 0 | 0 | 0.021473051 | 0 |
| Eubacterium:K01255 | 0 | 0 | 0 | 0 | 0 | 0 | 0 | 0 | 0 | 0 | 0 | 0.0001138 | 0 | 0.002088 | 0.000208 | 0.000326 | 0 | 0 | 0 | 0 | 0 |
| Eubacterium:K01338 | 0 | 0 | 0 | 0 | 0 | 0.00012 | 0 | 0 | 0 | 0.0267603 | 0.004773 | 0.0168478 | 0.008941 | 0.01378 | 0.009861 | 0.017842 | 0.015795 | 0 | 0 | 0.021473051 | 0 |
| Eubacterium:K01478 | 0 | 0 | 0 | 0.0016 | 0 | 0 | 0 | 0 | 0 | 0.0010704 | 0 | 0.0002277 | 0 | 0 | 0.000311 | 0.000109 | 0.000987 | 0 | 0 | 0 | 0 |
| Eubacterium:K01488 | 0 | 0 | 0 | 0 | 0 | 0 | 0 | 0 | 0 | 0 | 0.000955 | 0.0018214 | 0 | 0.000835 | 0 | 0.00272 | 0.000987 | 0 | 0 | 0 | 0 |
| Eubacterium:K01629 | 0 | 0 | 0.0356 | 0 | 0 | 0 | 0 | 0 | 0 | 0.0107041 | 0.002864 | 0.0021629 | 0.002235 | 0.000835 | 0.000519 | 0.006963 | 0.002962 | 0 | 0 | 0 | 0 |
| Eubacterium:K01679 | 0 | 0 | 0 | 0 | 0 | 0 | 0 | 0 | 0 | 0 | 0 | 0 | 0 | 0.000418 | 0.000104 | 0.000109 | 0.000987 | 0 | 0 | 0 | 0 |
| Eubacterium:K01698 | 0 | 0 | 0 | 0 | 0 | 0 | 0 | 0 | 0 | 0.0032112 | 0.000477 | 0.001366 | 0.002235 | 0.004176 | 0.001038 | 0.004569 | 0.002962 | 0 | 0 | 0 | 0 |
| Eubacterium:K01712 | 0 | 0 | 0 | 0.0016 | 0 | 0 | 0 | 0 | 0 | 0.0010704 | 0.000955 | 0.0003415 | 0 | 0 | 0.000104 | 0 | 0 | 0 | 0 | 0 | 0 |
| Eubacterium:K01737 | 0 | 0 | 0 | 0 | 0 | 0 | 0 | 0 | 0 | 0.0021408 | 0 | 0.0003415 | 0 | 0 | 0.000208 | 0.002394 | 0 | 0 | 0 | 0 | 0 |
| Eubacterium:K01749 | 0 | 0 | 0 | 0 | 0 | 0 | 0 | 0 | 0 | 0.0021408 | 0 | 0.0019352 | 0.001118 | 0.002923 | 0.000415 | 0.00272 | 0 | 0 | 0 | 0 | 0 |
| Eubacterium:K01845 | 0 | 0 | 0 | 0 | 0 | 4.15E-05 | 0 | 0 | 0 | 0.0010704 | 0.002386 | 0.004895 | 0.003353 | 0.003758 | 0.001038 | 0.010336 | 0.000987 | 0 | 0 | 0 | 0 |
| Eubacterium:K01895 | 0 | 0 | 0 | 0 | 0 | 0 | 0 | 0 | 0 | 0.0149858 | 0.001432 | 0.0020491 | 0 | 0.004593 | 0 | 0 | 0 | 0 | 0 | 0 | 0 |
| Eubacterium:K01915 | 0 | 0 | 0 | 0.0048 | 0 | 0.00025 | 0 | 0 | 0 | 0.0214082 | 0.009068 | 0.0166202 | 0.012294 | 0.01044 | 0.012352 | 0.025893 | 0.005923 | 0 | 0 | 0.042946103 | 0 |
| Eubacterium:K01958 | 0 | 0 | 0 | 0 | 0 | 8.29E-05 | 0 | 0 | 0 | 0 | 0.003341 | 0.004895 | 0.002235 | 0.007517 | 0.004463 | 0.003917 | 0.001974 | 0 | 0 | 0 | 0 |
| Eubacterium:K01959 | 0 | 0 | 0 | 0 | 0 | 0 | 0 | 0 | 0 | 0 | 0 | 0 | 0 | 0 | 0.000208 | 0 | 0.000987 | 0 | 0 | 0 | 0 |
| Eubacterium:K01960 | 0 | 0 | 0 | 0.0032 | 0 | 0 | 0 | 0 | 0 | 0 | 0 | 0 | 0 | 0 | 0 | 0 | 0 | 0 | 0 | 0 | 0 |
| Eubacterium:K01961 | 0 | 0 | 0 | 0 | 0 | 0.00025 | 0 | 0 | 0.001025 | 0.0107041 | 0.004295 | 0.0093346 | 0.001118 | 0.010857 | 0.005709 | 0.005766 | 0.009872 | 0 | 0 | 0 | 0 |
| Eubacterium:K01962 | 0 | 0 | 0 | 0 | 0 | 0 | 0 | 0 | 0 | 0.0064225 | 0.001432 | 0.0045535 | 0.001118 | 0.002923 | 0.000208 | 0.003373 | 0.000987 | 0 | 0 | 0 | 0 |
| Eubacterium:K02000 | 0 | 0 | 0 | 0 | 0 | 0 | 0 | 0 | 0 | 0 | 0 | 0.0001138 | 0 | 0 | 0 | 0 | 0 | 0 | 0 | 0 | 0 |
| Eubacterium:K02047 | 0 | 0 | 0 | 0 | 0 | 0 | 0 | 0 | 0 | 0.0032112 | 0.000955 | 0.0061472 | 0 | 0.000835 | 0 | 0.002176 | 0.000987 | 0 | 0 | 0 | 0 |
| Eubacterium:K02048 | 0 | 0 | 0 | 0 | 0 | 4.15E-05 | 0 | 0 | 0 | 0.0021408 | 0.001432 | 0.0047811 | 0 | 0.002506 | 0 | 0.002937 | 0.005923 | 0 | 0 | 0 | 0 |
| Eubacterium:K02049 | 0 | 0 | 0 | 0 | 0 | 0 | 0 | 0 | 0 | 0.0021408 | 0.001909 | 0.0019352 | 0.001118 | 0.002506 | 0.000519 | 0.004352 | 0.000987 | 0 | 0 | 0 | 0 |
| Eubacterium:K02051 | 0 | 0 | 0 | 0 | 0 | 4.15E-05 | 0 | 0 | 0 | 0.0074929 | 0.00525 | 0.0043258 | 0.002235 | 0.008352 | 0.000519 | 0.018713 | 0.000987 | 0 | 0 | 0 | 0 |
| Eubacterium:K02203 | 0 | 0 | 0 | 0 | 0 | 0 | 0 | 0 | 0 | 0 | 0.000955 | 0.0019352 | 0.001118 | 0.000835 | 0.000104 | 0.001197 | 0 | 0 | 0 | 0 | 0 |
| Eubacterium:K02372 | 0 | 0 | 0 | 0 | 0 | 0 | 0 | 0 | 0 | 0.0042816 | 0 | 0.001366 | 0.002235 | 0.011692 | 0.000623 | 0.002285 | 0.000987 | 0 | 0 | 0 | 0 |
| Eubacterium:K02400 | 0 | 0 | 0 | 0 | 0 | 0 | 0 | 0 | 0 | 0.0203378 | 0.009545 | 0.0044396 | 0.006706 | 0.007517 | 0.00301 | 0.008377 | 0 | 0 | 0 | 0 | 0 |
| Eubacterium:K02405 | 0 | 0 | 0 | 0 | 0 | 0 | 0 | 0 | 0 | 0.0032112 | 0 | 0.0017076 | 0 | 0.002923 | 0.000104 | 0.00185 | 0 | 0 | 0 | 0 | 0 |
| Eubacterium:K02409 | 0 | 0 | 0 | 0 | 0 | 0 | 0 | 0 | 0 | 0.0064225 | 0.000477 | 0.0028459 | 0 | 0.007517 | 0.001246 | 0.003264 | 0.000987 | 0 | 0 | 0 | 0 |

| | | | | | | | | | | | | | | | | | | | | | |
|---|---|---|---|---|---|---|---|---|---|---|---|---|---|---|---|---|---|---|---|---|---|
| Eubacterium:K02410 | 0 | 0 | 0 | 0 | 0 | 0 | 0 | 0 | 0 | 0.0042816 | 0.007159 | 0.0020491 | 0.00447 | 0.00167 | 0.001246 | 0.000544 | 0.000987 | 0 | 0 | 0 | 0 |
| Eubacterium:K02412 | 0 | 0 | 0 | 0 | 0 | 0 | 0 | 0 | 0 | 0.0096337 | 0.003818 | 0.0030736 | 0.007823 | 0.004593 | 0.002699 | 0.004025 | 0 | 0 | 0 | 0.021473051 | 0.007893906 |
| Eubacterium:K02416 | 0 | 0 | 0 | 0 | 0 | 0 | 0 | 0 | 0 | 0.0021408 | 0.001909 | 0.0010245 | 0 | 0.001253 | 0.002284 | 0.003481 | 0.000987 | 0 | 0 | 0.021473051 | 0 |
| Eubacterium:K02492 | 0 | 0 | 0 | 0 | 0 | 0 | 0 | 0 | 0 | 0.0010704 | 0.000477 | 0.0028459 | 0.001118 | 0.000418 | 0.000208 | 0.002067 | 0 | 0 | 0 | 0 | 0 |
| Eubacterium:K02556 | 0 | 0 | 0 | 0 | 0 | 0 | 0 | 0 | 0 | 0 | 0 | 0.0011384 | 0 | 0.000835 | 0.000208 | 0.000762 | 0 | 0 | 0 | 0 | 0 |
| Eubacterium:K02557 | 0 | 0 | 0 | 0 | 0 | 0 | 0 | 0 | 0 | 0.0064225 | 0 | 0.0019352 | 0 | 0.005846 | 0.000623 | 0.000653 | 0 | 0 | 0 | 0.021473051 | 0 |
| Eubacterium:K03092 | 0 | 0 | 0 | 0 | 0 | 0 | 0 | 0 | 0 | 0 | 0 | 0.0002277 | 0 | 0 | 0.000208 | 0.000653 | 0.000987 | 0 | 0 | 0 | 0 |
| Eubacterium:K03182 | 0 | 0 | 0 | 0 | 0 | 0 | 0 | 0 | 0 | 0.0021408 | 0 | 0.0004553 | 0 | 0.000418 | 0.000104 | 0.000218 | 0 | 0 | 0 | 0 | 0 |
| Eubacterium:K03394 | 0 | 0 | 0 | 0 | 0 | 0 | 0 | 0 | 0 | 0 | 0.000477 | 0.000683 | 0 | 0.000835 | 0.000519 | 0.001197 | 0 | 0 | 0 | 0 | 0 |
| Eubacterium:K03407 | 0 | 0 | 0 | 0 | 0 | 0 | 0 | 0 | 0 | 0.0149858 | 0.009068 | 0.0061472 | 0.015646 | 0.008352 | 0.002491 | 0.003808 | 0 | 0 | 0 | 0.021473051 | 0 |
| Eubacterium:K03408 | 0 | 0 | 0 | 0 | 0 | 0 | 0 | 0 | 0 | 0.0021408 | 0 | 0.0010245 | 0.001118 | 0 | 0.000208 | 0.000653 | 0 | 0 | 0 | 0 | 0 |
| Eubacterium:K03412 | 0 | 0 | 0 | 0 | 0 | 0 | 0 | 0 | 0 | 0.0010704 | 0 | 0.0012522 | 0 | 0.000418 | 0.001349 | 0.006419 | 0 | 0 | 0 | 0 | 0 |
| Eubacterium:K03415 | 0 | 0 | 0 | 0 | 0 | 4.15E-05 | 0 | 0 | 0 | 0.0053521 | 0.000477 | 0.0011384 | 0 | 0.002088 | 0.00083 | 0.002067 | 0 | 0 | 0 | 0.021473051 | 0 |
| Eubacterium:K05343 | 0 | 0 | 0 | 0 | 0 | 0 | 0 | 0 | 0 | 0.0032112 | 0.000955 | 0.0017076 | 0.006706 | 0.005429 | 0.000104 | 0.000979 | 0.000987 | 0 | 0 | 0 | 0 |
| Eubacterium:K05592 | 0 | 0 | 0 | 0 | 0 | 0 | 0 | 0 | 0 | 0.0053521 | 0.001432 | 0.0071717 | 0.003353 | 0.005429 | 0.005813 | 0.011641 | 0.004936 | 0 | 0 | 0 | 0 |
| Eubacterium:K05847 | 0 | 0 | 0 | 0 | 0 | 0 | 0 | 0 | 0 | 0 | 0 | 0.000683 | 0 | 0 | 0.000311 | 0.000653 | 0 | 0 | 0 | 0 | 0 |
| Eubacterium:K06148 | 0 | 0 | 0 | 0 | 0 | 4.15E-05 | 0 | 0 | 0 | 0.0021408 | 0.003818 | 0.0035289 | 0.011176 | 0.015868 | 0.01692 | 0.008268 | 0.042449 | 0 | 0 | 0.021473051 | 0 |
| Eubacterium:K10536 | 0 | 0 | 0 | 0 | 0 | 0 | 0 | 0 | 0 | 0.0032112 | 0.000477 | 0.0029598 | 0.005588 | 0.001253 | 0.000104 | 0.004352 | 0 | 0 | 0 | 0 | 0 |
| Eubacterium:K12573 | 0 | 0 | 0 | 0 | 0 | 4.15E-05 | 0 | 0 | 0 | 0.0107041 | 0.002386 | 0.0063749 | 0.002235 | 0.00167 | 0.003529 | 0.009574 | 0.001974 | 0 | 0 | 0 | 0 |
| Flavobacterium:K00029 | 0 | 0 | 0 | 0.0032 | 0 | 0 | 0 | 0 | 0 | 0 | 0 | 0 | 0 | 0.000418 | 0 | 0 | 0 | 0 | 0 | 0 | 0 |
| Flavobacterium:K00104 | 0 | 0.0205 | 0 | 0.0081 | 0 | 0 | 0 | 0 | 0 | 0 | 0 | 0 | 0 | 0 | 0 | 0 | 0 | 0 | 0 | 0 | 0 |
| Flavobacterium:K00111 | 0 | 0 | 0 | 0.0016 | 0 | 0 | 0 | 0 | 0 | 0 | 0 | 0 | 0 | 0 | 0 | 0 | 0 | 0 | 0 | 0 | 0 |
| Flavobacterium:K00116 | 0 | 0 | 0 | 0.0016 | 0 | 0 | 0 | 0 | 0 | 0 | 0 | 0 | 0 | 0 | 0 | 0 | 0 | 0 | 0 | 0 | 0 |
| Flavobacterium:K00128 | 0 | 0.0409 | 0.0713 | 0.0048 | 0 | 0 | 0 | 0 | 0 | 0 | 0 | 0 | 0 | 0 | 0 | 0 | 0 | 0 | 0 | 0 | 0 |
| Flavobacterium:K00131 | 0 | 0 | 0 | 0 | 0 | 0 | 0 | 0 | 0.001025 | 0 | 0 | 0 | 0 | 0 | 0 | 0 | 0 | 0 | 0 | 0 | 0 |
| Flavobacterium:K00161 | 0 | 0.0409 | 0 | 0.0226 | 0 | 0 | 0 | 0 | 0 | 0 | 0 | 0 | 0 | 0 | 0 | 0 | 0 | 0 | 0 | 0 | 0 |
| Flavobacterium:K00162 | 0 | 0.0409 | 0 | 0.0129 | 0 | 0 | 0 | 0 | 0.001025 | 0 | 0 | 0 | 0 | 0 | 0 | 0 | 0 | 0 | 0 | 0 | 0 |
| Flavobacterium:K00164 | 0 | 0.0409 | 0 | 0.0436 | 0 | 0 | 0 | 0 | 0 | 0 | 0 | 0 | 0 | 0 | 0 | 0 | 0 | 0 | 0 | 0 | 0 |
| Flavobacterium:K00228 | 0 | 0.0409 | 0.0356 | 0.0113 | 0 | 0 | 0 | 0 | 0 | 0 | 0 | 0 | 0 | 0 | 0 | 0 | 0 | 0 | 0 | 0 | 0 |
| Flavobacterium:K00252 | 0 | 0.2456 | 0.1782 | 0.0856 | 0 | 0 | 0 | 0 | 0.001025 | 0 | 0 | 0 | 0 | 0 | 0 | 0 | 0 | 0 | 0 | 0 | 0 |
| Flavobacterium:K00257 | 0 | 0.0205 | 0 | 0.0162 | 0 | 0 | 0 | 0 | 0 | 0 | 0 | 0 | 0 | 0 | 0 | 0 | 0 | 0 | 0 | 0 | 0 |
| Flavobacterium:K00281 | 0 | 0.1228 | 0.0356 | 0.0275 | 0 | 0 | 0 | 0 | 0 | 0 | 0.000477 | 0 | 0 | 0 | 0 | 0 | 0 | 0 | 0 | 0 | 0 |
| Flavobacterium:K00331 | 0 | 0.0819 | 0.0356 | 0.0129 | 0 | 0 | 0 | 0 | 0 | 0 | 0 | 0 | 0 | 0 | 0 | 0 | 0 | 0 | 0 | 0 | 0 |
| Flavobacterium:K00334 | 0 | 0.0205 | 0.0356 | 0.0129 | 0 | 0 | 0 | 0 | 0 | 0 | 0 | 0 | 0 | 0 | 0 | 0 | 0 | 0 | 0 | 0 | 0 |
| Flavobacterium:K00335 | 0 | 0.1433 | 0.0713 | 0.0372 | 0 | 0 | 0 | 0 | 0 | 0 | 0 | 0 | 0 | 0 | 0 | 0 | 0 | 0 | 0 | 0 | 0 |
| Flavobacterium:K00336 | 0 | 0.0614 | 0.0356 | 0.0226 | 0 | 0 | 0.002207 | 0.0007694 | 0.001025 | 0 | 0 | 0 | 0 | 0 | 0 | 0 | 0 | 0 | 0 | 0 | 0 |
| Flavobacterium:K00337 | 0 | 0.1023 | 0.0356 | 0.0178 | 0 | 0 | 0 | 0 | 0.001025 | 0 | 0 | 0 | 0 | 0 | 0 | 0 | 0 | 0 | 0 | 0 | 0 |
| Flavobacterium:K00338 | 0 | 0.0614 | 0.0356 | 0.0032 | 0 | 0 | 0 | 0 | 0 | 0 | 0 | 0 | 0 | 0 | 0 | 0 | 0 | 0 | 0 | 0 | 0 |
| Flavobacterium:K00341 | 0 | 0 | 0 | 0.0065 | 0 | 0 | 0 | 0 | 0.001025 | 0 | 0 | 0 | 0 | 0 | 0 | 0 | 0 | 0 | 0 | 0 | 0 |
| Flavobacterium:K00372 | 0 | 0.0205 | 0 | 0 | 0 | 0 | 0 | 0 | 0.001025 | 0 | 0 | 0 | 0 | 0 | 0 | 0 | 0 | 0 | 0 | 0 | 0 |
| Flavobacterium:K00406 | 0 | 0 | 0 | 0.0032 | 0 | 0 | 0 | 0 | 0 | 0 | 0 | 0 | 0 | 0 | 0 | 0 | 0 | 0 | 0 | 0 | 0 |
| Flavobacterium:K00425 | 0 | 0 | 0 | 0 | 0 | 0 | 0 | 0 | 0.001025 | 0 | 0 | 0 | 0 | 0 | 0 | 0 | 0 | 0 | 0 | 0 | 0 |
| Flavobacterium:K00426 | 0 | 0 | 0 | 0 | 0 | 0 | 0 | 0.0007694 | 0 | 0 | 0 | 0 | 0 | 0 | 0 | 0 | 0 | 0 | 0 | 0 | 0 |
| Flavobacterium:K00432 | 0 | 0 | 0 | 0 | 0 | 0 | 0 | 0.0007694 | 0 | 0 | 0 | 0 | 0 | 0 | 0 | 0 | 0 | 0 | 0 | 0 | 0 |
| Flavobacterium:K00451 | 0 | 0.0205 | 0.0356 | 0 | 0 | 0 | 0 | 0 | 0 | 0 | 0 | 0 | 0 | 0 | 0 | 0 | 0 | 0 | 0 | 0 | 0 |
| Flavobacterium:K00457 | 0 | 0.1228 | 0.1069 | 0.0517 | 0 | 0 | 0.002207 | 0.0007694 | 0 | 0 | 0 | 0 | 0 | 0 | 0 | 0 | 0 | 0 | 0 | 0 | 0 |
| Flavobacterium:K00525 | 0 | 0.0205 | 0.1426 | 0.0113 | 0 | 4.15E-05 | 0 | 0 | 0.001025 | 0 | 0 | 0 | 0 | 0 | 0 | 0 | 0 | 0 | 0 | 0 | 0 |
| Flavobacterium:K00526 | 0 | 0.0409 | 0.0713 | 0.0048 | 0 | 4.15E-05 | 0 | 0.0007694 | 0 | 0 | 0 | 0 | 0 | 0 | 0 | 0 | 0 | 0 | 0 | 0 | 0 |
| Flavobacterium:K00548 | 0 | 0 | 0 | 0.0097 | 0 | 0 | 0 | 0.0007694 | 0 | 0 | 0 | 0 | 0 | 0 | 0 | 0 | 0 | 0 | 0 | 0 | 0 |
| Flavobacterium:K00600 | 0 | 0 | 0.0713 | 0.0178 | 0 | 4.15E-05 | 0.002207 | 0 | 0.001025 | 0 | 0 | 0.0001138 | 0 | 0 | 0 | 0 | 0 | 0 | 0 | 0 | 0 |
| Flavobacterium:K00626 | 0 | 0 | 0 | 0.0113 | 0 | 0 | 0 | 0 | 0 | 0 | 0 | 0 | 0 | 0 | 0 | 0 | 0 | 0 | 0 | 0 | 0 |
| Flavobacterium:K00627 | 0 | 0 | 0.0713 | 0.0081 | 0 | 0 | 0 | 0 | 0 | 0 | 0 | 0 | 0 | 0 | 0 | 0 | 0 | 0 | 0 | 0 | 0 |
| Flavobacterium:K00632 | 0 | 0.0614 | 0.2851 | 0.042 | 0 | 0 | 0 | 0 | 0.003076 | 0 | 0 | 0 | 0 | 0 | 0 | 0 | 0 | 0 | 0 | 0 | 0 |
| Flavobacterium:K00647 | 0 | 0.0205 | 0 | 0.0097 | 0 | 0 | 0 | 0.0007694 | 0 | 0 | 0 | 0.0001138 | 0.002235 | 0 | 0 | 0 | 0 | 0 | 0 | 0 | 0 |
| Flavobacterium:K00648 | 0 | 0.0205 | 0 | 0.0048 | 0 | 0 | 0 | 0.0007694 | 0 | 0 | 0 | 0 | 0 | 0 | 0 | 0 | 0 | 0 | 0 | 0 | 0 |
| Flavobacterium:K00658 | 0 | 0.2251 | 0.0713 | 0.0711 | 0 | 0 | 0 | 0 | 0 | 0 | 0 | 0 | 0 | 0 | 0 | 0 | 0 | 0 | 0 | 0 | 0 |
| Flavobacterium:K00677 | 0 | 0 | 0.0713 | 0.0162 | 0 | 0 | 0 | 0 | 0 | 0 | 0 | 0 | 0 | 0 | 0 | 0 | 0 | 0 | 0 | 0 | 0 |
| Flavobacterium:K00681 | 0 | 0 | 0 | 0 | 0 | 0 | 0 | 0 | 0.001025 | 0 | 0 | 0 | 0 | 0 | 0 | 0 | 0 | 0 | 0 | 0 | 0 |
| Flavobacterium:K00748 | 0 | 0.0205 | 0 | 0.0016 | 0 | 0 | 0 | 0 | 0 | 0 | 0 | 0 | 0 | 0 | 0 | 0 | 0 | 0 | 0 | 0 | 0 |
| Flavobacterium:K00819 | 0 | 0.0205 | 0 | 0.0097 | 0 | 0 | 0 | 0.0007694 | 0 | 0 | 0 | 0 | 0 | 0 | 0 | 0 | 0 | 0 | 0 | 0 | 0 |

| | | | | | | | | | | | | | | | | | | | | | |
|---|---|---|---|---|---|---|---|---|---|---|---|---|---|---|---|---|---|---|---|---|---|
| Flavobacterium:K00833 | 0 | 0 | 0 | 0 | 0 | 0 | 0.002207 | 0 | 0 | 0 | 0 | 0 | 0 | 0 | 0 | 0 | 0 | 0 | 0 | 0 | 0 |
| Flavobacterium:K00940 | 0 | 0.0409 | 0.0356 | 0.0048 | 0 | 0 | 0 | 0 | 0 | 0 | 0 | 0 | 0 | 0 | 0 | 0 | 0 | 0 | 0 | 0 | 0 |
| Flavobacterium:K01029 | 0 | 0.0205 | 0 | 0.0032 | 0 | 0 | 0 | 0 | 0 | 0 | 0 | 0 | 0 | 0 | 0 | 0 | 0 | 0 | 0 | 0 | 0 |
| Flavobacterium:K01142 | 0 | 0.0205 | 0.0356 | 0.021 | 0 | 0 | 0.004413 | 0.0007694 | 0.001025 | 0 | 0 | 0 | 0 | 0 | 0 | 0 | 0 | 0 | 0 | 0 | 0 |
| Flavobacterium:K01251 | 0 | 0.2456 | 0.1782 | 0.0614 | 0 | 0 | 0 | 0 | 0.001025 | 0 | 0 | 0 | 0 | 0 | 0 | 0 | 0 | 0 | 0 | 0 | 0 |
| Flavobacterium:K01338 | 0 | 0 | 0 | 0 | 0 | 0 | 0 | 0.0007694 | 0.001025 | 0 | 0 | 0.0004553 | 0 | 0 | 0 | 0 | 0 | 0 | 0 | 0 | 0 |
| Flavobacterium:K01426 | 0 | 0 | 0 | 0 | 0 | 0 | 0 | 0.0007694 | 0 | 0 | 0 | 0 | 0 | 0 | 0 | 0 | 0 | 0 | 0 | 0 | 0 |
| Flavobacterium:K01428 | 0 | 0 | 0 | 0.0016 | 0 | 0 | 0 | 0 | 0 | 0 | 0 | 0 | 0 | 0 | 0 | 0 | 0 | 0 | 0 | 0 | 0 |
| Flavobacterium:K01433 | 0 | 0 | 0 | 0 | 0 | 0 | 0 | 0.0007694 | 0 | 0 | 0 | 0 | 0 | 0 | 0 | 0 | 0 | 0 | 0 | 0 | 0 |
| Flavobacterium:K01585 | 0 | 0.1433 | 0.2138 | 0.0759 | 0 | 0 | 0 | 0 | 0 | 0 | 0 | 0 | 0 | 0 | 0 | 0 | 0 | 0 | 0 | 0 | 0 |
| Flavobacterium:K01627 | 0 | 0.1023 | 0.0713 | 0.0275 | 0 | 0 | 0 | 0 | 0.001025 | 0 | 0 | 0 | 0 | 0 | 0 | 0 | 0 | 0 | 0 | 0 | 0 |
| Flavobacterium:K01637 | 0 | 0 | 0 | 0.0016 | 0 | 4.15E-05 | 0 | 0 | 0 | 0 | 0 | 0 | 0 | 0 | 0 | 0 | 0 | 0 | 0 | 0 | 0 |
| Flavobacterium:K01676 | 0 | 0.0819 | 0 | 0.0226 | 0 | 4.15E-05 | 0 | 0 | 0.001025 | 0 | 0 | 0 | 0 | 0 | 0 | 0 | 0 | 0 | 0 | 0 | 0 |
| Flavobacterium:K01679 | 0 | 0.0205 | 0.0713 | 0.0275 | 0 | 0 | 0 | 0.0007694 | 0 | 0 | 0 | 0 | 0 | 0 | 0 | 0 | 0 | 0 | 0 | 0 | 0 |
| Flavobacterium:K01698 | 0 | 0.0409 | 0.0356 | 0.0339 | 0 | 0 | 0 | 0 | 0 | 0 | 0 | 0 | 0 | 0 | 0 | 0 | 0 | 0 | 0 | 0 | 0 |
| Flavobacterium:K01712 | 0 | 0.0205 | 0.0713 | 0.0275 | 0 | 4.15E-05 | 0 | 0 | 0 | 0 | 0.000955 | 0.0002277 | 0 | 0 | 0 | 0.000109 | 0 | 0 | 0 | 0 | 0 |
| Flavobacterium:K01737 | 0 | 0 | 0 | 0.0097 | 0 | 0 | 0 | 0 | 0 | 0 | 0 | 0 | 0 | 0 | 0 | 0 | 0 | 0 | 0 | 0 | 0 |
| Flavobacterium:K01745 | 0 | 0.1023 | 0.0356 | 0.0307 | 0 | 0 | 0 | 0 | 0.001025 | 0 | 0.000477 | 0 | 0 | 0 | 0 | 0 | 0 | 0 | 0 | 0 | 0 |
| Flavobacterium:K01749 | 0 | 0.0614 | 0 | 0.0355 | 0 | 4.15E-05 | 0.002207 | 0.0007694 | 0 | 0 | 0 | 0 | 0 | 0 | 0 | 0 | 0 | 0 | 0 | 0 | 0 |
| Flavobacterium:K01772 | 0 | 0.0409 | 0.0356 | 0.0258 | 0 | 0 | 0 | 0 | 0 | 0 | 0 | 0 | 0 | 0 | 0 | 0 | 0 | 0 | 0 | 0 | 0 |
| Flavobacterium:K01845 | 0 | 0 | 0.0356 | 0.0081 | 0 | 0 | 0 | 0 | 0 | 0 | 0 | 0 | 0 | 0 | 0 | 0 | 0 | 0 | 0 | 0 | 0 |
| Flavobacterium:K01895 | 0 | 0.0409 | 0.0713 | 0.0032 | 0 | 0 | 0 | 0.0007694 | 0.002051 | 0 | 0 | 0 | 0 | 0 | 0 | 0 | 0 | 0 | 0 | 0 | 0 |
| Flavobacterium:K01902 | 0 | 0.0819 | 0.0713 | 0.0242 | 0 | 0 | 0 | 0 | 0 | 0 | 0 | 0 | 0 | 0 | 0 | 0 | 0 | 0 | 0 | 0 | 0 |
| Flavobacterium:K01903 | 0 | 0.0819 | 0.0356 | 0.0339 | 0 | 0 | 0 | 0 | 0.002051 | 0 | 0 | 0 | 0 | 0 | 0 | 0 | 0 | 0 | 0 | 0 | 0 |
| Flavobacterium:K01915 | 0 | 0.0409 | 0.1069 | 0.0339 | 0 | 0 | 0 | 0.0007694 | 0 | 0 | 0 | 0 | 0 | 0 | 0 | 0 | 0 | 0 | 0 | 0 | 0 |
| Flavobacterium:K01961 | 0 | 0 | 0 | 0.0065 | 0 | 4.15E-05 | 0 | 0 | 0 | 0 | 0 | 0 | 0 | 0 | 0 | 0 | 0 | 0 | 0 | 0 | 0 |
| Flavobacterium:K01962 | 0 | 0 | 0 | 0.0016 | 0 | 0 | 0 | 0 | 0 | 0 | 0 | 0 | 0 | 0 | 0 | 0 | 0 | 0 | 0 | 0 | 0 |
| Flavobacterium:K02274 | 0 | 0.2047 | 0.1426 | 0.0662 | 0 | 0 | 0 | 0 | 0.001025 | 0 | 0 | 0 | 0 | 0 | 0 | 0 | 0 | 0 | 0 | 0 | 0 |
| Flavobacterium:K02275 | 0 | 0 | 0 | 0.0065 | 0 | 0 | 0 | 0 | 0 | 0 | 0 | 0 | 0 | 0 | 0 | 0 | 0 | 0 | 0 | 0 | 0 |
| Flavobacterium:K02276 | 0 | 0.0205 | 0.0356 | 0 | 0 | 0 | 0.002207 | 0 | 0 | 0 | 0 | 0 | 0 | 0 | 0 | 0 | 0 | 0 | 0 | 0 | 0 |
| Flavobacterium:K02301 | 0 | 0 | 0 | 0.0048 | 0 | 0 | 0 | 0 | 0.001025 | 0 | 0 | 0 | 0 | 0 | 0 | 0 | 0 | 0 | 0 | 0 | 0 |
| Flavobacterium:K02372 | 0 | 0 | 0 | 0 | 0 | 0 | 0 | 0 | 0.001025 | 0 | 0 | 0 | 0 | 0 | 0 | 0 | 0 | 0 | 0 | 0 | 0 |
| Flavobacterium:K02492 | 0 | 0.0205 | 0.0356 | 0 | 0 | 4.15E-05 | 0 | 0 | 0 | 0 | 0 | 0 | 0 | 0 | 0 | 0 | 0 | 0 | 0 | 0 | 0 |
| Flavobacterium:K02523 | 0 | 0 | 0 | 0 | 0 | 0 | 0 | 0 | 0.001025 | 0 | 0 | 0 | 0 | 0 | 0 | 0 | 0 | 0 | 0 | 0 | 0 |
| Flavobacterium:K02557 | 0 | 0 | 0 | 0 | 0 | 0 | 0 | 0.0007694 | 0 | 0 | 0 | 0 | 0 | 0 | 0 | 0 | 0 | 0 | 0 | 0 | 0 |
| Flavobacterium:K03092 | 0 | 0.0614 | 0 | 0.0145 | 0 | 0 | 0 | 0.0007694 | 0.001025 | 0 | 0 | 0.0001138 | 0 | 0 | 0 | 0 | 0 | 0 | 0 | 0 | 0 |
| Flavobacterium:K03183 | 0 | 0 | 0 | 0.0016 | 0 | 0 | 0 | 0 | 0 | 0 | 0 | 0 | 0 | 0 | 0 | 0 | 0 | 0 | 0 | 0 | 0 |
| Flavobacterium:K03590 | 0 | 0.0409 | 0 | 0.0178 | 0 | 0 | 0 | 0 | 0 | 0 | 0 | 0 | 0 | 0 | 0 | 0 | 0 | 0 | 0 | 0 | 0 |
| Flavobacterium:K03644 | 0 | 0 | 0.0356 | 0.0032 | 0 | 0 | 0 | 0 | 0.002051 | 0 | 0 | 0 | 0 | 0 | 0 | 0 | 0 | 0 | 0 | 0 | 0 |
| Flavobacterium:K03781 | 0 | 0 | 0 | 0.0065 | 0 | 0 | 0 | 0 | 0 | 0 | 0 | 0 | 0 | 0 | 0 | 0 | 0 | 0 | 0 | 0 | 0 |
| Flavobacterium:K03841 | 0 | 0.0614 | 0.0356 | 0.0097 | 0 | 0 | 0.002207 | 0 | 0 | 0 | 0 | 0 | 0 | 0 | 0 | 0 | 0 | 0 | 0 | 0 | 0 |
| Flavobacterium:K03885 | 0 | 0.0819 | 0 | 0.0113 | 0 | 0 | 0 | 0 | 0 | 0 | 0 | 0 | 0 | 0 | 0 | 0 | 0 | 0 | 0 | 0 | 0 |
| Flavobacterium:K04561 | 0 | 0.0205 | 0.0356 | 0.021 | 0 | 0 | 0 | 0.0015388 | 0 | 0 | 0 | 0 | 0 | 0 | 0 | 0 | 0 | 0 | 0 | 0 | 0 |
| Flavobacterium:K04564 | 0 | 0 | 0 | 0.0016 | 0 | 0 | 0 | 0 | 0 | 0 | 0 | 0 | 0 | 0 | 0 | 0 | 0 | 0 | 0 | 0 | 0 |
| Flavobacterium:K05592 | 0 | 0.1637 | 0.0356 | 0.0242 | 0 | 0 | 0 | 0.0007694 | 0 | 0 | 0 | 0 | 0 | 0 | 0 | 0 | 0 | 0 | 0 | 0 | 0 |
| Flavobacterium:K06861 | 0 | 0.1023 | 0.0356 | 0.0226 | 0 | 0 | 0 | 0.0007694 | 0 | 0 | 0 | 0 | 0 | 0 | 0 | 0 | 0 | 0 | 0 | 0 | 0 |
| Flavobacterium:K07516 | 0 | 0.1433 | 0.0713 | 0.0711 | 0 | 0 | 0 | 0 | 0 | 0 | 0 | 0 | 0 | 0 | 0 | 0 | 0 | 0 | 0 | 0 | 0 |
| Flavobacterium:K10536 | 0 | 0 | 0 | 0 | 0 | 0 | 0.002207 | 0.0007694 | 0 | 0 | 0 | 0 | 0 | 0 | 0 | 0 | 0 | 0 | 0 | 0 | 0 |
| Flavobacterium:K11927 | 0 | 0 | 0 | 0.0048 | 0 | 0 | 0 | 0 | 0 | 0 | 0 | 0 | 0 | 0 | 0 | 0 | 0 | 0 | 0 | 0 | 0 |
| Flavobacterium:K12573 | 0 | 0.0819 | 0.0713 | 0.0355 | 0 | 0 | 0.002207 | 0 | 0 | 0 | 0 | 0 | 0 | 0 | 0 | 0 | 0 | 0 | 0 | 0 | 0 |
| Klebsiella:K00029 | 0 | 0 | 0 | 0 | 0 | 0 | 0 | 0 | 0 | 0.0042816 | 0 | 0 | 0 | 0 | 0 | 0 | 0 | 0 | 0 | 0 | 0 |
| Klebsiella:K00036 | 0 | 0 | 0 | 0 | 0 | 0 | 0 | 0 | 0 | 0.0021408 | 0 | 0 | 0 | 0 | 0 | 0 | 0 | 0 | 0 | 0 | 0 |
| Klebsiella:K00108 | 0 | 0 | 0 | 0 | 0 | 0 | 0 | 0 | 0.001025 | 0.0021408 | 0 | 0 | 0 | 0 | 0 | 0 | 0 | 0 | 0 | 0 | 0 |
| Klebsiella:K00111 | 0 | 0 | 0 | 0 | 0 | 0 | 0 | 0 | 0 | 0.0064225 | 0 | 0.0001138 | 0 | 0 | 0 | 0 | 0 | 0 | 0 | 0 | 0 |
| Klebsiella:K00116 | 0 | 0 | 0 | 0 | 0 | 0 | 0 | 0 | 0 | 0.0085633 | 0 | 0.0001138 | 0.001118 | 0 | 0 | 0 | 0 | 0 | 0 | 0 | 0 |
| Klebsiella:K00117 | 0 | 0 | 0 | 0 | 0 | 0 | 0 | 0 | 0 | 0.0042816 | 0.000477 | 0.0001138 | 0 | 0 | 0 | 0 | 0 | 0 | 0 | 0 | 0 |
| Klebsiella:K00123 | 0 | 0 | 0 | 0 | 0 | 0 | 0 | 0 | 0 | 0.0278307 | 0 | 0.0002277 | 0 | 0 | 0 | 0.000109 | 0.000987 | 0 | 0 | 0 | 0 |
| Klebsiella:K00130 | 0 | 0 | 0 | 0.0016 | 0 | 0 | 0 | 0 | 0 | 0.0042816 | 0 | 0.0001138 | 0 | 0 | 0 | 0 | 0 | 0 | 0 | 0 | 0 |
| Klebsiella:K00135 | 0 | 0 | 0 | 0.0032 | 0 | 0 | 0 | 0 | 0 | 0.0085633 | 0.000477 | 0.0002277 | 0 | 0.000418 | 0 | 0 | 0 | 0 | 0 | 0 | 0 |
| Klebsiella:K00140 | 0 | 0 | 0 | 0.0016 | 0 | 0 | 0 | 0 | 0 | 0.0053521 | 0 | 0.0001138 | 0 | 0 | 0 | 0 | 0 | 0 | 0 | 0 | 0 |

| | | | | | | | | | | | | | | | | | | | | | |
|---|---|---|---|---|---|---|---|---|---|---|---|---|---|---|---|---|---|---|---|---|---|
| Klebsiella:K00161 | 0 | 0 | 0 | 0 | 0 | 0 | 0 | 0 | 0 | 0.0074929 | 0 | 0 | 0 | 0 | 0 | 0 | 0 | 0 | 0 | 0 | 0 |
| Klebsiella:K00162 | 0 | 0 | 0 | 0 | 0 | 0 | 0 | 0 | 0 | 0.0010704 | 0 | 0 | 0 | 0 | 0 | 0 | 0 | 0 | 0 | 0 | 0 |
| Klebsiella:K00163 | 0 | 0 | 0 | 0 | 0 | 0 | 0 | 0 | 0 | 0.0032112 | 0 | 0.0001138 | 0 | 0 | 0 | 0 | 0 | 0 | 0 | 0 | 0 |
| Klebsiella:K00164 | 0 | 0 | 0 | 0 | 0 | 0 | 0 | 0 | 0 | 0.0021408 | 0 | 0 | 0 | 0 | 0 | 0.000109 | 0 | 0 | 0 | 0 | 0 |
| Klebsiella:K00281 | 0 | 0 | 0 | 0 | 0 | 0 | 0 | 0.0007694 | 0 | 0.0032112 | 0 | 0.0003415 | 0 | 0 | 0 | 0 | 0 | 0 | 0 | 0 | 0 |
| Klebsiella:K00285 | 0 | 0 | 0 | 0 | 0 | 0 | 0 | 0 | 0 | 0.0032112 | 0 | 0.0001138 | 0 | 0 | 0 | 0 | 0 | 0 | 0 | 0 | 0 |
| Klebsiella:K00322 | 0 | 0 | 0 | 0 | 0 | 0 | 0 | 0 | 0 | 0.0032112 | 0 | 0.0005692 | 0 | 0 | 0 | 0 | 0 | 0 | 0 | 0 | 0 |
| Klebsiella:K00331 | 0 | 0.0205 | 0 | 0 | 0 | 0 | 0 | 0 | 0 | 0 | 0 | 0 | 0 | 0 | 0 | 0 | 0 | 0 | 0 | 0 | 0 |
| Klebsiella:K00335 | 0 | 0 | 0 | 0 | 0 | 0 | 0 | 0 | 0 | 0.0064225 | 0 | 0 | 0 | 0 | 0 | 0 | 0 | 0 | 0 | 0 | 0 |
| Klebsiella:K00336 | 0 | 0 | 0 | 0.0016 | 0 | 0 | 0 | 0 | 0 | 0.0032112 | 0 | 0.0001138 | 0 | 0 | 0 | 0 | 0.000987 | 0 | 0 | 0 | 0 |
| Klebsiella:K00338 | 0 | 0 | 0 | 0 | 0 | 0 | 0 | 0.0007694 | 0 | 0 | 0 | 0 | 0 | 0 | 0 | 0 | 0 | 0 | 0 | 0 | 0 |
| Klebsiella:K00341 | 0 | 0 | 0 | 0 | 0 | 0 | 0 | 0 | 0 | 0.0064225 | 0 | 0 | 0 | 0 | 0 | 0 | 0 | 0 | 0 | 0 | 0 |
| Klebsiella:K00342 | 0 | 0 | 0 | 0 | 0 | 0 | 0 | 0 | 0 | 0.0032112 | 0 | 0 | 0 | 0 | 0 | 0 | 0 | 0 | 0 | 0 | 0 |
| Klebsiella:K00362 | 0 | 0 | 0 | 0 | 0 | 0 | 0 | 0 | 0 | 0.0149858 | 0.000477 | 0 | 0 | 0 | 0 | 0 | 0 | 0 | 0 | 0 | 0 |
| Klebsiella:K00370 | 0 | 0 | 0 | 0 | 0 | 0 | 0 | 0 | 0.001025 | 0.0160562 | 0 | 0 | 0 | 0 | 0 | 0 | 0 | 0 | 0 | 0 | 0 |
| Klebsiella:K00371 | 0 | 0 | 0 | 0 | 0 | 0 | 0.002207 | 0 | 0 | 0.0042816 | 0 | 0 | 0 | 0 | 0 | 0 | 0 | 0 | 0 | 0 | 0 |
| Klebsiella:K00372 | 0 | 0 | 0 | 0 | 0 | 0 | 0 | 0 | 0 | 0.0032112 | 0 | 0 | 0 | 0 | 0 | 0 | 0 | 0 | 0 | 0 | 0 |
| Klebsiella:K00381 | 0 | 0 | 0 | 0 | 0 | 0 | 0 | 0 | 0 | 0.0021408 | 0 | 0 | 0 | 0 | 0 | 0 | 0 | 0 | 0 | 0 | 0 |
| Klebsiella:K00383 | 0 | 0 | 0 | 0 | 0 | 0 | 0 | 0 | 0 | 0.0021408 | 0 | 0 | 0 | 0 | 0 | 0 | 0 | 0 | 0 | 0 | 0 |
| Klebsiella:K00390 | 0 | 0 | 0 | 0 | 0 | 0 | 0 | 0 | 0 | 0.0010704 | 0 | 0 | 0 | 0 | 0 | 0 | 0 | 0 | 0 | 0 | 0 |
| Klebsiella:K00425 | 0 | 0 | 0 | 0.0016 | 0 | 0 | 0 | 0 | 0 | 0.0160562 | 0.000477 | 0.0001138 | 0 | 0 | 0 | 0 | 0 | 0 | 0 | 0 | 0 |
| Klebsiella:K00426 | 0 | 0 | 0 | 0 | 0 | 0 | 0 | 0 | 0 | 0.0032112 | 0 | 0.0001138 | 0 | 0 | 0 | 0 | 0 | 0 | 0 | 0 | 0 |
| Klebsiella:K00432 | 0 | 0 | 0 | 0 | 0 | 0 | 0 | 0 | 0 | 0.0042816 | 0 | 0 | 0 | 0 | 0 | 0 | 0 | 0 | 0 | 0 | 0 |
| Klebsiella:K00457 | 0 | 0 | 0 | 0 | 0 | 0 | 0 | 0 | 0 | 0.0053521 | 0 | 0.0001138 | 0 | 0 | 0 | 0 | 0 | 0 | 0 | 0 | 0 |
| Klebsiella:K00525 | 0 | 0 | 0 | 0 | 0 | 0 | 0 | 0 | 0 | 0.0160562 | 0 | 0.0003415 | 0 | 0 | 0 | 0 | 0 | 0 | 0 | 0 | 0 |
| Klebsiella:K00526 | 0 | 0 | 0 | 0 | 0 | 0 | 0 | 0 | 0 | 0.0010704 | 0 | 0.0001138 | 0 | 0 | 0 | 0 | 0 | 0 | 0 | 0 | 0 |
| Klebsiella:K00548 | 0 | 0 | 0 | 0 | 0 | 0 | 0 | 0 | 0 | 0.0160562 | 0 | 0.0001138 | 0.001118 | 0 | 0 | 0 | 0 | 0 | 0 | 0 | 0 |
| Klebsiella:K00568 | 0 | 0 | 0 | 0 | 0 | 0 | 0 | 0 | 0 | 0.0021408 | 0 | 0.0001138 | 0 | 0 | 0 | 0 | 0 | 0 | 0 | 0 | 0 |
| Klebsiella:K00626 | 0 | 0 | 0 | 0 | 0 | 0 | 0 | 0 | 0 | 0.0021408 | 0 | 0.0001138 | 0 | 0 | 0 | 0 | 0 | 0 | 0 | 0 | 0 |
| Klebsiella:K00627 | 0 | 0 | 0 | 0 | 0 | 0 | 0 | 0 | 0 | 0.0053521 | 0 | 0 | 0 | 0 | 0 | 0 | 0 | 0 | 0 | 0 | 0 |
| Klebsiella:K00631 | 0 | 0 | 0 | 0 | 0 | 0 | 0 | 0 | 0 | 0.0053521 | 0 | 0 | 0 | 0 | 0 | 0 | 0 | 0 | 0 | 0 | 0 |
| Klebsiella:K00632 | 0 | 0 | 0 | 0 | 0 | 0 | 0 | 0 | 0 | 0.0107041 | 0 | 0 | 0 | 0 | 0 | 0.000109 | 0 | 0 | 0 | 0 | 0 |
| Klebsiella:K00640 | 0 | 0 | 0 | 0 | 0 | 0 | 0 | 0 | 0 | 0 | 0 | 0 | 0 | 0 | 0 | 0 | 0.000987 | 0 | 0 | 0 | 0 |
| Klebsiella:K00647 | 0 | 0 | 0 | 0 | 0 | 0 | 0 | 0 | 0 | 0.0021408 | 0 | 0 | 0 | 0 | 0 | 0 | 0 | 0 | 0 | 0 | 0 |
| Klebsiella:K00648 | 0 | 0 | 0 | 0 | 0 | 0 | 0 | 0 | 0 | 0.0021408 | 0 | 0 | 0 | 0 | 0 | 0 | 0 | 0 | 0 | 0 | 0 |
| Klebsiella:K00658 | 0 | 0 | 0 | 0 | 0 | 0 | 0 | 0 | 0 | 0.0010704 | 0 | 0 | 0 | 0 | 0 | 0 | 0 | 0 | 0 | 0 | 0 |
| Klebsiella:K00677 | 0 | 0 | 0 | 0 | 0 | 0 | 0 | 0 | 0 | 0.0032112 | 0 | 0 | 0 | 0 | 0 | 0 | 0 | 0 | 0 | 0 | 0 |
| Klebsiella:K00681 | 0 | 0 | 0 | 0 | 0 | 0 | 0 | 0 | 0 | 0.0074929 | 0 | 0 | 0 | 0 | 0 | 0 | 0.000987 | 0 | 0 | 0 | 0 |
| Klebsiella:K00697 | 0 | 0 | 0 | 0 | 0 | 0 | 0 | 0 | 0 | 0.0053521 | 0 | 0.0002277 | 0 | 0 | 0 | 0 | 0 | 0 | 0 | 0 | 0 |
| Klebsiella:K00748 | 0 | 0 | 0 | 0 | 0 | 0 | 0 | 0 | 0 | 0.0010704 | 0 | 0 | 0 | 0 | 0 | 0 | 0 | 0 | 0 | 0 | 0 |
| Klebsiella:K00798 | 0 | 0 | 0 | 0 | 0 | 0 | 0 | 0 | 0 | 0.0010704 | 0 | 0 | 0 | 0 | 0 | 0 | 0 | 0 | 0 | 0 | 0 |
| Klebsiella:K00799 | 0 | 0 | 0 | 0 | 0 | 0 | 0 | 0 | 0 | 0.0128449 | 0 | 0.0001138 | 0.001118 | 0 | 0 | 0 | 0.000987 | 0 | 0 | 0 | 0 |
| Klebsiella:K00833 | 0 | 0 | 0 | 0 | 0 | 0 | 0 | 0 | 0 | 0.0042816 | 0 | 0 | 0 | 0 | 0 | 0 | 0 | 0 | 0 | 0 | 0 |
| Klebsiella:K00836 | 0 | 0 | 0 | 0 | 0 | 0 | 0 | 0 | 0 | 0.0021408 | 0 | 0 | 0 | 0 | 0 | 0 | 0 | 0 | 0 | 0 | 0 |
| Klebsiella:K00864 | 0 | 0 | 0 | 0 | 0 | 0 | 0 | 0 | 0 | 0.0064225 | 0 | 0 | 0 | 0 | 0 | 0.000109 | 0 | 0 | 0 | 0 | 0 |
| Klebsiella:K01011 | 0 | 0 | 0 | 0 | 0 | 0 | 0 | 0 | 0 | 0.0032112 | 0 | 0.0001138 | 0 | 0 | 0 | 0 | 0 | 0 | 0 | 0 | 0 |
| Klebsiella:K01029 | 0 | 0 | 0 | 0 | 0 | 4.15E-05 | 0 | 0 | 0 | 0.0010704 | 0 | 0 | 0 | 0 | 0 | 0 | 0 | 0 | 0 | 0 | 0 |
| Klebsiella:K01070 | 0 | 0.0205 | 0 | 0 | 0 | 0 | 0 | 0 | 0 | 0.0021408 | 0 | 0 | 0 | 0 | 0 | 0 | 0 | 0 | 0 | 0 | 0 |
| Klebsiella:K01139 | 0 | 0 | 0 | 0.0016 | 0 | 0 | 0 | 0 | 0 | 0.0010704 | 0 | 0 | 0 | 0 | 0 | 0 | 0 | 0 | 0 | 0 | 0 |
| Klebsiella:K01141 | 0 | 0 | 0 | 0 | 0 | 0 | 0 | 0 | 0 | 0.0032112 | 0 | 0 | 0 | 0 | 0 | 0 | 0 | 0 | 0 | 0 | 0 |
| Klebsiella:K01142 | 0 | 0 | 0 | 0.0065 | 0 | 0 | 0 | 0 | 0 | 0.0010704 | 0 | 0 | 0 | 0 | 0 | 0 | 0 | 0 | 0 | 0 | 0 |
| Klebsiella:K01241 | 0 | 0 | 0 | 0 | 0 | 0 | 0 | 0 | 0 | 0.0064225 | 0 | 0 | 0 | 0 | 0 | 0 | 0 | 0 | 0 | 0 | 0 |
| Klebsiella:K01255 | 0 | 0 | 0 | 0 | 0 | 0 | 0 | 0 | 0 | 0.0021408 | 0 | 0 | 0 | 0 | 0 | 0 | 0 | 0 | 0 | 0 | 0 |
| Klebsiella:K01338 | 0 | 0 | 0 | 0.0016 | 0 | 0 | 0 | 0 | 0 | 0.0139154 | 0.000477 | 0.0001138 | 0 | 0 | 0 | 0 | 0 | 0 | 0 | 0 | 0 |
| Klebsiella:K01428 | 0 | 0 | 0.0356 | 0.0065 | 0 | 0 | 0 | 0.0007694 | 0 | 0.0064225 | 0 | 0 | 0 | 0.000418 | 0 | 0.000109 | 0 | 0 | 0 | 0 | 0 |
| Klebsiella:K01433 | 0 | 0 | 0 | 0 | 0 | 0 | 0 | 0 | 0 | 0.0010704 | 0 | 0 | 0 | 0 | 0 | 0 | 0 | 0 | 0 | 0 | 0 |
| Klebsiella:K01488 | 0 | 0 | 0 | 0 | 0 | 0 | 0 | 0 | 0 | 0.0021408 | 0 | 0 | 0 | 0 | 0 | 0 | 0 | 0 | 0 | 0 | 0 |
| Klebsiella:K01497 | 0 | 0 | 0 | 0 | 0 | 0 | 0 | 0 | 0 | 0 | 0.000477 | 0.0001138 | 0 | 0 | 0 | 0 | 0 | 0 | 0 | 0 | 0 |
| Klebsiella:K01581 | 0 | 0 | 0 | 0.0016 | 0 | 0 | 0 | 0 | 0 | 0.0085633 | 0 | 0.0001138 | 0 | 0 | 0 | 0 | 0 | 0 | 0 | 0 | 0 |
| Klebsiella:K01585 | 0 | 0 | 0 | 0 | 0 | 0 | 0 | 0 | 0 | 0.0085633 | 0 | 0 | 0 | 0 | 0 | 0 | 0 | 0 | 0 | 0 | 0 |

| | | | | | | | | | | | | | | | | | | | | | |
|---|---|---|---|---|---|---|---|---|---|---|---|---|---|---|---|---|---|---|---|---|---|
| Klebsiella:K01613 | 0 | 0 | 0 | 0 | 0 | 0 | 0 | 0 | 0 | 0.0010704 | 0 | 0.0001138 | 0 | 0 | 0 | 0 | 0 | 0 | 0 | 0 | 0 |
| Klebsiella:K01629 | 0 | 0 | 0 | 0 | 0 | 0 | 0 | 0 | 0 | 0.0021408 | 0 | 0.0002277 | 0 | 0 | 0 | 0.000109 | 0 | 0 | 0 | 0 | 0 |
| Klebsiella:K01637 | 0 | 0 | 0 | 0 | 0 | 0 | 0 | 0 | 0 | 0.0021408 | 0.000477 | 0 | 0 | 0 | 0 | 0 | 0 | 0 | 0 | 0 | 0 |
| Klebsiella:K01676 | 0 | 0 | 0 | 0 | 0 | 0 | 0 | 0 | 0 | 0.0032112 | 0.002864 | 0.0001138 | 0 | 0 | 0 | 0 | 0 | 0 | 0 | 0 | 0 |
| Klebsiella:K01679 | 0 | 0 | 0 | 0 | 0 | 0 | 0 | 0 | 0.001025 | 0.0032112 | 0 | 0 | 0 | 0 | 0 | 0 | 0 | 0 | 0 | 0 | 0 |
| Klebsiella:K01690 | 0 | 0 | 0 | 0 | 0 | 0 | 0 | 0 | 0 | 0.0021408 | 0.000477 | 0.0001138 | 0 | 0 | 0 | 0 | 0 | 0 | 0 | 0 | 0 |
| Klebsiella:K01692 | 0 | 0 | 0 | 0 | 0 | 0 | 0 | 0 | 0 | 0.0021408 | 0 | 0 | 0 | 0 | 0 | 0 | 0 | 0 | 0 | 0 | 0 |
| Klebsiella:K01698 | 0 | 0 | 0 | 0 | 0 | 0 | 0 | 0 | 0 | 0.0021408 | 0 | 0 | 0 | 0 | 0 | 0 | 0 | 0 | 0 | 0 | 0 |
| Klebsiella:K01712 | 0 | 0 | 0 | 0 | 0 | 0 | 0 | 0 | 0 | 0.0010704 | 0 | 0 | 0 | 0 | 0 | 0 | 0 | 0 | 0 | 0 | 0 |
| Klebsiella:K01737 | 0 | 0 | 0 | 0 | 0 | 0 | 0 | 0 | 0 | 0.0021408 | 0 | 0 | 0 | 0 | 0 | 0 | 0 | 0 | 0 | 0 | 0 |
| Klebsiella:K01745 | 0 | 0 | 0 | 0 | 0 | 0 | 0 | 0 | 0 | 0.0074929 | 0 | 0 | 0 | 0 | 0 | 0 | 0 | 0 | 0 | 0 | 0 |
| Klebsiella:K01749 | 0 | 0 | 0 | 0 | 0 | 0 | 0 | 0 | 0 | 0.0042816 | 0 | 0 | 0 | 0 | 0 | 0 | 0 | 0 | 0 | 0 | 0 |
| Klebsiella:K01772 | 0 | 0 | 0 | 0 | 0 | 0 | 0 | 0 | 0 | 0.0032112 | 0 | 0.0001138 | 0 | 0 | 0 | 0 | 0 | 0 | 0 | 0 | 0 |
| Klebsiella:K01825 | 0 | 0 | 0 | 0 | 0 | 0 | 0 | 0 | 0 | 0.0096337 | 0 | 0.0003415 | 0 | 0 | 0 | 0 | 0 | 0 | 0 | 0 | 0 |
| Klebsiella:K01845 | 0 | 0 | 0 | 0 | 0 | 0 | 0 | 0.0007694 | 0 | 0.0042816 | 0 | 0 | 0 | 0 | 0 | 0 | 0 | 0 | 0 | 0 | 0 |
| Klebsiella:K01878 | 0 | 0 | 0 | 0 | 0 | 0 | 0 | 0 | 0 | 0.0010704 | 0 | 0 | 0 | 0 | 0 | 0 | 0 | 0 | 0 | 0 | 0 |
| Klebsiella:K01879 | 0 | 0 | 0 | 0 | 0 | 0 | 0 | 0 | 0 | 0.0096337 | 0.000477 | 0.0002277 | 0 | 0 | 0 | 0 | 0 | 0 | 0 | 0 | 0 |
| Klebsiella:K01895 | 0 | 0 | 0 | 0 | 0 | 0 | 0 | 0 | 0 | 0.0053521 | 0 | 0 | 0 | 0 | 0 | 0 | 0 | 0 | 0 | 0 | 0 |
| Klebsiella:K01915 | 0 | 0 | 0 | 0 | 0 | 0 | 0 | 0 | 0 | 0.0010704 | 0 | 0 | 0 | 0 | 0 | 0 | 0 | 0 | 0 | 0 | 0 |
| Klebsiella:K01920 | 0 | 0 | 0 | 0 | 0 | 0 | 0 | 0 | 0 | 0.0010704 | 0.000477 | 0 | 0 | 0 | 0 | 0 | 0 | 0 | 0 | 0 | 0 |
| Klebsiella:K01941 | 0 | 0 | 0 | 0 | 0 | 0 | 0.002207 | 0.0007694 | 0 | 0 | 0 | 0 | 0 | 0 | 0 | 0 | 0 | 0 | 0 | 0 | 0 |
| Klebsiella:K01961 | 0 | 0 | 0 | 0 | 0 | 0 | 0 | 0 | 0 | 0.0032112 | 0 | 0 | 0 | 0 | 0 | 0 | 0 | 0 | 0 | 0 | 0 |
| Klebsiella:K01962 | 0 | 0 | 0 | 0 | 0 | 0 | 0 | 0 | 0 | 0.0010704 | 0 | 0 | 0 | 0 | 0 | 0 | 0 | 0 | 0 | 0 | 0 |
| Klebsiella:K02000 | 0 | 0 | 0 | 0 | 0 | 0 | 0 | 0 | 0 | 0.0021408 | 0.000477 | 0 | 0 | 0 | 0 | 0 | 0 | 0 | 0 | 0 | 0 |
| Klebsiella:K02002 | 0 | 0 | 0 | 0 | 0 | 0 | 0 | 0 | 0 | 0.0042816 | 0 | 0 | 0 | 0 | 0 | 0 | 0 | 0 | 0 | 0 | 0 |
| Klebsiella:K02011 | 0 | 0 | 0 | 0 | 0 | 0 | 0 | 0 | 0 | 0.0032112 | 0 | 0 | 0 | 0 | 0 | 0 | 0 | 0 | 0 | 0 | 0 |
| Klebsiella:K02012 | 0 | 0 | 0 | 0 | 0 | 0 | 0 | 0 | 0 | 0.0010704 | 0 | 0 | 0 | 0 | 0 | 0 | 0 | 0 | 0 | 0 | 0 |
| Klebsiella:K02047 | 0 | 0 | 0 | 0 | 0 | 0 | 0 | 0 | 0 | 0.0010704 | 0 | 0 | 0 | 0 | 0 | 0 | 0 | 0 | 0 | 0 | 0 |
| Klebsiella:K02048 | 0 | 0 | 0 | 0 | 0 | 0 | 0 | 0 | 0 | 0.0107041 | 0 | 0.0001138 | 0 | 0 | 0 | 0 | 0.000987 | 0 | 0 | 0 | 0 |
| Klebsiella:K02049 | 0 | 0 | 0 | 0 | 0 | 0 | 0 | 0 | 0 | 0.0021408 | 0 | 0 | 0 | 0 | 0 | 0 | 0 | 0 | 0 | 0 | 0 |
| Klebsiella:K02051 | 0 | 0 | 0 | 0 | 0 | 0 | 0 | 0 | 0 | 0.0021408 | 0 | 0.0001138 | 0 | 0 | 0 | 0 | 0 | 0 | 0 | 0 | 0 |
| Klebsiella:K02301 | 0 | 0 | 0 | 0 | 0 | 0 | 0 | 0 | 0 | 0.0032112 | 0 | 0.0001138 | 0 | 0 | 0 | 0 | 0 | 0 | 0 | 0 | 0 |
| Klebsiella:K02454 | 0 | 0 | 0 | 0 | 0 | 0 | 0 | 0 | 0 | 0.0042816 | 0 | 0 | 0 | 0 | 0 | 0 | 0 | 0 | 0 | 0 | 0 |
| Klebsiella:K02492 | 0 | 0 | 0 | 0 | 0 | 0 | 0 | 0 | 0 | 0.0021408 | 0 | 0 | 0 | 0 | 0 | 0 | 0 | 0 | 0 | 0 | 0 |
| Klebsiella:K02517 | 0 | 0 | 0 | 0 | 0 | 4.15E-05 | 0 | 0 | 0 | 0.0032112 | 0 | 0.0002277 | 0 | 0 | 0 | 0 | 0 | 0 | 0 | 0 | 0 |
| Klebsiella:K02523 | 0 | 0 | 0 | 0 | 0 | 0 | 0 | 0 | 0 | 0.0042816 | 0 | 0.0002277 | 0 | 0 | 0 | 0 | 0 | 0 | 0 | 0 | 0 |
| Klebsiella:K03072 | 0 | 0 | 0 | 0 | 0 | 0 | 0 | 0 | 0 | 0.0042816 | 0 | 0 | 0 | 0 | 0 | 0 | 0 | 0 | 0 | 0 | 0 |
| Klebsiella:K03087 | 0 | 0 | 0 | 0 | 0 | 0 | 0 | 0 | 0 | 0.0053521 | 0 | 0.0001138 | 0.001118 | 0 | 0 | 0 | 0 | 0 | 0 | 0 | 0 |
| Klebsiella:K03092 | 0 | 0 | 0 | 0 | 0 | 0 | 0 | 0 | 0 | 0.0042816 | 0 | 0 | 0 | 0 | 0 | 0 | 0 | 0 | 0 | 0 | 0 |
| Klebsiella:K03182 | 0 | 0 | 0 | 0 | 0 | 0 | 0 | 0.0007694 | 0 | 0.0021408 | 0 | 0 | 0 | 0 | 0 | 0 | 0 | 0 | 0 | 0 | 0 |
| Klebsiella:K03183 | 0 | 0.0205 | 0.0356 | 0.0065 | 0 | 0 | 0 | 0 | 0 | 0.0107041 | 0 | 0 | 0 | 0 | 0 | 0.000109 | 0 | 0 | 0 | 0 | 0 |
| Klebsiella:K03338 | 0 | 0 | 0 | 0 | 0 | 0 | 0 | 0 | 0 | 0.0042816 | 0 | 0 | 0 | 0 | 0 | 0 | 0 | 0 | 0 | 0 | 0 |
| Klebsiella:K03381 | 0 | 0 | 0 | 0 | 0 | 0 | 0 | 0 | 0 | 0.0010704 | 0 | 0 | 0 | 0 | 0 | 0 | 0 | 0 | 0 | 0 | 0 |
| Klebsiella:K03582 | 0 | 0 | 0 | 0 | 0 | 0 | 0 | 0 | 0 | 0.0042816 | 0 | 0.0001138 | 0 | 0 | 0 | 0 | 0 | 0 | 0 | 0 | 0 |
| Klebsiella:K03583 | 0 | 0 | 0 | 0 | 0 | 0 | 0 | 0 | 0 | 0.0085633 | 0.000477 | 0 | 0 | 0 | 0 | 0 | 0 | 0 | 0 | 0 | 0 |
| Klebsiella:K03590 | 0 | 0 | 0 | 0 | 0 | 0 | 0 | 0 | 0 | 0.0010704 | 0 | 0 | 0 | 0 | 0 | 0 | 0 | 0 | 0 | 0 | 0 |
| Klebsiella:K03644 | 0 | 0 | 0 | 0 | 0 | 0 | 0 | 0 | 0 | 0 | 0 | 0.0001138 | 0 | 0 | 0 | 0 | 0 | 0 | 0 | 0 | 0 |
| Klebsiella:K03732 | 0 | 0 | 0 | 0 | 0 | 0 | 0 | 0 | 0 | 0.0042816 | 0 | 0.0001138 | 0 | 0 | 0 | 0.000109 | 0 | 0 | 0 | 0 | 0 |
| Klebsiella:K03735 | 0 | 0 | 0 | 0 | 0 | 0 | 0 | 0 | 0 | 0.0053521 | 0 | 0 | 0.001118 | 0 | 0 | 0 | 0 | 0 | 0 | 0 | 0 |
| Klebsiella:K03781 | 0 | 0 | 0 | 0 | 0 | 0 | 0 | 0 | 0 | 0.0053521 | 0 | 0 | 0 | 0 | 0 | 0 | 0 | 0 | 0 | 0 | 0 |
| Klebsiella:K03841 | 0 | 0 | 0 | 0 | 0 | 0 | 0 | 0 | 0 | 0.0021408 | 0.000477 | 0.0001138 | 0 | 0 | 0 | 0 | 0 | 0 | 0 | 0 | 0 |
| Klebsiella:K03885 | 0 | 0 | 0 | 0 | 0 | 0 | 0 | 0 | 0 | 0.0032112 | 0 | 0 | 0 | 0 | 0 | 0 | 0 | 0 | 0 | 0 | 0 |
| Klebsiella:K04564 | 0 | 0 | 0 | 0 | 0 | 0 | 0 | 0 | 0 | 0.0032112 | 0 | 0 | 0 | 0 | 0 | 0.000109 | 0 | 0 | 0 | 0 | 0 |
| Klebsiella:K05343 | 0 | 0 | 0.0356 | 0 | 0 | 0 | 0 | 0.0007694 | 0 | 0.0042816 | 0 | 0 | 0 | 0 | 0 | 0 | 0.000987 | 0 | 0 | 0 | 0 |
| Klebsiella:K05365 | 0 | 0 | 0 | 0 | 0 | 0 | 0 | 0 | 0 | 0.0021408 | 0 | 0.0001138 | 0 | 0.000418 | 0 | 0 | 0 | 0 | 0 | 0 | 0 |
| Klebsiella:K05592 | 0 | 0 | 0 | 0 | 0 | 0 | 0 | 0 | 0 | 0.0010704 | 0 | 0.0001138 | 0 | 0 | 0 | 0 | 0 | 0 | 0 | 0 | 0 |
| Klebsiella:K05846 | 0 | 0 | 0 | 0 | 0 | 0 | 0 | 0 | 0 | 0.0042816 | 0 | 0 | 0 | 0 | 0 | 0 | 0 | 0 | 0 | 0 | 0 |
| Klebsiella:K05847 | 0 | 0 | 0 | 0 | 0 | 0 | 0 | 0 | 0 | 0.0064225 | 0 | 0 | 0 | 0 | 0 | 0 | 0 | 0 | 0 | 0 | 0 |
| Klebsiella:K06445 | 0 | 0 | 0 | 0 | 0 | 0 | 0 | 0 | 0 | 0.0074929 | 0 | 0.0001138 | 0 | 0 | 0 | 0 | 0 | 0 | 0 | 0 | 0 |
| Klebsiella:K06861 | 0 | 0 | 0 | 0 | 0 | 0 | 0 | 0 | 0 | 0.0010704 | 0 | 0.0001138 | 0 | 0 | 0 | 0 | 0 | 0 | 0 | 0 | 0 |

| | | | | | | | | | | | | | | | | | | | | | |
|---|---|---|---|---|---|---|---|---|---|---|---|---|---|---|---|---|---|---|---|---|---|
| Klebsiella:K07712 | 0 | 0 | 0 | 0 | 0 | 0 | 0 | 0 | 0 | 0.0021408 | 0 | 0 | 0.001118 | 0 | 0 | 0 | 0 | 0 | 0 | 0 | 0 |
| Klebsiella:K07788 | 0 | 0 | 0 | 0 | 0 | 0 | 0 | 0 | 0 | 0.0042816 | 0 | 0 | 0 | 0 | 0 | 0 | 0 | 0 | 0 | 0 | 0 |
| Klebsiella:K08484 | 0 | 0 | 0 | 0 | 0 | 0 | 0 | 0 | 0 | 0.0096337 | 0 | 0 | 0 | 0 | 0 | 0 | 0.000987 | 0 | 0 | 0 | 0 |
| Klebsiella:K11073 | 0 | 0 | 0 | 0 | 0 | 0 | 0 | 0 | 0 | 0.0021408 | 0 | 0 | 0 | 0 | 0 | 0 | 0 | 0 | 0 | 0 | 0 |
| Klebsiella:K11076 | 0 | 0 | 0 | 0 | 0 | 0 | 0 | 0 | 0 | 0.0021408 | 0 | 0 | 0 | 0 | 0 | 0 | 0 | 0 | 0 | 0 | 0 |
| Klebsiella:K11891 | 0 | 0 | 0 | 0 | 0 | 0 | 0 | 0 | 0 | 0.0128449 | 0.000477 | 0 | 0 | 0 | 0 | 0 | 0.001974 | 0 | 0 | 0 | 0 |
| Klebsiella:K11904 | 0 | 0 | 0 | 0 | 0 | 4.15E-05 | 0.00662 | 0.0030776 | 0.002051 | 0.0107041 | 0 | 0.0005692 | 0 | 0 | 0 | 0 | 0 | 0 | 0 | 0 | 0 |
| Klebsiella:K11907 | 0 | 0 | 0 | 0 | 0 | 0 | 0.004413 | 0 | 0 | 0.0042816 | 0 | 0 | 0 | 0 | 0 | 0 | 0 | 0 | 0 | 0 | 0 |
| Klebsiella:K11927 | 0 | 0 | 0 | 0.0032 | 0 | 0 | 0 | 0 | 0 | 0.0053521 | 0 | 0 | 0 | 0 | 0 | 0 | 0 | 0 | 0 | 0 | 0 |
| Klebsiella:K11959 | 0 | 0 | 0 | 0 | 0 | 0 | 0.002207 | 0.0007694 | 0 | 0.0010704 | 0 | 0 | 0 | 0 | 0 | 0 | 0 | 0 | 0 | 0 | 0 |
| Klebsiella:K11962 | 0 | 0 | 0 | 0 | 0 | 0 | 0 | 0 | 0 | 0.0010704 | 0 | 0 | 0 | 0 | 0 | 0 | 0 | 0 | 0 | 0 | 0 |
| Klebsiella:K11963 | 0 | 0 | 0 | 0 | 0 | 0 | 0 | 0 | 0 | 0.0042816 | 0 | 0 | 0 | 0 | 0 | 0 | 0 | 0 | 0 | 0 | 0 |
| Klebsiella:K12339 | 0 | 0 | 0 | 0 | 0 | 0 | 0 | 0 | 0 | 0 | 0 | 0 | 0.001118 | 0 | 0 | 0 | 0 | 0 | 0 | 0 | 0 |
| Klebsiella:K12368 | 0 | 0 | 0 | 0 | 0 | 0 | 0 | 0.0007694 | 0.001025 | 0.0074929 | 0.000477 | 0 | 0 | 0 | 0 | 0 | 0 | 0 | 0 | 0 | 0 |
| Klebsiella:K12573 | 0 | 0 | 0 | 0 | 0 | 0 | 0 | 0 | 0 | 0.0042816 | 0 | 0.0001138 | 0 | 0 | 0 | 0 | 0 | 0 | 0 | 0 | 0 |
| Klebsiella:K13378 | 0 | 0 | 0 | 0 | 0 | 0 | 0 | 0 | 0 | 0.0021408 | 0 | 0 | 0 | 0 | 0 | 0 | 0 | 0 | 0 | 0 | 0 |
| Klebsiella:K13895 | 0 | 0 | 0 | 0 | 0 | 0 | 0 | 0 | 0 | 0.0064225 | 0 | 0.0001138 | 0 | 0 | 0 | 0 | 0 | 0 | 0 | 0 | 0 |
| Lactobacillus:K00036 | 0 | 0 | 0 | 0 | 0 | 4.15E-05 | 0.004413 | 0 | 0.002051 | 0 | 0 | 0 | 0 | 0.000418 | 0 | 0 | 0 | 0.0373413 | 0.065965351 | 0 | 0.110514683 |
| Lactobacillus:K00111 | 0 | 0 | 0 | 0 | 0 | 8.29E-05 | 0.002207 | 0.0023082 | 0.001025 | 0 | 0 | 0 | 0 | 0 | 0 | 0 | 0 | 0 | 0 | 0 | 0 |
| Lactobacillus:K00131 | 0 | 0 | 0 | 0 | 0 | 0.0012 | 0 | 0 | 0 | 0 | 0 | 0 | 0 | 0 | 0 | 0 | 0 | 0 | 0 | 0 | 0 |
| Lactobacillus:K00135 | 0 | 0 | 0 | 0 | 0 | 0 | 0 | 0 | 0 | 0.0010704 | 0 | 0 | 0.001118 | 0.000418 | 0 | 0 | 0 | 0.186706497 | 0.090268375 | 0 | 0.047363435 |
| Lactobacillus:K00140 | 0 | 0 | 0 | 0 | 0 | 0 | 0 | 0 | 0 | 0 | 0 | 0 | 0 | 0 | 0 | 0 | 0.000987 | 0 | 0 | 0 | 0 |
| Lactobacillus:K00161 | 0 | 0 | 0 | 0 | 0 | 4.15E-05 | 0 | 0 | 0 | 0 | 0 | 0 | 0.002235 | 0 | 0 | 0.000109 | 0 | 0 | 0.013887442 | 0 | 0.007893906 |
| Lactobacillus:K00162 | 0 | 0 | 0 | 0 | 0 | 4.15E-05 | 0.00662 | 0 | 0 | 0 | 0 | 0 | 0 | 0 | 0 | 0 | 0 | 0 | 0.003471861 | 0 | 0 |
| Lactobacillus:K00383 | 0 | 0 | 0 | 0 | 0 | 0 | 0.002207 | 0.0015388 | 0 | 0 | 0 | 0 | 0 | 0 | 0 | 0.000109 | 0 | 0 | 0 | 0 | 0 |
| Lactobacillus:K00425 | 0 | 0 | 0 | 0.0016 | 0 | 0 | 0 | 0 | 0 | 0 | 0 | 0 | 0.001118 | 0 | 0 | 0.000109 | 0 | 0 | 0.006943721 | 0 | 0 |
| Lactobacillus:K00525 | 0 | 0.0205 | 0 | 0.0016 | 0 | 4.15E-05 | 0.00662 | 0.0007694 | 0 | 0 | 0 | 0 | 0.00447 | 0 | 0 | 0 | 0 | 0.149365198 | 0.107627678 | 0 | 0.126302495 |
| Lactobacillus:K00526 | 0 | 0 | 0 | 0.0016 | 0 | 4.15E-05 | 0.002207 | 0 | 0 | 0 | 0 | 0 | 0.002235 | 0 | 0 | 0 | 0 | 0.112023898 | 0.065965351 | 0 | 0.110514683 |
| Lactobacillus:K00575 | 0 | 0 | 0 | 0 | 0 | 0 | 0 | 0 | 0 | 0 | 0 | 0 | 0 | 0 | 0 | 0 | 0 | 0.186706497 | 0.013887442 | 0 | 0.023681718 |
| Lactobacillus:K00600 | 0 | 0 | 0 | 0 | 0 | 4.15E-05 | 0.002207 | 0 | 0 | 0.0032112 | 0 | 0.0004553 | 0.00447 | 0.002506 | 0 | 0.000109 | 0 | 0 | 0.010415582 | 0 | 0 |
| Lactobacillus:K00626 | 0 | 0 | 0 | 0 | 0 | 4.15E-05 | 0 | 0 | 0 | 0 | 0 | 0 | 0 | 0.000418 | 0 | 0 | 0 | 0 | 0.003471861 | 0 | 0 |
| Lactobacillus:K00627 | 0 | 0 | 0 | 0 | 0 | 0 | 0.002207 | 0 | 0.001025 | 0 | 0 | 0 | 0 | 0 | 0 | 0 | 0 | 0 | 0 | 0 | 0.007893906 |
| Lactobacillus:K00640 | 0 | 0 | 0 | 0 | 0 | 0 | 0 | 0 | 0 | 0.0010704 | 0 | 0 | 0 | 0.000418 | 0 | 0 | 0 | 0 | 0 | 0 | 0 |
| Lactobacillus:K00648 | 0 | 0 | 0 | 0 | 0 | 8.29E-05 | 0 | 0.0007694 | 0.001025 | 0 | 0 | 0 | 0.001118 | 0.000418 | 0 | 0.000109 | 0 | 0 | 0.034718606 | 0 | 0.078939059 |
| Lactobacillus:K00864 | 0 | 0 | 0 | 0.0032 | 0 | 8.29E-05 | 0.008827 | 0 | 0.001025 | 0 | 0.000955 | 0.0012522 | 0.003353 | 0.000835 | 0.000104 | 0.000435 | 0.000987 | 0 | 0.017359303 | 0 | 0 |
| Lactobacillus:K00940 | 0 | 0 | 0 | 0 | 0 | 0 | 0 | 0 | 0 | 0 | 0 | 0 | 0 | 0 | 0 | 0 | 0 | 0 | 0.003471861 | 0 | 0 |
| Lactobacillus:K01142 | 0 | 0 | 0 | 0.0016 | 0 | 0 | 0.00662 | 0 | 0 | 0.0010704 | 0.002864 | 0.0018214 | 0.003353 | 0 | 0.000104 | 0.000979 | 0 | 0.0373413 | 0.041662327 | 0 | 0.063151247 |
| Lactobacillus:K01338 | 0 | 0 | 0 | 0 | 0 | 0 | 0 | 0 | 0 | 0 | 0.000477 | 0 | 0.001118 | 0 | 0 | 0.000109 | 0 | 0 | 0 | 0 | 0 |
| Lactobacillus:K01426 | 0 | 0 | 0 | 0 | 0 | 0 | 0 | 0 | 0 | 0.0010704 | 0 | 0 | 0 | 0 | 0 | 0 | 0 | 0.261389096 | 0.124986981 | 0 | 0.165772024 |
| Lactobacillus:K01478 | 0 | 0 | 0 | 0.0016 | 0 | 0.00012 | 0.00662 | 0 | 0 | 0 | 0 | 0 | 0.001118 | 0 | 0 | 0 | 0 | 0.074682599 | 0.013887442 | 0 | 0.015787812 |
| Lactobacillus:K01488 | 0 | 0 | 0 | 0 | 0 | 0 | 0 | 0 | 0 | 0 | 0.000477 | 0.000683 | 0.001118 | 0.000835 | 0 | 0 | 0 | 0 | 0 | 0 | 0 |
| Lactobacillus:K01581 | 0 | 0 | 0 | 0 | 0 | 0 | 0 | 0 | 0 | 0 | 0 | 0 | 0 | 0 | 0 | 0 | 0 | 0 | 0.003471861 | 0 | 0 |
| Lactobacillus:K01627 | 0 | 0 | 0 | 0 | 0 | 0 | 0 | 0 | 0 | 0 | 0 | 0 | 0 | 0 | 0 | 0 | 0 | 0 | 0.003471861 | 0 | 0 |
| Lactobacillus:K01629 | 0 | 0 | 0 | 0 | 0 | 0 | 0 | 0.0007694 | 0 | 0 | 0 | 0 | 0 | 0 | 0 | 0 | 0 | 0.074682599 | 0.086796514 | 0 | 0.03946953 |
| Lactobacillus:K01679 | 0 | 0 | 0 | 0 | 0 | 0 | 0 | 0 | 0 | 0 | 0 | 0 | 0 | 0 | 0 | 0 | 0 | 0.0373413 | 0.006943721 | 0 | 0.007893906 |
| Lactobacillus:K01878 | 0 | 0 | 0 | 0.0016 | 0 | 4.15E-05 | 0 | 0 | 0 | 0 | 0 | 0 | 0 | 0 | 0 | 0 | 0 | 0.074682599 | 0.083324654 | 0 | 0.102620777 |
| Lactobacillus:K01879 | 0 | 0 | 0 | 0 | 0 | 0 | 0.015447 | 0.0007694 | 0.001025 | 0.0010704 | 0 | 0 | 0.001118 | 0 | 0.000104 | 0 | 0 | 0.074682599 | 0.048606048 | 0 | 0.047363435 |
| Lactobacillus:K01915 | 0 | 0 | 0 | 0.0048 | 0 | 4.15E-05 | 0.004413 | 0 | 0 | 0 | 0 | 0.0003415 | 0.001118 | 0 | 0 | 0 | 0 | 0.224047797 | 0.114571399 | 0 | 0.197347648 |
| Lactobacillus:K01958 | 0 | 0 | 0 | 0 | 0 | 0 | 0 | 0 | 0 | 0 | 0 | 0.0001138 | 0 | 0 | 0 | 0 | 0 | 0.0373413 | 0.031246745 | 0 | 0.063151247 |
| Lactobacillus:K01961 | 0 | 0 | 0 | 0 | 0 | 4.15E-05 | 0.011033 | 0.0015388 | 0 | 0 | 0 | 0.0002277 | 0 | 0.000835 | 0.000104 | 0 | 0 | 0.112023898 | 0.097212096 | 0 | 0.102620777 |
| Lactobacillus:K01962 | 0 | 0 | 0 | 0 | 0 | 0 | 0.002207 | 0 | 0.002051 | 0.0010704 | 0 | 0 | 0 | 0 | 0 | 0.000326 | 0 | 0.074682599 | 0.045134187 | 0 | 0.03946953 |
| Lactobacillus:K02000 | 0 | 0 | 0 | 0 | 0 | 8.29E-05 | 0.00662 | 0 | 0.001025 | 0 | 0 | 0 | 0 | 0 | 0 | 0 | 0 | 0 | 0 | 0 | 0 |
| Lactobacillus:K02002 | 0 | 0 | 0 | 0 | 0 | 0 | 0 | 0 | 0 | 0 | 0.000477 | 0 | 0 | 0.000418 | 0 | 0 | 0 | 0 | 0 | 0 | 0 |
| Lactobacillus:K02011 | 0 | 0 | 0 | 0 | 0 | 0 | 0 | 0 | 0 | 0 | 0 | 0.0001138 | 0 | 0 | 0 | 0 | 0 | 0 | 0 | 0 | 0 |
| Lactobacillus:K02372 | 0 | 0 | 0 | 0.0032 | 0 | 0 | 0.002207 | 0 | 0 | 0 | 0 | 0.000683 | 0.001118 | 0.000418 | 0 | 0.000435 | 0 | 0.074682599 | 0.06249349 | 0 | 0.063151247 |
| Lactobacillus:K02400 | 0 | 0 | 0 | 0 | 0 | 0 | 0 | 0 | 0 | 0 | 0 | 0 | 0 | 0 | 0 | 0 | 0 | 0 | 0.045134187 | 0 | 0.055257341 |
| Lactobacillus:K02405 | 0 | 0 | 0 | 0 | 0 | 0 | 0 | 0 | 0 | 0 | 0 | 0 | 0 | 0 | 0 | 0 | 0 | 0 | 0.013887442 | 0 | 0.015787812 |
| Lactobacillus:K02409 | 0 | 0 | 0 | 0 | 0 | 0 | 0 | 0 | 0 | 0 | 0 | 0 | 0 | 0 | 0 | 0 | 0 | 0 | 0.038190466 | 0 | 0.102620777 |
| Lactobacillus:K02410 | 0 | 0 | 0 | 0 | 0 | 0 | 0 | 0 | 0 | 0 | 0 | 0 | 0 | 0 | 0 | 0 | 0 | 0.074682599 | 0.048606048 | 0 | 0.047363435 |
| Lactobacillus:K02412 | 0 | 0 | 0 | 0 | 0 | 0 | 0 | 0 | 0 | 0 | 0 | 0 | 0 | 0 | 0 | 0 | 0 | 0.112023898 | 0.055549769 | 0 | 0.094726871 |

| | | | | | | | | | | | | | | | | | | | | | |
|---|---|---|---|---|---|---|---|---|---|---|---|---|---|---|---|---|---|---|---|---|---|
| Lactobacillus:K02416 | 0 | 0 | 0 | 0 | 0 | 0 | 0 | 0 | 0 | 0.0010704 | 0 | 0 | 0 | 0 | 0 | 0 | 0 | 0.074682599 | 0.06249349 | 0 | 0.149984212 |
| Lactobacillus:K02556 | 0 | 0 | 0 | 0 | 0 | 0 | 0 | 0 | 0 | 0 | 0 | 0 | 0 | 0 | 0 | 0 | 0 | 0 | 0.048606048 | 0 | 0.086832965 |
| Lactobacillus:K02557 | 0 | 0 | 0 | 0 | 0 | 0 | 0 | 0 | 0 | 0 | 0.000477 | 0 | 0 | 0 | 0 | 0 | 0 | 0.074682599 | 0.024303024 | 0 | 0.102620777 |
| Lactobacillus:K03092 | 0 | 0 | 0 | 0 | 0 | 4.15E-05 | 0 | 0 | 0 | 0 | 0 | 0 | 0 | 0 | 0 | 0 | 0 | 0 | 0.020831163 | 0 | 0.071045153 |
| Lactobacillus:K03183 | 0 | 0 | 0 | 0 | 0 | 0 | 0 | 0 | 0 | 0 | 0 | 0 | 0.001118 | 0 | 0 | 0 | 0 | 0 | 0.006943721 | 0 | 0 |
| Lactobacillus:K03407 | 0 | 0 | 0 | 0 | 0 | 0 | 0 | 0 | 0 | 0 | 0 | 0 | 0 | 0 | 0 | 0 | 0 | 0 | 0.031246745 | 0 | 0.078939059 |
| Lactobacillus:K03408 | 0 | 0 | 0 | 0 | 0 | 4.15E-05 | 0 | 0 | 0 | 0 | 0 | 0 | 0 | 0 | 0 | 0 | 0 | 0.074682599 | 0.010415582 | 0 | 0.03946953 |
| Lactobacillus:K03412 | 0 | 0 | 0 | 0 | 0 | 8.29E-05 | 0 | 0 | 0 | 0 | 0 | 0 | 0 | 0 | 0 | 0 | 0 | 0.0373413 | 0.017359303 | 0 | 0.007893906 |
| Lactobacillus:K03590 | 0 | 0 | 0 | 0 | 0 | 0 | 0.004413 | 0.0007694 | 0 | 0 | 0 | 0 | 0 | 0 | 0 | 0 | 0 | 0 | 0.093740235 | 0 | 0.110514683 |
| Lactobacillus:K03781 | 0 | 0 | 0 | 0 | 0 | 0 | 0.002207 | 0 | 0.001025 | 0 | 0 | 0 | 0 | 0.000418 | 0 | 0 | 0 | 0 | 0 | 0 | 0 |
| Lactobacillus:K03885 | 0 | 0 | 0 | 0 | 0 | 0 | 0 | 0 | 0 | 0 | 0 | 0 | 0.001118 | 0 | 0.000104 | 0 | 0 | 0 | 0.003471861 | 0 | 0 |
| Lactobacillus:K04564 | 0 | 0 | 0 | 0 | 0 | 4.15E-05 | 0.00662 | 0 | 0 | 0 | 0 | 0 | 0 | 0 | 0 | 0 | 0 | 0 | 0 | 0 | 0 |
| Lactobacillus:K05592 | 0 | 0 | 0 | 0.0016 | 0 | 8.29E-05 | 0.004413 | 0.0007694 | 0 | 0.0010704 | 0.000477 | 0.0001138 | 0 | 0 | 0 | 0 | 0 | 0 | 0.038190466 | 0 | 0.007893906 |
| Lactobacillus:K05846 | 0 | 0 | 0 | 0 | 0 | 0.00012 | 0 | 0 | 0 | 0 | 0 | 0.0001138 | 0 | 0.000835 | 0.000104 | 0 | 0 | 0 | 0.006943721 | 0 | 0.015787812 |
| Lactobacillus:K05847 | 0 | 0 | 0 | 0.0016 | 0 | 4.15E-05 | 0.008827 | 0.0007694 | 0.001025 | 0 | 0 | 0.0002277 | 0 | 0.000835 | 0 | 0 | 0 | 0 | 0.006943721 | 0 | 0.031575624 |
| Lactobacillus:K06148 | 0 | 0 | 0 | 0 | 0 | 0 | 0 | 0 | 0 | 0.0010704 | 0.001909 | 0 | 0 | 0.000418 | 0.000104 | 0.000109 | 0 | 0 | 0.027774885 | 0 | 0 |
| Lactobacillus:K10536 | 0 | 0 | 0 | 0.0016 | 0 | 0 | 0.002207 | 0 | 0.001025 | 0 | 0 | 0 | 0 | 0 | 0 | 0.00087 | 0 | 0 | 0 | 0 | 0 |
| Lactobacillus:K12573 | 0 | 0 | 0 | 0 | 0 | 0 | 0.004413 | 0 | 0.001025 | 0.0010704 | 0.000477 | 0.0005692 | 0.002235 | 0.001253 | 0.000104 | 0.001306 | 0 | 0.0373413 | 0.104155817 | 0 | 0.1341964 |
| Lactococcus:K00383 | 0 | 0 | 0 | 0 | 0 | 0 | 0 | 0 | 0 | 0.0010704 | 0 | 0 | 0 | 0 | 0 | 0 | 0 | 0 | 0 | 0 | 0 |
| Lactococcus:K00425 | 0 | 0 | 0 | 0 | 0 | 4.15E-05 | 0 | 0.0007694 | 0 | 0 | 0 | 0 | 0 | 0 | 0 | 0 | 0 | 0 | 0 | 0 | 0 |
| Lactococcus:K00432 | 0 | 0 | 0 | 0.0016 | 0 | 0 | 0 | 0 | 0.001025 | 0 | 0 | 0 | 0 | 0 | 0 | 0 | 0 | 0 | 0 | 0 | 0 |
| Lactococcus:K00526 | 0 | 0 | 0 | 0 | 0 | 0 | 0 | 0 | 0 | 0 | 0 | 0 | 0.001118 | 0 | 0 | 0 | 0 | 0 | 0 | 0 | 0 |
| Lactococcus:K00600 | 0 | 0 | 0 | 0 | 0 | 0 | 0 | 0 | 0 | 0.0032112 | 0.000477 | 0 | 0 | 0 | 0 | 0 | 0 | 0 | 0 | 0 | 0 |
| Lactococcus:K00640 | 0 | 0 | 0 | 0 | 0 | 0 | 0 | 0 | 0 | 0.0010704 | 0 | 0 | 0 | 0 | 0 | 0 | 0 | 0 | 0 | 0 | 0 |
| Lactococcus:K00648 | 0 | 0 | 0 | 0 | 0 | 4.15E-05 | 0 | 0.0015388 | 0 | 0 | 0 | 0 | 0 | 0 | 0 | 0 | 0 | 0 | 0 | 0 | 0 |
| Lactococcus:K00864 | 0 | 0 | 0 | 0.0016 | 0 | 0 | 0 | 0 | 0 | 0.0010704 | 0.001432 | 0 | 0 | 0.000418 | 0 | 0 | 0 | 0 | 0 | 0 | 0 |
| Lactococcus:K01426 | 0 | 0 | 0 | 0 | 0 | 0 | 0 | 0 | 0 | 0.0010704 | 0 | 0 | 0 | 0 | 0 | 0 | 0 | 0 | 0 | 0 | 0 |
| Lactococcus:K01478 | 0 | 0 | 0 | 0 | 0 | 4.15E-05 | 0 | 0 | 0 | 0 | 0 | 0 | 0 | 0.001253 | 0 | 0 | 0 | 0 | 0 | 0 | 0 |
| Lactococcus:K01915 | 0 | 0 | 0 | 0 | 0 | 0 | 0 | 0 | 0 | 0 | 0 | 0 | 0 | 0 | 0.000104 | 0 | 0 | 0 | 0 | 0 | 0 |
| Lactococcus:K01958 | 0 | 0 | 0 | 0.0016 | 0 | 0 | 0 | 0 | 0 | 0 | 0.000477 | 0 | 0 | 0 | 0 | 0 | 0 | 0 | 0 | 0 | 0 |
| Lactococcus:K01961 | 0 | 0 | 0 | 0 | 0 | 0 | 0 | 0 | 0 | 0 | 0 | 0 | 0 | 0.000418 | 0 | 0 | 0 | 0 | 0 | 0 | 0 |
| Lactococcus:K01962 | 0 | 0 | 0 | 0 | 0 | 0 | 0 | 0 | 0.001025 | 0 | 0 | 0 | 0 | 0 | 0 | 0 | 0 | 0 | 0 | 0 | 0 |
| Lactococcus:K02002 | 0 | 0 | 0 | 0.0016 | 0 | 0 | 0 | 0 | 0 | 0 | 0 | 0 | 0 | 0 | 0 | 0 | 0 | 0 | 0 | 0 | 0 |
| Lactococcus:K02372 | 0 | 0 | 0 | 0 | 0 | 0 | 0 | 0 | 0 | 0.0010704 | 0 | 0 | 0 | 0 | 0 | 0 | 0 | 0 | 0 | 0 | 0 |
| Lactococcus:K03590 | 0 | 0 | 0 | 0 | 0 | 0 | 0 | 0 | 0 | 0.0010704 | 0 | 0 | 0 | 0 | 0 | 0 | 0 | 0 | 0 | 0 | 0 |
| Lactococcus:K05846 | 0 | 0 | 0 | 0 | 0 | 4.15E-05 | 0 | 0 | 0 | 0 | 0 | 0 | 0 | 0 | 0 | 0 | 0 | 0 | 0 | 0 | 0 |
| Lactococcus:K10536 | 0 | 0 | 0 | 0 | 0 | 4.15E-05 | 0 | 0 | 0 | 0.0010704 | 0.001432 | 0 | 0.003353 | 0.003341 | 0 | 0 | 0 | 0 | 0 | 0 | 0 |
| Methanobrevibacter:K00123 | 0 | 0.4707 | 0.2495 | 0.3941 | 0 | 0.00012 | 0 | 0 | 0 | 0 | 0.003818 | 0.0026182 | 0.002235 | 0.022132 | 0 | 0.000544 | 0 | 0 | 0 | 0 | 0 |
| Methanobrevibacter:K00383 | 0 | 0.0409 | 0 | 0.0581 | 0 | 0 | 0 | 0 | 0 | 0 | 0 | 0 | 0 | 0 | 0 | 0 | 0 | 0 | 0 | 0 | 0 |
| Methanobrevibacter:K00432 | 0 | 0.0205 | 0.1069 | 0.0275 | 0 | 0 | 0 | 0 | 0 | 0 | 0 | 0 | 0 | 0 | 0 | 0 | 0 | 0 | 0 | 0 | 0 |
| Methanobrevibacter:K00600 | 0 | 0.0819 | 0.0356 | 0.1405 | 0 | 0 | 0 | 0 | 0 | 0.0010704 | 0.002386 | 0.0012522 | 0.003353 | 0.016286 | 0 | 0.000109 | 0 | 0 | 0 | 0 | 0 |
| Methanobrevibacter:K00626 | 0 | 0.0205 | 0.0713 | 0.1551 | 0 | 4.15E-05 | 0 | 0 | 0 | 0.0010704 | 0.000955 | 0.0012522 | 0.002235 | 0.017539 | 0 | 0.000326 | 0 | 0 | 0 | 0 | 0 |
| Methanobrevibacter:K00640 | 0 | 0.0409 | 0 | 0.0678 | 0 | 0 | 0 | 0 | 0 | 0 | 0.001909 | 0.0011384 | 0.001118 | 0.003341 | 0 | 0 | 0 | 0 | 0 | 0 | 0 |
| Methanobrevibacter:K00641 | 0 | 0.0409 | 0.1069 | 0.0905 | 0 | 4.15E-05 | 0 | 0 | 0 | 0.0032112 | 0.003341 | 0.0025044 | 0.002235 | 0.015868 | 0 | 0.000653 | 0 | 0 | 0 | 0 | 0 |
| Methanobrevibacter:K00833 | 0 | 0.0409 | 0 | 0.1131 | 0 | 0 | 0 | 0 | 0 | 0 | 0 | 0 | 0 | 0 | 0 | 0 | 0 | 0 | 0 | 0 | 0 |
| Methanobrevibacter:K00940 | 0 | 0 | 0 | 0.0388 | 0 | 0 | 0 | 0 | 0 | 0 | 0.000955 | 0 | 0.001118 | 0.003758 | 0 | 0.000109 | 0 | 0 | 0 | 0 | 0 |
| Methanobrevibacter:K01142 | 0 | 0.0819 | 0.0713 | 0.0581 | 0 | 4.15E-05 | 0 | 0 | 0 | 0.0010704 | 0.001432 | 0.0007969 | 0.001118 | 0.010857 | 0 | 0.000109 | 0 | 0 | 0 | 0 | 0 |
| Methanobrevibacter:K01251 | 0 | 0.1228 | 0.2138 | 0.2035 | 0 | 0 | 0 | 0 | 0 | 0.0010704 | 0.004773 | 0.0022767 | 0.001118 | 0.032572 | 0 | 0.000653 | 0 | 0 | 0 | 0 | 0 |
| Methanobrevibacter:K01338 | 0 | 0 | 0 | 0.0016 | 0 | 0 | 0 | 0 | 0 | 0 | 0 | 0 | 0 | 0 | 0 | 0.000326 | 0 | 0 | 0 | 0 | 0 |
| Methanobrevibacter:K01698 | 0 | 0 | 0 | 0.0291 | 0 | 4.15E-05 | 0 | 0 | 0 | 0.0010704 | 0.003341 | 0.0012522 | 0 | 0.010857 | 0 | 0.000326 | 0 | 0 | 0 | 0 | 0 |
| Methanobrevibacter:K01737 | 0 | 0.0205 | 0.1069 | 0.0678 | 0 | 0 | 0 | 0 | 0 | 0.0010704 | 0.001432 | 0.0001138 | 0.001118 | 0.003758 | 0 | 0.000109 | 0 | 0 | 0 | 0 | 0 |
| Methanobrevibacter:K01749 | 0 | 0 | 0.0356 | 0.0598 | 0 | 0 | 0 | 0 | 0 | 0.0010704 | 0.002386 | 0.001366 | 0.001118 | 0.010022 | 0 | 0 | 0 | 0 | 0 | 0 | 0 |
| Methanobrevibacter:K01845 | 0 | 0.0205 | 0 | 0.0953 | 0 | 0 | 0 | 0 | 0 | 0.0053521 | 0.003341 | 0.000683 | 0.001118 | 0.009604 | 0 | 0.000109 | 0 | 0 | 0 | 0 | 0 |
| Methanobrevibacter:K01895 | 0 | 0.1023 | 0.0713 | 0.1712 | 0 | 0 | 0 | 0 | 0 | 0.0042816 | 0.004295 | 0.0018214 | 0.00447 | 0.018791 | 0 | 0.000435 | 0 | 0 | 0 | 0.021473051 | 0 |
| Methanobrevibacter:K01902 | 0 | 0.0205 | 0 | 0.0291 | 0 | 0 | 0 | 0 | 0 | 0.0021408 | 0.001909 | 0.0003415 | 0 | 0.007934 | 0 | 0 | 0 | 0 | 0 | 0 | 0 |
| Methanobrevibacter:K01903 | 0 | 0.0205 | 0.0356 | 0.0614 | 0 | 0 | 0 | 0 | 0 | 0.0010704 | 0.001909 | 0.0009107 | 0.001118 | 0.009187 | 0 | 0.000218 | 0 | 0 | 0 | 0 | 0 |
| Methanobrevibacter:K01915 | 0 | 0.307 | 0.3207 | 0.4022 | 0 | 0 | 0 | 0 | 0 | 0.0010704 | 0.002386 | 0.0017076 | 0 | 0.020462 | 0 | 0.000326 | 0 | 0 | 0 | 0 | 0 |
| Methanobrevibacter:K01959 | 0 | 0.0205 | 0.0713 | 0.1744 | 0 | 0 | 0 | 0 | 0 | 0.0010704 | 0.004295 | 0.0011384 | 0 | 0.015033 | 0 | 0.000326 | 0 | 0 | 0 | 0 | 0 |
| Methanobrevibacter:K01960 | 0 | 0.1228 | 0.1426 | 0.2067 | 0 | 4.15E-05 | 0 | 0 | 0 | 0.0010704 | 0.001432 | 0.0018214 | 0.001118 | 0.014615 | 0 | 0.000109 | 0 | 0 | 0 | 0 | 0 |
| Methanobrevibacter:K01961 | 0 | 0 | 0 | 0.0016 | 0 | 0 | 0 | 0 | 0 | 0 | 0 | 0 | 0 | 0 | 0 | 0 | 0 | 0 | 0 | 0 | 0 |

| | | | | | | | | | | | | | | | | | | | | | |
|---|---|---|---|---|---|---|---|---|---|---|---|---|---|---|---|---|---|---|---|---|---|
| Methanobrevibacter:K02049 | 0 | 0.0614 | 0.0356 | 0.0824 | 0 | 0 | 0 | 0 | 0 | 0 | 0.002386 | 0.0005692 | 0 | 0.007099 | 0 | 0 | 0 | 0 | 0 | 0 | 0 |
| Methanobrevibacter:K02051 | 0 | 0.0409 | 0.0713 | 0.0307 | 0 | 4.15E-05 | 0 | 0 | 0 | 0 | 0.000477 | 0.000683 | 0.002235 | 0.004176 | 0 | 0.000109 | 0 | 0 | 0 | 0 | 0 |
| Methanobrevibacter:K02492 | 0 | 0 | 0.0356 | 0.0242 | 0 | 0 | 0 | 0 | 0 | 0.0053521 | 0.003818 | 0.0010245 | 0.001118 | 0.007934 | 0 | 0.000109 | 0 | 0 | 0 | 0 | 0 |
| Methanobrevibacter:K03072 | 0 | 0 | 0.0356 | 0.0485 | 0 | 0 | 0 | 0 | 0 | 0 | 0 | 0 | 0 | 0 | 0 | 0 | 0 | 0 | 0 | 0 | 0 |
| Methanobrevibacter:K03182 | 0 | 0 | 0 | 0.0016 | 0 | 0 | 0 | 0 | 0 | 0 | 0 | 0 | 0 | 0 | 0 | 0 | 0 | 0 | 0 | 0 | 0 |
| Methanobrevibacter:K03394 | 0 | 0.0409 | 0.0356 | 0.021 | 0 | 0 | 0 | 0 | 0 | 0 | 0.000955 | 0.0009107 | 0.001118 | 0.008769 | 0 | 0.000109 | 0 | 0 | 0 | 0 | 0 |
| Methanobrevibacter:K05592 | 0 | 0 | 0 | 0.0032 | 0 | 0 | 0 | 0 | 0 | 0.0010704 | 0.003818 | 0.0015937 | 0 | 0.011692 | 0 | 0 | 0 | 0 | 0 | 0 | 0 |
| Methanobrevibacter:K05846 | 0 | 0 | 0 | 0 | 0 | 0 | 0 | 0 | 0 | 0 | 0.001909 | 0.0012522 | 0.001118 | 0.015868 | 0 | 0.000109 | 0 | 0 | 0 | 0 | 0 |
| Methanobrevibacter:K05847 | 0 | 0 | 0 | 0 | 0 | 0 | 0 | 0 | 0 | 0 | 0 | 0.0010245 | 0.001118 | 0.007099 | 0 | 0.000218 | 0 | 0 | 0 | 0 | 0 |
| Mogibacterium:K00248 | 0 | 0 | 0 | 0 | 0 | 0 | 0 | 0 | 0 | 0 | 0 | 0.0001138 | 0 | 0 | 0 | 0 | 0 | 0 | 0 | 0 | 0 |
| Mogibacterium:K00336 | 0 | 0 | 0 | 0 | 0 | 0 | 0 | 0 | 0 | 0 | 0 | 0 | 0 | 0 | 0 | 0.000109 | 0 | 0 | 0 | 0 | 0 |
| Mogibacterium:K01142 | 0 | 0 | 0 | 0.0016 | 0 | 0 | 0 | 0 | 0 | 0 | 0 | 0 | 0 | 0 | 0 | 0 | 0 | 0 | 0 | 0 | 0 |
| Mogibacterium:K01915 | 0 | 0 | 0 | 0 | 0 | 4.15E-05 | 0 | 0 | 0 | 0 | 0 | 0.0001138 | 0 | 0 | 0 | 0 | 0 | 0 | 0 | 0 | 0 |
| Mogibacterium:K02203 | 0 | 0 | 0 | 0 | 0 | 0 | 0.002207 | 0 | 0 | 0 | 0 | 0 | 0 | 0 | 0 | 0.000109 | 0 | 0 | 0 | 0 | 0 |
| Mogibacterium:K12573 | 0 | 0 | 0 | 0 | 0 | 0 | 0 | 0 | 0 | 0 | 0 | 0 | 0 | 0 | 0 | 0.000109 | 0 | 0 | 0 | 0 | 0 |
| Oribacterium:K00104 | 0 | 0 | 0 | 0 | 0 | 0 | 0 | 0 | 0 | 0 | 0.001432 | 0.0011384 | 0 | 0 | 0 | 0.000544 | 0 | 0 | 0 | 0 | 0 |
| Oribacterium:K00111 | 0 | 0 | 0 | 0 | 0 | 0 | 0 | 0 | 0 | 0 | 0.000477 | 0.0001138 | 0 | 0 | 0 | 0 | 0 | 0 | 0 | 0 | 0 |
| Oribacterium:K00140 | 0 | 0 | 0 | 0 | 0 | 0 | 0 | 0 | 0 | 0 | 0 | 0.0009107 | 0 | 0.000418 | 0 | 0 | 0.009872 | 0 | 0 | 0 | 0 |
| Oribacterium:K00526 | 0 | 0 | 0 | 0 | 0 | 0 | 0 | 0 | 0 | 0 | 0 | 0.0001138 | 0 | 0.000835 | 0 | 0 | 0 | 0 | 0 | 0 | 0 |
| Oribacterium:K00548 | 0 | 0 | 0 | 0 | 0 | 0 | 0 | 0 | 0 | 0 | 0 | 0.0001138 | 0 | 0 | 0 | 0 | 0 | 0 | 0 | 0 | 0 |
| Oribacterium:K00640 | 0 | 0 | 0 | 0 | 0 | 0 | 0 | 0 | 0 | 0 | 0 | 0 | 0 | 0.000418 | 0 | 0 | 0 | 0 | 0 | 0 | 0 |
| Oribacterium:K01142 | 0 | 0 | 0 | 0 | 0 | 0 | 0 | 0 | 0 | 0 | 0.000477 | 0 | 0 | 0 | 0 | 0.000326 | 0 | 0 | 0 | 0 | 0 |
| Oribacterium:K01338 | 0 | 0 | 0 | 0 | 0 | 0 | 0 | 0 | 0 | 0 | 0 | 0.0002277 | 0 | 0.000418 | 0 | 0.000326 | 0 | 0 | 0 | 0 | 0 |
| Oribacterium:K01679 | 0 | 0 | 0 | 0 | 0 | 0 | 0 | 0 | 0 | 0 | 0 | 0 | 0 | 0 | 0 | 0.000109 | 0 | 0 | 0 | 0 | 0 |
| Oribacterium:K01915 | 0 | 0 | 0 | 0 | 0 | 0 | 0 | 0 | 0 | 0 | 0 | 0 | 0 | 0.000418 | 0.000104 | 0.000653 | 0.001974 | 0 | 0 | 0 | 0 |
| Oribacterium:K01962 | 0 | 0 | 0 | 0 | 0 | 0 | 0 | 0 | 0 | 0 | 0 | 0 | 0 | 0 | 0 | 0 | 0.000987 | 0 | 0 | 0 | 0 |
| Oribacterium:K02011 | 0 | 0 | 0 | 0 | 0 | 0 | 0 | 0 | 0 | 0 | 0.000477 | 0.0028459 | 0.001118 | 0 | 0 | 0.000653 | 0 | 0 | 0 | 0 | 0 |
| Oribacterium:K02012 | 0 | 0 | 0 | 0 | 0 | 0 | 0 | 0 | 0 | 0 | 0 | 0.0001138 | 0 | 0 | 0 | 0 | 0 | 0 | 0 | 0 | 0 |
| Oribacterium:K03338 | 0 | 0 | 0 | 0 | 0 | 0 | 0 | 0 | 0 | 0 | 0 | 0.0001138 | 0 | 0 | 0 | 0 | 0 | 0 | 0 | 0 | 0 |
| Oribacterium:K03644 | 0 | 0 | 0 | 0 | 0 | 0 | 0 | 0 | 0 | 0 | 0 | 0 | 0 | 0 | 0 | 0.000218 | 0 | 0 | 0 | 0 | 0 |
| Oscillibacter:K00161 | 0 | 0 | 0 | 0 | 0 | 0 | 0 | 0 | 0 | 0.0096337 | 0.000955 | 0.0089931 | 0.00447 | 0.003341 | 0.000311 | 0.001306 | 0 | 0 | 0 | 0.021473051 | 0 |
| Oscillibacter:K00162 | 0 | 0 | 0 | 0 | 0 | 4.15E-05 | 0 | 0 | 0 | 0.0171266 | 0 | 0.0149126 | 0.013411 | 0.005429 | 0.000934 | 0.00185 | 0 | 0 | 0 | 0 | 0 |
| Oscillibacter:K00248 | 0 | 0 | 0 | 0 | 0 | 0.00017 | 0 | 0 | 0 | 0.0706472 | 0.01575 | 0.032785 | 0.05141 | 0.009187 | 0.000727 | 0.005549 | 0 | 0 | 0 | 0 | 0 |
| Oscillibacter:K00253 | 0 | 0 | 0 | 0 | 0 | 0 | 0 | 0 | 0 | 0 | 0 | 0.0001138 | 0 | 0.000418 | 0 | 0 | 0 | 0 | 0 | 0 | 0 |
| Oscillibacter:K00257 | 0 | 0 | 0 | 0 | 0 | 0 | 0 | 0 | 0 | 0 | 0 | 0 | 0 | 0 | 0 | 0.000109 | 0 | 0 | 0 | 0 | 0 |
| Oscillibacter:K00334 | 0 | 0 | 0 | 0 | 0 | 0 | 0 | 0 | 0 | 0.0042816 | 0.004295 | 0.0036428 | 0.00447 | 0.003758 | 0.000104 | 0.001414 | 0 | 0 | 0 | 0 | 0 |
| Oscillibacter:K00335 | 0 | 0 | 0 | 0 | 0 | 4.15E-05 | 0 | 0 | 0 | 0.0149858 | 0.00525 | 0.0116113 | 0.013411 | 0.002923 | 0.000311 | 0.004461 | 0 | 0 | 0 | 0 | 0 |
| Oscillibacter:K00336 | 0 | 0.0205 | 0 | 0 | 0 | 8.29E-05 | 0 | 0 | 0 | 0.0385348 | 0.01575 | 0.0225397 | 0.041351 | 0.041758 | 0.00083 | 0.015014 | 0 | 0 | 0 | 0 | 0 |
| Oscillibacter:K00390 | 0 | 0 | 0 | 0.0016 | 0 | 0 | 0 | 0 | 0 | 0 | 0.000477 | 0.0001138 | 0 | 0.000835 | 0 | 0 | 0 | 0 | 0 | 0 | 0 |
| Oscillibacter:K00548 | 0 | 0 | 0 | 0 | 0 | 0 | 0 | 0 | 0 | 0 | 0 | 0 | 0 | 0 | 0 | 0.000109 | 0 | 0 | 0 | 0 | 0 |
| Oscillibacter:K00575 | 0 | 0 | 0 | 0 | 0 | 0 | 0 | 0 | 0 | 0 | 0 | 0.0002277 | 0.001118 | 0 | 0 | 0 | 0 | 0 | 0 | 0 | 0 |
| Oscillibacter:K00600 | 0 | 0 | 0 | 0 | 0 | 0.00017 | 0 | 0 | 0 | 0.0171266 | 0.002386 | 0.0067164 | 0.018999 | 0.001253 | 0.000623 | 0.007398 | 0 | 0 | 0 | 0.021473051 | 0 |
| Oscillibacter:K00626 | 0 | 0 | 0 | 0 | 0 | 4.15E-05 | 0 | 0 | 0 | 0.018197 | 0.007636 | 0.0124082 | 0.030175 | 0.020879 | 0.000727 | 0.006201 | 0 | 0 | 0 | 0.021473051 | 0 |
| Oscillibacter:K00627 | 0 | 0 | 0 | 0 | 0 | 0 | 0 | 0 | 0 | 0.0053521 | 0.000955 | 0.0029598 | 0.00447 | 0.003758 | 0.000415 | 0.000979 | 0 | 0 | 0 | 0 | 0 |
| Oscillibacter:K00640 | 0 | 0 | 0 | 0 | 0 | 0 | 0 | 0 | 0 | 0.0021408 | 0 | 0.0003415 | 0 | 0.001253 | 0 | 0.000435 | 0 | 0 | 0 | 0 | 0 |
| Oscillibacter:K00648 | 0 | 0 | 0 | 0 | 0 | 0 | 0 | 0 | 0 | 0.0010704 | 0.000477 | 0.0002277 | 0.001118 | 0.000418 | 0 | 0 | 0 | 0 | 0 | 0 | 0 |
| Oscillibacter:K00681 | 0 | 0 | 0 | 0 | 0 | 0 | 0 | 0 | 0 | 0.0021408 | 0 | 0.0007969 | 0.001118 | 0 | 0 | 0.000109 | 0 | 0 | 0 | 0 | 0 |
| Oscillibacter:K00798 | 0 | 0 | 0 | 0 | 0 | 0 | 0 | 0 | 0 | 0.0010704 | 0 | 0.0004553 | 0 | 0 | 0.000104 | 0 | 0 | 0 | 0 | 0 | 0 |
| Oscillibacter:K00833 | 0 | 0 | 0 | 0 | 0 | 0 | 0 | 0 | 0 | 0 | 0.000477 | 0.0001138 | 0 | 0 | 0 | 0 | 0.001974 | 0 | 0 | 0 | 0 |
| Oscillibacter:K00864 | 0 | 0 | 0 | 0 | 0 | 4.15E-05 | 0 | 0 | 0 | 0.0085633 | 0.004295 | 0.004895 | 0.006706 | 0.005429 | 0.000208 | 0.005331 | 0 | 0 | 0 | 0.021473051 | 0 |
| Oscillibacter:K01142 | 0 | 0 | 0 | 0 | 0 | 0 | 0 | 0 | 0 | 0.0074929 | 0.001432 | 0.0014799 | 0.005588 | 0 | 0.000415 | 0.001197 | 0 | 0 | 0 | 0 | 0 |
| Oscillibacter:K01338 | 0 | 0 | 0 | 0.0048 | 0 | 8.29E-05 | 0 | 0 | 0 | 0.0171266 | 0.010023 | 0.0081962 | 0.016764 | 0.006681 | 0.000208 | 0.002937 | 0.000987 | 0 | 0 | 0 | 0 |
| Oscillibacter:K01478 | 0 | 0 | 0 | 0 | 0 | 4.15E-05 | 0 | 0 | 0 | 0.0042816 | 0.006205 | 0.0064887 | 0.015646 | 0.007517 | 0.001246 | 0.00359 | 0.001974 | 0 | 0 | 0.021473051 | 0 |
| Oscillibacter:K01679 | 0 | 0 | 0 | 0 | 0 | 0 | 0 | 0 | 0 | 0 | 0 | 0.000683 | 0 | 0.000835 | 0 | 0.000109 | 0 | 0 | 0 | 0 | 0 |
| Oscillibacter:K01712 | 0 | 0 | 0 | 0.0081 | 0 | 0 | 0 | 0 | 0 | 0.0449573 | 0.018614 | 0.020946 | 0.015646 | 0.029231 | 0.002076 | 0.012511 | 0 | 0 | 0 | 0 | 0 |
| Oscillibacter:K01745 | 0 | 0 | 0 | 0 | 0 | 0 | 0 | 0 | 0 | 0.0171266 | 0.002386 | 0.0061472 | 0.006706 | 0.001253 | 0.000104 | 0.003808 | 0 | 0 | 0 | 0 | 0 |
| Oscillibacter:K01915 | 0 | 0.0409 | 0 | 0.0065 | 0 | 4.15E-05 | 0 | 0 | 0 | 0.0192674 | 0.007159 | 0.0105868 | 0.02794 | 0.011275 | 0.000415 | 0.007833 | 0.000987 | 0 | 0 | 0 | 0 |
| Oscillibacter:K01958 | 0 | 0 | 0 | 0.021 | 0 | 0 | 0 | 0 | 0 | 0.0139154 | 0.002864 | 0.0056918 | 0.016764 | 0.009187 | 0.002076 | 0.006745 | 0.000987 | 0 | 0 | 0 | 0 |
| Oscillibacter:K01960 | 0 | 0 | 0 | 0 | 0 | 0 | 0 | 0 | 0 | 0.0010704 | 0 | 0 | 0.001118 | 0 | 0 | 0.000109 | 0 | 0 | 0 | 0 | 0 |
| Oscillibacter:K01961 | 0 | 0 | 0 | 0 | 0 | 0 | 0 | 0 | 0 | 0 | 0 | 0.0007969 | 0 | 0 | 0 | 0.00087 | 0 | 0 | 0 | 0 | 0 |

| | | | | | | | | | | | | | | | | | | | | | |
|---|---|---|---|---|---|---|---|---|---|---|---|---|---|---|---|---|---|---|---|---|---|
| Oscillibacter:K01962 | 0 | 0 | 0 | 0 | 0 | 8.29E-05 | 0 | 0 | 0 | 0 | 0.000477 | 0.0001138 | 0.001118 | 0.001253 | 0.000208 | 0.00087 | 0 | 0 | 0 | 0 | 0 |
| Oscillibacter:K02011 | 0 | 0 | 0 | 0 | 0 | 0 | 0 | 0 | 0 | 0.0074929 | 0.006682 | 0.0050088 | 0.011176 | 0.002088 | 0.000104 | 0.001306 | 0 | 0 | 0 | 0 | 0 |
| Oscillibacter:K02012 | 0 | 0 | 0 | 0 | 0 | 0 | 0 | 0 | 0 | 0.0032112 | 0.000477 | 0.0031874 | 0.00447 | 0.002088 | 0.000415 | 0.001197 | 0 | 0 | 0 | 0.042946103 | 0 |
| Oscillibacter:K02049 | 0 | 0 | 0 | 0 | 0 | 8.29E-05 | 0 | 0 | 0 | 0.0021408 | 0.001909 | 0.0023906 | 0.00447 | 0.003341 | 0 | 0.000218 | 0 | 0 | 0 | 0 | 0 |
| Oscillibacter:K02051 | 0 | 0 | 0 | 0 | 0 | 0.00012 | 0 | 0 | 0 | 0.0203378 | 0.004295 | 0.0107007 | 0.021235 | 0.012528 | 0.000104 | 0.00359 | 0 | 0 | 0 | 0.021473051 | 0 |
| Oscillibacter:K02203 | 0 | 0 | 0 | 0 | 0 | 0 | 0 | 0 | 0 | 0.0021408 | 0.001909 | 0.0004553 | 0.001118 | 0 | 0 | 0 | 0 | 0 | 0 | 0 | 0 |
| Oscillibacter:K02400 | 0 | 0 | 0 | 0 | 0 | 0 | 0 | 0 | 0 | 0.0010704 | 0.000477 | 0.0012522 | 0.002235 | 0 | 0 | 0 | 0 | 0 | 0 | 0 | 0 |
| Oscillibacter:K02405 | 0 | 0 | 0 | 0 | 0 | 0 | 0 | 0 | 0 | 0 | 0 | 0.0011384 | 0 | 0 | 0 | 0 | 0 | 0 | 0 | 0 | 0 |
| Oscillibacter:K02409 | 0 | 0 | 0 | 0 | 0 | 0 | 0 | 0 | 0 | 0 | 0 | 0.0001138 | 0 | 0 | 0 | 0.000109 | 0 | 0 | 0 | 0 | 0 |
| Oscillibacter:K02410 | 0 | 0 | 0 | 0 | 0 | 0 | 0 | 0 | 0 | 0 | 0 | 0.0007969 | 0 | 0 | 0 | 0.000109 | 0.000987 | 0 | 0 | 0 | 0 |
| Oscillibacter:K02412 | 0 | 0 | 0 | 0 | 0 | 0 | 0 | 0 | 0 | 0 | 0 | 0.0009107 | 0 | 0.000418 | 0 | 0.000109 | 0 | 0 | 0 | 0 | 0 |
| Oscillibacter:K02416 | 0 | 0 | 0 | 0 | 0 | 0 | 0 | 0 | 0 | 0 | 0.000477 | 0.0003415 | 0.001118 | 0 | 0 | 0 | 0 | 0 | 0 | 0 | 0 |
| Oscillibacter:K02556 | 0 | 0 | 0 | 0 | 0 | 0 | 0 | 0 | 0 | 0.0021408 | 0.000477 | 0.0014799 | 0.001118 | 0.000835 | 0 | 0.000109 | 0 | 0 | 0 | 0 | 0 |
| Oscillibacter:K02557 | 0 | 0 | 0 | 0 | 0 | 0 | 0 | 0 | 0 | 0 | 0 | 0.0002277 | 0 | 0 | 0 | 0 | 0 | 0 | 0 | 0 | 0 |
| Oscillibacter:K03092 | 0 | 0 | 0 | 0 | 0 | 0 | 0 | 0 | 0 | 0.0042816 | 0.000477 | 0.000683 | 0.001118 | 0 | 0 | 0.000435 | 0 | 0 | 0 | 0.021473051 | 0.007893906 |
| Oscillibacter:K03407 | 0 | 0 | 0 | 0 | 0 | 0 | 0 | 0 | 0 | 0.0010704 | 0 | 0.0007969 | 0.001118 | 0.000418 | 0.000104 | 0.000435 | 0.000987 | 0 | 0 | 0 | 0 |
| Oscillibacter:K03412 | 0 | 0 | 0 | 0 | 0 | 0 | 0 | 0 | 0 | 0 | 0 | 0.0002277 | 0 | 0 | 0 | 0.000109 | 0 | 0 | 0 | 0 | 0 |
| Oscillibacter:K03415 | 0 | 0 | 0 | 0 | 0 | 0 | 0 | 0 | 0 | 0 | 0 | 0.000683 | 0 | 0 | 0 | 0 | 0 | 0 | 0 | 0 | 0 |
| Oscillibacter:K04564 | 0 | 0 | 0 | 0 | 0 | 0 | 0 | 0 | 0 | 0 | 0 | 0 | 0.001118 | 0.002506 | 0.000415 | 0.000762 | 0 | 0 | 0 | 0 | 0 |
| Oscillibacter:K05592 | 0 | 0 | 0 | 0 | 0 | 0 | 0 | 0 | 0 | 0 | 0 | 0.0001138 | 0 | 0 | 0 | 0 | 0 | 0 | 0 | 0 | 0 |
| Oscillibacter:K05846 | 0 | 0 | 0 | 0 | 0 | 0 | 0 | 0 | 0 | 0.0074929 | 0.001909 | 0.0014799 | 0.007823 | 0.005429 | 0.000311 | 0.006528 | 0.000987 | 0 | 0 | 0.021473051 | 0 |
| Oscillibacter:K05847 | 0 | 0 | 0 | 0 | 0 | 0 | 0 | 0 | 0 | 0 | 0.001432 | 0.0022767 | 0.002235 | 0.000835 | 0.000104 | 0.003264 | 0 | 0 | 0 | 0 | 0 |
| Oscillibacter:K06148 | 0 | 0 | 0 | 0 | 0 | 0 | 0 | 0 | 0 | 0 | 0 | 0.0002277 | 0 | 0 | 0.000104 | 0 | 0 | 0 | 0 | 0 | 0 |
| Oscillibacter:K10536 | 0 | 0 | 0 | 0 | 0 | 0 | 0 | 0 | 0 | 0.0074929 | 0.000955 | 0.0030736 | 0.007823 | 0.000418 | 0 | 0.005222 | 0 | 0 | 0 | 0 | 0 |
| Pedobacter:K00029 | 0 | 0 | 0 | 0.0194 | 0 | 4.15E-05 | 0 | 0 | 0 | 0 | 0 | 0 | 0 | 0 | 0 | 0 | 0 | 0 | 0 | 0 | 0 |
| Pedobacter:K00036 | 0 | 0 | 0 | 0 | 0 | 0 | 0.002207 | 0 | 0 | 0 | 0 | 0 | 0 | 0 | 0 | 0 | 0 | 0 | 0 | 0 | 0 |
| Pedobacter:K00116 | 0 | 0 | 0 | 0 | 0 | 4.15E-05 | 0 | 0 | 0 | 0 | 0 | 0 | 0 | 0 | 0 | 0 | 0 | 0 | 0 | 0 | 0 |
| Pedobacter:K00117 | 0 | 0 | 0 | 0.0016 | 0 | 0 | 0 | 0 | 0 | 0 | 0 | 0 | 0 | 0 | 0 | 0 | 0 | 0 | 0 | 0 | 0 |
| Pedobacter:K00128 | 0 | 0 | 0 | 0 | 0 | 4.15E-05 | 0 | 0 | 0 | 0 | 0 | 0 | 0 | 0 | 0 | 0 | 0 | 0 | 0 | 0 | 0 |
| Pedobacter:K00162 | 0 | 0.0205 | 0.0356 | 0.0113 | 0 | 4.15E-05 | 0 | 0 | 0 | 0 | 0 | 0 | 0 | 0 | 0 | 0 | 0 | 0 | 0 | 0 | 0 |
| Pedobacter:K00164 | 0 | 0 | 0 | 0 | 0 | 8.29E-05 | 0 | 0 | 0 | 0 | 0 | 0 | 0 | 0 | 0 | 0 | 0 | 0 | 0 | 0 | 0 |
| Pedobacter:K00252 | 0 | 0 | 0 | 0.0032 | 0 | 0.00012 | 0.002207 | 0.0007694 | 0 | 0 | 0 | 0 | 0 | 0 | 0 | 0 | 0 | 0 | 0 | 0 | 0 |
| Pedobacter:K00281 | 0 | 0 | 0 | 0.0065 | 0 | 4.15E-05 | 0 | 0 | 0 | 0 | 0 | 0 | 0 | 0 | 0 | 0 | 0 | 0 | 0 | 0 | 0 |
| Pedobacter:K00331 | 0 | 0 | 0 | 0 | 0 | 0 | 0.002207 | 0 | 0 | 0 | 0 | 0 | 0 | 0 | 0 | 0 | 0 | 0 | 0 | 0 | 0 |
| Pedobacter:K00334 | 0 | 0.0205 | 0 | 0.0097 | 0 | 4.15E-05 | 0 | 0 | 0 | 0 | 0 | 0 | 0 | 0 | 0 | 0 | 0 | 0 | 0 | 0 | 0 |
| Pedobacter:K00335 | 0 | 0 | 0.0356 | 0.0048 | 0 | 0 | 0 | 0.0007694 | 0 | 0 | 0 | 0 | 0 | 0 | 0 | 0 | 0 | 0 | 0 | 0 | 0 |
| Pedobacter:K00336 | 0 | 0 | 0 | 0.0032 | 0 | 4.15E-05 | 0 | 0 | 0 | 0 | 0 | 0 | 0 | 0 | 0 | 0 | 0 | 0 | 0 | 0 | 0 |
| Pedobacter:K00338 | 0 | 0 | 0 | 0 | 0 | 4.15E-05 | 0 | 0 | 0 | 0 | 0 | 0 | 0 | 0 | 0 | 0 | 0 | 0 | 0 | 0 | 0 |
| Pedobacter:K00341 | 0 | 0 | 0 | 0.0016 | 0 | 4.15E-05 | 0 | 0 | 0.001025 | 0 | 0 | 0 | 0 | 0 | 0 | 0 | 0 | 0 | 0 | 0 | 0 |
| Pedobacter:K00390 | 0 | 0 | 0 | 0.0016 | 0 | 0 | 0 | 0 | 0 | 0 | 0 | 0 | 0 | 0 | 0 | 0 | 0 | 0 | 0 | 0 | 0 |
| Pedobacter:K00425 | 0 | 0 | 0 | 0 | 0 | 0 | 0 | 0 | 0.001025 | 0 | 0 | 0 | 0 | 0 | 0 | 0 | 0 | 0 | 0 | 0 | 0 |
| Pedobacter:K00432 | 0 | 0 | 0 | 0 | 0 | 4.15E-05 | 0 | 0 | 0 | 0 | 0 | 0 | 0 | 0 | 0 | 0 | 0 | 0 | 0 | 0 | 0 |
| Pedobacter:K00451 | 0 | 0.0205 | 0 | 0.0048 | 0 | 0 | 0 | 0 | 0 | 0 | 0 | 0 | 0 | 0 | 0 | 0 | 0 | 0 | 0 | 0 | 0 |
| Pedobacter:K00457 | 0 | 0 | 0 | 0.0048 | 0 | 0 | 0 | 0 | 0 | 0 | 0 | 0 | 0 | 0 | 0 | 0 | 0 | 0 | 0 | 0 | 0 |
| Pedobacter:K00548 | 0 | 0 | 0 | 0 | 0 | 4.15E-05 | 0 | 0 | 0 | 0 | 0 | 0 | 0 | 0 | 0 | 0 | 0 | 0 | 0 | 0 | 0 |
| Pedobacter:K00575 | 0 | 0 | 0 | 0.0129 | 0 | 0.00012 | 0 | 0.0007694 | 0 | 0 | 0 | 0 | 0 | 0 | 0 | 0 | 0 | 0 | 0 | 0 | 0 |
| Pedobacter:K00600 | 0 | 0 | 0 | 0.0016 | 0 | 4.15E-05 | 0 | 0 | 0 | 0 | 0 | 0 | 0 | 0 | 0 | 0 | 0 | 0 | 0 | 0 | 0 |
| Pedobacter:K00627 | 0 | 0 | 0 | 0.0016 | 0 | 0 | 0 | 0 | 0 | 0 | 0 | 0 | 0 | 0 | 0 | 0 | 0 | 0 | 0 | 0 | 0 |
| Pedobacter:K00641 | 0 | 0 | 0 | 0 | 0 | 4.15E-05 | 0 | 0 | 0 | 0 | 0 | 0 | 0 | 0 | 0 | 0 | 0 | 0 | 0 | 0 | 0 |
| Pedobacter:K00648 | 0 | 0 | 0 | 0.0065 | 0 | 8.29E-05 | 0 | 0 | 0.001025 | 0 | 0 | 0 | 0 | 0 | 0 | 0 | 0 | 0 | 0 | 0 | 0 |
| Pedobacter:K00658 | 0 | 0 | 0 | 0.0016 | 0 | 0 | 0.002207 | 0 | 0 | 0 | 0 | 0 | 0 | 0 | 0 | 0 | 0 | 0 | 0 | 0 | 0 |
| Pedobacter:K00677 | 0 | 0 | 0 | 0.0016 | 0 | 0 | 0 | 0 | 0 | 0 | 0 | 0 | 0 | 0 | 0 | 0 | 0 | 0 | 0 | 0 | 0 |
| Pedobacter:K00681 | 0 | 0 | 0 | 0.0016 | 0 | 0 | 0 | 0 | 0 | 0 | 0 | 0 | 0 | 0 | 0 | 0 | 0 | 0 | 0 | 0 | 0 |
| Pedobacter:K00748 | 0 | 0 | 0 | 0 | 0 | 4.15E-05 | 0 | 0 | 0 | 0 | 0 | 0 | 0 | 0 | 0 | 0 | 0 | 0 | 0 | 0 | 0 |
| Pedobacter:K00940 | 0 | 0 | 0 | 0 | 0 | 4.15E-05 | 0 | 0 | 0 | 0 | 0 | 0 | 0 | 0 | 0 | 0 | 0 | 0 | 0 | 0 | 0 |
| Pedobacter:K01241 | 0 | 0 | 0.0356 | 0.0016 | 0 | 0 | 0 | 0 | 0 | 0 | 0 | 0 | 0 | 0 | 0 | 0 | 0 | 0 | 0 | 0 | 0 |
| Pedobacter:K01251 | 0 | 0 | 0 | 0.0016 | 0 | 0 | 0 | 0 | 0 | 0 | 0 | 0 | 0 | 0 | 0 | 0 | 0 | 0 | 0 | 0 | 0 |
| Pedobacter:K01338 | 0 | 0.0205 | 0 | 0.0048 | 0 | 4.15E-05 | 0 | 0 | 0.001025 | 0.0010704 | 0.000477 | 0 | 0 | 0 | 0 | 0 | 0 | 0 | 0 | 0 | 0 |
| Pedobacter:K01585 | 0 | 0.1433 | 0.1782 | 0.0905 | 0 | 0.00012 | 0 | 0 | 0 | 0 | 0 | 0 | 0 | 0 | 0 | 0 | 0 | 0 | 0 | 0 | 0 |
| Pedobacter:K01627 | 0 | 0.0205 | 0 | 0.0065 | 0 | 0 | 0 | 0 | 0 | 0 | 0 | 0 | 0 | 0 | 0 | 0 | 0 | 0 | 0 | 0 | 0 |

| | | | | | | | | | | | | | | | | | | | | | |
|---|---|---|---|---|---|---|---|---|---|---|---|---|---|---|---|---|---|---|---|---|---|
| Pedobacter:K01679 | 0 | 0 | 0 | 0 | 0 | 4.15E-05 | 0 | 0 | 0 | 0 | 0 | 0 | 0 | 0 | 0 | 0 | 0 | 0 | 0 | 0 | 0 |
| Pedobacter:K01895 | 0 | 0 | 0 | 0.0032 | 0 | 4.15E-05 | 0 | 0 | 0.001025 | 0 | 0 | 0 | 0 | 0 | 0 | 0 | 0 | 0 | 0 | 0 | 0 |
| Pedobacter:K01902 | 0 | 0.0614 | 0 | 0.0145 | 0 | 4.15E-05 | 0 | 0 | 0 | 0 | 0 | 0 | 0 | 0 | 0 | 0 | 0 | 0 | 0 | 0 | 0 |
| Pedobacter:K01903 | 0 | 0 | 0 | 0.0032 | 0 | 4.15E-05 | 0 | 0 | 0 | 0 | 0 | 0 | 0 | 0 | 0 | 0 | 0 | 0 | 0 | 0 | 0 |
| Pedobacter:K01915 | 0 | 0 | 0 | 0.0016 | 0 | 0 | 0.002207 | 0 | 0 | 0 | 0 | 0 | 0 | 0 | 0 | 0 | 0 | 0 | 0 | 0 | 0 |
| Pedobacter:K01961 | 0 | 0.0205 | 0 | 0.0129 | 0 | 8.29E-05 | 0 | 0.0023082 | 0 | 0 | 0 | 0 | 0 | 0 | 0 | 0 | 0 | 0 | 0 | 0 | 0 |
| Pedobacter:K01962 | 0 | 0 | 0 | 0.0032 | 0 | 8.29E-05 | 0 | 0 | 0 | 0 | 0 | 0 | 0 | 0 | 0 | 0 | 0 | 0 | 0 | 0 | 0 |
| Pedobacter:K02274 | 0 | 0 | 0 | 0 | 0 | 4.15E-05 | 0 | 0 | 0 | 0 | 0 | 0 | 0 | 0 | 0 | 0 | 0 | 0 | 0 | 0 | 0 |
| Pedobacter:K02276 | 0 | 0 | 0 | 0 | 0 | 0 | 0 | 0.0007694 | 0 | 0 | 0 | 0 | 0 | 0 | 0 | 0 | 0 | 0 | 0 | 0 | 0 |
| Pedobacter:K02492 | 0 | 0 | 0 | 0 | 0 | 8.29E-05 | 0 | 0 | 0 | 0 | 0 | 0 | 0 | 0 | 0 | 0 | 0 | 0 | 0 | 0 | 0 |
| Pedobacter:K03092 | 0 | 0 | 0 | 0.0032 | 0 | 0 | 0 | 0 | 0.001025 | 0 | 0 | 0 | 0 | 0 | 0 | 0 | 0 | 0 | 0 | 0 | 0 |
| Pedobacter:K03182 | 0 | 0 | 0 | 0.0016 | 0 | 8.29E-05 | 0 | 0 | 0 | 0 | 0 | 0 | 0 | 0 | 0 | 0 | 0 | 0 | 0 | 0 | 0 |
| Pedobacter:K03412 | 0 | 0 | 0 | 0 | 0 | 4.15E-05 | 0 | 0 | 0 | 0 | 0 | 0 | 0 | 0 | 0 | 0 | 0 | 0 | 0 | 0 | 0 |
| Pedobacter:K03590 | 0 | 0 | 0 | 0 | 0 | 4.15E-05 | 0 | 0 | 0 | 0 | 0 | 0 | 0 | 0 | 0 | 0 | 0 | 0 | 0 | 0 | 0 |
| Pedobacter:K03644 | 0 | 0 | 0.0356 | 0.0178 | 0 | 4.15E-05 | 0 | 0 | 0 | 0 | 0 | 0 | 0 | 0 | 0 | 0 | 0 | 0 | 0 | 0 | 0 |
| Pedobacter:K03841 | 0 | 0.0409 | 0.0356 | 0.0032 | 0 | 4.15E-05 | 0 | 0 | 0 | 0 | 0 | 0 | 0 | 0 | 0 | 0 | 0 | 0 | 0 | 0 | 0 |
| Pedobacter:K04564 | 0 | 0 | 0 | 0.0016 | 0 | 0 | 0 | 0 | 0.001025 | 0 | 0 | 0 | 0 | 0 | 0 | 0 | 0 | 0 | 0 | 0 | 0 |
| Pedobacter:K05592 | 0 | 0 | 0 | 0 | 0 | 0 | 0 | 0.0007694 | 0 | 0 | 0 | 0 | 0 | 0 | 0 | 0 | 0 | 0 | 0 | 0 | 0 |
| Pedobacter:K06861 | 0 | 0 | 0 | 0.0016 | 0 | 0 | 0 | 0 | 0 | 0 | 0 | 0 | 0 | 0 | 0 | 0 | 0 | 0 | 0 | 0 | 0 |
| Pedobacter:K07516 | 0 | 0.0205 | 0 | 0.0113 | 0 | 4.15E-05 | 0 | 0 | 0 | 0 | 0 | 0 | 0 | 0 | 0 | 0 | 0 | 0 | 0 | 0 | 0 |
| Pedobacter:K11927 | 0 | 0 | 0 | 0 | 0 | 0.00017 | 0 | 0 | 0 | 0 | 0 | 0 | 0 | 0 | 0 | 0 | 0 | 0 | 0 | 0 | 0 |
| Pedobacter:K12339 | 0 | 0.0205 | 0 | 0.0065 | 0 | 0 | 0 | 0.0007694 | 0 | 0 | 0 | 0 | 0 | 0 | 0 | 0 | 0 | 0 | 0 | 0 | 0 |
| Pedobacter:K12573 | 0 | 0 | 0 | 0.0032 | 0 | 0.00012 | 0 | 0 | 0 | 0 | 0 | 0 | 0 | 0 | 0 | 0 | 0 | 0 | 0 | 0 | 0 |
| Prevotella:K00029 | 0 | 0 | 0 | 0.0032 | 0 | 0.00012 | 0.002207 | 0 | 0 | 0.0503093 | 0.039614 | 0.1095109 | 0.07488 | 0.000418 | 0.007266 | 0.002176 | 0 | 0.0373413 | 0 | 0.042946103 | 0 |
| Prevotella:K00111 | 0 | 0 | 0 | 0 | 0 | 0 | 0 | 0 | 0 | 0.0032112 | 0.000955 | 0.001366 | 0 | 0 | 0 | 0 | 0 | 0 | 0 | 0 | 0 |
| Prevotella:K00257 | 0 | 0 | 0 | 0.0016 | 0 | 0 | 0 | 0 | 0 | 0 | 0 | 0 | 0 | 0 | 0 | 0 | 0 | 0 | 0 | 0 | 0 |
| Prevotella:K00331 | 0 | 0 | 0 | 0 | 0 | 4.15E-05 | 0 | 0 | 0 | 0.0321124 | 0.023864 | 0.0445102 | 0.022352 | 0 | 0.006021 | 0.001088 | 0 | 0 | 0 | 0 | 0 |
| Prevotella:K00337 | 0 | 0 | 0 | 0 | 0 | 0.00012 | 0 | 0.0007694 | 0 | 0.0481685 | 0.084 | 0.0819625 | 0.058115 | 0.000835 | 0.012871 | 0.002829 | 0 | 0.0373413 | 0 | 0.042946103 | 0.007893906 |
| Prevotella:K00338 | 0 | 0 | 0 | 0 | 0 | 4.15E-05 | 0 | 0 | 0 | 0.0096337 | 0.040091 | 0.029256 | 0.014529 | 0 | 0.00083 | 0.000762 | 0 | 0 | 0 | 0.021473051 | 0 |
| Prevotella:K00341 | 0 | 0 | 0 | 0 | 0 | 0.00046 | 0 | 0 | 0 | 0.0995483 | 0.140796 | 0.1955715 | 0.112878 | 0.00167 | 0.013494 | 0.000762 | 0.000987 | 0 | 0 | 0.021473051 | 0 |
| Prevotella:K00342 | 0 | 0.0205 | 0.0356 | 0 | 0 | 8.29E-05 | 0 | 0 | 0 | 0.0331828 | 0.077318 | 0.1134952 | 0.043587 | 0.00167 | 0.004048 | 0.005875 | 0 | 0 | 0 | 0.042946103 | 0.015787812 |
| Prevotella:K00390 | 0 | 0 | 0 | 0 | 0 | 0 | 0 | 0 | 0 | 0.0042816 | 0.00525 | 0.004212 | 0.00447 | 0 | 0 | 0 | 0 | 0.074682599 | 0 | 0.042946103 | 0.031575624 |
| Prevotella:K00425 | 0 | 0 | 0 | 0 | 0 | 0.00017 | 0 | 0 | 0 | 0.0941962 | 0.044387 | 0.0817348 | 0.070409 | 0.000835 | 0.009861 | 0.005113 | 0 | 0.373412995 | 0 | 0.107365257 | 0.236817177 |
| Prevotella:K00426 | 0 | 0 | 0 | 0.0016 | 0 | 0 | 0 | 0 | 0 | 0.0331828 | 0.028636 | 0.0410951 | 0.024587 | 0.000418 | 0.003218 | 0.000979 | 0 | 0.0373413 | 0 | 0.085892205 | 0.015787812 |
| Prevotella:K00432 | 0 | 0 | 0 | 0 | 0 | 8.29E-05 | 0 | 0 | 0 | 0.0299715 | 0.028159 | 0.0387045 | 0.026823 | 0.000418 | 0.008616 | 0.003155 | 0 | 0.0373413 | 0 | 0.085892205 | 0.031575624 |
| Prevotella:K00525 | 0 | 0 | 0 | 0 | 0 | 0 | 0 | 0 | 0.001025 | 0.0128449 | 0.013841 | 0.0716033 | 0.022352 | 0.003341 | 0.000727 | 0.004025 | 0.000987 | 0 | 0 | 0.021473051 | 0 |
| Prevotella:K00526 | 0 | 0 | 0 | 0 | 0 | 0 | 0 | 0 | 0 | 0 | 0.000477 | 0 | 0.001118 | 0.000835 | 0 | 0.000109 | 0.000987 | 0 | 0 | 0 | 0 |
| Prevotella:K00548 | 0 | 0 | 0 | 0 | 0 | 4.15E-05 | 0 | 0 | 0 | 0.036394 | 0.075409 | 0.1226022 | 0.031293 | 0 | 0.001246 | 0.006093 | 0 | 0 | 0 | 0.064419154 | 0 |
| Prevotella:K00600 | 0 | 0 | 0 | 0.0016 | 0 | 0.00017 | 0 | 0 | 0 | 0.036394 | 0.07875 | 0.0907279 | 0.037999 | 0 | 0.001453 | 0.004461 | 0 | 0 | 0 | 0.085892205 | 0 |
| Prevotella:K00640 | 0 | 0 | 0 | 0 | 0 | 0 | 0 | 0 | 0 | 0.0246195 | 0.044387 | 0.0270931 | 0.005588 | 0.00167 | 0.000727 | 0.000435 | 0 | 0 | 0 | 0.021473051 | 0.023681718 |
| Prevotella:K00641 | 0 | 0 | 0 | 0 | 0 | 0 | 0 | 0 | 0 | 0 | 0.000955 | 0 | 0 | 0 | 0 | 0 | 0 | 0 | 0 | 0 | 0 |
| Prevotella:K00647 | 0 | 0 | 0 | 0.0016 | 0 | 0 | 0 | 0 | 0 | 0.0042816 | 0.003818 | 0.0034151 | 0.003353 | 0 | 0 | 0 | 0 | 0 | 0 | 0.085892205 | 0 |
| Prevotella:K00648 | 0 | 0 | 0 | 0.0016 | 0 | 0.00012 | 0 | 0.0007694 | 0 | 0.0235491 | 0.076841 | 0.0702373 | 0.017882 | 0.005846 | 0.001765 | 0.008812 | 0 | 0 | 0 | 0.064419154 | 0 |
| Prevotella:K00677 | 0 | 0 | 0 | 0.0016 | 0 | 4.15E-05 | 0 | 0 | 0 | 0.0246195 | 0.068728 | 0.0509989 | 0.032411 | 0 | 0.002076 | 0.00359 | 0 | 0 | 0 | 0.021473051 | 0 |
| Prevotella:K00748 | 0 | 0 | 0 | 0.0016 | 0 | 0 | 0 | 0 | 0 | 0.0149858 | 0.013841 | 0.0297114 | 0.02347 | 0 | 0.005502 | 0.000979 | 0 | 0 | 0 | 0 | 0 |
| Prevotella:K00864 | 0 | 0 | 0 | 0 | 0 | 0 | 0 | 0 | 0 | 0.0053521 | 0.011455 | 0.0036428 | 0.00447 | 0 | 0 | 0 | 0.000987 | 0 | 0 | 0.021473051 | 0 |
| Prevotella:K00940 | 0 | 0 | 0 | 0 | 0 | 0 | 0 | 0 | 0 | 0 | 0.001909 | 0 | 0 | 0 | 0 | 0 | 0 | 0 | 0 | 0 | 0 |
| Prevotella:K00955 | 0 | 0 | 0 | 0 | 0 | 0 | 0 | 0 | 0 | 0 | 0 | 0.0001138 | 0 | 0 | 0 | 0 | 0 | 0 | 0 | 0 | 0 |
| Prevotella:K01142 | 0 | 0.0409 | 0 | 0.0097 | 0 | 0.00021 | 0 | 0 | 0 | 0.0385348 | 0.0315 | 0.0616995 | 0.05141 | 0.003758 | 0.004879 | 0.003481 | 0 | 0.074682599 | 0 | 0.042946103 | 0.023681718 |
| Prevotella:K01338 | 0 | 0 | 0 | 0.0081 | 0 | 8.29E-05 | 0 | 0 | 0.001025 | 0.0759992 | 0.079705 | 0.1544765 | 0.082703 | 0.000418 | 0.007681 | 0.011424 | 0 | 0.0373413 | 0.003471861 | 0.128838308 | 0.023681718 |
| Prevotella:K01433 | 0 | 0 | 0 | 0.0016 | 0 | 4.15E-05 | 0 | 0 | 0 | 0.0021408 | 0.048682 | 0.0109283 | 0.002235 | 0 | 0 | 0.001523 | 0 | 0 | 0 | 0 | 0 |
| Prevotella:K01585 | 0 | 0 | 0 | 0.0016 | 0 | 0.00021 | 0 | 0 | 0 | 0.1102524 | 0.201887 | 0.2003527 | 0.129642 | 0.006681 | 0.016608 | 0.013273 | 0 | 0.074682599 | 0 | 0.042946103 | 0.023681718 |
| Prevotella:K01613 | 0 | 0 | 0 | 0.0016 | 0 | 0 | 0 | 0 | 0 | 0.0107041 | 0.042477 | 0.0308498 | 0.007823 | 0 | 0.000311 | 0.001958 | 0 | 0 | 0 | 0 | 0 |
| Prevotella:K01627 | 0 | 0 | 0 | 0 | 0 | 4.15E-05 | 0 | 0 | 0 | 0.0107041 | 0.022432 | 0.0217428 | 0.020117 | 0.00167 | 0.000104 | 0.000544 | 0 | 0 | 0 | 0.021473051 | 0 |
| Prevotella:K01629 | 0 | 0.0205 | 0 | 0.0032 | 0 | 0 | 0 | 0 | 0 | 0.0042816 | 0.013841 | 0.0162787 | 0.005588 | 0 | 0 | 0.000653 | 0 | 0 | 0 | 0 | 0 |
| Prevotella:K01676 | 0 | 0.0205 | 0 | 0.0065 | 0 | 0.00017 | 0 | 0.0007694 | 0 | 0.1027595 | 0.141751 | 0.1751947 | 0.112878 | 0.001253 | 0.018062 | 0.003264 | 0 | 0 | 0 | 0 | 0 |
| Prevotella:K01698 | 0 | 0 | 0 | 0 | 0 | 0 | 0 | 0 | 0 | 0.0021408 | 0.000477 | 0.0029598 | 0.001118 | 0 | 0 | 0 | 0 | 0 | 0 | 0 | 0 |
| Prevotella:K01712 | 0 | 0.0819 | 0 | 0.021 | 0 | 8.29E-05 | 0 | 0.0007694 | 0 | 0.0074929 | 0.017659 | 0.0399567 | 0.020117 | 0.000418 | 0.000104 | 0.002394 | 0 | 0 | 0 | 0 | 0 |
| Prevotella:K01737 | 0 | 0 | 0 | 0 | 0 | 0 | 0 | 0 | 0 | 0 | 0.000477 | 0.000683 | 0 | 0 | 0 | 0 | 0 | 0 | 0 | 0 | 0 |
| Prevotella:K01745 | 0 | 0 | 0 | 0.0016 | 0 | 4.15E-05 | 0 | 0 | 0 | 0.0074929 | 0.010023 | 0.016734 | 0.001118 | 0 | 0 | 0.00087 | 0 | 0 | 0 | 0 | 0 |

| | | | | | | | | | | | | | | | | | | | | | |
|---|---|---|---|---|---|---|---|---|---|---|---|---|---|---|---|---|---|---|---|---|---|
| Prevotella:K01845 | 0 | 0 | 0 | 0 | 0 | 0 | 0 | 0 | 0 | 0.0010704 | 0.000477 | 0.0036428 | 0 | 0 | 0 | 0 | 0 | 0 | 0 | 0 | 0 |
| Prevotella:K01895 | 0 | 0 | 0 | 0 | 0 | 0.00029 | 0 | 0 | 0 | 0.0203378 | 0.144137 | 0.1327336 | 0.029058 | 0.004593 | 0.001765 | 0.010336 | 0.000987 | 0 | 0 | 0.064419154 | 0 |
| Prevotella:K01902 | 0 | 0 | 0 | 0 | 0 | 0 | 0 | 0 | 0 | 0 | 0 | 0 | 0.001118 | 0 | 0 | 0 | 0 | 0 | 0.003471861 | 0 | 0 |
| Prevotella:K01903 | 0 | 0 | 0 | 0 | 0 | 0 | 0 | 0 | 0 | 0 | 0 | 0 | 0.002235 | 0 | 0 | 0 | 0 | 0 | 0 | 0 | 0 |
| Prevotella:K01915 | 0 | 0 | 0 | 0.0016 | 0 | 0.00041 | 0 | 0 | 0.001025 | 0.1916037 | 0.272524 | 0.2426999 | 0.207875 | 0.001253 | 0.02429 | 0.021324 | 0.000987 | 0.0373413 | 0 | 0.128838308 | 0.015787812 |
| Prevotella:K01960 | 0 | 0 | 0 | 0.0032 | 0 | 0.00029 | 0 | 0 | 0 | 0.0492389 | 0.055841 | 0.1015424 | 0.05588 | 0.002088 | 0.003737 | 0.005766 | 0 | 0 | 0 | 0.064419154 | 0 |
| Prevotella:K02047 | 0 | 0 | 0 | 0 | 0 | 0 | 0 | 0 | 0 | 0 | 0 | 0 | 0 | 0 | 0 | 0 | 0 | 0 | 0 | 0.021473051 | 0 |
| Prevotella:K02051 | 0 | 0 | 0 | 0 | 0 | 0 | 0 | 0 | 0 | 0.0010704 | 0.002864 | 0.0130912 | 0.002235 | 0 | 0.000311 | 0.000435 | 0 | 0.0373413 | 0 | 0 | 0 |
| Prevotella:K02517 | 0 | 0 | 0 | 0 | 0 | 0 | 0 | 0 | 0 | 0.0139154 | 0.025773 | 0.0225397 | 0.014529 | 0.000418 | 0 | 0.000109 | 0 | 0 | 0 | 0.085892205 | 0 |
| Prevotella:K02523 | 0 | 0 | 0 | 0 | 0 | 4.15E-05 | 0 | 0 | 0 | 0.0053521 | 0.012409 | 0.027207 | 0.003353 | 0 | 0.000727 | 0.000653 | 0 | 0 | 0 | 0.021473051 | 0 |
| Prevotella:K02535 | 0 | 0 | 0 | 0 | 0 | 0 | 0 | 0 | 0 | 0 | 0 | 0.0001138 | 0 | 0 | 0 | 0 | 0 | 0 | 0 | 0 | 0 |
| Prevotella:K02557 | 0 | 0 | 0 | 0 | 0 | 0.00012 | 0 | 0 | 0 | 0.036394 | 0.021955 | 0.0772951 | 0.048057 | 0 | 0.008408 | 0.000435 | 0 | 0 | 0 | 0.021473051 | 0.031575624 |
| Prevotella:K03072 | 0 | 0 | 0 | 0 | 0 | 0 | 0 | 0 | 0 | 0 | 0.001909 | 0.0003415 | 0 | 0 | 0 | 0 | 0 | 0 | 0 | 0 | 0 |
| Prevotella:K03092 | 0 | 0 | 0 | 0 | 0 | 0.00012 | 0 | 0.0007694 | 0 | 0.0428165 | 0.035318 | 0.0657976 | 0.029058 | 0 | 0.009965 | 0.005331 | 0 | 0.112023898 | 0 | 0.042946103 | 0.078939059 |
| Prevotella:K03183 | 0 | 0 | 0 | 0 | 0 | 0.00012 | 0 | 0 | 0 | 0.0203378 | 0.039614 | 0.0449655 | 0.018999 | 0 | 0.000311 | 0.000435 | 0 | 0 | 0 | 0.042946103 | 0 |
| Prevotella:K03590 | 0 | 0 | 0 | 0 | 0 | 0 | 0 | 0 | 0 | 0.0235491 | 0.019091 | 0.0589674 | 0.013411 | 0.002506 | 0.001038 | 0.004787 | 0 | 0 | 0 | 0.064419154 | 0 |
| Prevotella:K03644 | 0 | 0 | 0 | 0 | 0 | 0 | 0 | 0 | 0 | 0 | 0 | 0 | 0.002235 | 0 | 0 | 0.001414 | 0 | 0 | 0 | 0 | 0 |
| Prevotella:K03885 | 0 | 0 | 0 | 0 | 0 | 0 | 0 | 0 | 0 | 0.0074929 | 0.001432 | 0.0030736 | 0.002235 | 0 | 0.001038 | 0.000435 | 0 | 0 | 0 | 0 | 0 |
| Prevotella:K05343 | 0 | 0 | 0 | 0 | 0 | 4.15E-05 | 0 | 0 | 0 | 0.0235491 | 0.021477 | 0.0178724 | 0.033528 | 0 | 0 | 0.000218 | 0 | 0 | 0 | 0 | 0 |
| Prevotella:K05592 | 0 | 0 | 0 | 0 | 0 | 0.00017 | 0 | 0 | 0 | 0.0417461 | 0.035318 | 0.0869713 | 0.048057 | 0.00167 | 0.00519 | 0.004569 | 0 | 0.0373413 | 0 | 0.042946103 | 0.007893906 |
| Prevotella:K06861 | 0 | 0 | 0 | 0 | 0 | 0.00012 | 0 | 0 | 0.001025 | 0.0406756 | 0.054887 | 0.0517957 | 0.034646 | 0.000418 | 0.015467 | 0.003155 | 0 | 0 | 0 | 0 | 0.007893906 |
| Prevotella:K12573 | 0 | 0 | 0 | 0.0016 | 0 | 0.00012 | 0 | 0 | 0 | 0.0342532 | 0.060614 | 0.1153166 | 0.025705 | 0.000418 | 0.002699 | 0.006093 | 0 | 0 | 0 | 0.021473051 | 0 |
| Prevotella:K13378 | 0 | 0 | 0 | 0 | 0 | 0.00012 | 0 | 0 | 0 | 0.0267603 | 0.095455 | 0.1092833 | 0.044704 | 0 | 0.00519 | 0.001632 | 0 | 0.0373413 | 0 | 0.021473051 | 0 |
| Ruminococcus:K00029 | 0 | 0 | 0 | 0 | 0 | 0 | 0 | 0 | 0 | 0 | 0 | 0.0001138 | 0 | 0 | 0 | 0 | 0 | 0 | 0 | 0 | 0 |
| Ruminococcus:K00111 | 0 | 0 | 0 | 0 | 0 | 0 | 0 | 0 | 0 | 0.0010704 | 0.005727 | 0.0096761 | 0.007823 | 0.000835 | 0.000519 | 0.000979 | 0.000987 | 0 | 0 | 0 | 0 |
| Ruminococcus:K00128 | 0 | 0 | 0 | 0 | 0 | 0 | 0 | 0 | 0 | 0 | 0 | 0.0004553 | 0 | 0 | 0 | 0 | 0.003949 | 0 | 0 | 0 | 0 |
| Ruminococcus:K00303 | 0 | 0 | 0 | 0 | 0 | 0 | 0 | 0 | 0 | 0.0010704 | 0 | 0.0002277 | 0 | 0 | 0 | 0 | 0.000987 | 0 | 0 | 0 | 0 |
| Ruminococcus:K00334 | 0 | 0.0205 | 0 | 0 | 0 | 0 | 0 | 0 | 0 | 0.0021408 | 0.000955 | 0.0035289 | 0.002235 | 0.002506 | 0 | 0.011206 | 0 | 0 | 0 | 0 | 0 |
| Ruminococcus:K00335 | 0 | 0 | 0 | 0.0065 | 0 | 0.00012 | 0 | 0 | 0 | 0.0021408 | 0.001432 | 0.0022767 | 0 | 0.002088 | 0.00083 | 0.006201 | 0.001974 | 0 | 0 | 0 | 0 |
| Ruminococcus:K00336 | 0 | 0 | 0.0356 | 0.0065 | 0 | 0 | 0 | 0 | 0 | 0.0021408 | 0.000477 | 0.001366 | 0.002235 | 0.000418 | 0.000104 | 0.014579 | 0.000987 | 0 | 0 | 0 | 0 |
| Ruminococcus:K00390 | 0 | 0 | 0 | 0 | 0 | 0 | 0 | 0 | 0 | 0.0021408 | 0.000477 | 0.0002277 | 0 | 0 | 0 | 0 | 0 | 0 | 0 | 0 | 0 |
| Ruminococcus:K00432 | 0 | 0 | 0 | 0.0016 | 0 | 4.15E-05 | 0 | 0 | 0 | 0 | 0 | 0 | 0 | 0 | 0 | 0 | 0 | 0 | 0 | 0 | 0 |
| Ruminococcus:K00526 | 0 | 0 | 0 | 0 | 0 | 0 | 0 | 0 | 0 | 0 | 0 | 0 | 0 | 0.001253 | 0.000415 | 0.002611 | 0 | 0 | 0 | 0 | 0 |
| Ruminococcus:K00548 | 0 | 0 | 0 | 0.0016 | 0 | 4.15E-05 | 0 | 0 | 0 | 0.0203378 | 0.032455 | 0.0086516 | 0.008941 | 0.014198 | 0.012768 | 0.037861 | 0.040474 | 0 | 0 | 0.021473051 | 0 |
| Ruminococcus:K00575 | 0 | 0 | 0 | 0 | 0 | 0 | 0 | 0 | 0 | 0 | 0 | 0 | 0 | 0 | 0 | 0.002937 | 0 | 0 | 0 | 0 | 0 |
| Ruminococcus:K00600 | 0 | 0 | 0.0356 | 0.0048 | 0 | 0.00012 | 0 | 0 | 0 | 0.0235491 | 0.037227 | 0.0085378 | 0.010058 | 0.005846 | 0.014117 | 0.041886 | 0.019744 | 0 | 0 | 0.042946103 | 0 |
| Ruminococcus:K00640 | 0 | 0 | 0 | 0.0032 | 0 | 8.29E-05 | 0 | 0 | 0 | 0.0021408 | 0.000955 | 0.0030736 | 0.001118 | 0.012528 | 0.000623 | 0.023935 | 0.004936 | 0 | 0 | 0 | 0 |
| Ruminococcus:K00648 | 0 | 0 | 0 | 0 | 0 | 0 | 0 | 0 | 0 | 0 | 0.000955 | 0.000683 | 0.001118 | 0.000835 | 0.00083 | 0.008268 | 0 | 0 | 0 | 0 | 0 |
| Ruminococcus:K00798 | 0 | 0 | 0 | 0 | 0 | 0 | 0 | 0 | 0 | 0.0032112 | 0.012886 | 0.0004553 | 0 | 0.002506 | 0.001038 | 0.001414 | 0.004936 | 0 | 0 | 0 | 0 |
| Ruminococcus:K00833 | 0 | 0 | 0 | 0 | 0 | 0 | 0 | 0 | 0 | 0 | 0 | 0.000683 | 0.001118 | 0.001253 | 0.000208 | 0.000762 | 0.000987 | 0 | 0 | 0 | 0 |
| Ruminococcus:K00836 | 0 | 0 | 0 | 0 | 0 | 4.15E-05 | 0 | 0 | 0 | 0.0010704 | 0 | 0.0001138 | 0 | 0.00167 | 0 | 0 | 0 | 0 | 0 | 0 | 0 |
| Ruminococcus:K00864 | 0 | 0 | 0 | 0 | 0 | 0 | 0 | 0 | 0 | 0.0010704 | 0.000477 | 0.0009107 | 0 | 0.000835 | 0.001453 | 0.024588 | 0.00691 | 0 | 0 | 0 | 0 |
| Ruminococcus:K00955 | 0 | 0 | 0 | 0 | 0 | 0 | 0 | 0 | 0 | 0 | 0 | 0 | 0 | 0 | 0 | 0.000326 | 0 | 0 | 0 | 0 | 0 |
| Ruminococcus:K01142 | 0 | 0 | 0 | 0.0048 | 0 | 0 | 0 | 0 | 0 | 0.0053521 | 0.010023 | 0.001366 | 0.006706 | 0.000835 | 0.000208 | 0 | 0.000987 | 0 | 0 | 0 | 0 |
| Ruminococcus:K01251 | 0 | 0 | 0 | 0.0016 | 0 | 0.00017 | 0 | 0 | 0 | 0.0085633 | 0.006682 | 0.0034151 | 0.00447 | 0.002506 | 0.000104 | 0.035794 | 0.000987 | 0 | 0 | 0 | 0 |
| Ruminococcus:K01338 | 0 | 0 | 0 | 0 | 0 | 0.00017 | 0 | 0 | 0 | 0.0224786 | 0.071114 | 0.0240196 | 0.02347 | 0.017121 | 0.028338 | 0.023391 | 0.030602 | 0 | 0 | 0 | 0 |
| Ruminococcus:K01426 | 0 | 0 | 0 | 0 | 0 | 0 | 0 | 0 | 0 | 0 | 0.002386 | 0 | 0.003353 | 0 | 0 | 0 | 0 | 0 | 0 | 0 | 0 |
| Ruminococcus:K01428 | 0 | 0 | 0 | 0 | 0 | 0 | 0 | 0 | 0 | 0 | 0 | 0 | 0 | 0 | 0 | 0.007181 | 0.001974 | 0 | 0 | 0 | 0 |
| Ruminococcus:K01455 | 0 | 0 | 0 | 0 | 0 | 0 | 0 | 0 | 0 | 0 | 0 | 0 | 0 | 0 | 0.000934 | 0 | 0 | 0 | 0 | 0 | 0 |
| Ruminococcus:K01488 | 0 | 0 | 0 | 0 | 0 | 0 | 0 | 0 | 0 | 0 | 0 | 0 | 0 | 0 | 0.001868 | 0 | 0 | 0 | 0 | 0 | 0.015787812 |
| Ruminococcus:K01613 | 0 | 0 | 0 | 0 | 0 | 0 | 0 | 0 | 0 | 0.0032112 | 0.010023 | 0.0005692 | 0 | 0.000835 | 0.001246 | 0.015667 | 0 | 0 | 0 | 0 | 0 |
| Ruminococcus:K01629 | 0 | 0 | 0 | 0 | 0 | 4.15E-05 | 0 | 0 | 0 | 0.0064225 | 0.003341 | 0.0071717 | 0.002235 | 0.008769 | 0.004879 | 0.004352 | 0.021718 | 0 | 0 | 0 | 0 |
| Ruminococcus:K01679 | 0 | 0.0205 | 0.0356 | 0.0081 | 0 | 0.00017 | 0 | 0 | 0 | 0.090985 | 0.067296 | 0.0169617 | 0.060351 | 0.004593 | 0.000208 | 0.000218 | 0 | 0 | 0 | 0 | 0 |
| Ruminococcus:K01698 | 0 | 0 | 0 | 0 | 0 | 4.15E-05 | 0 | 0 | 0 | 0.0053521 | 0.004773 | 0.001366 | 0.003353 | 0.001253 | 0.000104 | 0.048414 | 0.007897 | 0 | 0 | 0 | 0 |
| Ruminococcus:K01737 | 0 | 0 | 0 | 0 | 0 | 4.15E-05 | 0 | 0.0007694 | 0 | 0 | 0 | 0 | 0 | 0 | 0 | 0 | 0 | 0 | 0 | 0 | 0 |
| Ruminococcus:K01749 | 0 | 0 | 0 | 0 | 0 | 0 | 0 | 0 | 0 | 0 | 0 | 0.0003415 | 0 | 0 | 0 | 0.00087 | 0 | 0 | 0 | 0 | 0 |
| Ruminococcus:K01845 | 0 | 0 | 0 | 0 | 0 | 0 | 0 | 0 | 0 | 0.0010704 | 0.000477 | 0.0004553 | 0 | 0.001253 | 0.000415 | 0.00816 | 0.002962 | 0 | 0 | 0 | 0 |
| Ruminococcus:K01895 | 0 | 0 | 0 | 0.0178 | 0 | 8.29E-05 | 0 | 0 | 0 | 0.0460277 | 0.11025 | 0.006944 | 0.030175 | 0.019626 | 0 | 0.055268 | 0.009872 | 0 | 0 | 0.021473051 | 0 |
| Ruminococcus:K01915 | 0 | 0 | 0 | 0.0048 | 0 | 0.00033 | 0 | 0 | 0 | 0.0385348 | 0.111205 | 0.0326711 | 0.030175 | 0.043429 | 0.043701 | 0.084425 | 0.079961 | 0 | 0 | 0.042946103 | 0 |
| Ruminococcus:K01941 | 0 | 0 | 0 | 0 | 0 | 0 | 0 | 0 | 0 | 0 | 0 | 0 | 0 | 0 | 0 | 0.000109 | 0 | 0 | 0 | 0 | 0 |

| | | | | | | | | | | | | | | | | | | | | | |
|---|---|---|---|---|---|---|---|---|---|---|---|---|---|---|---|---|---|---|---|---|---|
| Ruminococcus:K01958 | 0 | 0 | 0 | 0 | 0 | 0 | 0 | 0 | 0 | 0 | 0 | 0 | 0 | 0 | 0.000623 | 0 | 0 | 0 | 0 | 0 | 0 |
| Ruminococcus:K01959 | 0 | 0 | 0 | 0 | 0 | 0 | 0 | 0 | 0 | 0 | 0 | 0 | 0 | 0 | 0.000415 | 0 | 0 | 0 | 0 | 0 | 0 |
| Ruminococcus:K01960 | 0 | 0 | 0 | 0 | 0 | 0 | 0 | 0 | 0 | 0 | 0 | 0.0001138 | 0 | 0 | 0 | 0.000326 | 0 | 0 | 0 | 0 | 0 |
| Ruminococcus:K01961 | 0 | 0 | 0 | 0 | 0 | 0 | 0 | 0 | 0 | 0 | 0.012409 | 0.0058057 | 0 | 0.002506 | 0.006436 | 0.035903 | 0.007897 | 0 | 0 | 0 | 0 |
| Ruminococcus:K01962 | 0 | 0 | 0 | 0 | 0 | 0 | 0 | 0 | 0 | 0.0042816 | 0.008114 | 0.0004553 | 0.003353 | 0.001253 | 0 | 0.014035 | 0 | 0 | 0 | 0 | 0 |
| Ruminococcus:K02047 | 0 | 0 | 0 | 0 | 0 | 0 | 0 | 0 | 0 | 0 | 0.001432 | 0.0001138 | 0.001118 | 0.004593 | 0.000104 | 0.000109 | 0.000987 | 0 | 0 | 0 | 0 |
| Ruminococcus:K02048 | 0 | 0 | 0 | 0 | 0 | 0 | 0 | 0 | 0 | 0.0010704 | 0.000477 | 0.0014799 | 0.001118 | 0.000418 | 0.00083 | 0.000326 | 0.003949 | 0 | 0 | 0 | 0 |
| Ruminococcus:K02049 | 0 | 0.0409 | 0 | 0.0065 | 0 | 8.29E-05 | 0 | 0 | 0.001025 | 0.0042816 | 0.023386 | 0.0040981 | 0.00447 | 0.001253 | 0.003114 | 0.00816 | 0 | 0 | 0 | 0 | 0 |
| Ruminococcus:K02051 | 0 | 0 | 0 | 0 | 0 | 0 | 0 | 0 | 0 | 0.0042816 | 0.03675 | 0.0027321 | 0.003353 | 0.00167 | 0.007785 | 0.001306 | 0.012833 | 0 | 0 | 0 | 0 |
| Ruminococcus:K02203 | 0 | 0 | 0 | 0 | 0 | 4.15E-05 | 0 | 0 | 0 | 0 | 0.002386 | 0.0030736 | 0.005588 | 0.002088 | 0.000311 | 0.017407 | 0.000987 | 0 | 0 | 0.021473051 | 0 |
| Ruminococcus:K02372 | 0 | 0 | 0 | 0 | 0 | 0 | 0 | 0 | 0 | 0 | 0.006205 | 0.0029598 | 0.002235 | 0.002506 | 0.006124 | 0.01806 | 0.004936 | 0 | 0 | 0 | 0 |
| Ruminococcus:K02492 | 0 | 0 | 0 | 0 | 0 | 0 | 0 | 0 | 0 | 0 | 0 | 0.0001138 | 0 | 0 | 0 | 0.000109 | 0 | 0 | 0 | 0 | 0 |
| Ruminococcus:K03072 | 0 | 0 | 0 | 0 | 0 | 0 | 0 | 0 | 0 | 0.0021408 | 0 | 0.0001138 | 0.001118 | 0.00167 | 0 | 0.006419 | 0 | 0 | 0 | 0 | 0 |
| Ruminococcus:K03092 | 0 | 0 | 0 | 0 | 0 | 0 | 0 | 0 | 0 | 0 | 0 | 0 | 0 | 0.001253 | 0.002076 | 0.000979 | 0 | 0 | 0 | 0 | 0 |
| Ruminococcus:K03394 | 0 | 0 | 0 | 0 | 0 | 0 | 0 | 0 | 0 | 0 | 0.000955 | 0.0002277 | 0 | 0 | 0.000415 | 0.000218 | 0 | 0 | 0 | 0 | 0 |
| Ruminococcus:K03407 | 0 | 0 | 0 | 0 | 0 | 0 | 0 | 0 | 0 | 0.0010704 | 0.000955 | 0 | 0.001118 | 0.000418 | 0 | 0.007072 | 0 | 0 | 0 | 0 | 0 |
| Ruminococcus:K03408 | 0 | 0 | 0 | 0 | 0 | 0 | 0 | 0 | 0 | 0 | 0 | 0.0002277 | 0 | 0.000835 | 0 | 0.029484 | 0 | 0 | 0 | 0 | 0 |
| Ruminococcus:K03412 | 0 | 0 | 0 | 0 | 0 | 0 | 0 | 0 | 0 | 0 | 0 | 0 | 0 | 0 | 0 | 0.009574 | 0 | 0 | 0 | 0 | 0 |
| Ruminococcus:K05343 | 0 | 0 | 0 | 0 | 0 | 0 | 0 | 0 | 0 | 0.0010704 | 0 | 0.0003415 | 0 | 0 | 0.000104 | 0.000544 | 0 | 0 | 0 | 0 | 0 |
| Ruminococcus:K05592 | 0 | 0 | 0 | 0 | 0 | 0 | 0 | 0 | 0 | 0.0074929 | 0.012886 | 0.0003415 | 0.003353 | 0.001253 | 0.000311 | 0.000109 | 0 | 0 | 0 | 0.021473051 | 0 |
| Ruminococcus:K05847 | 0 | 0 | 0 | 0 | 0 | 0 | 0 | 0 | 0 | 0 | 0 | 0 | 0 | 0.000418 | 0 | 0.000653 | 0 | 0 | 0 | 0 | 0 |
| Ruminococcus:K06148 | 0 | 0 | 0 | 0 | 0 | 0 | 0 | 0 | 0 | 0 | 0.000477 | 0.0003415 | 0.001118 | 0 | 0.000208 | 0.000435 | 0 | 0 | 0 | 0 | 0 |
| Ruminococcus:K10536 | 0 | 0 | 0 | 0 | 0 | 0 | 0 | 0 | 0 | 0 | 0 | 0.0001138 | 0.001118 | 0 | 0 | 0.000218 | 0 | 0 | 0 | 0 | 0 |
| Ruminococcus:K11963 | 0 | 0 | 0 | 0 | 0 | 0 | 0 | 0 | 0 | 0 | 0.001909 | 0 | 0.003353 | 0 | 0 | 0 | 0 | 0 | 0 | 0 | 0 |
| Ruminococcus:K12573 | 0 | 0 | 0 | 0 | 0 | 0 | 0 | 0 | 0 | 0.0010704 | 0.002864 | 0.0009107 | 0.001118 | 0.000418 | 0.003633 | 0.010988 | 0.002962 | 0 | 0 | 0 | 0 |
| Slackia:K00104 | 0 | 0 | 0 | 0.0016 | 0 | 0 | 0 | 0 | 0 | 0 | 0 | 0.0004553 | 0 | 0.003758 | 0 | 0.000218 | 0.001974 | 0 | 0 | 0 | 0 |
| Slackia:K00111 | 0 | 0 | 0 | 0.0016 | 0 | 0 | 0 | 0 | 0 | 0 | 0 | 0 | 0 | 0 | 0 | 0 | 0 | 0 | 0 | 0 | 0 |
| Slackia:K00123 | 0 | 0.0205 | 0 | 0.0824 | 0 | 4.15E-05 | 0 | 0 | 0 | 0 | 0 | 0.0005692 | 0 | 0.000418 | 0 | 0 | 0.000987 | 0 | 0 | 0 | 0 |
| Slackia:K00334 | 0 | 0.0205 | 0.0356 | 0.0242 | 0 | 0 | 0 | 0 | 0 | 0 | 0 | 0 | 0 | 0 | 0 | 0 | 0 | 0 | 0 | 0 | 0 |
| Slackia:K00335 | 0 | 0.0205 | 0 | 0.1244 | 0 | 0 | 0 | 0 | 0 | 0.0032112 | 0.000477 | 0.0001138 | 0 | 0 | 0 | 0 | 0 | 0 | 0 | 0 | 0 |
| Slackia:K00336 | 0 | 0 | 0 | 0 | 0 | 0 | 0 | 0 | 0 | 0 | 0 | 0.0001138 | 0 | 0 | 0 | 0 | 0 | 0 | 0 | 0 | 0 |
| Slackia:K00425 | 0 | 0 | 0 | 0 | 0 | 0 | 0 | 0 | 0 | 0 | 0 | 0.0001138 | 0 | 0 | 0 | 0 | 0 | 0 | 0 | 0 | 0 |
| Slackia:K00432 | 0 | 0 | 0 | 0.0081 | 0 | 0 | 0 | 0 | 0 | 0 | 0 | 0 | 0 | 0 | 0 | 0 | 0 | 0 | 0 | 0 | 0 |
| Slackia:K00548 | 0 | 0 | 0 | 0 | 0 | 0 | 0 | 0 | 0 | 0 | 0 | 0.000683 | 0 | 0 | 0.000104 | 0 | 0 | 0 | 0 | 0 | 0 |
| Slackia:K00600 | 0 | 0 | 0 | 0 | 0 | 0 | 0 | 0 | 0 | 0.0021408 | 0.000477 | 0.0002277 | 0 | 0 | 0 | 0 | 0 | 0 | 0 | 0 | 0 |
| Slackia:K00640 | 0 | 0 | 0 | 0 | 0 | 0 | 0 | 0 | 0 | 0 | 0 | 0.001366 | 0 | 0 | 0 | 0 | 0 | 0 | 0 | 0 | 0 |
| Slackia:K00864 | 0 | 0 | 0 | 0.0032 | 0 | 0 | 0 | 0 | 0 | 0 | 0.000477 | 0.0001138 | 0 | 0.000418 | 0.000104 | 0.000109 | 0 | 0 | 0 | 0 | 0 |
| Slackia:K00940 | 0 | 0 | 0 | 0 | 0 | 0 | 0 | 0 | 0 | 0 | 0 | 0.0005692 | 0 | 0 | 0 | 0 | 0.000987 | 0 | 0 | 0 | 0 |
| Slackia:K01251 | 0 | 0 | 0 | 0.0065 | 0 | 8.29E-05 | 0 | 0 | 0 | 0.0010704 | 0 | 0.000683 | 0.001118 | 0 | 0 | 0.000109 | 0 | 0 | 0 | 0 | 0 |
| Slackia:K01338 | 0 | 0 | 0 | 0.0032 | 0 | 0 | 0 | 0 | 0 | 0 | 0 | 0.0018214 | 0 | 0.002088 | 0 | 0.000109 | 0 | 0 | 0 | 0 | 0 |
| Slackia:K01426 | 0 | 0 | 0 | 0 | 0 | 0 | 0 | 0 | 0 | 0 | 0 | 0 | 0 | 0 | 0 | 0 | 0 | 0.0373413 | 0 | 0 | 0 |
| Slackia:K01478 | 0 | 0 | 0 | 0 | 0 | 0 | 0 | 0 | 0 | 0 | 0 | 0 | 0 | 0 | 0.000104 | 0 | 0 | 0 | 0 | 0 | 0 |
| Slackia:K01772 | 0 | 0 | 0 | 0 | 0 | 0 | 0 | 0 | 0 | 0.0010704 | 0 | 0 | 0 | 0 | 0 | 0 | 0 | 0 | 0 | 0 | 0 |
| Slackia:K01845 | 0 | 0 | 0 | 0.0032 | 0 | 0 | 0 | 0 | 0 | 0 | 0 | 0 | 0 | 0 | 0 | 0 | 0 | 0 | 0 | 0 | 0 |
| Slackia:K01878 | 0 | 0.0205 | 0 | 0.042 | 0 | 0 | 0 | 0 | 0 | 0.0010704 | 0 | 0.0004553 | 0 | 0.005429 | 0.000104 | 0.000109 | 0.000987 | 0 | 0 | 0 | 0 |
| Slackia:K01879 | 0 | 0 | 0 | 0.0016 | 0 | 0 | 0 | 0 | 0 | 0.0021408 | 0 | 0.0001138 | 0 | 0 | 0 | 0 | 0 | 0 | 0 | 0 | 0 |
| Slackia:K01895 | 0 | 0 | 0 | 0.0388 | 0 | 0 | 0 | 0 | 0 | 0 | 0 | 0.0002277 | 0 | 0 | 0 | 0 | 0 | 0 | 0 | 0 | 0 |
| Slackia:K01915 | 0 | 0 | 0 | 0 | 0 | 0 | 0 | 0 | 0 | 0 | 0 | 0.0007969 | 0.002235 | 0.00167 | 0 | 0 | 0 | 0 | 0 | 0 | 0 |
| Slackia:K01961 | 0 | 0 | 0 | 0.0032 | 0 | 0 | 0 | 0 | 0 | 0.0010704 | 0.002864 | 0.0010245 | 0.001118 | 0.002088 | 0 | 0.000326 | 0 | 0 | 0 | 0 | 0 |
| Slackia:K01962 | 0 | 0 | 0 | 0.0032 | 0 | 4.15E-05 | 0 | 0 | 0 | 0 | 0.000955 | 0.0014799 | 0.001118 | 0.000418 | 0 | 0.000109 | 0.000987 | 0 | 0 | 0 | 0 |
| Slackia:K02049 | 0 | 0 | 0 | 0 | 0 | 0 | 0 | 0 | 0 | 0 | 0 | 0.0001138 | 0.001118 | 0.000418 | 0 | 0 | 0 | 0 | 0 | 0 | 0 |
| Slackia:K02051 | 0 | 0 | 0 | 0.0016 | 0 | 0 | 0 | 0 | 0 | 0.0010704 | 0.000477 | 0.0001138 | 0 | 0 | 0 | 0 | 0 | 0 | 0 | 0 | 0 |
| Slackia:K02372 | 0 | 0 | 0 | 0 | 0 | 0 | 0 | 0 | 0 | 0 | 0.000477 | 0.0002277 | 0 | 0 | 0 | 0.000326 | 0.000987 | 0 | 0 | 0 | 0 |
| Slackia:K03183 | 0 | 0 | 0 | 0 | 0 | 0 | 0 | 0 | 0 | 0 | 0 | 0 | 0 | 0.000835 | 0 | 0 | 0 | 0 | 0 | 0 | 0 |
| Slackia:K03394 | 0 | 0 | 0 | 0.0032 | 0 | 0 | 0 | 0 | 0 | 0 | 0 | 0 | 0 | 0 | 0 | 0 | 0 | 0 | 0 | 0 | 0 |
| Slackia:K11927 | 0 | 0 | 0 | 0 | 0 | 0 | 0 | 0 | 0 | 0 | 0 | 0.0003415 | 0 | 0 | 0 | 0 | 0 | 0 | 0 | 0 | 0 |
| Slackia:K12573 | 0 | 0 | 0 | 0 | 0 | 0 | 0 | 0 | 0 | 0 | 0 | 0.0003415 | 0 | 0.000418 | 0 | 0 | 0 | 0 | 0 | 0 | 0 |
| Sphingobium:K00029 | 0 | 0 | 0 | 0.0016 | 0 | 0 | 0 | 0 | 0 | 0 | 0 | 0 | 0 | 0 | 0 | 0 | 0 | 0 | 0 | 0 | 0 |
| Sphingobium:K00108 | 0 | 0 | 0.0356 | 0.0032 | 0 | 0 | 0 | 0 | 0 | 0 | 0 | 0 | 0 | 0 | 0 | 0 | 0 | 0 | 0 | 0 | 0 |
| Sphingobium:K00121 | 0 | 0 | 0 | 0.0032 | 0 | 0 | 0 | 0 | 0 | 0 | 0 | 0 | 0 | 0 | 0 | 0 | 0 | 0 | 0 | 0 | 0 |

| | | | | | | | | | | | | | | | | | | | | | |
|---|---|---|---|---|---|---|---|---|---|---|---|---|---|---|---|---|---|---|---|---|---|
| Sphingobium:K00123 | 0 | 0.0205 | 0 | 0.0016 | 0 | 0 | 0 | 0 | 0 | 0 | 0 | 0 | 0 | 0 | 0 | 0 | 0 | 0 | 0 | 0 | 0 |
| Sphingobium:K00128 | 0 | 0 | 0 | 0.0081 | 0 | 0 | 0 | 0 | 0 | 0 | 0 | 0 | 0 | 0 | 0 | 0 | 0 | 0 | 0 | 0 | 0 |
| Sphingobium:K00140 | 0 | 0 | 0 | 0.0016 | 0 | 0 | 0 | 0 | 0 | 0 | 0 | 0 | 0 | 0 | 0 | 0 | 0 | 0 | 0 | 0 | 0 |
| Sphingobium:K00161 | 0 | 0.0205 | 0 | 0.0032 | 0 | 0 | 0 | 0 | 0 | 0 | 0 | 0 | 0 | 0 | 0 | 0 | 0 | 0 | 0 | 0 | 0 |
| Sphingobium:K00162 | 0 | 0 | 0 | 0.0032 | 0 | 0 | 0 | 0 | 0 | 0 | 0 | 0 | 0 | 0 | 0 | 0 | 0 | 0 | 0 | 0 | 0 |
| Sphingobium:K00163 | 0 | 0 | 0 | 0.0016 | 0 | 0 | 0 | 0 | 0 | 0 | 0 | 0 | 0 | 0 | 0 | 0 | 0 | 0 | 0 | 0 | 0 |
| Sphingobium:K00164 | 0 | 0 | 0 | 0.0016 | 0 | 0 | 0 | 0 | 0 | 0 | 0 | 0 | 0 | 0 | 0 | 0 | 0 | 0 | 0 | 0 | 0 |
| Sphingobium:K00253 | 0 | 0 | 0 | 0.0016 | 0 | 0 | 0 | 0 | 0 | 0 | 0 | 0 | 0 | 0 | 0 | 0 | 0 | 0 | 0 | 0 | 0 |
| Sphingobium:K00335 | 0 | 0 | 0.0356 | 0.0016 | 0 | 0 | 0 | 0 | 0 | 0 | 0 | 0 | 0 | 0 | 0 | 0 | 0 | 0 | 0 | 0 | 0 |
| Sphingobium:K00336 | 0 | 0 | 0 | 0.0016 | 0 | 0 | 0 | 0 | 0 | 0 | 0 | 0 | 0 | 0 | 0 | 0 | 0 | 0 | 0 | 0 | 0 |
| Sphingobium:K00341 | 0 | 0 | 0 | 0 | 0 | 4.15E-05 | 0 | 0 | 0 | 0 | 0 | 0 | 0 | 0 | 0 | 0 | 0 | 0 | 0 | 0 | 0 |
| Sphingobium:K00372 | 0 | 0 | 0 | 0.0016 | 0 | 0 | 0 | 0 | 0 | 0 | 0 | 0 | 0 | 0 | 0 | 0 | 0 | 0 | 0 | 0 | 0 |
| Sphingobium:K00383 | 0 | 0 | 0 | 0.0016 | 0 | 4.15E-05 | 0 | 0 | 0 | 0 | 0 | 0 | 0 | 0 | 0 | 0 | 0 | 0 | 0 | 0 | 0 |
| Sphingobium:K00405 | 0 | 0 | 0 | 0.0016 | 0 | 0 | 0 | 0 | 0 | 0 | 0 | 0 | 0 | 0 | 0 | 0 | 0 | 0 | 0 | 0 | 0 |
| Sphingobium:K00412 | 0 | 0.0205 | 0 | 0.0048 | 0 | 0 | 0 | 0 | 0 | 0 | 0 | 0 | 0 | 0 | 0 | 0 | 0 | 0 | 0 | 0 | 0 |
| Sphingobium:K00426 | 0 | 0 | 0 | 0.0016 | 0 | 0 | 0 | 0 | 0 | 0 | 0 | 0 | 0 | 0 | 0 | 0 | 0 | 0 | 0 | 0 | 0 |
| Sphingobium:K00457 | 0 | 0 | 0 | 0.0016 | 0 | 0 | 0 | 0 | 0 | 0 | 0 | 0 | 0 | 0 | 0 | 0 | 0 | 0 | 0 | 0 | 0 |
| Sphingobium:K00525 | 0 | 0 | 0 | 0.0032 | 0 | 0 | 0 | 0 | 0 | 0 | 0 | 0 | 0 | 0 | 0 | 0 | 0 | 0 | 0 | 0 | 0 |
| Sphingobium:K00526 | 0 | 0 | 0 | 0.0065 | 0 | 0 | 0 | 0 | 0 | 0 | 0 | 0 | 0 | 0 | 0 | 0 | 0 | 0 | 0 | 0 | 0 |
| Sphingobium:K00548 | 0 | 0 | 0 | 0.0016 | 0 | 0 | 0 | 0 | 0 | 0 | 0 | 0 | 0 | 0 | 0 | 0 | 0 | 0 | 0 | 0 | 0 |
| Sphingobium:K00600 | 0 | 0 | 0 | 0 | 0 | 0 | 0 | 0 | 0 | 0 | 0 | 0 | 0.001118 | 0 | 0 | 0 | 0 | 0 | 0 | 0 | 0 |
| Sphingobium:K00658 | 0 | 0 | 0 | 0.0032 | 0 | 0 | 0 | 0 | 0 | 0 | 0 | 0 | 0 | 0 | 0 | 0 | 0 | 0 | 0 | 0 | 0 |
| Sphingobium:K00681 | 0 | 0 | 0 | 0.0016 | 0 | 0 | 0 | 0 | 0 | 0 | 0 | 0 | 0 | 0 | 0 | 0 | 0 | 0 | 0 | 0 | 0 |
| Sphingobium:K00955 | 0 | 0 | 0 | 0.0016 | 0 | 0 | 0 | 0 | 0 | 0 | 0 | 0 | 0 | 0 | 0 | 0 | 0 | 0 | 0 | 0 | 0 |
| Sphingobium:K01029 | 0 | 0 | 0 | 0.0016 | 0 | 0 | 0 | 0 | 0 | 0 | 0 | 0 | 0 | 0 | 0 | 0 | 0 | 0 | 0 | 0 | 0 |
| Sphingobium:K01241 | 0 | 0 | 0 | 0.0016 | 0 | 0 | 0 | 0 | 0 | 0 | 0 | 0 | 0 | 0 | 0 | 0 | 0 | 0 | 0 | 0 | 0 |
| Sphingobium:K01251 | 0 | 0 | 0 | 0.0032 | 0 | 0 | 0 | 0 | 0 | 0 | 0 | 0 | 0 | 0 | 0 | 0 | 0 | 0 | 0 | 0 | 0 |
| Sphingobium:K01255 | 0 | 0 | 0 | 0.0016 | 0 | 0 | 0 | 0 | 0 | 0 | 0 | 0 | 0 | 0 | 0 | 0 | 0 | 0 | 0 | 0 | 0 |
| Sphingobium:K01338 | 0 | 0.0205 | 0 | 0.0016 | 0 | 0 | 0 | 0 | 0 | 0 | 0 | 0 | 0 | 0 | 0 | 0 | 0 | 0 | 0 | 0 | 0 |
| Sphingobium:K01676 | 0 | 0 | 0 | 0.0016 | 0 | 0 | 0 | 0 | 0 | 0 | 0 | 0 | 0 | 0 | 0 | 0 | 0 | 0 | 0 | 0 | 0 |
| Sphingobium:K01690 | 0 | 0 | 0 | 0.0016 | 0 | 0 | 0 | 0 | 0 | 0 | 0 | 0 | 0 | 0 | 0 | 0 | 0 | 0 | 0 | 0 | 0 |
| Sphingobium:K01692 | 0 | 0 | 0 | 0.0016 | 0 | 0 | 0 | 0 | 0 | 0 | 0 | 0 | 0 | 0 | 0 | 0 | 0 | 0 | 0 | 0 | 0 |
| Sphingobium:K01895 | 0 | 0 | 0 | 0.0032 | 0 | 0 | 0 | 0 | 0 | 0 | 0 | 0 | 0 | 0 | 0 | 0 | 0 | 0 | 0 | 0 | 0 |
| Sphingobium:K01915 | 0 | 0 | 0 | 0.0016 | 0 | 0 | 0 | 0 | 0 | 0 | 0 | 0 | 0 | 0 | 0 | 0 | 0 | 0 | 0 | 0 | 0 |
| Sphingobium:K01941 | 0 | 0 | 0 | 0.0016 | 0 | 0 | 0 | 0 | 0 | 0 | 0 | 0 | 0 | 0 | 0 | 0 | 0 | 0 | 0 | 0 | 0 |
| Sphingobium:K01962 | 0 | 0 | 0 | 0.0016 | 0 | 0 | 0 | 0 | 0 | 0 | 0 | 0 | 0 | 0 | 0 | 0 | 0 | 0 | 0 | 0 | 0 |
| Sphingobium:K01969 | 0 | 0 | 0 | 0 | 0 | 0 | 0 | 0.0007694 | 0 | 0 | 0 | 0 | 0 | 0 | 0 | 0 | 0 | 0 | 0 | 0 | 0 |
| Sphingobium:K02274 | 0 | 0 | 0 | 0.0032 | 0 | 0 | 0 | 0 | 0 | 0 | 0 | 0 | 0 | 0 | 0 | 0 | 0 | 0 | 0 | 0 | 0 |
| Sphingobium:K02276 | 0 | 0 | 0 | 0.0016 | 0 | 0 | 0 | 0 | 0 | 0 | 0 | 0 | 0 | 0 | 0 | 0 | 0 | 0 | 0 | 0 | 0 |
| Sphingobium:K02412 | 0 | 0 | 0 | 0.0016 | 0 | 0 | 0 | 0 | 0 | 0 | 0 | 0 | 0 | 0 | 0 | 0 | 0 | 0 | 0 | 0 | 0 |
| Sphingobium:K02523 | 0 | 0 | 0 | 0.0016 | 0 | 0 | 0 | 0 | 0 | 0 | 0 | 0 | 0 | 0 | 0 | 0 | 0 | 0 | 0 | 0 | 0 |
| Sphingobium:K03412 | 0 | 0 | 0 | 0 | 0 | 4.15E-05 | 0 | 0 | 0 | 0 | 0 | 0 | 0 | 0 | 0 | 0 | 0 | 0 | 0 | 0 | 0 |
| Sphingobium:K03590 | 0 | 0 | 0 | 0.0016 | 0 | 0 | 0 | 0 | 0 | 0 | 0 | 0 | 0 | 0 | 0 | 0 | 0 | 0 | 0 | 0 | 0 |
| Sphingobium:K03644 | 0 | 0 | 0 | 0.0016 | 0 | 0 | 0 | 0 | 0 | 0 | 0 | 0 | 0 | 0 | 0 | 0 | 0 | 0 | 0 | 0 | 0 |
| Variovorax:K00029 | 0 | 0 | 0 | 0 | 0 | 0.00025 | 0 | 0 | 0 | 0 | 0 | 0 | 0 | 0 | 0 | 0 | 0 | 0 | 0 | 0 | 0 |
| Variovorax:K00104 | 0 | 0 | 0 | 0 | 0 | 0.00017 | 0 | 0.0007694 | 0 | 0 | 0 | 0 | 0 | 0 | 0 | 0 | 0 | 0 | 0 | 0 | 0 |
| Variovorax:K00108 | 0 | 0 | 0 | 0.0048 | 0 | 0 | 0.002207 | 0 | 0 | 0 | 0 | 0 | 0 | 0 | 0 | 0 | 0 | 0 | 0 | 0 | 0 |
| Variovorax:K00111 | 0 | 0 | 0 | 0 | 0 | 8.29E-05 | 0 | 0 | 0.001025 | 0 | 0 | 0 | 0 | 0 | 0 | 0 | 0 | 0 | 0 | 0 | 0 |
| Variovorax:K00117 | 0 | 0 | 0 | 0.0016 | 0 | 0 | 0 | 0 | 0 | 0 | 0 | 0 | 0 | 0 | 0 | 0 | 0 | 0 | 0 | 0 | 0 |
| Variovorax:K00128 | 0 | 0 | 0 | 0 | 0 | 0.00025 | 0 | 0 | 0 | 0 | 0 | 0 | 0 | 0 | 0 | 0 | 0 | 0 | 0 | 0 | 0 |
| Variovorax:K00130 | 0 | 0 | 0 | 0.0016 | 0 | 8.29E-05 | 0 | 0 | 0 | 0 | 0 | 0 | 0 | 0 | 0 | 0 | 0 | 0 | 0 | 0 | 0 |
| Variovorax:K00140 | 0 | 0 | 0 | 0.0016 | 0 | 4.15E-05 | 0 | 0 | 0 | 0 | 0 | 0 | 0 | 0 | 0 | 0 | 0 | 0 | 0 | 0 | 0 |
| Variovorax:K00162 | 0 | 0 | 0 | 0.0016 | 0 | 8.29E-05 | 0.002207 | 0 | 0 | 0 | 0 | 0 | 0 | 0 | 0 | 0 | 0 | 0 | 0 | 0 | 0 |
| Variovorax:K00163 | 0 | 0 | 0 | 0.0016 | 0 | 0.00017 | 0 | 0.0007694 | 0 | 0 | 0 | 0 | 0 | 0 | 0 | 0 | 0 | 0 | 0 | 0 | 0 |
| Variovorax:K00164 | 0 | 0 | 0 | 0.0048 | 0 | 4.15E-05 | 0 | 0.0007694 | 0 | 0 | 0 | 0 | 0 | 0 | 0 | 0 | 0 | 0 | 0 | 0 | 0 |
| Variovorax:K00248 | 0 | 0 | 0 | 0 | 0 | 0.00021 | 0 | 0 | 0 | 0 | 0 | 0 | 0 | 0 | 0 | 0 | 0 | 0 | 0 | 0 | 0 |
| Variovorax:K00249 | 0 | 0.0205 | 0 | 0.0016 | 0 | 0.00017 | 0 | 0 | 0 | 0 | 0 | 0 | 0 | 0 | 0 | 0 | 0 | 0 | 0 | 0 | 0 |
| Variovorax:K00253 | 0 | 0 | 0 | 0 | 0 | 0.00017 | 0 | 0.0007694 | 0.001025 | 0 | 0 | 0 | 0 | 0 | 0 | 0 | 0 | 0 | 0 | 0 | 0 |
| Variovorax:K00257 | 0 | 0.0205 | 0 | 0.0016 | 0 | 0.00075 | 0 | 0 | 0 | 0 | 0 | 0 | 0 | 0 | 0 | 0 | 0 | 0 | 0 | 0 | 0 |
| Variovorax:K00281 | 0 | 0 | 0 | 0 | 0 | 4.15E-05 | 0 | 0 | 0 | 0 | 0 | 0 | 0 | 0 | 0 | 0 | 0 | 0 | 0 | 0 | 0 |

| | | | | | | | | | | | | | | | | | | | | | |
|---|---|---|---|---|---|---|---|---|---|---|---|---|---|---|---|---|---|---|---|---|---|
| Variovorax:K00285 | 0 | 0 | 0 | 0.0016 | 0 | 0.00012 | 0 | 0 | 0 | 0 | 0 | 0 | 0 | 0 | 0 | 0 | 0 | 0 | 0 | 0 | 0 |
| Variovorax:K00334 | 0 | 0.0205 | 0 | 0.0048 | 0 | 0 | 0 | 0.0007694 | 0 | 0 | 0 | 0 | 0 | 0 | 0 | 0 | 0 | 0 | 0 | 0 | 0 |
| Variovorax:K00335 | 0 | 0 | 0 | 0 | 0 | 4.15E-05 | 0 | 0 | 0 | 0 | 0 | 0 | 0 | 0 | 0 | 0 | 0 | 0 | 0 | 0 | 0 |
| Variovorax:K00336 | 0 | 0 | 0 | 0 | 0 | 0.00012 | 0 | 0 | 0 | 0 | 0 | 0 | 0 | 0 | 0 | 0 | 0 | 0 | 0 | 0 | 0 |
| Variovorax:K00337 | 0 | 0 | 0 | 0 | 0 | 8.29E-05 | 0 | 0 | 0 | 0 | 0 | 0 | 0 | 0 | 0 | 0 | 0 | 0 | 0 | 0 | 0 |
| Variovorax:K00341 | 0 | 0 | 0 | 0 | 0 | 4.15E-05 | 0 | 0 | 0 | 0 | 0 | 0 | 0 | 0 | 0 | 0 | 0 | 0 | 0 | 0 | 0 |
| Variovorax:K00342 | 0 | 0 | 0 | 0 | 0 | 4.15E-05 | 0 | 0 | 0 | 0 | 0 | 0 | 0 | 0 | 0 | 0 | 0 | 0 | 0 | 0 | 0 |
| Variovorax:K00362 | 0 | 0 | 0 | 0 | 0 | 0.00025 | 0 | 0 | 0.001025 | 0 | 0 | 0 | 0 | 0 | 0 | 0 | 0 | 0 | 0 | 0 | 0 |
| Variovorax:K00372 | 0 | 0 | 0 | 0 | 0 | 8.29E-05 | 0 | 0 | 0.001025 | 0 | 0 | 0 | 0 | 0 | 0 | 0 | 0 | 0 | 0 | 0 | 0 |
| Variovorax:K00381 | 0 | 0 | 0 | 0 | 0 | 8.29E-05 | 0 | 0 | 0 | 0 | 0 | 0 | 0 | 0 | 0 | 0 | 0 | 0 | 0 | 0 | 0 |
| Variovorax:K00383 | 0 | 0 | 0 | 0 | 0 | 8.29E-05 | 0 | 0 | 0 | 0 | 0 | 0 | 0 | 0 | 0 | 0 | 0 | 0 | 0 | 0 | 0 |
| Variovorax:K00412 | 0 | 0 | 0 | 0.0016 | 0 | 0.00012 | 0 | 0 | 0 | 0 | 0 | 0 | 0 | 0 | 0 | 0 | 0 | 0 | 0 | 0 | 0 |
| Variovorax:K00425 | 0 | 0 | 0 | 0.0016 | 0 | 0 | 0 | 0 | 0 | 0 | 0 | 0 | 0 | 0 | 0 | 0 | 0 | 0 | 0 | 0 | 0 |
| Variovorax:K00426 | 0 | 0 | 0 | 0.0016 | 0 | 0 | 0 | 0 | 0 | 0 | 0 | 0 | 0 | 0 | 0 | 0 | 0 | 0 | 0 | 0 | 0 |
| Variovorax:K00432 | 0 | 0 | 0 | 0 | 0 | 4.15E-05 | 0 | 0 | 0 | 0 | 0 | 0 | 0 | 0 | 0 | 0 | 0 | 0 | 0 | 0 | 0 |
| Variovorax:K00451 | 0 | 0 | 0 | 0 | 0 | 4.15E-05 | 0 | 0 | 0 | 0 | 0 | 0 | 0 | 0 | 0 | 0 | 0 | 0 | 0 | 0 | 0 |
| Variovorax:K00457 | 0 | 0 | 0 | 0 | 0 | 0.00017 | 0 | 0 | 0 | 0 | 0 | 0 | 0 | 0 | 0 | 0 | 0 | 0 | 0 | 0 | 0 |
| Variovorax:K00525 | 0 | 0 | 0 | 0.0016 | 0 | 0.00021 | 0 | 0 | 0 | 0 | 0 | 0 | 0 | 0 | 0 | 0 | 0 | 0 | 0 | 0 | 0 |
| Variovorax:K00548 | 0 | 0 | 0 | 0 | 0 | 0.00012 | 0 | 0 | 0 | 0 | 0 | 0 | 0 | 0 | 0 | 0 | 0 | 0 | 0 | 0 | 0 |
| Variovorax:K00568 | 0 | 0 | 0 | 0 | 0 | 8.29E-05 | 0 | 0 | 0 | 0 | 0 | 0 | 0 | 0 | 0 | 0 | 0 | 0 | 0 | 0 | 0 |
| Variovorax:K00575 | 0 | 0 | 0 | 0.0016 | 0 | 0 | 0 | 0 | 0 | 0 | 0 | 0 | 0 | 0 | 0 | 0 | 0 | 0 | 0 | 0 | 0 |
| Variovorax:K00600 | 0 | 0 | 0 | 0.0048 | 0 | 4.15E-05 | 0 | 0 | 0 | 0 | 0 | 0 | 0 | 0 | 0 | 0 | 0 | 0 | 0 | 0 | 0 |
| Variovorax:K00626 | 0 | 0 | 0 | 0.0032 | 0 | 0.00029 | 0 | 0 | 0 | 0 | 0 | 0 | 0 | 0 | 0 | 0 | 0 | 0 | 0 | 0 | 0 |
| Variovorax:K00627 | 0 | 0 | 0 | 0 | 0 | 4.15E-05 | 0 | 0 | 0 | 0 | 0 | 0 | 0 | 0 | 0 | 0 | 0 | 0 | 0 | 0 | 0 |
| Variovorax:K00632 | 0 | 0 | 0 | 0 | 0 | 8.29E-05 | 0 | 0.0007694 | 0 | 0 | 0 | 0 | 0 | 0 | 0 | 0 | 0 | 0 | 0 | 0 | 0 |
| Variovorax:K00641 | 0 | 0.0205 | 0 | 0.0065 | 0 | 4.15E-05 | 0 | 0 | 0 | 0 | 0 | 0 | 0 | 0 | 0 | 0 | 0 | 0 | 0 | 0 | 0 |
| Variovorax:K00647 | 0 | 0 | 0 | 0 | 0 | 8.29E-05 | 0.002207 | 0 | 0 | 0 | 0 | 0 | 0 | 0 | 0 | 0 | 0 | 0 | 0 | 0 | 0 |
| Variovorax:K00648 | 0 | 0 | 0 | 0.0032 | 0 | 8.29E-05 | 0 | 0 | 0 | 0 | 0 | 0 | 0 | 0 | 0 | 0 | 0 | 0 | 0 | 0 | 0 |
| Variovorax:K00658 | 0 | 0 | 0 | 0 | 0 | 8.29E-05 | 0 | 0 | 0 | 0 | 0 | 0 | 0 | 0 | 0 | 0 | 0 | 0 | 0 | 0 | 0 |
| Variovorax:K00681 | 0 | 0 | 0 | 0.0016 | 0 | 0.00021 | 0 | 0 | 0 | 0 | 0 | 0 | 0 | 0 | 0 | 0 | 0 | 0 | 0 | 0 | 0 |
| Variovorax:K00697 | 0 | 0 | 0 | 0 | 0 | 0.00012 | 0 | 0 | 0 | 0 | 0 | 0 | 0 | 0 | 0 | 0 | 0 | 0 | 0 | 0 | 0 |
| Variovorax:K00798 | 0 | 0 | 0 | 0 | 0 | 0.00012 | 0 | 0 | 0 | 0 | 0 | 0 | 0 | 0 | 0 | 0 | 0 | 0 | 0 | 0 | 0 |
| Variovorax:K00799 | 0 | 0 | 0 | 0.0016 | 0 | 0.00033 | 0 | 0 | 0 | 0 | 0 | 0 | 0 | 0 | 0 | 0 | 0 | 0 | 0 | 0 | 0 |
| Variovorax:K00833 | 0 | 0 | 0 | 0 | 0 | 0 | 0 | 0 | 0 | 0 | 0 | 0 | 0.001118 | 0 | 0 | 0 | 0 | 0 | 0 | 0 | 0 |
| Variovorax:K00864 | 0 | 0 | 0 | 0.0016 | 0 | 8.29E-05 | 0 | 0.0007694 | 0 | 0 | 0 | 0 | 0 | 0 | 0 | 0 | 0 | 0 | 0 | 0 | 0 |
| Variovorax:K01142 | 0 | 0 | 0 | 0 | 0 | 0.00017 | 0 | 0 | 0 | 0 | 0 | 0 | 0 | 0 | 0 | 0 | 0 | 0 | 0 | 0 | 0 |
| Variovorax:K01251 | 0 | 0 | 0.0356 | 0.0016 | 0 | 0 | 0 | 0 | 0 | 0 | 0 | 0 | 0 | 0 | 0 | 0 | 0 | 0 | 0 | 0 | 0 |
| Variovorax:K01255 | 0 | 0 | 0 | 0 | 0 | 4.15E-05 | 0 | 0 | 0 | 0 | 0 | 0 | 0 | 0 | 0 | 0 | 0 | 0 | 0 | 0 | 0 |
| Variovorax:K01338 | 0 | 0 | 0 | 0 | 0 | 0.00012 | 0 | 0 | 0 | 0 | 0 | 0 | 0 | 0 | 0 | 0 | 0 | 0 | 0 | 0 | 0 |
| Variovorax:K01426 | 0 | 0 | 0 | 0.0016 | 0 | 8.29E-05 | 0 | 0 | 0 | 0 | 0 | 0 | 0 | 0 | 0 | 0 | 0 | 0 | 0 | 0 | 0 |
| Variovorax:K01428 | 0 | 0 | 0 | 0 | 0 | 4.15E-05 | 0 | 0 | 0 | 0 | 0 | 0 | 0 | 0 | 0 | 0 | 0 | 0 | 0 | 0 | 0 |
| Variovorax:K01433 | 0 | 0 | 0 | 0 | 0 | 0.00012 | 0 | 0 | 0 | 0 | 0 | 0 | 0 | 0 | 0 | 0 | 0 | 0 | 0 | 0 | 0 |
| Variovorax:K01455 | 0 | 0 | 0 | 0 | 0 | 4.15E-05 | 0 | 0 | 0 | 0 | 0 | 0 | 0 | 0 | 0 | 0 | 0 | 0 | 0 | 0 | 0 |
| Variovorax:K01584 | 0 | 0 | 0 | 0.0016 | 0 | 0.00017 | 0 | 0 | 0.001025 | 0 | 0 | 0 | 0 | 0 | 0 | 0 | 0 | 0 | 0 | 0 | 0 |
| Variovorax:K01613 | 0 | 0 | 0 | 0 | 0 | 8.29E-05 | 0 | 0 | 0 | 0 | 0 | 0 | 0 | 0 | 0 | 0 | 0 | 0 | 0 | 0 | 0 |
| Variovorax:K01627 | 0 | 0 | 0 | 0 | 0 | 0 | 0 | 0 | 0 | 0 | 0 | 0 | 0 | 0.000835 | 0 | 0 | 0 | 0 | 0 | 0 | 0 |
| Variovorax:K01637 | 0 | 0 | 0 | 0 | 0 | 8.29E-05 | 0 | 0 | 0 | 0 | 0 | 0 | 0 | 0 | 0 | 0 | 0 | 0 | 0 | 0 | 0 |
| Variovorax:K01676 | 0 | 0 | 0 | 0.0016 | 0 | 4.15E-05 | 0 | 0 | 0 | 0 | 0 | 0 | 0 | 0 | 0 | 0 | 0 | 0 | 0 | 0 | 0 |
| Variovorax:K01679 | 0 | 0 | 0 | 0 | 0 | 8.29E-05 | 0 | 0 | 0 | 0 | 0 | 0 | 0 | 0 | 0 | 0 | 0 | 0 | 0 | 0 | 0 |
| Variovorax:K01692 | 0 | 0 | 0 | 0 | 0 | 0.00012 | 0 | 0 | 0 | 0 | 0 | 0 | 0 | 0 | 0 | 0 | 0 | 0 | 0 | 0 | 0 |
| Variovorax:K01712 | 0 | 0.0205 | 0 | 0 | 0 | 0 | 0 | 0.0007694 | 0 | 0 | 0 | 0 | 0 | 0 | 0 | 0 | 0 | 0 | 0 | 0 | 0 |
| Variovorax:K01745 | 0 | 0 | 0 | 0 | 0 | 4.15E-05 | 0 | 0 | 0 | 0 | 0 | 0 | 0 | 0 | 0 | 0 | 0 | 0 | 0 | 0 | 0 |
| Variovorax:K01749 | 0 | 0 | 0 | 0 | 0 | 4.15E-05 | 0 | 0 | 0 | 0 | 0 | 0 | 0 | 0 | 0 | 0 | 0 | 0 | 0 | 0 | 0 |
| Variovorax:K01772 | 0 | 0 | 0 | 0 | 0 | 4.15E-05 | 0 | 0 | 0 | 0 | 0 | 0 | 0 | 0 | 0 | 0 | 0 | 0 | 0 | 0 | 0 |
| Variovorax:K01845 | 0 | 0 | 0 | 0 | 0 | 4.15E-05 | 0 | 0 | 0 | 0 | 0 | 0 | 0 | 0 | 0 | 0 | 0 | 0 | 0 | 0 | 0 |
| Variovorax:K01878 | 0 | 0 | 0 | 0 | 0 | 4.15E-05 | 0 | 0 | 0 | 0 | 0 | 0 | 0 | 0 | 0 | 0 | 0 | 0 | 0 | 0 | 0 |
| Variovorax:K01879 | 0 | 0 | 0 | 0 | 0 | 8.29E-05 | 0 | 0 | 0.001025 | 0 | 0 | 0 | 0 | 0 | 0 | 0 | 0 | 0 | 0 | 0 | 0 |
| Variovorax:K01895 | 0 | 0 | 0 | 0.0016 | 0 | 4.15E-05 | 0 | 0 | 0 | 0 | 0 | 0 | 0 | 0 | 0 | 0 | 0 | 0 | 0 | 0 | 0 |
| Variovorax:K01907 | 0 | 0 | 0 | 0.0016 | 0 | 8.29E-05 | 0 | 0.0007694 | 0 | 0 | 0 | 0 | 0 | 0 | 0 | 0 | 0 | 0 | 0 | 0 | 0 |
| Variovorax:K01920 | 0 | 0 | 0 | 0.0016 | 0 | 8.29E-05 | 0 | 0 | 0 | 0 | 0 | 0 | 0 | 0 | 0 | 0 | 0 | 0 | 0 | 0 | 0 |

| | | | | | | | | | | | | | | | | | | | | | |
|---|---|---|---|---|---|---|---|---|---|---|---|---|---|---|---|---|---|---|---|---|---|
| Variovorax:K01961 | 0 | 0 | 0 | 0 | 0 | 0.00033 | 0 | 0.0007694 | 0 | 0 | 0 | 0 | 0 | 0 | 0 | 0 | 0 | 0 | 0 | 0 | 0 |
| Variovorax:K01962 | 0 | 0 | 0 | 0.0016 | 0 | 0 | 0 | 0 | 0 | 0 | 0 | 0 | 0 | 0 | 0 | 0 | 0 | 0 | 0 | 0 | 0 |
| Variovorax:K01969 | 0 | 0.0205 | 0 | 0.0032 | 0 | 8.29E-05 | 0 | 0.0007694 | 0 | 0 | 0 | 0 | 0 | 0 | 0 | 0 | 0 | 0 | 0 | 0 | 0 |
| Variovorax:K02000 | 0 | 0 | 0 | 0.0016 | 0 | 0 | 0 | 0 | 0 | 0 | 0 | 0 | 0 | 0 | 0 | 0 | 0 | 0 | 0 | 0 | 0 |
| Variovorax:K02002 | 0 | 0 | 0 | 0.0016 | 0 | 0 | 0 | 0 | 0 | 0 | 0 | 0 | 0 | 0 | 0 | 0 | 0 | 0 | 0 | 0 | 0 |
| Variovorax:K02011 | 0 | 0 | 0 | 0 | 0 | 4.15E-05 | 0.002207 | 0 | 0 | 0 | 0 | 0 | 0 | 0 | 0 | 0 | 0 | 0 | 0 | 0 | 0 |
| Variovorax:K02012 | 0 | 0 | 0 | 0 | 0 | 0.00012 | 0 | 0.0007694 | 0.002051 | 0 | 0 | 0 | 0 | 0 | 0 | 0 | 0 | 0 | 0 | 0 | 0 |
| Variovorax:K02047 | 0 | 0 | 0 | 0.0032 | 0 | 0.00012 | 0 | 0 | 0 | 0 | 0 | 0 | 0 | 0 | 0 | 0 | 0 | 0 | 0 | 0 | 0 |
| Variovorax:K02048 | 0 | 0 | 0 | 0.0097 | 0 | 0.00017 | 0 | 0 | 0 | 0 | 0 | 0 | 0 | 0 | 0 | 0 | 0 | 0 | 0 | 0 | 0 |
| Variovorax:K02049 | 0 | 0 | 0.0356 | 0 | 0 | 0.00033 | 0 | 0 | 0 | 0 | 0 | 0 | 0 | 0 | 0 | 0 | 0 | 0 | 0 | 0 | 0 |
| Variovorax:K02051 | 0 | 0.0205 | 0 | 0 | 0 | 0.00029 | 0 | 0 | 0 | 0 | 0 | 0 | 0 | 0 | 0 | 0 | 0 | 0 | 0 | 0 | 0 |
| Variovorax:K02274 | 0 | 0 | 0 | 0.0016 | 0 | 0 | 0 | 0 | 0 | 0 | 0 | 0 | 0 | 0 | 0 | 0 | 0 | 0 | 0 | 0 | 0 |
| Variovorax:K02276 | 0 | 0 | 0 | 0 | 0 | 8.29E-05 | 0.002207 | 0 | 0 | 0 | 0 | 0 | 0 | 0 | 0 | 0 | 0 | 0 | 0 | 0 | 0 |
| Variovorax:K02301 | 0 | 0 | 0 | 0 | 0 | 4.15E-05 | 0 | 0 | 0 | 0 | 0 | 0 | 0 | 0 | 0 | 0 | 0 | 0 | 0 | 0 | 0 |
| Variovorax:K02409 | 0 | 0 | 0 | 0 | 0 | 4.15E-05 | 0 | 0 | 0 | 0 | 0 | 0 | 0 | 0 | 0 | 0 | 0 | 0 | 0 | 0 | 0 |
| Variovorax:K02454 | 0 | 0 | 0 | 0 | 0 | 8.29E-05 | 0 | 0 | 0 | 0 | 0 | 0 | 0 | 0 | 0 | 0 | 0 | 0 | 0 | 0 | 0 |
| Variovorax:K02517 | 0 | 0 | 0 | 0 | 0 | 4.15E-05 | 0 | 0 | 0 | 0 | 0 | 0 | 0 | 0 | 0 | 0 | 0 | 0 | 0 | 0 | 0 |
| Variovorax:K02535 | 0 | 0 | 0 | 0 | 0 | 8.29E-05 | 0 | 0 | 0 | 0 | 0 | 0 | 0 | 0 | 0 | 0 | 0 | 0 | 0 | 0 | 0 |
| Variovorax:K03072 | 0 | 0 | 0 | 0 | 0 | 0.00012 | 0 | 0 | 0 | 0 | 0 | 0 | 0 | 0 | 0 | 0 | 0 | 0 | 0 | 0 | 0 |
| Variovorax:K03087 | 0 | 0 | 0 | 0 | 0 | 4.15E-05 | 0 | 0 | 0 | 0 | 0 | 0 | 0 | 0 | 0 | 0 | 0 | 0 | 0 | 0 | 0 |
| Variovorax:K03092 | 0 | 0 | 0 | 0 | 0 | 0.00012 | 0 | 0 | 0 | 0 | 0 | 0 | 0 | 0 | 0 | 0 | 0 | 0 | 0 | 0 | 0 |
| Variovorax:K03183 | 0 | 0 | 0 | 0 | 0 | 8.29E-05 | 0 | 0 | 0 | 0 | 0 | 0 | 0 | 0 | 0 | 0 | 0 | 0 | 0 | 0 | 0 |
| Variovorax:K03407 | 0 | 0 | 0 | 0.0016 | 0 | 0 | 0 | 0 | 0 | 0 | 0 | 0 | 0 | 0 | 0 | 0 | 0 | 0 | 0 | 0 | 0 |
| Variovorax:K03644 | 0 | 0 | 0 | 0 | 0 | 4.15E-05 | 0 | 0 | 0 | 0 | 0 | 0 | 0 | 0 | 0 | 0 | 0 | 0 | 0 | 0 | 0 |
| Variovorax:K03735 | 0 | 0 | 0 | 0 | 0 | 8.29E-05 | 0.002207 | 0 | 0 | 0 | 0 | 0 | 0 | 0 | 0 | 0 | 0 | 0 | 0 | 0 | 0 |
| Variovorax:K03781 | 0 | 0 | 0 | 0.0016 | 0 | 8.29E-05 | 0 | 0 | 0 | 0 | 0 | 0 | 0 | 0 | 0 | 0 | 0 | 0 | 0 | 0 | 0 |
| Variovorax:K03841 | 0 | 0 | 0 | 0.0081 | 0 | 0 | 0 | 0 | 0 | 0 | 0 | 0 | 0 | 0 | 0 | 0 | 0 | 0 | 0 | 0 | 0 |
| Variovorax:K03885 | 0 | 0 | 0 | 0 | 0 | 4.15E-05 | 0 | 0 | 0 | 0 | 0 | 0 | 0 | 0 | 0 | 0 | 0 | 0 | 0 | 0 | 0 |
| Variovorax:K04564 | 0 | 0 | 0 | 0.0016 | 0 | 0 | 0 | 0 | 0 | 0 | 0 | 0 | 0 | 0 | 0 | 0 | 0 | 0 | 0 | 0 | 0 |
| Variovorax:K05343 | 0 | 0 | 0 | 0.0016 | 0 | 8.29E-05 | 0 | 0 | 0 | 0 | 0 | 0 | 0 | 0 | 0 | 0 | 0 | 0 | 0 | 0 | 0 |
| Variovorax:K05846 | 0 | 0 | 0 | 0.0016 | 0 | 8.29E-05 | 0 | 0 | 0 | 0 | 0 | 0 | 0 | 0 | 0 | 0 | 0 | 0 | 0 | 0 | 0 |
| Variovorax:K06596 | 0 | 0 | 0 | 0 | 0 | 0.00017 | 0 | 0 | 0 | 0 | 0 | 0 | 0 | 0 | 0 | 0 | 0 | 0 | 0 | 0 | 0 |
| Variovorax:K06861 | 0 | 0 | 0 | 0.0065 | 0 | 4.15E-05 | 0 | 0 | 0 | 0 | 0 | 0 | 0 | 0 | 0 | 0 | 0 | 0 | 0 | 0 | 0 |
| Variovorax:K07516 | 0 | 0 | 0 | 0.0016 | 0 | 0.00046 | 0 | 0.0007694 | 0 | 0 | 0 | 0 | 0 | 0 | 0 | 0 | 0 | 0 | 0 | 0 | 0 |
| Variovorax:K07649 | 0 | 0 | 0 | 0 | 0 | 8.29E-05 | 0 | 0 | 0 | 0 | 0 | 0 | 0 | 0 | 0 | 0 | 0 | 0 | 0 | 0 | 0 |
| Variovorax:K07712 | 0 | 0 | 0 | 0.0016 | 0 | 0 | 0 | 0 | 0 | 0 | 0 | 0 | 0 | 0 | 0 | 0 | 0 | 0 | 0 | 0 | 0 |
| Variovorax:K07788 | 0 | 0 | 0 | 0 | 0 | 4.15E-05 | 0 | 0 | 0 | 0 | 0 | 0 | 0 | 0 | 0 | 0 | 0 | 0 | 0 | 0 | 0 |
| Variovorax:K08311 | 0 | 0 | 0 | 0.0016 | 0 | 0 | 0 | 0 | 0 | 0 | 0 | 0 | 0 | 0 | 0 | 0 | 0 | 0 | 0 | 0 | 0 |
| Variovorax:K11891 | 0 | 0 | 0 | 0 | 0 | 0.00029 | 0 | 0 | 0.001025 | 0 | 0 | 0 | 0 | 0 | 0 | 0 | 0 | 0 | 0 | 0 | 0 |
| Variovorax:K11903 | 0 | 0 | 0 | 0 | 0 | 4.15E-05 | 0 | 0 | 0 | 0 | 0 | 0 | 0 | 0 | 0 | 0 | 0 | 0 | 0 | 0 | 0 |
| Variovorax:K11904 | 0 | 0 | 0 | 0.0032 | 0 | 8.29E-05 | 0 | 0 | 0.001025 | 0 | 0 | 0 | 0 | 0 | 0 | 0 | 0 | 0 | 0 | 0 | 0 |
| Variovorax:K11907 | 0 | 0 | 0 | 0 | 0 | 0.00017 | 0.002207 | 0 | 0 | 0 | 0.000477 | 0 | 0 | 0 | 0 | 0 | 0 | 0 | 0 | 0 | 0 |
| Variovorax:K11927 | 0 | 0 | 0 | 0.0016 | 0 | 4.15E-05 | 0 | 0 | 0 | 0 | 0 | 0 | 0 | 0 | 0 | 0 | 0 | 0 | 0 | 0 | 0 |
| Variovorax:K11959 | 0 | 0 | 0 | 0 | 0 | 4.15E-05 | 0 | 0 | 0 | 0 | 0 | 0 | 0 | 0 | 0 | 0 | 0 | 0 | 0 | 0 | 0 |
| Variovorax:K11962 | 0 | 0 | 0 | 0 | 0 | 4.15E-05 | 0 | 0 | 0 | 0 | 0 | 0 | 0 | 0 | 0 | 0 | 0 | 0 | 0 | 0 | 0 |
| Variovorax:K11963 | 0 | 0 | 0 | 0 | 0 | 4.15E-05 | 0 | 0 | 0 | 0 | 0 | 0 | 0 | 0 | 0 | 0 | 0 | 0 | 0 | 0 | 0 |
| Variovorax:K12368 | 0 | 0 | 0 | 0 | 0 | 4.15E-05 | 0 | 0 | 0.001025 | 0 | 0 | 0 | 0 | 0 | 0 | 0 | 0 | 0 | 0 | 0 | 0 |
| Variovorax:K12573 | 0 | 0 | 0 | 0 | 0 | 0.00012 | 0 | 0 | 0 | 0 | 0 | 0 | 0 | 0 | 0 | 0 | 0 | 0 | 0 | 0 | 0 |
| Variovorax:K13482 | 0 | 0 | 0 | 0 | 0 | 0.00012 | 0 | 0 | 0.001025 | 0 | 0 | 0 | 0 | 0 | 0 | 0 | 0 | 0 | 0 | 0 | 0 |

**Supplementary Table 10.** Comparison of relative abundance of antibiotic resistance genes in the BRS and the ancient (Ötzi) and modern (Italian, Hadza and Malawian) comparative IM cohorts.

| KEGG | BRS4 | Otzi_LPLI | Otzi_SI | Otzi_UPLI2 | Otzi_UPLI | Hadza_H11 | Hadza_H15 | Hadza_H17 | Hadza_H9 | Italian_IT2 | Italian_IT5 | Italian_IT6 |
|---|---|---|---|---|---|---|---|---|---|---|---|---|
| *phoP* | 0 | 0 | 0 | 0 | 0 | 6 | 0 | 0 | 0 | 0 | 0 | 0 |
| *SOXS* | 1 | 0 | 0 | 0 | 0 | 13 | 0 | 0 | 0 | 0 | 0 | 1 |
| *CRP* | 0 | 0 | 0 | 0 | 0 | 20 | 0 | 0 | 0 | 0 | 0 | 0 |
| *parE* | 73 | 0 | 0 | 0 | 209 | 27 | 0 | 0 | 0 | 0 | 0 | 0 |
| *EMRA* | 8 | 0 | 0 | 0 | 0 | 19 | 0 | 0 | 0 | 0 | 0 | 0 |
| *EMRB* | 9 | 0 | 0 | 0 | 0 | 24 | 0 | 0 | 0 | 0 | 0 | 0 |
| *APH3-PRIME* | 0 | 0 | 0 | 0 | 35 | 0 | 0 | 0 | 0 | 73 | 44 | 11 |
| *cat* | 0 | 0 | 0 | 0 | 0 | 0 | 0 | 0 | 0 | 83 | 32 | 90 |
| *GADX* | 10 | 0 | 0 | 0 | 0 | 17 | 0 | 0 | 0 | 0 | 0 | 0 |
| *EMRK* | 9 | 0 | 0 | 0 | 0 | 11 | 0 | 0 | 0 | 0 | 0 | 0 |
| *ERMB* | 0 | 0 | 0 | 0 | 0 | 0 | 0 | 0 | 0 | 153 | 1 | 129 |
| *CTX* | 0 | 0 | 0 | 0 | 0 | 92 | 0 | 0 | 0 | 0 | 0 | 0 |
| *pmrA* | 6 | 0 | 0 | 0 | 0 | 14 | 0 | 0 | 0 | 0 | 0 | 0 |
| *OMPF* | 9 | 0 | 0 | 0 | 0 | 16 | 0 | 0 | 0 | 0 | 0 | 0 |
| *TET32* | 0 | 0 | 0 | 0 | 0 | 29 | 172 | 140 | 226 | 387 | 0 | 49 |
| *ERMF* | 0 | 0 | 0 | 0 | 0 | 9 | 0 | 0 | 0 | 14 | 385 | 26 |
| *EMRR* | 3 | 0 | 0 | 0 | 0 | 12 | 0 | 0 | 0 | 0 | 0 | 0 |
| *ERMG* | 0 | 0 | 0 | 0 | 0 | 0 | 0 | 0 | 0 | 0 | 2 | 80 |
| *PHOP* | 0 | 0 | 0 | 0 | 0 | 10 | 0 | 0 | 0 | 0 | 0 | 0 |
| *PHOQ* | 13 | 0 | 0 | 0 | 0 | 22 | 0 | 0 | 0 | 0 | 0 | 0 |
| *AMPH* | 11 | 0 | 0 | 0 | 0 | 15 | 0 | 0 | 0 | 0 | 0 | 0 |
| *TET39* | 0 | 4 | 0 | 42 | 50 | 0 | 0 | 0 | 0 | 0 | 0 | 0 |
| *OXA* | 0 | 23 | 0 | 390 | 478 | 0 | 0 | 0 | 0 | 0 | 0 | 0 |
| *EMRY* | 15 | 0 | 0 | 0 | 0 | 18 | 0 | 0 | 0 | 0 | 0 | 0 |
| *PARC* | 0 | 0 | 0 | 0 | 0 | 32 | 0 | 0 | 0 | 0 | 0 | 0 |
| *tufA/B* | 871 | 1023 | 0 | 86 | 81 | 50 | 0 | 0 | 35 | 0 | 0 | 55 |
| *PARE* | 75 | 0 | 0 | 224 | 0 | 0 | 0 | 0 | 0 | 0 | 0 | 0 |
| *rpoB* | 547 | 1312 | 2377 | 1541 | 1346 | 46 | 0 | 0 | 0 | 0 | 0 | 0 |
| *VANTG* | 0 | 0 | 0 | 0 | 0 | 0 | 11 | 5 | 0 | 0 | 0 | 0 |
| *MDFA* | 10 | 0 | 0 | 0 | 0 | 14 | 0 | 0 | 0 | 0 | 0 | 0 |
| *BACA* | 0 | 0 | 0 | 0 | 0 | 17 | 0 | 0 | 0 | 0 | 0 | 0 |
| *EVGA* | 4 | 0 | 0 | 0 | 0 | 5 | 0 | 0 | 0 | 0 | 0 | 0 |
| *ACI* | 0 | 0 | 0 | 0 | 0 | 0 | 0 | 0 | 0 | 0 | 20 | 0 |
| *PMRB* | 8 | 0 | 0 | 0 | 0 | 23 | 0 | 0 | 0 | 0 | 0 | 0 |
| *PBP2* | 0 | 0 | 0 | 0 | 0 | 29 | 0 | 0 | 0 | 0 | 0 | 0 |
| *PMRC* | 15 | 0 | 0 | 0 | 0 | 38 | 0 | 0 | 0 | 0 | 0 | 0 |
| *PMRF* | 8 | 0 | 0 | 0 | 0 | 19 | 0 | 0 | 0 | 0 | 0 | 0 |
| *DFRF* | 0 | 0 | 0 | 0 | 0 | 0 | 2 | 16 | 6 | 10 | 22 | 11 |
| *LNUC* | 0 | 0 | 0 | 0 | 0 | 0 | 0 | 0 | 0 | 12 | 0 | 0 |
| *EVGS* | 26 | 0 | 0 | 0 | 0 | 42 | 0 | 0 | 0 | 0 | 0 | 0 |
| *BAER* | 7 | 0 | 0 | 0 | 0 | 11 | 0 | 0 | 0 | 0 | 0 | 0 |
| *BAES* | 6 | 0 | 0 | 0 | 0 | 17 | 0 | 0 | 0 | 0 | 0 | 0 |
| *marA* | 1 | 0 | 0 | 0 | 0 | 6 | 0 | 0 | 0 | 0 | 0 | 0 |

| | | | | | | | | | | | | |
|---|---|---|---|---|---|---|---|---|---|---|---|---|
| *MEL* | 0 | 0 | 0 | 0 | 0 | 0 | 0 | 0 | 0 | 0 | 0 | 48 |
| *BLAEC* | 7 | 0 | 0 | 0 | 0 | 16 | 0 | 0 | 0 | 0 | 0 | 0 |
| *TETO* | 0 | 0 | 0 | 0 | 0 | 64 | 257 | 249 | 422 | 247 | 140 | 233 |
| *GyrA* | 249 | 0 | 0 | 0 | 0 | 22 | 0 | 0 | 0 | 0 | 0 | 0 |
| *CATP* | 0 | 0 | 0 | 0 | 0 | 0 | 0 | 0 | 0 | 0 | 39 | 0 |
| *SAT* | 0 | 0 | 0 | 0 | 0 | 0 | 0 | 0 | 0 | 11 | 4 | 0 |
| *GYRB* | 39 | 0 | 0 | 0 | 0 | 37 | 0 | 0 | 0 | 0 | 0 | 0 |
| *ASMA* | 13 | 0 | 0 | 0 | 0 | 26 | 0 | 0 | 0 | 0 | 0 | 0 |
| *MEFA* | 0 | 0 | 0 | 0 | 0 | 0 | 0 | 0 | 0 | 0 | 8 | 105 |
| *CBLA* | 0 | 0 | 0 | 0 | 0 | 0 | 0 | 0 | 0 | 0 | 27 | 32 |
| *TETQ* | 0 | 0 | 0 | 0 | 0 | 396 | 1001 | 865 | 779 | 52 | 1162 | 419 |
| *TETR* | 0 | 14 | 0 | 21 | 46 | 12 | 0 | 0 | 0 | 0 | 0 | 0 |
| *CPXAR* | 0 | 0 | 0 | 0 | 0 | 23 | 0 | 0 | 0 | 0 | 0 | 0 |
| *HNS* | 4 | 0 | 0 | 0 | 0 | 4 | 0 | 0 | 0 | 0 | 0 | 0 |
| *AAC6-PRIME* | 0 | 1 | 0 | 73 | 132 | 0 | 0 | 0 | 0 | 0 | 0 | 0 |
| *MDTA* | 8 | 0 | 0 | 0 | 0 | 20 | 0 | 0 | 0 | 0 | 0 | 0 |
| *TETW* | 0 | 0 | 0 | 0 | 0 | 435 | 757 | 954 | 851 | 1277 | 109 | 601 |
| *TET40* | 0 | 0 | 0 | 0 | 0 | 24 | 69 | 85 | 39 | 58 | 0 | 0 |
| *MDTB* | 39 | 0 | 0 | 0 | 0 | 28 | 0 | 0 | 0 | 0 | 0 | 0 |
| *TETX* | 0 | 0 | 0 | 0 | 0 | 0 | 0 | 0 | 0 | 0 | 181 | 74 |
| *mdtC* | 66 | 0 | 0 | 0 | 0 | 89 | 0 | 0 | 0 | 0 | 0 | 0 |
| *MDTD* | 6 | 0 | 0 | 0 | 0 | 18 | 0 | 0 | 0 | 0 | 0 | 0 |
| *MDTE* | 10 | 0 | 0 | 0 | 0 | 23 | 0 | 0 | 0 | 0 | 0 | 0 |
| *TET44* | 0 | 0 | 0 | 0 | 0 | 18 | 14 | 0 | 0 | 0 | 0 | 0 |
| *MARR* | 4 | 0 | 0 | 0 | 0 | 4 | 0 | 0 | 0 | 0 | 0 | 0 |
| *MDTG* | 7 | 0 | 0 | 0 | 0 | 17 | 0 | 0 | 0 | 0 | 0 | 0 |
| *MDTH* | 7 | 0 | 0 | 0 | 0 | 12 | 0 | 0 | 0 | 0 | 0 | 0 |
| *ACRD* | 21 | 0 | 0 | 0 | 0 | 45 | 0 | 0 | 0 | 0 | 0 | 0 |
| *Cfx* | 0 | 0 | 0 | 0 | 0 | 24 | 94 | 63 | 19 | 0 | 201 | 263 |
| *ACRE* | 11 | 0 | 0 | 0 | 0 | 23 | 0 | 0 | 0 | 0 | 0 | 0 |
| *MDTK* | 16 | 0 | 0 | 0 | 0 | 16 | 0 | 0 | 0 | 0 | 0 | 0 |
| *ACRF* | 31 | 0 | 0 | 0 | 0 | 68 | 0 | 0 | 0 | 0 | 0 | 0 |
| *MDTL* | 11 | 0 | 0 | 0 | 0 | 26 | 0 | 0 | 0 | 0 | 0 | 0 |
| *TOLC* | 10 | 0 | 0 | 0 | 0 | 25 | 0 | 0 | 0 | 0 | 0 | 0 |
| *MDTN* | 11 | 0 | 0 | 0 | 0 | 11 | 0 | 0 | 0 | 0 | 0 | 0 |
| *MDTO* | 24 | 0 | 0 | 0 | 0 | 24 | 0 | 0 | 0 | 0 | 0 | 0 |
| *MDTP* | 0 | 0 | 0 | 0 | 0 | 24 | 0 | 0 | 0 | 0 | 0 | 0 |
| *ARNA* | 18 | 0 | 0 | 0 | 0 | 35 | 0 | 0 | 0 | 0 | 0 | 0 |
| *CPXA* | 0 | 0 | 0 | 0 | 0 | 4 | 0 | 0 | 0 | 0 | 0 | 0 |
| *MSRD* | 0 | 0 | 0 | 0 | 0 | 0 | 0 | 0 | 0 | 0 | 0 | 150 |
| *APH6* | 0 | 0 | 0 | 0 | 0 | 1 | 0 | 0 | 0 | 0 | 0 | 0 |
| *ACRS* | 7 | 0 | 0 | 0 | 0 | 11 | 0 | 0 | 0 | 0 | 0 | 0 |
| *PBP4B* | 8 | 0 | 0 | 0 | 0 | 18 | 0 | 0 | 0 | 0 | 0 | 0 |
| *FOLP* | 5 | 0 | 0 | 0 | 0 | 11 | 0 | 0 | 0 | 0 | 0 | 0 |

**Supplementary Table 11.** Raw and filtered high-quality sequence read counts as related to the BRS and the ancient (Ötzi) and modern (Italian, Hadza and Malawian) comparative IM datasets.

| Sample | Raw reads | Raw bases | Filtered reads | Filtered bases | BLASTn assigned reads | BLASTx assigned reads | % BLASTn assignment | %BLASTx assignement |
|---|---|---|---|---|---|---|---|---|
| BRS1 (sediment) | 17979669 | 1456353189 | 563981 | 42423854 | 29291 | 27914 | 5.19 | 4.95 |
| BRS2 (interior) | 36050926 | 2920125006 | 3773449 | 285580025 | 295759 | 441778 | 7.84 | 11.71 |
| BRS3 (interior) | 8754692 | 709130052 | 1835294 | 140687394 | 121925 | 216567 | 6.64 | 11.80 |
| BRS4 (interior) | 117683541 | 9532366821 | 39149751 | 2968477951 | 2880169 | 4485273 | 7.36 | 11.46 |
| BRS5 (sediment) | 110485402 | 894931740 | 1426974 | 109596591 | 41704 | 39372 | 2.92 | 2.76 |
| Italian IT8_F | 3801021 | 380102100 | 3679972 | 366382762 | 1223400 | 1421338 | 33.24 | 38.62 |
| Italian IT5_F | 16744111 | 1674411100 | 16400101 | 1633075868 | 7153759 | 10282078 | 43.62 | 62.70 |
| Italian IT6_F | 23917309 | 2391730900 | 21193228 | 2107364176 | 7619871 | 12069668 | 35.95 | 56.95 |
| Italian IT2_F | 9974264 | 997426400 | 9423584 | 938267232 | 3257428 | 4235481 | 34.57 | 44.95 |
| Hadza_H17_F | 38920714 | 3892071400 | 35711833 | 3552002824 | 3382760 | 14397581 | 9.47 | 40.32 |
| Hadza_H11_F | 8271548 | 827154800 | 8198629 | 815836774 | 1242394 | 2724587 | 15.15 | 33.23 |
| Hadza_H15_F | 10431372 | 1043137200 | 10158369 | 1011507644 | 983029 | 3133139 | 9.68 | 30.84 |
| HadzaH_9_F | 8006701 | 800670100 | 7896108 | 785798043 | 1094654 | 2754683 | 13.86 | 34.89 |
| Otzi_SI_F | 20871329 | 2108004229 | 19819371 | 1611496115 | 7106155 | 3628829 | 35.85 | 18.31 |
| Otzi_UPLI | 60954954 | 6156450354 | 35549389 | 2635224622 | 5413130 | 3729770 | 15.23 | 10.49 |
| Otzi_UPLI2 | 63743693 | 6438112993 | 35669617 | 2678808549 | 5729827 | 4148223 | 16.06 | 11.63 |
| Otzi_LPLI | 33452784 | 3378731184 | 28363635 | 2416377839 | 9782299 | 901180 | 34.49 | 3.18 |
| Malawi_mgm4461162 | 170067 | 71848878 | 170067 | 71848878 | 37980 | 131180 | 22.33 | 77.13 |
| Malawi_mgm4461165 | 304035 | 120341771 | 304035 | 120341771 | 176041 | 272672 | 57.90 | 89.68 |
| Malawi_mgm4461146 | 193786 | 83497735 | 193786 | 83497735 | 122470 | 178850 | 63.20 | 92.29 |
| Malawi_mgm4461161 | 349265 | 144828528 | 349265 | 144828528 | 314312 | 333400 | 89.99 | 95.46 |
| *Total* | *591061183* | *46021426480* | *279830438* | *24519425175* | *58008357* | *69553563* | – | – |

**Supplementary Table 12.** Information concerning the comparative NCBI genomes used during this study.

| Taxonomic classification | Genomes downloaded | Date of download | Accession numbers |
|---|---|---|---|
| **Environmental taxa (species)** | | | |
| *Babesia bovis* | 1 | 09/25/18 | GCA_000165395.1 |
| *Lactococcus lactis* | 149 | 09/25/18 | – |
| *Ficus sur* | NA | NA | NA |
| *Parastrongyloides trichosuri* | 1 | 09/25/18 | GCA_000941615.1 |
| *Podospora anserina* | 1 | 09/25/18 | GCA_000226545.1 |
| *Sorghum bicolor* | 1 | 09/12/18 | ABXC00000000.3 |
| *Triticum aestivum* | 1 | 09/15/18 | GCA_900519105.1 |
| *Zea mays* | 1 | 09/12/18 | GCA_000005005.6 |
| *Bos taurus* | 1 | 09/21/18 | GCA_000005005.6 |
| *Capra hircus* | 1 | 09/25/18 | GCA_001704415.1 |
| *Equus caballus* | 1 | 09/15/18 | LWLT00000000.1 |
| *Homo sapiens* | 1 | 03/20/17 | GCA_000001405.2 |
| *Macaca mulatta* | 1 | 09/25/18 | GCA_000772875.3 |
| *Ovis aries* | 1 | 09/25/18 | GCA_000298735.2 |
| *Pan troglodytes* | 1 | 09/25/18 | GCA_002880755.3 |
| **Commensal taxa (genera)** | | | |
| *Methanobrevibacter* | 9 | 09/11/18 | – |
| *Bifidobacterium* | 107 | 09/11/18 | – |
| *Cellulomonas* | 45 | 09/12/18 | – |
| *Slackia* | 6 | 09/12/18 | – |
| *Alistipes* | 53 | 09/12/18 | – |
| *Bacteroides* | 442 | 09/12/18 | – |
| *Flavobacterium* | 312 | 09/12/18 | – |
| *Parabacteroides* | 63 | 09/12/18 | – |
| *Pedobacter* | 70 | 09/12/18 | – |
| *Prevotella* | 494 | 09/12/18 | – |
| *Blautia* | 81 | 09/12/18 | – |
| *Butyrivibrio* | 62 | 09/12/18 | – |
| *Clostridium* | 905 | 09/12/18 | – |

| | | | |
|---|---|---|---|
| *Coprococcus* | 38 | 09/12/18 | – |
| *Dorea* | 25 | 09/12/18 | – |
| *Enterococcus* | 91 | 09/12/18 | – |
| *Eubacterium* | 103 | 09/12/18 | – |
| *Faecalibacterium* | 59 | 09/12/18 | – |
| *Lactobacillus* | 271 | 09/12/18 | – |
| *Lactococcus* | 195 | 09/12/18 | – |
| *Mogibacterium* | 4 | 09/12/18 | – |
| *Mollicutes* | 248 | 09/12/18 | – |
| *Oribacterium* | 19 | 09/12/18 | – |
| *Oscillibacter* | 15 | 09/12/18 | – |
| *Phascolarctobacterium* | 8 | 09/12/18 | – |
| *Romboutsia* | 9 | 09/12/18 | – |
| *Roseburia* | 77 | 09/12/18 | – |
| *Ruminiclostridium* | 9 | 09/12/18 | – |
| *Ruminococcus* | 165 | 09/12/18 | – |
| *Sarcina* | 1 | 09/12/18 | GCA_900101015.1 |
| *Tissierellia* | 131 | 09/12/18 | – |
| *Enterobacter* | 1456 | 09/12/18 | – |
| *Klebsiella* | 304 | 09/12/18 | – |
| *Neisseria* | 123 | 09/12/18 | – |
| *Sphingobium* | 99 | 09/12/18 | – |
| *Variovorax* | 82 | 09/12/18 | – |
| *Treponema* | 189 | 09/12/18 | – |
| **Pathogenic taxa (genera)** | | | |
| *Aeromonas* | 68 | 08/30/18 | – |
| *Achromobacter* | 141 | 09/06/18 | – |
| *Bordetella* | 409 | 09/06/18 | – |
| *Burkholderia* | 228 | 09/06/18 | – |
| *Comamonas* | 57 | 09/11/18 | – |
| *Delftia* | 36 | 09/06/18 | – |
| *Elizabethkingia* | 123 | 09/06/18 | – |
| *Listeria* | 178 | 09/06/18 | – |

| | | | |
|---|---|---|---|
| *Lysobacter* | 39 | 09/06/18 | – |
| *Mycobacterium* | 2342 | 09/06/18 | – |
| *Shigella* | 90 | 09/06/18 | – |
| *Vibrio* | 162 | 09/07/18 | – |
| *Weeksella* | 4 | 09/07/18 | – |
